# Viromes in Marine Ecosystems Reveal Remarkable Invertebrate RNA Virus Diversity

**DOI:** 10.1101/2021.04.21.440720

**Authors:** Yu-Yi Zhang, Yicong Chen, Xiaoman Wei, Jie Cui

## Abstract

Ocean viromes remain poorly understood and little is known about the ecological factors driving aquatic RNA virus evolution. In this study, we used a meta-transcriptomic approach to characterize the viromes of 58 marine invertebrate species across three seas. This revealed the presence of 315 newly identified RNA viruses in nine viral families or orders (*Durnavirales, Totiviridae, Bunyavirales, Hantaviridae, Picornavirales, Flaviviridae, Hepelivirales, Solemoviridae* and *Tombusviridae*), with most of them are sufficiently divergent to the documented viruses. With special notice that we first time revealed an ocean virus rooting to mammalian hantaviruses. We also found evidence for possible host sharing and switch events during virus evolution. In sum, we demonstrated the hidden diversity of marine invertebrate RNA viruses.

## Introduction

Viruses are ubiquitous and exist in every living species (Koonin, et al. 2006; Zhang, et al. 2019). They are mainly studied as agents of disease in humans, animals with biosafety and economical importance. However, pathogenic viruses represent only a minor proportion of the virosphere (Geoghegan and Holmes 2017; Middelboe and Brussaard 2017; Zhang, et al. 2018; Salazar, et al. 2019; Zhang, et al. 2019). Advances in meta-genomics and meta-transcriptomics led to the discovery of an enormous amount of viruses, most of which are distinct from presently well-defined pathogenic viruses (Shi, et al. 2016; Shi, Lin, et al. 2018; Chang, Eden, et al. 2020; Chang, Li, et al. 2020; Pettersson, et al. 2020; Wu, et al. 2020). These findings have filled important gaps in virus evolution and reflected the fact that RNA viruses with relatively small genome size could also have huge diversity in genomic elasticity (Qin, et al. 2014; Zhang, et al. 2018). The International Committee on Taxonomy of Viruses (ICTV) has announced to add the viruses discovered by metagenomic and meta-transcriptomic into the formal classification of viruses (Simmonds, et al. 2017).

Invertebrate organisms are considered as ancient species and represent the majority of animal biodiversity, especially in the oceans (Lopez, et al. 2019). Marine invertebrates are important food source for aquatic higher animals and are significant to maintain the homeostasis of ecosystem. The oceans, which cover 71% of the Earth, are the most valuable resource for investigating the ecological and geographical diversity of viruses. However, to date, aquatic viromic studies tended to study phages but already showed enormous genetic diversity and ecological distributions (Roux, et al. 2016; Vlok, et al. 2019; Guajardo-Leiva, et al. 2020; Gulino, et al. 2020; Moon, et al. 2020; Zhang, et al. 2020). Although some studies included aquatic invertebrates (Magbanua, et al. 2000; Thakur, et al. 2002; Shi, et al. 2016; Rosani, et al. 2019), there is a lack of systematic study of these animals in a whole.

To address key questions on the transmission and evolutionary patterns of viruses in aquatic invertebrates, we conducted an extensive analysis of RNA viromes in more than 50 aquatic invertebrates acquired in three different seas. In particular, by measuring and comparing the composition and structure of the viral community within different seas, we aimed to understand the virus-host interaction among them. Such exploration in this study could not only help us to better understand how viruses spread between oceans and form the virosphere, but also provide a theoretical basis for the exploitation and utilization of marine resources.

## Results

### Characterization of RNA viromes across different seas

Fifty-eight aquatic invertebrate species of 696 individuals were acquired from five locations in three different seas (the South China Sea, the East China Sea, and the Yellow Sea). These species represented six classes (*Polychaeta, Crustacea, Hexanauplia, Bivalvia, Gastropoda*, and *Cephalopoda*) of three phyla (*Annelida, Arthropoda* and *Mollusca*) (Table S1). Fifty-eight RNA sequencing libraries were generated and sequenced to a high depth and assembled *de novo*. In total, 2,242,428,860 paired-end (PE) reads were generated. The library size of species ranged from 28,501,148 to 64,776,106 PE reads (Fig. 1 and Table S1).

**Fig. 1.**
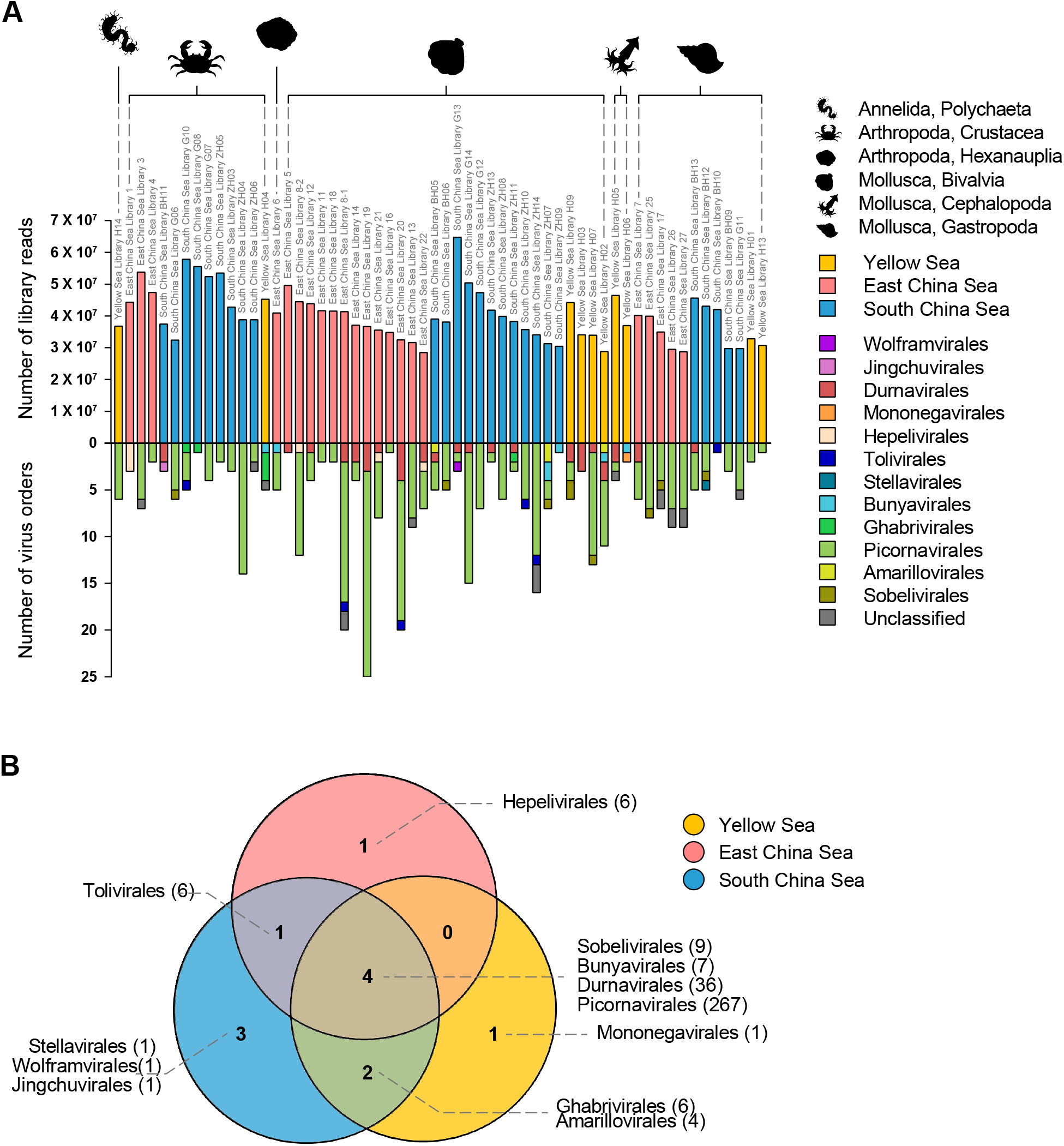
The overall characterization of virus distribution. **A)** The distribution and diversity of virus in invertebrate transcriptomes. The top graph shows the number of reads in each library. The colors of the bars indicate the location of sample collecting, Yellow Sea (yellow), East China Sea (red), South China Sea (blue). The full name of each library is shown on top of each bar, while major host classifications are shown above the bar graph. The bottom graph shows a summary of classification of virus species found in this study. **B)** Overlap of RNA viral families in aquatic invertebrates across different seas. The number of viruses in each order or family is shown in brackets. Circles are color coded according to the location of sample collecting, Yellow Sea (yellow), East China Sea (red), South China Sea (blue).

Using a series of protein sequence similarity-based BLAST searches, 363 RNA viral (or partial) genomes were identified, 117 of which contained RNA-dependent RNA polymerase (RdRp) regions. Previous studies showed endogenous viral elements (EVEs) existed in the meta-transcriptome generated sequences (Shi, et al. 2016). It is unlikely that the viruses in this study acted as endogenized forms, as they showed limited similarity to animal genome sequences and contained opening reading frames (ORFs) without any premature stop codon. A similarity comparison of RdRp region indicated that 106 viruses were novel and distinct to publicly available viruses in Genbank. Ninety-seven fell within known viral families or orders, including double-stranded RNA viruses (*Durnavirales* and *Totiviridae*) (Fig. 2), negative-sense single-stranded RNA viruses (*Bunyavirales* and *Hantaviridae*) (Fig, 3), and positive-sense single-stranded RNA viruses (*Picornavirales, Flaviviridae, Hepelivirales, Solemoviridae* and *Tombusviridae*) (Fig. 4). In addition, for the absence of RdRp conserved domain, 246 aquatic invertebrate viruses identified here were unable to be classified. The viruses identified in each species ranged from 1 to 26 (East China Sea), 1–16 (South China Sea), and 1–13 (Yellow Sea). The number of viruses identified was uneven across different water areas in different species (Fig. 1A). There were multiple groups of viruses common to all three seas, including *Sobelivirales, Bunyavirales, Durnavirales*, and *Picornavirales*, which comprised more than half of the viruses discovered in our study (Fig. 1B). And *Tolivirales* were discovered both in South China Sea and East China Sea, while *Ghabrivirales* and *Amarillovirales* were discovered in South China Sea and Yellow Sea. Viruses were heterogeneous in different seas, as each sea harbored unique viruses. However, as the viral abundance in different seas much relied on their host distribution, such that description above should be treated with caution.

**Fig. 2.**
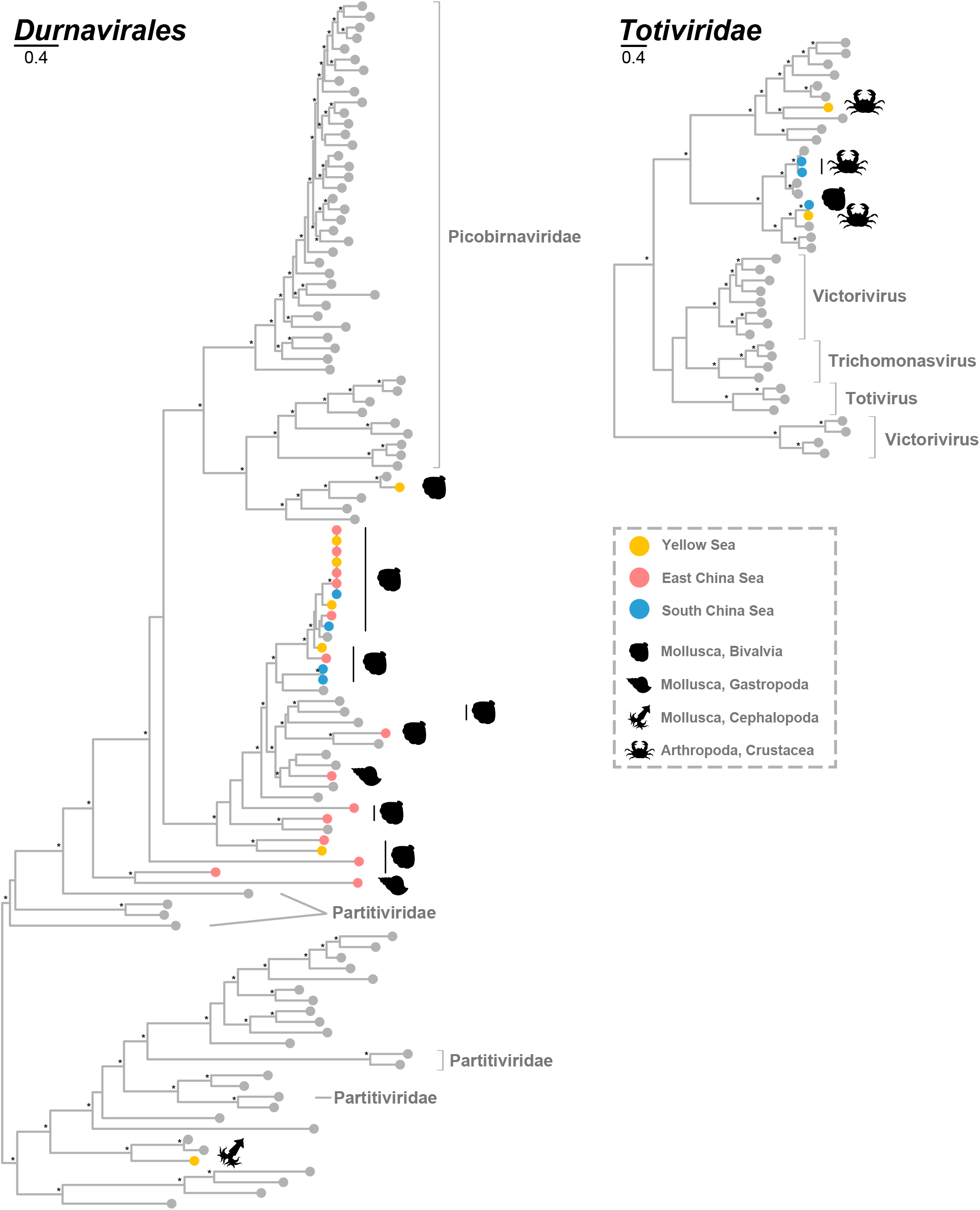
Phylogenetic trees of the dsRNA viruses, including the viruses identified in this study and related representative viruses. The trees are inferred using amino acid sequences of the RdRp gene and flanking conserved domain. These trees are midpoint rooted for clarity only. Viruses identified in this study are denoted with a filled colored circle based on the areas where their hosts were acquired, Yellow Sea (yellow), East China Sea (red), South China Sea (blue), while other representative publicly available viruses are denoted with a grey circle. An asterisk indicates node SH-aLRT support >70%. The scale bar indicates the number of amino acid changes per site.

**Fig. 3.**
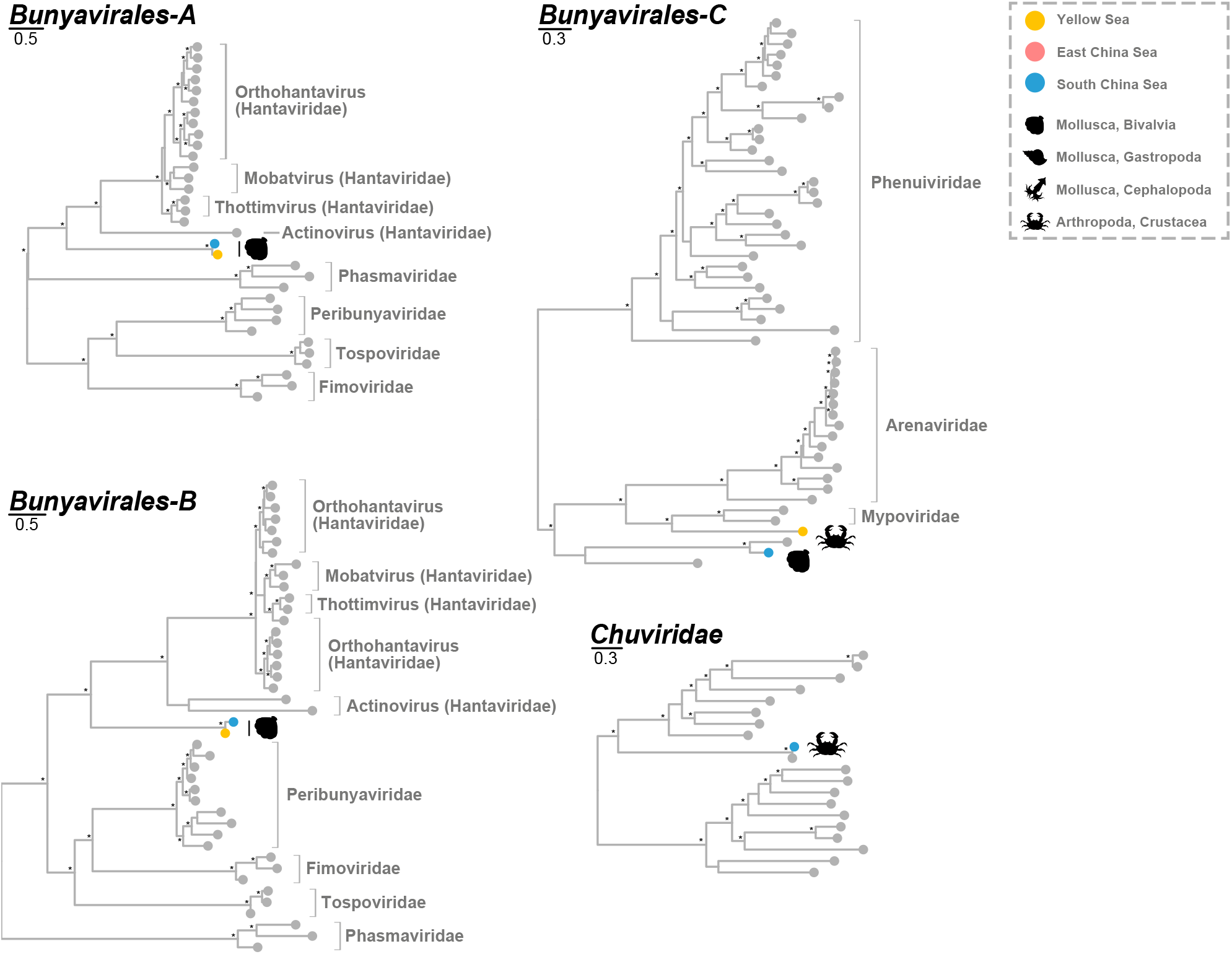
Phylogenetic trees of the -ssRNA viruses, including the viruses identified in this study and related representative viruses. The trees are inferred using amino acid sequences of the RdRp gene and flanking conserved domain. These trees are midpoint rooted for clarity only. Viruses identified in this study are denoted with a filled colored circle based on the area where their hosts were acquired, Yellow Sea (yellow), East China Sea (red), South China Sea (blue), while other representative publicly available viruses which are denoted with a grey circle. An asterisk indicates node SH-aLRT support >70%. The scale bar indicates the number of amino acid changes per site.

**Fig. 4.**
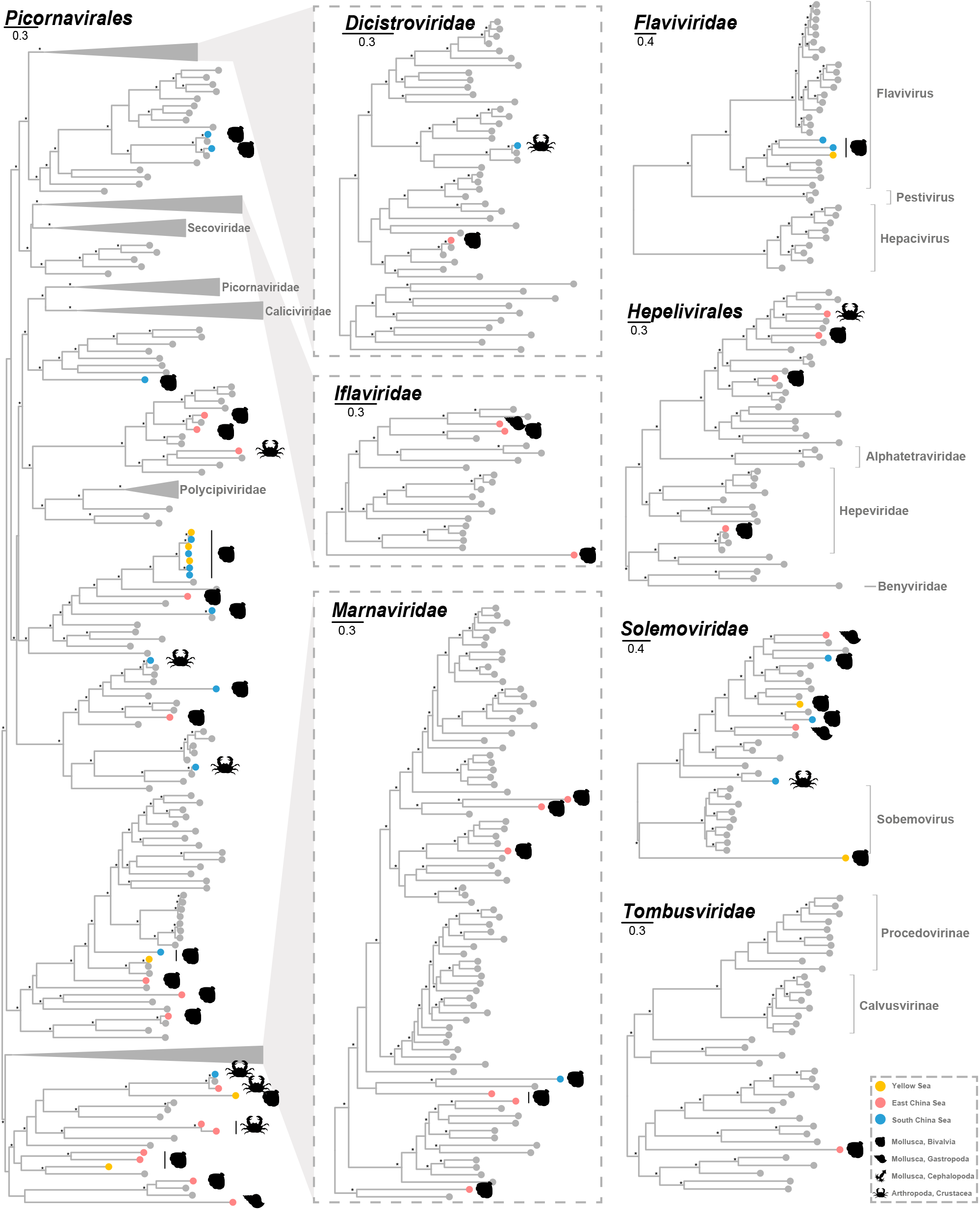
Phylogenetic trees of the +ssRNA viruses, including the viruses identified in this study and related representative viruses. The trees are inferred using amino acid sequences of the RdRp gene and flanking conserved domain. These trees are midpoint rooted for clarity only. Viruses identified in this study are denoted with a filled colored circle based on the area where their hosts were acquired, Yellow Sea (yellow), East China Sea (red), South China Sea (blue), while other representative publicly available viruses are denoted with a grey circle. An asterisk indicates node SH-aLRT support >70%. The scale bar indicates the number of amino acid changes per site.

### Characterization of double-strand RNA viruses

Thirty dsRNA virus genomes were characterized that belonged to one virus order and virus family (Fig. 2, S1 and S2).

Twenty-five viruses were classified into *Durnavirales* (Fig. 2 and S1). Phylogenetic analysis revealed that Bivalvia Durna-like virus H2 was closely related to Dralkin virus (Wille, et al. 2020), a penguin virus in a clade related to vertebrate-specific genogroup 1, indicating that it was a possible potential zoonotic pathogen. Twelve viruses clustered strongly in the supposedly invertebrate-specific genogroup 3. Importantly, among these viruses, thirteen viruses, namely, Bivalvia Durna-like viruses D1, H7, D7, H5, D2, D14, N4, H1, D16, N3, H8, D15, and Beihai picobirna-like virus 10, formed a close monophyletic group with high amino acid similarity and short branch lengths indicating possible host sharing across different seas. In addition, Bivalvia Durna-like viruses D11 and D5 and Gastropoda Durna-like virus D2 were distantly related to *Picobirnaviridae*, showing that they were unclassified picobirna-like viruses. Cephalopoda Durna-like virus H2 fell outside of well-defined viruses from *Partitiviridae* and clustered with other unclassified partiti-like viruses, indicating that it belonged to the unclassified partiti-like viruses. The capsid-encoded segment was not identified in Cephalopoda Durna-like virus H2. Although, members of the family *Partitiviridae* were believed to contain two genomic segments (Vainio, et al. 2018).

Five Toti-like viruses were characterized in *Crustacea* and *Bivalvia* from the South China Sea and *Crustacea* from the Yellow Sea (Fig. 2 and S2). These viruses showed limited sequence similarity with the recognized totiviruses. Their genome size ranged from 1 kb to 8 kb, which was dissimilar to the typical virus genome length (4.6–7.0 kb) in *Totiviridae* (Fig. 5). Phylogenetic analysis revealed that Bivalvia Ghabri-like virus N1 from South China Sea clustered with Crustacea Ghabri-like virus H3 from the Yellow Sea, which indicated possible host sharing between different seas during evolution. Host sharing was also observed within the same sea, as Crustacea Ghabri-like virus N1 found in *Charybdis feriata* and Crustacea Ghabri-like virus N2 found in *Scylla olivacea* clustered together and shared high similarity.

**Fig. 5.**
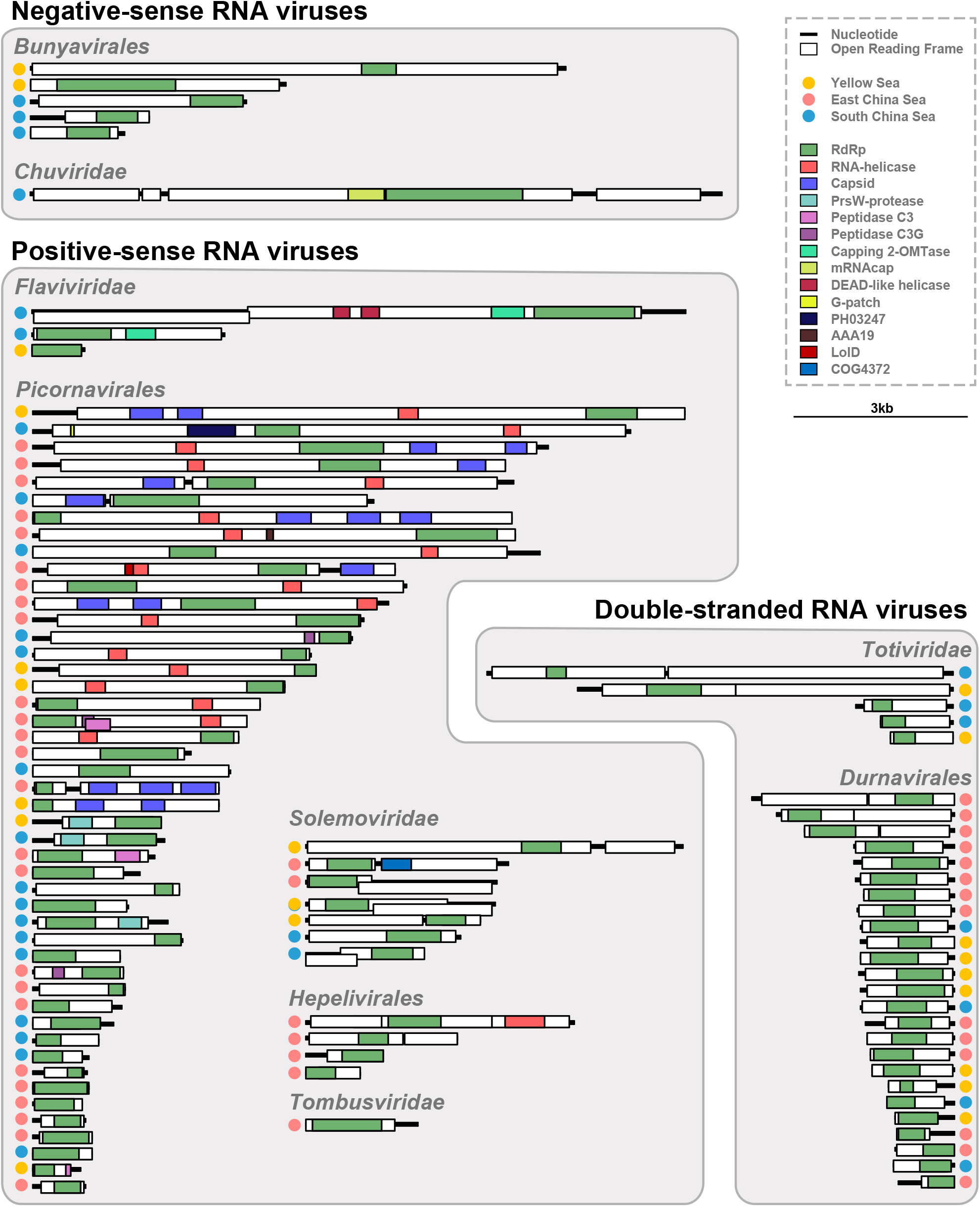
Genome organization of representative viruses within major viral clades. The contigs and genomes are drawn as lines and boxes to scale, respectively. The predicted regions that encode major functional proteins or domains are labelled with colored boxes. The filled colored circles indicate the area where their hosts were acquired, Yellow Sea (yellow), East China Sea (red), South China Sea (blue).

### Characterization of negative-sense RNA viruses

Nine ssRNA(-) viruses were identified within 6 of the 58 libraries present in this study (Fig. 3 and S3-S5). Three viruses in Bivalvia acquired from the Yellow Sea and South China Sea were closed related to *Hantaviridae*, a family of segmented RNA viruses (Fig. 2 and S3) (Laenen, et al. 2019). However, the viruses here only contained single L segments (Fig. 5) containing RdRp regions. Phylogenetic analysis revealed that these viruses fell out of the *Hantaviridae* cluster. However, a BLAST similarity search indicated that Bivalvia Bunya-like virus H1 and Bivalvia Bunya-like virus N2 showed low similarity (25% and 28%, respectively) to Wenling minipizza batfish hantavirus (Actinovirus), whereas Bivalvia Bunya-like virus N1 showed low similarity to Camp Ripley virus (24%). Thus, these viruses were deemed to be novel distinct hanta-like viruses. It was noteworthy that these viruses showed high similarity with each other (78%-86%), indicating host sharing between different seas.

Crustacea Bunya-like virus H1 was characterized and classified as a mypo-like virus, as this virus fell into the clustered group containing *Mypoviridae* and the best BLAST hit was Hubei myriapoda virus 5 with 21% protein similarity (Fig. 2 and S4). Bivalvia Bunya-like virus N3 in *Bunyavirales* was not able to be classified into family, as this virus showed limited similarity to any well-defined family in *Bunyavirales*. Instead, it was grouped with unclassified *Bunyavirales* including Beihai bunya-like virus 3 and Wuhan snail virus.

Crustacea Jingchu-like virus N1 from South China Sea fell within the recently established *Chuviridae* (Fig. 3 and S5). It exhibited typical chuvirus structure, encoding glycoprotein protein and containing RdRp and RNA capping domain (Fig. 5). However, the reverse location of RdRp and RNA capping domain also elucidated the diverse genome structure of chuviruses. It showed 97% nucleotide similarity with previously identified Beihai hermit crab virus 3, which was acquired in the same sea. They also formed a well-supported monophyletic group within the mivirus clade, which was compatible with the idea that these two viruses might have a single origin.

### Positive-sense RNA viruses

Our study contained genomic evidence for the presence of 61 ssRNA(+) viruses in *Bivalvia, Gastropoda, Cephalopoda*, and *Crustacea* across three seas and these viruses were from three viral families and two orders (Fig. 4 and S6-S11).

The most well-represented order of viruses discovered in our survey is *Picornavales* (Fig. 4 and S6-S7). There were 46 viruses from three seas widely distributed in the *Picornavales* phylogenetic tree and most of these viruses were isolated from South China Sea and East China Sea (Table S1-S2). The elasticity of their genome structures illustrated the diversity of *Picornavales* (Fig. 5). Two viruses (Crustacea Picorna-like virus N14 from South China Sea and Bivalvia Picorna-like virus D23 from East China Sea) fell into the *Dicistroviridae* clade. Both of them shared high similarity (80% and 97%, respectively) to their closely related viruses from South China Sea, showing possible host sharing across or within different seas. Three viruses from East China Sea fell within *Iflaviridae* cluster. The phylogenetic tree revealed that all of these viruses were distantly related to other viruses. Seven viruses fell within *Marnaviridae*, all of which were also distantly related to other viruses. Other viruses did not cluster with any well-defined family but clustered with other unclassified picorna-like viruses. These 34 viruses widely distributed in the phylogeny of *Picornavales*. Several host-sharing events could be characterized involving Bivalvia Picorna-like viruses H13, N7, H15, N20, H14, N56, and N21, Bivalvia Picorna-like virus N58, and Wenzhou picorna-like virus 38.

Three viruses in *Bivalvia* (Bivalvia Amarillo-like virus N1, Bivalvia Amarillo-like virus N3, and Bivalvia Amarillo-like virus H1) formed a single cluster that fell within the *Flaviridae* clade, indicating that they were flaviviruses (Fig. 4 and S8). Compared with the flaviviruses in invertebrates, they were most closely related to squid flaviviruses, although they only exhibited 26%–36% similarity to southern pygmy squid flavivirus. Unexpectedly, DEAD-like helicase C (cl38915), which was involved in ATP-dependent RNA or DNA unwinding and existed in eukaryotic cells and in many bacteria and Archaea (Linder and Jankowsky 2011), was identified in Bivalvia Amarillo-like virus N1, lending additional support to the idea that gene exchange occurs between host and virus.

Four viruses from East China Sea were identified as *Hepelivirales* (Fig. 4 and S9). Phylogenetic analysis revealed that Bivalvia Hepeli-like virus D2 belonged to *Hepeviridae* and a similarity search showed that it was identical to Barns Ness breadcrumb sponge hepe-like virus 2, indicating a possible host-switching event during evolution. Another three viruses (Bivalvia Hepeli-like virus D1 and D3 and Crustacea Hepeli-like virus D1) showed limited similarity to well-defined classified viruses. They and other invertebrate hepeli-like viruses formed a cluster related to *Alphatetraviridae* that was once thought to have a narrow host range (larvae of lepidopteran insect species, such as moth and butterfly) and tissue tropism (Dorrington, et al. 2020).

Seven viruses across all three seas were related to *Solemoviridae* (plant/invertebrate-associated) (Fig. 4 and S10). All of these viruses showed limited similarity to sobemoviruses, which were once thought to be plant specific, with long branches, probably giving support to the idea that virus may have switched the hosts on different trophic levels during evolution, as some Bivalvia and Gastropoda fed on aquatic plants (Harder, et al. 2016; Iturriza-Gomara and O’Brien 2016; Wu, et al. 2020).

One virus (Bivalvia Toli-like virus D1) only had a partial genome containing the RdRp region and fell within *Tombusviridae* (Fig. 4-5 and S11). The amino acid sequence similarity to the most closely related virus species (Sanxia tombus-like virus 5) was 30%. Both viruses are distantly related to viruses from *Calvusvirinae* and *Procedovirinae*, indicating that Bivalvia Toli-like virus D1 is a tombus-like virus.

### Likelihood of Host-sharing and switching pattern accordance to the ecology

A large number of diverse viruses (including >300 novel species) were discovered in three different seas (Yellow Sea, South China Sea, and East China Sea). As the phylogenetic analysis combined with genome comparison indicated, a large amount of host sharing events occurred between these viruses. For example, 1) the distribution of Bivalvia Durna-like virus among three ocean areas; 2) similar Bunya-like viruses shared by the South China Sea and the Yellow Sea; and 3) the sharing of Picorna-like virus in different hosts, as we described above. However, these viruses only comprised a tiny portion of these viruses found in these marine areas.

In order to gain a more complete view of patterns in RNA virus evolution in these areas, we built a dataset comprised of RNA viruses identified in these seas and in two more distinct areas, freshwater and terrestrial, with certain hosts. Finally, 2671 RNA viruses with certain hosts were added into the dataset. Network analysis indicated that there was an enormous number of host-sharing events occurring across different habitats during evolution (Fig. 6A) and number of viruses had a positive correlation with host-sharing frequency (Fig. 6B). It was noteworthy that the filtration of viruses across different seas carried by a variety of hosts.

**Fig. 6.**
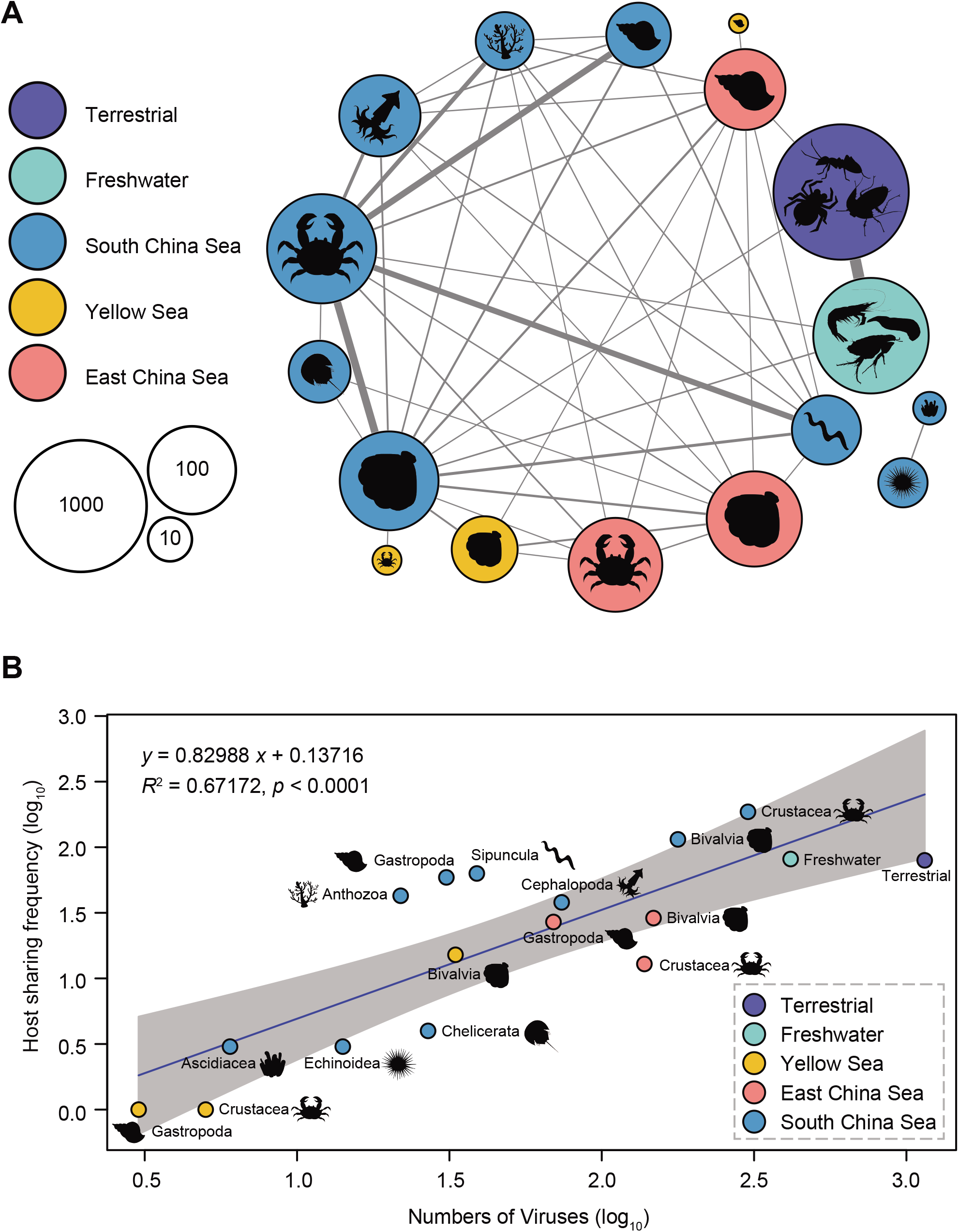
Possible host-sharing pattern of viruses in major invertebrate habitats. **A)** The viral network indicating the unevenly distributed of host sharing. Nodes represent the class of the host groups. Node size is proportional to number of viruses collected into the analysis while node color is related to the areas where the hosts live. Edge width is proportional to the putative host sharing frequency. **B)** Correlation of host sharing frequency ratio, calculated using host sharing frequency with number of viruses in each type of host in different seas. Best-fit lines with 95% confidence intervals from linear regression are plotted. The filled colored circles indicate the area where their hosts were acquired, terrestrial (bluish violet), freshwater (blue green), Yellow Sea (yellow), East China Sea (red), South China Sea (blue).

## Discussion

In this study, we performed viromics to investigate the virus community in aquatic invertebrates across three different seas. Analysis of the meta-transcriptomes of 58 aquatic species led to the discovery of the capacity of aquatic invertebrates to carry widely unknown viruses, in particular a total of 315 novel RNA viruses were characterized (Table S2). More than half of the newly identified RNA viruses showed 20%–50% similarity to their most closely related viruses, indicating the hidden genetic diversity of marine viruses. We were able to identify multiple domains (e.g., DEAD-like helicase C) that have not been previously observed in specific viral families. This revealed that the genome structures of marine viruses could be elastic (Fig. S12) and diverse than previously thought. However, the biological function of such domains still needs to be experimentally confirmed. As virus identification mostly depended on similarity-based searching, there may have been a failure to discover extremely divergent viruses (Zhang, et al. 2018). Hence, further research using more advanced viral identification methods/pipelines and metagenomic technology is merited.

Viruses were characterized in all marine invertebrate samples from this study and no visible lesions or illness were observed in any of the samples we acquired. This provided further evidence for the idea that disease-causing viruses are probably the exception rather than the rule (Junglen and Drosten 2013; Li, et al. 2015; Marklewitz, et al. 2015; Webster, et al. 2015). Interestingly, some invertebrate viruses seemed more ancestral than vertebrate viruses, i.e., rooted in the phylogeny. For example, Bivalvia Bunya-like virus H1, N1 and N2 were basal to other hantaviruses found in vertebrates (Fig. 3 and S3), suggesting a probable marine origin of *Hantaviridae*. However, large data may be needed to address the true co-divergence of this viral family.

Different seas seemingly contained sea-specific virus groups as shown in Fig. 1. This probably indicated the restrained heterogeneity of viral communities in different seas. However, no statistical methods were used to determine sample size and distribution. Thus, this conclusion should be treated with caution and an in-depth study of these viruses remains necessary. An amount of cross-ocean transmission and host sharing events were observed in this study (Fig. 6), suggesting evolutionary connectivity of marine viruses between seas. The possible transmission of viruses across different seas might involve natural factors such as moving of marine species. More importantly, we observed some viruses crossing the water–land interface during evolution, indicating that the ocean is not a boundary for host sharing or host switching. In sum, we showed the hidden diversity of RNA viruses in marine invertebrates and revealed the geographical structure of viruses and transmission dynamic of the viruses in multiple marine environments.

## Materials and Methods

### Sample collection and preparation

58 marine invertebrate species (in the phylum of *Arthropoda* and *Mollusca*) were collected from the seafood markets or from the fishermen in 5 different locations (Table S1). No statistical methods were used to determine sample size and distribution. Twelve individuals of each species were collected, then raised in artificial seawater for 24h if conditions permitted. Tissues of viscera for all the individuals were dissected and pooled together to increase the virus abundance. Samples were stored in RNAlater Stabilization Solution (Invitrogen) at room temperature according to the instruction before transferred to a -80°C freezer.

### Sequence library construction and sequencing

To construct each library, samples were homogenized first and total RNA was extracted using TRIzol Reagent (Invitrogen). rRNA was removed using Ribo-Zero rRNA Removal Kit (Human/Mouse/Rat) and Ribo-Zero rRNA Removal Kit (Bacteria) (Illumina). In order to reduce the impact of the host transcriptome on subsequent analysis, we enriched viral nucleic acid by negative selection targeted RNA with poly A tails, and then prepare the library using Nextera XT DNA Library Preparation Kit (Illumina). Paired-end (150bp) sequencing of each library was performed on the NovaSeq 6000 (Illumina).

### Virus discovery and annotation

For each library, sequencing reads were quality trimmed and then assembled *de novo* using Trinity program (Haas, et al. 2013) with default settings. The assembled (consensus) contigs were then screened against the non-redundat protein (nr) database using Diamond blastx (Buchfink, et al. 2015) with a cut-off e-value of 1E-5. We excluded the hits which showed similarity to the host, plant, bacterial, and fungal sequences to reduce potential internal or external contaminants. The remaining viral hits were further filtered to remove viruses with host of plant, bacteria and fungi as described previously (Wille, et al. 2019). All the sequences were searched for the RdRp region with a RdRp dataset collected from NCBI refseq database (Table S2). A virus was considered novel if the RdRp region showed < 90% amino acid similarity to any previously identified virus (Shi, et al. 2016; Chang, Eden, et al. 2020). All the sequence data could be accessed from National Genomics Data Center (https://bigd.big.ac.cn/) with BioProject Accession PRJCA003705 (also see Table S2) and should be used with judgement. To study the genome structure, we predicted the open reading frames (ORF) with the website of ORF finder provided by NCBI (https://www.ncbi.nlm.nih.gov/orffinder), and performed domain-based search with NCBI conserved domain database (CDD) (Lu, et al. 2020) with an expect value threshold of 0.01.

### Confirmation of viral contigs

Viral contigs were confirmed by RT-PCR method with multiple primers (Table S4) designed according to the assembled sequences. Amplification products were identified by Sanger sequencing and by aligning to the raw contigs (identity > 95%).

### Host species confirmation

The cytochrome c oxidase subunit I (COI) gene was amplified by PCR method with the primers LCO1490 and HCO2198 reported before (Folmer, et al. 1994), and then sequenced by Sanger sequencing. They were subsequently compared against the nr database to confirm the host species (identity > 90%). For those specimens that failed PCR experiment due to low specificity of general primers, we used COI protein sequence from related host (download from NCBI refseq database) as a bait to obtain the sequences from the assembled contigs with tblastn. However, in some samples, the COI sequences showed low identity to sequences from other species. This probably suggested possible high diversity within the species or the misclassification at the species level as mentioned previously (Metzger, et al. 2018).

### Phylogenetic analysis

To inferring the evolutionary history of all RNA viruses identified in this study, sequences of viruses identified here, combining with protein sequences obtained from GenBank using the top search results from BLAST (Altschul, et al. 1990), and representative viruses in ICTV were collected. Then, the RdRp protein sequences of these viruses were respectively aligned in accordance to their classified orders or families using MAFFT 7.271 (Katoh and Standley 2013). In order to reduce the alignment uncertainty, regions that aligned poorly were removed using TrimAL (Capella-Gutiérrez, et al. 2009) with the default setting and confirmed manually using MEGA X (Kumar, et al. 2018). the sequence was excluded if its length was less than 1/3 of the alignment. In total, 97 of 117 viruses were included in phylogenetic analysis. The best-fit model for each alignment was selected and maximum likelihood (ML) phylogenetic trees were constructed using IQ-Tree 1.6.12 (Nguyen, et al. 2015), incorporating 1000 replicates of SH-like approximate likelihood ratio test (SH-aLRT) to assess node robustness. Phylogenetic trees were viewed and annotated in FigTree V1.4.3 (https://github.com/rambaut/figtree/).

### Host sharing event characterization

To determine the potential host sharing events, all the reported publicly available viruses with certain host from five specific area (South China Sea, East China Sea, Yellow Sea, freshwater and terrestrial) were collected into the dataset. The dataset comprised of 2607 RNA viruses with certain hosts (Table S3). Nucleotide sequences were first aligned with MAFFT 7.271 (Katoh and Standley 2013) and manually confirmed. Comparison of genetic diversity between the different viruses was undertaken by computing the number of base differences per site averaged over all sequence pairs between pairwise viruses using MEGA X (Kumar, et al. 2018). Pairwise viruses from different areas with a distance less than 0.2 as one host sharing event (Wille, et al. 2019; Mahar, et al. 2020; Wille, et al. 2020) and the host sharing network was drawn with Cytoscape 3.8.1 (Shannon, et al. 2003).

### Compliance and ethics

The authors declare that they have no conflict of interest.

## Supporting information

Host and geographic information and data output for each library of aquatic invertebrate samples

The information about viruses identified in this study

The information about viruses used for host sharing estimation

The information of primers used for PCR confirmation

## Acknowledgments

This work was supported by National Natural Science Foundation of China (31970176), Collaborative Research Grant (KLMVI-OP-202002) of CAS Key Laboratory of Molecular Virology & Immunology, Institut Pasteur of Shanghai, Chinese Academy of Sciences, Guangdong Provincial Key Laboratory of Fishery Ecology and Environment (FEEL-2019-6), and CAS Pioneer Hundred Talents Program.

## References

Altschul SF, Gish W, Miller W, Myers EW, Lipman DJ. 1990. Basic local alignment search tool. J Mol Biol 215:403–410.

Buchfink B, Xie C, Huson DH. 2015. Fast and sensitive protein alignment using DIAMOND. Nat Methods 12:59–60.

Capella-Gutiérrez S, Silla-Martínez JM,Gabaldón T. 2009. trimAl: a tool for automated alignment trimming in large-scale phylogenetic analyses. Bioinformatics 25:1972–1973.

Chang WS, Eden JS, Hall J, Shi M, Rose K, Holmes EC. 2020. Metatranscriptomic Analysis of Virus Diversity in Urban Wild Birds with Paretic Disease. J Virol 94.

Chang WS, Li CX, Hall J, Eden JS, Hyndman TH, Holmes EC, Rose K. 2020. Meta-Transcriptomic Discovery of a Divergent Circovirus and a Chaphamaparvovirus in Captive Reptiles with Proliferative Respiratory Syndrome. Viruses 12.

Dorrington RA, Jiwaji M, Awando JA, Bruyn MM. 2020. Advances in Tetravirus Research: New Insight Into the Infectious Virus Lifecycle and an Expanding Host Range. Curr Issues Mol Biol 34:145–162.

Folmer O, Black M, Hoeh W, Lutz R, Vrijenhoek R. 1994. DNA primers for amplification of mitochondrial cytochrome c oxidase subunit I from diverse metazoan invertebrates. Mol Mar Biol Biotechnol 3:294–299.

Geoghegan JL, Holmes EC. 2017. Predicting virus emergence amid evolutionary noise. Open Biol 7.

Guajardo-Leiva S, Chnaiderman J, Gaggero A, Díez B. 2020. Metagenomic Insights into the Sewage RNA Virosphere of a Large City. Viruses 12.

Gulino K, Rahman J, Badri M, Morton J, Bonneau R, Ghedin E. 2020. Initial Mapping of the New York City Wastewater Virome. mSystems 5.

Haas BJ, Papanicolaou A, Yassour M, Grabherr M, Blood PD, Bowden J, Couger MB, Eccles D, Li B, Lieber M, et al. 2013. De novo transcript sequence reconstruction from RNA-seq using the Trinity platform for reference generation and analysis. Nat Protoc 8:1494–1512.

Harder TC, Buda S, Hengel H, Beer M, Mettenleiter TC. 2016. Poultry food products--a source of avian influenza virus transmission to humans? Clin Microbiol Infect 22:141–146.

Iturriza-Gomara M, O’Brien SJ. 2016. Foodborne viral infections. Curr Opin Infect Dis 29:495–501.

Junglen S, Drosten C. 2013. Virus discovery and recent insights into virus diversity in arthropods. Curr Opin Microbiol 16:507–513.

Katoh K, Standley DM. 2013. MAFFT multiple sequence alignment software version 7: improvements in performance and usability. Mol Biol Evol 30:772–780.

Koonin EV, Senkevich TG, Dolja VV. 2006. The ancient Virus World and evolution of cells. Biol Direct 1:29.

Kumar S, Stecher G, Li M, Knyaz C, Tamura K. 2018. MEGA X: Molecular Evolutionary Genetics Analysis across Computing Platforms. Mol Biol Evol 35:1547–1549.

Laenen L, Vergote V, Calisher CH, Klempa B, Klingström J, Kuhn JH, Maes P. 2019. Hantaviridae: Current Classification and Future Perspectives. Viruses 11.

Li CX, Shi M, Tian JH, Lin XD, Kang YJ, Chen LJ, Qin XC, Xu J, Holmes EC, Zhang YZ. 2015. Unprecedented genomic diversity of RNA viruses in arthropods reveals the ancestry of negative-sense RNA viruses. Elife 4.

Linder P, Jankowsky E. 2011. From unwinding to clamping - the DEAD box RNA helicase family. Nat Rev Mol Cell Biol 12:505–516.

Lopez JV, Kamel B, Medina M, Collins T, Baums IB. 2019. Multiple Facets of Marine Invertebrate Conservation Genomics. Annu Rev Anim Biosci 7:473–497.

Lu S, Wang J, Chitsaz F, Derbyshire MK, Geer RC, Gonzales NR, Gwadz M, Hurwitz DI, Marchler GH, Song JS, et al. 2020. CDD/SPARCLE: the conserved domain database in 2020. Nucleic Acids Res 48:D265–d268.

Magbanua FO, Natividad KT, Migo VP, Alfafara CG, de la Pena FO, Miranda RO, Albaladejo JD, Nadala EC, Jr., Loh PC, Mahilum-Tapay L. 2000. White spot syndrome virus (WSSV) in cultured Penaeus monodon in the Philippines. Dis Aquat Organ 42:77–82.

Mahar JE, Shi M, Hall RN, Strive T, Holmes EC. 2020. Comparative Analysis of RNA Virome Composition in Rabbits and Associated Ectoparasites. J Virol 94.

Marklewitz M, Zirkel F, Kurth A, Drosten C, Junglen S. 2015. Evolutionary and phenotypic analysis of live virus isolates suggests arthropod origin of a pathogenic RNA virus family. Proc Natl Acad Sci U S A 112:7536–7541.

Metzger MJ, Paynter AN, Siddall ME, Goff SP. 2018. Horizontal transfer of retrotransposons between bivalves and other aquatic species of multiple phyla. Proc Natl Acad Sci U S A 115:E4227–e4235.

Middelboe M, Brussaard CPD. 2017. Marine Viruses: Key Players in Marine Ecosystems. Viruses 9.

Moon K, Jeon JH, Kang I, Park KS, Lee K, Cha CJ, Lee SH, Cho JC. 2020. Freshwater viral metagenome reveals novel and functional phage-borne antibiotic resistance genes. Microbiome 8:75.

Nguyen LT, Schmidt HA, von Haeseler A, Minh BQ. 2015. IQ-TREE: a fast and effective stochastic algorithm for estimating maximum-likelihood phylogenies. Mol Biol Evol 32:268–274.

Pettersson JH, Ellström P, Ling J, Nilsson I, Bergström S, González-Acuña D, Olsen B, Holmes EC. 2020. Circumpolar diversification of the Ixodes uriae tick virome. PLoS Pathog 16:e1008759.

Qin XC, Shi M, Tian JH, Lin XD, Gao DY, He JR, Wang JB, Li CX, Kang YJ, Yu B, et al. 2014. A tick-borne segmented RNA virus contains genome segments derived from unsegmented viral ancestors. Proc Natl Acad Sci U S A 111:6744–6749.

Rosani U, Shapiro M, Venier P, Allam B. 2019. A Needle in A Haystack: Tracing Bivalve-Associated Viruses in High-Throughput Transcriptomic Data. Viruses 11.

Roux S, Brum JR, Dutilh BE, Sunagawa S, Duhaime MB, Loy A, Poulos BT, Solonenko N, Lara E, Poulain J, et al. 2016. Ecogenomics and potential biogeochemical impacts of globally abundant ocean viruses. Nature 537:689–693.

Salazar G, Paoli L, Alberti A, Huerta-Cepas J, Ruscheweyh HJ, Cuenca M, Field CM, Coelho LP, Cruaud C, Engelen S, et al. 2019. Gene Expression Changes and Community Turnover Differentially Shape the Global Ocean Metatranscriptome. Cell 179:1068–1083 e1021.

Shannon P, Markiel A, Ozier O, Baliga NS, Wang JT, Ramage D, Amin N, Schwikowski B, Ideker T. 2003. Cytoscape: a software environment for integrated models of biomolecular interaction networks. Genome Res 13:2498–2504.

Shi M, Lin XD, Chen X, Tian JH, Chen LJ, Li K, Wang W, Eden JS, Shen JJ, Liu L, et al. 2018. The evolutionary history of vertebrate RNA viruses. Nature 556:197–202.

Shi M, Lin XD, Tian JH, Chen LJ, Chen X, Li CX, Qin XC, Li J, Cao JP, Eden JS, et al. 2016. Redefining the invertebrate RNA virosphere. Nature 540:539–543.

Shi M, Zhang YZ, Holmes EC. 2018. Meta-transcriptomics and the evolutionary biology of RNA viruses. Virus Res 243:83–90.

Simmonds P, Adams MJ, Benkő M, Breitbart M, Brister JR, Carstens EB, Davison AJ, Delwart E, Gorbalenya AE, Harrach B, et al. 2017. Consensus statement: Virus taxonomy in the age of metagenomics. Nat Rev Microbiol 15:161–168.

Thakur PC, Corsin F, Turnbull JF, Shankar KM, Hao NV, Padiyar PA, Madhusudhan M, Morgan KL, Mohan CV. 2002. Estimation of prevalence of white spot syndrome virus (WSSV) by polymerase chain reaction in Penaeus monodon postlarvae at time of stocking in shrimp farms of Karnataka, India: a population-based study. Dis Aquat Organ 49:235–243.

Vainio EJ, Chiba S, Ghabrial SA, Maiss E, Roossinck M, Sabanadzovic S, Suzuki N, Xie J, Nibert M, Ictv Report C. 2018. ICTV Virus Taxonomy Profile: Partitiviridae. J Gen Virol 99:17–18.

Vlok M, Gibbs AJ, Suttle CA. 2019. Metagenomes of a Freshwater Charavirus from British Columbia Provide a Window into Ancient Lineages of Viruses. Viruses 11.

Webster CL, Waldron FM, Robertson S, Crowson D, Ferrari G, Quintana JF, Brouqui JM, Bayne EH, Longdon B, Buck AH, et al. 2015. The Discovery, Distribution, and Evolution of Viruses Associated with Drosophila melanogaster. PLoS Biol 13:e1002210.

Wille M, Harvey E, Shi M, Gonzalez-Acuña D, Holmes EC, Hurt AC. 2020. Sustained RNA virome diversity in Antarctic penguins and their ticks. ISME J 14:1768–1782.

Wille M, Shi M, Klaassen M, Hurt AC, Holmes EC. 2019. Virome heterogeneity and connectivity in waterfowl and shorebird communities. ISME J 13:2603–2616.

Wu H, Pang R, Cheng T, Xue L, Zeng H, Lei T, Chen M, Wu S, Ding Y, Zhang J, et al. 2020. Abundant and Diverse RNA Viruses in Insects Revealed by RNA-Seq Analysis: Ecological and Evolutionary Implications. mSystems 5:e00039–00020.

Zhang Y-Z, Chen Y-M, Wang W, Qin X-C, Holmes EC. 2019. Expanding the RNA Virosphere by Unbiased Metagenomics. Annual Review of Virology 6:119–139.

Zhang YZ, Shi M, Holmes EC. 2018. Using Metagenomics to Characterize an Expanding Virosphere. Cell 172:1168–1172.

Zhang Z, Qin F, Chen F, Chu X, Luo H, Zhang R, Du S, Tian Z, Zhao Y. 2020. Culturing novel and abundant pelagiphages in the ocean. Environ Microbiol.

