## Supplementary material for "Viromes in Marine Ecosystems Reveal Remarkable Invertebrate RNA Virus Diversity": Fig S1-S12

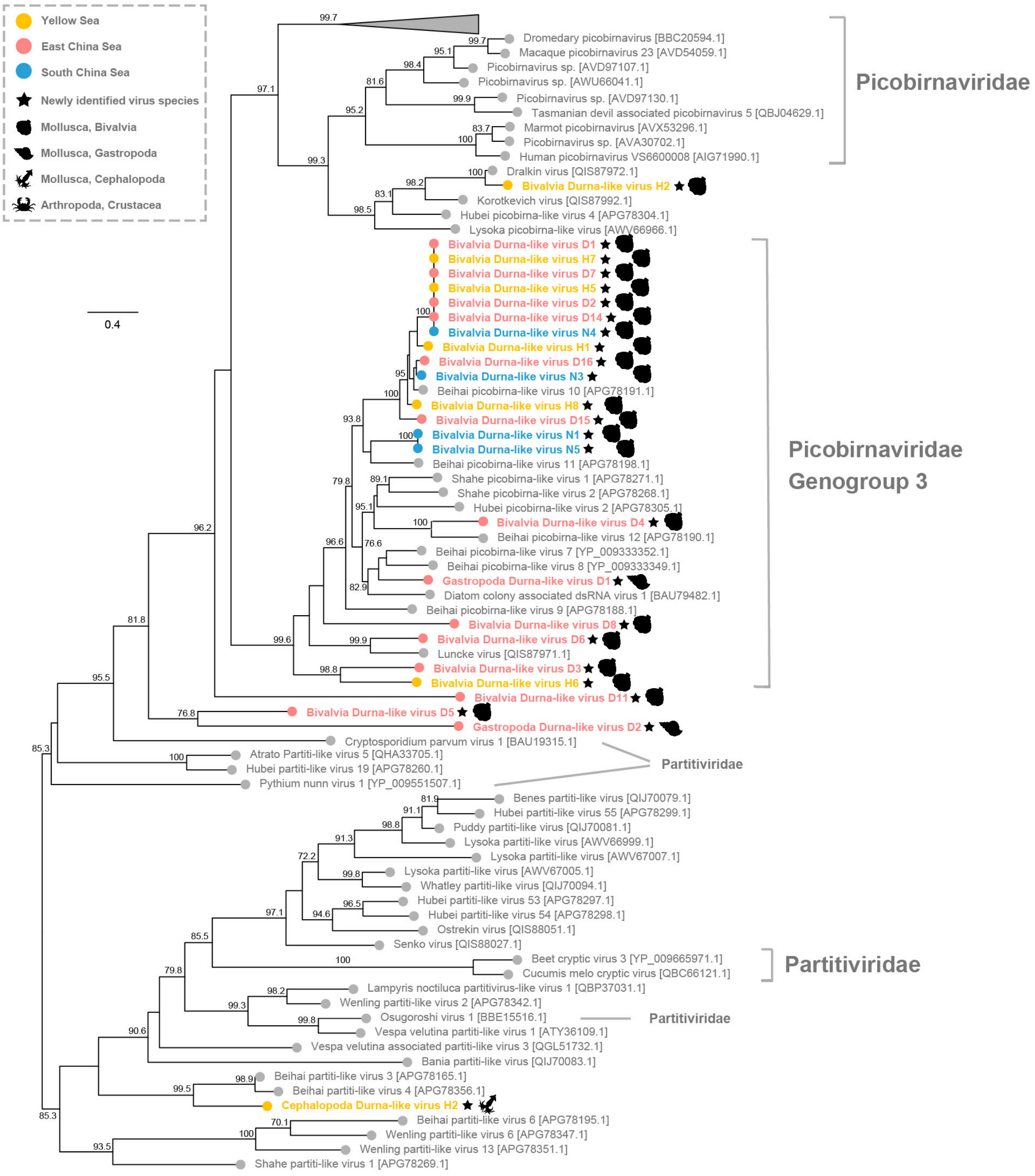


**Fig. S1 Phylogenetic trees of the Durnavirales viruses, including the viruses identified in this study and related representative viruses.** The tree is inferred using amino acid sequences of the RdRp gene and flanking conserved domain. The tree is midpoint rooted for clarity only. Viruses identified in this study are denoted with a filled colored circle based on the area where their hosts were acquired, Yellow Sea (yellow), East China Sea (red), South China Sea (blue), while other representative publicly available viruses are denoted with a grey circle. The newly identified viruses are distinguished from previously published viruses with black solid asterisk at the end of their names. SH-aLRT supports less than 70% are not shown. The scale bar indicates the number of amino acid changes per site.

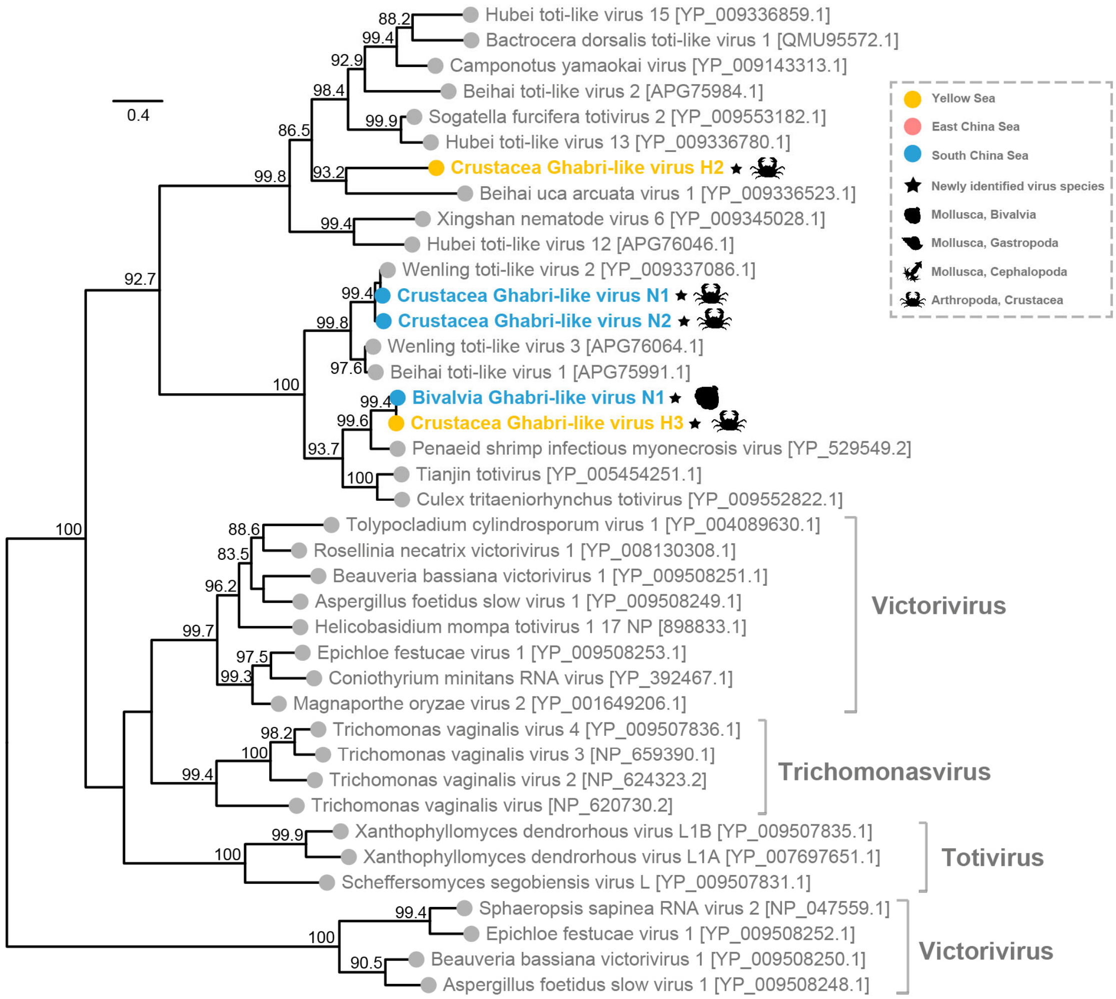


**Fig. S2 Phylogenetic trees of the Totiviridae viruses, including the viruses identified in this study and related representative viruses.** The tree is inferred using amino acid sequences of the RdRp gene and flanking conserved domain. The tree is midpoint rooted for clarity only. Viruses identified in this study are denoted with a filled colored circle based on the area where their hosts were acquired, Yellow Sea (yellow), East China Sea (red), South China Sea (blue), while other representative publicly available viruses are denoted with a grey circle. The newly identified viruses are distinguished from previously published viruses with black solid asterisk at the end of their names. SH-aLRT supports less than 70% are not shown. The scale bar indicates the number of amino acid changes per site.

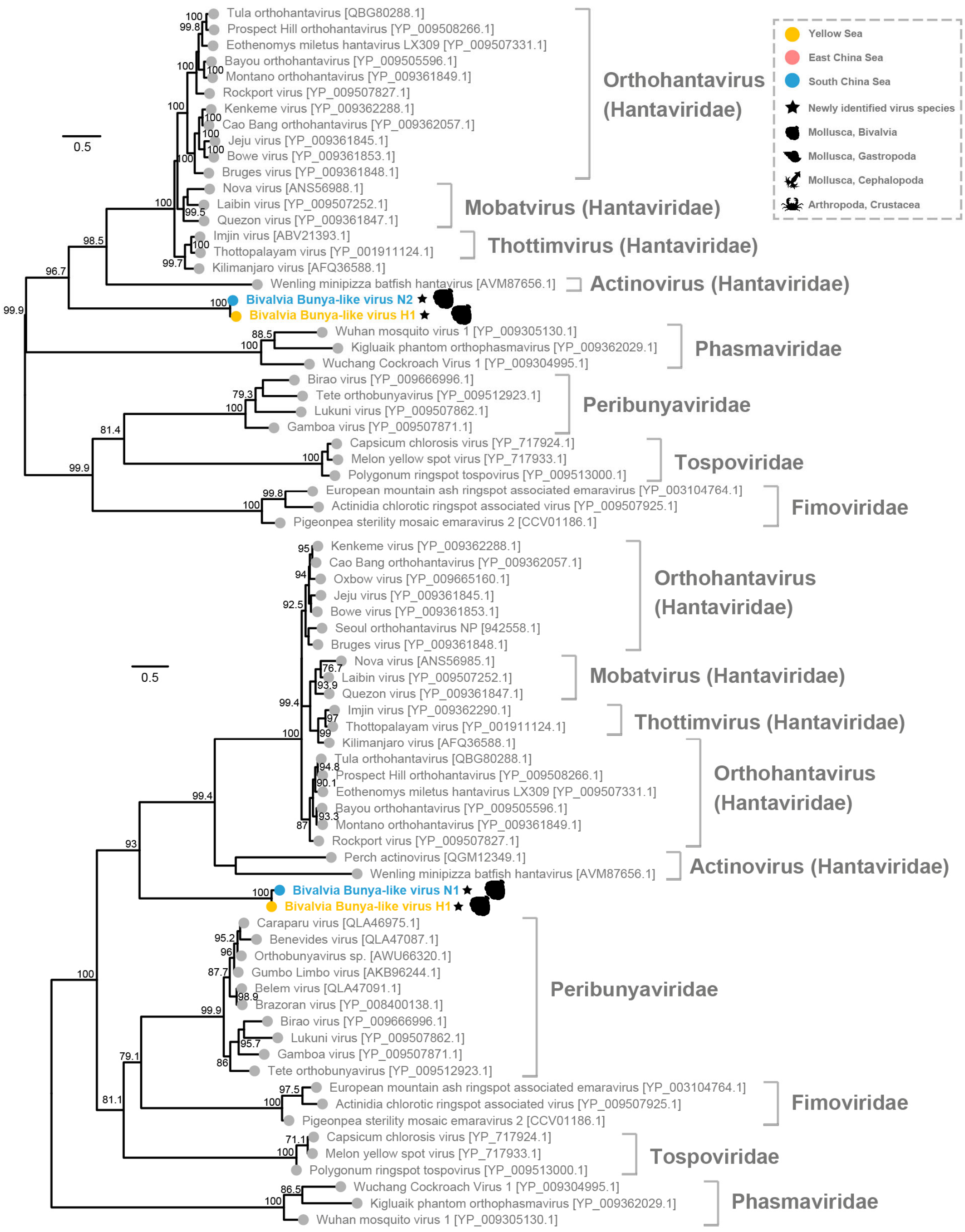


**Fig. S3 Phylogenetic trees of the Bunyavirales viruses (A, B), including the viruses identified in this study and related representative viruses.** These trees are inferred using amino acid sequences of the RdRp gene and flanking conserved domain. The trees are midpoint rooted for clarity only. Viruses identified in this study are denoted with a filled colored circle based on the area where their hosts were acquired, Yellow Sea (yellow), East China Sea (red), South China Sea (blue), while other representative publicly available viruses are denoted with a grey circle. The newly identified viruses are distinguished from previously published viruses with black solid asterisk at the end of their names. SH-aLRT supports less than 70% are not shown. The scale bar indicates the number of amino acid changes per site.

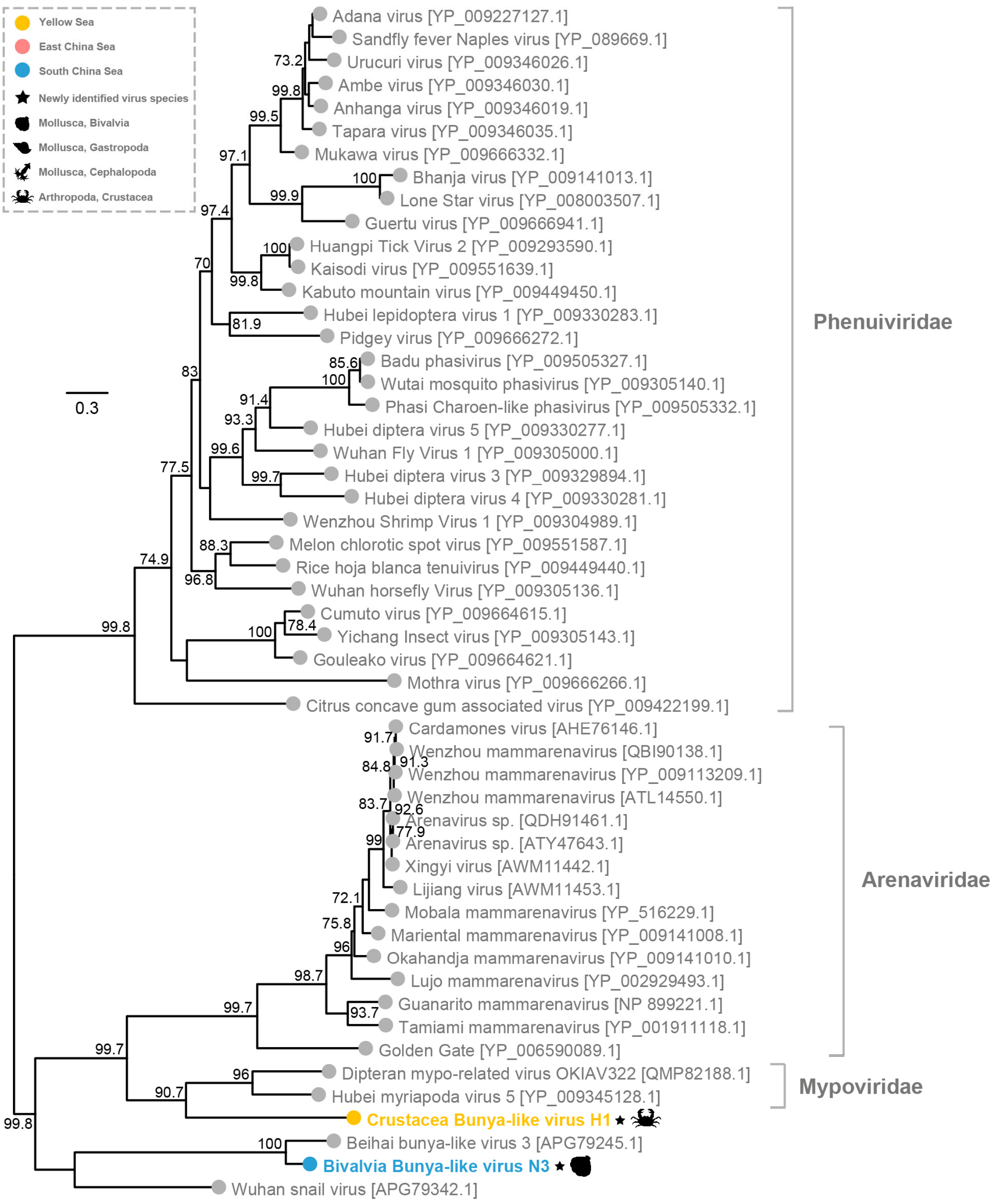


**Fig. S4 Phylogenetic trees of the Bunyavirales viruses (C), including the viruses identified in this study and related representative viruses.** The tree is inferred using amino acid sequences of the RdRp gene and flanking conserved domain. The tree is midpoint rooted for clarity only. Viruses identified in this study are denoted with a filled colored circle based on the area where their hosts were acquired, Yellow Sea (yellow), East China Sea (red), South China Sea (blue), while other representative publicly available viruses are denoted with a grey circle. The newly identified viruses are distinguished from previously published viruses with black solid asterisk at the end of their names. SH-aLRT supports less than 70% are not shown. The scale bar indicates the number of amino acid changes per site.

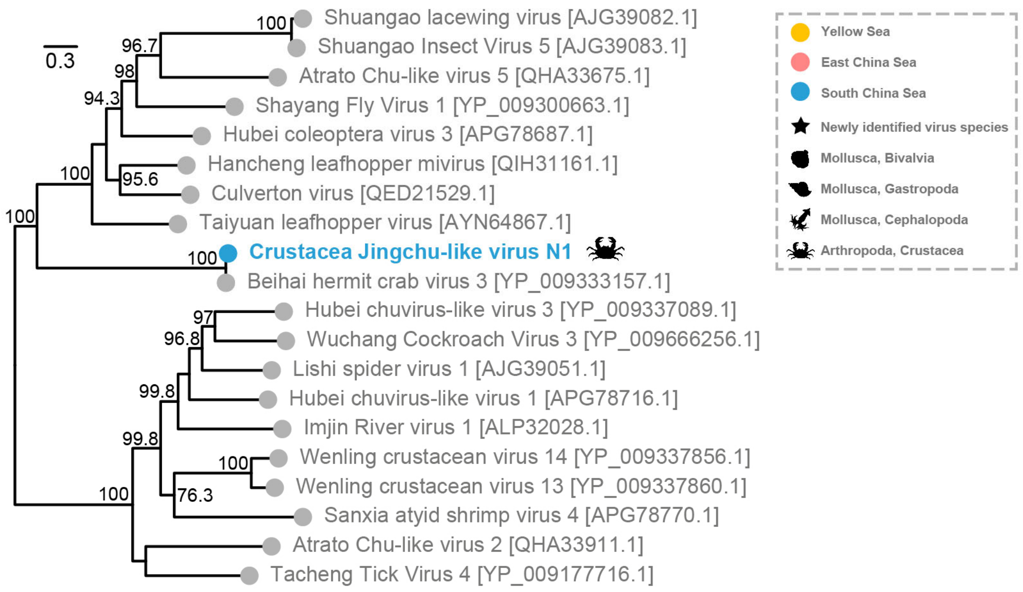


**Fig. S5 Phylogenetic trees of the Chuviridae viruses, including the viruses identified in this study and related representative viruses.** The tree is inferred using amino acid sequences of the RdRp gene and flanking conserved domain. The tree is midpoint rooted for clarity only. Viruses identified in this study are denoted with a filled colored circle based on the area where their hosts were acquired, Yellow Sea (yellow), East China Sea (red), South China Sea (blue), while other representative publicly available viruses are denoted with a grey circle. The newly identified viruses are distinguished from previously published viruses with black solid asterisk at the end of their names. SH-aLRT supports less than 70% are not shown. The scale bar indicates the number of amino acid changes per site.

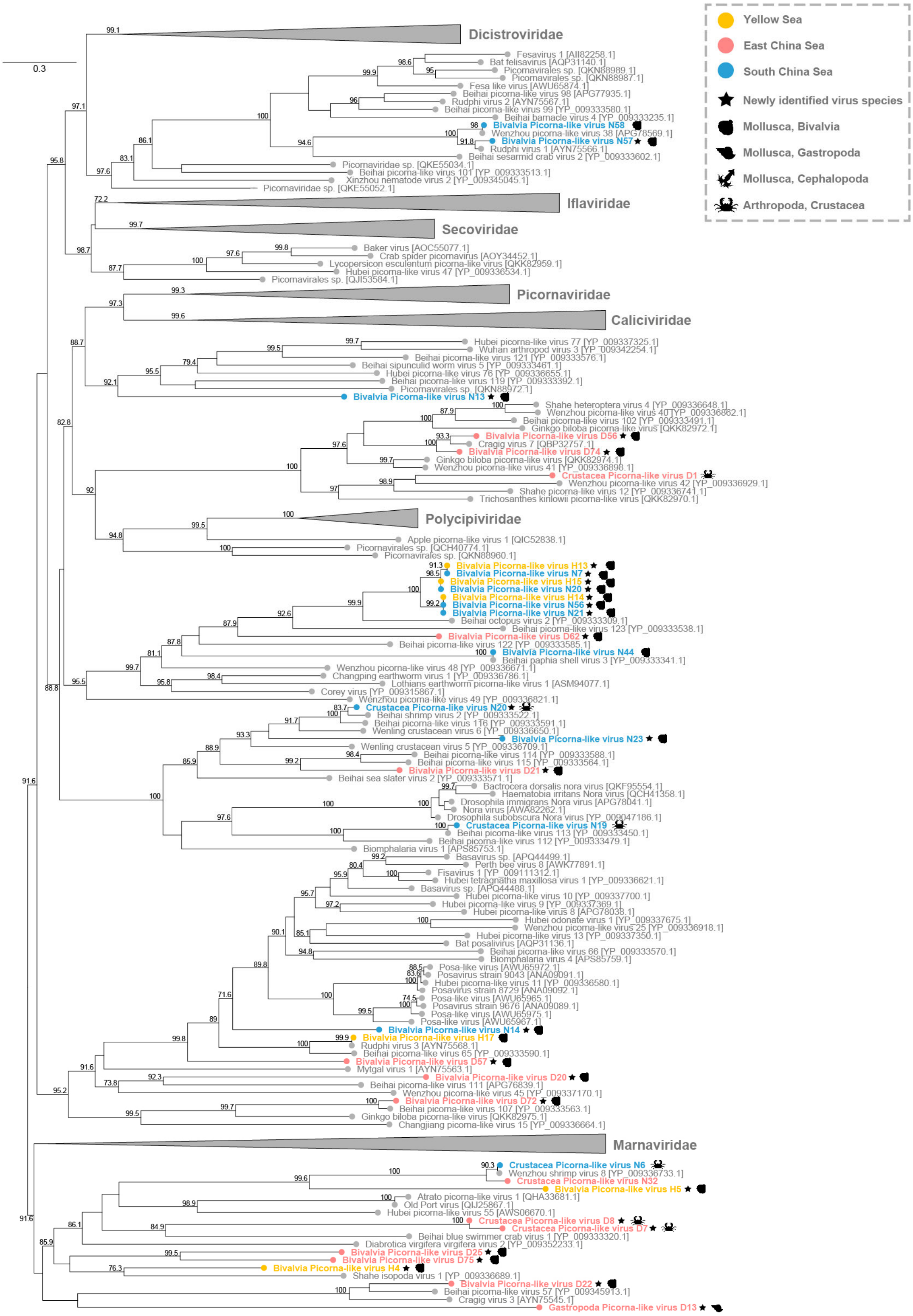


**Fig. S6 Phylogenetic trees of the Picornavirales viruses, including the viruses identified in this study and related representative viruses.** The tree is inferred using amino acid sequences of the RdRp gene and flanking conserved domain. The tree is midpoint rooted for clarity only. Viruses identified in this study are denoted with a filled colored circle based on the area where their hosts were acquired, Yellow Sea (yellow), East China Sea (red), South China Sea (blue), while other representative publicly available viruses are denoted with a grey circle. The newly identified viruses are distinguished from previously published viruses with black solid asterisk at the end of their names. SH-aLRT supports less than 70% are not shown. The scale bar indicates the number of amino acid changes per site.

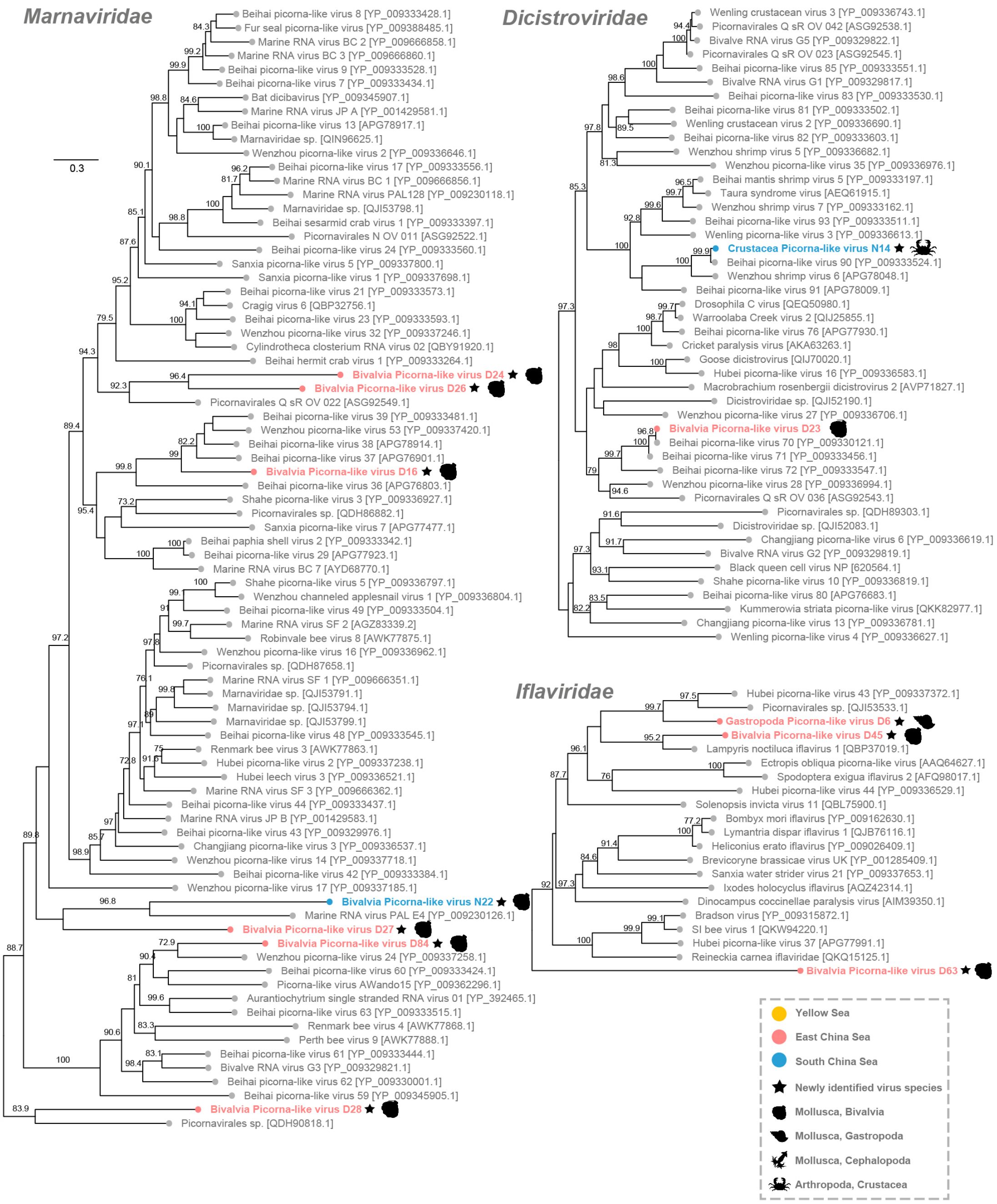


**Fig. S7 Phylogenetic trees of the Dicistroviridae viruses, Iflaviridae viruses and Marnaviridae viruses from the Picornavirales tree in Fig. S6, including the viruses identified in this study and related representative viruses.** Figure legend follows Fig. S6.

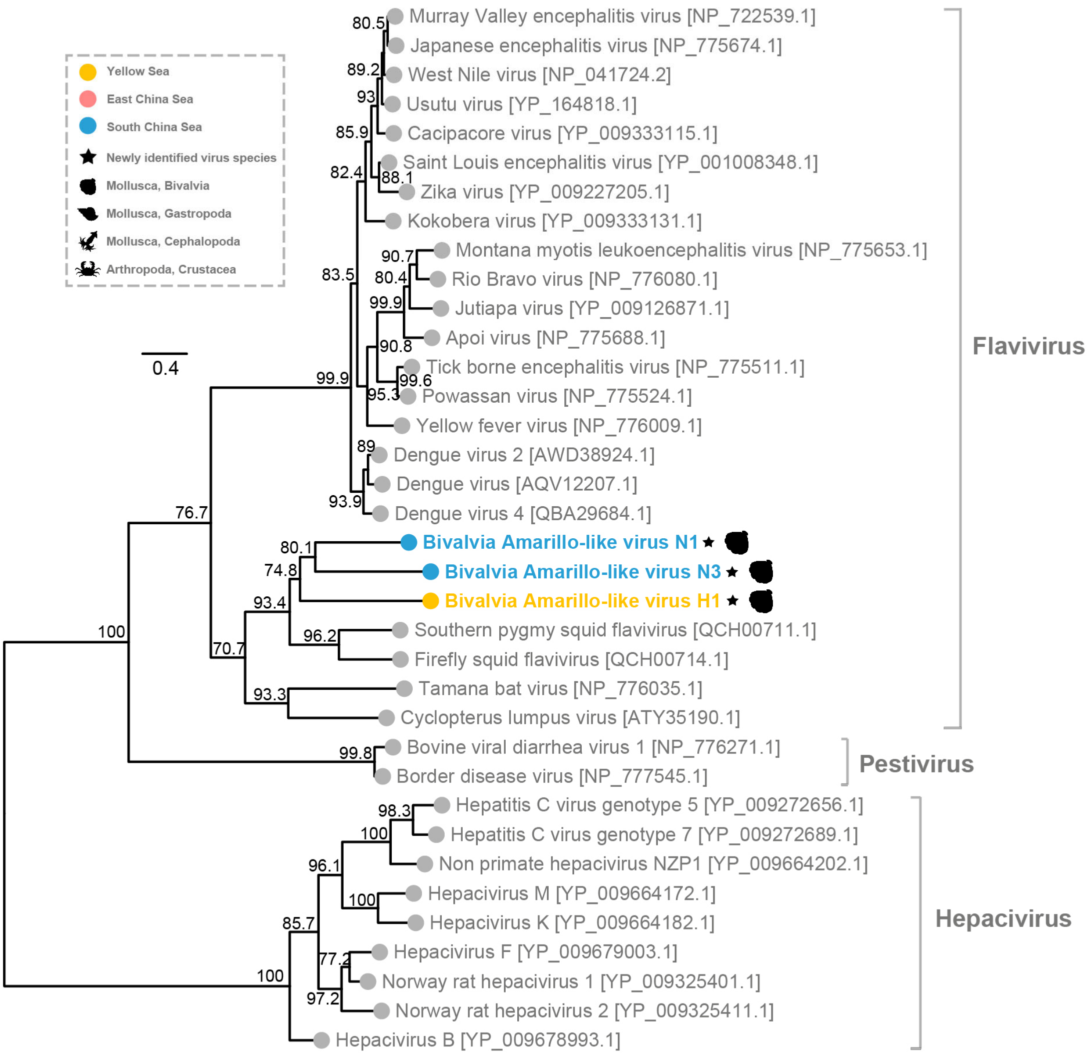


**Fig. S8 Phylogenetic trees of the Flaviviridae viruses, including the viruses identified in this study and related representative viruses.** The tree is inferred using amino acid sequences of the RdRp gene and flanking conserved domain. The tree is midpoint rooted for clarity only. Viruses identified in this study are denoted with a filled colored circle based on the area where their hosts were acquired, Yellow Sea (yellow), East China Sea (red), South China Sea (blue), while other representative publicly available viruses are denoted with a grey circle. The newly identified viruses are distinguished from previously published viruses with black solid asterisk at the end of their names. SH-aLRT supports less than 70% are not shown. The scale bar indicates the number of amino acid changes per site.

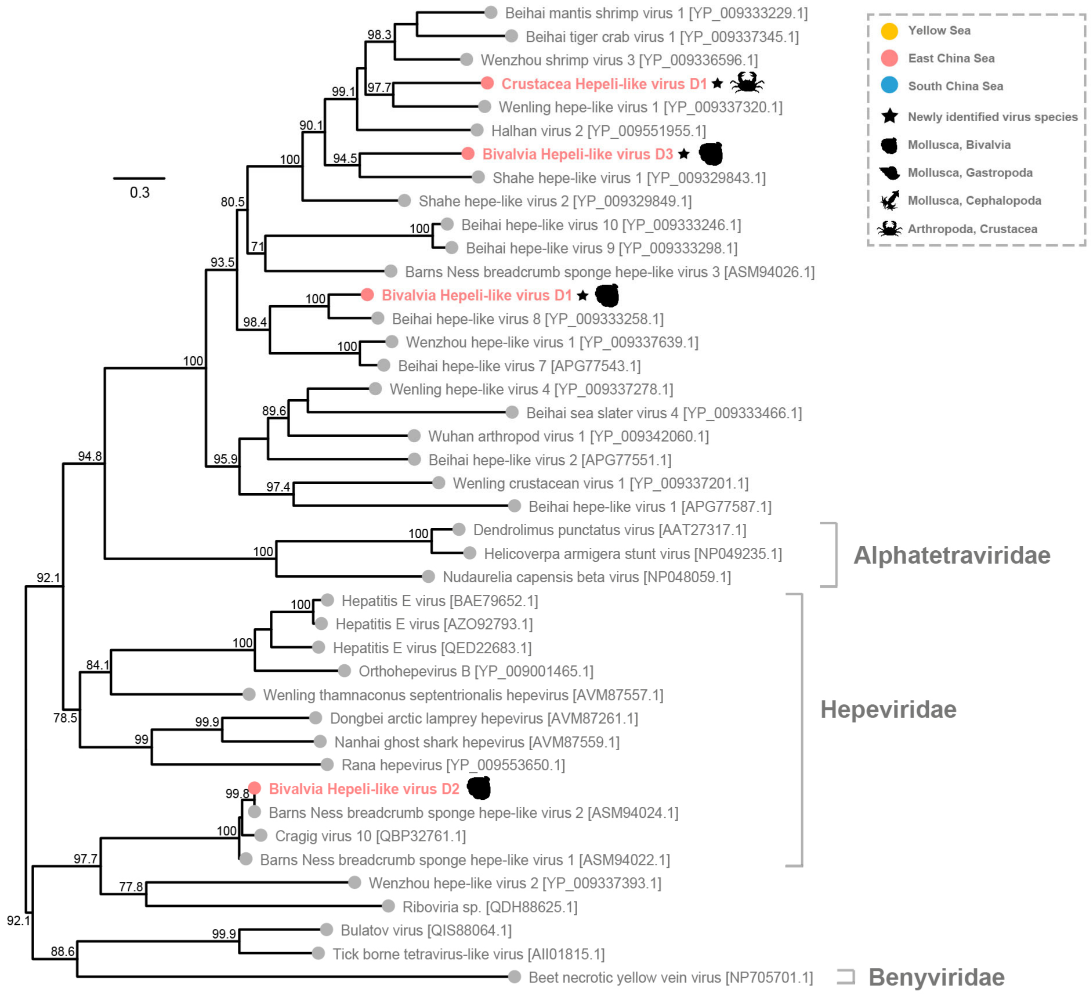


**Fig. S9 Phylogenetic trees of the Hepelivirales viruses, including the viruses identified in this study and related representative viruses.** The tree is inferred using amino acid sequences of the RdRp gene and flanking conserved domain. The tree is midpoint rooted for clarity only. Viruses identified in this study are denoted with a filled colored circle based on the area where their hosts were acquired, Yellow Sea (yellow), East China Sea (red), South China Sea (blue), while other representative publicly available viruses are denoted with a grey circle. The newly identified viruses are distinguished from previously published viruses with black solid asterisk at the end of their names. SH-aLRT supports less than 70% are not shown. The scale bar indicates the number of amino acid changes per site.

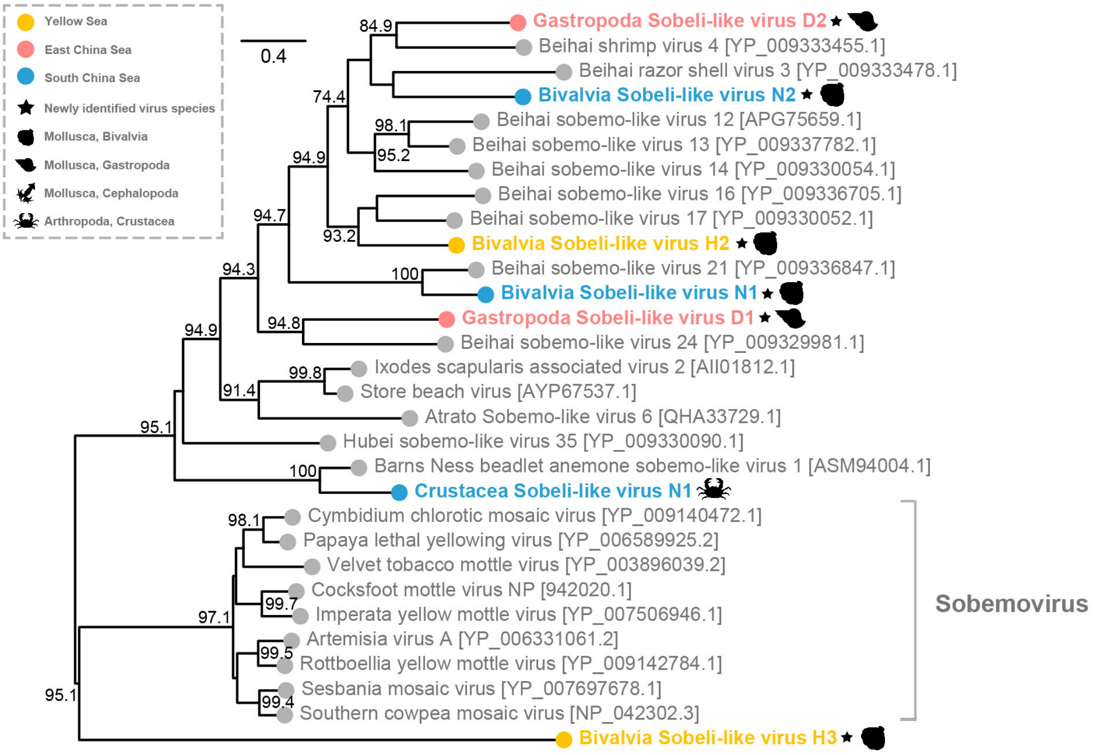


**Fig. S10 Phylogenetic trees of the Solemoviridae viruses, including the viruses identified in this study and related representative viruses.** The tree is inferred using amino acid sequences of the RdRp gene and flanking conserved domain. The tree is midpoint rooted for clarity only. Viruses identified in this study are denoted with a filled colored circle based on the area where their hosts were acquired, Yellow Sea (yellow), East China Sea (red), South China Sea (blue), while other representative publicly available viruses are denoted with a grey circle. The newly identified viruses are distinguished from previously published viruses with black solid asterisk at the end of their names. SH-aLRT supports less than 70% are not shown. The scale bar indicates the number of amino acid changes per site.

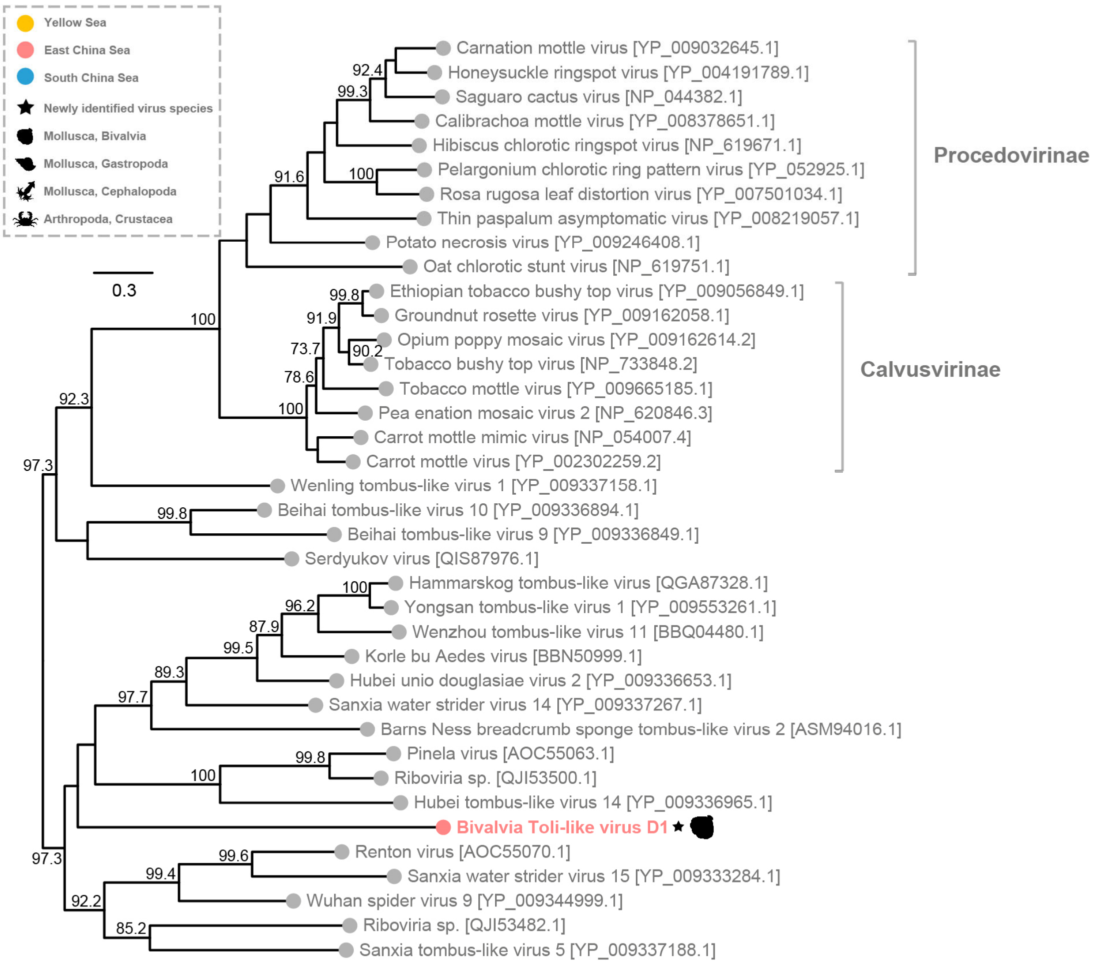


**Fig. S11 Phylogenetic trees of the Tombusviridae viruses, including the viruses identified in this study and related representative viruses.** The tree is inferred using amino acid sequences of the RdRp gene and flanking conserved domain. The tree is midpoint rooted for clarity only. Viruses identified in this study are denoted with a filled colored circle based on the area where their hosts were acquired, Yellow Sea (yellow), East China Sea (red), South China Sea (blue), while other representative publicly available viruses are denoted with a grey circle. The newly identified viruses are distinguished from previously published viruses with black solid asterisk at the end of their names. SH-aLRT supports less than 70% are not shown. The scale bar indicates the number of amino acid changes per site.

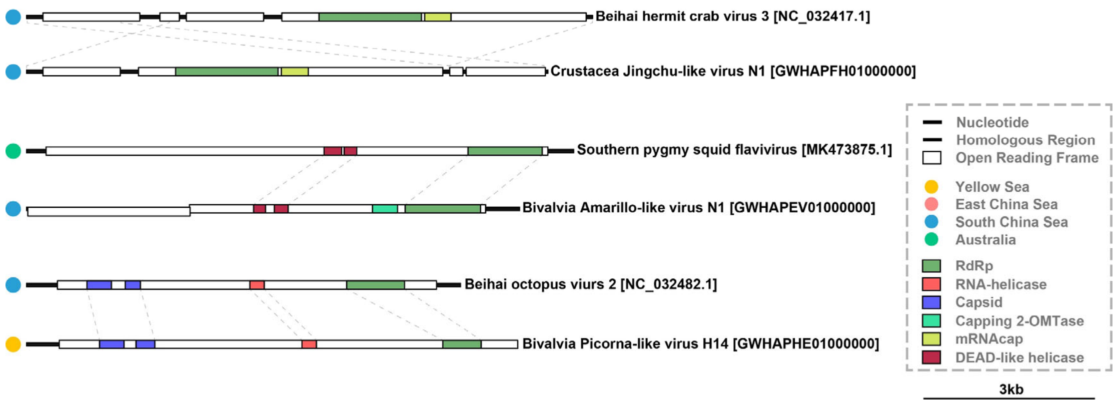


**Fig. S12 Genomic elasticity of representative viruses and the reference.** The contigs and genomes are drawn as lines and boxes to scale, respectively. The predicted regions that encode major functional proteins or domains are labelled with colored boxes. The filled colored circles indicate the area where their hosts were acquired, Yellow Sea (yellow), East China Sea (red), South China Sea (blue).
