## Supplementary material for "Viromes in Marine Ecosystems Reveal Remarkable Invertebrate RNA Virus Diversity": Host and geographic information and data output for each library of aquatic invertebrate samples

**Table S1 Host and geographic information and data output for each library of aquatic invertebrate samples**

| Library name | Sampling date | Host classification | Host species (blast nucleotide identity) | Tissue | No. of individuals | Location (city, province) | No. of reads | No. of viruses found |
|---|---|---|---|---|---|---|---|---|
| East China Sea Library 1 | Apr, 2019 | Arthropoda, Crustacea | *Melicertus canaliculatus (99%)* | Liverpancreas, gill | 12 | Zhoushan, Zhejiang | 44,295,625 | 3 |
| East China Sea Library 3 | Apr, 2019 | Arthropoda, Crustacea | *Litopenaeus vannamei (100%)* | Liverpancreas, gill | 12 | Zhoushan, Zhejiang | 53,761,512 | 7 |
| East China Sea Library 4 | Apr, 2019 | Arthropoda, Crustacea | *Exopalaemon carinicauda (99%)* | Liverpancreas, gill | 12 | Zhoushan, Zhejiang | 47,370,250 | 2 |
| East China Sea Library 5 | Apr, 2019 | Mollusca, Bivalvia | *Mytilus coruscus (100%)* | Liver and gill | 12 | Zhoushan, Zhejiang | 49,565,413 | 1 |
| East China Sea Library 6 | Apr, 2019 | Arthropoda, Hexanauplia | *Megabalanus volcano (99%)* | Individual | 12 | Zhoushan, Zhejiang | 40,932,034 | 5 |
| East China Sea Library 7 | Apr, 2019 | Mollusca, Gastropoda | *Bullacta exarata (99%)* | Individual | 12 | Zhoushan, Zhejiang | 40,158,578 | 6 |
| East China Sea Library 8-1 | Apr, 2019 | Mollusca, Bivalvia | *Corbula amurensis (100%)* | Liver and gill | 12 | Zhoushan, Zhejiang | 41,318,913 | 20 |
| East China Sea Library 8-2 | Apr, 2019 | Mollusca, Bivalvia | *Moerella iridescens (98%)* | Liver and gill | 12 | Zhoushan, Zhejiang | 44,456,340 | 12 |
| East China Sea Library 11 | Apr, 2019 | Mollusca, Bivalvia | *Mya japonica (100%)* | Liver and gill | 12 | Zhoushan, Zhejiang | 41,655,327 | 2 |
| East China Sea Library 12 | Apr, 2019 | Mollusca, Bivalvia | *Crassostrea gigas (100%)* | Liver and gill | 12 | Zhoushan, Zhejiang | 43,804,863 | 4 |
| East China Sea Library 13 | Apr, 2019 | Mollusca, Bivalvia | *Cyclina sinensis (100%)* | Liver and gill | 12 | Zhoushan, Zhejiang | 31,624,508 | 9 |
| East China Sea Library 14 | Apr, 2019 | Mollusca, Bivalvia | *Mimachlamys nobilis (100%)* | Liver and gill | 12 | Zhoushan, Zhejiang | 37,065,174 | 4 |
| East China Sea Library 16 | Apr, 2019 | Mollusca, Bivalvia | *Mercenaria mercenaria (100%)* | Liver and gill | 12 | Zhoushan, Zhejiang | 34,827,772 | 1 |
| East China Sea Library 17 | Apr, 2019 | Mollusca, Gastropoda | *Fusinus longicaudus (99%)* | Individual | 12 | Zhoushan, Zhejiang | 34,949,818 | 7 |
| East China Sea Library 18 | Apr, 2019 | Mollusca, Bivalvia | *Scapharca subcrenata (100%)* | Liver and gill | 12 | Zhoushan, Zhejiang | 41,515,918 | 2 |
| East China Sea Library 19 | Apr, 2019 | Mollusca, Bivalvia | *Sinonovacula constricta (99%)* | Liver and gill | 12 | Zhoushan, Zhejiang | 36,665,257 | 26 |
| East China Sea Library 20 | Apr, 2019 | Mollusca, Bivalvia | *Ruditapes philippinarum (99%)* | Liver and gill | 12 | Zhoushan, Zhejiang | 32,435,144 | 20 |
| East China Sea Library 21 | Apr, 2019 | Mollusca, Bivalvia | *Meretrix meretrix (100%)* | Liver and gill | 12 | Zhoushan, Zhejiang | 35,524,821 | 8 |
| East China Sea Library 22 | Apr, 2019 | Mollusca, Bivalvia | *Mercenaria mercenaria (100%)* | Liver and gill | 12 | Zhoushan, Zhejiang | 28,501,148 | 7 |
| East China Sea Library 25 | Apr, 2019 | Mollusca, Gastropoda | *Omphalius rusticus (99%)* | Individual | 12 | Zhoushan, Zhejiang | 39,941,909 | 8 |
| East China Sea Library 26 | Apr, 2019 | Mollusca, Gastropoda | *Monodonta labio (100%)* | Individual | 12 | Zhoushan, Zhejiang | 29,476,988 | 9 |
| East China Sea Library 27 | Apr, 2019 | Mollusca, Gastropoda | *Bufonaria rana (100%)* | Individual | 12 | Zhoushan, Zhejiang | 28,696,315 | 9 |
| South China Sea Library BH05 | Dec, 2019 | Mollusca, Bivalvia | *Mimachlamys nobilis (99%)* | Liver and gill | 12 | Beihai, Guangxi | 39,051,204 | 5 |
| South China Sea Library BH06 | Dec, 2019 | Mollusca, Bivalvia | *Venerupis philippinarum (99%)* | Liver and gill | 12 | Beihai, Guangxi | 38,093,765 | 5 |
| South China Sea Library BH09 | Dec, 2019 | Mollusca, Gastropoda | *Tectus pyramis (100%)* | Individual | 12 | Beihai, Guangxi | 29,718,126 | 3 |
| South China Sea Library BH10 | Dec, 2019 | Mollusca, Gastropoda | *Babylonia lutosa (99%)* | Individual | 12 | Beihai, Guangxi | 41,979,359 | 1 |
| South China Sea Library BH11 | Dec, 2019 | Arthropoda, Crustacea | *Aegla castro (85%)* | Liverpancreas, gill | 12 | Beihai, Guangxi | 37,401,824 | 3 |
| South China Sea Library BH12 | Dec, 2019 | Mollusca, Gastropoda | *Nerita albicilla (99%)* | Individual | 12 | Beihai, Guangxi | 43,054,485 | 5 |
| South China Sea Library BH13 | Dec, 2019 | Mollusca, Gastropoda | *Ergalatax margariticola (99%)* | Individual | 12 | Beihai, Guangxi | 45,606,534 | 5 |
| South China Sea Library G06 | Jun, 2019 | Arthropoda, Crustacea | *Penaeus monodon (99%)* | Liverpancreas, gill | 12 | Guangzhou, Guangdong | 32,374,573 | 6 |

| | | | | | | | | |
|---|---|---|---|---|---|---|---|---|
| South China Sea Library G07 | Jun, 2019 | Arthropoda, Crustacea | *Litopenaeus vannamei (94%)* | Liverpancreas, gill | 12 | Guangzhou, Guangdong | 52,372,546 | 4 |
| South China Sea Library G08 | Jun, 2019 | Arthropoda, Crustacea | *Charybdis feriata (100%)* | Liverpancreas, gill | 12 | Guangzhou, Guangdong | 55,486,394 | 1 |
| South China Sea Library G10 | Jun, 2019 | Arthropoda, Crustacea | *Scylla olivacea (100%)* | Liverpancreas, gill | 12 | Guangzhou, Guangdong | 57,837,428 | 5 |
| South China Sea Library G11 | Jun, 2019 | Mollusca, Gastropoda | *Babylonia spirata (100%)* | Individual | 12 | Guangzhou, Guangdong | 29,723,495 | 6 |
| South China Sea Library G12 | Jun, 2019 | Mollusca, Bivalvia | *Meretrix lyrata (100%)* | Liver and gill | 12 | Guangzhou, Guangdong | 47,331,867 | 7 |
| South China Sea Library G13 | Jun, 2019 | Mollusca, Bivalvia | *Mizuhopecten yessoensis (99%)* | Liver and gill | 12 | Guangzhou, Guangdong | 64,776,106 | 3 |
| South China Sea Library G14 | Jun, 2019 | Mollusca, Bivalvia | *Venerupis philippinarum (99%)* | Liver and gill | 12 | Guangzhou, Guangdong | 50,432,062 | 15 |
| South China Sea Library ZH03 | Dec, 2019 | Arthropoda, Crustacea | *Metapenaeus ensis (99%)* | Liverpancreas, gill | 12 | Zhuhai, Guangdong | 42,792,152 | 3 |
| South China Sea Library ZH04 | Dec, 2019 | Arthropoda, Crustacea | *Parapenaeopsis tenella (84%)* | Liverpancreas, gill | 12 | Zhuhai, Guangdong | 38,821,790 | 14 |
| South China Sea Library ZH05 | Dec, 2019 | Arthropoda, Crustacea | *Oratosquilla oratoria (100%)* | Liverpancreas, gill | 12 | Zhuhai, Guangdong | 53,506,132 | 2 |
| South China Sea Library ZH06 | Dec, 2019 | Arthropoda, Crustacea | *Metapenaeus affinis (99%)* | Liverpancreas, gill | 12 | Zhuhai, Guangdong | 38,805,810 | 3 |
| South China Sea Library ZH07 | Dec, 2019 | Mollusca, Bivalvia | *Solen lamarckii (99%)* | Liver and gill | 12 | Zhuhai, Guangdong | 31,248,906 | 7 |
| South China Sea Library ZH08 | Dec, 2019 | Mollusca, Bivalvia | *Meretrix petechialis (99%)* | Liver and gill | 12 | Zhuhai, Guangdong | 39,849,402 | 6 |
| South China Sea Library ZH09 | Dec, 2019 | Mollusca, Bivalvia | *Lutraria arcuata (99%)* | Liver and gill | 12 | Zhuhai, Guangdong | 30,450,217 | 1 |
| South China Sea Library ZH10 | Dec, 2019 | Mollusca, Bivalvia | *Paphia undulata (100%)* | Liver and gill | 12 | Zhuhai, Guangdong | 35,725,446 | 7 |
| South China Sea Library ZH11 | Dec, 2019 | Mollusca, Bivalvia | *Mytilus coruscus (100%)* | Liver and gill | 12 | Zhuhai, Guangdong | 38,272,297 | 3 |
| South China Sea Library ZH13 | Dec, 2019 | Mollusca, Bivalvia | *Meretrix lyrata (100%)* | Liver and gill | 12 | Zhuhai, Guangdong | 41,841,201 | 2 |
| South China Sea Library ZH14 | Dec, 2019 | Mollusca, Bivalvia | *Venerupis philippinarum (99%)* | Liver and gill | 12 | Zhuhai, Guangdong | 34,071,956 | 16 |
| Yellow Sea Library H01 | May, 2019 | Mollusca, Gastropoda | *Haliotis discus hannai (99%)* | Individual | 12 | Qingdao, Shandong | 32,781,450 | 2 |
| Yellow Sea Library H02 | May, 2019 | Mollusca, Bivalvia | *Solen grandis (85%)* | Liver and gill | 12 | Qingdao, Shandong | 28,787,966 | 11 |
| Yellow Sea Library H03 | May, 2019 | Mollusca, Bivalvia | *Chlamys farreri (99%)* | Liver and gill | 12 | Qingdao, Shandong | 34,028,189 | 3 |
| Yellow Sea Library H04 | May, 2019 | Arthropoda, Crustacea | *Oratosquilla oratoria (100%)* | Liverpancreas, gill | 12 | Qingdao, Shandong | 45,266,480 | 5 |
| Yellow Sea Library H05 | May, 2019 | Mollusca, Cephalopoda | *Doryteuthis gahi (100%)* | Gill | 12 | Qingdao, Shandong | 46,441,631 | 4 |
| Yellow Sea Library H06 | May, 2019 | Mollusca, Cephalopoda | *Octopus ocellatus (99%)* | Gill | 12 | Qingdao, Shandong | 36,941,115 | 2 |
| Yellow Sea Library H07 | May, 2019 | Mollusca, Bivalvia | *Venerupis philippinarum (99%)* | Liver and gill | 12 | Qingdao, Shandong | 33,949,947 | 13 |
| Yellow Sea Library H09 | May, 2019 | Mollusca, Bivalvia | *Scapharca subcrenata (97%)* | Liver and gill | 12 | Qingdao, Shandong | 44,213,243 | 6 |
| Yellow Sea Library H13 | May, 2019 | Mollusca, Gastropoda | *Turritella bacillum (99%)* | Individual | 12 | Qingdao, Shandong | 30,722,697 | 1 |
| Yellow Sea Library H14 | May, 2019 | Annelida, Polychaeta | *Urechis unicinctus (99%)* | Gill | 12 | Qingdao, Shandong | 36,797,636 | 6 |
