## Supplementary material for "Viromes in Marine Ecosystems Reveal Remarkable Invertebrate RNA Virus Diversity": The information about viruses identified in this study

### Table S2 The information about viruses identified in this study

| Library | Contig ID | Length | Order | Family | BioProject | BioSample | Accession |
|---|---|---|---|---|---|---|---|
| East China Sea Library 13 | 13_TRINITY_DN106217_c0_g2_i1 | 1049 | Picornavirales | Marnaviridae | PRJCA003705 | SAMC253645 | GWHAOXV01000000 |
| East China Sea Library 13 | 13_TRINITY_DN10961_c0_g1_i4 | 3876 | Picornavirales | Marnaviridae | PRJCA003705 | SAMC253646 | GWHAOXW01000000 |
| East China Sea Library 13 | 13_TRINITY_DN1254_c0_g1_i7 | 1278 | Picornavirales | Marnaviridae | PRJCA003705 | SAMC253647 | GWHAOXX01000000 |
| East China Sea Library 13 | 13_TRINITY_DN1723_c0_g1_i6 | 2103 | Picornavirales | Dicistroviridae | PRJCA003705 | SAMC253648 | GWHAOXY01000000 |
| East China Sea Library 13 | 13_TRINITY_DN19948_c0_g1_i16 | 5254 | Unclassified | Unclassified | PRJCA003705 | SAMC253649 | GWHAOXZ01000000 |
| East China Sea Library 13 | 13_TRINITY_DN34518_c0_g1_i2 | 1113 | Picornavirales | Dicistroviridae | PRJCA003705 | SAMC253650 | GWHAOYA01000000 |
| East China Sea Library 13 | 13_TRINITY_DN43256_c0_g2_i1 | 935 | Picornavirales | Dicistroviridae | PRJCA003705 | SAMC253651 | GWHAOYB01000000 |
| East China Sea Library 13 | 13_TRINITY_DN7096_c0_g1_i1 | 1060 | Picornavirales | Unclassified | PRJCA003705 | SAMC253652 | GWHAOYC01000000 |
| East China Sea Library 13 | 13_TRINITY_DN7205_c0_g1_i6 | 894 | Picornavirales | Dicistroviridae | PRJCA003705 | SAMC253653 | GWHAOYD01000000 |
| East China Sea Library 14 | 14_TRINITY_DN1066_c0_g1_i4 | 3511 | Durnavirales | Picobirnaviridae | PRJCA003705 | SAMC253654 | GWHAOYE01000000 |
| East China Sea Library 14 | 14_TRINITY_DN127071_c0_g1_i1 | 831 | Picornavirales | Unclassified | PRJCA003705 | SAMC253655 | GWHAOYF01000000 |
| East China Sea Library 14 | 14_TRINITY_DN2705_c0_g1_i3 | 1728 | Durnavirales | Picobirnaviridae | PRJCA003705 | SAMC253656 | GWHAOYG01000000 |
| East China Sea Library 14 | 14_TRINITY_DN51343_c0_g2_i1 | 866 | Picornavirales | Marnaviridae | PRJCA003705 | SAMC253657 | GWHAOYH01000000 |
| Yellow Sea Library H07 | H07_TRINITY_DN27_c0_g1_i1 | 11286 | Picornavirales | Unclassified | PRJCA003705 | SAMC253888 | GWHAPHE01000000 |
| East China Sea Library 17 | 17_TRINITY_DN105295_c0_g1_i1 | 802 | Picornavirales | Unclassified | PRJCA003705 | SAMC253659 | GWHAOYJ01000000 |
| South China Sea Library G07 | G07_TRINITY_DN120_c0_g1_i4 | 10343 | Picornavirales | Unclassified | PRJCA003705 | SAMC253858 | GWHAPGA01000000 |
| East China Sea Library 19 | 19_TRINITY_DN182296_c0_g1_i1 | 1000 | Durnavirales | Picobirnaviridae | PRJCA003705 | SAMC253661 | GWHAOYL01000000 |
| East China Sea Library 8-1 | 8-1_TRINITY_DN23155_c0_g1_i1 | 8924 | Picornavirales | Marnaviridae | PRJCA003705 | SAMC253982 | GWHAPKU01000000 |
| South China Sea Library ZH06 | ZH06_TRINITY_DN546_c0_g1_i1 | 8788 | Picornavirales | Unclassified | PRJCA003705 | SAMC253800 | GWHAPDU01000000 |
| East China Sea Library 4 | 4_TRINITY_DN612_c0_g1_i2 | 8365 | Picornavirales | Unclassified | PRJCA003705 | SAMC253749 | GWHAPBV01000000 |
| East China Sea Library 19 | 19_TRINITY_DN3718_c0_g1_i4 | 1014 | Picornavirales | Marnaviridae | PRJCA003705 | SAMC253665 | GWHAOYP01000000 |
| East China Sea Library 19 | 19_TRINITY_DN39273_c0_g1_i6 | 952 | Picornavirales | Marnaviridae | PRJCA003705 | SAMC253666 | GWHAOYQ01000000 |
| East China Sea Library 19 | 19_TRINITY_DN5_c0_g1_i9 | 3090 | Durnavirales | Picobirnaviridae | PRJCA003705 | SAMC253667 | GWHAOYR01000000 |

| | | | | | | | |
|---|---|---|---|---|---|---|---|
| East China Sea Library 19 | 19_TRINITY_DN58075_c0_g1_i7 | 2075 | Picornavirales | Marnaviridae | PRJCA003705 | SAMC253668 | GWHAOYS01000000 |
| East China Sea Library 19 | 19_TRINITY_DN58808_c0_g1_i1 | 1550 | Picornavirales | Unclassified | PRJCA003705 | SAMC253669 | GWHAOYT01000000 |
| East China Sea Library 8-1 | 8-1_TRINITY_DN62_c0_g1_i1 | 8339 | Picornavirales | Polycipiviridae | PRJCA003705 | SAMC253912 | GWHAPIC01000000 |
| East China Sea Library 25 | 25_TRINITY_DN1564_c0_g1_i5 | 8299 | Picornavirales | Iflaviridae | PRJCA003705 | SAMC253720 | GWHAPAS01000000 |
| East China Sea Library 8-1 | 8-1_TRINITY_DN46313_c0_g1_i2 | 8178 | Picornavirales | Unclassified | PRJCA003705 | SAMC253910 | GWHAPIA01000000 |
| East China Sea Library 19 | 19_TRINITY_DN68127_c0_g1_i2 | 817 | Picornavirales | Dicistroviridae | PRJCA003705 | SAMC253673 | GWHAOYX01000000 |
| East China Sea Library 19 | 19_TRINITY_DN71737_c0_g1_i2 | 1685 | Picornavirales | Marnaviridae | PRJCA003705 | SAMC253674 | GWHAOYY01000000 |
| East China Sea Library 19 | 19_TRINITY_DN72857_c0_g1_i1 | 1646 | Picornavirales | Unclassified | PRJCA003705 | SAMC253675 | GWHAOYZ01000000 |
| East China Sea Library 19 | 19_TRINITY_DN3173_c0_g1_i4 | 6463 | Picornavirales | Unclassified | PRJCA003705 | SAMC253664 | GWHAOYO01000000 |
| East China Sea Library 19 | 19_TRINITY_DN894_c0_g1_i6 | 2076 | Unclassified | Unclassified | PRJCA003705 | SAMC253677 | GWHAOZB01000000 |
| East China Sea Library 21 | 21_TRINITY_DN3628_c0_g1_i5 | 6277 | Picornavirales | Polycipiviridae | PRJCA003705 | SAMC253703 | GWHAPAB01000000 |
| East China Sea Library 19 | 19_TRINITY_DN9125_c0_g1_i8 | 1828 | Picornavirales | Marnaviridae | PRJCA003705 | SAMC253679 | GWHAOZD01000000 |
| East China Sea Library 20 | 20_TRINITY_DN10818_c0_g1_i3 | 1943 | Tolivirales | Tombusviridae | PRJCA003705 | SAMC253680 | GWHAOZE01000000 |
| East China Sea Library 20 | 20_TRINITY_DN119_c0_g1_i1 | 3552 | Durnavirales | Picobirnaviridae | PRJCA003705 | SAMC253681 | GWHAOZF01000000 |
| East China Sea Library 19 | 19_TRINITY_DN24867_c0_g1_i2 | 6152 | Picornavirales | Marnaviridae | PRJCA003705 | SAMC253663 | GWHAOYN01000000 |
| East China Sea Library 20 | 20_TRINITY_DN162235_c0_g1_i1 | 922 | Picornavirales | Unclassified | PRJCA003705 | SAMC253683 | GWHAOZH01000000 |
| East China Sea Library 20 | 20_TRINITY_DN1696_c0_g1_i7 | 6800 | Picornavirales | Unclassified | PRJCA003705 | SAMC253684 | GWHAOZI01000000 |
| East China Sea Library 20 | 20_TRINITY_DN195005_c0_g1_i1 | 1064 | Picornavirales | Marnaviridae | PRJCA003705 | SAMC253685 | GWHAOZJ01000000 |
| East China Sea Library 20 | 20_TRINITY_DN28415_c0_g1_i1 | 1523 | Picornavirales | Unclassified | PRJCA003705 | SAMC253686 | GWHAOZK01000000 |
| East China Sea Library 20 | 20_TRINITY_DN3500_c0_g1_i13 | 1441 | Picornavirales | Unclassified | PRJCA003705 | SAMC253687 | GWHAOZL01000000 |
| East China Sea Library 20 | 20_TRINITY_DN3714_c0_g1_i2 | 1431 | Durnavirales | Picobirnaviridae | PRJCA003705 | SAMC253688 | GWHAOZM01000000 |
| East China Sea Library 20 | 20_TRINITY_DN4026_c0_g1_i1 | 2367 | Picornavirales | Unclassified | PRJCA003705 | SAMC253689 | GWHAOZN01000000 |
| East China Sea Library 20 | 20_TRINITY_DN49_c0_g2_i1 | 2632 | Picornavirales | Marnaviridae | PRJCA003705 | SAMC253690 | GWHAOZO01000000 |
| East China Sea Library 20 | 20_TRINITY_DN493_c0_g1_i5 | 2497 | Picornavirales | Unclassified | PRJCA003705 | SAMC253691 | GWHAOZP01000000 |
| East China Sea Library 20 | 20_TRINITY_DN67_c0_g1_i4 | 1722 | Durnavirales | Picobirnaviridae | PRJCA003705 | SAMC253692 | GWHAOZQ01000000 |
| East China Sea Library 20 | 20_TRINITY_DN67819_c0_g1_i1 | 1625 | Picornavirales | Marnaviridae | PRJCA003705 | SAMC253693 | GWHAOZR01000000 |
| East China Sea Library 20 | 20_TRINITY_DN703_c0_g1_i9 | 898 | Picornavirales | Unclassified | PRJCA003705 | SAMC253694 | GWHAOZS01000000 |

| | | | | | | | |
|---|---|---|---|---|---|---|---|
| East China Sea Library 20 | 20_TRINITY_DN75302_c0_g2_i2 | 1082 | Picornavirales | Marnaviridae | PRJCA003705 | SAMC253695 | GWHAOZT01000000 |
| East China Sea Library 20 | 20_TRINITY_DN75695_c0_g1_i1 | 1498 | Picornavirales | Marnaviridae | PRJCA003705 | SAMC253696 | GWHAOZU01000000 |
| East China Sea Library 20 | 20_TRINITY_DN76783_c0_g1_i3 | 1569 | Picornavirales | Marnaviridae | PRJCA003705 | SAMC253697 | GWHAOZV01000000 |
| East China Sea Library 20 | 20_TRINITY_DN76892_c0_g1_i1 | 1565 | Durnavirales | Picobirnaviridae | PRJCA003705 | SAMC253698 | GWHAOZW01000000 |
| East China Sea Library 20 | 20_TRINITY_DN898_c0_g1_i16 | 1092 | Picornavirales | Marnaviridae | PRJCA003705 | SAMC253699 | GWHAOZX01000000 |
| East China Sea Library 21 | 21_TRINITY_DN1218_c0_g1_i8 | 4661 | Hepelivirales | Alphatetraviridae | PRJCA003705 | SAMC253700 | GWHAOZY01000000 |
| East China Sea Library 21 | 21_TRINITY_DN2044_c0_g1_i3 | 2500 | Picornavirales | Unclassified | PRJCA003705 | SAMC253701 | GWHAOZZ01000000 |
| East China Sea Library 21 | 21_TRINITY_DN23877_c0_g2_i2 | 1545 | Durnavirales | Picobirnaviridae | PRJCA003705 | SAMC253702 | GWHAPAA01000000 |
| South China Sea Library G12 | G12_TRINITY_DN1475_c0_g1_i1 | 5912 | Picornavirales | Unclassified | PRJCA003705 | SAMC253866 | GWHAPGI01000000 |
| East China Sea Library 21 | 21_TRINITY_DN45675_c0_g2_i1 | 827 | Picornavirales | Marnaviridae | PRJCA003705 | SAMC253704 | GWHAPAC01000000 |
| East China Sea Library 21 | 21_TRINITY_DN66916_c0_g1_i3 | 1088 | Picornavirales | Unclassified | PRJCA003705 | SAMC253705 | GWHAPAD01000000 |
| East China Sea Library 21 | 21_TRINITY_DN66916_c0_g2_i1 | 1531 | Picornavirales | Unclassified | PRJCA003705 | SAMC253706 | GWHAPAE01000000 |
| East China Sea Library 19 | 19_TRINITY_DN1578_c0_g1_i14 | 5724 | Picornavirales | Marnaviridae | PRJCA003705 | SAMC253660 | GWHAOYK01000000 |
| East China Sea Library 22 | 22_TRINITY_DN11118_c0_g1_i5 | 1157 | Picornavirales | Marnaviridae | PRJCA003705 | SAMC253708 | GWHAPAG01000000 |
| East China Sea Library 22 | 22_TRINITY_DN15190_c0_g1_i1 | 860 | Picornavirales | Marnaviridae | PRJCA003705 | SAMC253709 | GWHAPAH01000000 |
| South China Sea Library ZH03 | ZH03_TRINITY_DN1649_c0_g1_i3 | 5535 | Picornavirales | Dicistroviridae | PRJCA003705 | SAMC253779 | GWHAPCZ01000000 |
| East China Sea Library 22 | 22_TRINITY_DN28221_c0_g1_i1 | 1704 | Durnavirales | Picobirnaviridae | PRJCA003705 | SAMC253711 | GWHAPAJ01000000 |
| East China Sea Library 22 | 22_TRINITY_DN33039_c0_g1_i1 | 945 | Hepelivirales | Hepeviridae | PRJCA003705 | SAMC253712 | GWHAPAK01000000 |
| East China Sea Library 22 | 22_TRINITY_DN37492_c0_g1_i2 | 834 | Durnavirales | Picobirnaviridae | PRJCA003705 | SAMC253713 | GWHAPAL01000000 |
| Yellow Sea Library H02 | H02_TRINITY_DN1250_c0_g1_i3 | 4905 | Picornavirales | Unclassified | PRJCA003705 | SAMC253766 | GWHAPCM01000000 |
| East China Sea Library 25 | 25_TRINITY_DN1292_c0_g1_i7 | 3304 | Sobelivirales | Solemoviridae | PRJCA003705 | SAMC253715 | GWHAPAN01000000 |
| East China Sea Library 25 | 25_TRINITY_DN14243_c0_g2_i2 | 1365 | Picornavirales | Marnaviridae | PRJCA003705 | SAMC253716 | GWHAPAO01000000 |
| East China Sea Library 25 | 25_TRINITY_DN14243_c0_g3_i1 | 1688 | Picornavirales | Marnaviridae | PRJCA003705 | SAMC253717 | GWHAPAP01000000 |
| East China Sea Library 25 | 25_TRINITY_DN14399_c0_g1_i1 | 839 | Picornavirales | Unclassified | PRJCA003705 | SAMC253718 | GWHAPAQ01000000 |
| East China Sea Library 25 | 25_TRINITY_DN15438_c0_g3_i2 | 898 | Picornavirales | Marnaviridae | PRJCA003705 | SAMC253719 | GWHAPAR01000000 |
| South China Sea Library ZH04 | ZH04_TRINITY_DN1912_c0_g1_i1 | 4804 | Picornavirales | Unclassified | PRJCA003705 | SAMC253785 | GWHAPDF01000000 |
| East China Sea Library 25 | 25_TRINITY_DN4029_c0_g1_i1 | 1403 | Picornavirales | Unclassified | PRJCA003705 | SAMC253721 | GWHAPAT01000000 |

| | | | | | | | |
|---|---|---|---|---|---|---|---|
| East China Sea Library 25 | 25_TRINITY_DN5970_c0_g1_i8 | 1965 | Picornavirales | Dicistroviridae | PRJCA003705 | SAMC253722 | GWHAPAU01000000 |
| Yellow Sea Library H07 | H07_TRINITY_DN205_c0_g1_i10 | 4358 | Picornavirales | Unclassified | PRJCA003705 | SAMC253887 | GWHAPHD01000000 |
| East China Sea Library 26 | 26_TRINITY_DN32994_c0_g1_i1 | 1678 | Unclassified | Unclassified | PRJCA003705 | SAMC253724 | GWHAPAW01000000 |
| East China Sea Library 26 | 26_TRINITY_DN38383_c0_g1_i3 | 1452 | Picornavirales | Unclassified | PRJCA003705 | SAMC253725 | GWHAPAX01000000 |
| East China Sea Library 26 | 26_TRINITY_DN38671_c0_g2_i1 | 1410 | Picornavirales | Dicistroviridae | PRJCA003705 | SAMC253726 | GWHAPAY01000000 |
| East China Sea Library 26 | 26_TRINITY_DN45248_c0_g2_i2 | 836 | Picornavirales | Dicistroviridae | PRJCA003705 | SAMC253727 | GWHAPAZ01000000 |
| East China Sea Library 26 | 26_TRINITY_DN5270_c0_g1_i5 | 2307 | Picornavirales | Unclassified | PRJCA003705 | SAMC253728 | GWHAPBA01000000 |
| East China Sea Library 26 | 26_TRINITY_DN63407_c0_g1_i1 | 1572 | Picornavirales | Marnaviridae | PRJCA003705 | SAMC253729 | GWHAPBB01000000 |
| East China Sea Library 26 | 26_TRINITY_DN67898_c0_g1_i1 | 2615 | Unclassified | Unclassified | PRJCA003705 | SAMC253730 | GWHAPBC01000000 |
| East China Sea Library 26 | 26_TRINITY_DN7040_c0_g2_i4 | 4980 | Picornavirales | Unclassified | PRJCA003705 | SAMC253731 | GWHAPBD01000000 |
| East China Sea Library 27 | 27_TRINITY_DN1621_c0_g1_i12 | 1275 | Picornavirales | Dicistroviridae | PRJCA003705 | SAMC253732 | GWHAPBE01000000 |
| East China Sea Library 27 | 27_TRINITY_DN1864_c0_g1_i2 | 1375 | Picornavirales | Unclassified | PRJCA003705 | SAMC253733 | GWHAPBF01000000 |
| East China Sea Library 27 | 27_TRINITY_DN19287_c0_g1_i2 | 871 | Picornavirales | Dicistroviridae | PRJCA003705 | SAMC253734 | GWHAPBG01000000 |
| East China Sea Library 27 | 27_TRINITY_DN2607_c0_g1_i3 | 813 | Picornavirales | Dicistroviridae | PRJCA003705 | SAMC253735 | GWHAPBH01000000 |
| East China Sea Library 27 | 27_TRINITY_DN43834_c0_g1_i6 | 1041 | Picornavirales | Dicistroviridae | PRJCA003705 | SAMC253736 | GWHAPBI01000000 |
| East China Sea Library 27 | 27_TRINITY_DN5962_c0_g1_i1 | 1808 | Picornavirales | Dicistroviridae | PRJCA003705 | SAMC253737 | GWHAPBJ01000000 |
| East China Sea Library 27 | 27_TRINITY_DN5962_c1_g1_i4 | 1033 | Picornavirales | Unclassified | PRJCA003705 | SAMC253738 | GWHAPBK01000000 |
| East China Sea Library 27 | 27_TRINITY_DN61220_c0_g1_i1 | 827 | Unclassified | Unclassified | PRJCA003705 | SAMC253739 | GWHAPBL01000000 |
| East China Sea Library 27 | 27_TRINITY_DN75078_c0_g1_i4 | 1386 | Unclassified | Unclassified | PRJCA003705 | SAMC253740 | GWHAPBM01000000 |
| East China Sea Library 22 | 22_TRINITY_DN17786_c0_g1_i8 | 3933 | Picornavirales | Unclassified | PRJCA003705 | SAMC253710 | GWHAPAI01000000 |
| East China Sea Library 3 | 3_TRINITY_DN141_c0_g1_i11 | 4515 | Picornavirales | Unclassified | PRJCA003705 | SAMC253742 | GWHAPBO01000000 |
| East China Sea Library 3 | 3_TRINITY_DN19160_c0_g1_i1 | 2856 | Unclassified | Unclassified | PRJCA003705 | SAMC253743 | GWHAPBP01000000 |
| East China Sea Library 3 | 3_TRINITY_DN35153_c0_g1_i1 | 2919 | Picornavirales | Dicistroviridae | PRJCA003705 | SAMC253744 | GWHAPBQ01000000 |
| East China Sea Library 3 | 3_TRINITY_DN4609_c0_g1_i3 | 963 | Picornavirales | Unclassified | PRJCA003705 | SAMC253745 | GWHAPBR01000000 |
| East China Sea Library 3 | 3_TRINITY_DN61560_c0_g1_i1 | 1223 | Picornavirales | Marnaviridae | PRJCA003705 | SAMC253746 | GWHAPBS01000000 |
| East China Sea Library 3 | 3_TRINITY_DN84776_c0_g1_i1 | 954 | Picornavirales | Marnaviridae | PRJCA003705 | SAMC253747 | GWHAPBT01000000 |
| East China Sea Library 19 | 19_TRINITY_DN64758_c0_g1_i1 | 3714 | Picornavirales | Dicistroviridae | PRJCA003705 | SAMC253670 | GWHAOYU01000000 |

| | | | | | | | |
|---|---|---|---|---|---|---|---|
| East China Sea Library 21 | 21_TRINITY_DN8467_c0_g1_i2 | 3565 | Picornavirales | Unclassified | PRJCA003705 | SAMC253707 | GWHAPAF01000000 |
| East China Sea Library 5 | 5_TRINITY_DN64381_c0_g1_i1 | 977 | Durnavirales | Picobirnaviridae | PRJCA003705 | SAMC253750 | GWHAPBW01000000 |
| East China Sea Library 6 | 6_TRINITY_DN104099_c0_g1_i1 | 1827 | Picornavirales | Dicistroviridae | PRJCA003705 | SAMC253751 | GWHAPBX01000000 |
| East China Sea Library 6 | 6_TRINITY_DN13006_c0_g1_i1 | 12715 | Bunyavirales | Unclassified | PRJCA003705 | SAMC253752 | GWHAPBY01000000 |
| South China Sea Library G10 | G10_TRINITY_DN88610_c0_g1_i1 | 1143 | Picornavirales | Dicistroviridae | PRJCA003705 | SAMC253753 | GWHAPBZ01000000 |
| South China Sea Library G11 | G11_TRINITY_DN111_c0_g1_i7 | 5396 | Picornavirales | Dicistroviridae | PRJCA003705 | SAMC253754 | GWHAPCA01000000 |
| South China Sea Library G11 | G11_TRINITY_DN1500_c0_g1_i3 | 3127 | Picornavirales | Dicistroviridae | PRJCA003705 | SAMC253755 | GWHAPCB01000000 |
| South China Sea Library G11 | G11_TRINITY_DN1861_c0_g1_i1 | 1215 | Picornavirales | Dicistroviridae | PRJCA003705 | SAMC253756 | GWHAPCC01000000 |
| South China Sea Library G11 | G11_TRINITY_DN5796_c0_g1_i2 | 929 | Picornavirales | Dicistroviridae | PRJCA003705 | SAMC253757 | GWHAPCD01000000 |
| South China Sea Library ZH14 | ZH14_TRINITY_DN7333_c0_g1_i6 | 3417 | Picornavirales | Unclassified | PRJCA003705 | SAMC254004 | GWHAPLQ01000000 |
| East China Sea Library 26 | 26_TRINITY_DN16980_c0_g1_i8 | 3223 | Picornavirales | Unclassified | PRJCA003705 | SAMC253723 | GWHAPAV01000000 |
| South China Sea Library G14 | G14_TRINITY_DN33145_c0_g2_i1 | 2338 | Picornavirales | Dicistroviridae | PRJCA003705 | SAMC253760 | GWHAPCG01000000 |
| South China Sea Library G14 | G14_TRINITY_DN7_c0_g1_i3 | 1459 | Picornavirales | Unclassified | PRJCA003705 | SAMC253761 | GWHAPCH01000000 |
| South China Sea Library G14 | G14_TRINITY_DN9013_c0_g1_i2 | 942 | Picornavirales | Unclassified | PRJCA003705 | SAMC253762 | GWHAPCI01000000 |
| South China Sea Library G14 | G14_TRINITY_DN92252_c0_g2_i2 | 1049 | Picornavirales | Marnaviridae | PRJCA003705 | SAMC253763 | GWHAPCJ01000000 |
| Yellow Sea Library H01 | H01_TRINITY_DN119159_c0_g2_i | 910 | Picornavirales | Marnaviridae | PRJCA003705 | SAMC253764 | GWHAPCK01000000 |
| Yellow Sea Library H01 | H01_TRINITY_DN60146_c0_g5_i1 | 1263 | Picornavirales | Marnaviridae | PRJCA003705 | SAMC253765 | GWHAPCL01000000 |
| Yellow Sea Library H07 | H07_TRINITY_DN5785_c0_g1_i1 | 3214 | Picornavirales | Unclassified | PRJCA003705 | SAMC253944 | GWHAPJI01000000 |
| Yellow Sea Library H02 | H02_TRINITY_DN12567_c0_g1_i4 | 1436 | Durnavirales | Picobirnaviridae | PRJCA003705 | SAMC253767 | GWHAPCN01000000 |
| East China Sea Library 19 | 19_TRINITY_DN8971_c0_g1_i2 | 2749 | Picornavirales | Dicistroviridae | PRJCA003705 | SAMC253678 | GWHAOZC01000000 |
| Yellow Sea Library H02 | H02_TRINITY_DN4_c0_g1_i2 | 1637 | Durnavirales | Picobirnaviridae | PRJCA003705 | SAMC253769 | GWHAPCP01000000 |
| Yellow Sea Library H09 | H09_TRINITY_DN29861_c0_g1_i3 | 1613 | Durnavirales | Picobirnaviridae | PRJCA003705 | SAMC253770 | GWHAPCQ01000000 |
| Yellow Sea Library H09 | H09_TRINITY_DN55407_c0_g1_i4 | 3028 | Sobelivirales | Solemoviridae | PRJCA003705 | SAMC253771 | GWHAPCR01000000 |
| Yellow Sea Library H09 | H09_TRINITY_DN59781_c0_g1_i2 | 983 | Picornavirales | Marnaviridae | PRJCA003705 | SAMC253772 | GWHAPCS01000000 |
| Yellow Sea Library H14 | H14_TRINITY_DN15661_c0_g1_i2 | 1633 | Picornavirales | Unclassified | PRJCA003705 | SAMC253773 | GWHAPCT01000000 |
| Yellow Sea Library H14 | H14_TRINITY_DN20355_c0_g1_i3 | 1825 | Picornavirales | Unclassified | PRJCA003705 | SAMC253774 | GWHAPCU01000000 |
| Yellow Sea Library H14 | H14_TRINITY_DN20824_c0_g1_i1 | 971 | Picornavirales | Dicistroviridae | PRJCA003705 | SAMC253775 | GWHAPCV01000000 |

| | | | | | | | |
|---|---|---|---|---|---|---|---|
| Yellow Sea Library H14 | H14_TRINITY_DN2661_c0_g1_i1 | 1844 | Picornavirales | Unclassified | PRJCA003705 | SAMC253776 | GWHAPCW01000000 |
| Yellow Sea Library H14 | H14_TRINITY_DN32077_c0_g1_i1 | 800 | Picornavirales | Dicistroviridae | PRJCA003705 | SAMC253777 | GWHAPCX01000000 |
| Yellow Sea Library H14 | H14_TRINITY_DN32628_c0_g1_i2 | 1056 | Picornavirales | Marnaviridae | PRJCA003705 | SAMC253778 | GWHAPCY01000000 |
| South China Sea Library ZH14 | ZH14_TRINITY_DN20009_c0_g1_i | 2581 | Picornavirales | Dicistroviridae | PRJCA003705 | SAMC253808 | GWHAPEC01000000 |
| South China Sea Library ZH03 | ZH03_TRINITY_DN1695_c0_g1_i1 | 3005 | Picornavirales | Dicistroviridae | PRJCA003705 | SAMC253780 | GWHAPDA01000000 |
| South China Sea Library ZH03 | ZH03_TRINITY_DN1759_c0_g1_i6 | 1598 | Picornavirales | Unclassified | PRJCA003705 | SAMC253781 | GWHAPDB01000000 |
| South China Sea Library ZH04 | ZH04_TRINITY_DN12125_c0_g1_i | 1266 | Picornavirales | Unclassified | PRJCA003705 | SAMC253782 | GWHAPDC01000000 |
| South China Sea Library ZH04 | ZH04_TRINITY_DN14124_c0_g1_i | 1107 | Picornavirales | Unclassified | PRJCA003705 | SAMC253783 | GWHAPDD01000000 |
| South China Sea Library ZH04 | ZH04_TRINITY_DN17951_c0_g1_i | 1338 | Picornavirales | Unclassified | PRJCA003705 | SAMC253784 | GWHAPDE01000000 |
| South China Sea Library G14 | G14_TRINITY_DN125892_c0_g1_i | 2528 | Picornavirales | Unclassified | PRJCA003705 | SAMC253868 | GWHAPGK01000000 |
| South China Sea Library ZH04 | ZH04_TRINITY_DN25765_c0_g1_i | 883 | Picornavirales | Unclassified | PRJCA003705 | SAMC253786 | GWHAPDG01000000 |
| South China Sea Library ZH04 | ZH04_TRINITY_DN4100_c0_g1_i2 | 2778 | Picornavirales | Unclassified | PRJCA003705 | SAMC253787 | GWHAPDH01000000 |
| South China Sea Library ZH04 | ZH04_TRINITY_DN450_c0_g1_i9 | 3799 | Picornavirales | Dicistroviridae | PRJCA003705 | SAMC253788 | GWHAPDI01000000 |
| South China Sea Library BH06 | BH06_TRINITY_DN5672_c0_g1_i5 | 2335 | Picornavirales | Unclassified | PRJCA003705 | SAMC253833 | GWHAPFB01000000 |
| South China Sea Library ZH04 | ZH04_TRINITY_DN494_c0_g1_i5 | 1414 | Picornavirales | Dicistroviridae | PRJCA003705 | SAMC253790 | GWHAPDK01000000 |
| South China Sea Library ZH04 | ZH04_TRINITY_DN641_c0_g1_i16 | 2573 | Picornavirales | Dicistroviridae | PRJCA003705 | SAMC253791 | GWHAPDL01000000 |
| South China Sea Library ZH04 | ZH04_TRINITY_DN7212_c0_g1_i4 | 2111 | Picornavirales | Unclassified | PRJCA003705 | SAMC253792 | GWHAPDM01000000 |
| South China Sea Library ZH04 | ZH04_TRINITY_DN782_c0_g1_i5 | 3238 | Picornavirales | Dicistroviridae | PRJCA003705 | SAMC253793 | GWHAPDN01000000 |
| South China Sea Library ZH04 | ZH04_TRINITY_DN903_c0_g1_i5 | 1059 | Picornavirales | Dicistroviridae | PRJCA003705 | SAMC253794 | GWHAPDO01000000 |
| South China Sea Library ZH04 | ZH04_TRINITY_DN960_c0_g1_i9 | 1154 | Picornavirales | Dicistroviridae | PRJCA003705 | SAMC253795 | GWHAPDP01000000 |
| South China Sea Library ZH05 | ZH05_TRINITY_DN22607_c0_g1_i | 2322 | Picornavirales | Unclassified | PRJCA003705 | SAMC253796 | GWHAPDQ01000000 |
| South China Sea Library ZH05 | ZH05_TRINITY_DN3251_c0_g1_i5 | 2243 | Picornavirales | Unclassified | PRJCA003705 | SAMC253797 | GWHAPDR01000000 |
| South China Sea Library ZH06 | ZH06_TRINITY_DN18407_c0_g1_i | 1107 | Picornavirales | Unclassified | PRJCA003705 | SAMC253798 | GWHAPDS01000000 |
| South China Sea Library ZH06 | ZH06_TRINITY_DN2721_c0_g1_i9 | 1429 | Unclassified | Unclassified | PRJCA003705 | SAMC253799 | GWHAPDT01000000 |
| South China Sea Library G14 | G14_TRINITY_DN1548_c0_g1_i3 | 2275 | Picornavirales | Unclassified | PRJCA003705 | SAMC253933 | GWHAPIX01000000 |
| South China Sea Library ZH07 | ZH07_TRINITY_DN11327_c0_g1_i | 3348 | Amarillovirales | Flaviviridae | PRJCA003705 | SAMC253801 | GWHAPDV01000000 |
| South China Sea Library ZH10 | ZH10_TRINITY_DN563_c0_g1_i6 | 1978 | Tolivirales | Tombusviridae | PRJCA003705 | SAMC253802 | GWHAPDW01000000 |

| | | | | | | | |
|---|---|---|---|---|---|---|---|
| South China Sea Library ZH10 | ZH10_TRINITY_DN79622_c0_g1_i | 854 | Picornavirales | Unclassified | PRJCA003705 | SAMC253803 | GWHAPDX01000000 |
| South China Sea Library ZH13 | ZH13_TRINITY_DN28503_c0_g1_i | 2559 | Picornavirales | Dicistroviridae | PRJCA003705 | SAMC253804 | GWHAPDY01000000 |
| South China Sea Library ZH13 | ZH13_TRINITY_DN92793_c0_g1_i | 1630 | Durnavirales | Picobirnaviridae | PRJCA003705 | SAMC253805 | GWHAPDZ01000000 |
| South China Sea Library ZH14 | ZH14_TRINITY_DN10501_c0_g1_i | 4207 | Picornavirales | Unclassified | PRJCA003705 | SAMC253806 | GWHAPEA01000000 |
| South China Sea Library ZH14 | ZH14_TRINITY_DN10919_c0_g1_i | 2219 | Picornavirales | Dicistroviridae | PRJCA003705 | SAMC253807 | GWHAPEB01000000 |
| Yellow Sea Library H07 | H07_TRINITY_DN579_c0_g1_i13 | 2211 | Picornavirales | Unclassified | PRJCA003705 | SAMC253945 | GWHAPJJ01000000 |
| South China Sea Library ZH14 | ZH14_TRINITY_DN2139_c0_g1_i7 | 870 | Unclassified | Unclassified | PRJCA003705 | SAMC253809 | GWHAPED01000000 |
| South China Sea Library ZH14 | ZH14_TRINITY_DN2243_c0_g1_i6 | 2526 | Picornavirales | Unclassified | PRJCA003705 | SAMC253810 | GWHAPEE01000000 |
| South China Sea Library ZH14 | ZH14_TRINITY_DN23605_c0_g2_i | 897 | Picornavirales | Dicistroviridae | PRJCA003705 | SAMC253811 | GWHAPEF01000000 |
| South China Sea Library ZH14 | ZH14_TRINITY_DN248443_c0_g1_ | 808 | Unclassified | Unclassified | PRJCA003705 | SAMC253812 | GWHAPEG01000000 |
| South China Sea Library ZH14 | ZH14_TRINITY_DN32447_c0_g1_i | 1420 | Picornavirales | Unclassified | PRJCA003705 | SAMC253813 | GWHAPEH01000000 |
| South China Sea Library ZH14 | ZH14_TRINITY_DN34716_c0_g1_i | 1649 | Tolivirales | Tombusviridae | PRJCA003705 | SAMC253814 | GWHAPEI01000000 |
| East China Sea Library 1 | 1_TRINITY_DN8232_c0_g1_i1 | 4982 | Hepelivirales | Alphatetraviridae | PRJCA003705 | SAMC253815 | GWHAPEJ01000000 |
| East China Sea Library 1 | 1_TRINITY_DN83828_c0_g1_i1 | 997 | Hepelivirales | Alphatetraviridae | PRJCA003705 | SAMC253816 | GWHAPEK01000000 |
| East China Sea Library 11 | 11_TRINITY_DN101233_c0_g1_i3 | 838 | Picornavirales | Marnaviridae | PRJCA003705 | SAMC253817 | GWHAPEL01000000 |
| East China Sea Library 12 | 12_TRINITY_DN27119_c0_g1_i17 | 2243 | Picornavirales | Marnaviridae | PRJCA003705 | SAMC253818 | GWHAPEM01000000 |
| East China Sea Library 12 | 12_TRINITY_DN47756_c0_g1_i2 | 1856 | Picornavirales | Unclassified | PRJCA003705 | SAMC253819 | GWHAPEN01000000 |
| East China Sea Library 12 | 12_TRINITY_DN80727_c0_g1_i4 | 1956 | Picornavirales | Unclassified | PRJCA003705 | SAMC253820 | GWHAPEO01000000 |
| East China Sea Library 19 | 19_TRINITY_DN10_c0_g1_i3 | 1094 | Picornavirales | Marnaviridae | PRJCA003705 | SAMC253821 | GWHAPEP01000000 |
| East China Sea Library 19 | 19_TRINITY_DN138578_c0_g1_i2 | 1157 | Picornavirales | Dicistroviridae | PRJCA003705 | SAMC253822 | GWHAPEQ01000000 |
| East China Sea Library 19 | 19_TRINITY_DN1449_c0_g1_i3 | 3088 | Picornavirales | Unclassified | PRJCA003705 | SAMC253823 | GWHAPER01000000 |
| East China Sea Library 19 | 19_TRINITY_DN1456_c0_g1_i5 | 2602 | Durnavirales | Picobirnaviridae | PRJCA003705 | SAMC253824 | GWHAPES01000000 |
| East China Sea Library 8-1 | 8-1_TRINITY_DN38244_c0_g1_i1 | 1296 | Picornavirales | Marnaviridae | PRJCA003705 | SAMC253825 | GWHAPET01000000 |
| East China Sea Library 8-2 | 8-2_TRINITY_DN148753_c0_g3_i1 | 1051 | Picornavirales | Marnaviridae | PRJCA003705 | SAMC253826 | GWHAPEU01000000 |
| South China Sea Library BH05 | BH05_TRINITY_DN2720_c0_g1_i2 | 11329 | Amarillovirales | Flaviviridae | PRJCA003705 | SAMC253827 | GWHAPEV01000000 |
| South China Sea Library BH05 | BH05_TRINITY_DN83599_c0_g1_i | 989 | Picornavirales | Marnaviridae | PRJCA003705 | SAMC253828 | GWHAPEW01000000 |
| South China Sea Library BH05 | BH05_TRINITY_DN9263_c0_g2_i1 | 893 | Picornavirales | Marnaviridae | PRJCA003705 | SAMC253829 | GWHAPEX01000000 |

| | | | | | | | |
|---|---|---|---|---|---|---|---|
| South China Sea Library BH06 | BH06_TRINITY_DN1764_c0_g1_i2 | 1134 | Picornavirales | Unclassified | PRJCA003705 | SAMC253830 | GWHAPEY01000000 |
| South China Sea Library BH06 | BH06_TRINITY_DN17841_c0_g1_i | 958 | Picornavirales | Unclassified | PRJCA003705 | SAMC253831 | GWHAPEZ01000000 |
| South China Sea Library BH06 | BH06_TRINITY_DN4704_c0_g1_i2 | 2144 | Picornavirales | Unclassified | PRJCA003705 | SAMC253832 | GWHAPFA01000000 |
| East China Sea Library 3 | 3_TRINITY_DN13688_c0_g1_i1 | 2115 | Picornavirales | Polycipiviridae | PRJCA003705 | SAMC253741 | GWHAPBN01000000 |
| South China Sea Library BH06 | BH06_TRINITY_DN89577_c0_g1_i | 3275 | Sobelivirales | Solemoviridae | PRJCA003705 | SAMC253834 | GWHAPFC01000000 |
| South China Sea Library BH09 | BH09_TRINITY_DN1818_c0_g1_i9 | 1488 | Picornavirales | Marnaviridae | PRJCA003705 | SAMC253835 | GWHAPFD01000000 |
| South China Sea Library BH09 | BH09_TRINITY_DN25791_c0_g1_i | 843 | Picornavirales | Marnaviridae | PRJCA003705 | SAMC253836 | GWHAPFE01000000 |
| South China Sea Library BH09 | BH09_TRINITY_DN30_c0_g1_i6 | 1143 | Picornavirales | Marnaviridae | PRJCA003705 | SAMC253837 | GWHAPFF01000000 |
| South China Sea Library BH10 | BH10_TRINITY_DN92075_c0_g1_i | 1058 | Tolivirales | Tombusviridae | PRJCA003705 | SAMC253838 | GWHAPFG01000000 |
| South China Sea Library BH11 | BH11_TRINITY_DN113_c0_g1_i4 | 11974 | Jingchuvirales | Chuviridae | PRJCA003705 | SAMC253839 | GWHAPFH01000000 |
| South China Sea Library BH11 | BH11_TRINITY_DN16288_c0_g1_i | 1293 | Durnavirales | Picobirnaviridae | PRJCA003705 | SAMC253840 | GWHAPFI01000000 |
| South China Sea Library BH11 | BH11_TRINITY_DN75_c0_g1_i12 | 2015 | Durnavirales | Picobirnaviridae | PRJCA003705 | SAMC253841 | GWHAPFJ01000000 |
| South China Sea Library BH12 | BH12_TRINITY_DN116276_c0_g1_ | 969 | Stellavirales | Astroviridae | PRJCA003705 | SAMC253842 | GWHAPFK01000000 |
| South China Sea Library BH12 | BH12_TRINITY_DN11840_c0_g1_i | 3027 | Picornavirales | Dicistroviridae | PRJCA003705 | SAMC253843 | GWHAPFL01000000 |
| South China Sea Library BH12 | BH12_TRINITY_DN38608_c0_g2_i | 836 | Picornavirales | Marnaviridae | PRJCA003705 | SAMC253844 | GWHAPFM01000000 |
| South China Sea Library BH12 | BH12_TRINITY_DN89012_c0_g1_i | 1014 | Picornavirales | Marnaviridae | PRJCA003705 | SAMC253845 | GWHAPFN01000000 |
| South China Sea Library BH12 | BH12_TRINITY_DN91413_c0_g1_i | 1382 | Sobelivirales | Solemoviridae | PRJCA003705 | SAMC253846 | GWHAPFO01000000 |
| South China Sea Library BH13 | BH13_TRINITY_DN15351_c0_g1_i | 2214 | Picornavirales | Unclassified | PRJCA003705 | SAMC253847 | GWHAPFP01000000 |
| South China Sea Library BH13 | BH13_TRINITY_DN16706_c0_g1_i | 1595 | Picornavirales | Unclassified | PRJCA003705 | SAMC253848 | GWHAPFQ01000000 |
| South China Sea Library BH13 | BH13_TRINITY_DN262263_c0_g1_ | 822 | Picornavirales | Dicistroviridae | PRJCA003705 | SAMC253849 | GWHAPFR01000000 |
| South China Sea Library BH13 | BH13_TRINITY_DN405_c0_g1_i3 | 2038 | Picornavirales | Marnaviridae | PRJCA003705 | SAMC253850 | GWHAPFS01000000 |
| South China Sea Library BH13 | BH13_TRINITY_DN48022_c0_g1_i | 993 | Durnavirales | Picobirnaviridae | PRJCA003705 | SAMC253851 | GWHAPFT01000000 |
| South China Sea Library G06 | G06_TRINITY_DN10475_c0_g1_i3 | 1448 | Picornavirales | Dicistroviridae | PRJCA003705 | SAMC253852 | GWHAPFU01000000 |
| South China Sea Library G06 | G06_TRINITY_DN4195_c0_g1_i3 | 1217 | Picornavirales | Dicistroviridae | PRJCA003705 | SAMC253853 | GWHAPFV01000000 |
| South China Sea Library G06 | G06_TRINITY_DN42055_c0_g1_i1 | 807 | Picornavirales | Dicistroviridae | PRJCA003705 | SAMC253854 | GWHAPFW01000000 |
| South China Sea Library G06 | G06_TRINITY_DN5219_c0_g1_i2 | 1131 | Picornavirales | Dicistroviridae | PRJCA003705 | SAMC253855 | GWHAPFX01000000 |
| South China Sea Library G06 | G06_TRINITY_DN705_c0_g1_i3 | 2676 | Sobelivirales | Solemoviridae | PRJCA003705 | SAMC253856 | GWHAPFY01000000 |

| | | | | | | | |
|---|---|---|---|---|---|---|---|
| South China Sea Library G06 | G06_TRINITY_DN8627_c0_g1_i1 | 897 | Picornavirales | Dicistroviridae | PRJCA003705 | SAMC253857 | GWHAPFZ01000000 |
| East China Sea Library 4 | 4_TRINITY_DN10052_c0_g1_i1 | 1854 | Picornavirales | Unclassified | PRJCA003705 | SAMC253748 | GWHAPBU01000000 |
| South China Sea Library G07 | G07_TRINITY_DN3356_c0_g1_i2 | 940 | Picornavirales | Unclassified | PRJCA003705 | SAMC253859 | GWHAPGB01000000 |
| South China Sea Library G07 | G07_TRINITY_DN3356_c0_g2_i2 | 1061 | Picornavirales | Unclassified | PRJCA003705 | SAMC253860 | GWHAPGC01000000 |
| South China Sea Library G10 | G10_TRINITY_DN3803_c0_g1_i5 | 1692 | Ghabrivirales | Totiviridae | PRJCA003705 | SAMC253861 | GWHAPGD01000000 |
| South China Sea Library G10 | G10_TRINITY_DN55839_c0_g1_i1 | 1000 | Picornavirales | Dicistroviridae | PRJCA003705 | SAMC253862 | GWHAPGE01000000 |
| South China Sea Library G11 | G11_TRINITY_DN11242_c0_g1_i1 | 1054 | Picornavirales | Unclassified | PRJCA003705 | SAMC253863 | GWHAPGF01000000 |
| South China Sea Library G11 | G11_TRINITY_DN1339_c0_g1_i3 | 1407 | Unclassified | Unclassified | PRJCA003705 | SAMC253864 | GWHAPGG01000000 |
| South China Sea Library G12 | G12_TRINITY_DN13497_c0_g1_i2 | 2007 | Picornavirales | Dicistroviridae | PRJCA003705 | SAMC253865 | GWHAPGH01000000 |
| South China Sea Library G14 | G14_TRINITY_DN128452_c0_g1_i | 1657 | Picornavirales | Marnaviridae | PRJCA003705 | SAMC253869 | GWHAPGL01000000 |
| South China Sea Library G12 | G12_TRINITY_DN36494_c0_g1_i1 | 1186 | Picornavirales | Marnaviridae | PRJCA003705 | SAMC253867 | GWHAPGJ01000000 |
| East China Sea Library 19 | 19_TRINITY_DN76323_c0_g1_i1 | 1584 | Picornavirales | Marnaviridae | PRJCA003705 | SAMC253676 | GWHAOZA01000000 |
| East China Sea Library 8-2 | 8-2_TRINITY_DN41441_c0_g1_i2 | 1562 | Picornavirales | Marnaviridae | PRJCA003705 | SAMC253922 | GWHAPIM01000000 |
| South China Sea Library G14 | G14_TRINITY_DN89352_c0_g2_i2 | 1425 | Picornavirales | Unclassified | PRJCA003705 | SAMC253870 | GWHAPGM01000000 |
| Yellow Sea Library H02 | H02_TRINITY_DN67462_c0_g1_i2 | 904 | Amarillovirales | Flaviviridae | PRJCA003705 | SAMC253871 | GWHAPGN01000000 |
| Yellow Sea Library H02 | H02_TRINITY_DN7318_c0_g1_i15 | 2403 | Picornavirales | Unclassified | PRJCA003705 | SAMC253872 | GWHAPGO01000000 |
| Yellow Sea Library H02 | H02_TRINITY_DN84_c0_g1_i3 | 4427 | Bunyavirales | Hantaviridae | PRJCA003705 | SAMC253873 | GWHAPGP01000000 |
| Yellow Sea Library H03 | H03_TRINITY_DN4623_c0_g1_i1 | 1639 | Durnavirales | Picobirnaviridae | PRJCA003705 | SAMC253874 | GWHAPGQ01000000 |
| Yellow Sea Library H03 | H03_TRINITY_DN72722_c0_g1_i1 | 889 | Durnavirales | Picobirnaviridae | PRJCA003705 | SAMC253875 | GWHAPGR01000000 |
| Yellow Sea Library H03 | H03_TRINITY_DN78573_c0_g1_i1 | 3468 | Durnavirales | Picobirnaviridae | PRJCA003705 | SAMC253876 | GWHAPGS01000000 |
| Yellow Sea Library H04 | H04_TRINITY_DN100462_c0_g1_i | 1093 | Ghabrivirales | Totiviridae | PRJCA003705 | SAMC253877 | GWHAPGT01000000 |
| Yellow Sea Library H04 | H04_TRINITY_DN202600_c0_g1_i | 9269 | Bunyavirales | Mypoviridae | PRJCA003705 | SAMC253878 | GWHAPGU01000000 |
| Yellow Sea Library H04 | H04_TRINITY_DN38222_c0_g1_i6 | 2089 | Ghabrivirales | Totiviridae | PRJCA003705 | SAMC253879 | GWHAPGV01000000 |
| Yellow Sea Library H04 | H04_TRINITY_DN993_c0_g1_i1 | 6494 | Ghabrivirales | Totiviridae | PRJCA003705 | SAMC253880 | GWHAPGW01000000 |
| Yellow Sea Library H05 | H05_TRINITY_DN1216_c0_g1_i1 | 845 | Durnavirales | Picobirnaviridae | PRJCA003705 | SAMC253881 | GWHAPGX01000000 |
| Yellow Sea Library H05 | H05_TRINITY_DN2042_c0_g1_i13 | 1068 | Unclassified | Unclassified | PRJCA003705 | SAMC253882 | GWHAPGY01000000 |
| Yellow Sea Library H05 | H05_TRINITY_DN3806_c0_g1_i3 | 1021 | Durnavirales | Partitiviridae | PRJCA003705 | SAMC253883 | GWHAPGZ01000000 |

| | | | | | | | |
|---|---|---|---|---|---|---|---|
| Yellow Sea Library H05 | H05_TRINITY_DN61_c0_g1_i10 | 2681 | Picornavirales | Dicistroviridae | PRJCA003705 | SAMC253884 | GWHAPHA01000000 |
| Yellow Sea Library H06 | H06_TRINITY_DN19025_c0_g2_i1 | 1159 | Mononegavirales | Rhabdoviridae | PRJCA003705 | SAMC253885 | GWHAPHB01000000 |
| Yellow Sea Library H06 | H06_TRINITY_DN2012_c0_g1_i1 | 1208 | Bunyavirales | Phenuiviridae | PRJCA003705 | SAMC253886 | GWHAPHC01000000 |
| East China Sea Library 22 | 22_TRINITY_DN5657_c0_g1_i7 | 1531 | Picornavirales | Iflaviridae | PRJCA003705 | SAMC253714 | GWHAPAM01000000 |
| South China Sea Library G14 | G14_TRINITY_DN1548_c1_g1_i3 | 1509 | Picornavirales | Unclassified | PRJCA003705 | SAMC253759 | GWHAPCF01000000 |
| Yellow Sea Library H07 | H07_TRINITY_DN338148_c0_g1_i | 1125 | Picornavirales | Unclassified | PRJCA003705 | SAMC253889 | GWHAPHF01000000 |
| Yellow Sea Library H07 | H07_TRINITY_DN339057_c0_g1_i | 1128 | Durnavirales | Picobirnaviridae | PRJCA003705 | SAMC253890 | GWHAPHG01000000 |
| Yellow Sea Library H07 | H07_TRINITY_DN3435_c0_g4_i4 | 1152 | Picornavirales | Unclassified | PRJCA003705 | SAMC253891 | GWHAPHH01000000 |
| Yellow Sea Library H09 | H09_TRINITY_DN40946_c0_g1_i1 | 6529 | Sobelivirales | Solemoviridae | PRJCA003705 | SAMC253892 | GWHAPHI01000000 |
| Yellow Sea Library H13 | H13_TRINITY_DN7_c0_g1_i3 | 1914 | Picornavirales | Marnaviridae | PRJCA003705 | SAMC253893 | GWHAPHJ01000000 |
| South China Sea Library ZH07 | ZH07_TRINITY_DN473_c0_g1_i9 | 2043 | Sobelivirales | Solemoviridae | PRJCA003705 | SAMC253894 | GWHAPHK01000000 |
| South China Sea Library ZH07 | ZH07_TRINITY_DN8577_c0_g1_i4 | 3727 | Bunyavirales | Hantaviridae | PRJCA003705 | SAMC253895 | GWHAPHL01000000 |
| South China Sea Library ZH08 | ZH08_TRINITY_DN79368_c0_g1_i | 909 | Picornavirales | Marnaviridae | PRJCA003705 | SAMC253896 | GWHAPHM01000000 |
| South China Sea Library ZH08 | ZH08_TRINITY_DN81541_c0_g1_i | 1551 | Picornavirales | Dicistroviridae | PRJCA003705 | SAMC253897 | GWHAPHN01000000 |
| South China Sea Library ZH09 | ZH09_TRINITY_DN73110_c0_g1_i | 1613 | Bunyavirales | Unclassified | PRJCA003705 | SAMC253898 | GWHAPHO01000000 |
| South China Sea Library ZH10 | ZH10_TRINITY_DN12413_c0_g1_i | 3585 | Picornavirales | Unclassified | PRJCA003705 | SAMC253899 | GWHAPHP01000000 |
| South China Sea Library ZH10 | ZH10_TRINITY_DN22369_c0_g1_i | 1409 | Picornavirales | Unclassified | PRJCA003705 | SAMC253900 | GWHAPHQ01000000 |
| South China Sea Library ZH10 | ZH10_TRINITY_DN2577_c0_g1_i5 | 941 | Picornavirales | Unclassified | PRJCA003705 | SAMC253901 | GWHAPHR01000000 |
| South China Sea Library ZH10 | ZH10_TRINITY_DN31476_c0_g1_i | 1224 | Picornavirales | Unclassified | PRJCA003705 | SAMC253902 | GWHAPHS01000000 |
| South China Sea Library ZH10 | ZH10_TRINITY_DN4534_c0_g1_i1 | 1721 | Picornavirales | Unclassified | PRJCA003705 | SAMC253903 | GWHAPHT01000000 |
| South China Sea Library ZH11 | ZH11_TRINITY_DN12131_c0_g3_i | 2037 | Picornavirales | Unclassified | PRJCA003705 | SAMC253904 | GWHAPHU01000000 |
| South China Sea Library ZH11 | ZH11_TRINITY_DN15550_c0_g1_i | 1169 | Durnavirales | Picobirnaviridae | PRJCA003705 | SAMC253905 | GWHAPHV01000000 |
| South China Sea Library ZH11 | ZH11_TRINITY_DN212836_c0_g1_ | 1258 | Ghabrivirales | Totiviridae | PRJCA003705 | SAMC253906 | GWHAPHW01000000 |
| East China Sea Library 19 | 19_TRINITY_DN125270_c0_g1_i2 | 820 | Picornavirales | Marnaviridae | PRJCA003705 | SAMC253907 | GWHAPHX01000000 |
| East China Sea Library 19 | 19_TRINITY_DN1338_c0_g1_i6 | 3833 | Picornavirales | Unclassified | PRJCA003705 | SAMC253908 | GWHAPHY01000000 |
| East China Sea Library 8-1 | 8-1_TRINITY_DN4556_c0_g1_i1 | 1685 | Durnavirales | Picobirnaviridae | PRJCA003705 | SAMC253909 | GWHAPHZ01000000 |
| South China Sea Library ZH14 | ZH14_TRINITY_DN22339_c0_g1_i | 1129 | Picornavirales | Dicistroviridae | PRJCA003705 | SAMC253953 | GWHAPJR01000000 |

| | | | | | | | |
|---|---|---|---|---|---|---|---|
| East China Sea Library 8-1 | 8-1_TRINITY_DN51474_c0_g1_i3 | 2084 | Picornavirales | Unclassified | PRJCA003705 | SAMC253911 | GWHAPIB01000000 |
| South China Sea Library G12 | G12_TRINITY_DN134571_c0_g2_i | 1034 | Picornavirales | Unclassified | PRJCA003705 | SAMC253758 | GWHAPCE01000000 |
| East China Sea Library 8-1 | 8-1_TRINITY_DN66185_c0_g3_i1 | 2610 | Picornavirales | Marnaviridae | PRJCA003705 | SAMC253913 | GWHAPID01000000 |
| East China Sea Library 8-1 | 8-1_TRINITY_DN7678_c0_g2_i1 | 1028 | Picornavirales | Marnaviridae | PRJCA003705 | SAMC253914 | GWHAPIE01000000 |
| East China Sea Library 8-1 | 8-1_TRINITY_DN86975_c0_g1_i2 | 1048 | Unclassified | Unclassified | PRJCA003705 | SAMC253915 | GWHAPIF01000000 |
| East China Sea Library 8-1 | 8-1_TRINITY_DN96678_c0_g1_i1 | 1040 | Durnavirales | Picobirnaviridae | PRJCA003705 | SAMC253916 | GWHAPIG01000000 |
| East China Sea Library 8-2 | 8-2_TRINITY_DN122090_c0_g2_i5 | 2954 | Picornavirales | Marnaviridae | PRJCA003705 | SAMC253917 | GWHAPIH01000000 |
| East China Sea Library 8-2 | 8-2_TRINITY_DN128170_c0_g1_i2 | 1336 | Hepelivirales | Alphatetraviridae | PRJCA003705 | SAMC253918 | GWHAPII01000000 |
| East China Sea Library 8-2 | 8-2_TRINITY_DN320196_c0_g1_i1 | 1306 | Picornavirales | Unclassified | PRJCA003705 | SAMC253919 | GWHAPIJ01000000 |
| East China Sea Library 8-2 | 8-2_TRINITY_DN358270_c0_g1_i1 | 853 | Picornavirales | Marnaviridae | PRJCA003705 | SAMC253920 | GWHAPIK01000000 |
| East China Sea Library 8-2 | 8-2_TRINITY_DN369983_c0_g1_i1 | 806 | Picornavirales | Marnaviridae | PRJCA003705 | SAMC253921 | GWHAPIL01000000 |
| East China Sea Library 19 | 19_TRINITY_DN67445_c0_g1_i2 | 1026 | Picornavirales | Marnaviridae | PRJCA003705 | SAMC253671 | GWHAOYV01000000 |
| East China Sea Library 8-2 | 8-2_TRINITY_DN44242_c0_g3_i2 | 1115 | Picornavirales | Marnaviridae | PRJCA003705 | SAMC253923 | GWHAPIN01000000 |
| East China Sea Library 8-2 | 8-2_TRINITY_DN50636_c0_g1_i2 | 856 | Picornavirales | Marnaviridae | PRJCA003705 | SAMC253924 | GWHAPIO01000000 |
| East China Sea Library 8-2 | 8-2_TRINITY_DN60228_c0_g1_i1 | 959 | Picornavirales | Unclassified | PRJCA003705 | SAMC253925 | GWHAPIP01000000 |
| East China Sea Library 8-2 | 8-2_TRINITY_DN88231_c0_g2_i1 | 1465 | Picornavirales | Marnaviridae | PRJCA003705 | SAMC253926 | GWHAPIQ01000000 |
| East China Sea Library 8-2 | BH05_TRINITY_DN217842_c0_g1_ | 897 | Picornavirales | Marnaviridae | PRJCA003705 | SAMC253927 | GWHAPIR01000000 |
| South China Sea Library BH05 | BH05_TRINITY_DN238840_c0_g1_ | 1058 | Durnavirales | Picobirnaviridae | PRJCA003705 | SAMC253928 | GWHAPIS01000000 |
| South China Sea Library BH05 | BH05_TRINITY_DN3549_c0_g1_i8 | 2993 | Picornavirales | Dicistroviridae | PRJCA003705 | SAMC253929 | GWHAPIT01000000 |
| South China Sea Library G07 | G07_TRINITY_DN4331_c0_g1_i3 | 954 | Picornavirales | Unclassified | PRJCA003705 | SAMC253930 | GWHAPIU01000000 |
| South China Sea Library G12 | G12_TRINITY_DN45079_c0_g1_i3 | 1772 | Picornavirales | Marnaviridae | PRJCA003705 | SAMC253931 | GWHAPIV01000000 |
| South China Sea Library G14 | G14_TRINITY_DN143872_c0_g3_i | 1096 | Picornavirales | Marnaviridae | PRJCA003705 | SAMC253932 | GWHAPIW01000000 |
| East China Sea Library 19 | 19_TRINITY_DN67864_c0_g1_i1 | 975 | Picornavirales | Marnaviridae | PRJCA003705 | SAMC253672 | GWHAOYW01000000 |
| Yellow Sea Library H02 | H02_TRINITY_DN43443_c0_g1_i1 | 1284 | Picornavirales | Unclassified | PRJCA003705 | SAMC253934 | GWHAPIY01000000 |
| Yellow Sea Library H02 | H02_TRINITY_DN5051_c0_g1_i3 | 1316 | Picornavirales | Marnaviridae | PRJCA003705 | SAMC253935 | GWHAPIZ01000000 |
| Yellow Sea Library H02 | H02_TRINITY_DN5152_c0_g1_i1 | 870 | Picornavirales | Unclassified | PRJCA003705 | SAMC253936 | GWHAPJA01000000 |
| Yellow Sea Library H02 | H02_TRINITY_DN60471_c0_g1_i1 | 924 | Picornavirales | Marnaviridae | PRJCA003705 | SAMC253937 | GWHAPJB01000000 |

| | | | | | | | |
|---|---|---|---|---|---|---|---|
| Yellow Sea Library H04 | H04_TRINITY_DN134457_c0_g1_i | 938 | Unclassified | Unclassified | PRJCA003705 | SAMC253938 | GWHAPJC01000000 |
| Yellow Sea Library H07 | H07_TRINITY_DN208108_c0_g2_i | 1281 | Picornavirales | Marnaviridae | PRJCA003705 | SAMC253939 | GWHAPJD01000000 |
| Yellow Sea Library H07 | H07_TRINITY_DN352121_c0_g1_i | 1126 | Picornavirales | Unclassified | PRJCA003705 | SAMC253940 | GWHAPJE01000000 |
| Yellow Sea Library H07 | H07_TRINITY_DN38393_c0_g1_i1 | 1172 | Picornavirales | Unclassified | PRJCA003705 | SAMC253941 | GWHAPJF01000000 |
| Yellow Sea Library H07 | H07_TRINITY_DN38542_c0_g1_i3 | 3358 | Sobelivirales | Solemoviridae | PRJCA003705 | SAMC253942 | GWHAPJG01000000 |
| Yellow Sea Library H07 | H07_TRINITY_DN5264_c0_g1_i7 | 1320 | Picornavirales | Unclassified | PRJCA003705 | SAMC253943 | GWHAPJH01000000 |
| South China Sea Library ZH04 | ZH04_TRINITY_DN45473_c0_g1_i | 959 | Picornavirales | Unclassified | PRJCA003705 | SAMC253789 | GWHAPDJ01000000 |
| East China Sea Library 20 | 20_TRINITY_DN147687_c0_g1_i3 | 942 | Picornavirales | Iflaviridae | PRJCA003705 | SAMC253682 | GWHAOZG01000000 |
| Yellow Sea Library H07 | H07_TRINITY_DN8311_c0_g1_i5 | 990 | Picornavirales | Unclassified | PRJCA003705 | SAMC253946 | GWHAPJK01000000 |
| Yellow Sea Library H09 | H09_TRINITY_DN214833_c0_g1_i | 1652 | Durnavirales | Picobirnaviridae | PRJCA003705 | SAMC253947 | GWHAPJL01000000 |
| South China Sea Library ZH07 | ZH07_TRINITY_DN8886_c0_g1_i5 | 8265 | Amarillovirales | Flaviviridae | PRJCA003705 | SAMC253948 | GWHAPJM01000000 |
| South China Sea Library ZH08 | ZH08_TRINITY_DN168754_c0_g1_ | 1196 | Picornavirales | Marnaviridae | PRJCA003705 | SAMC253949 | GWHAPJN01000000 |
| South China Sea Library ZH08 | ZH08_TRINITY_DN176904_c0_g1_ | 886 | Picornavirales | Marnaviridae | PRJCA003705 | SAMC253950 | GWHAPJO01000000 |
| South China Sea Library ZH08 | ZH08_TRINITY_DN20506_c0_g1_i | 2714 | Picornavirales | Marnaviridae | PRJCA003705 | SAMC253951 | GWHAPJP01000000 |
| South China Sea Library ZH08 | ZH08_TRINITY_DN61069_c0_g1_i | 1294 | Picornavirales | Marnaviridae | PRJCA003705 | SAMC253952 | GWHAPJQ01000000 |
| East China Sea Library 19 | 19_TRINITY_DN197365_c0_g1_i1 | 913 | Picornavirales | Unclassified | PRJCA003705 | SAMC253662 | GWHAOYM01000000 |
| East China Sea Library 1 | 1_TRINITY_DN13810_c0_g1_i1 | 2625 | Hepelivirales | Alphatetraviridae | PRJCA003705 | SAMC253954 | GWHAPJS01000000 |
| East China Sea Library 11 | 11_TRINITY_DN2073_c0_g1_i2 | 2327 | Picornavirales | Marnaviridae | PRJCA003705 | SAMC253955 | GWHAPJT01000000 |
| East China Sea Library 17 | 17_TRINITY_DN106166_c0_g1_i2 | 811 | Unclassified | Unclassified | PRJCA003705 | SAMC253956 | GWHAPJU01000000 |
| East China Sea Library 17 | 17_TRINITY_DN1818_c0_g1_i6 | 2378 | Picornavirales | Dicistroviridae | PRJCA003705 | SAMC253957 | GWHAPJV01000000 |
| East China Sea Library 17 | 17_TRINITY_DN24209_c0_g1_i2 | 3497 | Sobelivirales | Solemoviridae | PRJCA003705 | SAMC253958 | GWHAPJW01000000 |
| East China Sea Library 17 | 17_TRINITY_DN24637_c0_g1_i1 | 1283 | Picornavirales | Dicistroviridae | PRJCA003705 | SAMC253959 | GWHAPJX01000000 |
| East China Sea Library 17 | 17_TRINITY_DN6981_c0_g1_i2 | 1117 | Picornavirales | Marnaviridae | PRJCA003705 | SAMC253960 | GWHAPJY01000000 |
| East China Sea Library 17 | 17_TRINITY_DN87839_c0_g1_i2 | 1740 | Unclassified | Unclassified | PRJCA003705 | SAMC253961 | GWHAPJZ01000000 |
| East China Sea Library 18 | 18_TRINITY_DN46869_c0_g1_i1 | 986 | Picornavirales | Marnaviridae | PRJCA003705 | SAMC253962 | GWHAPKA01000000 |
| East China Sea Library 18 | 18_TRINITY_DN95612_c0_g1_i4 | 1436 | Picornavirales | Unclassified | PRJCA003705 | SAMC253963 | GWHAPKB01000000 |
| East China Sea Library 6 | 6_TRINITY_DN2661_c0_g3_i1 | 1069 | Picornavirales | Dicistroviridae | PRJCA003705 | SAMC253964 | GWHAPKC01000000 |

| | | | | | | | |
|---|---|---|---|---|---|---|---|
| East China Sea Library 6 | 6_TRINITY_DN51494_c0_g1_i1 | 1190 | Picornavirales | Dicistroviridae | PRJCA003705 | SAMC253965 | GWHAPKD01000000 |
| East China Sea Library 6 | 6_TRINITY_DN80338_c0_g3_i1 | 1640 | Picornavirales | Dicistroviridae | PRJCA003705 | SAMC253966 | GWHAPKE01000000 |
| East China Sea Library 7 | 7_TRINITY_DN105202_c0_g1_i1 | 846 | Picornavirales | Marnaviridae | PRJCA003705 | SAMC253967 | GWHAPKF01000000 |
| East China Sea Library 7 | 7_TRINITY_DN111908_c0_g2_i1 | 878 | Picornavirales | Marnaviridae | PRJCA003705 | SAMC253968 | GWHAPKG01000000 |
| East China Sea Library 7 | 7_TRINITY_DN217501_c0_g1_i1 | 1055 | Picornavirales | Marnaviridae | PRJCA003705 | SAMC253969 | GWHAPKH01000000 |
| East China Sea Library 7 | 7_TRINITY_DN22197_c0_g1_i1 | 1458 | Durnavirales | Picobirnaviridae | PRJCA003705 | SAMC253970 | GWHAPKI01000000 |
| East China Sea Library 7 | 7_TRINITY_DN65869_c0_g1_i2 | 1521 | Durnavirales | Picobirnaviridae | PRJCA003705 | SAMC253971 | GWHAPKJ01000000 |
| East China Sea Library 7 | 7_TRINITY_DN91604_c0_g1_i1 | 4551 | Picornavirales | Marnaviridae | PRJCA003705 | SAMC253972 | GWHAPKK01000000 |
| East China Sea Library 8-1 | 8-1_TRINITY_DN118249_c0_g4_i1 | 967 | Picornavirales | Marnaviridae | PRJCA003705 | SAMC253973 | GWHAPKL01000000 |
| East China Sea Library 8-1 | 8-1_TRINITY_DN124617_c0_g2_i1 | 897 | Unclassified | Unclassified | PRJCA003705 | SAMC253974 | GWHAPKM01000000 |
| East China Sea Library 8-1 | 8-1_TRINITY_DN15500_c0_g3_i1 | 1158 | Picornavirales | Marnaviridae | PRJCA003705 | SAMC253975 | GWHAPKN01000000 |
| East China Sea Library 8-1 | 8-1_TRINITY_DN160154_c0_g2_i1 | 921 | Picornavirales | Marnaviridae | PRJCA003705 | SAMC253976 | GWHAPKO01000000 |
| East China Sea Library 8-1 | 8-1_TRINITY_DN163267_c0_g1_i1 | 826 | Picornavirales | Marnaviridae | PRJCA003705 | SAMC253977 | GWHAPKP01000000 |
| East China Sea Library 8-1 | 8-1_TRINITY_DN166016_c0_g1_i1 | 820 | Picornavirales | Marnaviridae | PRJCA003705 | SAMC253978 | GWHAPKQ01000000 |
| East China Sea Library 8-1 | 8-1_TRINITY_DN182373_c0_g1_i1 | 954 | Picornavirales | Marnaviridae | PRJCA003705 | SAMC253979 | GWHAPKR01000000 |
| East China Sea Library 8-1 | 8-1_TRINITY_DN18524_c0_g1_i1 | 1556 | Tolivirales | Tombusviridae | PRJCA003705 | SAMC253980 | GWHAPKS01000000 |
| East China Sea Library 8-1 | 8-1_TRINITY_DN190690_c0_g1_i1 | 1187 | Picornavirales | Marnaviridae | PRJCA003705 | SAMC253981 | GWHAPKT01000000 |
| East China Sea Library 16 | 16_TRINITY_DN68194_c0_g1_i1 | 863 | Picornavirales | Marnaviridae | PRJCA003705 | SAMC253658 | GWHAOYI01000000 |
| East China Sea Library 8-1 | 8-1_TRINITY_DN24911_c0_g2_i1 | 1579 | Picornavirales | Marnaviridae | PRJCA003705 | SAMC253983 | GWHAPKV01000000 |
| South China Sea Library G08 | G08_TRINITY_DN464_c0_g1_i1 | 8081 | Ghabrivirales | Totiviridae | PRJCA003705 | SAMC253984 | GWHAPKW01000000 |
| South China Sea Library G10 | G10_TRINITY_DN11011_c0_g1_i1 | 1230 | Picornavirales | Dicistroviridae | PRJCA003705 | SAMC253985 | GWHAPKX01000000 |
| South China Sea Library G10 | G10_TRINITY_DN2373_c0_g1_i8 | 5141 | Tolivirales | Tombusviridae | PRJCA003705 | SAMC253986 | GWHAPKY01000000 |
| South China Sea Library G12 | G12_TRINITY_DN73867_c0_g1_i1 | 801 | Picornavirales | Marnaviridae | PRJCA003705 | SAMC253987 | GWHAPKZ01000000 |
| South China Sea Library G12 | G12_TRINITY_DN810_c0_g1_i5 | 2275 | Picornavirales | Unclassified | PRJCA003705 | SAMC253988 | GWHAPLA01000000 |
| South China Sea Library G13 | G13_TRINITY_DN20817_c0_g1_i1 | 1650 | Picornavirales | Unclassified | PRJCA003705 | SAMC253989 | GWHAPLB01000000 |
| South China Sea Library G13 | G13_TRINITY_DN7700_c0_g1_i6 | 1216 | Wolframvirales | Narnaviridae | PRJCA003705 | SAMC253990 | GWHAPLC01000000 |
| South China Sea Library G13 | G13_TRINITY_DN81622_c0_g1_i1 | 852 | Durnavirales | Picobirnaviridae | PRJCA003705 | SAMC253991 | GWHAPLD01000000 |

| | | | | | | | |
|---|---|---|---|---|---|---|---|
| South China Sea Library G14 | G14_TRINITY_DN11170_c0_g1_i1 | 1618 | Durnavirales | Picobirnaviridae | PRJCA003705 | SAMC253992 | GWHAPLE01000000 |
| South China Sea Library G14 | G14_TRINITY_DN125042_c0_g1_i | 1223 | Picornavirales | Unclassified | PRJCA003705 | SAMC253993 | GWHAPLF01000000 |
| South China Sea Library G14 | G14_TRINITY_DN134_c0_g1_i12 | 10002 | Picornavirales | Unclassified | PRJCA003705 | SAMC253994 | GWHAPLG01000000 |
| South China Sea Library G14 | G14_TRINITY_DN134_c1_g1_i4 | 1828 | Picornavirales | Unclassified | PRJCA003705 | SAMC253995 | GWHAPLH01000000 |
| South China Sea Library G14 | G14_TRINITY_DN135456_c0_g1_i | 875 | Picornavirales | Picornaviridae | PRJCA003705 | SAMC253996 | GWHAPLI01000000 |
| Yellow Sea Library H09 | H09_TRINITY_DN38653_c0_g1_i1 | 1317 | Picornavirales | Marnaviridae | PRJCA003705 | SAMC253997 | GWHAPLJ01000000 |
| South China Sea Library ZH07 | ZH07_TRINITY_DN19819_c0_g1_i | 1117 | Bunyavirales | Hantaviridae | PRJCA003705 | SAMC253998 | GWHAPLK01000000 |
| South China Sea Library ZH07 | ZH07_TRINITY_DN22405_c0_g1_i | 2473 | Picornavirales | Unclassified | PRJCA003705 | SAMC253999 | GWHAPLL01000000 |
| South China Sea Library ZH07 | ZH07_TRINITY_DN40582_c0_g1_i | 1315 | Picornavirales | Unclassified | PRJCA003705 | SAMC254000 | GWHAPLM01000000 |
| South China Sea Library ZH14 | ZH14_TRINITY_DN4424_c0_g1_i1 | 815 | Picornavirales | Unclassified | PRJCA003705 | SAMC254001 | GWHAPLN01000000 |
| South China Sea Library ZH14 | ZH14_TRINITY_DN520_c0_g1_i6 | 8721 | Picornavirales | Unclassified | PRJCA003705 | SAMC254002 | GWHAPLO01000000 |
| South China Sea Library ZH14 | ZH14_TRINITY_DN60154_c0_g2_i | 2001 | Picornavirales | Marnaviridae | PRJCA003705 | SAMC254003 | GWHAPLP01000000 |
| Yellow Sea Library H02 | H02_TRINITY_DN34742_c0_g1_i2 | 829 | Picornavirales | Unclassified | PRJCA003705 | SAMC253768 | GWHAPCO01000000 |
| South China Sea Library ZH14 | ZH14_TRINITY_DN7640_c0_g1_i2 | 2133 | Unclassified | Unclassified | PRJCA003705 | SAMC254005 | GWHAPLR01000000 |
| South China Sea Library ZH14 | ZH14_TRINITY_DN7791_c0_g1_i7 | 3856 | Picornavirales | Unclassified | PRJCA003705 | SAMC254006 | GWHAPLS01000000 |
| East China Sea Library 12 | 12_TRINITY_DN962_c0_g1_i2 | 1725 | Durnavirales | Picobirnaviridae | PRJCA003705 | SAMC253644 | GWHAPLT01000000 |

| Virus Name | Whether in tree | Region | Whether novel | Blastx hits on known viruses (Blast amino acid identity) | PCR |
|---|---|---|---|---|---|
| Bivalvia Picorna-like virus D13 | | Capsid | Yes | Wenzhou picorna-like virus 53 (28%) | Yes |
| Bivalvia Picorna-like virus D7 | | RNA-helicase | Yes | Beihai picorna-like virus 36 (46%) | Yes |
| Bivalvia Picorna-like virus D8 | | | Yes | Marine RNA virus BC-1 (50%) | Yes |
| Bivalvia Picorna-like virus D9 | | RNA-helicase | Yes | Wenzhou picorna-like virus 29 (31%) | Yes |
| Bivalvia Unclassified virus D1 | | RNA-helicase, Tryp SPc | Yes | Halhan virus 1 (72%) | Yes |
| Bivalvia Picorna-like virus D10 | | | | Wenzhou picorna-like virus 29 (100%) | Yes |
| Bivalvia Picorna-like virus D11 | | RNA-helicase | | Wenzhou picorna-like virus 29 (100%) | Yes |
| Bivalvia Picorna-like virus D6 | | | Yes | Beihai narna-like virus 26 (40%) | Yes |
| Bivalvia Picorna-like virus D12 | | Capsid | Yes | Wenzhou picorna-like virus 29 (49%) | Yes |
| Bivalvia Durna-like virus D3 | Yes | RdRp | Yes | Beihai picobirna-like virus 13 (50%) | Yes |
| Bivalvia Picorna-like virus D14 | | | Yes | Tioga picorna-like virus 1 (38%) | Yes |
| Bivalvia Durna-like virus D2 | Yes | RdRp | Yes | Beihai picobirna-like virus 10 (73%) | Yes |
| Bivalvia Picorna-like virus D15 | | RNA-helicase | Yes | Wenzhou picorna-like virus 2 (66%) | Yes |
| Bivalvia Picorna-like virus H14 | Yes | RdRp, Capsid, RNA-helicase | Yes | Beihai octopus virus 2 (26%) | Yes |
| Gastropoda Picorna-like virus D4 | | RdRp | Yes | Wenzhou picorna-like virus 48 (43%) | Yes |
| Crustacea Picorna-like virus N6 | Yes | RdRp, RNA-helicase, PHA03247, G-patch | | Wenzhou shrimp virus 8 (92%) | Yes |
| Bivalvia Durna-like virus D5 | Yes | RdRp | Yes | Cryptosporidium parvum virus 1 (29%) | Yes |
| Bivalvia Picorna-like virus D72 | Yes | RdRp, RNA-helicase, Capsid | Yes | Beihai picorna-like virus 107 (70%) | Yes |
| Crustacea Picorna-like virus N32 | Yes | RdRp, RNA-helicase, PHA03247 | Yes | Wenzhou shrimp virus 8 (74%) | Yes |
| Crustacea Picorna-like virus D8 | Yes | RdRp, RNA-helicase, AAA 19 | Yes | Beihai blue swimmer crab virus 1 (26%) | Yes |
| Bivalvia Picorna-like virus D37 | | | Yes | Marine RNA virus PAL438 (35%) | Yes |
| Bivalvia Picorna-like virus D30 | | | Yes | Beihai picorna-like virus 29 (37%) | Yes |
| Bivalvia Durna-like virus D4 | Yes | RdRp | Yes | Beihai picobirna-like virus 12 (47%) | Yes |

| | | | | | |
|---|---|---|---|---|---|
| Bivalvia Picorna-like virus D32 | | | Yes | Beihai picorna-like virus 41 (31%) | Yes |
| Bivalvia Picorna-like virus D29 | | RNA-helicase | Yes | Beihai picorna-like virus 122 (31%) | Yes |
| Bivalvia Picorna-like virus D74 | Yes | RdRp, RNA-helicase, Capsid | Yes | Cragig virus 7 (60%) | Yes |
| Gastropoda Picorna-like virus D6 | Yes | Capsid, RNA-helicase, RdRp | Yes | Hubei picorna-like virus 43 (32%) | Yes |
| Bivalvia Picorna-like virus D75 | Yes | RdRp, Capsid, RNA-helicase | Yes | Picornavirales sp. (24%) | Yes |
| Bivalvia Picorna-like virus D34 | | | | Beihai picorna-like virus 70 (93%) | Yes |
| Bivalvia Picorna-like virus D40 | | RNA-helicase | Yes | Zehneria japonica marnavirus (30%) | Yes |
| Bivalvia Picorna-like virus D38 | | Capsid | Yes | Picornavirales sp. (36%) | Yes |
| Bivalvia Picorna-like virus D20 | Yes | RdRp, RNA-helicase | Yes | Beihai picorna-like virus 111 (23%) | Yes |
| Bivalvia Unclassified virus D2 | | RNA-helicase | Yes | Cylindrotheca closterium RNA virus 02 (47%) | Yes |
| Bivalvia Picorna-like virus D56 | Yes | RdRp, RNA-helicase, P-loop_NTPase, Capsid | Yes | Cragig virus 7 (69%) | Yes |
| Bivalvia Picorna-like virus D31 | | RNA-helicase | Yes | Beihai picorna-like virus 36 (53%) | Yes |
| Bivalvia Toli-like virus D1 | Yes | RdRp | Yes | Sanxia tombus-like virus 5 (30%) | Yes |
| Bivalvia Durna-like virus D9 | | | Yes | Beihai picobirna-like virus 13 (44%) | Yes |
| Bivalvia Picorna-like virus D22 | Yes | RdRp, Capsid | Yes | Beihai picorna-like virus 57 (42%) | Yes |
| Bivalvia Picorna-like virus D55 | | | Yes | Wenzhou picorna-like virus 4 (60%) | Yes |
| Bivalvia Picorna-like virus D42 | | Capsid, RNA-helicase | Yes | Beihai octopus virus 2 (24%) | Yes |
| Bivalvia Picorna-like virus D46 | | | Yes | Beihai picorna-like virus 14 (72%) | Yes |
| Bivalvia Picorna-like virus D41 | | | Yes | Beihai narna-like virus 10 (42%) | Yes |
| Bivalvia Picorna-like virus D43 | | RdRp | Yes | Beihai octopus virus 2 (35%) | Yes |
| Bivalvia Durna-like virus D10 | | | Yes | Lysoka partiti-like virus (22%) | Yes |
| Bivalvia Picorna-like virus D44 | | RdRp | Yes | Beihai octopus virus 2 (31%) | Yes |
| Bivalvia Picorna-like virus D50 | | RNA-helicase | Yes | Beihai sesarmid crab virus 1 (40%) | Yes |
| Bivalvia Picorna-like virus D53 | | Capsid | Yes | Picornavirales sp. (39%) | Yes |
| Bivalvia Durna-like virus D7 | Yes | RdRp | Yes | Beihai picobirna-like virus 10 (73%) | Yes |
| Bivalvia Picorna-like virus D52 | | RNA-helicase | Yes | Marine RNA virus BC-4 (56%) | Yes |
| Bivalvia Picorna-like virus D54 | | | | Wenzhou gastropodes virus 2 (97%) | |

| | | | | | |
|---|---|---|---|---|---|
| Bivalvia Picorna-like virus D49 | | | Yes | Beihai picorna-like virus 4 (51%) | Yes |
| Bivalvia Picorna-like virus D48 | | Capsid | Yes | Beihai picorna-like virus 23 (54%) | Yes |
| Bivalvia Picorna-like virus D51 | | | Yes | Marine RNA virus BC-1 (25%) | Yes |
| Bivalvia Durna-like virus D8 | Yes | RdRp | Yes | Beihai picobirna-like virus 7 (35%) | Yes |
| Bivalvia Picorna-like virus D47 | | | Yes | Beihai picorna-like virus 17 (51%) | Yes |
| Bivalvia Hepeli-like virus D1 | Yes | RdRp, RNA-helicase | Yes | Beihai hepe-like virus 8 (49%) | Yes |
| Bivalvia Picorna-like virus D61 | | RNA-helicase | Yes | Picornavirales N OV 003 (55%) | Yes |
| Bivalvia Durna-like virus D11 | Yes | RdRp | Yes | Picobirnavirus sp. (26%) | Yes |
| Bivalvia Picorna-like virus N13 | Yes | RdRp, Capsid | Yes | Beihai picorna-like virus 121 (26%) | Yes |
| Bivalvia Picorna-like virus D58 | | | Yes | Beihai picorna-like virus 14 (73%) | Yes |
| Bivalvia Picorna-like virus D60 | | Capsid | Yes | Perth bee virus 8 (38%) | Yes |
| Bivalvia Picorna-like virus D59 | | Capsid | Yes | Beihai picorna-like virus 69 (30%) | Yes |
| Bivalvia Picorna-like virus D28 | Yes | RdRp, RNA-helicase | Yes | Picornavirales sp. (25%) | Yes |
| Bivalvia Picorna-like virus D64 | | RNA-helicase | Yes | Beihai picorna-like virus 32 (30%) | Yes |
| Bivalvia Picorna-like virus D65 | | | Yes | Wenzhou picorna-like virus 9 (35%) | Yes |
| Crustacea Picorna-like virus N14 | Yes | RNA-helicase, RdRp, Tryp SPc | Yes | Beihai picorna-like virus 90 (80%) | Yes |
| Bivalvia Durna-like virus D13 | | | Yes | Shahe picobirna-like virus 2 (40%) | Yes |
| Bivalvia Hepeli-like virus D2 | Yes | RdRp | | Barns Ness breadcrumb sponge hepe-like virus 2 (100%) | Yes |
| Bivalvia Durna-like virus D12 | | | Yes | Shahe picobirna-like virus 1 (53%) | Yes |
| Bivalvia Picorna-like virus H5 | Yes | RNA-helicase, RdRp | Yes | Wenzhou picorna-like virus 45 (30%) | Yes |
| Gastropoda Sobeli-like virus D2 | Yes | RdRp | Yes | Beihai sobemo-like virus 20 (45%) | Yes |
| Gastropoda Picorna-like virus D11 | | RNA-helicase | Yes | Wenzhou picorna-like virus 6 (38%) | Yes |
| Gastropoda Picorna-like virus D9 | | Peptidase C3 | Yes | Shahe picorna-like virus 2 (27%) | Yes |
| Gastropoda Picorna-like virus D8 | | RNA-helicase | Yes | Picornavirales sp. (31%) | Yes |
| Gastropoda Picorna-like virus D5 | | | Yes | Beihai paphia shell virus 2 (40%) | Yes |
| Crustacea Picorna-like virus N20 | Yes | RNA-helicase, RdRp | Yes | Beihai shrimp virus 2 (63%) | Yes |
| Gastropoda Picorna-like virus D10 | | | Yes | Wenzhou picorna-like virus 4 (66%) | Yes |

| | | | | | |
|---|---|---|---|---|---|
| Gastropoda Picorna-like virus D7 | | RNA-helicase | Yes | Beihai picorna-like virus 77 (31%) | Yes |
| Bivalvia Picorna-like virus H13 | Yes | RNA-helicase, RdRp | Yes | Beihai octopus virus 2 (28%) | Yes |
| Gastropoda Unclassified virus D3 | | RNA-helicase | Yes | Basavirus sp. (32%) | Yes |
| Gastropoda Picorna-like virus D14 | | RNA-helicase | Yes | Picornavirales N OV 003 (46%) | Yes |
| Gastropoda Picorna-like virus D16 | | Capsid | Yes | Wenling picorna-like virus 5 (31%) | Yes |
| Gastropoda Picorna-like virus D17 | | RNA-helicase | Yes | Wenzhou picorna-like virus 28 (26%) | Yes |
| Gastropoda Picorna-like virus D12 | | RNA-helicase | Yes | Barns Ness breadcrumb sponge aquatic picorna-like virus 1 (26%) | Yes |
| Gastropoda Picorna-like virus D18 | | | Yes | Wenzhou picorna-like virus 7 (57%) | Yes |
| Gastropoda Unclassified virus D4 | | RdRp, Capsid | Yes | Fisavirus 1 (29%) | Yes |
| Gastropoda Picorna-like virus D15 | | RNA-helicase, Peptidase C3 | Yes | Picornavirales sp. (21%) | Yes |
| Gastropoda Picorna-like virus D23 | | RNA-helicase | Yes | Bivalve RNA virus G5 (43%) | Yes |
| Gastropoda Picorna-like virus D25 | | RdRp | Yes | Wenzhou picorna-like virus 48 (39%) | Yes |
| Gastropoda Picorna-like virus D22 | | Capsid | Yes | Beihai picorna-like virus 72 (52%) | Yes |
| Gastropoda Picorna-like virus D20 | | | | Beihai picorna-like virus 70 (92%) | Yes |
| Gastropoda Picorna-like virus D24 | | Capsid | Yes | Wenzhou picorna-like virus 26 (86%) | Yes |
| Gastropoda Picorna-like virus D21 | | RNA-helicase | Yes | Beihai picorna-like virus 70 (82%) | Yes |
| Gastropoda Picorna-like virus D19 | | Peptidase C3 | Yes | Atrpec virus 1 (79%) | Yes |
| Gastropoda Unclassified virus D5 | | helicase | Yes | Daeseongdong virus 1 (45%) | Yes |
| Gastropoda Unclassified virus D6 | | | Yes | Loreto virus (38%) | Yes |
| Bivalvia Picorna-like virus D62 | Yes | RNA-helicase, RdRp | Yes | Beihai octopus virus 2 (27%) | Yes |
| Crustacea Picorna-like virus D5 | | RNA-helicase | | Wenzhou shrimp virus 8 (96%) | Yes |
| Crustacea Unclassified virus D1 | | RNA-helicase | Yes | Rainier virus (26%) | Yes |
| Crustacea Picorna-like virus D3 | | Capsid | Yes | Hubei picorna-like virus 23 (23%) | Yes |
| Crustacea Picorna-like virus D6 | | | | Wenzhou shrimp virus 8 (99%) | Yes |
| Crustacea Picorna-like virus D2 | | RNA-helicase | Yes | Beihai picorna-like virus 107 (62%) | Yes |
| Crustacea Picorna-like virus D4 | | | Yes | Marine RNA virus BC-1 (36%) | Yes |
| Bivalvia Picorna-like virus D23 | Yes | RNA-helicase, RdRp, Peptidase C3 | | Beihai picorna-like virus 70 (97%) | Yes |

| | | | | | |
|---|---|---|---|---|---|
| Bivalvia Picorna-like virus D57 | Yes | RNA-helicase, RdRp | Yes | Hubei odonate virus 1 (26%) | Yes |
| Bivalvia Durna-like virus D14 | Yes | RdRp | Yes | Beihai picobirna-like virus 10 (72%) | Yes |
| Hexanauplia Picorna-like virus D1 | | Capsid, VP4 | Yes | Beihai picorna-like virus 84 (37%) | Yes |
| Hexanauplia Bunya-like virus D1 | | | Yes | Shahe bunya-like virus 4 (24%) | Yes |
| Crustacea Picorna-like virus N12 | | RNA-helicase | Yes | Wenzhou shrimp virus 4 (37%) | Yes |
| Gastropoda Picorna-like virus N12 | | RNA-helicase, DSRM SF, 235kDa-fam, Peptidase C3 | Yes | Wenling crustacean virus 2 (36%) | Yes |
| Gastropoda Picorna-like virus N14 | | RNA-helicase | | Wenzhou picorna-like virus 28 (99%) | Yes |
| Gastropoda Picorna-like virus N15 | | Capsid | | Wenzhou picorna-like virus 28 (100%) | Yes |
| Gastropoda Picorna-like virus N13 | | Peptidase C3 | Yes | Wenling crustacean virus 2 (33%) | Yes |
| Bivalvia Picorna-like virus N56 | Yes | RdRp | Yes | Beihai octopus virus 2 (35%) | Yes |
| Gastropoda Picorna-like virus D13 | Yes | Capsid, RdRp | Yes | Shahe picorna-like virus 10 (25%) | Yes |
| Bivalvia Picorna-like virus N29 | | RNA-helicase | | Wenzhou picorna-like virus 38 (95%) | Yes |
| Bivalvia Picorna-like virus N27 | | | Yes | Wenzhou gastropodes virus 2 (84%) | Yes |
| Bivalvia Picorna-like virus N28 | | | | Wenzhou gastropodes virus 2 (97%) | Yes |
| Bivalvia Picorna-like virus N19 | | PRK01122 | Yes | Beihai picorna-like virus 17 (33%) | Yes |
| Gastropoda Picorna-like virus H1 | | | Yes | Beihai paphia shell virus 3 (33%) | Yes |
| Gastropoda Picorna-like virus H2 | | RNA-helicase | Yes | Cragig virus 1 (38%) | Yes |
| Bivalvia Picorna-like virus H17 | Yes | Capsid, RdRp | | Wenzhou gastropodes virus 2 (92%) | Yes |
| Bivalvia Durna-like virus H2 | Yes | RdRp | Yes | Dralkin virus (79%) | Yes |
| Bivalvia Picorna-like virus D25 | Yes | RdRp, Capsid | Yes | Beihai picorna-like virus 82 (27%) | Yes |
| Bivalvia Durna-like virus H1 | Yes | RdRp | Yes | Beihai picobirna-like virus 10 (73%) | Yes |
| Bivalvia Durna-like virus H8 | Yes | RdRp | Yes | Beihai picobirna-like virus 10 (78%) | Yes |
| Bivalvia Sobeli-like virus H2 | Yes | RdRp | Yes | Beihai sobemo-like virus 17 (48%) | Yes |
| Bivalvia Picorna-like virus H20 | | | | Wenzhou picorna-like virus 10 (97%) | Yes |
| Polychaeta Picorna-like virus H6 | | RdRp, Capsid | Yes | Posavirus strain 9043 (31%) | Yes |
| Polychaeta Picorna-like virus H5 | | Capsid | Yes | Posavirus 1 (23%) | Yes |
| Polychaeta Picorna-like virus H2 | | Capsid | Yes | Beihai picorna-like virus 70 (67%) | Yes |

| | | | | | |
|---|---|---|---|---|---|
| Polychaeta Picorna-like virus H4 | | RNA-helicase | Yes | Paroligolophus agrestis posalike virus 1 (27%) | Yes |
| Polychaeta Picorna-like virus H3 | | | Yes | Beihai picorna-like virus 70 (48%) | Yes |
| Polychaeta Picorna-like virus H1 | | | Yes | Beihai picorna-like virus 15 (55%) | Yes |
| Bivalvia Picorna-like virus N58 | Yes | RdRp, DUF4455 | | Wenzhou picorna-like virus 38 (99%) | Yes |
| Crustacea Picorna-like virus N13 | | Capsid | Yes | Beihai picorna-like virus 90 (84%) | Yes |
| Crustacea Picorna-like virus N15 | | RNA-helicase | Yes | Wenzhou shrimp virus 10 (79%) | Yes |
| Crustacea Picorna-like virus N27 | | | Yes | Wenzhou shrimp virus 8 (63%) | Yes |
| Crustacea Picorna-like virus N16 | | | Yes | Beihai picorna-like virus 113 (82%) | Yes |
| Crustacea Picorna-like virus N28 | | | Yes | Wenzhou shrimp virus 8 (66%) | Yes |
| Bivalvia Picorna-like virus N23 | Yes | RdRp | Yes | Wenling crustacean virus 6 (34%) | Yes |
| Crustacea Picorna-like virus N17 | | | Yes | Beihai picorna-like virus 113 (89%) | Yes |
| Crustacea Picorna-like virus N29 | | PRK04195 | Yes | Wenzhou shrimp virus 8 (70%) | Yes |
| Crustacea Picorna-like virus N22 | | RNA-helicase | Yes | Wenzhou shrimp virus 5 (44%) | Yes |
| Bivalvia Picorna-like virus N7 | Yes | RdRp, PrsW-protease | Yes | Beihai octopus virus 2 (42%) | Yes |
| Crustacea Picorna-like virus N21 | | Capsid | Yes | Wenzhou shrimp virus 4 (60%) | Yes |
| Crustacea Picorna-like virus N24 | | | Yes | Wenzhou shrimp virus 6 (52%) | Yes |
| Crustacea Picorna-like virus N18 | | RNA-helicase | Yes | Beihai picorna-like virus 113 (78%) | Yes |
| Crustacea Picorna-like virus N25 | | Peptidase C3 | Yes | Wenzhou shrimp virus 6 (64%) | Yes |
| Crustacea Picorna-like virus N26 | | | Yes | Wenzhou shrimp virus 6 (47%) | Yes |
| Crustacea Picorna-like virus N23 | | | Yes | Wenzhou shrimp virus 5 (56%) | Yes |
| Crustacea Picorna-like virus N30 | | | | Beihai picorna-like virus 112 (97%) | Yes |
| Crustacea Picorna-like virus N31 | | | | Beihai picorna-like virus 112 (99%) | Yes |
| Crustacea Picorna-like virus N33 | | G-patch | Yes | Wenzhou shrimp virus 8 (60%) | |
| Crustacea Unclassified virus N1 | | | Yes | Kuiper virus (56%) | Yes |
| Bivalvia Picorna-like virus N20 | Yes | RdRp, PrsW-protease | Yes | Beihai octopus virus 2 (42%) | Yes |
| Bivalvia Amarillo-like virus N3 | Yes | RdRp, capping 2-OMTase | Yes | Southern pygmy squid flavivirus (26%) | Yes |
| Bivalvia Toli-like virus N1 | | PHA03255 | Yes | Beihai tombus-like virus 10 (61%) | Yes |

| | | | | | |
|---|---|---|---|---|---|
| Bivalvia Picorna-like virus N43 | | RdRp | Yes | Picornaviridae sp. (34%) | Yes |
| Bivalvia Picorna-like virus N46 | | RNA-helicase | Yes | Wenling picorna-like virus 2 (39%) | Yes |
| Bivalvia Durna-like virus N5 | Yes | RdRp | Yes | Beihai picobirna-like virus 10 (53%) | Yes |
| Bivalvia Picorna-like virus N52 | | RNA-helicase, CAF-1 p150, MFS | Yes | Wenzhou gastropodes virus 2 (89%) | Yes |
| Bivalvia Picorna-like virus N54 | | RNA-helicase | | Wenzhou picorna-like virus 38 (94%) | Yes |
| Bivalvia Picorna-like virus H15 | Yes | RdRp, PrsW-protease | Yes | Beihai octopus virus 2 (40%) | Yes |
| Bivalvia Unclassified virus N2 | | | Yes | Rudphi virus 1 (53%) | Yes |
| Bivalvia Picorna-like virus N47 | | RNA-helicase | Yes | Beihai octopus virus 2 (29%) | Yes |
| Bivalvia Picorna-like virus N55 | | RdRp | | Wenzhou picorna-like virus 38 (97%) | Yes |
| Bivalvia Unclassified virus N1 | | | Yes | Mytedu virus 1 (81%) | Yes |
| Bivalvia Picorna-like virus N48 | | Capsid, Capsid | Yes | Beihai octopus virus 2 (33%) | Yes |
| Bivalvia Toli-like virus N2 | | | Yes | Beihai tombus-like virus 4 (71%) | Yes |
| Crustacea Hepeli-like virus D2 | | Macro OAADPr deacetylase, NADAR | Yes | Wenling hepe-like virus 1 (27%) | Yes |
| Crustacea Hepeli-like virus D3 | | | Yes | Wenling hepe-like virus 1 (37%) | Yes |
| Bivalvia Picorna-like virus D2 | | | Yes | Beihai picorna-like virus 29 (46%) | Yes |
| Bivalvia Picorna-like virus D3 | | RNA-helicase | | Beihai mollusks virus 1 (98%) | Yes |
| Bivalvia Picorna-like virus D4 | | | Yes | Wenzhou picorna-like virus 45 (35%) | Yes |
| Bivalvia Picorna-like virus D5 | | Capsid | Yes | Wenzhou picorna-like virus 45 (32%) | Yes |
| Bivalvia Picorna-like virus D19 | | | Yes | Beihai hermit crab virus 1 (38%) | Yes |
| Bivalvia Picorna-like virus D33 | | Capsid | | Beihai picorna-like virus 70 (98%) | Yes |
| Bivalvia Picorna-like virus D39 | | RNA-helicase | Yes | Trichosanthes kirilowii picorna-like virus (21%) | Yes |
| Bivalvia Durna-like virus D6 | Yes | RdRp | Yes | Luncke virus (51%) | Yes |
| Bivalvia Picorna-like virus D79 | | RNA-helicase | Yes | Shahe picorna-like virus 4 (33%) | Yes |
| Bivalvia Picorna-like virus D81 | | | Yes | Beihai picorna-like virus 39 (47%) | Yes |
| Bivalvia Amarillo-like virus N1 | Yes | RdRp, capping 2-OMTase, RNA-helicase | Yes | Southern pygmy squid flavivirus (27%) | Yes |
| Bivalvia Picorna-like virus N2 | | | | Beihai picorna-like virus 56 (97%) | Yes |
| Bivalvia Picorna-like virus N4 | | | | Wenzhou picorna-like virus 50 (97%) | Yes |

| | | | | | |
|---|---|---|---|---|---|
| Bivalvia Picorna-like virus N8 | | | | Wenzhou gastropodes virus 2 (90%) | Yes |
| Bivalvia Picorna-like virus N5 | | Capsid | Yes | Beihai octopus virus 2 (35%) | Yes |
| Bivalvia Picorna-like virus N6 | | RNA-helicase | Yes | Beihai octopus virus 2 (39%) | Yes |
| Crustacea Picorna-like virus D1 | Yes | RdRp, Peptidase C3 | Yes | Wenzhou picorna-like virus 42 (34%) | Yes |
| Bivalvia Sobeli-like virus N1 | Yes | RdRp | Yes | Beihai sobemo-like virus 21 (41%) | Yes |
| Gastropoda Picorna-like virus N1 | | | Yes | Beihai mollusks virus 2 (44%) | Yes |
| Gastropoda Picorna-like virus N3 | | | Yes | Beihai picorna-like virus 29 (31%) | Yes |
| Gastropoda Picorna-like virus N2 | | | Yes | Beihai picorna-like virus 15 (79%) | Yes |
| Gastropoda Toli-like virus N1 | | RdRp | | Wenling tombus-like virus 1 (100%) | |
| Crustacea Jingchu-like virus N1 | Yes | RdRp, mRNAcap | | Beihai hermit crab virus 3 (97%) | Yes |
| Crustacea Durna-like virus N1 | | | Yes | Beihai picobirna-like virus 1 (60%) | Yes |
| Crustacea Durna-like virus N2 | | | | Beihai picobirna-like virus 5 (91%) | Yes |
| Gastropoda Stella-like virus N1 | | | Yes | Astroviridae sp. (28%) | Yes |
| Gastropoda Picorna-like virus N5 | | RNA-helicase | Yes | Changjiang picorna-like virus 13 (31%) | Yes |
| Gastropoda Picorna-like virus N6 | | RNA-helicase | Yes | Marnaviridae sp. (35%) | Yes |
| Gastropoda Picorna-like virus N4 | | | Yes | Beihai picorna-like virus 46 (29%) | Yes |
| Gastropoda Sobeli-like virus N1 | | Tryp SPc | Yes | Beihai sobemo-like virus 13 (26%) | Yes |
| Gastropoda Picorna-like virus N9 | | RNA-helicase | Yes | Biomphalaria virus 3 (28%) | Yes |
| Gastropoda Picorna-like virus N10 | | RNA-helicase | Yes | Biomphalaria virus 3 (27%) | Yes |
| Gastropoda Picorna-like virus N8 | | RdRp | Yes | Beihai picorna-like virus 70 (45%) | Yes |
| Gastropoda Picorna-like virus N7 | | | Yes | Beihai picorna-like virus 15 (76%) | Yes |
| Gastropoda Durna-like virus N1 | | | Yes | Hubei picobirna-like virus 2 (34%) | Yes |
| Crustacea Picorna-like virus N1 | | Capsid, VP4 | | Wenzhou shrimp virus 5 (95%) | Yes |
| Crustacea Picorna-like virus N2 | | | | Wenzhou shrimp virus 5 (97%) | Yes |
| Crustacea Picorna-like virus N5 | | Capsid | Yes | Wenzhou shrimp virus 6 (63%) | Yes |
| Crustacea Picorna-like virus N3 | | | | Wenzhou shrimp virus 5 (94%) | Yes |
| Crustacea Sobeli-like virus N1 | Yes | RdRp | | Wenzhou shrimp virus 9 (100%) | Yes |

| | | | | | |
|---|---|---|---|---|---|
| Crustacea Picorna-like virus N4 | | Peptidase C3 | Yes | Wenzhou shrimp virus 5 (88%) | Yes |
| Crustacea Picorna-like virus D7 | Yes | RdRp | Yes | Beihai blue swimmer crab virus 1 (25%) | Yes |
| Crustacea Picorna-like virus N7 | | | Yes | Wenzhou shrimp virus 8 (76%) | Yes |
| Crustacea Picorna-like virus N8 | | | Yes | Wenzhou shrimp virus 8 (77%) | Yes |
| Crustacea Ghabri-like virus N2 | Yes | RdRp | Yes | Wenling toti-like virus 2 (75%) | Yes |
| Crustacea Picorna-like virus N11 | | Capsid | Yes | Wenzhou shrimp virus 4 (40%) | Yes |
| Gastropoda Picorna-like virus N11 | | | Yes | Beihai picorna-like virus 114 (39%) | Yes |
| Gastropoda Unclassified virus N1 | | | Yes | Corey virus (26%) | Yes |
| Bivalvia Picorna-like virus N9 | | Capsid | | Beihai picorna-like virus 80 (93%) | Yes |
| Bivalvia Picorna-like virus N22 | Yes | RdRp | Yes | Marine RNA virus PAL E4 (28%) | Yes |
| Bivalvia Picorna-like virus N15 | | | Yes | Marine RNA virus BC-1 (33%) | Yes |
| Bivalvia Picorna-like virus D26 | Yes | RdRp | Yes | Beihai picorna-like virus 9 (35%) | Yes |
| Bivalvia Picorna-like virus D84 | Yes | RdRp, Peptidase C3G | Yes | Cragig virus 2 (37%) | Yes |
| Bivalvia Picorna-like virus N26 | | RdRp | Yes | Wenling crustacean virus 6 (38%) | Yes |
| Bivalvia Amarillo-like virus H1 | Yes | RdRp | Yes | Southern pygmy squid flavivirus (36%) | Yes |
| Bivalvia Picorna-like virus H1 | | RNA-helicase | Yes | Beihai narna-like virus 18 (35%) | Yes |
| Bivalvia Bunya-like virus H1 | Yes | RdRp | Yes | Wenling minipizza batfish hantavirus (25%) | Yes |
| Bivalvia Durna-like virus H5 | Yes | RdRp | Yes | Beihai picobirna-like virus 10 (73%) | Yes |
| Bivalvia Durna-like virus H3 | | | Yes | Beihai picobirna-like virus 10 (53%) | Yes |
| Bivalvia Durna-like virus H4 | | | Yes | Beihai picobirna-like virus 13 (44%) | Yes |
| Crustacea Ghabri-like virus H3 | Yes | RdRp | Yes | Penaeid shrimp infectious myonecrosis virus (59%) | Yes |
| Crustacea Bunya-like virus H1 | Yes | RdRp | Yes | Hubei myriapoda virus 5 (21%) | Yes |
| Crustacea Ghabri-like virus H1 | | RdRp | Yes | Murri virus (29%) | Yes |
| Crustacea Ghabri-like virus H2 | Yes | RdRp | Yes | Beihai uca arcuata virus 1 (30%) | Yes |
| Cephalopoda Durna-like virus H1 | | | Yes | Beihai picobirna-like virus 10 (80%) | Yes |
| Cephalopoda Unclassified virus H1 | | RNA-helicase | Yes | Halhan virus 1 (54%) | Yes |
| Cephalopoda Durna-like virus H2 | Yes | RdRp | Yes | Beihai partiti-like virus 4 (48%) | Yes |

| | | | | | |
|---|---|---|---|---|---|
| Cephalopoda Picorna-like virus H1 | | | Yes | Wenling picorna-like virus 2 (45%) | Yes |
| Cephalopoda Mononega-like virus H1 | | | Yes | Eel virus European X (40%) | Yes |
| Cephalopoda Bunya-like virus H1 | | | Yes | Cumuto virus (28%) | |
| Bivalvia Picorna-like virus D63 | Yes | RdRp | Yes | Picornaviridae sp. (24%) | Yes |
| Bivalvia Picorna-like virus N21 | Yes | RdRp | Yes | Beihai octopus virus 2 (39%) | Yes |
| Bivalvia Picorna-like virus H18 | | | Yes | Wenzhou picorna-like virus 4 (41%) | Yes |
| Bivalvia Durna-like virus H6 | Yes | RdRp | Yes | Beihai picobirna-like virus 13 (44%) | Yes |
| Bivalvia Picorna-like virus H8 | | | Yes | Beihai octopus virus 2 (28%) | |
| Bivalvia Sobeli-like virus H3 | Yes | RdRp | Yes | Beihai sobemo-like virus 7 (27%) | Yes |
| Gastropoda Picorna-like virus H3 | | | Yes | Beihai picorna-like virus 35 (61%) | Yes |
| Bivalvia Sobeli-like virus N2 | Yes | RdRp | Yes | Beihai sobemo-like virus 13 (35%) | Yes |
| Bivalvia Bunya-like virus N1 | Yes | RdRp | Yes | Camp Ripley virus (24%) | Yes |
| Bivalvia Picorna-like virus N37 | | | Yes | Lindernia crustacea marnavirus (31%) | Yes |
| Bivalvia Picorna-like virus N38 | | | Yes | Wenzhou picorna-like virus 29 (41%) | Yes |
| Bivalvia Bunya-like virus N3 | Yes | RdRp | Yes | Beihai bunya-like virus 3 (59%) | Yes |
| Bivalvia Picorna-like virus N40 | | RNA-helicase | | Beihai paphia shell virus 3 (98%) | Yes |
| Bivalvia Picorna-like virus N44 | Yes | RdRp | | Beihai paphia shell virus 3 (97%) | Yes |
| Bivalvia Picorna-like virus N39 | | | Yes | Barns Ness breadcrumb sponge aquatic picorna-like virus 3 (42%) | Yes |
| Bivalvia Picorna-like virus N42 | | RNA-helicase | Yes | Picornaviridae sp. (28%) | Yes |
| Bivalvia Picorna-like virus N41 | | Capsid, Capsid | | Beihai paphia shell virus 3 (98%) | Yes |
| Bivalvia Picorna-like virus N45 | | Capsid | Yes | Perth bee virus 8 (23%) | Yes |
| Bivalvia Durna-like virus N4 | Yes | RdRp | Yes | Beihai picobirna-like virus 10 (71%) | Yes |
| Bivalvia Ghabri-like virus N1 | Yes | RdRp | Yes | Penaeid shrimp infectious myonecrosis virus (58%) | Yes |
| Bivalvia Picorna-like virus D35 | | RNA-helicase | Yes | Cragig virus 1 (39%) | Yes |
| Bivalvia Picorna-like virus D36 | | RNA-helicase | Yes | Eel picornavirus Eel/31945 (39%) | Yes |
| Bivalvia Durna-like virus D15 | Yes | RdRp | Yes | Beihai picobirna-like virus 10 (69%) | Yes |
| Bivalvia Picorna-like virus N57 | Yes | RdRp | Yes | Rudphi virus 1 (74%) | Yes |

| | | | | | |
|---|---|---|---|---|---|
| Bivalvia Picorna-like virus D66 | | RNA-helicase | Yes | Antarctic picorna-like virus (35%) | Yes |
| Bivalvia Picorna-like virus N14 | Yes | RdRp | Yes | Biomphalaria virus 4 (32%) | Yes |
| Bivalvia Picorna-like virus D78 | | RNA-helicase | Yes | Marine RNA virus BC-4 (79%) | Yes |
| Bivalvia Picorna-like virus D71 | | | Yes | Beihai picorna-like virus 29 (88%) | Yes |
| Bivalvia Unclassified virus D4 | | RNA-helicase | Yes | Mytedu virus 1 (58%) | Yes |
| Bivalvia Durna-like virus D16 | Yes | RdRp | Yes | Beihai picobirna-like virus 10 (79%) | Yes |
| Bivalvia Picorna-like virus D87 | | RNA-helicase, PTZ00121 | Yes | Wenzhou picorna-like virus 24 (35%) | Yes |
| Bivalvia Hepeli-like virus D3 | Yes | RdRp | Yes | Shahe hepe-like virus 1 (33%) | Yes |
| Bivalvia Picorna-like virus D86 | | RdRp | Yes | Renmark bee virus 3 (47%) | Yes |
| Bivalvia Picorna-like virus D88 | | RNA-helicase | Yes | Wenzhou picorna-like virus 7 (52%) | Yes |
| Bivalvia Picorna-like virus D83 | | Capsid | Yes | Beihai picorna-like virus 62 (38%) | Yes |
| Bivalvia Picorna-like virus D27 | Yes | RdRp | Yes | Dicistroviridae sp. (27%) | Yes |
| Bivalvia Picorna-like virus D89 | | RNA-helicase | Yes | Wenzhou picorna-like virus 7 (70%) | Yes |
| Bivalvia Picorna-like virus D90 | | | Yes | Wenzhou picorna-like virus 7 (36%) | Yes |
| Bivalvia Picorna-like virus D85 | | RdRp | Yes | Picornavirales sp. (38%) | Yes |
| Bivalvia Picorna-like virus D82 | | RdRp | Yes | Beihai picorna-like virus 39 (40%) | Yes |
| Bivalvia Picorna-like virus N1 | | | | Beihai picorna-like virus 56 (100%) | Yes |
| Bivalvia Durna-like virus N1 | Yes | RdRp | Yes | Beihai picobirna-like virus 10 (57%) | Yes |
| Bivalvia Picorna-like virus N3 | | RNA-helicase, RILP-like | Yes | Wenling picorna-like virus 2 (45%) | Yes |
| Crustacea Picorna-like virus N9 | | | | Wenzhou shrimp virus 8 (90%) | Yes |
| Bivalvia Picorna-like virus N10 | | | | Beihai razor shell virus 1 (90%) | Yes |
| Bivalvia Picorna-like virus N30 | | Peptidase C3, RdRp | Yes | Wenzhou picorna-like virus 7 (55%) | Yes |
| Bivalvia Picorna-like virus D24 | Yes | RdRp | Yes | Beihai picorna-like virus 8 (29%) | Yes |
| Bivalvia Picorna-like virus H2 | | Capsid | Yes | Beihai picorna-like virus 111 (36%) | Yes |
| Bivalvia Picorna-like virus H3 | | | Yes | Beihai picorna-like virus 39 (43%) | Yes |
| Bivalvia Picorna-like virus H7 | | | Yes | Renmark bee virus 3 (32%) | Yes |
| Bivalvia Picorna-like virus H6 | | Capsid | Yes | Marnaviridae sp. (39%) | Yes |

| | | | | | |
|---|---|---|---|---|---|
| Crustacea Unclassified virus H1 | | | Yes | San Bernardo virus (25%) | Yes |
| Bivalvia Picorna-like virus H12 | | | Yes | Beihai picorna-like virus 17 (39%) | Yes |
| Bivalvia Picorna-like virus H16 | | Capsid | Yes | Posavirus strain 8729 (24%) | Yes |
| Bivalvia Picorna-like virus H9 | | Capsid | Yes | Beihai octopus virus 2 (34%) | Yes |
| Bivalvia Sobeli-like virus H1 | | K trans | Yes | Hubei sobemo-like virus 3 (35%) | Yes |
| Bivalvia Picorna-like virus H10 | | Capsid | Yes | Beihai octopus virus 2 (35%) | Yes |
| Crustacea Picorna-like virus N19 | Yes | RdRp | | Beihai picorna-like virus 113 (94%) | Yes |
| Bivalvia Picorna-like virus D45 | Yes | RdRp | Yes | Picornavirales sp. (35%) | Yes |
| Bivalvia Picorna-like virus H11 | | RNA-helicase | Yes | Beihai picorna-like virus 115 (34%) | Yes |
| Bivalvia Durna-like virus H7 | Yes | RdRp | Yes | Beihai picobirna-like virus 10 (73%) | Yes |
| Bivalvia Amarillo-like virus N2 | | RNA-helicase, DEAD-like helicase C, glycoprot | Yes | Firefly squid flavivirus (21%) | Yes |
| Bivalvia Picorna-like virus N36 | | | Yes | Cragig virus 6 (48%) | Yes |
| Bivalvia Picorna-like virus N34 | | RNA-helicase, DUF3808 | Yes | Beihai picorna-like virus 17 (32%) | Yes |
| Bivalvia Picorna-like virus N35 | | RNA-helicase | Yes | Beihai picorna-like virus 41 (45%) | Yes |
| Bivalvia Picorna-like virus N33 | | | Yes | Beihai picorna-like virus 14 (32%) | Yes |
| Bivalvia Picorna-like virus D21 | Yes | RdRp | Yes | Beihai picorna-like virus 115 (43%) | Yes |
| Crustacea Hepeli-like virus D1 | Yes | RdRp | Yes | Wenling hepe-like virus 1 (40%) | Yes |
| Bivalvia Picorna-like virus D1 | | | Yes | Bat dicibavirus (23%) | Yes |
| Gastropoda Unclassified virus D1 | | | Yes | Halhan virus 3 (28%) | Yes |
| Gastropoda Picorna-like virus D3 | | RNA-helicase | Yes | Wenzhou picorna-like virus 39 (28%) | Yes |
| Gastropoda Sobeli-like virus D1 | Yes | RdRp | Yes | Wenling sobemo-like virus 1 (46%) | Yes |
| Gastropoda Picorna-like virus D2 | | Peptidase C3 | Yes | Beihai picorna-like virus 70 (31%) | Yes |
| Gastropoda Picorna-like virus D1 | | | | Beihai picorna-like virus 33 (98%) | Yes |
| Gastropoda Unclassified virus D2 | | RNA-helicase | Yes | Linepithema humile virus 1 (35%) | Yes |
| Bivalvia Picorna-like virus D17 | | Capsid | | Beihai picorna-like virus 23 (100%) | Yes |
| Bivalvia Picorna-like virus D18 | | Capsid, VP4 | Yes | Picornavirales N OV 001 (54%) | Yes |
| Hexanauplia Picorna-like virus D2 | | | Yes | Beihai picorna-like virus 84 (38%) | Yes |

| | | | | | |
|---|---|---|---|---|---|
| Hexanauplia Picorna-like virus D3 | | RNA-helicase | Yes | Beihai picorna-like virus 84 (44%) | Yes |
| Hexanauplia Picorna-like virus D4 | | Peptidase C3 | Yes | Beihai picorna-like virus 84 (36%) | Yes |
| Gastropoda Picorna-like virus D29 | | | Yes | Marnaviridae sp. (57%) | Yes |
| Gastropoda Picorna-like virus D27 | | RNA-helicase | Yes | Beihai picorna-like virus 34 (32%) | Yes |
| Gastropoda Picorna-like virus D28 | | | Yes | Marine RNA virus BC-1 (44%) | Yes |
| Gastropoda Durna-like virus D2 | Yes | RdRp | Yes | Benes partiti-like virus (22%) | Yes |
| Gastropoda Durna-like virus D1 | Yes | RdRp | Yes | Beihai picobirna-like virus 7 (55%) | Yes |
| Gastropoda Picorna-like virus D26 | | RNA-helicase | Yes | Beihai picorna-like virus 29 (25%) | Yes |
| Bivalvia Picorna-like virus D73 | | | Yes | Beihai picorna-like virus 39 (44%) | Yes |
| Bivalvia Unclassified virus D3 | | | Yes | Guinardia delicatula RNA virus 01 (47%) | Yes |
| Bivalvia Picorna-like virus D76 | | | Yes | Chaetoceros species RNA virus 02 (56%) | Yes |
| Bivalvia Picorna-like virus D69 | | | Yes | Beihai picorna-like virus 18 (38%) | Yes |
| Bivalvia Picorna-like virus D80 | | RdRp | Yes | Wenzhou channeled applesnail virus 1 (27%) | Yes |
| Bivalvia Picorna-like virus D68 | | | Yes | Beihai picorna-like virus 14 (92%) | Yes |
| Bivalvia Picorna-like virus D67 | | RNA-helicase | | Beihai picorna-like virus 107 (98%) | Yes |
| Bivalvia Toli-like virus D2 | | | Yes | Beihai tombus-like virus 4 (72%) | Yes |
| Bivalvia Picorna-like virus D70 | | | | Beihai picorna-like virus 29 (95%) | Yes |
| Bivalvia Picorna-like virus D16 | Yes | RdRp | Yes | Beihai picorna-like virus 38 (48%) | Yes |
| Bivalvia Picorna-like virus D77 | | | Yes | Chaetoceros species RNA virus 02 (62%) | Yes |
| Crustacea Ghabri-like virus N1 | Yes | RdRp | Yes | Wenling toti-like virus 2 (64%) | Yes |
| Crustacea Picorna-like virus N10 | | | Yes | Wenzhou shrimp virus 4 (31%) | Yes |
| Crustacea Toli-like virus N1 | | RdRp, Capsid, PHA03247 | Yes | Wenzhou tombus-like virus 18 (87%) | Yes |
| Bivalvia Picorna-like virus N11 | | | Yes | Beihai razor shell virus 1 (85%) | Yes |
| Bivalvia Picorna-like virus N12 | | | Yes | Beihai sipunculid worm virus 5 (29%) | Yes |
| Bivalvia Picorna-like virus N16 | | RNA-helicase | Yes | Beihai narna-like virus 19 (41%) | Yes |
| Bivalvia Wolfram-like virus N1 | | RdRp | Yes | Barns Ness serrated wrack narna-like virus 2 (27%) | Yes |
| Bivalvia Durna-like virus N2 | | | Yes | Beihai picobirna-like virus 10 (80%) | Yes |

| | | | | | |
|---|---|---|---|---|---|
| Bivalvia Durna-like virus N3 | Yes | RdRp | Yes | Beihai picobirna-like virus 10 (82%) | Yes |
| Bivalvia Picorna-like virus N25 | | RNA-helicase | Yes | Wenling crustacean virus 5 (38%) | Yes |
| Bivalvia Picorna-like virus N17 | | Capsid, RNA-helicase | Yes | Beihai octopus virus 2 (23%) | Yes |
| Bivalvia Picorna-like virus N18 | | RNA-helicase | Yes | Beihai octopus virus 2 (30%) | Yes |
| Bivalvia Picorna-like virus N24 | | | Yes | Fur seal picorna-like virus (44%) | Yes |
| Bivalvia Picorna-like virus H19 | | | | Wenzhou picorna-like virus 10 (99%) | Yes |
| Bivalvia Bunya-like virus N2 | Yes | RdRp | Yes | Wenling minipizza batfish hantavirus (28%) | Yes |
| Bivalvia Picorna-like virus N31 | | RNA-helicase | Yes | Beihai picorna-like virus 123 (25%) | Yes |
| Bivalvia Picorna-like virus N32 | | Capsid | Yes | Beihai picorna-like virus 123 (31%) | Yes |
| Bivalvia Picorna-like virus N49 | | RdRp | Yes | Beihai octopus virus 2 (49%) | Yes |
| Bivalvia Picorna-like virus N50 | | Capsid, RNA-helicase | Yes | Beihai octopus virus 2 (23%) | Yes |
| Bivalvia Picorna-like virus N51 | | | | Wenzhou gastropodes virus 1 (99%) | Yes |
| Bivalvia Picorna-like virus H4 | Yes | RdRp | Yes | Sanxia water strider virus 21 (29%) | Yes |
| Bivalvia Unclassified virus N3 | | | Yes | Rudphi virus 1 (55%) | Yes |
| Bivalvia Picorna-like virus N53 | | | | Wenzhou gastropodes virus 2 (93%) | Yes |
| Bivalvia Durna-like virus D1 | Yes | RdRp | Yes | Beihai picobirna-like virus 10 (73%) | Yes |
