## Supplementary material for "Viromes in Marine Ecosystems Reveal Remarkable Invertebrate RNA Virus Diversity": The information about viruses used for host sharing estimation

### Table S3 The information about viruses used for host sharing estimation

| No. | Virus Name | Accession | Location | Host Phylum | Host Class |
|---|---|---|---|---|---|
| 1 | Bivalvia Durna-like virus D1 | GWHAPLT01000000 | East China Sea | Mollusca | Bivalvia |
| 2 | Bivalvia Durna-like virus H5 | GWHAPGQ01000000 | Yellow China Sea | Mollusca | Bivalvia |
| 3 | Bivalvia Durna-like virus D7 | GWHAOZQ01000000 | East China Sea | Mollusca | Bivalvia |
| 4 | Bivalvia Durna-like virus N4 | GWHAPHV01000000 | South China Sea | Mollusca | Bivalvia |
| 5 | Beihai mantis shrimp virus 3 strain BHXG26678 hypothetical protein gene complete cds | KX884471.1 | South China Sea | Arthropoda | Crustacea |
| 6 | Bivalvia Picorna-like virus N16 | GWHAPLB01000000 | South China Sea | Mollusca | Bivalvia |
| 7 | Hubei picorna-like virus 6 strain WHBL73048 hypothetical protein 1 and hypothetical protein 2 genes complete cds | KX884262.1 | Freshwater | Mollusca | Bivalvia |
| 8 | Bivalvia Picorna-like virus D85 | GWHAPIP01000000 | East China Sea | Mollusca | Bivalvia |
| 9 | Hubei picorna-like virus 62 strain spider133936 hypothetical protein gene complete cds | KX883686.1 | Terrestrial | Arthropoda | Chelicerata |
| 10 | Hubei picorna-like virus 63 strain insectZJ97544 hypothetical protein gene complete cds | KX883911.1 | Terrestrial | Arthropoda | Insecta |
| 11 | Hubei tetragnatha maxillosa virus 8 strain arthropodmix13417 segment Seg 2 Capsid gene complete cds | KX884047.1 | Terrestrial | Arthropoda | Mix |
| 12 | Hubei tetragnatha maxillosa virus 8 strain QTM24815 segment Seg 2 Capsid gene complete cds | KX884122.1 | Terrestrial | Arthropoda | Insecta |
| 13 | Hubei tetragnatha maxillosa virus 8 strain SSZZ3448 segment Seg 2 Capsid gene complete cds | KX884174.1 | Freshwater | Arthropoda | Chelicerata |
| 14 | Hubei tetragnatha maxillosa virus 8 strain arthropodmix13417 segment Seg 1 RdRp gene complete cds | KX884046.1 | Terrestrial | Arthropoda | Mix |
| 15 | Hubei tetragnatha maxillosa virus 8 strain QTM24815 segment Seg 1 RdRp gene complete cds | KX884121.1 | Terrestrial | Arthropoda | Insecta |
| 16 | Hubei tetragnatha maxillosa virus 8 strain SSZZ3448 segment Seg 1 RdRp gene complete cds | KX884173.1 | Freshwater | Arthropoda | Chelicerata |
| 17 | Gastropoda Picorna-like virus D7 | GWHAPAU01000000 | East China Sea | Mollusca | Gastropoda |
| 18 | Hubei Poty-like virus 1 strain SCM51506 polyprotein gene complete cds | KX884568.1 | Terrestrial | Arthropoda | Insecta |
| 19 | Shangao tombus-like virus 1 strain insectZJ94961 hypothetical protein 1 gene complete cds and hypothetical protei | KX883118.1 | Terrestrial | Arthropoda | Insecta |
| 20 | Wenling picorna-like virus 4 strain WLJQ100990 hypothetical protein 1 and hypothetical protein 2 genes complete | KX884310.1 | East China Sea | Arthropoda | Crustacea |
| 21 | Hubei partiti-like virus 52 strain spider134151 RdRp gene complete cds | KX884165.1 | Terrestrial | Arthropoda | Chelicerata |
| 22 | Hubei virga-like virus 9 strain SCM51642 RdRp hypothetical protein putative capsid protein and hypothetical prote | KX883792.1 | Terrestrial | Arthropoda | Insecta |
| 23 | Banna virus strain QTM104536 Banna VP6 gene complete cds | KX884643.1 | Terrestrial | Arthropoda | Insecta |
| 24 | Bivalvia Picorna-like virus N35 | GWHAPJP01000000 | South China Sea | Mollusca | Bivalvia |
| 25 | Beihai picorna-like virus 41 strain BHJJX25957 hypothetical protein 1 and hypothetical protein 2 genes complete c | KX883376.1 | South China Sea | Arthropoda | Crustacea |
| 26 | Bivalvia Durna-like virus D2 | GWHAOYG0100000 | East China Sea | Mollusca | Bivalvia |
| 27 | Bivalvia Durna-like virus D14 | GWHAPBW0100000 | East China Sea | Mollusca | Bivalvia |
| 28 | Bivalvia Durna-like virus H7 | GWHAPJL01000000 | Yellow China Sea | Mollusca | Bivalvia |
| 29 | Banna virus strain QTM104536 Banna VP8 gene complete cds | KX884645.1 | Terrestrial | Arthropoda | Insecta |
| 30 | Beihai picorna-like virus 52 strain BHNXC41485 hypothetical protein gene complete cds | KX883405.1 | South China Sea | Sipuncula | Sipuncula |
| 31 | Bivalvia Durna-like virus D16 | GWHAPIG01000000 | East China Sea | Mollusca | Bivalvia |
| 32 | Beihai hepe-like virus 4 strain BHZY60842 replicase and hypothetical protein genes complete cds | KX883758.1 | South China Sea | Mollusca | Cephalopoda |
| 33 | Yichang Insect virus strain YCYC01 nucleopasid protein (N) gene complete cds | KM817763.1 | Terrestrial | Arthropoda | Insecta |

| | | | | | |
|---|---|---|---|---|---|
| 34 | Bivalvia Picorna-like virus N34 | GWHAPJO01000000 | South China Sea | Mollusca | Bivalvia |
| 35 | Shahe heteroptera virus 1 strain SHWC01c3606 hypothetical protein gene complete cds | KX883639.1 | Freshwater | Arthropoda | Insecta |
| 36 | Shahe heteroptera virus 1 strain SHWC13624 hypothetical protein gene complete cds | KX883668.1 | Freshwater | Arthropoda | Mix |
| 37 | Bivalvia Unclassified virus N1 | GWHAPEG01000000 | South China Sea | Mollusca | Bivalvia |
| 38 | Wenling hepe-like virus 2 strain WLJQ99021 replicase putative glycoprotein and hypothetical protein genes comple | KX883831.1 | East China Sea | Arthropoda | Crustacea |
| 39 | Crustacea Picorna-like virus N17 | GWHAPDG01000000 | South China Sea | Arthropoda | Crustacea |
| 40 | Bivalvia Picorna-like virus D43 | GWHAOZL01000000 | East China Sea | Mollusca | Bivalvia |
| 41 | Bivalvia Picorna-like virus N21 | GWHAPCF01000000 | South China Sea | Mollusca | Bivalvia |
| 42 | Bivalvia Picorna-like virus H14 | GWHAPHE01000000 | Yellow China Sea | Mollusca | Bivalvia |
| 43 | Bivalvia Picorna-like virus N18 | GWHAPLH01000000 | South China Sea | Mollusca | Bivalvia |
| 44 | Bivalvia Picorna-like virus D73 | GWHAPKL01000000 | East China Sea | Mollusca | Bivalvia |
| 45 | Wuhan insect virus 14 strain WHZM10168 polyprotein gene complete cds | KX883007.1 | Terrestrial | Arthropoda | Insecta |
| 46 | Bivalvia Unclassified virus D3 | GWHAPKM0100000( | East China Sea | Mollusca | Bivalvia |
| 47 | Beihai paphia shell virus 1 strain BHZY60696 hypothetical protein 1 and hypothetical protein 2 genes complete cds | KX884512.1 | South China Sea | Mollusca | Cephalopoda |
| 48 | Beihai paphia shell virus 1 strain BWBFG40339 hypothetical protein 1 and hypothetical protein 2 genes complete c | KX884527.1 | South China Sea | Mollusca | Bivalvia |
| 49 | Hubei picorna-like virus 67 strain WGML147056 hypothetical protein 1 hypothetical protein 2 and hypothetical pro | KX884255.1 | Terrestrial | Arthropoda | Myriapoda |
| 50 | Bivalvia Picorna-like virus D6 | GWHAOYC01000000 | East China Sea | Mollusca | Bivalvia |
| 51 | Bole Tick Virus 2 strain BL076 nucleocapsid (N) putative phosphoprotein (ORF2) matrix protein (M) glycoprotein | KM817629.1 | Terrestrial | Arthropoda | Chelicerata |
| 52 | Crustacea Picorna-like virus N33 | GWHAPDS01000000 | South China Sea | Arthropoda | Crustacea |
| 53 | Crustacea Picorna-like virus N6 | GWHAPGA01000000 | South China Sea | Arthropoda | Crustacea |
| 54 | Crustacea Picorna-like virus N8 | GWHAPGC01000000 | South China Sea | Arthropoda | Crustacea |
| 55 | Crustacea Picorna-like virus N7 | GWHAPGB01000000 | South China Sea | Arthropoda | Crustacea |
| 56 | Bivalvia Picorna-like virus D33 | GWHAPEQ01000000 | East China Sea | Mollusca | Bivalvia |
| 57 | Beihai picorna-like virus 70 strain BHWZXX15680 hypothetical protein 1 and hypothetical protein 2 genes comple | KX883433.1 | South China Sea | Arthropoda | Crustacea |
| 58 | Beihai picorna-like virus 70 strain BHHK131042 hypothetical protein 1 and hypothetical protein 2 genes complete | KX883350.1 | South China Sea | Cnidaria | Anthozoa |
| 59 | Beihai picorna-like virus 70 strain BHJJX25952 hypothetical protein 1 and hypothetical protein 2 genes complete c | KX883375.1 | South China Sea | Arthropoda | Crustacea |
| 60 | Beihai picorna-like virus 70 strain BHBei76899 hypothetical protein 1 gene complete cds and hypothetical protein 2 | KX883315.1 | South China Sea | Mollusca | Bivalvia |
| 61 | Beihai picorna-like virus 70 strain shrimp14548 hypothetical protein 1 and hypothetical protein 2 genes complete c | KX883986.1 | East China Sea | Arthropoda | Crustacea |
| 62 | Gastropoda Picorna-like virus D20 | GWHAPBH01000000 | East China Sea | Mollusca | Gastropoda |
| 63 | Gastropoda Picorna-like virus D27 | GWHAPKG01000000 | East China Sea | Mollusca | Gastropoda |
| 64 | Beihai barnacle virus 7 strain BHTH16013 hypothetical protein 1 hypothetical protein 2 hypothetical protein 3 puta | KX884411.1 | South China Sea | Arthropoda | Crustacea |
| 65 | Bivalvia Durna-like virus H3 | GWHAPGR01000000 | Yellow China Sea | Mollusca | Bivalvia |
| 66 | Gastropoda Picorna-like virus D21 | GWHAPBJ01000000 | East China Sea | Mollusca | Gastropoda |
| 67 | Gastropoda Picorna-like virus D24 | GWHAPBI01000000 | East China Sea | Mollusca | Gastropoda |
| 68 | Wenzhou picorna-like virus 26 strain WZSLuoII93478 hypothetical protein 1 and hypothetical protein 2 genes com | KX884379.1 | East China Sea | Mollusca | Gastropoda |
| 69 | Beihai picorna-like virus 71 strain BHXG26494 hypothetical protein 1 and hypothetical protein 2 genes complete c | KX884469.1 | South China Sea | Arthropoda | Crustacea |
| 70 | Polychaeta Picorna-like virus H3 | GWHAPCX01000000 | Yellow China Sea | Annelida | Polychaeta |

| | | | | | |
|---|---|---|---|---|---|
| 71 | Beihai picorna-like virus 85 strain BHHK129719 hypothetical protein 1 and hypothetical protein 2 genes complete | KX883346.1 | South China Sea | Cnidaria | Anthozoa |
| 72 | Beihai sipunculid worm virus 2 strain BHNXC41483 hypothetical protein 1 and hypothetical protein 2 genes compl | KX883404.1 | South China Sea | Sipuncula | Sipuncula |
| 73 | Beihai shrimp virus 5 strain BHWZXX15615 segment 1 hypothetical protein 1 and hypothetical protein 2 genes cor | KX882792.1 | South China Sea | Arthropoda | Crustacea |
| 74 | Beihai shrimp virus 5 strain BHBJDX18804 segment 1 hypothetical protein 2 and hypothetical protein 1 genes parti | KX882784.1 | South China Sea | Arthropoda | Crustacea |
| 75 | Bivalvia Picorna-like virus D77 | GWHAPKV01000000 | East China Sea | Mollusca | Bivalvia |
| 76 | Beihai picorna-like virus 4 strain BHZY60671 hypothetical protein 1 and hypothetical protein 2 genes complete cds | KX884510.1 | South China Sea | Mollusca | Cephalopoda |
| 77 | Hubei diptera virus 21 strain SCM95013 hypothetical protein 4 gene complete cds | KX884700.1 | Terrestrial | Arthropoda | Insecta |
| 78 | Wuhan house centipede virus 2 strain arthropodmix22554 hypothetical protein gene complete cds | KX883295.1 | Terrestrial | Arthropoda | Mix |
| 79 | Wuhan house centipede virus 2 strain WHYY32989 hypothetical protein gene complete cds | KX884299.1 | Terrestrial | Arthropoda | Myriapoda |
| 80 | Wenzhou picorna-like virus 37 strain WZSBei69579 hypothetical protein 1 and hypothetical protein 2 genes compl | KX884349.1 | East China Sea | Mollusca | Bivalvia |
| 81 | Hubei diptera virus 21 strain SCM95013 hypothetical protein 5 gene complete cds | KX884701.1 | Terrestrial | Arthropoda | Insecta |
| 82 | Hubei diptera virus 18 strain SCM37169 segment Seg 1 RdRp gene complete cds | KX884150.1 | Terrestrial | Arthropoda | Insecta |
| 83 | Hubei diptera virus 18 strain QTM25917 segment Seg 1 RdRp gene complete cds | KX884112.1 | Terrestrial | Arthropoda | Insecta |
| 84 | Bivalvia Picorna-like virus N10 | GWHAPIV01000000 | South China Sea | Mollusca | Bivalvia |
| 85 | Beihai razor shell virus 1 strain BHZC37272 hypothetical protein 1 and hypothetical protein 2 genes complete cds | KX884486.1 | South China Sea | Mollusca | Bivalvia |
| 86 | Beihai razor shell virus 1 strain BHBei76693 hypothetical protein genes complete cds | KX883312.1 | South China Sea | Mollusca | Bivalvia |
| 87 | Beihai razor shell virus 1 strain BHBJDX18719 hypothetical protein 1 and hypothetical protein 2 genes complete c | KX883333.1 | South China Sea | Arthropoda | Crustacea |
| 88 | Wenling partiti-like virus 3 strain WLJQ124021 RdRp gene complete cds | KX884230.1 | East China Sea | Arthropoda | Crustacea |
| 89 | Beihai sipunculid worm virus 1 strain BHNXC40689 hypothetical protein 1 and hypothetical protein 2 genes compl | KX883388.1 | South China Sea | Sipuncula | Sipuncula |
| 90 | Gastropoda Picorna-like virus D1 | GWHAPJY01000000 | East China Sea | Mollusca | Gastropoda |
| 91 | Beihai picorna-like virus 33 strain BWBFG40845 hypothetical protein 1 and hypothetical protein 2 genes complete | KX884530.1 | South China Sea | Mollusca | Bivalvia |
| 92 | Gastropoda Picorna-like virus H3 | GWHAPHJ01000000 | Yellow China Sea | Mollusca | Gastropoda |
| 93 | Beihai picorna-like virus 35 strain BHTH15623 hypothetical protein 1 and hypothetical protein 2 genes complete c | KX883412.1 | South China Sea | Arthropoda | Crustacea |
| 94 | Beihai picorna-like virus 35 strain BHWZXX15273 hypothetical protein 1 and hypothetical protein 2 genes comple | KX883425.1 | South China Sea | Arthropoda | Crustacea |
| 95 | Beihai picorna-like virus 35 strain SCJXSX39018 hypothetical protein 1 and hypothetical protein 2 genes complete | KX883960.1 | South China Sea | Arthropoda | Crustacea |
| 96 | Beihai picorna-like virus 35 strain YYSZX17573 hypothetical protein 1 and hypothetical protein 2 genes complete | KX884388.1 | South China Sea | Arthropoda | Crustacea |
| 97 | Beihai picorna-like virus 35 strain BHTSS17888 hypothetical protein 1 and hypothetical protein 2 genes complete c | KX883420.1 | South China Sea | Arthropoda | Crustacea |
| 98 | Beihai picorna-like virus 35 strain BHXG26244 hypothetical protein 1 and hypothetical protein 2 genes complete c | KX884467.1 | South China Sea | Arthropoda | Crustacea |
| 99 | Beihai picorna-like virus 35 strain BHJJX23572 hypothetical protein 1 and hypothetical protein 2 genes complete c | KX883363.1 | South China Sea | Arthropoda | Crustacea |
| 100 | Beihai picorna-like virus 35 strain BHHK128006 hypothetical protein 1 and hypothetical protein 2 genes complete | KX883345.1 | South China Sea | Cnidaria | Anthozoa |
| 101 | Beihai picorna-like virus 35 strain BHBJDX17224 hypothetical protein 1 and hypothetical protein 2 genes complete | KX883323.1 | South China Sea | Arthropoda | Crustacea |
| 102 | Beihai picorna-like virus 35 strain BHNXC40527 hypothetical protein 1 and hypothetical protein 2 genes complete | KX883386.1 | South China Sea | Sipuncula | Sipuncula |
| 103 | Beihai picorna-like virus 35 strain BZL82652 hypothetical protein 1 and hypothetical protein 2 genes complete cds | KX884534.1 | South China Sea | Mollusca | Gastropoda |
| 104 | Beihai picorna-like virus 35 strain HWRTX12817 hypothetical protein 1 and hypothetical protein 2 genes complete | KX883907.1 | South China Sea | Arthropoda | Crustacea |
| 105 | Beihai picorna-like virus 35 strain BHZC37213 hypothetical protein 1 and hypothetical protein 2 genes complete cd | KX884483.1 | South China Sea | Mollusca | Bivalvia |
| 106 | Aphid lethal paralysis virus strain QTM16 hypothetical protein 1 and hypothetical protein 2 genes complete cds | KX883929.1 | Terrestrial | Arthropoda | Insecta |
| 107 | Aphid lethal paralysis virus strain SCM51524 hypothetical protein 1 and hypothetical protein 2 genes complete cds | KX883977.1 | Terrestrial | Arthropoda | Insecta |

| | | | | | |
|---|---|---|---|---|---|
| ***108*** | Aphid lethal paralysis virus strain spider133997 hypothetical protein 1 and hypothetical protein 2 genes complete cd | KX883690.1 | Terrestrial | Arthropoda | Chelicerata |
| ***109*** | Aphid lethal paralysis virus strain WHCCII13327 hypothetical protein 1 and hypothetical protein 2 genes complete | KX884276.1 | Terrestrial | Arthropoda | Insecta |
| ***110*** | Hubei partiti-like virus 24 strain WHYY17134 RdRp gene partial cds | KX884214.1 | Terrestrial | Arthropoda | Myriapoda |
| ***111*** | Wuhan fly virus 5 strain fly34516 segment Seg 2 hypothetical protein gene complete cds | KX884092.1 | Terrestrial | Arthropoda | Insecta |
| ***112*** | Kadipiro virus strain QTM27331 Kadipiro VP6 gene complete cds | KX884655.1 | Terrestrial | Arthropoda | Insecta |
| ***113*** | Bivalvia Toli-like virus N2 | GWHAPEI01000000 | South China Sea | Mollusca | Bivalvia |
| ***114*** | Beihai tombus-like virus 4 strain YYSZX17783 hypothetical protein 1 hypothetical protein 2 and hypothetical prote | KX883281.1 | South China Sea | Arthropoda | Crustacea |
| ***115*** | Beihai tombus-like virus 4 strain BHZY58951 hypothetical protein 1 hypothetical protein 2 and hypothetical protein | KX883069.1 | South China Sea | Mollusca | Cephalopoda |
| ***116*** | Beihai tombus-like virus 4 strain BZL81307 hypothetical protein 1 and hypothetical protein 2 genes complete cds | KX883079.1 | South China Sea | Mollusca | Gastropoda |
| ***117*** | Hubei Wuhan insect virus 9 strain QTM27230 hypothetical protein genes complete cds | KX883782.1 | Terrestrial | Arthropoda | Insecta |
| ***118*** | Wuhan insect virus 9 strain WHCCII13328 RdRp and hypothetical protein genes complete cds | KX883816.1 | Terrestrial | Arthropoda | Insecta |
| ***119*** | Hubei Wuhan insect virus 9 strain QTM27230 hypothetical protein genes complete cds | KX883781.1 | Terrestrial | Arthropoda | Insecta |
| ***120*** | Hubei virga-like virus 7 strain QTM27262 hypothetical protein genes complete cds | KX883783.1 | Terrestrial | Arthropoda | Insecta |
| ***121*** | Bivalvia Picorna-like virus D19 | GWHAPEP01000000 | East China Sea | Mollusca | Bivalvia |
| ***122*** | Bivalvia Picorna-like virus N36 | GWHAPJN01000000 | South China Sea | Mollusca | Bivalvia |
| ***123*** | Wenzhou picorna-like virus 52 strain beimix75583 hypothetical protein 1 and hypothetical protein 2 genes complete | KX883300.1 | East China Sea | Mollusca | Bivalvia |
| ***124*** | Bivalvia Unclassified virus D4 | GWHAPIF01000000 | East China Sea | Mollusca | Bivalvia |
| ***125*** | Hubei virga-like virus 18 strain fly97447 RdRp and hypothetical protein genes complete cds | KX883767.1 | Terrestrial | Arthropoda | Insecta |
| ***126*** | Bivalvia Picorna-like virus H12 | GWHAPJD01000000 | Yellow China Sea | Mollusca | Bivalvia |
| ***127*** | Wenling crustacean virus 5 strain WLJQ103138 hypothetical protein 1 and hypothetical protein 2 genes complete cd | KX884317.1 | East China Sea | Arthropoda | Crustacea |
| ***128*** | Crustacea Picorna-like virus N18 | GWHAPDM0100000C | South China Sea | Arthropoda | Crustacea |
| ***129*** | Beihai picorna-like virus 113 strain BHXG26415 hypothetical protein 1 and hypothetical protein 2 genes complete | KX884468.1 | South China Sea | Arthropoda | Crustacea |
| ***130*** | Crustacea Hepeli-like virus D3 | GWHAPEK01000000 | East China Sea | Arthropoda | Crustacea |
| ***131*** | Bivalvia Ghabri-like virus N1 | GWHAPHW0100000( | South China Sea | Mollusca | Bivalvia |
| ***132*** | Crustacea Ghabri-like virus H3 | GWHAPGT01000000 | Yellow China Sea | Arthropoda | Crustacea |
| ***133*** | Wenzhou picorna-like virus 19 strain WZSLuoII97616 hypothetical protein 1 and hypothetical protein 2 genes comp | KX884381.1 | East China Sea | Mollusca | Gastropoda |
| ***134*** | Bivalvia Picorna-like virus H18 | GWHAPHF01000000 | Yellow China Sea | Mollusca | Bivalvia |
| ***135*** | Wenzhou picorna-like virus 44 strain WZSLuoII91200 hypothetical protein gene complete cds | KX884378.1 | East China Sea | Mollusca | Gastropoda |
| ***136*** | Bivalvia Picorna-like virus H19 | GWHAPLJ01000000 | Yellow China Sea | Mollusca | Bivalvia |
| ***137*** | Wenzhou picorna-like virus 10 strain WZSLuoI86114 hypothetical protein 1 and hypothetical protein 2 genes comp | KX884376.1 | East China Sea | Mollusca | Gastropoda |
| ***138*** | Wenzhou picorna-like virus 10 strain WZRBX43164 hypothetical protein 1 and hypothetical protein 2 genes compl | KX884346.1 | East China Sea | Arthropoda | Crustacea |
| ***139*** | Bivalvia Picorna-like virus D11 | GWHAOYB01000000 | East China Sea | Mollusca | Bivalvia |
| ***140*** | Wenzhou picorna-like virus 29 strain BHBei77092 hypothetical protein 1 and hypothetical protein 2 genes complete | KX883319.1 | South China Sea | Mollusca | Bivalvia |
| ***141*** | Wenzhou picorna-like virus 29 strain WZSBei69577 hypothetical protein 1 and hypothetical protein 2 genes compl | KX884350.1 | East China Sea | Mollusca | Bivalvia |
| ***142*** | Gastropoda Picorna-like virus D16 | GWHAPAY01000000 | East China Sea | Mollusca | Gastropoda |
| ***143*** | Wuhan spirurian nematodes virus 1 strain WHSWHC143244 hypothetical protein gene complete cds | KX884291.1 | Terrestrial | Nematoda | Chromadorea |
| ***144*** | Beihai picorna-like virus 26 strain BZL93356 hypothetical protein 1 and hypothetical protein 2 genes complete cds | KX884536.1 | South China Sea | Mollusca | Gastropoda |

| | | | | | |
|---|---|---|---|---|---|
| 145 | Beihai partiti-like virus 2 strain BHZY60797 segment Seg 1 RdRp gene complete cds | KX884083.1 | South China Sea | Mollusca | Cephalopoda |
| 146 | Hubei partiti-like virus 20 strain QTM74833 RdRp gene complete cds | KX884140.1 | Terrestrial | Arthropoda | Insecta |
| 147 | Crustacea Ghabri-like virus N2 | GWHAPGD01000000 | South China Sea | Arthropoda | Crustacea |
| 148 | Wenling toti-like virus 2 strain WLJQ104148 hypothetical protein 1 and hypothetical protein 2 genes complete cds | KX882992.1 | East China Sea | Arthropoda | Crustacea |
| 149 | Wenzhou shrimp virus 9 strain shrimp6189 segment 2 putative capsid protein gene complete cds | KX882861.1 | East China Sea | Arthropoda | Crustacea |
| 150 | Gastropoda Picorna-like virus N2 | GWHAPFF01000000 | South China Sea | Mollusca | Gastropoda |
| 151 | Gastropoda Picorna-like virus N7 | GWHAPFS01000000 | South China Sea | Mollusca | Gastropoda |
| 152 | Beihai picorna-like virus 15 strain BZL81977 hypothetical protein 1 and hypothetical protein 2 genes complete cds | KX884533.1 | South China Sea | Mollusca | Gastropoda |
| 153 | Beihai picorna-like virus 15 strain YYSZX17489 hypothetical protein 1 and hypothetical protein 2 genes complete ( | KX884387.1 | South China Sea | Arthropoda | Crustacea |
| 154 | Beihai picorna-like virus 15 strain BHZC36638 hypothetical protein 1 and hypothetical protein 2 genes complete cd | KX884479.1 | South China Sea | Mollusca | Bivalvia |
| 155 | Beihai picorna-like virus 15 strain BHZY59344 hypothetical protein 1 and hypothetical protein 2 genes complete cc | KX884506.1 | South China Sea | Mollusca | Cephalopoda |
| 156 | Beihai picorna-like virus 15 strain HWRTX12823 hypothetical protein 1 and hypothetical protein 2 genes complete | KX883908.1 | South China Sea | Arthropoda | Crustacea |
| 157 | Beihai picorna-like virus 15 strain BHBJDX18577 hypothetical protein 1 and hypothetical protein 2 genes complete | KX883330.1 | South China Sea | Arthropoda | Crustacea |
| 158 | Beihai picorna-like virus 15 strain BHTSS17870 hypothetical protein 1 and hypothetical protein 2 genes complete c | KX883418.1 | South China Sea | Arthropoda | Crustacea |
| 159 | Beihai picorna-like virus 15 strain BHCL77416 hypothetical protein 1 and hypothetical protein 2 genes complete cd | KX883339.1 | South China Sea | Mollusca | Gastropoda |
| 160 | Beihai picorna-like virus 15 strain BWBFG37850 hypothetical protein 1 and hypothetical protein 2 genes complete | KX884524.1 | South China Sea | Mollusca | Bivalvia |
| 161 | Beihai picorna-like virus 15 strain BHXG24735 hypothetical protein 1 and hypothetical protein 2 genes complete c | KX884464.1 | South China Sea | Arthropoda | Crustacea |
| 162 | Beihai picorna-like virus 15 strain BHXun32793 hypothetical protein 1 and hypothetical protein 2 genes complete c | KX884474.1 | South China Sea | Arthropoda | Crustacea |
| 163 | Beihai picorna-like virus 15 strain BHNXC41227 hypothetical protein 1 and hypothetical protein 2 genes complete | KX883391.1 | South China Sea | Sipuncula | Sipuncula |
| 164 | Beihai picorna-like virus 15 strain BHJJX25823 hypothetical protein 1 and hypothetical protein 2 genes complete c | KX883370.1 | South China Sea | Arthropoda | Crustacea |
| 165 | Beihai picorna-like virus 15 strain BHWZXX15642 hypothetical protein 1 and hypothetical protein 2 genes comple | KX883431.1 | South China Sea | Arthropoda | Crustacea |
| 166 | Bivalvia Picorna-like virus D2 | GWHAPEL01000000 | East China Sea | Mollusca | Bivalvia |
| 167 | Bivalvia Picorna-like virus D60 | GWHAPAD01000000 | East China Sea | Mollusca | Bivalvia |
| 168 | Gastropoda Picorna-like virus D19 | GWHAPBK01000000 | East China Sea | Mollusca | Gastropoda |
| 169 | Beihai barnacle virus 12 strain BHTH16091 segment Seg 2 putative capsid gene complete cds | KX884068.1 | South China Sea | Arthropoda | Crustacea |
| 170 | Xingshan cricket virus strain XSXS-2 polyprotein gene complete cds | KR902738.1 | Terrestrial | Arthropoda | Insecta |
| 171 | Bivalvia Picorna-like virus D84 | GWHAPIM01000000 | East China Sea | Mollusca | Bivalvia |
| 172 | Wenling nido-like virus 1 strain WLJQ99370 hypothetical protein putative glycoprotein 1 and putative glycoprotein | KX883636.1 | East China Sea | Arthropoda | Crustacea |
| 173 | Bivalvia Picorna-like virus N37 | GWHAPHM0100000( | South China Sea | Mollusca | Bivalvia |
| 174 | Bivalvia Unclassified virus N2 | GWHAPED01000000 | South China Sea | Mollusca | Bivalvia |
| 175 | Polychaeta Picorna-like virus H2 | GWHAPCV01000000 | Yellow China Sea | Annelida | Polychaeta |
| 176 | Tacheng Tick Virus 7 strain TCRP-3 ORF1 (ORF1) ORF2 (ORF2) ORF3 (ORF3) ORF4 (ORF4) glycoprotein (G) | KM817642.1 | Terrestrial | Arthropoda | Chelicerata |
| 177 | Wenling crustacean virus 9 strain WLJQ100911 putative nucleoprotein gene complete cds | KX884858.1 | East China Sea | Arthropoda | Crustacea |
| 178 | Gastropoda Picorna-like virus N6 | GWHAPFM01000000 | South China Sea | Mollusca | Gastropoda |
| 179 | Shayang ascaridia galli virus 2 strain SYJFC48550 putative nucleoprotein hypothetical protein putative glycoprotei | KX884440.1 | Terrestrial | Nematoda | Chromadorea |
| 180 | Shayang ascaridia galli virus 2 strain HC21241 hypothetical protein 1 hypothetical protein 2 hypothetical protein 3 l | KX884414.1 | Terrestrial | Nematoda | Rhabditida |
| 181 | Banna virus strain QTM104536 Banna VP11 gene complete cds | KX884649.1 | Terrestrial | Arthropoda | Insecta |

| | | | | | |
|---|---|---|---|---|---|
| 182 | Gastropoda Stella-like virus N1 | GWHAPFK01000000 | South China Sea | Mollusca | Gastropoda |
| 183 | Crustacea Picorna-like virus N1 | GWHAPFU01000000 | South China Sea | Arthropoda | Crustacea |
| 184 | Wenzhou shrimp virus 5 strain shrimp14323 hypothetical protein 1 and hypothetical protein 2 genes complete cds | KX883979.1 | East China Sea | Arthropoda | Crustacea |
| 185 | Crustacea Picorna-like virus N2 | GWHAPFV01000000 | South China Sea | Arthropoda | Crustacea |
| 186 | Crustacea Picorna-like virus N4 | GWHAPFZ01000000 | South China Sea | Arthropoda | Crustacea |
| 187 | Hubei rhabdo-like virus 9 strain WHZHC73015 hypothetical protein 1 hypothetical protein 2 hypothetical protein 3 | KX884448.1 | Terrestrial | Nematoda | Chromadorea |
| 188 | Gastropoda Picorna-like virus N11 | GWHAPGF01000000 | South China Sea | Mollusca | Gastropoda |
| 189 | Wuhan Mosquito Virus 9 strain JX1-13 ORF1 (ORF1) ORF2 (ORF2) ORF3 (ORF3) glycoprotein (G) and RNA-de | KM817659.1 | Terrestrial | Arthropoda | Insecta |
| 190 | Wuhan Fly Virus 1 strain SYY1-9 nucleopasid protein (N) gene complete cds | KM817749.1 | Terrestrial | Arthropoda | Insecta |
| 191 | Wuhan Insect virus 5 strain YCYC02 nucleocapsid (N) phosphoprotein (P) 4b protein (4b) matrix protein (M) glyco | KM817651.1 | Terrestrial | Arthropoda | Insecta |
| 192 | Gastropoda Picorna-like virus D4 | GWHAOYJ01000000 | East China Sea | Mollusca | Gastropoda |
| 193 | Wuhan Tick Virus 2 strain X78-1 complete genome | KM817611.1 | Terrestrial | Arthropoda | Chelicerata |
| 194 | Bivalvia Picorna-like virus N17 | GWHAPLG01000000 | South China Sea | Mollusca | Bivalvia |
| 195 | Bivalvia Picorna-like virus H8 | GWHAPHH01000000 | Yellow China Sea | Mollusca | Bivalvia |
| 196 | Beihai hepe-like virus 9 strain HOU157038 hypothetical protein genes complete cds | KX883769.1 | South China Sea | Arthropoda | Chelicerata |
| 197 | Beihai rhabdo-like virus 3 strain BHNXC39077 putative nucleoprotein hypothetical protein 1 hypothetical protein 2 | KX884408.1 | South China Sea | Sipuncula | Sipuncula |
| 198 | Wuhan Louse Fly Virus 9 strain BFJSC-7 nucleocapsid (N) putative phosphoprotein (ORF2) matrix protein (M) gly | KM817656.1 | Terrestrial | Arthropoda | Insecta |
| 199 | Xinzhou Spider Virus strain XZZZ-2 nucleopasid protein (N) gene complete cds | KM817762.1 | Terrestrial | Arthropoda | Chelicerata |
| 200 | Hubei orthoptera virus 6 strain ZCM21154 segment 4 hemagglutinin gene complete cds | KX883887.1 | Terrestrial | Arthropoda | Insecta |
| 201 | Hubei orthomyxo-like virus 2 strain QTM27225 segment 1 polymerase PB2 gene complete cds | KX883869.1 | Terrestrial | Arthropoda | Insecta |
| 202 | Beihai picorna-like virus 20 strain BHJJX25607 hypothetical protein 1 and hypothetical protein 2 genes complete c | KX883365.1 | South China Sea | Arthropoda | Crustacea |
| 203 | Bivalvia Picorna-like virus H16 | GWHAPJE01000000 | Yellow China Sea | Mollusca | Bivalvia |
| 204 | Bivalvia Picorna-like virus D12 | GWHAOYD0100000 | East China Sea | Mollusca | Bivalvia |
| 205 | Beihai picorna-like virus 121 strain BHHZL10364 hypothetical protein 1 and hypothetical protein 2 genes complete | KX883360.1 | South China Sea | Arthropoda | Crustacea |
| 206 | Bivalvia Picorna-like virus H3 | GWHAPIZ01000000 | Yellow China Sea | Mollusca | Bivalvia |
| 207 | Beihai picorna-like virus 8 strain BHNXC41454 hypothetical protein 1 and hypothetical protein 2 genes complete c | KX883401.1 | South China Sea | Sipuncula | Sipuncula |
| 208 | Bivalvia Picorna-like virus D70 | GWHAPKT01000000 | East China Sea | Mollusca | Bivalvia |
| 209 | Beihai picorna-like virus 29 strain HOU158675 hypothetical protein 1 gene complete cds and hypothetical protein 2 | KX883898.1 | South China Sea | Arthropoda | Chelicerata |
| 210 | Bivalvia Unclassified virus D2 | GWHAOZB01000000 | East China Sea | Mollusca | Bivalvia |
| 211 | Crustacea Unclassified virus N1 | GWHAPDT01000000 | South China Sea | Arthropoda | Crustacea |
| 212 | Bivalvia Picorna-like virus D30 | GWHAOYQ0100000 | East China Sea | Mollusca | Bivalvia |
| 213 | Beihai sipunculid worm virus 4 strain BHBJDX17813 hypothetical protein 1 and hypothetical protein 2 genes comp | KX883325.1 | South China Sea | Arthropoda | Crustacea |
| 214 | Beihai sipunculid worm virus 4 strain BHNXC41347 hypothetical protein 1 and hypothetical protein 2 genes compl | KX883396.1 | South China Sea | Sipuncula | Sipuncula |
| 215 | Beihai sipunculid worm virus 4 strain BHZC37368 hypothetical protein 1 and hypothetical protein 2 genes complete | KX884491.1 | South China Sea | Mollusca | Bivalvia |
| 216 | Gastropoda Picorna-like virus D5 | GWHAPAR01000000 | East China Sea | Mollusca | Gastropoda |
| 217 | Xinzhou nematode virus 5 strain XZSJSC65765 hypothetical protein and RNA-dependent RNA polymerase genes c | KX884461.1 | Terrestrial | Nematoda | Mix |
| 218 | Bivalvia Picorna-like virus D72 | GWHAPKU01000000 | East China Sea | Mollusca | Bivalvia |

| | | | | | |
|---|---|---|---|---|---|
| 219 | Crustacea Picorna-like virus D2 | GWHAPBS01000000 | East China Sea | Arthropoda | Crustacea |
| 220 | Beihai picorna-like virus 19 strain BHJJX24523 hypothetical protein 1 and hypothetical protein 2 genes complete c | KX883364.1 | South China Sea | Arthropoda | Crustacea |
| 221 | Gastropoda Picorna-like virus D11 | GWHAPAO01000000 | East China Sea | Mollusca | Gastropoda |
| 222 | Hubei tetragnatha maxillosa virus 4 strain arthropodmix13651 hypothetical protein gene complete cds | KX883289.1 | Terrestrial | Arthropoda | Mix |
| 223 | Hubei tetragnatha maxillosa virus 4 strain SSZZ3380 hypothetical protein gene complete cds | KX883697.1 | Freshwater | Arthropoda | Chelicerata |
| 224 | Hubei partiti-like virus 48 strain QCM147194 RdRp gene complete cds | KX884105.1 | Terrestrial | Arthropoda | Insecta |
| 225 | Bivalvia Picorna-like virus N11 | GWHAPKZ01000000 | South China Sea | Mollusca | Bivalvia |
| 226 | Hubei virga-like virus 10 strain WHYY22294 hypothetical protein gene complete cds and hypothetical protein gene | KX883824.1 | Terrestrial | Arthropoda | Myriapoda |
| 227 | Hubei tetragnatha maxillosa virus 9 strain QTM27093 hypothetical protein 5 gene complete cds | KX884680.1 | Terrestrial | Arthropoda | Insecta |
| 228 | Hubei tetragnatha maxillosa virus 9 strain arthropodmix14049 hypothetical protein 5 gene complete cds | KX884603.1 | Terrestrial | Arthropoda | Mix |
| 229 | Crustacea Picorna-like virus N24 | GWHAPDL01000000 | South China Sea | Arthropoda | Crustacea |
| 230 | Crustacea Picorna-like virus N26 | GWHAPDO01000000 | South China Sea | Arthropoda | Crustacea |
| 231 | Wuhan insect virus 23 strain WHCCII13263 segment Seg 2 hypothetical protein gene complete cds | KX884197.1 | Terrestrial | Arthropoda | Insecta |
| 232 | Beihai picorna-like virus 39 strain BHSC164089 hypothetical protein 1 and hypothetical protein 2 genes complete c | KX883410.1 | South China Sea | Annelida | Polychaeta |
| 233 | Bivalvia Picorna-like virus D7 | GWHAOXW0100000( | East China Sea | Mollusca | Bivalvia |
| 234 | Bivalvia Picorna-like virus D20 | GWHAOYO0100000C | East China Sea | Mollusca | Bivalvia |
| 235 | Crustacea Picorna-like virus N21 | GWHAPDK01000000 | South China Sea | Arthropoda | Crustacea |
| 236 | Crustacea Picorna-like virus N23 | GWHAPDP01000000 | South China Sea | Arthropoda | Crustacea |
| 237 | Bivalvia Picorna-like virus D64 | GWHAPAG01000000 | East China Sea | Mollusca | Bivalvia |
| 238 | Bivalvia Picorna-like virus D54 | GWHAOZS01000000 | East China Sea | Mollusca | Bivalvia |
| 239 | Bivalvia Picorna-like virus N52 | GWHAPEA01000000 | South China Sea | Mollusca | Bivalvia |
| 240 | Bivalvia Picorna-like virus N27 | GWHAPCH01000000 | South China Sea | Mollusca | Bivalvia |
| 241 | Hubei virga-like virus 23 strain mosHB236486 RdRp hypothetical protein putative glycoprotein and hypothetical pr | KX883774.1 | Terrestrial | Arthropoda | Insecta |
| 242 | Bivalvia Durna-like virus N2 | GWHAPLD01000000 | South China Sea | Mollusca | Bivalvia |
| 243 | Hubei coleoptera virus 2 strain QCM76287 hypothetical protein gene complete cds | KX883925.1 | Terrestrial | Arthropoda | Insecta |
| 244 | Gastropoda Picorna-like virus D23 | GWHAPBE01000000 | East China Sea | Mollusca | Gastropoda |
| 245 | Hubei earwig virus 3 strain GCM10213 hypothetical protein 1 and hypothetical protein 2 genes complete cds | KX883892.1 | Terrestrial | Arthropoda | Insecta |
| 246 | Hubei earwig virus 3 strain ZCM20657 hypothetical protein 1 and hypothetical protein 2 genes complete cds | KX884397.1 | Terrestrial | Arthropoda | Insecta |
| 247 | Hubei earwig virus 3 strain QCM145712 hypothetical protein 1 and hypothetical protein 2 genes complete cds | KX883922.1 | Terrestrial | Arthropoda | Insecta |
| 248 | Hubei earwig virus 3 strain QTM27170 hypothetical protein 1 and hypothetical protein 2 genes complete cds | KX883942.1 | Terrestrial | Arthropoda | Insecta |
| 249 | Hubei earwig virus 3 strain WGML147931 hypothetical protein 1 and hypothetical protein 2 genes complete cds | KX884259.1 | Terrestrial | Arthropoda | Myriapoda |
| 250 | Hubei earwig virus 3 strain WHYCM1140 hypothetical protein 1 and hypothetical protein 2 genes complete cds | KX884296.1 | Terrestrial | Arthropoda | Insecta |
| 251 | Hubei earwig virus 3 strain WHXS4867 hypothetical protein 1 and hypothetical protein 2 genes complete cds | KX884294.1 | Terrestrial | Arthropoda | Insecta |
| 252 | Hubei earwig virus 3 strain arthropodmix12795 hypothetical protein 1 and hypothetical protein 2 genes complete cd | KX883286.1 | Terrestrial | Arthropoda | Mix |
| 253 | Wuhan heteroptera virus 3 strain WHYCM468 hypothetical protein 6 gene complete cds | KX884728.1 | Terrestrial | Arthropoda | Insecta |
| 254 | Wuhan heteroptera virus 3 strain arthropodmix44000 hypothetical protein 7 gene complete cds | KX884595.1 | Terrestrial | Arthropoda | Mix |
| 255 | Bivalvia Picorna-like virus D13 | GWHAOXV0100000C | East China Sea | Mollusca | Bivalvia |

| | | | | | |
|---|---|---|---|---|---|
| 256 | Beihai hermit crab virus 1 strain BHTSS17051 hypothetical protein 1 and hypothetical protein 2 genes complete cds | KX883417.1 | South China Sea | Arthropoda | Crustacea |
| 257 | Beihai hermit crab virus 1 strain SCJXSX39363 hypothetical protein 1 and hypothetical protein 2 genes complete cc | KX883966.1 | South China Sea | Arthropoda | Crustacea |
| 258 | Beihai hermit crab virus 1 strain BHJJX25619 hypothetical protein 1 and hypothetical protein 2 genes complete cds | KX883366.1 | South China Sea | Arthropoda | Crustacea |
| 259 | Beihai hermit crab virus 1 strain BHWZXX15637 hypothetical protein 1 and hypothetical protein 2 genes complete | KX883430.1 | South China Sea | Arthropoda | Crustacea |
| 260 | Beihai hermit crab virus 1 strain BHBJDX18472 hypothetical protein 1 and hypothetical protein 2 genes complete c | KX883327.1 | South China Sea | Arthropoda | Crustacea |
| 261 | Gastropoda Picorna-like virus N4 | GWHAPFN01000000 | South China Sea | Mollusca | Gastropoda |
| 262 | Gastropoda Picorna-like virus D14 | GWHAPAX01000000 | East China Sea | Mollusca | Gastropoda |
| 263 | Hubei picorna-like virus 73 strain WGML140238 hypothetical protein 1 hypothetical protein 2 and hypothetical pro | KX883735.1 | Terrestrial | Arthropoda | Myriapoda |
| 264 | Gastropoda Picorna-like virus D29 | GWHAPKF01000000 | East China Sea | Mollusca | Gastropoda |
| 265 | Wenzhou picorna-like virus 45 strain WZSBei69068 hypothetical protein gene complete cds | KX884357.1 | East China Sea | Mollusca | Bivalvia |
| 266 | Beihai picorna-like virus 51 strain SCJXSX38939 hypothetical protein 1 and hypothetical protein 2 genes complete | KX883958.1 | South China Sea | Arthropoda | Crustacea |
| 267 | Bivalvia Picorna-like virus N7 | GWHAPFB01000000 | South China Sea | Mollusca | Bivalvia |
| 268 | Gastropoda Unclassified virus D1 | GWHAPJU01000000 | East China Sea | Mollusca | Gastropoda |
| 269 | Crustacea Picorna-like virus N31 | GWHAPDR01000000 | South China Sea | Arthropoda | Crustacea |
| 270 | Beihai picorna-like virus 112 strain BHXG26692 hypothetical protein 1 and hypothetical protein 2 genes complete ( | KX884473.1 | South China Sea | Arthropoda | Crustacea |
| 271 | Wuhan insect virus 10 strain WHCCII13330 hypothetical protein gene complete cds | KX884277.1 | Terrestrial | Arthropoda | Insecta |
| 272 | Bivalvia Durna-like virus D10 | GWHAOZM0100000( | East China Sea | Mollusca | Bivalvia |
| 273 | Wenzhou picorna-like virus 28 strain WZRBX42853 hypothetical protein 1 and hypothetical protein 2 genes compl | KX884345.1 | East China Sea | Arthropoda | Crustacea |
| 274 | Bivalvia Durna-like virus N1 | GWHAPIS01000000 | South China Sea | Mollusca | Bivalvia |
| 275 | Bivalvia Durna-like virus N5 | GWHAPDZ01000000 | South China Sea | Mollusca | Bivalvia |
| 276 | Beihai picorna-like virus 28 strain BHZY60695 hypothetical protein 1 and hypothetical protein 2 genes complete cc | KX884511.1 | South China Sea | Mollusca | Cephalopoda |
| 277 | Bivalvia Picorna-like virus H11 | GWHAPJK01000000 | Yellow China Sea | Mollusca | Bivalvia |
| 278 | Beihai octopus viurs 2 strain BHZY60909 hypothetical protein gene complete cds | KX884520.1 | South China Sea | Mollusca | Cephalopoda |
| 279 | Sanxia water strider virus 21 strain SXSSP2576 hypothetical protein gene complete cds | KX883706.1 | Freshwater | Arthropoda | Insecta |
| 280 | Hubei diptera virus 14 strain QTM46268 segment 2 hypothetical protein 1 and hypothetical protein 2 genes complet | KX882852.1 | Terrestrial | Arthropoda | Insecta |
| 281 | Hubei diptera virus 14 strain SCM51203 segment 2 hypothetical protein 1 gene complete cds | KX882857.1 | Terrestrial | Arthropoda | Insecta |
| 282 | Bivalvia Picorna-like virus D8 | GWHAOXX0100000( | East China Sea | Mollusca | Bivalvia |
| 283 | Hubei picorna-like virus 12 strain HC20950 hypothetical protein gene complete cds | KX883896.1 | Terrestrial | Nematoda | Rhabditida |
| 284 | Hubei picorna-like virus 12 strain WHZHC35617 hypothetical protein gene complete cds | KX884305.1 | Terrestrial | Nematoda | Chromadorea |
| 285 | Bivalvia Picorna-like virus D37 | GWHAOYP01000000 | East China Sea | Mollusca | Bivalvia |
| 286 | Jingmen picorna-like virus strain JMYJCC71227 hypothetical protein gene complete cds | KX883913.1 | Terrestrial | Nematoda | Chromadorea |
| 287 | Shahe picorna-like virus 7 strain SHWC13621 hypothetical protein gene complete cds | KX883665.1 | Freshwater | Arthropoda | Mix |
| 288 | Shahe picorna-like virus 7 strain SHWC0209c12710 hypothetical protein gene complete cds | KX883652.1 | Freshwater | Arthropoda | Crustacea |
| 289 | Hubei insect virus 2 strain arthropodmix13736 hypothetical protein 8 gene complete cds | KX884587.1 | Terrestrial | Arthropoda | Mix |
| 290 | Hubei partiti-like virus 16 strain WGML112875 RdRp gene complete cds | KX884186.1 | Terrestrial | Arthropoda | Myriapoda |
| 291 | Hubei partiti-like virus 16 strain WHSFII9387 RdRp gene complete cds | KX884201.1 | Terrestrial | Arthropoda | Crustacea |
| 292 | Bivalvia Picorna-like virus D39 | GWHAPER01000000 | East China Sea | Mollusca | Bivalvia |

| | | | | | |
|---|---|---|---|---|---|
| 293 | Hubei myriapoda virus 8 strain WGML66308 putative glycoprotein hypothetical protein 1 hypothetical protein 2 an | KX884444.1 | Terrestrial | Arthropoda | Myriapoda |
| 294 | Wuhan Louse Fly Virus 1 strain BFJSC-10 nucleopasid protein (N) and nonstructural protein (NSs) genes complete | KM817755.1 | Terrestrial | Arthropoda | Insecta |
| 295 | Wenling hepe-like virus 1 strain WLJQ103981 replicase hypothetical protein putative structural protein and hypoth | KX883830.1 | East China Sea | Arthropoda | Crustacea |
| 296 | Bivalvia Picorna-like virus N14 | GWHAPCE01000000 | South China Sea | Mollusca | Bivalvia |
| 297 | Wuhan House Fly Virus 1 strain SCM39215 putative nucleoprotein hypothetical protein putative X protein putative | KX884432.1 | Terrestrial | Arthropoda | Insecta |
| 298 | Hubei dimarhabdovirus virus 1 strain SCM51525 putative nucleoprotein hypothetical protein putative matrix protei | KX884431.1 | Terrestrial | Arthropoda | Insecta |
| 299 | Bivalvia Picorna-like virus N48 | GWHAPEH01000000 | South China Sea | Mollusca | Bivalvia |
| 300 | Wuhan House Fly Virus 1 strain SYY2-4 nucleocapsid (N) putative phosphoprotein (ORF2) putative X protein (OR | KM817648.1 | Terrestrial | Arthropoda | Insecta |
| 301 | Beihai orthomyxo-like virus 1 strain BHTSS17872 segment 3 polymerase PA gene complete cds | KX883846.1 | South China Sea | Arthropoda | Crustacea |
| 302 | Hubei myriapoda virus 7 strain WGML146453 hypothetical protein 1 hypothetical protein 2 hypothetical protein 3 l | KX884443.1 | Terrestrial | Arthropoda | Myriapoda |
| 303 | Jingshan Fly Virus 1 strain SYY1-7 segment PA polymerase PA gene complete cds | KX883856.1 | Terrestrial | Arthropoda | Insecta |
| 304 | Jingshan Fly Virus 1 strain SCM40889 segment PB2 polymerase PB2 gene complete cds | KX883876.1 | Terrestrial | Arthropoda | Insecta |
| 305 | Hubei odonate virus 11 strain QTM161788 RNA-dependent RNA polymerase putative glycoprotein and hypothetic | KX884419.1 | Terrestrial | Arthropoda | Insecta |
| 306 | Bivalvia Picorna-like virus N23 | GWHAPGK01000000 | South China Sea | Mollusca | Bivalvia |
| 307 | Shayang Fly Virus 3 strain SYY1-1 ORF1 (ORF1) ORF2 (ORF2) ORF3 (ORF3) glycoprotein (G) and RNA-depen | KM817636.1 | Terrestrial | Arthropoda | Insecta |
| 308 | Bivalvia Picorna-like virus N43 | GWHAPDX01000000 | South China Sea | Mollusca | Bivalvia |
| 309 | Wuhan Insect virus 6 strain SXCC01-1 nucleocapsid (N) phosphoprotein (P) 4b protein (4b) matrix protein (M) gly | KM817652.1 | Terrestrial | Arthropoda | Insecta |
| 310 | Wuchang Cockraoch Virus 2 strain WCZL-3 PB1 (PB1) gene partial cds | KM817628.1 | Terrestrial | Arthropoda | Insecta |
| 311 | Bivalvia Picorna-like virus N26 | GWHAPGM0100000 | South China Sea | Mollusca | Bivalvia |
| 312 | Wenzhou Crab Virus 3 strain RBX9 glycoprotein (G1) putative nucleoprotein (N) glycoprotein (G2) and polymeras | KM817603.1 | East China Sea | Arthropoda | Crustacea |
| 313 | Jingshan Fly Virus 1 strain SCM40889 segment PA PA gene complete cds | KX883877.1 | Terrestrial | Arthropoda | Insecta |
| 314 | Jingshan Fly Virus 1 strain SYY1-7 segment PB2 polymerase PB2 gene complete cds | KX883855.1 | Terrestrial | Arthropoda | Insecta |
| 315 | Wuhan horsefly Virus strain JJ2-1 nonstructural protein (NSs) gene complete cds | KM817750.1 | Terrestrial | Arthropoda | Insecta |
| 316 | Wuhan horsefly Virus strain JJ2-1 nucleopasid protein (N) gene complete cds | KM817751.1 | Terrestrial | Arthropoda | Insecta |
| 317 | Hubei odonate virus 10 strain QTM133243 hypothetical protein 1 hypothetical protein 2 hypothetical protein 3 puta | KX884418.1 | Terrestrial | Arthropoda | Insecta |
| 318 | Wuhan House Fly Virus 2 strain SYY4-5 ORF1 (ORF1) ORF2 (ORF2) ORF3 (ORF3) glycoprotein (G) and RNA-c | KM817649.1 | Terrestrial | Arthropoda | Chelicerata |
| 319 | Hubei orthomyxo-like virus 2 strain QTM27225 segment 5 hemagglutinin gene complete cds | KX883872.1 | Terrestrial | Arthropoda | Insecta |
| 320 | Hubei blood fluke virus 1 strain HBXXC56361 putative nucleoprotein gene complete cds | KX884764.1 | Freshwater | Platyhelminthes | Trematoda |
| 321 | Hubei blood fluke virus 1 strain JSC85519 putative nucleoprotein gene complete cds | KX884771.1 | Freshwater | Platyhelminthes | Trematoda |
| 322 | Hubei blood fluke virus 1 strain RBXXC303595 putative nucleoprotein gene complete cds | KX884793.1 | Freshwater | Platyhelminthes | Trematoda |
| 323 | Xincheng Mosquito Virus strain XC1-6 ORF1 (ORF1) ORF2 (ORF2) glycoprotein (G) and RNA-dependent RNA p | KM817661.1 | Terrestrial | Arthropoda | Insecta |
| 324 | Bivalvia Picorna-like virus N28 | GWHAPCI01000000 | South China Sea | Mollusca | Bivalvia |
| 325 | Wenzhou gastropodes virus 2 strain WZSLuoII97238 hypothetical protein gene complete cds | KX884380.1 | East China Sea | Mollusca | Gastropoda |
| 326 | Wenzhou gastropodes virus 2 strain WZSLuoI86086 hypothetical protein gene complete cds | KX884374.1 | East China Sea | Mollusca | Gastropoda |
| 327 | Hubei diptera virus 10 strain SCM43656 putative nucleoprotein hypothetical protein putative X protein putative ma | KX884433.1 | Terrestrial | Arthropoda | Insecta |
| 328 | Hubei coleoptera virus 3 strain QCM109726 putative glycoprotein hypothetical protein and RNA-dependent RNA p | KX884416.1 | Terrestrial | Arthropoda | Insecta |
| 329 | Beihai orthomyxo-like virus 1 strain BHTSS17872 segment 2 polymerase PB1 gene complete cds | KX883844.1 | South China Sea | Arthropoda | Crustacea |

| | | | | | |
|---|---|---|---|---|---|
| ***330*** | Sanxia Water Strider Virus 4 strain SXSSP12 ORF1 (ORF1) ORF2 (ORF2) glycoprotein (G) and RNA-dependent I | KM817633.1 | Freshwater | Arthropoda | Insecta |
| ***331*** | Beihai orthomyxo-like virus 1 strain BHTSS17872 segment 1 polymerase PB2 gene complete cds | KX883845.1 | South China Sea | Arthropoda | Crustacea |
| ***332*** | Crustacea Picorna-like virus N10 | GWHAPKX01000000 | South China Sea | Arthropoda | Crustacea |
| ***333*** | Shayang Fly Virus 2 strain SYY1-8 nucleocapsid (N) ORF2 (ORF2) X protein (X) matrix protein (M) glycoprotein | KM817635.1 | Terrestrial | Arthropoda | Insecta |
| ***334*** | Wuhan insect virus 16 strain WHZM9931 putative nucleoprotein gene complete cds | KX884855.1 | Terrestrial | Arthropoda | Insecta |
| ***335*** | Wuhan insect virus 16 strain arthropodmix13442 nucleoprotein gene complete cds | KX884734.1 | Terrestrial | Arthropoda | Mix |
| ***336*** | Jingshan Fly Virus 1 strain SYY1-7 PB1 (PB1) gene complete cds | KM817615.1 | Terrestrial | Arthropoda | Insecta |
| ***337*** | Jingshan Fly Virus 1 strain SCM40889 segment PB1 polymerase PB1 gene complete cds | KX883875.1 | Terrestrial | Arthropoda | Insecta |
| ***338*** | Hubei diptera virus 9 strain SCM172232 putative nucleoprotein hypothetical protein 2 putative X protein putative m | KX884429.1 | Terrestrial | Arthropoda | Insecta |
| ***339*** | Jiujie Fly Virus strain JJ3-6 segment PB2 polymerase PB2 gene complete cds | KX883863.1 | Terrestrial | Arthropoda | Insecta |
| ***340*** | Wuhan spider virus 4 strain spider133854 hypothetical protein 1 hypothetical protein 2 hypothetical protein 3 and h | KX883684.1 | Terrestrial | Arthropoda | Chelicerata |
| ***341*** | Hubei orthomyxo-like virus 2 strain QTM27225 segment 4 nucleocapsid protein gene complete cds | KX883871.1 | Terrestrial | Arthropoda | Insecta |
| ***342*** | Hubei picorna-like virus 75 strain WGML145097 hypothetical protein 1 hypothetical protein 2 hypothetical protein | KX884251.1 | Terrestrial | Arthropoda | Myriapoda |
| ***343*** | Wuhan heteroptera virus 3 strain WHYCM468 hypothetical protein 9 gene complete cds | KX884730.1 | Terrestrial | Arthropoda | Insecta |
| ***344*** | Wuhan heteroptera virus 3 strain arthropodmix44000 hypothetical protein 9 gene complete cds | KX884597.1 | Terrestrial | Arthropoda | Mix |
| ***345*** | Xingshan nematode virus 4 strain XSNXC32924 putative nucleoprotein hypothetical protein 1 hypothetical protein | KX884459.1 | Terrestrial | Nematoda | Chromadorea |
| ***346*** | Shuangao Insect Virus 2 strain QSA03 nucleopasid protein (S1) gene complete cds | KM817740.1 | Terrestrial | Arthropoda | Insecta |
| ***347*** | Hubei orthomyxo-like virus 2 strain QTM27225 segment 2 polymerase PB1 gene complete cds | KX883868.1 | Terrestrial | Arthropoda | Insecta |
| ***348*** | Wuhan Louse Fly Virus 10 strain BFJSC-8 nucleocapsid (N) putative phosphoprotein (ORF2) matrix protein (M) gl | KM817657.1 | Terrestrial | Arthropoda | Insecta |
| ***349*** | Wuhan horsefly Virus strain JJ2-1 glycoprotein precursor (G) gene complete cds | KM817723.1 | Terrestrial | Arthropoda | Insecta |
| ***350*** | Shayang Spider Virus 1 strain SYZZ-4 RNA-dependent RNA polymerase (L) gene complete cds | KM817676.1 | Terrestrial | Arthropoda | Chelicerata |
| ***351*** | Hubei picorna-like virus 69 strain WGML147773 hypothetical protein 1 hypothetical protein 2 and hypothetical pro | KX884257.1 | Terrestrial | Arthropoda | Myriapoda |
| ***352*** | Hubei orthomyxo-like virus 2 strain QTM27225 segment 3 polymerase PA gene complete cds | KX883870.1 | Terrestrial | Arthropoda | Insecta |
| ***353*** | Wuhan pillworm virus 2 strain WHSFII14871 putative nucleoprotein hypothetical protein putative matrix protein pu | KX884445.1 | Terrestrial | Arthropoda | Crustacea |
| ***354*** | Wuhan pillworm virus 2 strain WGML144272 putative nucleoprotein hypothetical protein putative matrix protein p | KX884442.1 | Terrestrial | Arthropoda | Myriapoda |
| ***355*** | Hubei earwig virus 1 strain GCM19162 segment 1 polymerase PB2 gene complete cds | KX883859.1 | Terrestrial | Arthropoda | Insecta |
| ***356*** | Hubei blood fluke virus 2 strain RBXXC137009 putative nucleoprotein gene complete cds | KX884791.1 | Freshwater | Platyhelminthes | Trematoda |
| ***357*** | Hubei blood fluke virus 2 strain HBXXC60124 putative nucleoprotein gene complete cds | KX884762.1 | Freshwater | Platyhelminthes | Trematoda |
| ***358*** | Hubei blood fluke virus 2 strain JSC69896 putative nucleoprotein gene complete cds | KX884769.1 | Freshwater | Platyhelminthes | Trematoda |
| ***359*** | Hubei rhabdo-like virus 2 strain WHSWHC60518 hypothetical protein 1 hypothetical protein 2 hypothetical protein | KX884447.1 | Terrestrial | Nematoda | Chromadorea |
| ***360*** | Hubei earwig virus 1 strain GCM19162 segment 3 polymerase PA gene complete cds | KX883860.1 | Terrestrial | Arthropoda | Insecta |
| ***361*** | Wuhan heteroptera virus 3 strain WHYCM468 hypothetical protein 3 and hypothetical protein 4 genes complete cd | KX884726.1 | Terrestrial | Arthropoda | Insecta |
| ***362*** | Wuhan heteroptera virus 3 strain arthropodmix44000 hypothetical protein 3 and hypothetical protein 4 genes compl | KX884593.1 | Terrestrial | Arthropoda | Mix |
| ***363*** | Wuhan nido-like virus 1 strain SCM49454 1ab hypothetical protein and putative structural polyprotein genes compl | KX883630.1 | Terrestrial | Arthropoda | Insecta |
| ***364*** | Wuhan insect virus 19 strain WHCCII13334 1ab hypothetical protein and putative structural polyprotein genes com | KX883634.1 | Terrestrial | Arthropoda | Insecta |
| ***365*** | Wenzhou toti-like virus 2 strain beimix73231 hypothetical protein gene partial cds | KX882941.1 | East China Sea | Mollusca | Bivalvia |
| ***366*** | Wuhan heteroptera virus 3 strain arthropodmix44000 Inner core protein gene complete cds | KX884591.1 | Terrestrial | Arthropoda | Mix |

| | | | | | |
|---|---|---|---|---|---|
| 367 | Wuhan heteroptera virus 3 strain WHYCM468 Inner core protein gene complete cds | KX884724.1 | Terrestrial | Arthropoda | Insecta |
| 368 | Bivalvia Picorna-like virus D52 | GWHAOZR01000000 | East China Sea | Mollusca | Bivalvia |
| 369 | Beihai picorna-like virus 42 strain SCJXSX39263 hypothetical protein 1 and hypothetical protein 2 genes complete | KX883963.1 | South China Sea | Arthropoda | Crustacea |
| 370 | Hubei picorna-like virus 9 strain QTM26755 hypothetical protein gene complete cds | KX883937.1 | Terrestrial | Arthropoda | Insecta |
| 371 | Hubei permutotetra-like virus 11 strain mosHB234479 RdRp and putative capsid protein genes complete cds | KX883440.1 | Terrestrial | Arthropoda | Insecta |
| 372 | Hubei permutotetra-like virus 11 strain 3mos6479 hypothetical protein gene partial cds | KX883435.1 | Terrestrial | Arthropoda | Insecta |
| 373 | Wuhan Louse Fly Virus 11 strain BFJSC-9 RNA-dependent RNA polymerase (L) gene complete cds | KM817658.1 | Terrestrial | Arthropoda | Insecta |
| 374 | Beihai narna-like virus 26 strain BHHQ66335 RNA-dependent RNA polymerase gene complete cds | KX883467.1 | South China Sea | Tunicata | Ascidiacea |
| 375 | Hubei orthoptera virus 3 strain ZCM18544 hypothetical protein 1 and hypothetical protein 2 genes complete cds | KX884395.1 | Terrestrial | Arthropoda | Insecta |
| 376 | Hubei odonate virus 13 strain QTM74844 RdRp gene complete cds | KX884141.1 | Terrestrial | Arthropoda | Insecta |
| 377 | Hubei odonate virus 13 strain WGML31721 RdRp gene complete cds | KX884188.1 | Terrestrial | Arthropoda | Myriapoda |
| 378 | Wenzhou picorna-like virus 5 strain WZSLuoI84329 hypothetical protein 1 and hypothetical protein 2 genes comple | KX884369.1 | East China Sea | Mollusca | Gastropoda |
| 379 | Bivalvia Durna-like virus D8 | GWHAOZW0100000( | East China Sea | Mollusca | Bivalvia |
| 380 | Hubei sobemo-like virus 37 strain QTM25849 hypothetical protein 1 hypothetical protein 2 and hypothetical protei | KX882842.1 | Terrestrial | Arthropoda | Insecta |
| 381 | Wenzhou picorna-like virus 14 strain WZSLuoII97619 hypothetical protein 1 and hypothetical protein 2 genes com | KX884382.1 | East China Sea | Mollusca | Gastropoda |
| 382 | Bivalvia Picorna-like virus D3 | GWHAPEM01000000 | East China Sea | Mollusca | Bivalvia |
| 383 | Beihai mollusks virus 1 strain WZSLuoI86140 hypothetical protein 1 and hypothetical protein 2 genes complete cds | KX884377.1 | East China Sea | Mollusca | Gastropoda |
| 384 | Beihai mollusks virus 1 strain BHXG26682 hypothetical protein 1 and hypothetical protein 2 genes complete cds | KX884472.1 | South China Sea | Arthropoda | Crustacea |
| 385 | Beihai mollusks virus 1 strain BHNXC41338 hypothetical protein 1 gene complete cds and hypothetical protein 2 g | KX883395.1 | South China Sea | Sipuncula | Sipuncula |
| 386 | Beihai mollusks virus 1 strain BHHK131425 hypothetical protein 1 and hypothetical protein 2 genes complete cds | KX883351.1 | South China Sea | Cnidaria | Anthozoa |
| 387 | Beihai mollusks virus 1 strain YYSZX17807 hypothetical protein 1 and hypothetical protein 2 genes complete cds | KX884391.1 | South China Sea | Arthropoda | Crustacea |
| 388 | Beihai mollusks virus 1 strain BHZY60617 hypothetical protein 1 and hypothetical protein 2 genes complete cds | KX884509.1 | South China Sea | Mollusca | Cephalopoda |
| 389 | Beihai mollusks virus 1 strain BWBFG40783 hypothetical protein 1 and hypothetical protein 2 genes complete cds | KX884528.1 | South China Sea | Mollusca | Bivalvia |
| 390 | Wenzhou picorna-like virus 9 strain WZSBei69430 hypothetical protein 1 and hypothetical protein 2 genes complet | KX884359.1 | East China Sea | Mollusca | Bivalvia |
| 391 | Wenzhou picorna-like virus 20 strain WZSBei68985 hypothetical protein 1 and hypothetical protein 2 genes comple | KX884356.1 | East China Sea | Mollusca | Bivalvia |
| 392 | Beihai picorna-like virus 17 strain BHZY55665 hypothetical protein 1 and hypothetical protein 2 genes complete cd | KX884505.1 | South China Sea | Mollusca | Cephalopoda |
| 393 | Crustacea Picorna-like virus D1 | GWHAPBN01000000 | East China Sea | Arthropoda | Crustacea |
| 394 | Beihai anemone virus 1 strain BHHK129576 hypothetical protein genes complete cds | KX883744.1 | South China Sea | Cnidaria | Anthozoa |
| 395 | Hubei picorna-like virus 56 strain SCM49537 hypothetical protein 1 and hypothetical protein 2 genes complete cds | KX883969.1 | Terrestrial | Arthropoda | Insecta |
| 396 | Wuhan heteroptera virus 3 strain arthropodmix44000 hypothetical protein 6 gene complete cds | KX884594.1 | Terrestrial | Arthropoda | Mix |
| 397 | Wuhan heteroptera virus 3 strain WHYCM468 hypothetical protein 5 gene complete cds | KX884727.1 | Terrestrial | Arthropoda | Insecta |
| 398 | Wuhan heteroptera virus 1 strain WHYCM964 RdRp and hypothetical protein genes complete cds | KX883821.1 | Terrestrial | Arthropoda | Insecta |
| 399 | Wuhan heteroptera virus 1 strain arthropodmix8462 RdRp and hypothetical protein genes complete cds | KX883741.1 | Terrestrial | Arthropoda | Mix |
| 400 | Hubei reo-like virus 10 strain QTM136127 Minor structural protein gene complete cds | KX884670.1 | Terrestrial | Arthropoda | Insecta |
| 401 | Hubei picorna-like virus 27 strain QCM155345 hypothetical protein gene complete cds | KX883924.1 | Terrestrial | Arthropoda | Insecta |
| 402 | Hubei diptera virus 11 strain SCM51501 hypothetical protein 1 hypothetical protein 2 putative glycoprotein and RN | KX884430.1 | Terrestrial | Arthropoda | Insecta |
| 403 | Shuangao Fly Virus 2 strain QSA05 RNA-dependent RNA polymerase (L) gene partial cds | KM817638.1 | Terrestrial | Arthropoda | Insecta |

| | | | | | |
|---|---|---|---|---|---|
| 404 | Wuhan spider virus 5 strain spider133684 hypothetical protein 1 hypothetical protein 2 hypothetical protein 3 and h | KX883681.1 | Terrestrial | Arthropoda | Chelicerata |
| 405 | Hubei diptera virus 20 strain SCM51404 RNA-dependent RNA polymerase gene complete cds | KX884693.1 | Terrestrial | Arthropoda | Insecta |
| 406 | Hubei picorna-like virus 70 strain WGML134035 hypothetical protein 1 hypothetical protein 2 hypothetical protein | KX883733.1 | Terrestrial | Arthropoda | Myriapoda |
| 407 | Wuhan heteroptera virus 3 strain WHYCM468 hypothetical protein 2 gene complete cds | KX884725.1 | Terrestrial | Arthropoda | Insecta |
| 408 | Wuhan heteroptera virus 3 strain arthropodmix44000 hypothetical protein 2 gene complete cds | KX884592.1 | Terrestrial | Arthropoda | Mix |
| 409 | Wuhan heteroptera virus 3 strain arthropodmix44000 Major core protein gene complete cds | KX884589.1 | Terrestrial | Arthropoda | Mix |
| 410 | Wuhan heteroptera virus 3 strain WHYCM468 Major core protein gene complete cds | KX884722.1 | Terrestrial | Arthropoda | Insecta |
| 411 | Wuhan arthropod virus 1 strain WHCC81862 hypothetical protein genes complete cds | KX883810.1 | Terrestrial | Arthropoda | Insecta |
| 412 | Wuhan heteroptera virus 3 strain arthropodmix44000 hypothetical protein 1 gene complete cds | KX884590.1 | Terrestrial | Arthropoda | Mix |
| 413 | Wuhan heteroptera virus 3 strain WHYCM468 hypothetical protein 1 gene complete cds | KX884723.1 | Terrestrial | Arthropoda | Insecta |
| 414 | Wuhan coneheads virus 2 strain ZCM13613 hypothetical protein 1 hypothetical protein 2 and hypothetical protein 3 | KX884394.1 | Terrestrial | Arthropoda | Insecta |
| 415 | Wuhan coneheads virus 2 strain WHLC4766 hypothetical protein 1 hypothetical protein 2 and hypothetical protein 3 | KX884280.1 | Terrestrial | Arthropoda | Insecta |
| 416 | Wuhan spider virus 2 strain spider133995 hypothetical protein gene complete cds | KX883689.1 | Terrestrial | Arthropoda | Chelicerata |
| 417 | Hubei diptera virus 21 strain SCM95013 hypothetical protein 2 gene complete cds | KX884698.1 | Terrestrial | Arthropoda | Insecta |
| 418 | Wuhan fly virus 5 strain fly34516 segment Seg 1 RdRp gene complete cds | KX884091.1 | Terrestrial | Arthropoda | Insecta |
| 419 | Xinzhou nematode virus 1 strain XZSJSC65770 hypothetical protein genes complete cds | KX883838.1 | Terrestrial | Nematoda | Mix |
| 420 | Bivalvia Durna-like virus D13 | GWHAPAJ01000000 | East China Sea | Mollusca | Bivalvia |
| 421 | Gastropoda Durna-like virus D2 | GWHAPKI01000000 | East China Sea | Mollusca | Gastropoda |
| 422 | Sanxia picorna-like virus 9 strain SXXX36560 hypothetical protein 1 and hypothetical protein 2 genes complete cds | KX883717.1 | Freshwater | Arthropoda | Crustacea |
| 423 | Hubei picorna-like virus 72 strain WHSFII16052 hypothetical protein 1 hypothetical protein 2 and hypothetical prot | KX884289.1 | Terrestrial | Arthropoda | Crustacea |
| 424 | Gastropoda Picorna-like virus N1 | GWHAPFD01000000 | South China Sea | Mollusca | Gastropoda |
| 425 | Shahe picorna-like virus 6 strain SHWC13532 hypothetical protein 1 and hypothetical protein 2 genes complete cds | KX883655.1 | Freshwater | Arthropoda | Mix |
| 426 | Shahe picorna-like virus 6 strain SHWC0209c11084 hypothetical protein 1 and hypothetical protein 2 genes comple | KX883653.1 | Freshwater | Arthropoda | Crustacea |
| 427 | Beihai mollusks virus 2 strain BHZC37403 hypothetical protein 1 and hypothetical protein 2 genes complete cds | KX884502.1 | South China Sea | Mollusca | Bivalvia |
| 428 | Beihai mollusks virus 2 strain WZSLuoI86113 hypothetical protein 1 and hypothetical protein 2 genes complete cds | KX884375.1 | East China Sea | Mollusca | Gastropoda |
| 429 | Beihai mollusks virus 2 strain BHZY60889 hypothetical protein 1 and hypothetical protein 2 genes complete cds | KX884518.1 | South China Sea | Mollusca | Cephalopoda |
| 430 | Beihai picorna-like virus 34 strain HOU158088 hypothetical protein 1 gene complete cds and hypothetical protein 2 | KX883902.1 | South China Sea | Arthropoda | Chelicerata |
| 431 | Hubei sobemo-like virus 39 strain mosHB229649 hypothetical protein 2 and hypothetical protein 1 genes complete | KX882828.1 | Terrestrial | Arthropoda | Insecta |
| 432 | Hubei sobemo-like virus 12 strain spider127544 hypothetical protein 1 and hypothetical protein 2 genes complete c | KX882863.1 | Terrestrial | Arthropoda | Chelicerata |
| 433 | Hubei partiti-like virus 40 strain WHCCII13098 RdRp gene complete cds | KX884195.1 | Terrestrial | Arthropoda | Insecta |
| 434 | Bivalvia Picorna-like virus N4 | GWHAPEX01000000 | South China Sea | Mollusca | Bivalvia |
| 435 | Gastropoda Durna-like virus N1 | GWHAPFT01000000 | South China Sea | Mollusca | Gastropoda |
| 436 | Beihai sea slater virus 1 strain BHHZL10310 hypothetical protein gene complete cds | KX883359.1 | South China Sea | Arthropoda | Crustacea |
| 437 | Hubei partiti-like virus 57 strain QTM22037 RdRp gene complete cds | KX884116.1 | Terrestrial | Arthropoda | Insecta |
| 438 | Beihai picobirna-like virus 10 strain BHZC36972 RdRp gene complete cds | KX884079.1 | South China Sea | Mollusca | Bivalvia |
| 439 | Gastropoda Picorna-like virus D25 | GWHAPBF01000000 | East China Sea | Mollusca | Gastropoda |
| 440 | Beihai mantis shrimp virus 4 strain BHXG23944 hypothetical protein 1 and hypothetical protein 2 genes complete c | KX883434.1 | South China Sea | Arthropoda | Crustacea |

| | | | | | |
|---|---|---|---|---|---|
| 441 | Bivalvia Picorna-like virus N29 | GWHAPCG01000000 | South China Sea | Mollusca | Bivalvia |
| 442 | Wenzhou picorna-like virus 38 strain WZSLuoI83635 hypothetical protein gene partial cds | KX884368.1 | East China Sea | Mollusca | Gastropoda |
| 443 | Beihai picorna-like virus 48 strain BHZC37443 hypothetical protein gene complete cds | KX884498.1 | South China Sea | Mollusca | Bivalvia |
| 444 | Banna virus strain QTM104536 Banna VP3 gene complete cds | KX884640.1 | Terrestrial | Arthropoda | Insecta |
| 445 | Wenzhou gastropodes virus 1 strain WZSLuoI86017 hypothetical protein 1 and hypothetical protein 2 genes comple | KX884372.1 | East China Sea | Mollusca | Gastropoda |
| 446 | Wuhan fly virus 6 strain fly116610 RdRp gene complete cds | KX884090.1 | Terrestrial | Arthropoda | Insecta |
| 447 | Wenzhou picorna-like virus 23 strain WZFSL85852 hypothetical protein gene complete cds | KX884324.1 | Freshwater | Mollusca | Gastropoda |
| 448 | Beihai picobirna-like virus 9 strain BHZC32520 RdRp gene partial cds | KX884077.1 | South China Sea | Mollusca | Bivalvia |
| 449 | Bivalvia Wolfram-like virus N1 | GWHAPLC01000000 | South China Sea | Mollusca | Bivalvia |
| 450 | Bivalvia Durna-like virus H6 | GWHAPHG01000000 | Yellow China Sea | Mollusca | Bivalvia |
| 451 | Beihai picobirna-like virus 2 strain BHJJX66797 RdRp gene complete cds | KX884061.1 | South China Sea | Arthropoda | Crustacea |
| 452 | Sanxia picorna-like virus 3 strain SXXX36208 hypothetical protein 1 and hypothetical protein 2 genes complete cds | KX883718.1 | Freshwater | Arthropoda | Crustacea |
| 453 | Beihai picorna-like virus 16 strain BHJJX25950 hypothetical protein gene complete cds | KX883374.1 | South China Sea | Arthropoda | Crustacea |
| 454 | Beihai picorna-like virus 16 strain BHHK123579 hypothetical protein gene complete cds | KX883343.1 | South China Sea | Cnidaria | Anthozoa |
| 455 | Beihai picorna-like virus 16 strain BHBei73558 hypothetical protein gene complete cds | KX883308.1 | South China Sea | Mollusca | Bivalvia |
| 456 | Wenzhou picorna-like virus 3 strain WZSBei69560 hypothetical protein gene complete cds | KX884363.1 | East China Sea | Mollusca | Bivalvia |
| 457 | Beihai picobirna-like virus 11 strain BWBFG40030 RdRp gene complete cds | KX884085.1 | South China Sea | Mollusca | Bivalvia |
| 458 | Jingmen picobirna-like virus 1 strain JMYJCC46554 RdRp gene partial cds | KX884101.1 | Terrestrial | Nematoda | Chromadorea |
| 459 | Sanxia Water Strider Virus 1 strain SXSSP08 nucleopasid protein (N) gene complete cds | KM817737.1 | Freshwater | Arthropoda | Insecta |
| 460 | Sanxia Water Strider Virus 1 strain SXXX37846 putative nucleoprotein gene complete cds | KX884834.1 | Freshwater | Arthropoda | Crustacea |
| 461 | Wuhan large pig roundworm virus 1 strain WHZHC73278 segment Seg 1 RdRp gene complete cds | KX884219.1 | Terrestrial | Nematoda | Chromadorea |
| 462 | Hubei diptera virus 17 strain SCM172187 segment Seg 2 hypothetical protein gene complete cds | KX884146.1 | Terrestrial | Arthropoda | Insecta |
| 463 | Sanxia water strider virus 10 strain SXSSP2571 segment 2 hypothetical protein 1 gene complete cds | KX882890.1 | Freshwater | Arthropoda | Insecta |
| 464 | Wuhan insect virus 24 strain CC64001 RdRp gene complete cds | KX884086.1 | Terrestrial | Arthropoda | Insecta |
| 465 | Bivalvia Hepeli-like virus D1 | GWHAOZY01000000 | East China Sea | Mollusca | Bivalvia |
| 466 | Bivalvia Picorna-like virus D62 | GWHAPAI01000000 | East China Sea | Mollusca | Bivalvia |
| 467 | Bivalvia Unclassified virus N3 | GWHAPLR01000000 | South China Sea | Mollusca | Bivalvia |
| 468 | Bivalvia Picorna-like virus D34 | GWHAOYX0100000C | East China Sea | Mollusca | Bivalvia |
| 469 | Hubei astro-like virus strain WGML136555 putative protease hypothetical protein RNA-dependent RNA polymeras | KX884572.1 | Terrestrial | Arthropoda | Myriapoda |
| 470 | Beihai picorna-like virus 31 strain BHZC37445 hypothetical protein 1 gene complete cds and hypothetical protein 2 | KX884499.1 | South China Sea | Mollusca | Bivalvia |
| 471 | Hubei partiti-like virus 44 strain QTM29853 RdRp gene complete cds | KX884135.1 | Terrestrial | Arthropoda | Insecta |
| 472 | Hubei partiti-like virus 29 strain QTM26744 RdRp gene complete cds | KX884131.1 | Terrestrial | Arthropoda | Insecta |
| 473 | Wenzhou picorna-like virus 43 strain WZSBei69573 hypothetical protein gene complete cds | KX884365.1 | East China Sea | Mollusca | Bivalvia |
| 474 | Sanxia water strider virus 18 strain SXSSP2538 RdRp gene complete cds | KX884178.1 | Freshwater | Arthropoda | Insecta |
| 475 | Sanxia water strider virus 18 strain SXXX36890 RdRp gene complete cds | KX884181.1 | Freshwater | Arthropoda | Crustacea |
| 476 | Hubei partiti-like virus 32 strain QTM46741 RdRp gene complete cds | KX884138.1 | Terrestrial | Arthropoda | Insecta |
| 477 | Wuhan Louse Fly Virus 6 strain BFJSC-2 segment 1 complete sequence | KM817606.1 | Terrestrial | Arthropoda | Insecta |

| | | | | | |
|---|---|---|---|---|---|
| 478 | Hubei picorna-like virus 21 strain WHBL73090 hypothetical protein 1 and hypothetical protein 2 genes complete cc | KX884263.1 | Freshwater | Mollusca | Bivalvia |
| 479 | Hubei partiti-like virus 21 strain QTM27155 RdRp gene partial cds | KX884134.1 | Terrestrial | Arthropoda | Insecta |
| 480 | Beihai partiti-like virus 9 strain BHXG11739 RdRp gene complete cds | KX884075.1 | South China Sea | Arthropoda | Crustacea |
| 481 | Wuhan Louse Fly Virus 7 strain BFJSC-3 segment 1 complete sequence | KM817609.1 | Terrestrial | Arthropoda | Insecta |
| 482 | Gastropoda Picorna-like virus D9 | GWHAPAP01000000 | East China Sea | Mollusca | Gastropoda |
| 483 | Hubei partiti-like virus 4 strain QCM950973 RdRp gene complete cds | KX884111.1 | Terrestrial | Arthropoda | Insecta |
| 484 | Changjiang sobemo-like virus 2 strain CJLX29674 replicase hypothetical protein 2 and putative capsid protein gene | KX882818.1 | Freshwater | Arthropoda | Crustacea |
| 485 | Wuhan insect virus 22 strain arthropodmix13806 segment Seg 2 hypothetical protein gene complete cds | KX884051.1 | Terrestrial | Arthropoda | Mix |
| 486 | Wuhan insect virus 22 strain WHZM10073 segment Seg 2 hypothetical protein gene complete cds | KX884222.1 | Terrestrial | Arthropoda | Insecta |
| 487 | Wuhan insect virus 22 strain spider98584 segment Seg 2 hypothetical protein gene complete cds | KX884172.1 | Terrestrial | Arthropoda | Chelicerata |
| 488 | Hubei picorna-like virus 57 strain SCM38770 hypothetical protein gene partial cds | KX883968.1 | Terrestrial | Arthropoda | Insecta |
| 489 | Hubei orthoptera virus 4 strain ZCM21011 capsid protein gene complete cds | KX883284.1 | Terrestrial | Arthropoda | Insecta |
| 490 | Beihai picorna-like virus 95 strain BHBJDX18836 hypothetical protein gene partial cds | KX883337.1 | South China Sea | Arthropoda | Crustacea |
| 491 | Wenling sobemo-like virus 1 strain WLJQ103508 hypothetical protein 1 and hypothetical protein 2 genes complete | KX882930.1 | East China Sea | Arthropoda | Crustacea |
| 492 | Beihai picorna-like virus 98 strain HWRTX12368 hypothetical protein gene partial cds | KX883906.1 | South China Sea | Arthropoda | Crustacea |
| 493 | Wenzhou picorna-like virus 11 strain WZFSL83389 hypothetical protein gene complete cds | KX884333.1 | Freshwater | Mollusca | Gastropoda |
| 494 | Wuhan aphid virus 2 strain WHYC-2 segment 2 complete sequence | KR902726.1 | Terrestrial | Arthropoda | Insecta |
| 495 | Banna virus strain QTM104536 Banna VP2 gene complete cds | KX884639.1 | Terrestrial | Arthropoda | Insecta |
| 496 | Shuangao Lacewing Virus strain QSA01 polymerase (L) gene complete cds | KM817613.1 | Terrestrial | Arthropoda | Insecta |
| 497 | Changjiang Jingmen-like virus strain CJLX28200 NS3-like protein gene complete cds | KX883003.1 | Freshwater | Arthropoda | Crustacea |
| 498 | Wuchang Cockraoch Virus 3 strain WCZL-1 segment 1 complete sequence | KM817604.1 | Terrestrial | Arthropoda | Insecta |
| 499 | Bivalvia Durna-like virus H2 | GWHAPCN01000000 | Yellow China Sea | Mollusca | Bivalvia |
| 500 | Wenzhou picorna-like virus 31 strain WZFSL72092 hypothetical protein 1 and hypothetical protein 2 genes comple | KX884341.1 | Freshwater | Mollusca | Gastropoda |
| 501 | Beihai picorna-like virus 72 strain BHJJX25705 hypothetical protein 1 and hypothetical protein 2 genes complete cc | KX883368.1 | South China Sea | Arthropoda | Crustacea |
| 502 | Beihai picorna-like virus 72 strain BHHK126587 hypothetical protein 1 and hypothetical protein 2 genes complete ( | KX883344.1 | South China Sea | Cnidaria | Anthozoa |
| 503 | Beihai orthomyxo-like virus 2 strain BHHQ71863 polymerase PB1 gene complete cds | KX883843.1 | South China Sea | Tunicata | Ascidiacea |
| 504 | Hubei partiti-like virus 13 strain spider122140 RdRp gene complete cds | KX884160.1 | Terrestrial | Arthropoda | Chelicerata |
| 505 | Beihai sesarmid crab virus 1 strain SCJXSX39422 hypothetical protein 1 and hypothetical protein 2 genes complete | KX883967.1 | South China Sea | Arthropoda | Crustacea |
| 506 | Changjiang picorna-like virus 4 strain CJLX29654 hypothetical protein 1 and hypothetical protein 2 genes complete | KX884543.1 | Freshwater | Arthropoda | Crustacea |
| 507 | Shahe picorna-like virus 1 strain SHWC13622 hypothetical protein 1 and hypothetical protein 2 genes complete cds | KX883666.1 | Freshwater | Arthropoda | Mix |
| 508 | Hubei sobemo-like virus 38 strain QTM27202 hypothetical protein 1 hypothetical protein 2 and hypothetical protein | KX882846.1 | Terrestrial | Arthropoda | Insecta |
| 509 | Hubei tetragnatha maxillosa virus 1 strain SSZZ1726 hypothetical protein gene complete cds | KX883693.1 | Freshwater | Arthropoda | Chelicerata |
| 510 | Hubei tetragnatha maxillosa virus 1 strain arthropodmix23158 hypothetical protein gene complete cds | KX883296.1 | Terrestrial | Arthropoda | Mix |
| 511 | Shahe picorna-like virus 5 strain SHWC13542 hypothetical protein 1 and hypothetical protein 2 genes complete cds | KX883656.1 | Freshwater | Arthropoda | Mix |
| 512 | Wuhan Mosquito Virus 1 strain WT3-15 nucleopasid protein (N) gene complete cds | KM817758.1 | Terrestrial | Arthropoda | Insecta |
| 513 | Changjiang picorna-like virus 6 strain CJLX30470 hypothetical protein 1 and hypothetical protein 2 genes complete | KX884548.1 | Freshwater | Arthropoda | Crustacea |
| 514 | Beihai picorna-like virus 115 strain BHHK131520 hypothetical protein 1 and hypothetical protein 2 genes complete | KX883353.1 | South China Sea | Cnidaria | Anthozoa |

| | | | | | |
|---|---|---|---|---|---|
| 515 | Hubei partiti-like virus 11 strain QTM25188 segment Seg 2 hypothetical protein gene complete cds | KX884127.1 | Terrestrial | Arthropoda | Insecta |
| 516 | Wenzhou picorna-like virus 8 strain beimix75836 hypothetical protein 1 gene complete cds and hypothetical proteir | KX883303.1 | East China Sea | Mollusca | Bivalvia |
| 517 | Beihai picobirna-like virus 5 strain BHJJX25000 RdRp gene complete cds | KX884060.1 | South China Sea | Arthropoda | Crustacea |
| 518 | Kadipiro virus strain QTM27331 Kadipiro VP10 gene complete cds | KX884659.1 | Terrestrial | Arthropoda | Insecta |
| 519 | Beihai picorna-like virus 18 strain BHSC157282 hypothetical protein 1 and hypothetical protein 2 genes complete c | KX883409.1 | South China Sea | Annelida | Polychaeta |
| 520 | Jiujie Fly Virus strain JJ3-6 PB1 (PB1) gene complete cds | KM817616.1 | Terrestrial | Arthropoda | Insecta |
| 521 | Wenzhou picorna-like virus 4 strain WZSLuoI86003 hypothetical protein 1 and hypothetical protein 2 genes comple | KX884371.1 | East China Sea | Mollusca | Gastropoda |
| 522 | Hubei picobirna-like virus 4 strain WHBL66087 RdRp gene complete cds | KX884189.1 | Freshwater | Mollusca | Bivalvia |
| 523 | Hubei diptera virus 18 strain QTM25917 segment Seg 2 hypothetical protein gene complete cds | KX884113.1 | Terrestrial | Arthropoda | Insecta |
| 524 | Hubei diptera virus 18 strain SCM37169 segment Seg 2 hypothetical protein gene complete cds | KX884151.1 | Terrestrial | Arthropoda | Insecta |
| 525 | Hubei odonate virus 2 strain QTM74832 hypothetical protein gene complete cds | KX883926.1 | Terrestrial | Arthropoda | Insecta |
| 526 | Hubei picorna-like virus 14 strain QTM26191 hypothetical protein 1 and hypothetical protein 2 genes complete cds | KX883934.1 | Terrestrial | Arthropoda | Insecta |
| 527 | Beihai picobirna-like virus 8 strain BHNXC71065 segment Seg 2 hypothetical protein gene complete cds | KX884065.1 | South China Sea | Sipuncula | Sipuncula |
| 528 | Wuhan house centipede virus 7 strain WHYY9725 RdRp gene complete cds | KX884216.1 | Terrestrial | Arthropoda | Myriapoda |
| 529 | Changjiang crawfish virus 3 strain CJLX30786 hypothetical protein 1 and hypothetical protein 2 genes complete cds | KX884561.1 | Freshwater | Arthropoda | Crustacea |
| 530 | Hubei sobemo-like virus 14 strain QTM6420 hypothetical protein 1 and hypothetical protein 2 genes complete cds | KX882850.1 | Terrestrial | Arthropoda | Insecta |
| 531 | Sanxia atyid shrimp virus 3 strain SXXX37589 hypothetical protein 1 and hypothetical protein 2 genes complete cd | KX883714.1 | Freshwater | Arthropoda | Crustacea |
| 532 | Hubei sobemo-like virus 10 strain spider133871 hypothetical protein 1 and hypothetical protein 2 genes complete c | KX882878.1 | Terrestrial | Arthropoda | Chelicerata |
| 533 | Wuhan house centipede virus 5 strain WHYY23902 segment 2 putative capsid protein gene complete cds | KX882922.1 | Terrestrial | Arthropoda | Myriapoda |
| 534 | Hubei partiti-like virus 7 strain QTM23269 RdRp gene complete cds | KX884117.1 | Terrestrial | Arthropoda | Insecta |
| 535 | Wenzhou picorna-like virus 18 strain shrimp14534 hypothetical protein gene complete cds | KX883983.1 | East China Sea | Arthropoda | Crustacea |
| 536 | Hubei unio douglasiae virus 1 strain WHBL72336 capsid protein gene complete cds | KX883208.1 | Freshwater | Mollusca | Bivalvia |
| 537 | Hubei bunya-like virus 5 strain SCM50476 RNA-dependent RNA polymerase gene partial cds | KX884812.1 | Terrestrial | Arthropoda | Insecta |
| 538 | Wuhan spider virus 7 strain spider133569 RNA-dependent RNA polymerase gene complete cds | KX883564.1 | Terrestrial | Arthropoda | Chelicerata |
| 539 | Crustacea Hepeli-like virus D2 | GWHAPEJ01000000 | East China Sea | Arthropoda | Crustacea |
| 540 | Hubei tombus-like virus 23 strain WHCCII13249 hypothetical protein 1 and hypothetical protein 2 genes complete | KX883229.1 | Terrestrial | Arthropoda | Insecta |
| 541 | Hubei sobemo-like virus 33 strain LCM101922 hypothetical protein 1 and hypothetical protein 2 genes complete cd | KX882827.1 | Terrestrial | Arthropoda | Insecta |
| 542 | Beihai sobemo-like virus 18 strain BWBFG40814 hypothetical protein 1 and hypothetical protein 2 genes complete | KX882810.1 | South China Sea | Mollusca | Bivalvia |
| 543 | Bivalvia Picorna-like virus H5 | GWHAPCM01000000 | Yellow China Sea | Mollusca | Bivalvia |
| 544 | Changjiang tombus-like virus 2 strain CJLX30692 hypothetical protein 1 and hypothetical protein 2 genes complete | KX883098.1 | Freshwater | Arthropoda | Crustacea |
| 545 | Hubei tombus-like virus 31 strain WHBL72828 hypothetical protein 1 hypothetical protein 2 and hypothetical prote | KX883217.1 | Freshwater | Mollusca | Bivalvia |
| 546 | Beihai tombus-like virus 1 strain SCJXSX39078 hypothetical protein 1 hypothetical protein 2 and hypothetical prot | KX883144.1 | South China Sea | Arthropoda | Crustacea |
| 547 | Beihai levi-like virus 17 strain BHTH16066 hypothetical protein genes complete cds | KX883485.1 | South China Sea | Arthropoda | Crustacea |
| 548 | Beihai picorna-like virus 7 strain BHNXC41478 hypothetical protein 1 hypothetical protein 2 and hypothetical prot | KX883403.1 | South China Sea | Sipuncula | Sipuncula |
| 549 | Beihai Nido-like virus 1 strain BZL87871 hypothetical protein and putative structural polyprotein genes complete c | KX883629.1 | South China Sea | Mollusca | Gastropoda |
| 550 | Gill-associated virus strain shrimp14090 1ab gene partial cds | KX883632.1 | East China Sea | Arthropoda | Crustacea |
| 551 | Beihai tombus-like virus 7 strain BHBJDX18773 hypothetical protein 1 hypothetical protein 2 hypothetical protein | KX883021.1 | South China Sea | Arthropoda | Crustacea |

| | | | | | |
|---|---|---|---|---|---|
| 552 | Beihai tombus-like virus 7 strain BHWZXX15571 hypothetical protein 1 hypothetical protein 2 and hypothetical pr | KX883048.1 | South China Sea | Arthropoda | Crustacea |
| 553 | Beihai zhaovirus-like virus 5 strain BHNXC41311 RNA-dependent RNA polymerase gene complete cds | KX884036.1 | South China Sea | Sipuncula | Sipuncula |
| 554 | Beihai sobemo-like virus 4 strain BHZY59438 hypothetical protein 1 and hypothetical protein 2 genes complete cds | KX882801.1 | South China Sea | Mollusca | Cephalopoda |
| 555 | Hubei lepidoptera virus 5 strain LCM63289 hypothetical protein 2 gene complete cds | KX884625.1 | Terrestrial | Arthropoda | Insecta |
| 556 | Beihai charybdis crab virus 1 strain BHXun33339 RdRp and putative capsid protein genes complete cds | KX883753.1 | South China Sea | Arthropoda | Crustacea |
| 557 | Hubei noda-like virus 25 strain QTM19461 hypothetical protein gene complete cds | KX883128.1 | Terrestrial | Arthropoda | Insecta |
| 558 | Wenling noda-like virus 4 strain WLJQ84133 hypothetical protein gene complete cds | KX883256.1 | East China Sea | Arthropoda | Crustacea |
| 559 | Beihai horseshoe crab virus 1 strain HOU157708 hypothetical protein genes complete cds | KX883114.1 | South China Sea | Arthropoda | Chelicerata |
| 560 | Hubei tombus-like virus 14 strain spider118341 hypothetical protein 1 hypothetical protein 2 and hypothetical prote | KX883159.1 | Terrestrial | Arthropoda | Chelicerata |
| 561 | Beihai noda-like virus 6 strain BHWZXX9026 hypothetical protein 1 and hypothetical protein 2 genes complete cds | KX883055.1 | South China Sea | Arthropoda | Crustacea |
| 562 | Beihai levi-like virus 18 strain BHWZXX15152 hypothetical protein genes complete cds | KX883497.1 | South China Sea | Arthropoda | Crustacea |
| 563 | Wenzhou narna-like virus 9 strain WZSLuoI85295 RNA-dependent RNA polymerase gene complete cds | KX883619.1 | East China Sea | Mollusca | Gastropoda |
| 564 | Beihai noda-like virus 26 strain BHWZXX15658 hypothetical protein gene complete cds | KX883052.1 | South China Sea | Arthropoda | Crustacea |
| 565 | Beihai noda-like virus 26 strain BHBJDX18528 hypothetical protein gene complete cds | KX883019.1 | South China Sea | Arthropoda | Crustacea |
| 566 | Beihai noda-like virus 30 strain BHJJX6144 hypothetical protein gene complete cds | KX883031.1 | South China Sea | Arthropoda | Crustacea |
| 567 | Hubei Beny-like virus 1 strain SCM50761 hypothetical protein gene complete cds | KX883789.1 | Terrestrial | Arthropoda | Insecta |
| 568 | Beihai levi-like virus 9 strain BHTSS17966 hypothetical protein genes complete cds | KX883493.1 | South China Sea | Arthropoda | Crustacea |
| 569 | Hubei tombus-like virus 17 strain QTM26798 hypothetical protein 1 hypothetical protein 2 and hypothetical protein | KX883136.1 | Terrestrial | Arthropoda | Insecta |
| 570 | Hubei narna-like virus 5 strain SSZZ3007 RNA-dependent RNA polymerase gene complete cds | KX883565.1 | Freshwater | Arthropoda | Chelicerata |
| 571 | Sanxia tombus-like virus 2 strain SXXX29202 hypothetical protein 1 hypothetical protein 2 and hypothetical protein | KX883186.1 | Freshwater | Arthropoda | Crustacea |
| 572 | Melon necrotic spot virus strain SCM43651 hypothetical protein 1 hypothetical protein 2 and hypothetical protein 3 | KX883146.1 | Terrestrial | Arthropoda | Insecta |
| 573 | Bivalvia Picorna-like virus N13 | GWHAPGI01000000 | South China Sea | Mollusca | Bivalvia |
| 574 | Crustacea Picorna-like virus N25 | GWHAPDN01000000 | South China Sea | Arthropoda | Crustacea |
| 575 | Bivalvia Picorna-like virus D36 | GWHAPHY01000000 | East China Sea | Mollusca | Bivalvia |
| 576 | Beihai reo-like virus 2 strain BHBJDX18427 RdRp gene complete cds | KX884605.1 | South China Sea | Arthropoda | Crustacea |
| 577 | Jingmen tombus-like virus 2 strain JMYJCC11049 hypothetical protein 1 hypothetical protein 2 and hypothetical pr | KX883123.1 | Terrestrial | Nematoda | Chromadorea |
| 578 | Wenzhou tombus-like virus 10 strain WZFSL57346 hypothetical protein 1 and hypothetical protein 2 genes comple | KX883273.1 | Freshwater | Mollusca | Gastropoda |
| 579 | Crustacea Picorna-like virus N13 | GWHAPDA01000000 | South China Sea | Arthropoda | Crustacea |
| 580 | Hubei unio douglasiae virus 2 strain WHBL72658 hypothetical protein 1 hypothetical protein 2 and hypothetical pr | KX883215.1 | Freshwater | Mollusca | Bivalvia |
| 581 | Hubei tombus-like virus 8 strain WHCC116238 hypothetical protein 1 hypothetical protein 2 and hypothetical prote | KX883226.1 | Terrestrial | Arthropoda | Insecta |
| 582 | Sanxia tombus-like virus 4 strain SXXX7002 hypothetical protein 1 hypothetical protein 2 and hypothetical protein | KX883180.1 | Freshwater | Arthropoda | Crustacea |
| 583 | Bivalvia Sobeli-like virus H1 | GWHAPJG01000000 | Yellow China Sea | Mollusca | Bivalvia |
| 584 | Hubei narna-like virus 10 strain WHBL60499 hypothetical protein genes complete cds and hypothetical protein gen | KX883580.1 | Freshwater | Mollusca | Bivalvia |
| 585 | Hubei macula-like virus 1 strain QTM27280 hypothetical protein genes complete cds | KX883784.1 | Terrestrial | Arthropoda | Insecta |
| 586 | Beihai levi-like virus 20 strain BHJJX20688 hypothetical protein genes complete cds | KX883469.1 | South China Sea | Arthropoda | Crustacea |
| 587 | Wenzhou tombus-like virus 16 strain WZFSL78689 hypothetical protein gene complete cds | KX883259.1 | Freshwater | Mollusca | Gastropoda |
| 588 | Sanxia sobemo-like virus 5 strain SXSSP2561 hypothetical protein 1 and hypothetical protein 2 genes complete cds | KX882888.1 | Freshwater | Arthropoda | Insecta |

| | | | | | |
|---|---|---|---|---|---|
| 589 | Wenzhou tombus-like virus 9 strain WZSLuoI85212 hypothetical protein 1 hypothetical protein 2 and hypothetical | KX883279.1 | East China Sea | Mollusca | Gastropoda |
| 590 | Hubei sobemo-like virus 13 strain WHYY18840 hypothetical protein 1 hypothetical protein 2 and hypothetical prot | KX882928.1 | Terrestrial | Arthropoda | Myriapoda |
| 591 | Hubei noda-like virus 3 strain WGML145075 hypothetical protein gene complete cds | KX883197.1 | Terrestrial | Arthropoda | Myriapoda |
| 592 | Wenling narna-like virus 3 strain WLJQ100768 RNA-dependent RNA polymerase gene complete cds | KX883601.1 | East China Sea | Arthropoda | Crustacea |
| 593 | Beihai weivirus-like virus 9 strain BWBFG39428 RNA-dependent RNA polymerase and hypothetical protein genes | KX884026.1 | South China Sea | Mollusca | Bivalvia |
| 594 | Wenzhou sobemo-like virus 2 strain mosZJ15137 hypothetical protein 1 and hypothetical protein 2 genes complete | KX882829.1 | Terrestrial | Arthropoda | Insecta |
| 595 | Beihai zhaovirus-like virus 1 strain BHZY59866 polyprotein gene complete cds | KX884038.1 | South China Sea | Mollusca | Cephalopoda |
| 596 | Beihai picorna-like virus 32 strain BHZC37269 hypothetical protein gene complete cds | KX884485.1 | South China Sea | Mollusca | Bivalvia |
| 597 | Beihai picorna-like virus 32 strain BHNXC41402 hypothetical protein gene complete cds | KX883397.1 | South China Sea | Sipuncula | Sipuncula |
| 598 | Hubei partiti-like virus 10 strain QTM133141 RdRp gene complete cds | KX884114.1 | Terrestrial | Arthropoda | Insecta |
| 599 | Beihai tombus-like virus 14 strain BHZC37000 hypothetical protein gene complete cds | KX883060.1 | South China Sea | Mollusca | Bivalvia |
| 600 | Wuhan insect virus 20 strain WHCCII13322 hypothetical protein 1 hypothetical protein 2 and hypothetical protein | KX883231.1 | Terrestrial | Arthropoda | Insecta |
| 601 | Wenzhou tombus-like virus 6 strain WZFSL85189 hypothetical protein 1 hypothetical protein 2 and hypothetical pr | KX883261.1 | Freshwater | Mollusca | Gastropoda |
| 602 | Wuhan pillworm virus 3 strain WHSFII20254 segment 2 Capsid gene complete cds | KX882914.1 | Terrestrial | Arthropoda | Crustacea |
| 603 | Wenling toga-like virus strain WLJQ101932 hypothetical protein genes complete cds | KX883827.1 | East China Sea | Arthropoda | Crustacea |
| 604 | Hubei noda-like virus 4 strain WHBL72570 hypothetical protein 1 and hypothetical protein 2 genes complete cds | KX883214.1 | Freshwater | Mollusca | Bivalvia |
| 605 | Beihai sobemo-like virus 5 strain BHZY60175 hypothetical protein 1 and hypothetical protein 2 genes complete cds | KX882802.1 | South China Sea | Mollusca | Cephalopoda |
| 606 | Wenzhou qinvirus-like virus 2 strain WZRBX42684 RNA-dependent RNA polymerase gene complete cds | KX884000.1 | East China Sea | Arthropoda | Crustacea |
| 607 | Crustacea Hepeli-like virus D1 | GWHAPJS01000000 | East China Sea | Arthropoda | Crustacea |
| 608 | Beihai razor shell virus 4 strain BHZC37363 hypothetical protein gene complete cds | KX882949.1 | South China Sea | Mollusca | Bivalvia |
| 609 | Hubei toti-like virus 6 strain arthropodmix282 hypothetical protein 1 gene complete cds and hypothetical protein 2 | KX882940.1 | Terrestrial | Arthropoda | Mix |
| 610 | Wenzhou picorna-like virus 46 strain WZFSL85813 hypothetical protein gene complete cds | KX884326.1 | Freshwater | Mollusca | Gastropoda |
| 611 | Hubei orthoptera virus 4 strain ZCM21011 RNA-dependent RNA polymerase gene complete cds | KX883283.1 | Terrestrial | Arthropoda | Insecta |
| 612 | Hubei partiti-like virus 38 strain SCM245982 RdRp gene complete cds | KX884148.1 | Terrestrial | Arthropoda | Insecta |
| 613 | Wenzhou picorna-like virus 34 strain WZFSL80163 hypothetical protein 1 gene complete cds and hypothetical prot | KX884335.1 | Freshwater | Mollusca | Gastropoda |
| 614 | Beihai partiti-like virus 1 strain BHTSS8738 RdRp gene complete cds | KX884072.1 | South China Sea | Arthropoda | Crustacea |
| 615 | Wuhan pillworm virus 3 strain WHSFII20254 segment 1 hypothetical protein 1 and hypothetical protein 2 genes co | KX882913.1 | Terrestrial | Arthropoda | Crustacea |
| 616 | Hubei diptera virus 22 strain SCM45279 hypothetical protein 1 and hypothetical protein 2 genes complete cds | KX882972.1 | Terrestrial | Arthropoda | Insecta |
| 617 | Beihai partiti-like virus 3 strain BHHK112924 RdRp gene complete cds | KX884055.1 | South China Sea | Cnidaria | Anthozoa |
| 618 | Hubei partiti-like virus 41 strain QTM24887 RdRp gene complete cds | KX884125.1 | Terrestrial | Arthropoda | Insecta |
| 619 | Gastropoda Unclassified virus N1 | GWHAPGG01000000 | South China Sea | Mollusca | Gastropoda |
| 620 | Wenling narna-like virus 5 strain WLJQ94194 RNA-dependent RNA polymerase gene complete cds | KX883600.1 | East China Sea | Arthropoda | Crustacea |
| 621 | Beihai sipunculid worm virus 3 strain BHZC37455 hypothetical protein 1 and hypothetical protein 2 genes complet | KX884501.1 | South China Sea | Mollusca | Bivalvia |
| 622 | Beihai sipunculid worm virus 3 strain BHNXC41487 hypothetical protein 1 gene complete cds and hypothetical pro | KX883406.1 | South China Sea | Sipuncula | Sipuncula |
| 623 | Beihai picorna-like virus 62 strain BHBei74566 hypothetical protein gene complete cds | KX883309.1 | South China Sea | Mollusca | Bivalvia |
| 624 | Shahe hepe-like virus 2 strain SHWC13612 hypothetical protein genes complete cds | KX883797.1 | Freshwater | Arthropoda | Mix |
| 625 | Hubei orthomyxo-like virus 5 strain WHBL85127 polymerase PB1 gene complete cds | KX883883.1 | Freshwater | Mollusca | Bivalvia |

| | | | | | |
|---|---|---|---|---|---|
| 626 | Shahe hepe-like virus 1 strain SHWC13572 replicase hypothetical protein putative structural protein and hypothetic | KX883796.1 | Freshwater | Arthropoda | Mix |
| 627 | Hubei picorna-like virus 46 strain spider133332 hypothetical protein gene complete cds | KX883677.1 | Terrestrial | Arthropoda | Chelicerata |
| 628 | Wuhan Louse Fly Virus 3 strain BFJSC-1 PB1 (PB1) gene complete cds | KM817620.1 | Terrestrial | Arthropoda | Insecta |
| 629 | Shahe bunya-like virus 3 strain SHWC0209c12636 RNA-dependent RNA polymerase gene complete cds | KX884824.1 | Freshwater | Arthropoda | Crustacea |
| 630 | Beihai picorna-like virus 22 strain HOU126951 hypothetical protein 1 gene complete cds and hypothetical protein 2 | KX883905.1 | South China Sea | Arthropoda | Chelicerata |
| 631 | Hubei diptera virus 17 strain SCM172187 segment Seg 1 RdRp gene complete cds | KX884145.1 | Terrestrial | Arthropoda | Insecta |
| 632 | Hubei picorna-like virus 20 strain WGML146806 hypothetical protein 1 and hypothetical protein 2 genes complete | KX884254.1 | Terrestrial | Arthropoda | Myriapoda |
| 633 | Crustacea Ghabri-like virus H1 | GWHAPGV01000000 | Yellow China Sea | Arthropoda | Crustacea |
| 634 | Hubei picorna-like virus 43 strain QTM27027 hypothetical protein gene complete cds | KX883938.1 | Terrestrial | Arthropoda | Insecta |
| 635 | Wuhan insect virus 22 strain WHZM10073 segment Seg 1 RdRp gene complete cds | KX884221.1 | Terrestrial | Arthropoda | Insecta |
| 636 | Wuhan insect virus 22 strain spider98584 segment Seg 1 RdRp gene complete cds | KX884171.1 | Terrestrial | Arthropoda | Chelicerata |
| 637 | Wuhan insect virus 22 strain arthropodmix13806 segment Seg 1 RdRp gene complete cds | KX884050.1 | Terrestrial | Arthropoda | Mix |
| 638 | Shahe picorna-like virus 10 strain SHWC13586 hypothetical protein gene complete cds | KX883658.1 | Freshwater | Arthropoda | Mix |
| 639 | Cucumber green mottle mosaic virus strain spider132626 replicase movement protein and coat protein genes compl | KX883801.1 | Terrestrial | Arthropoda | Chelicerata |
| 640 | Cucumber green mottle mosaic virus strain BHZC36029 replicase movement protein and coat protein genes comple | KX883754.1 | South China Sea | Mollusca | Bivalvia |
| 641 | Cucumber green mottle mosaic virus strain CJLX28437 replicase movement protein and capsid genes complete cds | KX883761.1 | Freshwater | Arthropoda | Crustacea |
| 642 | Cucumber green mottle mosaic virus strain WZFSL84053 hypothetical protein genes complete cds | KX883833.1 | Freshwater | Mollusca | Gastropoda |
| 643 | Cucumber green mottle mosaic virus strain SCM50440 replicase movement protein and coat protein genes complete | KX883788.1 | Terrestrial | Arthropoda | Insecta |
| 644 | Beihai narna-like virus 2 strain BHZY58217 RNA-dependent RNA polymerase gene complete cds | KX883512.1 | South China Sea | Mollusca | Cephalopoda |
| 645 | Hubei dimarhabdovirus virus 4 strain SCM46012 RNA-dependent RNA polymerase gene complete cds | KX884435.1 | Terrestrial | Arthropoda | Insecta |
| 646 | Hubei narna-like virus 2 strain WHBL67117 RNA-dependent RNA polymerase gene complete cds | KX883578.1 | Freshwater | Mollusca | Bivalvia |
| 647 | Wenling crustacean virus 9 strain WLJQ100911 RNA-dependent RNA polymerase gene complete cds | KX884856.1 | East China Sea | Arthropoda | Crustacea |
| 648 | Sanxia tombus-like virus 1 strain SXXX36383 hypothetical protein 3 hypothetical protein 2 and hypothetical protei | KX883185.1 | Freshwater | Arthropoda | Crustacea |
| 649 | Sanxia tombus-like virus 1 strain WHCCII13265 hypothetical protein 1 hypothetical protein 2 and hypothetical prot | KX883230.1 | Terrestrial | Arthropoda | Insecta |
| 650 | Jiangxia Mosquito Virus 1 strain JX1-12 RNA-dependent RNA polymerase (L) gene complete cds | KM817670.1 | Terrestrial | Arthropoda | Insecta |
| 651 | Gastropoda Unclassified virus D4 | GWHAPBC01000000 | East China Sea | Mollusca | Gastropoda |
| 652 | Wuhan flea virus strain WHZM segment 2 complete sequence | KR902714.1 | Terrestrial | Arthropoda | Mix |
| 653 | Lihan Tick Virus strain LH-1 RNA-dependent RNA polymerase (L) gene complete cds | KM817672.1 | Terrestrial | Arthropoda | Chelicerata |
| 654 | Wuhan Insect virus 3 strain WHYC05 RNA-dependent RNA polymerase (L) gene complete cds | KM817693.1 | Terrestrial | Arthropoda | Insecta |
| 655 | Changjiang astro-like virus strain CJLX30757 putative protease hypothetical protein RNA-dependent RNA polymei | KX884566.1 | Freshwater | Arthropoda | Crustacea |
| 656 | Hubei sobemo-like virus 48 strain QCM18154 segment 1 hypothetical protein 1 and hypothetical protein 2 genes co | KX882834.1 | Terrestrial | Arthropoda | Insecta |
| 657 | Hubei sobemo-like virus 49 strain QTM12655 hypothetical protein 1 and hypothetical protein 2 genes complete cds | KX882836.1 | Terrestrial | Arthropoda | Insecta |
| 658 | Wuhan insect virus 28 strain WHZM10169 hypothetical protein 1 and hypothetical protein 2 genes complete cds | KX882988.1 | Terrestrial | Arthropoda | Insecta |
| 659 | Beihai picorna-like virus 13 strain BHZC37435 hypothetical protein gene partial cds | KX884497.1 | South China Sea | Mollusca | Bivalvia |
| 660 | Hubei lepidoptera virus 5 strain LCM63289 RNA-dependent RNA polymerase gene complete cds | KX884623.1 | Terrestrial | Arthropoda | Insecta |
| 661 | Hubei lepidoptera virus 4 strain LCM63323 RNA-dependent RNA polymerase gene complete cds | KX884628.1 | Terrestrial | Arthropoda | Insecta |
| 662 | Beihai levi-like virus 12 strain HWRTX21601 hypothetical protein genes complete cds | KX883535.1 | South China Sea | Arthropoda | Crustacea |

| | | | | | |
|---|---|---|---|---|---|
| 663 | Hubei lepidoptera virus 4 strain LCM63323 hypothetical protein 1 gene complete cds | KX884629.1 | Terrestrial | Arthropoda | Insecta |
| 664 | Hubei lepidoptera virus 5 strain LCM63289 hypothetical protein 1 gene complete cds | KX884624.1 | Terrestrial | Arthropoda | Insecta |
| 665 | Bivalvia Picorna-like virus D66 | GWHAPIB01000000 | East China Sea | Mollusca | Bivalvia |
| 666 | Beihai picorna-like virus 88 strain BHBJDX16027 hypothetical protein gene partial cds | KX883322.1 | South China Sea | Arthropoda | Crustacea |
| 667 | Gastropoda Picorna-like virus D12 | GWHAPBA01000000 | East China Sea | Mollusca | Gastropoda |
| 668 | Hubei picorna-like virus 4 strain WHBL90007 hypothetical protein 1 and hypothetical protein 2 genes complete cds | KX884265.1 | Freshwater | Mollusca | Bivalvia |
| 669 | Beihai noda-like virus 15 strain BHZY56617 hypothetical protein gene partial cds | KX883066.1 | South China Sea | Mollusca | Cephalopoda |
| 670 | Bivalvia Picorna-like virus D87 | GWHAPIH01000000 | East China Sea | Mollusca | Bivalvia |
| 671 | Sanxia picorna-like virus 11 strain SXXX37695 hypothetical protein 1 and hypothetical protein 2 genes complete cc | KX883712.1 | Freshwater | Arthropoda | Crustacea |
| 672 | Changjiang picorna-like virus 7 strain CJLX30780 hypothetical protein 1 and hypothetical protein 2 genes complete | KX884560.1 | Freshwater | Arthropoda | Crustacea |
| 673 | Hubei picorna-like virus 19 strain SZmix17531 hypothetical protein 1 gene complete cds and hypothetical protein 2 | KX883724.1 | Freshwater | Annelida | Clitellata |
| 674 | Hubei sobemo-like virus 9 strain spider112041 hypothetical protein 1 and hypothetical protein 2 genes complete cds | KX882865.1 | Terrestrial | Arthropoda | Chelicerata |
| 675 | Hubei partiti-like virus 17 strain QTM75790 RdRp gene complete cds | KX884143.1 | Terrestrial | Arthropoda | Insecta |
| 676 | Hubei picorna-like virus 48 strain spider129651 hypothetical protein gene complete cds | KX883676.1 | Terrestrial | Arthropoda | Chelicerata |
| 677 | Hubei partiti-like virus 3 strain arthropodmix2414 RdRp gene complete cds | KX884052.1 | Terrestrial | Arthropoda | Mix |
| 678 | Hubei partiti-like virus 3 strain SSZZ4709 RdRp gene complete cds | KX884177.1 | Freshwater | Arthropoda | Chelicerata |
| 679 | Hubei partiti-like virus 3 strain QTM75369 RdRp gene complete cds | KX884142.1 | Terrestrial | Arthropoda | Insecta |
| 680 | Hubei picorna-like virus 13 strain QTM26736 hypothetical protein gene complete cds | KX883936.1 | Terrestrial | Arthropoda | Insecta |
| 681 | Hubei partiti-like virus 51 strain WHWN52466 RdRp gene complete cds | KX884206.1 | Freshwater | Mollusca | Gastropoda |
| 682 | Beihai barnacle virus 4 strain BHTH16136 hypothetical protein gene complete cds | KX883416.1 | South China Sea | Arthropoda | Crustacea |
| 683 | Sanxia water strider virus 11 strain SXSSP2582 hypothetical protein 1 and hypothetical protein 2 genes complete cc | KX882891.1 | Freshwater | Arthropoda | Insecta |
| 684 | Beihai picorna-like virus 99 strain BHJJX22326 hypothetical protein gene complete cds | KX883362.1 | South China Sea | Arthropoda | Crustacea |
| 685 | Wuhan insect virus 34 strain CC64594 hypothetical protein 1 and hypothetical protein 2 genes complete cds | KX882815.1 | Terrestrial | Arthropoda | Insecta |
| 686 | Shahe narna-like virus 7 strain SHWC0209c12713 RNA-dependent RNA polymerase gene complete cds | KX883560.1 | Freshwater | Arthropoda | Crustacea |
| 687 | Wuhan Louse Fly Virus 2 strain BFJSC-11 RNA-dependent RNA polymerase (L) gene partial cds | KM817694.1 | Terrestrial | Arthropoda | Insecta |
| 688 | Beihai narna-like virus 15 strain BHHZL10358 RNA-dependent RNA polymerase gene complete cds | KX883468.1 | South China Sea | Arthropoda | Crustacea |
| 689 | Hubei narna-like virus 4 strain WHYY20523 RNA-dependent RNA polymerase gene complete cds | KX883596.1 | Terrestrial | Arthropoda | Myriapoda |
| 690 | Wuhan Fly Virus 3 strain SYY2-5 RNA-dependent RNA polymerase (L) gene complete cds | KM817647.1 | Terrestrial | Arthropoda | Insecta |
| 691 | Sanxia Qinvirus-like virus 1 strain SXXX11646 RNA-dependent RNA polymerase gene complete cds | KX883996.1 | Freshwater | Arthropoda | Crustacea |
| 692 | Xinzhou partiti-like virus 1 strain XZSJSC65291 segment Seg 2 hypothetical protein gene complete cds | KX884239.1 | Terrestrial | Nematoda | Mix |
| 693 | Beihai partiti-like virus 4 strain YYSZX180 RdRp gene complete cds | KX884241.1 | South China Sea | Arthropoda | Crustacea |
| 694 | Hubei toti-like virus 16 strain spider123775 hypothetical protein 1 hypothetical protein 2 and hypothetical protein 3 | KX882975.1 | Terrestrial | Arthropoda | Chelicerata |
| 695 | Wuhan large pig roundworm virus 1 strain WHZHC73278 segment Seg 2 hypothetical protein gene complete cds | KX884220.1 | Terrestrial | Nematoda | Chromadorea |
| 696 | Beihai sesarmid crab virus 2 strain SCJXSX39002 hypothetical protein gene complete cds | KX883959.1 | South China Sea | Arthropoda | Crustacea |
| 697 | Jingmen tombus-like virus 1 strain JMYJCC68055 hypothetical protein genes complete cds | KX883122.1 | Terrestrial | Nematoda | Chromadorea |
| 698 | Wuhan arthropod virus 4 strain WHCC117028 segment 2 hypothetical protein gene complete cds | KX882907.1 | Terrestrial | Arthropoda | Insecta |
| 699 | Beihai picorna-like virus 36 strain BHJJX25939 hypothetical protein gene partial cds | KX883373.1 | South China Sea | Arthropoda | Crustacea |

| | | | | | |
|---|---|---|---|---|---|
| 700 | Hubei sobemo-like virus 48 strain QCM18154 segment 2 hypothetical protein gene complete cds | KX882835.1 | Terrestrial | Arthropoda | Insecta |
| 701 | Wenzhou picorna-like virus 30 strain WZSBei63630 hypothetical protein 1 gene complete cds and hypothetical pro | KX884354.1 | East China Sea | Mollusca | Bivalvia |
| 702 | Shuangao insect virus 7 strain SKC segment 2 complete sequence | KR902718.1 | Terrestrial | Arthropoda | Insecta |
| 703 | Bivalvia Picorna-like virus D28 | GWHAOYK0100000C | East China Sea | Mollusca | Bivalvia |
| 704 | Pepper mild mottle virus replicase gene partial cds and movement protein and coat protein genes complete cds | KX883742.1 | Terrestrial | Arthropoda | Mix |
| 705 | Pepper mild mottle virus strain WHZM9517 183 kDa protein and hypothetical protein genes complete cds | KX883825.1 | Terrestrial | Arthropoda | Insecta |
| 706 | Hubei partiti-like virus 8 strain WHCC37315 RdRp gene partial cds | KX884192.1 | Terrestrial | Arthropoda | Insecta |
| 707 | Hubei partiti-like virus 9 strain WHCC72771 RdRp gene complete cds | KX884193.1 | Terrestrial | Arthropoda | Insecta |
| 708 | Hubei partiti-like virus 55 strain SZmix21556 RdRp gene partial cds | KX884184.1 | Freshwater | Annelida | Clitellata |
| 709 | Cephalopoda Unclassified virus H1 | GWHAPGY01000000 | Yellow China Sea | Mollusca | Cephalopoda |
| 710 | Beihai picorna-like virus 40 strain BHNXC39087 hypothetical protein gene complete cds | KX883385.1 | South China Sea | Sipuncula | Sipuncula |
| 711 | Bivalvia Picorna-like virus D17 | GWHAPKA01000000 | East China Sea | Mollusca | Bivalvia |
| 712 | Hubei toti-like virus 13 strain QTM25941 hypothetical protein 1 and hypothetical protein 2 genes complete cds | KX882966.1 | Terrestrial | Arthropoda | Insecta |
| 713 | Bivalvia Picorna-like virus N22 | GWHAPGL01000000 | South China Sea | Mollusca | Bivalvia |
| 714 | Hubei toti-like virus 15 strain SCM51462 hypothetical protein 1 and hypothetical protein 2 genes complete cds | KX882974.1 | Terrestrial | Arthropoda | Insecta |
| 715 | Hubei tombus-like virus 30 strain spider132818 hypothetical protein 1 and hypothetical protein 2 genes complete cc | KX883166.1 | Terrestrial | Arthropoda | Chelicerata |
| 716 | Beihai partiti-like virus 11 strain BHWZXX14306 RdRp gene complete cds | KX884074.1 | South China Sea | Arthropoda | Crustacea |
| 717 | Beihai partiti-like virus 11 strain BHBJDX17847 RdRp gene complete cds | KX884054.1 | South China Sea | Arthropoda | Crustacea |
| 718 | Hubei picorna-like virus 30 strain QTM27289 hypothetical protein gene complete cds | KX883952.1 | Terrestrial | Arthropoda | Insecta |
| 719 | Bivalvia Picorna-like virus D1 | GWHAPJT01000000 | East China Sea | Mollusca | Bivalvia |
| 720 | Gastropoda Picorna-like virus D26 | GWHAPKK01000000 | East China Sea | Mollusca | Gastropoda |
| 721 | Hubei odonate virus 1 strain QTM27271 hypothetical protein gene complete cds | KX883947.1 | Terrestrial | Arthropoda | Insecta |
| 722 | Hubei sobemo-like virus 18 strain arthropodmix13949 hypothetical protein genes complete cds | KX882772.1 | Terrestrial | Arthropoda | Mix |
| 723 | Hubei sobemo-like virus 18 strain WHYY23868 hypothetical protein 1 and hypothetical protein 2 genes complete c | KX882925.1 | Terrestrial | Arthropoda | Myriapoda |
| 724 | Hubei picorna-like virus 32 strain QTM25882 hypothetical protein gene complete cds | KX883933.1 | Terrestrial | Arthropoda | Insecta |
| 725 | Hubei picorna-like virus 34 strain QCM148994 hypothetical protein gene complete cds | KX883923.1 | Terrestrial | Arthropoda | Insecta |
| 726 | Hubei sobemo-like virus 46 strain spider133700 hypothetical protein 1 and hypothetical protein 2 genes complete c | KX882875.1 | Terrestrial | Arthropoda | Chelicerata |
| 727 | Hubei sobemo-like virus 32 strain spider112797 hypothetical protein 1 and hypothetical protein 2 genes complete c | KX882866.1 | Terrestrial | Arthropoda | Chelicerata |
| 728 | Sanxia sobemo-like virus 3 strain SXSSP2531 hypothetical protein 1 and hypothetical protein 2 genes complete cds | KX882887.1 | Freshwater | Arthropoda | Insecta |
| 729 | Wenzhou picorna-like virus 16 strain WZFSL67369 hypothetical protein gene complete cds | KX884342.1 | Freshwater | Mollusca | Gastropoda |
| 730 | Hubei partiti-like virus 53 strain SZmix20674 RdRp gene complete cds | KX884182.1 | Freshwater | Annelida | Clitellata |
| 731 | Wenzhou picorna-like virus 54 strain WZSBei68749 hypothetical protein gene complete cds | KX884355.1 | East China Sea | Mollusca | Bivalvia |
| 732 | Hubei sobemo-like virus 11 strain spider125434 hypothetical protein 1 and hypothetical protein 2 genes complete c | KX882868.1 | Terrestrial | Arthropoda | Chelicerata |
| 733 | Beihai picobirna-like virus 4 strain BHJJX24179 RdRp gene complete cds | KX884059.1 | South China Sea | Arthropoda | Crustacea |
| 734 | Sanxia picorna-like virus 7 strain SXXX37313 hypothetical protein gene partial cds | KX883716.1 | Freshwater | Arthropoda | Crustacea |
| 735 | Wenzhou tombus-like virus 17 strain WZFSL82820 hypothetical protein 1 and hypothetical protein 2 genes comple | KX883265.1 | Freshwater | Mollusca | Gastropoda |
| 736 | Wenzhou picorna-like virus 7 strain shrimp14504 hypothetical protein 1 and hypothetical protein 2 genes complete | KX883982.1 | East China Sea | Arthropoda | Crustacea |

| | | | | | |
|---|---|---|---|---|---|
| 737 | Beihai partiti-like virus 2 strain BHZY60797 segment Seg 2 hypothetical protein gene complete cds | KX884084.1 | South China Sea | Mollusca | Cephalopoda |
| 738 | Beihai picorna-like virus 124 strain BHBJDX18598 hypothetical protein 1 and hypothetical protein 2 genes comple | KX883331.1 | South China Sea | Arthropoda | Crustacea |
| 739 | Hubei lepidoptera virus 5 strain LCM63289 hypothetical protein 4 gene complete cds | KX884627.1 | Terrestrial | Arthropoda | Insecta |
| 740 | Beihai picorna-like virus 64 strain BHJP54755 hypothetical protein 1 and hypothetical protein 2 genes complete cds | KX883382.1 | South China Sea | Echinodermata | Echinozoa |
| 741 | Beihai picorna-like virus 64 strain BHHQ137467 hypothetical protein 1 and hypothetical protein 2 genes complete | KX883355.1 | South China Sea | Tunicata | Ascidiacea |
| 742 | Hubei sobemo-like virus 23 strain QTM27250 hypothetical protein 1 and hypothetical protein 2 genes complete cds | KX882848.1 | Terrestrial | Arthropoda | Insecta |
| 743 | Hubei sobemo-like virus 47 strain spider133229 hypothetical protein 1 and hypothetical protein 2 genes complete c | KX882871.1 | Terrestrial | Arthropoda | Chelicerata |
| 744 | Hubei tick virus 3 strain tick108426 hypothetical protein gene complete cds | KX883728.1 | Terrestrial | Arthropoda | Chelicerata |
| 745 | Beihai picorna-like virus 69 strain HOU158647 hypothetical protein gene complete cds | KX883900.1 | South China Sea | Arthropoda | Chelicerata |
| 746 | Bivalvia Hepeli-like virus D2 | GWHAPAK01000000 | East China Sea | Mollusca | Bivalvia |
| 747 | Shahe picorna-like virus 2 strain SHWC0209c12507 hypothetical protein 1 and hypothetical protein 2 genes comple | KX883646.1 | Freshwater | Arthropoda | Crustacea |
| 748 | Hubei picorna-like virus 2 strain WHBL73152 hypothetical protein 1 and hypothetical protein 2 genes complete cds | KX884264.1 | Freshwater | Mollusca | Bivalvia |
| 749 | Banna virus strain QTM104536 Banna VP9 gene complete cds | KX884646.1 | Terrestrial | Arthropoda | Insecta |
| 750 | Bivalvia Picorna-like virus D83 | GWHAPIL01000000 | East China Sea | Mollusca | Bivalvia |
| 751 | Beihai octopus viurs 1 strain BHZY180716 hypothetical protein 1 and hypothetical protein 2 genes complete cds | KX884503.1 | South China Sea | Mollusca | Cephalopoda |
| 752 | Wenzhou picorna-like virus 1 strain WZSLuoI85940 hypothetical protein 1 and hypothetical protein 2 genes comple | KX884370.1 | East China Sea | Mollusca | Gastropoda |
| 753 | Wuhan insect virus 33 strain WHYY3291 hypothetical protein 1 and hypothetical protein 2 genes complete cds | KX884304.1 | Terrestrial | Arthropoda | Myriapoda |
| 754 | Hubei tetragnatha maxillosa virus 9 strain QTM27093 hypothetical protein 7 gene complete cds | KX884682.1 | Terrestrial | Arthropoda | Insecta |
| 755 | Hubei tetragnatha maxillosa virus 9 strain SSZZ3444 hypothetical protein 6 gene complete cds | KX884709.1 | Freshwater | Arthropoda | Chelicerata |
| 756 | Beihai picorna-like virus 27 strain BHNXC41439 hypothetical protein 1 and hypothetical protein 2 genes complete | KX883399.1 | South China Sea | Sipuncula | Sipuncula |
| 757 | Hubei tetragnatha maxillosa virus 2 strain QTM20713 hypothetical protein gene complete cds | KX883930.1 | Terrestrial | Arthropoda | Insecta |
| 758 | Hubei tetragnatha maxillosa virus 2 strain SSZZ3510 hypothetical protein gene complete cds | KX883694.1 | Freshwater | Arthropoda | Chelicerata |
| 759 | Hubei tetragnatha maxillosa virus 2 strain arthropodmix14091 hypothetical protein gene complete cds | KX883294.1 | Terrestrial | Arthropoda | Mix |
| 760 | Hubei odonate virus 15 strain QTM133059 hypothetical protein 3 gene complete cds | KX884667.1 | Terrestrial | Arthropoda | Insecta |
| 761 | Beihai partiti-like virus 5 strain YYSZX17448 RdRp gene complete cds | KX884240.1 | South China Sea | Arthropoda | Crustacea |
| 762 | Hubei partiti-like virus 49 strain spider118314 RdRp gene complete cds | KX884159.1 | Terrestrial | Arthropoda | Chelicerata |
| 763 | Beihai picorna-like virus 47 strain SCJXSX39359 hypothetical protein 1 and hypothetical protein 2 genes complete | KX883965.1 | South China Sea | Arthropoda | Crustacea |
| 764 | Sanxia picorna-like virus 2 strain SXXX37883 hypothetical protein 1 and hypothetical protein 2 genes complete cds | KX883709.1 | Freshwater | Arthropoda | Crustacea |
| 765 | Hubei partiti-like virus 25 strain QTM20591 RdRp gene complete cds | KX884115.1 | Terrestrial | Arthropoda | Insecta |
| 766 | Zhejiang mosquito virus 1 strain 3mos6204 hypothetical protein gene complete cds | KX883285.1 | Terrestrial | Arthropoda | Insecta |
| 767 | Zhejiang mosquito virus 1 strain mosHB236219 hypothetical protein gene complete cds | KX883916.1 | Terrestrial | Arthropoda | Insecta |
| 768 | Zhejiang mosquito virus 1 strain mosZJ35497 hypothetical protein gene complete cds | KX883919.1 | Terrestrial | Arthropoda | Insecta |
| 769 | Beihai picobirna-like virus 1 strain BHJJX25356 RdRp gene complete cds | KX884058.1 | South China Sea | Arthropoda | Crustacea |
| 770 | Hubei partiti-like virus 56 strain QTM26145 RdRp gene complete cds | KX884129.1 | Terrestrial | Arthropoda | Insecta |
| 771 | Hubei picorna-like virus 25 strain QTM27237 hypothetical protein 1 and hypothetical protein 2 genes complete cds | KX883945.1 | Terrestrial | Arthropoda | Insecta |
| 772 | Bivalvia Picorna-like virus D67 | GWHAPKR01000000 | East China Sea | Mollusca | Bivalvia |
| 773 | Beihai picorna-like virus 49 strain BHZC36855 hypothetical protein 1 and hypothetical protein 2 genes complete cd | KX884480.1 | South China Sea | Mollusca | Bivalvia |

| | | | | | |
|---|---|---|---|---|---|
| 774 | Hubei partiti-like virus 35 strain WHYY30311 RdRp gene complete cds | KX884218.1 | Terrestrial | Arthropoda | Myriapoda |
| 775 | Hubei partiti-like virus 46 strain QTM26281 RdRp gene complete cds | KX884130.1 | Terrestrial | Arthropoda | Insecta |
| 776 | Sanxia picorna-like virus 10 strain SXXX37528 hypothetical protein gene complete cds | KX883715.1 | Freshwater | Arthropoda | Crustacea |
| 777 | Bivalvia Picorna-like virus D49 | GWHAOZT01000000 | East China Sea | Mollusca | Bivalvia |
| 778 | Hubei picorna-like virus 53 strain GCM3912 hypothetical protein gene complete cds | KX883894.1 | Terrestrial | Arthropoda | Insecta |
| 779 | Hubei tetragnatha maxillosa virus 8 strain QTM24815 segment Seg 3 hypothetical protein gene complete cds | KX884123.1 | Terrestrial | Arthropoda | Insecta |
| 780 | Hubei tetragnatha maxillosa virus 8 strain arthropodmix13417 segment Seg 3 hypothetical protein gene complete cc | KX884048.1 | Terrestrial | Arthropoda | Mix |
| 781 | Hubei tetragnatha maxillosa virus 8 strain SSZZ3448 segment Seg 3 hypothetical protein gene complete cds | KX884175.1 | Freshwater | Arthropoda | Chelicerata |
| 782 | Beihai picorna-like virus 90 strain BHWZXX14572 hypothetical protein 1 and hypothetical protein 2 genes comple | KX883424.1 | South China Sea | Arthropoda | Crustacea |
| 783 | Banna virus strain QTM104536 Banna VP10 gene complete cds | KX884647.1 | Terrestrial | Arthropoda | Insecta |
| 784 | Bivalvia Picorna-like virus D86 | GWHAPIJ01000000 | East China Sea | Mollusca | Bivalvia |
| 785 | Hubei picorna-like virus 71 strain spider133937 hypothetical protein 1 hypothetical protein 2 hypothetical protein 3 | KX883687.1 | Terrestrial | Arthropoda | Chelicerata |
| 786 | Bivalvia Picorna-like virus N42 | GWHAPHS01000000 | South China Sea | Mollusca | Bivalvia |
| 787 | Xinzhou nematode virus 4 strain XZSJSC65771 putative nucleoprotein hypothetical protein 2 hypothetical protein 3 | KX884462.1 | Terrestrial | Nematoda | Mix |
| 788 | Gastropoda Picorna-like virus D28 | GWHAPKH01000000 | East China Sea | Mollusca | Gastropoda |
| 789 | Hubei picorna-like virus 37 strain QTM27228 hypothetical protein gene complete cds | KX883944.1 | Terrestrial | Arthropoda | Insecta |
| 790 | Hubei tetragnatha maxillosa virus 8 strain SSZZ3448 segment Seg 4 hypothetical protein gene complete cds | KX884176.1 | Freshwater | Arthropoda | Chelicerata |
| 791 | Hubei tetragnatha maxillosa virus 8 strain QTM24815 segment Seg 4 hypothetical protein gene complete cds | KX884124.1 | Terrestrial | Arthropoda | Insecta |
| 792 | Hubei tetragnatha maxillosa virus 8 strain arthropodmix13417 segment Seg 4 hypothetical protein gene complete cc | KX884049.1 | Terrestrial | Arthropoda | Mix |
| 793 | Beihai picorna-like virus 87 strain BHJJX25970 hypothetical protein gene complete cds | KX883378.1 | South China Sea | Arthropoda | Crustacea |
| 794 | Hubei orthoptera virus 6 strain ZCM21154 segment 2 polymerase PB1 gene complete cds | KX883884.1 | Terrestrial | Arthropoda | Insecta |
| 795 | Hubei lepidoptera virus 2 strain LCM101902 putative nucleoprotein hypothetical protein 2 putative membrane prote | KX884415.1 | Terrestrial | Arthropoda | Insecta |
| 796 | Wuhan Insect virus 7 strain WHYC02 nucleocapsid (N) putative phosphoprotein (ORF2) matrix protein (M) glycop | KM817653.1 | Terrestrial | Arthropoda | Insecta |
| 797 | Hubei earwig virus 1 strain GCM19162 segment 2 polymerase PB1 gene complete cds | KX883858.1 | Terrestrial | Arthropoda | Insecta |
| 798 | Hubei earwig virus 1 strain WGML148351 polymerase PB1 gene partial cds | KX883882.1 | Terrestrial | Arthropoda | Myriapoda |
| 799 | Wuhan Mothfly Virus strain WHDL03 PB1 (PB1) gene complete cds | KM817627.1 | Terrestrial | Arthropoda | Insecta |
| 800 | Beihai orthomyxo-like virus 1 strain BHTSS17872 segment 5 nucleocapsid protein gene complete cds | KX883848.1 | South China Sea | Arthropoda | Crustacea |
| 801 | Bivalvia Picorna-like virus D35 | GWHAPHX01000000 | East China Sea | Mollusca | Bivalvia |
| 802 | Beihai picorna-like virus 24 strain BHZY60459 hypothetical protein 1 and hypothetical protein 2 genes complete cc | KX884507.1 | South China Sea | Mollusca | Cephalopoda |
| 803 | Wuhan insect virus 33 strain WHCCII11871 hypothetical protein 1 and hypothetical protein 2 genes complete cds | KX884274.1 | Terrestrial | Arthropoda | Insecta |
| 804 | Hubei partiti-like virus 50 strain spider58532 RdRp gene complete cds | KX884169.1 | Terrestrial | Arthropoda | Chelicerata |
| 805 | Hubei partiti-like virus 34 strain WHWN68514 RdRp gene complete cds | KX884207.1 | Freshwater | Mollusca | Gastropoda |
| 806 | Hubei reo-like virus 4 strain WGML144712 hypothetical protein 3 gene partial cds | KX884715.1 | Terrestrial | Arthropoda | Myriapoda |
| 807 | Beihai sea slater virus 4 strain BHHZL10410 RdRp hypothetical protein putative structural protein and hypothetical | KX883746.1 | South China Sea | Arthropoda | Crustacea |
| 808 | Wenling crustacean virus 3 strain WLJQ142924 hypothetical protein 1 and hypothetical protein 2 genes complete co | KX884321.1 | East China Sea | Arthropoda | Crustacea |
| 809 | Bivalvia Picorna-like virus D69 | GWHAPKO01000000 | East China Sea | Mollusca | Bivalvia |
| 810 | Changjiang picorna-like virus 8 strain CJLX30776 hypothetical protein 1 and hypothetical protein 2 genes complete | KX884559.1 | Freshwater | Arthropoda | Crustacea |

| | | | | | |
|---|---|---|---|---|---|
| 811 | Gastropoda Picorna-like virus D10 | GWHAPAT01000000 | East China Sea | Mollusca | Gastropoda |
| 812 | Changping earthworm virus 1 strain CPQY105740 hypothetical protein gene complete cds | KX883890.1 | Terrestrial | Annelida | Clitellata |
| 813 | Shahe narna-like virus 5 strain SHWC0209c20055 RNA-dependent RNA polymerase gene complete cds | KX883555.1 | Freshwater | Arthropoda | Crustacea |
| 814 | Beihai picorna-like virus 5 strain BHNXC41415 hypothetical protein 1 and hypothetical protein 2 genes complete c | KX883398.1 | South China Sea | Sipuncula | Sipuncula |
| 815 | Shahe partiti-like virus 1 strain SHWC0209c27544 RdRp gene complete cds | KX884155.1 | Freshwater | Arthropoda | Crustacea |
| 816 | Shahe partiti-like virus 1 strain SHWCII3004 RdRp gene complete cds | KX884157.1 | Freshwater | Arthropoda | Mix |
| 817 | Hubei toti-like virus 8 strain QTM25279 hypothetical protein gene partial cds | KX882965.1 | Terrestrial | Arthropoda | Insecta |
| 818 | Sanxia water strider virus 8 strain SXSSP2580 hypothetical protein gene complete cds | KX883707.1 | Freshwater | Arthropoda | Insecta |
| 819 | Hubei picorna-like virus 41 strain QTM133099 hypothetical protein gene complete cds | KX883955.1 | Terrestrial | Arthropoda | Insecta |
| 820 | Hubei insect virus 2 strain arthropodmix13736 hypothetical protein 4 gene complete cds | KX884584.1 | Terrestrial | Arthropoda | Mix |
| 821 | Hubei diptera virus 14 strain SCM51203 segment 1 hypothetical protein 1 and hypothetical protein 2 genes complet | KX882856.1 | Terrestrial | Arthropoda | Insecta |
| 822 | Hubei diptera virus 14 strain QTM46268 segment 1 hypothetical protein 1 and hypothetical protein 2 genes comple | KX882851.1 | Terrestrial | Arthropoda | Insecta |
| 823 | Hubei picorna-like virus 17 strain horsefly124179 hypothetical protein 1 and hypothetical protein 2 genes complete | KX883897.1 | Terrestrial | Arthropoda | Insecta |
| 824 | Beihai picorna-like virus 9 strain BHBJDX18814 hypothetical protein 1 and hypothetical protein 2 genes complete | KX883336.1 | South China Sea | Arthropoda | Crustacea |
| 825 | Bivalvia Picorna-like virus D51 | GWHAOZV01000000 | East China Sea | Mollusca | Bivalvia |
| 826 | Gastropoda Picorna-like virus D18 | GWHAPBB01000000 | East China Sea | Mollusca | Gastropoda |
| 827 | Beihai picorna-like virus 30 strain BHNXC41477 hypothetical protein 1 and hypothetical protein 2 genes complete | KX883402.1 | South China Sea | Sipuncula | Sipuncula |
| 828 | Beihai picorna-like virus 38 strain BHZC37430 hypothetical protein 1 gene complete cds and hypothetical protein 2 | KX884495.1 | South China Sea | Mollusca | Bivalvia |
| 829 | Wenzhou picorna-like virus 13 strain WZSBei69459 hypothetical protein gene complete cds | KX884360.1 | East China Sea | Mollusca | Bivalvia |
| 830 | Hubei partiti-like virus 11 strain QTM25188 segment Seg 1 RdRp gene complete cds | KX884126.1 | Terrestrial | Arthropoda | Insecta |
| 831 | Wenzhou narna-like virus 10 strain WZSLuoI82025 RNA-dependent RNA polymerase gene partial cds | KX883624.1 | East China Sea | Mollusca | Gastropoda |
| 832 | Bivalvia Picorna-like virus D26 | GWHAOZA01000000 | East China Sea | Mollusca | Bivalvia |
| 833 | Gastropoda Toli-like virus N1 | GWHAPFG01000000 | South China Sea | Mollusca | Gastropoda |
| 834 | Beihai picorna-like virus 46 strain BHZC37388 hypothetical protein 1 and hypothetical protein 2 genes complete cd | KX884493.1 | South China Sea | Mollusca | Bivalvia |
| 835 | Hubei myriapoda virus 9 strain arthropodmix12082 segment 2 hypothetical protein gene complete cds | KX882771.1 | Terrestrial | Arthropoda | Mix |
| 836 | Hubei myriapoda virus 9 strain ZCM18719 segment 2 hypothetical protein genes complete cds | KX882936.1 | Terrestrial | Arthropoda | Insecta |
| 837 | Hubei myriapoda virus 9 strain WHYCM992 segment 2 hypothetical protein genes complete cds | KX882920.1 | Terrestrial | Arthropoda | Insecta |
| 838 | Hubei myriapoda virus 9 strain WGML401068 segment 2 putative capsid protein and hypothetical protein genes co | KX882899.1 | Terrestrial | Arthropoda | Myriapoda |
| 839 | Hubei myriapoda virus 9 strain WHXS4007 segment 2 hypothetical protein genes complete cds | KX882916.1 | Terrestrial | Arthropoda | Insecta |
| 840 | Wenling partiti-like virus 8 strain WLJQ123769 RdRp gene complete cds | KX884229.1 | East China Sea | Arthropoda | Crustacea |
| 841 | Wenling crustacean virus 1 strain WLJQ101409 RdRp hypothetical protein and putative structural protein genes co | KX883826.1 | East China Sea | Arthropoda | Crustacea |
| 842 | Hexanauplia Picorna-like virus D4 | GWHAPKE01000000 | East China Sea | Arthropoda | Hexanauplia |
| 843 | Shahe isopoda virus 2 strain SHWC0209c12758 hypothetical protein gene complete cds | KX883642.1 | Freshwater | Arthropoda | Crustacea |
| 844 | Shahe isopoda virus 2 strain SHWCII5330 hypothetical protein gene complete cds | KX883672.1 | Freshwater | Arthropoda | Mix |
| 845 | Shahe isopoda virus 2 strain SHWC13604 hypothetical protein gene complete cds | KX883660.1 | Freshwater | Arthropoda | Mix |
| 846 | Beihai barnacle virus 11 strain BHTH10280 capsid protein gene complete cds | KX883037.1 | South China Sea | Arthropoda | Crustacea |
| 847 | Hubei virga-like virus 2 strain QTM25157 hypothetical protein genes complete cds | KX883780.1 | Terrestrial | Arthropoda | Insecta |

| | | | | | |
|---|---|---|---|---|---|
| 848 | Hubei virga-like virus 2 strain mosHB235832 RdRp and putative coat protein genes complete cds | KX883772.1 | Terrestrial | Arthropoda | Insecta |
| 849 | Beihai barnacle virus 12 strain BHTH16091 segment Seg 1 RdRp gene complete cds | KX884067.1 | South China Sea | Arthropoda | Crustacea |
| 850 | Beihai barnacle virus 3 strain BHTH16145 replicase and hypothetical protein genes complete cds | KX883749.1 | South China Sea | Arthropoda | Crustacea |
| 851 | Beihai barnacle virus 13 strain BHTH16173 segment Seg 1 RdRp gene complete cds | KX884069.1 | South China Sea | Arthropoda | Crustacea |
| 852 | Xinzhou nematode virus 3 strain XZSJSC65579 hypothetical protein gene complete cds | KX884005.1 | Terrestrial | Nematoda | Mix |
| 853 | Bivalvia Picorna-like virus N33 | GWHAPJQ01000000 | South China Sea | Mollusca | Bivalvia |
| 854 | Beihai picorna-like virus 56 strain BWBFG40791 hypothetical protein gene complete cds | KX884529.1 | South China Sea | Mollusca | Bivalvia |
| 855 | Wenling partiti-like virus 6 strain WLJQ47818 RdRp gene complete cds | KX884232.1 | East China Sea | Arthropoda | Crustacea |
| 856 | Beihai picorna-like virus 57 strain BHJP51916 hypothetical protein gene complete cds | KX883380.1 | South China Sea | Echinodermata | Echinozoa |
| 857 | Beihai picorna-like virus 57 strain BHHQ57630 hypothetical protein gene complete cds | KX883356.1 | South China Sea | Tunicata | Ascidiacea |
| 858 | Beihai barnacle virus 13 strain BHTH16173 segment Seg 2 hypothetical protein gene complete cds | KX884070.1 | South China Sea | Arthropoda | Crustacea |
| 859 | Changjiang picorna-like virus 1 strain CJLX30149 hypothetical protein gene complete cds | KX884546.1 | Freshwater | Arthropoda | Crustacea |
| 860 | Beihai picorna-like virus 75 strain BHJP52955 hypothetical protein 1 and hypothetical protein 2 genes complete cds | KX883381.1 | South China Sea | Echinodermata | Echinozoa |
| 861 | Changjiang picorna-like virus 9 strain CJLX26226 hypothetical protein 1 and hypothetical protein 2 genes complete | KX884541.1 | Freshwater | Arthropoda | Crustacea |
| 862 | Hubei partiti-like virus 19 strain SCM172380 RdRp gene complete cds | KX884147.1 | Terrestrial | Arthropoda | Insecta |
| 863 | Wenling partiti-like virus 13 strain WLJQ101202 RdRp gene complete cds | KX884236.1 | East China Sea | Arthropoda | Crustacea |
| 864 | Bivalvia Picorna-like virus H2 | GWHAPIY01000000 | Yellow China Sea | Mollusca | Bivalvia |
| 865 | Yongjia Tick Virus 2 strain YJ1-2 nucleocapsid (N) putative phosphoprotein (ORF2) matrix protein (M) glycoprote | KM817662.1 | Terrestrial | Arthropoda | Chelicerata |
| 866 | Wuhan Mosquito Virus 3 strain XZ2-7 PB1 (PB1) gene complete cds | KM817622.1 | Terrestrial | Arthropoda | Insecta |
| 867 | Sanxia Water Strider Virus 3 strain SXSSP01 PB1 (PB1) gene complete cds | KM817617.1 | Freshwater | Arthropoda | Insecta |
| 868 | Hubei orthoptera virus 6 strain ZCM21154 segment 1 polymerase PB2 gene complete cds | KX883885.1 | Terrestrial | Arthropoda | Insecta |
| 869 | Beihai rhabdo-like virus 4 strain BHJP58499 putative nucleoprotein hypothetical protein and RNA-dependent RNA | KX884406.1 | South China Sea | Echinodermata | Echinozoa |
| 870 | Wenzhou partiti-like virus 1 strain WZRBX37312 RdRp gene complete cds | KX884237.1 | East China Sea | Arthropoda | Crustacea |
| 871 | Beihai rhabdo-like virus 6 strain BHJJX49420 putative nucleoprotein hypothetical protein putative glycoprotein anc | KX884405.1 | South China Sea | Arthropoda | Crustacea |
| 872 | Wuhan Mosquito Virus 5 strain XC1-7 PB1 (PB1) gene complete cds | KM817624.1 | Terrestrial | Arthropoda | Insecta |
| 873 | Beihai picorna-like virus 110 strain BHXG26677 hypothetical protein gene complete cds | KX884470.1 | South China Sea | Arthropoda | Crustacea |
| 874 | Hubei rhabdo-like virus 4 strain arthropodmix13990 hypothetical protein 1 hypothetical protein 2 and RNA-depend | KX884403.1 | Terrestrial | Arthropoda | Mix |
| 875 | Wuhan Mosquito Virus 4 strain XC3-4 segment PA polymerase PA gene complete cds | KX883866.1 | Terrestrial | Arthropoda | Insecta |
| 876 | Wuhan Mosquito Virus 4 strain 3mos5038 segment PA polymerase PA gene complete cds | KX883841.1 | Terrestrial | Arthropoda | Insecta |
| 877 | Shahe qinvirus-like virus 1 strain SHWC0209c11359 hypothetical protein gene complete cds | KX883995.1 | Freshwater | Arthropoda | Crustacea |
| 878 | Wenzhou Tick Virus strain TS1-2 RNA-dependent RNA polymerase (L) gene complete cds | KM817685.1 | Terrestrial | Arthropoda | Chelicerata |
| 879 | Wuhan Mosquito Virus 2 strain QN2-7 nucleopasid protein (N) gene complete cds | KM817759.1 | Terrestrial | Arthropoda | Insecta |
| 880 | Wuhan Millipede Virus 2 strain WHWG03 nucleopasid protein (N) gene complete cds | KM817757.2 | Terrestrial | Arthropoda | Insecta |
| 881 | Beihai shrimp virus 3 strain BHWZXX15649 RNA-dependent RNA polymerase gene complete cds | KX884751.1 | South China Sea | Arthropoda | Crustacea |
| 882 | Beihai shrimp virus 3 strain BHBJDX18821 RNA-dependent RNA polymerase gene complete cds | KX884737.1 | South China Sea | Arthropoda | Crustacea |
| 883 | Changping earthworm virus 2 strain CPQY18140 segment 4 putative glycoprotein 2 gene complete cds | KX883852.1 | Terrestrial | Annelida | Clitellata |
| 884 | Changping Tick Virus 3 strain CP1-3 complete genome | KM817595.1 | Terrestrial | Arthropoda | Chelicerata |

| | | | | | |
|---|---|---|---|---|---|
| 885 | Lihan Tick Virus strain LH-1 nucleopasid protein (N) gene complete cds | KM817736.1 | Terrestrial | Arthropoda | Chelicerata |
| 886 | Wenling partiti-like virus 5 strain WLJQ59225 RdRp gene complete cds | KX884233.1 | East China Sea | Arthropoda | Crustacea |
| 887 | Wuhan Mosquito Virus 4 strain XC3-4 segment PB2 polymerase PB2 gene complete cds | KX883865.1 | Terrestrial | Arthropoda | Insecta |
| 888 | Wuhan Mosquito Virus 4 strain 3mos5038 segment PB2 polymerase PB2 gene complete cds | KX883840.1 | Terrestrial | Arthropoda | Insecta |
| 889 | Beihai picorna-like virus 83 strain BHWZXX15514 hypothetical protein 1 hypothetical protein 2 and hypothetical p | KX883427.1 | South China Sea | Arthropoda | Crustacea |
| 890 | Shuangao permutotetra-like virus 1 strain insectZJ82042 RdRp gene partial cds | KX883439.1 | Terrestrial | Arthropoda | Insecta |
| 891 | Bivalvia Picorna-like virus D9 | GWHAOXY0100000( | East China Sea | Mollusca | Bivalvia |
| 892 | Gastropoda Picorna-like virus D6 | GWHAPAS01000000 | East China Sea | Mollusca | Gastropoda |
| 893 | Bivalvia Picorna-like virus N20 | GWHAPIX01000000 | South China Sea | Mollusca | Bivalvia |
| 894 | Bivalvia Picorna-like virus H15 | GWHAPJJ01000000 | Yellow China Sea | Mollusca | Bivalvia |
| 895 | Bivalvia Picorna-like virus N53 | GWHAPLS01000000 | South China Sea | Mollusca | Bivalvia |
| 896 | Wuhan insect virus 12 strain arthropodmix13526 hypothetical protein gene complete cds | KX883288.1 | Terrestrial | Arthropoda | Mix |
| 897 | Wuhan insect virus 12 strain WHZM10126 hypothetical protein gene complete cds | KX884308.1 | Terrestrial | Arthropoda | Insecta |
| 898 | Bivalvia Picorna-like virus D42 | GWHAOZI01000000 | East China Sea | Mollusca | Bivalvia |
| 899 | Bivalvia Picorna-like virus N47 | GWHAPEE01000000 | South China Sea | Mollusca | Bivalvia |
| 900 | Bivalvia Picorna-like virus N50 | GWHAPLO01000000 | South China Sea | Mollusca | Bivalvia |
| 901 | Bivalvia Picorna-like virus D44 | GWHAOZN01000000 | East China Sea | Mollusca | Bivalvia |
| 902 | Bivalvia Picorna-like virus N56 | GWHAPLQ01000000 | South China Sea | Mollusca | Bivalvia |
| 903 | Bivalvia Picorna-like virus N51 | GWHAPLP01000000 | South China Sea | Mollusca | Bivalvia |
| 904 | Hubei picorna-like virus 18 strain QTM26408 hypothetical protein 1 and hypothetical protein 2 genes complete cds | KX883935.1 | Terrestrial | Arthropoda | Insecta |
| 905 | Hubei picorna-like virus 77 strain WHCC115343 hypothetical protein 1 hypothetical protein 2 hypothetical protein | KX884269.1 | Terrestrial | Arthropoda | Insecta |
| 906 | Gastropoda Picorna-like virus D3 | GWHAPJV01000000 | East China Sea | Mollusca | Gastropoda |
| 907 | Hubei picorna-like virus 79 strain WHCCII12350 hypothetical protein 1 and hypothetical protein 2 genes complete | KX884275.1 | Terrestrial | Arthropoda | Insecta |
| 908 | Shahe picorna-like virus 13 strain SHWC0209c12284 hypothetical protein gene complete cds | KX883649.1 | Freshwater | Arthropoda | Crustacea |
| 909 | Shahe picorna-like virus 13 strain SHWC13556 hypothetical protein gene complete cds | KX883657.1 | Freshwater | Arthropoda | Mix |
| 910 | Xinzhou partiti-like virus 1 strain XZSJSC65291 segment Seg 1 RdRp gene complete cds | KX884238.1 | Terrestrial | Nematoda | Mix |
| 911 | Raphanus sativus cryptic virus 1 strain CJLX6222 RdRp gene partial cds | KX884089.1 | Freshwater | Arthropoda | Crustacea |
| 912 | Beihai picorna-like virus 118 strain BHHK118641 hypothetical protein 1 hypothetical protein 2 and hypothetical pr | KX883342.1 | South China Sea | Cnidaria | Anthozoa |
| 913 | Hubei partiti-like virus 26 strain QTM27118 RdRp gene complete cds | KX884133.1 | Terrestrial | Arthropoda | Insecta |
| 914 | Hubei diptera virus 15 strain SCM95219 hypothetical protein 1 and hypothetical protein 2 genes complete cds | KX883150.1 | Terrestrial | Arthropoda | Insecta |
| 915 | Beihai picorna-like virus 117 strain BHJP63029 hypothetical protein 1 hypothetical protein 2 and hypothetical prote | KX883384.1 | South China Sea | Echinodermata | Echinozoa |
| 916 | Beihai picorna-like virus 117 strain BHHQ50349 hypothetical protein 1 and hypothetical protein 2 genes complete ( | KX883354.1 | South China Sea | Tunicata | Ascidiacea |
| 917 | Beihai picorna-like virus 81 strain HOU149705 hypothetical protein 1 and hypothetical protein 2 genes complete cd | KX883904.1 | South China Sea | Arthropoda | Chelicerata |
| 918 | Bivalvia Picorna-like virus N54 | GWHAPEB01000000 | South China Sea | Mollusca | Bivalvia |
| 919 | Bivalvia Toli-like virus D2 | GWHAPKS01000000 | East China Sea | Mollusca | Bivalvia |
| 920 | Bivalvia Amarillo-like virus N2 | GWHAPJM01000000 | South China Sea | Mollusca | Bivalvia |
| 921 | Wuhan Millipede Virus 1 strain WHWG04 nucleoprotein gene complete cds | KX884851.1 | Terrestrial | Arthropoda | Myriapoda |

| | | | | | |
|---|---|---|---|---|---|
| 922 | Bivalvia Picorna-like virus N15 | GWHAPGJ01000000 | South China Sea | Mollusca | Bivalvia |
| 923 | Hubei noda-like virus 8 strain WHLC5312 capsid protein gene complete cds | KX883238.1 | Terrestrial | Arthropoda | Insecta |
| 924 | Crustacea Picorna-like virus D7 | GWHAPBU01000000 | East China Sea | Arthropoda | Crustacea |
| 925 | Wenzhou picorna-like virus 50 strain WZSLuoI83448 hypothetical protein 1 gene complete cds and hypothetical pr | KX884367.1 | East China Sea | Mollusca | Gastropoda |
| 926 | Wenling crustacean virus 13 strain WLJQ104251 RNA-dependent RNA polymerase gene complete cds | KX884453.1 | East China Sea | Arthropoda | Crustacea |
| 927 | Beihai picorna-like virus 125 strain BHZY54174 hypothetical protein gene complete cds | KX884504.1 | South China Sea | Mollusca | Cephalopoda |
| 928 | Hubei lepidoptera virus 4 strain LCM63323 hypothetical protein 3 gene complete cds | KX884631.1 | Terrestrial | Arthropoda | Insecta |
| 929 | Bivalvia Bunya-like virus N3 | GWHAPHO01000000 | South China Sea | Mollusca | Bivalvia |
| 930 | Hubei picorna-like virus 49 strain WGML112833 hypothetical protein gene complete cds | KX883732.1 | Terrestrial | Arthropoda | Myriapoda |
| 931 | Wenzhou shrimp virus 7 strain WZSBei42084 hypothetical protein 1 and hypothetical protein 2 genes complete cds | KX884352.1 | East China Sea | Mollusca | Bivalvia |
| 932 | Wenzhou shrimp virus 7 strain beimix73547 hypothetical protein 1 and hypothetical protein 2 genes complete cds | KX883298.1 | East China Sea | Mollusca | Bivalvia |
| 933 | Wenzhou shrimp virus 7 strain shrimp14434 hypothetical protein 1 and hypothetical protein 2 genes complete cds | KX883980.1 | East China Sea | Arthropoda | Crustacea |
| 934 | Crustacea Durna-like virus N2 | GWHAPFJ01000000 | South China Sea | Arthropoda | Crustacea |
| 935 | Crustacea Ghabri-like virus H2 | GWHAPGW0100000( | Yellow China Sea | Arthropoda | Crustacea |
| 936 | Wuhan house centipede virus 1 strain WHCCII11885 hypothetical protein genes complete cds | KX883813.1 | Terrestrial | Arthropoda | Insecta |
| 937 | Wuhan house centipede virus 1 strain WHYY23932 RdRp and hypothetical protein genes complete cds | KX883822.1 | Terrestrial | Arthropoda | Myriapoda |
| 938 | Wuhan house centipede virus 1 strain spider119908 hypothetical protein genes complete cds | KX883798.1 | Terrestrial | Arthropoda | Chelicerata |
| 939 | Wuhan house centipede virus 1 strain arthropodmix13957 hypothetical protein 1 hypothetical protein 2 hypothetical | KX883740.1 | Terrestrial | Arthropoda | Mix |
| 940 | Hubei earwig virus 2 strain WGML118279 RdRp gene complete cds | KX884185.1 | Terrestrial | Arthropoda | Myriapoda |
| 941 | Hubei negev-like virus 1 strain QTM162107 hypothetical protein gene complete cds and hypothetical protein gene p | KX883786.1 | Terrestrial | Arthropoda | Insecta |
| 942 | Hubei virga-like virus 19 strain SCM95808 hypothetical protein gene complete cds and hypothetical protein gene pa | KX883793.1 | Terrestrial | Arthropoda | Insecta |
| 943 | Wenling partiti-like virus 1 strain WLJQ97111 RdRp gene complete cds | KX884235.1 | East China Sea | Arthropoda | Crustacea |
| 944 | Hubei tombus-like virus 28 strain SSZZ1762 hypothetical protein 1 capsid protein and hypothetical protein 3 genes | KX883169.1 | Freshwater | Arthropoda | Chelicerata |
| 945 | Bivalvia Picorna-like virus N58 | GWHAPEC01000000 | South China Sea | Mollusca | Bivalvia |
| 946 | Bivalvia Picorna-like virus D10 | GWHAOYA0100000( | East China Sea | Mollusca | Bivalvia |
| 947 | Hubei toti-like virus 21 strain CC64629 hypothetical protein 1 and hypothetical protein 2 genes complete cds | KX882955.1 | Terrestrial | Arthropoda | Insecta |
| 948 | Hubei toti-like virus 23 strain CC64311 hypothetical protein gene partial cds | KX882953.1 | Terrestrial | Arthropoda | Insecta |
| 949 | Hubei partiti-like virus 1 strain WHCC258759 RdRp gene complete cds | KX884191.1 | Terrestrial | Arthropoda | Insecta |
| 950 | Wuhan insect virus 31 strain WHZM10182 hypothetical protein 1 and hypothetical protein 2 genes complete cds | KX882989.1 | Terrestrial | Arthropoda | Insecta |
| 951 | Wenling chuvirus-like virus 2 strain WLJQ104274 RNA-dependent RNA polymerase gene complete cds | KX884455.1 | East China Sea | Arthropoda | Crustacea |
| 952 | Hubei toti-like virus 3 strain ZL17544 hypothetical protein gene partial cds | KX882999.1 | Terrestrial | Arthropoda | Insecta |
| 953 | Beihai partiti-like virus 10 strain YYSZX26933 RdRp gene complete cds | KX884242.1 | South China Sea | Arthropoda | Crustacea |
| 954 | Hubei narna-like virus 24 strain SCM50734 RNA-dependent RNA polymerase gene complete cds | KX883547.1 | Terrestrial | Arthropoda | Insecta |
| 955 | Bivalvia Picorna-like virus D82 | GWHAPIQ01000000 | East China Sea | Mollusca | Bivalvia |
| 956 | Hubei sobemo-like virus 42 strain QTM26192 hypothetical protein 1 and hypothetical protein 2 genes complete cds | KX882843.1 | Terrestrial | Arthropoda | Insecta |
| 957 | Wuhan house centipede virus 5 strain WHYY23902 segment 1 hypothetical protein 1 and RdRp genes complete cds | KX882921.1 | Terrestrial | Arthropoda | Myriapoda |
| 958 | Beihai sesarmid crab virus 7 strain SCJXSX14567 hypothetical protein gene complete cds | KX882970.1 | South China Sea | Arthropoda | Crustacea |

| | | | | | |
|---|---|---|---|---|---|
| 959 | Bivalvia Picorna-like virus N38 | GWHAPHN01000000 | South China Sea | Mollusca | Bivalvia |
| 960 | Changjiang zhaovirus-like virus 1 strain CJLX30599 polyprotein gene complete cds | KX884041.1 | Freshwater | Arthropoda | Crustacea |
| 961 | Hubei tetragnatha maxillosa virus 9 strain QTM27093 hypothetical protein 4 gene complete cds | KX884679.1 | Terrestrial | Arthropoda | Insecta |
| 962 | Hubei tetragnatha maxillosa virus 9 strain SSZZ3444 hypothetical protein 5 gene complete cds | KX884708.1 | Freshwater | Arthropoda | Chelicerata |
| 963 | Hubei tetragnatha maxillosa virus 9 strain arthropodmix14049 hypothetical protein 4 gene complete cds | KX884602.1 | Terrestrial | Arthropoda | Mix |
| 964 | Hubei tetragnatha maxillosa virus 9 strain SSZZ3444 hypothetical protein 4 gene complete cds | KX884707.1 | Freshwater | Arthropoda | Chelicerata |
| 965 | Changjiang picorna-like virus 14 strain CJLX29953 hypothetical protein 1 and hypothetical protein 2 genes comple | KX884544.1 | Freshwater | Arthropoda | Crustacea |
| 966 | Hubei tetragnatha maxillosa virus 3 strain SSZZ2994 hypothetical protein gene complete cds | KX883696.1 | Freshwater | Arthropoda | Chelicerata |
| 967 | Bivalvia Sobeli-like virus H2 | GWHAPCR01000000 | Yellow China Sea | Mollusca | Bivalvia |
| 968 | Hubei bunya-like virus 13 strain QTM27285 RNA-dependent RNA polymerase gene complete cds | KX884784.1 | Terrestrial | Arthropoda | Insecta |
| 969 | Wuhan flea virus strain WHZM segment 1 complete sequence | KR902713.1 | Terrestrial | Arthropoda | Insecta |
| 970 | Bean yellow mosaic virus strain SCM51513 polyprotein gene partial cds | KX884569.1 | Terrestrial | Arthropoda | Insecta |
| 971 | Hubei Poty-like virus 1 strain QTM27268 polyprotein gene partial cds | KX884567.1 | Terrestrial | Arthropoda | Insecta |
| 972 | Wuhan aphid virus 2 strain WHYC-2 segment 3 complete sequence | KR902727.1 | Terrestrial | Arthropoda | Insecta |
| 973 | Sanxia sobemo-like virus 2 strain SXSSP2316 hypothetical protein 1 and hypothetical protein 2 genes complete cds | KX882886.1 | Freshwater | Arthropoda | Insecta |
| 974 | Crustacea Picorna-like virus D8 | GWHAPBV01000000 | East China Sea | Arthropoda | Crustacea |
| 975 | Bivalvia Toli-like virus N1 | GWHAPDW0100000( | South China Sea | Mollusca | Bivalvia |
| 976 | Beihai hepe-like virus 7 strain BHBJDX18438 hypothetical protein genes complete cds | KX883743.1 | South China Sea | Arthropoda | Crustacea |
| 977 | Wuhan poty-like virus 1 strain WHWN51517 polyprotein gene complete cds | KX884573.1 | Freshwater | Mollusca | Gastropoda |
| 978 | Hubei bunya-like virus 11 strain WGML134260 RNA-dependent RNA polymerase gene partial cds | KX884845.1 | Terrestrial | Arthropoda | Myriapoda |
| 979 | Hubei bunya-like virus 8 strain SCM48138 putative nucleoprotein gene complete cds | KX884803.1 | Terrestrial | Arthropoda | Insecta |
| 980 | Hubei bunya-like virus 2 strain QTM26385 RNA-dependent RNA polymerase gene complete cds | KX884780.1 | Terrestrial | Arthropoda | Insecta |
| 981 | Bivalvia Picorna-like virus D74 | GWHAPIC01000000 | East China Sea | Mollusca | Bivalvia |
| 982 | Hubei virga-like virus 17 strain QTM23703 RdRp and hypothetical protein genes complete cds | KX883779.1 | Terrestrial | Arthropoda | Insecta |
| 983 | Wenling crustacean virus 4 strain WLJQ79834 hypothetical protein gene complete cds | KX884322.1 | East China Sea | Arthropoda | Crustacea |
| 984 | Hubei virga-like virus 12 strain SCM49209 hypothetical protein genes complete cds | KX883787.1 | Terrestrial | Arthropoda | Insecta |
| 985 | Beihai picorna-like virus 6 strain BHZC37373 hypothetical protein 1 and hypothetical protein 2 genes complete cds | KX884492.1 | South China Sea | Mollusca | Bivalvia |
| 986 | Beihai picorna-like virus 6 strain BHNXC41284 hypothetical protein 1 and hypothetical protein 2 genes complete c | KX883393.1 | South China Sea | Sipuncula | Sipuncula |
| 987 | Wenzhou Tick Virus strain TS1-2 glycoprotein precursor (G) gene complete cds | KM817718.1 | Terrestrial | Arthropoda | Chelicerata |
| 988 | Beihai bunya-like virus 5 strain BHJP59844 RNA-dependent RNA polymerase gene complete cds | KX884744.1 | South China Sea | Echinodermata | Echinozoa |
| 989 | Wenzhou bunya-like virus 1 strain WZSLuoI85881 putative glycoprotein gene complete cds | KX884866.1 | East China Sea | Mollusca | Gastropoda |
| 990 | Jingmen birna-like virus strain JMYJCC68862 RNA-dependent RNA polymerase gene complete cds | KX883000.1 | Terrestrial | Nematoda | Chromadorea |
| 991 | Yongjia Tick Virus 1 strain YJ1-1 RNA-dependent RNA polymerase (L) gene complete cds | KM817704.1 | Terrestrial | Arthropoda | Chelicerata |
| 992 | Bivalvia Picorna-like virus D4 | GWHAPEN01000000 | East China Sea | Mollusca | Bivalvia |
| 993 | Shahe picorna-like virus 14 strain SHWC13617 hypothetical protein gene complete cds | KX883664.1 | Freshwater | Arthropoda | Mix |
| 994 | Shahe picorna-like virus 14 strain SHWC0209c12767 hypothetical protein gene complete cds | KX883647.1 | Freshwater | Arthropoda | Crustacea |
| 995 | Sanxia partiti-like virus 1 strain SXSSP515 RdRp gene complete cds | KX884179.1 | Freshwater | Arthropoda | Insecta |

| | | | | | |
|---|---|---|---|---|---|
| 996 | Shahe picorna-like virus 3 strain SHWC13623 hypothetical protein 1 and hypothetical protein 2 genes complete cds | KX883667.1 | Freshwater | Arthropoda | Mix |
| 997 | Hubei sobemo-like virus 21 strain CC64469 segment 2 hypothetical protein gene partial cds | KX882813.1 | Terrestrial | Arthropoda | Insecta |
| 998 | Beihai picorna-like virus 80 strain BHBei50452 hypothetical protein 1 and hypothetical protein 2 genes complete cc | KX883304.1 | South China Sea | Mollusca | Bivalvia |
| 999 | Beihai picorna-like virus 80 strain BHZC36213 hypothetical protein 2 genes complete cds | KX884478.1 | South China Sea | Mollusca | Bivalvia |
| 1000 | Beihai partiti-like virus 8 strain HWRTX13334 RdRp gene complete cds | KX884099.1 | South China Sea | Arthropoda | Crustacea |
| 1001 | Beihai picorna-like virus 58 strain BHBei77093 hypothetical protein 1 and hypothetical protein 2 genes complete cc | KX883320.1 | South China Sea | Mollusca | Bivalvia |
| 1002 | Hubei odonate virus 9 strain QTM74767 putative glycoprotein gene complete cds | KX884787.1 | Terrestrial | Arthropoda | Insecta |
| 1003 | Hubei picorna-like virus 1 strain WHSF7464 hypothetical protein 1 and hypothetical protein 2 genes complete cds | KX884285.1 | Terrestrial | Arthropoda | Crustacea |
| 1004 | Hubei odonate virus 8 strain QTM19232 putative nucleoprotein gene complete cds | KX884777.1 | Terrestrial | Arthropoda | Insecta |
| 1005 | Beihai picorna-like virus 105 strain BHHK79817 hypothetical protein 1 and hypothetical protein 2 genes complete | KX883341.1 | South China Sea | Cnidaria | Anthozoa |
| 1006 | Wuhan cricket virus strain WHXS-1 segment 2 complete sequence | KR902710.1 | Terrestrial | Arthropoda | Insecta |
| 1007 | Hubei picorna-like virus 68 strain WHWG56209 hypothetical protein gene complete cds | KX884292.1 | Terrestrial | Arthropoda | Myriapoda |
| 1008 | Changping earthworm virus 2 strain CPQY18140 segment 3 putative glycoprotein 1 gene complete cds | KX883851.1 | Terrestrial | Annelida | Clitellata |
| 1009 | Hubei partiti-like virus 45 strain WHYY15874 RdRp gene complete cds | KX884215.1 | Terrestrial | Arthropoda | Myriapoda |
| 1010 | Beihai picorna-like virus 23 strain BWBFG39274 hypothetical protein and hypothetical protein 2 genes complete cc | KX884526.1 | South China Sea | Mollusca | Bivalvia |
| 1011 | Hubei partiti-like virus 36 strain QTM3514 RdRp gene complete cds | KX884136.1 | Terrestrial | Arthropoda | Insecta |
| 1012 | Beihai picorna-like virus 55 strain BHZC37036 hypothetical protein gene complete cds | KX884482.1 | South China Sea | Mollusca | Bivalvia |
| 1013 | Beihai picorna-like virus 55 strain BHBei68636 hypothetical protein gene complete cds | KX883306.1 | South China Sea | Mollusca | Bivalvia |
| 1014 | Hubei leech virus 3 strain SZmix63518 hypothetical protein 1 and hypothetical protein 2 genes complete cds | KX883727.1 | Freshwater | Annelida | Clitellata |
| 1015 | Hubei partiti-like virus 33 strain QCM951148 RdRp gene complete cds | KX884109.1 | Terrestrial | Arthropoda | Insecta |
| 1016 | Hubei tetragnatha maxillosa virus 9 strain arthropodmix14049 hypothetical protein 3 gene complete cds | KX884601.1 | Terrestrial | Arthropoda | Mix |
| 1017 | Hubei tetragnatha maxillosa virus 9 strain SSZZ3444 hypothetical protein 3 gene complete cds | KX884706.1 | Freshwater | Arthropoda | Chelicerata |
| 1018 | Hubei tetragnatha maxillosa virus 9 strain QTM27093 hypothetical protein 3 gene complete cds | KX884678.1 | Terrestrial | Arthropoda | Insecta |
| 1019 | Beihai picorna-like virus 60 strain BHBei55726 hypothetical protein gene complete cds | KX883305.1 | South China Sea | Mollusca | Bivalvia |
| 1020 | Beihai picorna-like virus 61 strain BHBei76813 hypothetical protein gene complete cds | KX883313.1 | South China Sea | Mollusca | Bivalvia |
| 1021 | Beihai picorna-like virus 59 strain BHZC37451 hypothetical protein gene complete cds | KX884500.1 | South China Sea | Mollusca | Bivalvia |
| 1022 | Beihai picorna-like virus 59 strain BHBei77099 hypothetical protein gene complete cds | KX883321.1 | South China Sea | Mollusca | Bivalvia |
| 1023 | Hubei tetragnatha maxillosa virus 9 strain QTM27093 hypothetical protein 2 gene partial cds | KX884677.1 | Terrestrial | Arthropoda | Insecta |
| 1024 | Hubei tetragnatha maxillosa virus 9 strain SSZZ3444 hypothetical protein 1 gene complete cds | KX884704.1 | Freshwater | Arthropoda | Chelicerata |
| 1025 | Hubei tetragnatha maxillosa virus 9 strain arthropodmix14049 hypothetical protein 2 gene complete cds | KX884600.1 | Terrestrial | Arthropoda | Mix |
| 1026 | Xingshan nematode virus 2 strain XSNXC13152 RdRp and hypothetical protein genes complete cds | KX883836.1 | Terrestrial | Nematoda | Chromadorea |
| 1027 | Changjiang picorna-like virus 3 strain CJLX28786 hypothetical protein 1 and hypothetical protein 2 genes complete | KX884542.1 | Freshwater | Arthropoda | Crustacea |
| 1028 | Sanxia water strider virus 12 strain SXSSP2564 hypothetical protein 1 hypothetical protein 2 and hypothetical prote | KX882892.1 | Freshwater | Arthropoda | Insecta |
| 1029 | Hubei odonate virus 15 strain QTM133059 hypothetical protein 1 gene complete cds | KX884665.1 | Terrestrial | Arthropoda | Insecta |
| 1030 | Beihai picorna-like virus 123 strain BHCL81476 hypothetical protein gene complete cds | KX883340.1 | South China Sea | Mollusca | Gastropoda |
| 1031 | Hubei blood fluke virus 3 strain JSC69362 hypothetical protein 3 gene complete cds | KX884611.1 | Freshwater | Platyhelminthes | Trematoda |
| 1032 | Hubei blood fluke virus 3 strain RBXXC337523 hypothetical protein 3 gene complete cds | KX884690.1 | Freshwater | Platyhelminthes | Trematoda |

| | | | | | |
|---|---|---|---|---|---|
| 1033 | Hubei picobirna-like virus 1 strain WHSFII41391 RdRp gene complete cds | KX884200.1 | Terrestrial | Arthropoda | Crustacea |
| 1034 | Hubei odonate virus 3 strain QTM25153 hypothetical protein gene complete cds | KX883932.1 | Terrestrial | Arthropoda | Insecta |
| 1035 | Wenzhou picorna-like virus 21 strain WZSBei23778 hypothetical protein 1 and hypothetical protein 2 genes comple | KX884351.1 | East China Sea | Mollusca | Bivalvia |
| 1036 | Shahe picorna-like virus 11 strain SHWC13613 hypothetical protein 1 and hypothetical protein 2 genes complete cc | KX883663.1 | Freshwater | Arthropoda | Mix |
| 1037 | Bivalvia Picorna-like virus D16 | GWHAOYI01000000 | East China Sea | Mollusca | Bivalvia |
| 1038 | Crustacea Picorna-like virus D5 | GWHAPBO01000000 | East China Sea | Arthropoda | Crustacea |
| 1039 | Crustacea Picorna-like virus D6 | GWHAPBR01000000 | East China Sea | Arthropoda | Crustacea |
| 1040 | Wenzhou shrimp virus 8 strain shrimp14543 hypothetical protein gene complete cds | KX883984.1 | East China Sea | Arthropoda | Crustacea |
| 1041 | Crustacea Picorna-like virus N9 | GWHAPIU01000000 | South China Sea | Arthropoda | Crustacea |
| 1042 | Huangpi Tick Virus 3 strain H124-2 nucleocapsid (N) putative phosphoprotein (ORF2) putative matrix protein (OR | KM817630.1 | Terrestrial | Arthropoda | Chelicerata |
| 1043 | Hubei picorna-like virus 7 strain WHSF7327 hypothetical protein gene complete cds | KX884284.1 | Terrestrial | Arthropoda | Crustacea |
| 1044 | Hubei picorna-like virus 7 strain WHCC111241 hypothetical protein gene complete cds | KX884268.1 | Terrestrial | Arthropoda | Insecta |
| 1045 | Changjiang picorna-like virus 16 strain CJLX20820 hypothetical protein gene complete cds | KX884538.1 | Freshwater | Arthropoda | Crustacea |
| 1046 | Gastropoda Picorna-like virus D22 | GWHAPBG01000000 | East China Sea | Mollusca | Gastropoda |
| 1047 | Bivalvia Picorna-like virus D81 | GWHAPEU01000000 | East China Sea | Mollusca | Bivalvia |
| 1048 | Wenzhou shrimp virus 3 strain shrimp12824 replicase hypothetical protein putative structural protein and hypotheti | KX883794.1 | East China Sea | Arthropoda | Crustacea |
| 1049 | Bivalvia Picorna-like virus N2 | GWHAPEW01000000 | South China Sea | Mollusca | Bivalvia |
| 1050 | Bivalvia Picorna-like virus D24 | GWHAOYW0100000( | East China Sea | Mollusca | Bivalvia |
| 1051 | Wenling crustacean virus 15 strain WLJQ91782 RNA-dependent RNA polymrease putative glycoprotein and hypot | KX884458.1 | East China Sea | Arthropoda | Crustacea |
| 1052 | Beihai mantis shrimp virus 6 strain BHXG24422 capsid protein gene complete cds | KX883057.1 | South China Sea | Arthropoda | Crustacea |
| 1053 | Wenzhou picorna-like virus 49 strain WZFSL84024 hypothetical protein gene complete cds | KX884330.1 | Freshwater | Mollusca | Gastropoda |
| 1054 | Beihai noda-like virus 11 strain BHJJX49550 capsid protein gene complete cds | KX883033.1 | South China Sea | Arthropoda | Crustacea |
| 1055 | Changping Tick Virus 1 strain CP1-2 nucleopasid protein (N) gene partial cds | KM817732.1 | Terrestrial | Arthropoda | Chelicerata |
| 1056 | Bole tick virus 4 strain BLP-1 polyprotein gene complete cds | KR902736.1 | Terrestrial | Arthropoda | Chelicerata |
| 1057 | Wenzhou Tick Virus strain TS1-2 nucleopasid protein (N) gene complete cds | KM817745.1 | Terrestrial | Arthropoda | Chelicerata |
| 1058 | Changping Tick Virus 2 strain CP1-4 complete genome | KM817594.1 | Terrestrial | Arthropoda | Chelicerata |
| 1059 | Beihai barnacle virus 8 strain BHTH14652 hypothetical protein 1 hypothetical protein 2 putative glycoprotein and F | KX884410.1 | South China Sea | Arthropoda | Crustacea |
| 1060 | Wuhan cricket virus 2 strain ZCM19297 segment Seg 3 putative capsid protein gene complete cds | KX884245.1 | Terrestrial | Arthropoda | Insecta |
| 1061 | Wuhan cricket virus 2 strain WHXS3745 segment Seg 3 putative capsid protein gene complete cds | KX884210.1 | Terrestrial | Arthropoda | Insecta |
| 1062 | Beihai sphaeromadae virus 3 strain BHTSS17005 capsid protein gene complete cds | KX883040.1 | South China Sea | Arthropoda | Crustacea |
| 1063 | Beihai sphaeromadae virus 3 strain BHHK126836 capsid protein gene complete cds | KX883028.1 | South China Sea | Cnidaria | Anthozoa |
| 1064 | Bivalvia Picorna-like virus D31 | GWHAOZD01000000 | East China Sea | Mollusca | Bivalvia |
| 1065 | Beihai picorna-like virus 53 strain BHNXC41443 hypothetical protein 1 and hypothetical protein 2 genes complete | KX883400.1 | South China Sea | Sipuncula | Sipuncula |
| 1066 | Wenling partiti-like virus 11 strain WLJQ9539 RdRp gene complete cds | KX884225.1 | East China Sea | Arthropoda | Crustacea |
| 1067 | Hubei picorna-like virus 65 strain WHCC118065 hypothetical protein gene complete cds | KX884270.1 | Terrestrial | Arthropoda | Insecta |
| 1068 | Wuhan insect virus 23 strain spider96508 RdRp gene complete cds | KX884170.1 | Terrestrial | Arthropoda | Chelicerata |
| 1069 | Wuhan insect virus 23 strain WHCCII13263 segment Seg 1 RdRp gene complete cds | KX884196.1 | Terrestrial | Arthropoda | Insecta |

| | | | | | |
|---|---|---|---|---|---|
| 1070 | Wuhan insect virus 23 strain mosZJ33376 RdRp gene complete cds | KX884103.1 | Terrestrial | Arthropoda | Insecta |
| 1071 | Hubei picorna-like virus 44 strain WHYY20621 hypothetical protein gene complete cds | KX884301.1 | Terrestrial | Arthropoda | Myriapoda |
| 1072 | Crustacea Picorna-like virus N11 | GWHAPGE01000000 | South China Sea | Arthropoda | Crustacea |
| 1073 | Wenzhou picorna-like virus 15 strain WZFSL85504 hypothetical protein 1 and hypothetical protein 2 genes comple | KX884328.1 | Freshwater | Mollusca | Gastropoda |
| 1074 | Beihai zhaovirus-like virus 4 strain BHZY60633 polyprotein gene complete cds | KX884040.1 | South China Sea | Mollusca | Cephalopoda |
| 1075 | Wenling partiti-like virus 9 strain WLJQ23031 RdRp gene partial cds | KX884231.1 | East China Sea | Arthropoda | Crustacea |
| 1076 | Beihai tiger crab virus 1 strain HWRTX14333 replicase hypothetical protein putative structural protein and hypothe | KX883770.1 | South China Sea | Arthropoda | Crustacea |
| 1077 | Cephalopoda Mononega-like virus H1 | GWHAPHB01000000 | Yellow China Sea | Mollusca | Cephalopoda |
| 1078 | Wenling hepe-like virus 4 strain WLJQ102701 RdRp hypothetical protein putative structural protein and hypothetic | KX883829.1 | East China Sea | Arthropoda | Crustacea |
| 1079 | Hubei picorna-like virus 45 strain spider133521 hypothetical protein gene complete cds | KX883678.1 | Terrestrial | Arthropoda | Chelicerata |
| 1080 | Shahe picorna-like virus 9 strain SHWC0209c12638 hypothetical protein 1 and hypothetical protein 2 genes comple | KX883651.1 | Freshwater | Arthropoda | Crustacea |
| 1081 | Shahe picorna-like virus 9 strain SHWC13605 hypothetical protein 1 and hypothetical protein 2 genes complete cds | KX883661.1 | Freshwater | Arthropoda | Mix |
| 1082 | Shahe picorna-like virus 9 strain WHSFII15007 hypothetical protein 1 and hypothetical protein 2 genes complete cc | KX884290.1 | Terrestrial | Arthropoda | Crustacea |
| 1083 | Shahe picorna-like virus 9 strain CJLX30533 hypothetical protein 1 and hypothetical protein 2 genes complete cds | KX884550.1 | Freshwater | Arthropoda | Crustacea |
| 1084 | Beihai echinoderm virus 1 strain BHJP42036 hypothetical protein gene complete cds | KX883379.1 | South China Sea | Echinodermata | Echinozoa |
| 1085 | Changping earthworm virus 2 strain CPQY18140 segment 5 hypothetical protein gene complete cds | KX883853.1 | Terrestrial | Annelida | Clitellata |
| 1086 | Banna virus strain QTM104536 Banna VP5 gene complete cds | KX884642.1 | Terrestrial | Arthropoda | Insecta |
| 1087 | Hubei picorna-like virus 74 strain WGML145648 hypothetical protein 1 hypothetical protein 2 and hypothetical pro | KX884252.1 | Terrestrial | Arthropoda | Myriapoda |
| 1088 | Hubei partiti-like virus 31 strain spider131862 RdRp gene complete cds | KX884162.1 | Terrestrial | Arthropoda | Chelicerata |
| 1089 | Hubei virga-like virus 21 strain mosHB236537 hypothetical protein genes complete cds | KX883775.1 | Terrestrial | Arthropoda | Insecta |
| 1090 | Bivalvia Picorna-like virus D27 | GWHAOYV0100000( | East China Sea | Mollusca | Bivalvia |
| 1091 | Sanxia atyid shrimp virus 2 strain SXXX37884 hypothetical protein gene complete cds | KX883708.1 | Freshwater | Arthropoda | Crustacea |
| 1092 | Sanxia atyid shrimp virus 2 strain WHCCII13331 hypothetical protein gene complete cds | KX884278.1 | Terrestrial | Arthropoda | Insecta |
| 1093 | Changping earthworm virus 2 strain CPQY18140 segment 6 hypothetical protein genes complete cds | KX883854.1 | Terrestrial | Annelida | Clitellata |
| 1094 | Beihai rhabdo-like virus 5 strain BHJP63888 putative nucleoprotein hypothetical protein 1 hypothetical protein 2 ar | KX884407.1 | South China Sea | Echinodermata | Echinozoa |
| 1095 | Hubei lepidoptera virus 3 strain LCM141729 Cypovirus VP10 gene complete cds | KX884622.1 | Terrestrial | Arthropoda | Insecta |
| 1096 | Crustacea Picorna-like virus N5 | GWHAPFW01000000 | South China Sea | Arthropoda | Crustacea |
| 1097 | Hubei mosquito virus 1 strain mosHB236488 hypothetical protein gene complete cds | KX883917.1 | Terrestrial | Arthropoda | Insecta |
| 1098 | Hubei mosquito virus 1 strain mosZJ34341 hypothetical protein gene complete cds | KX883918.1 | Terrestrial | Arthropoda | Insecta |
| 1099 | Beihai sipunculid worm virus 6 strain BHNXC41400 polyprotein gene complete cds | KX883005.1 | South China Sea | Sipuncula | Sipuncula |
| 1100 | Bivalvia Picorna-like virus N1 | GWHAPIR01000000 | South China Sea | Mollusca | Bivalvia |
| 1101 | Beihai shrimp virus 5 strain BHBJDX18804 segment 2 hypothetical protein gene complete cds | KX882785.1 | South China Sea | Arthropoda | Crustacea |
| 1102 | Beihai shrimp virus 5 strain BHWZXX15615 segment 2 hypothetical protein gene complete cds | KX882793.1 | South China Sea | Arthropoda | Crustacea |
| 1103 | Bivalvia Picorna-like virus D32 | GWHAOYS01000000 | East China Sea | Mollusca | Bivalvia |
| 1104 | Hubei picorna-like virus 76 strain WGML143182 hypothetical protein 1 hypothetical protein 2 and hypothetical pro | KX883737.1 | Terrestrial | Arthropoda | Myriapoda |
| 1105 | Bivalvia Durna-like virus H1 | GWHAPCP01000000 | Yellow China Sea | Mollusca | Bivalvia |
| 1106 | Hubei odonate virus 7 strain QTM27298 hypothetical protein 1 hypothetical protein 2 and hypothetical protein 3 ge | KX883954.1 | Terrestrial | Arthropoda | Insecta |

| | | | | | |
|---|---|---|---|---|---|
| **1107** | Wutai Mosquito Virus strain QN3-5 nucleopasid protein (N) gene complete cds | KM817761.1 | Terrestrial | Arthropoda | Insecta |
| **1108** | Wenling narna-like virus 4 strain WLJQ103797 RNA-dependent RNA polymerase gene complete cds | KX883599.1 | East China Sea | Arthropoda | Crustacea |
| **1109** | Hubei myriapoda virus 2 strain WGML147749 hypothetical protein 1 hypothetical protein 2 and hypothetical protei | KX884256.1 | Terrestrial | Arthropoda | Myriapoda |
| **1110** | Gastropoda Picorna-like virus D2 | GWHAPJX01000000 | East China Sea | Mollusca | Gastropoda |
| **1111** | Sanxia picorna-like virus 1 strain SXXX37713 hypothetical protein 1 and hypothetical protein 2 genes complete cds | KX883711.1 | Freshwater | Arthropoda | Crustacea |
| **1112** | Bivalvia Picorna-like virus N24 | GWHAPLI01000000 | South China Sea | Mollusca | Bivalvia |
| **1113** | Beihai shrimp virus 2 strain BHWZXX12501 hypothetical protein 1 and hypothetical protein 2 genes complete cds | KX883423.1 | South China Sea | Arthropoda | Crustacea |
| **1114** | Beihai shrimp virus 2 strain BHBJDX18741 hypothetical protein 1 and hypothetical protein 2 genes complete cds | KX883334.1 | South China Sea | Arthropoda | Crustacea |
| **1115** | Gastropoda Unclassified virus D2 | GWHAPJZ01000000 | East China Sea | Mollusca | Gastropoda |
| **1116** | Wenzhou picorna-like virus 12 strain WZFSL74240 hypothetical protein 1 and hypothetical protein 2 genes comple | KX884340.1 | Freshwater | Mollusca | Gastropoda |
| **1117** | Hexanauplia Picorna-like virus D2 | GWHAPKC01000000 | East China Sea | Arthropoda | Hexanauplia |
| **1118** | Beihai hepe-like virus 10 strain BHZY60406 replicase hypothetical protein putative structural protein and hypotheti | KX883757.1 | South China Sea | Mollusca | Cephalopoda |
| **1119** | Beihai picorna-like virus 114 strain BHZY60932 hypothetical protein 1 and hypothetical protein 2 genes complete c | KX884522.1 | South China Sea | Mollusca | Cephalopoda |
| **1120** | Hubei partiti-like virus 58 strain QCM38725 RdRp gene complete cds | KX884110.1 | Terrestrial | Arthropoda | Insecta |
| **1121** | Bivalvia Durna-like virus D5 | GWHAOYL01000000 | East China Sea | Mollusca | Bivalvia |
| **1122** | Crustacea Picorna-like virus N12 | GWHAPBZ01000000 | South China Sea | Arthropoda | Crustacea |
| **1123** | Wuhan insect virus 8 strain WHCCII10272 hypothetical protein genes complete cds | KX883812.1 | Terrestrial | Arthropoda | Insecta |
| **1124** | Hubei picorna-like virus 15 strain arthropodmix13755 hypothetical protein 1 and hypothetical protein 2 genes comp | KX883290.1 | Terrestrial | Arthropoda | Mix |
| **1125** | Hubei picorna-like virus 15 strain WHXS4635 hypothetical protein 1 and hypothetical protein 2 genes complete cds | KX884295.1 | Terrestrial | Arthropoda | Insecta |
| **1126** | Hubei picorna-like virus 15 strain ZCM21299 hypothetical protein 1 and hypothetical protein 2 genes complete cds | KX884399.1 | Terrestrial | Arthropoda | Insecta |
| **1127** | Hubei picorna-like virus 15 strain WGML146626 hypothetical protein 1 and hypothetical protein 2 genes complete | KX884253.1 | Terrestrial | Arthropoda | Myriapoda |
| **1128** | Hubei picorna-like virus 15 strain GCM10717 hypothetical protein 1 and hypothetical protein 2 genes complete cds | KX883893.1 | Terrestrial | Arthropoda | Insecta |
| **1129** | Hubei picorna-like virus 15 strain WHYCM1097 hypothetical protein 1 and hypothetical protein 2 genes complete c | KX884298.1 | Terrestrial | Arthropoda | Insecta |
| **1130** | Hubei picorna-like virus 15 strain QTM27139 hypothetical protein 1 and hypothetical protein 2 genes complete cds | KX883941.1 | Terrestrial | Arthropoda | Insecta |
| **1131** | Kadipiro virus strain QTM27331 Kadipiro VP7 gene complete cds | KX884656.1 | Terrestrial | Arthropoda | Insecta |
| **1132** | Shahe narna-like virus 6 strain SHWC0209c11783 RNA-dependent RNA polymerase gene partial cds | KX883554.1 | Freshwater | Arthropoda | Crustacea |
| **1133** | Shuangao Insect Virus 1 strain QSA02 nucleopasid protein (N) gene complete cds | KM817739.1 | Terrestrial | Arthropoda | Insecta |
| **1134** | Changjiang crawfish virus 6 strain CJLX30496 hypothetical protein 1 and hypothetical protein 2 genes complete cds | KX884549.1 | Freshwater | Arthropoda | Crustacea |
| **1135** | Changjiang crawfish virus 6 strain SCM51510 hypothetical protein 1 and hypothetical protein 2 genes complete cds | KX883976.1 | Terrestrial | Arthropoda | Insecta |
| **1136** | Changjiang crawfish virus 6 strain QTM27188 hypothetical protein 1 and hypothetical protein 2 genes complete cds | KX883943.1 | Terrestrial | Arthropoda | Insecta |
| **1137** | Hubei reo-like virus 2 strain QTM25955 hypothetical protein gene partial cds | KX884663.1 | Terrestrial | Arthropoda | Insecta |
| **1138** | Beihai picorna-like virus 100 strain BHNXC41148 hypothetical protein 1 and hypothetical protein 2 genes complete | KX883389.1 | South China Sea | Sipuncula | Sipuncula |
| **1139** | Bivalvia Durna-like virus D11 | GWHAPAA01000000 | East China Sea | Mollusca | Bivalvia |
| **1140** | Sanxia water strider virus 7 strain SXSSP2499 hypothetical protein 1 hypothetical protein 2 and hypothetical protei | KX883705.1 | Freshwater | Arthropoda | Insecta |
| **1141** | Sanxia water strider virus 7 strain SXXX36023 hypothetical protein 1 hypothetical protein 2 and hypothetical protei | KX883719.1 | Freshwater | Arthropoda | Crustacea |
| **1142** | Beihai bunya-like virus 1 strain BHTSS17606 putative nucleoprotein gene complete cds | KX884750.1 | South China Sea | Arthropoda | Crustacea |
| **1143** | Hubei tetragnatha maxillosa virus 9 strain QTM27093 hypothetical protein 6 gene complete cds | KX884681.1 | Terrestrial | Arthropoda | Insecta |

| | | | | | |
|---|---|---|---|---|---|
| 1144 | Hubei tetragnatha maxillosa virus 9 strain SSZZ3444 hypothetical protein 7 gene complete cds | KX884710.1 | Freshwater | Arthropoda | Chelicerata |
| 1145 | Hubei tetragnatha maxillosa virus 9 strain arthropodmix14049 hypothetical protein 6 gene complete cds | KX884604.1 | Terrestrial | Arthropoda | Mix |
| 1146 | Wenzhou picorna-like virus 22 strain WZFSL83961 hypothetical protein gene complete cds | KX884331.1 | Freshwater | Mollusca | Gastropoda |
| 1147 | Hubei odonate virus 15 strain QTM133059 hypothetical protein 4 gene complete cds | KX884668.1 | Terrestrial | Arthropoda | Insecta |
| 1148 | Hubei picorna-like virus 29 strain arthropodmix14031 hypothetical protein gene complete cds | KX883292.1 | Terrestrial | Arthropoda | Mix |
| 1149 | Banna virus strain QTM104536 Banna VP12 gene complete cds | KX884648.1 | Terrestrial | Arthropoda | Insecta |
| 1150 | Wenling hepe-like virus 3 strain WLJQ102665 replicase and hypothetical protein genes complete cds | KX883828.1 | East China Sea | Arthropoda | Crustacea |
| 1151 | Kadipiro virus strain QTM27331 Kadipiro VP5 gene complete cds | KX884654.1 | Terrestrial | Arthropoda | Insecta |
| 1152 | Kadipiro virus strain QTM27331 Kadipiro VP9 gene complete cds | KX884658.1 | Terrestrial | Arthropoda | Insecta |
| 1153 | Hubei picorna-like virus 38 strain spider133826 hypothetical protein gene complete cds | KX883683.1 | Terrestrial | Arthropoda | Chelicerata |
| 1154 | Bivalvia Picorna-like virus N6 | GWHAPFA01000000 | South China Sea | Mollusca | Bivalvia |
| 1155 | Bivalvia Picorna-like virus H13 | GWHAPHD01000000 | Yellow China Sea | Mollusca | Bivalvia |
| 1156 | Hubei picorna-like virus 66 strain SSZZ2530 hypothetical protein 1 hypothetical protein 2 and hypothetical protein | KX883698.1 | Freshwater | Arthropoda | Chelicerata |
| 1157 | Hubei orthomyxo-like virus 1 strain SCM47652 polymerase PB1 gene partial cds | KX883873.1 | Terrestrial | Arthropoda | Insecta |
| 1158 | Beihai bunya-like virus 3 strain BWBFG40849 RNA-dependent RNA polymerase gene complete cds | KX884756.1 | South China Sea | Mollusca | Bivalvia |
| 1159 | Wenzhou picorna-like virus 32 strain WZSBei69487 hypothetical protein 1 and hypothetical protein 2 genes comple | KX884361.1 | East China Sea | Mollusca | Bivalvia |
| 1160 | Shuangao Insect Virus 2 strain QSA03 glycoprotein precursor (G) gene complete cds | KM817715.1 | Terrestrial | Arthropoda | Insecta |
| 1161 | Hubei picorna-like virus 47 strain WHYY21982 hypothetical protein gene complete cds | KX884302.1 | Terrestrial | Arthropoda | Myriapoda |
| 1162 | Wenzhou Crab Virus 1 strain RBX2 nucleocapsid (N) glycoprotein (G) and RNA-dependent RNA polymerase (L) g | KM817644.1 | East China Sea | Arthropoda | Crustacea |
| 1163 | Wenling crustacean virus 10 strain WLJQ101844 hypothetical protein 1 hypothetical protein 2 hypothetical protein | KX884450.1 | East China Sea | Arthropoda | Crustacea |
| 1164 | Shuangao Insect Virus 4 strain QSA07-1 PB1 (PB1) gene complete cds | KM817619.1 | Terrestrial | Arthropoda | Insecta |
| 1165 | Shayang Spider Virus 3 strain SYZZ-5 PB1 (PB1) gene complete cds | KM817618.1 | Terrestrial | Arthropoda | Chelicerata |
| 1166 | Sanxia Water Strider Virus 3 strain SXSSP01 segment PA polymerase PA gene complete cds | KX883880.1 | Freshwater | Arthropoda | Insecta |
| 1167 | Sanxia Water Strider Virus 3 strain SXSSP01 segment PB2 polymerase PB2 gene complete cds | KX883879.1 | Freshwater | Arthropoda | Insecta |
| 1168 | Hubei diptera virus 3 strain SCM17647 putative glycoprotein gene complete cds | KX884797.1 | Terrestrial | Arthropoda | Insecta |
| 1169 | Hubei orthomyxo-like virus 4 strain SCM51211 polymerase PB1 gene complete cds | KX883874.1 | Terrestrial | Arthropoda | Insecta |
| 1170 | Hubei picorna-like virus 37 strain ZCM296 hypothetical protein gene partial cds | KX884401.1 | Terrestrial | Arthropoda | Insecta |
| 1171 | Hubei picorna-like virus 37 strain WHLC4784 hypothetical protein gene complete cds | KX884281.1 | Terrestrial | Arthropoda | Insecta |
| 1172 | Hubei picorna-like virus 81 strain CJLX25805 hypothetical protein 1 and hypothetical protein 2 genes complete cds | KX884540.1 | Freshwater | Arthropoda | Crustacea |
| 1173 | Hubei picorna-like virus 81 strain QTM27117 hypothetical protein 1 hypothetical protein 2 and hypothetical proteir | KX883940.1 | Terrestrial | Arthropoda | Insecta |
| 1174 | Beihai picorna-like virus 91 strain SCJXSX38640 hypothetical protein 1 and hypothetical protein 2 genes complete | KX883957.1 | South China Sea | Arthropoda | Crustacea |
| 1175 | Beihai picorna-like virus 91 strain BHJJX25930 hypothetical protein 1 and hypothetical protein 2 genes complete c | KX883372.1 | South China Sea | Arthropoda | Crustacea |
| 1176 | Wuchang Cockraoch Virus 1 strain WCZL-5 glycoprotein precursor (G) gene complete cds | KM817721.1 | Terrestrial | Arthropoda | Insecta |
| 1177 | Wuhan Louse Fly Virus 4 strain BFJSC-4 PB1 (PB1) gene complete cds | KM817621.1 | Terrestrial | Arthropoda | Insecta |
| 1178 | Hubei picorna-like virus 23 strain spider70066 hypothetical protein 1 gene complete cds and hypothetical protein 2 | KX883692.1 | Terrestrial | Arthropoda | Chelicerata |
| 1179 | Hubei partiti-like virus 23 strain QTM24142 RdRp gene complete cds | KX884119.1 | Terrestrial | Arthropoda | Insecta |
| 1180 | Hubei picorna-like virus 60 strain SSZZ2616 hypothetical protein gene complete cds | KX883702.1 | Freshwater | Arthropoda | Chelicerata |

| | | | | | |
|---|---|---|---|---|---|
| 1181 | Hubei picorna-like virus 60 strain QTM27104 hypothetical protein gene complete cds | KX883939.1 | Terrestrial | Arthropoda | Insecta |
| 1182 | Wenling picobirna-like virus 1 strain WLJQ144192 RdRp gene partial cds | KX884223.1 | East China Sea | Arthropoda | Crustacea |
| 1183 | Wenzhou hepe-like virus 1 strain WZSLuoII97618 hypothetical protein genes complete cds | KX883835.1 | East China Sea | Mollusca | Gastropoda |
| 1184 | Wenzhou hepe-like virus 1 strain WZSLuoI86079 RdRp putative structural protein and hypothetical protein genes c | KX883834.1 | East China Sea | Mollusca | Gastropoda |
| 1185 | Hubei picobirna-like virus 3 strain WGML128211 RdRp gene complete cds | KX884187.1 | Terrestrial | Arthropoda | Myriapoda |
| 1186 | Wenling crustacean virus 6 strain WLJQ101491 hypothetical protein 1 and hypothetical protein 2 genes complete c | KX884312.1 | East China Sea | Arthropoda | Crustacea |
| 1187 | Jiujie Fly Virus strain JJ3-6 segment PA polymerase PA gene complete cds | KX883864.1 | Terrestrial | Arthropoda | Insecta |
| 1188 | Wenling partiti-like virus 7 strain WLJQ82808 RdRp gene complete cds | KX884234.1 | East China Sea | Arthropoda | Crustacea |
| 1189 | Wuhan spider virus 3 strain spider133889 hypothetical protein gene complete cds | KX883685.1 | Terrestrial | Arthropoda | Chelicerata |
| 1190 | Hubei earwig virus 2 strain GCM11219 RdRp gene complete cds | KX884097.1 | Terrestrial | Arthropoda | Insecta |
| 1191 | Zucchini yellow mosaic virus strain CJLX30535 polyprotein gene complete cds | KX884565.1 | Freshwater | Arthropoda | Crustacea |
| 1192 | Zucchini yellow mosaic virus strain spider131932 polyprotein gene complete cds | KX884570.1 | Terrestrial | Arthropoda | Chelicerata |
| 1193 | Hubei myriapoda virus 5 strain GCM10499 putative nucleoprotein gene complete cds | KX884760.1 | Terrestrial | Arthropoda | Insecta |
| 1194 | Hubei myriapoda virus 5 strain WGML140132 putative nucleoprotein gene complete cds | KX884842.1 | Terrestrial | Arthropoda | Myriapoda |
| 1195 | Bivalvia Picorna-like virus D14 | GWHAOYF01000000 | East China Sea | Mollusca | Bivalvia |
| 1196 | Hubei chuvirus-like virus 3 strain QTM26698 RNA-dependent RNA polymerase putative glycoprotein and hypothe | KX884427.1 | Terrestrial | Arthropoda | Insecta |
| 1197 | Bivalvia Picorna-like virus N55 | GWHAPEF01000000 | South China Sea | Mollusca | Bivalvia |
| 1198 | Wenzhou shrimp virus 6 strain shrimp14542 hypothetical protein 1 and hypothetical protein 2 genes complete cds | KX883978.1 | East China Sea | Arthropoda | Crustacea |
| 1199 | Wenzhou shrimp virus 6 strain WZRBX43423 hypothetical protein 1 and hypothetical protein 2 genes complete cds | KX884348.1 | East China Sea | Arthropoda | Crustacea |
| 1200 | Bivalvia Picorna-like virus D65 | GWHAPAH01000000 | East China Sea | Mollusca | Bivalvia |
| 1201 | Bivalvia Picorna-like virus H10 | GWHAPJH01000000 | Yellow China Sea | Mollusca | Bivalvia |
| 1202 | Wenzhou picorna-like virus 35 strain WZRBX39547 hypothetical protein 1 and hypothetical protein 2 genes compl | KX884344.1 | East China Sea | Arthropoda | Crustacea |
| 1203 | Kadipiro virus strain QTM27331 Kadipiro VP11 gene complete cds | KX884660.1 | Terrestrial | Arthropoda | Insecta |
| 1204 | Hubei picorna-like virus 36 strain SCM50248 hypothetical protein gene complete cds | KX883970.1 | Terrestrial | Arthropoda | Insecta |
| 1205 | Hubei earwig virus 1 strain GCM19162 segment 4 nucleocapsid protein gene complete cds | KX883861.1 | Terrestrial | Arthropoda | Insecta |
| 1206 | Bivalvia Picorna-like virus D58 | GWHAPAC01000000 | East China Sea | Mollusca | Bivalvia |
| 1207 | Taishun Tick Virus strain BL198 nucleocapsid (N) ORF2 (ORF2) and RNA-dependent RNA polymerase (L) genes | KM817643.1 | Terrestrial | Arthropoda | Chelicerata |
| 1208 | Bivalvia Picorna-like virus D80 | GWHAPKP01000000 | East China Sea | Mollusca | Bivalvia |
| 1209 | Beihai mantis shrimp virus 2 strain BHXG46195 replicase and hypothetical protein genes complete cds | KX883752.1 | South China Sea | Arthropoda | Crustacea |
| 1210 | Cephalopoda Bunya-like virus H1 | GWHAPHC01000000 | Yellow China Sea | Mollusca | Cephalopoda |
| 1211 | Hubei orthoptera virus 5 strain ZCM94262 hypothetical protein 1 hypothetical protein 2 hypothetical protein 3 and | KX884463.1 | Terrestrial | Arthropoda | Insecta |
| 1212 | Gastropoda Picorna-like virus D17 | GWHAPAZ01000000 | East China Sea | Mollusca | Gastropoda |
| 1213 | Hubei dimarhabdovirus virus 3 strain QTM26149 putative nucleoprotein hypothetical protein 2 hypothetical protein | KX884423.1 | Terrestrial | Arthropoda | Insecta |
| 1214 | Hubei bunya-like virus 13 strain QTM27285 putative nucleoprotein gene complete cds | KX884785.1 | Terrestrial | Arthropoda | Insecta |
| 1215 | Bole Tick Virus 3 strain BL199 complete genome | KM817593.1 | Terrestrial | Arthropoda | Chelicerata |
| 1216 | Beihai bunya-like virus 2 strain BHZC35426 putative nucleoprotein gene complete cds | KX884755.1 | South China Sea | Mollusca | Bivalvia |
| 1217 | Bivalvia Bunya-like virus N2 | GWHAPLK01000000 | South China Sea | Mollusca | Bivalvia |

| | | | | | |
|---|---|---|---|---|---|
| 1218 | Gastropoda Picorna-like virus N8 | GWHAPFR01000000 | South China Sea | Mollusca | Gastropoda |
| 1219 | Wenling picorna-like virus 2 strain WLJQ104117 hypothetical protein 1 and hypothetical protein 2 genes complete | KX884319.1 | East China Sea | Arthropoda | Crustacea |
| 1220 | Shahe heteroptera virus 3 strain SHWC0209c8767 putative nucleoprotein gene complete cds | KX884820.1 | Freshwater | Arthropoda | Crustacea |
| 1221 | Shahe heteroptera virus 3 strain SHWCII5319 putative nucleoprotein gene complete cds | KX884831.1 | Freshwater | Arthropoda | Mix |
| 1222 | Shahe heteroptera virus 3 strain SHWC01c3680 putative nucleoprotein gene complete cds | KX884815.1 | Freshwater | Arthropoda | Insecta |
| 1223 | Tacheng Tick Virus 5 strain TC254 complete genome | KM817600.1 | Terrestrial | Arthropoda | Chelicerata |
| 1224 | Sanxia Water Strider Virus 3 strain SXSSP01 segment NP nucleocapsid protein gene complete cds | KX883881.1 | Freshwater | Arthropoda | Insecta |
| 1225 | Beihai hermit crab virus 3 strain BHJJX21702 putative glycoprotein hypothetical protein 1 hypothetical protein 2 ar | KX884404.1 | South China Sea | Arthropoda | Crustacea |
| 1226 | Wuhan Tick Virus 1 strain X78-2 nucleocapsid (N) ORF2 (ORF2) ORF3 (ORF3) and RNA-dependent RNA polym | KM817660.1 | Terrestrial | Arthropoda | Chelicerata |
| 1227 | Bivalvia Amarillo-like virus H1 | GWHAPGN01000000 | Yellow China Sea | Mollusca | Bivalvia |
| 1228 | Tacheng Tick Virus 6 strain TCRP-2 ORF1 (ORF1) ORF2 (ORF2) ORF3 (ORF3) and RNA-dependent RNA polyn | KM817641.1 | Terrestrial | Arthropoda | Chelicerata |
| 1229 | Tacheng Tick Virus 4 strain TCRP-1 complete genome | KM817599.1 | Terrestrial | Arthropoda | Chelicerata |
| 1230 | Sanxia partiti-like virus 2 strain SXXX6738 RdRp gene complete cds | KX884180.1 | Freshwater | Arthropoda | Crustacea |
| 1231 | Gastropoda Unclassified virus D3 | GWHAPAW0100000( | East China Sea | Mollusca | Gastropoda |
| 1232 | Bivalvia Picorna-like virus D71 | GWHAPIE01000000 | East China Sea | Mollusca | Bivalvia |
| 1233 | Hubei rhabdo-like virus 1 strain QTM25107 hypothetical protein 1 hypothetical protein 2 hypothetical protein 3 put | KX884422.1 | Terrestrial | Arthropoda | Insecta |
| 1234 | Hubei tick virus 2 strain tick109131 hypothetical protein gene complete cds | KX883730.1 | Terrestrial | Arthropoda | Chelicerata |
| 1235 | Crustacea Picorna-like virus N27 | GWHAPDC01000000 | South China Sea | Arthropoda | Crustacea |
| 1236 | Crustacea Jingchu-like virus N1 | GWHAPFH01000000 | South China Sea | Arthropoda | Crustacea |
| 1237 | Whenzhou Shrimp Virus 1 strain BJDX-5 nucleopasid protein (N) gene complete cds | KM817746.1 | East China Sea | Arthropoda | Crustacea |
| 1238 | Bivalvia Hepeli-like virus D3 | GWHAPII01000000 | East China Sea | Mollusca | Bivalvia |
| 1239 | Hubei picorna-like virus 52 strain QCM127295 hypothetical protein gene complete cds | KX883920.1 | Terrestrial | Arthropoda | Insecta |
| 1240 | Hubei tombus-like virus 27 strain SSZZ3453 hypothetical protein 1 and hypothetical protein 2 genes complete cds | KX883171.1 | Freshwater | Arthropoda | Chelicerata |
| 1241 | Kadipiro virus strain QTM27331 Kadipiro VP8 gene complete cds | KX884657.1 | Terrestrial | Arthropoda | Insecta |
| 1242 | Wuchan romanomermis nematode virus 1 strain WCLSXC58279 hypothetical protein gene complete cds | KX883806.1 | Terrestrial | Nematoda | Enoplea |
| 1243 | Crustacea Unclassified virus H1 | GWHAPJC01000000 | Yellow China Sea | Arthropoda | Crustacea |
| 1244 | Beihai picorna-like virus 23 strain BHZY60791 hypothetical protein 1 and hypothetical protein 2 genes complete cc | KX884514.1 | South China Sea | Mollusca | Cephalopoda |
| 1245 | Beihai picorna-like virus 23 strain YYSZX17154 hypothetical protein 1 and hypothetical protein 2 genes complete ( | KX884386.1 | South China Sea | Arthropoda | Crustacea |
| 1246 | Beihai picorna-like virus 67 strain BHZY60873 hypothetical protein gene complete cds | KX884516.1 | South China Sea | Mollusca | Cephalopoda |
| 1247 | Changjiang partiti-like virus 1 strain CJLX57436 RdRp gene partial cds | KX884088.1 | Freshwater | Arthropoda | Crustacea |
| 1248 | Hubei picorna-like virus 64 strain SSZZ3507 hypothetical protein 1 and hypothetical protein 2 genes complete cds | KX883699.1 | Freshwater | Arthropoda | Chelicerata |
| 1249 | Wuhan house centipede virus 8 strain WHYY33011 RdRp gene complete cds | KX884217.1 | Terrestrial | Arthropoda | Myriapoda |
| 1250 | Beihai hypo-like virus 1 strain BHBei69634 polyprotein gene complete cds | KX883004.1 | South China Sea | Mollusca | Bivalvia |
| 1251 | Beihai hypo-like virus 1 strain BHZC36965 polyprotein gene complete cds | KX883006.1 | South China Sea | Mollusca | Bivalvia |
| 1252 | Hubei diptera virus 19 strain SCM51314 RdRp gene complete cds | KX884153.1 | Terrestrial | Arthropoda | Insecta |
| 1253 | Hubei diptera virus 19 strain QTM27098 RdRp gene complete cds | KX884132.1 | Terrestrial | Arthropoda | Insecta |
| 1254 | Hexanauplia Bunya-like virus D1 | GWHAPBY01000000 | East China Sea | Arthropoda | Hexanauplia |

| | | | | | |
|---|---|---|---|---|---|
| 1255 | Hubei odonate virus 9 strain QTM74767 putative nucleoprotein gene complete cds | KX884788.1 | Terrestrial | Arthropoda | Insecta |
| 1256 | Wuhan Millipede Virus 2 strain WHWG03 RNA-dependent RNA polymerase (L) gene complete cds | KM817696.2 | Terrestrial | Arthropoda | Insecta |
| 1257 | Wuhan Insect virus 2 strain WHDL02 nucleopasid protein (N) gene complete cds | KM817753.2 | Terrestrial | Arthropoda | Insecta |
| 1258 | Beihai hermit crab virus 2 strain BHJJX25757 RNA-dependent RNA polymerase gene complete cds | KX884741.1 | South China Sea | Arthropoda | Crustacea |
| 1259 | Hubei diptera virus 7 strain SCM173098 putative nucleoprotein gene complete cds | KX884808.1 | Terrestrial | Arthropoda | Insecta |
| 1260 | Huangpi Tick Virus 1 strain H124-1 nucleopasid protein (N) gene complete cds | KM817734.1 | Terrestrial | Arthropoda | Chelicerata |
| 1261 | Tacheng Tick Virus 1 strain TC253 RNA-dependent RNA polymerase (L) gene complete cds | KM817683.1 | Terrestrial | Arthropoda | Chelicerata |
| 1262 | Hubei diptera virus 5 strain SCM245062 putative nucleoprotein gene complete cds | KX884801.1 | Terrestrial | Arthropoda | Insecta |
| 1263 | Huangpi Tick Virus 1 strain H124-1 RNA-dependent RNA polymerase (L) gene complete cds | KM817667.1 | Terrestrial | Arthropoda | Chelicerata |
| 1264 | Shayang Spider Virus 1 strain SYZZ-4 nucleopasid protein (N) gene complete cds | KM817738.1 | Terrestrial | Arthropoda | Chelicerata |
| 1265 | Wuhan pillworm virus 1 strain WHSFII20185 RNA-dependent RNA polymerase gene complete cds | KX884848.1 | Terrestrial | Arthropoda | Crustacea |
| 1266 | Hubei orthoptera virus 6 strain ZCM21154 segment 3 polymerase PA gene complete cds | KX883886.1 | Terrestrial | Arthropoda | Insecta |
| 1267 | Cephalopoda Durna-like virus H2 | GWHAPGZ01000000 | Yellow China Sea | Mollusca | Cephalopoda |
| 1268 | Beihai blue swimmer crab virus 1 strain YYSZX13554 hypothetical protein gene complete cds | KX884384.1 | South China Sea | Arthropoda | Crustacea |
| 1269 | Wuhan cricket virus 2 strain WHXS3745 segment Seg 2 hypothetical protein gene complete cds | KX884209.1 | Terrestrial | Arthropoda | Insecta |
| 1270 | Wuhan cricket virus 2 strain ZCM19297 segment Seg 2 hypothetical protein gene complete cds | KX884244.1 | Terrestrial | Arthropoda | Insecta |
| 1271 | Hubei bunya-like virus 2 strain QTM26385 hypothetical protein gene complete cds | KX884781.1 | Terrestrial | Arthropoda | Insecta |
| 1272 | Bivalvia Picorna-like virus N32 | GWHAPLM01000000 | South China Sea | Mollusca | Bivalvia |
| 1273 | Hubei arthropod virus 1 strain SXSSP2046 hypothetical protein gene complete cds | KX883703.1 | Freshwater | Arthropoda | Insecta |
| 1274 | Hubei arthropod virus 1 strain SSZZ94 hypothetical protein gene complete cds | KX883695.1 | Freshwater | Arthropoda | Chelicerata |
| 1275 | Hubei arthropod virus 1 strain arthropodmix4509 hypothetical protein gene complete cds | KX883297.1 | Terrestrial | Arthropoda | Mix |
| 1276 | Hubei partiti-like virus 43 strain WHLC8707 RdRp gene complete cds | KX884198.1 | Terrestrial | Arthropoda | Insecta |
| 1277 | Hubei partiti-like virus 43 strain ZCM99354 RdRp gene complete cds | KX884249.1 | Terrestrial | Arthropoda | Insecta |
| 1278 | Gastropoda Picorna-like virus H1 | GWHAPCK01000000 | Yellow China Sea | Mollusca | Gastropoda |
| 1279 | Beihai sea slater virus 2 strain BHHZL10411 hypothetical protein 1 and hypothetical protein 2 genes complete cds | KX883358.1 | South China Sea | Arthropoda | Crustacea |
| 1280 | Beihai picorna-like virus 2 strain BHJJX25716 hypothetical protein 1 and hypothetical protein 2 genes complete cds | KX883369.1 | South China Sea | Arthropoda | Crustacea |
| 1281 | Beihai picorna-like virus 2 strain BHBJDX18281 hypothetical protein 1 and hypothetical protein 2 genes complete | KX883326.1 | South China Sea | Arthropoda | Crustacea |
| 1282 | Beihai picorna-like virus 2 strain BHHK129932 hypothetical protein 1 and hypothetical protein 2 genes complete c | KX883347.1 | South China Sea | Cnidaria | Anthozoa |
| 1283 | Beihai picorna-like virus 2 strain BZL93063 hypothetical protein 1 and hypothetical protein 2 genes complete cds | KX884535.1 | South China Sea | Mollusca | Gastropoda |
| 1284 | Beihai picorna-like virus 2 strain BHNXC40556 hypothetical protein 1 and hypothetical protein 2 genes complete c | KX883387.1 | South China Sea | Sipuncula | Sipuncula |
| 1285 | Beihai picorna-like virus 2 strain BHZC36110 hypothetical protein 1 and hypothetical protein 2 genes complete cds | KX884477.1 | South China Sea | Mollusca | Bivalvia |
| 1286 | Beihai picorna-like virus 2 strain BHZY60755 hypothetical protein 1 and hypothetical protein 2 genes complete cds | KX884513.1 | South China Sea | Mollusca | Cephalopoda |
| 1287 | Beihai picorna-like virus 2 strain BHWZXX15591 hypothetical protein 1 and hypothetical protein 2 genes complete | KX883428.1 | South China Sea | Arthropoda | Crustacea |
| 1288 | Beihai picorna-like virus 2 strain HWRTX12828 hypothetical protein 1 and hypothetical protein 2 genes complete c | KX883909.1 | South China Sea | Arthropoda | Crustacea |
| 1289 | Beihai mantis shrimp virus 5 strain BHXG25299 hypothetical protein 1 and hypothetical protein 2 genes complete c | KX884465.1 | South China Sea | Arthropoda | Crustacea |
| 1290 | Hubei picorna-like virus 28 strain QTM24820 hypothetical protein gene complete cds | KX883931.1 | Terrestrial | Arthropoda | Insecta |
| 1291 | Shahe heteroptera virus 2 strain SHWC13625 hypothetical protein gene complete cds | KX883669.1 | Freshwater | Arthropoda | Mix |

| | | | | | |
|---|---|---|---|---|---|
| 1292 | Shahe heteroptera virus 2 strain SHWCII5334 hypothetical protein gene complete cds | KX883674.1 | Freshwater | Arthropoda | Mix |
| 1293 | Shahe heteroptera virus 2 strain SHWC01c3694 hypothetical protein gene complete cds | KX883987.1 | Freshwater | Arthropoda | Insecta |
| 1294 | Shahe heteroptera virus 2 strain SHWC0209c10451 hypothetical protein gene complete cds | KX883644.1 | Freshwater | Arthropoda | Crustacea |
| 1295 | Beihai picobirna-like virus 8 strain BHNXC71065 segment Seg 1 RdRp gene complete cds | KX884064.1 | South China Sea | Sipuncula | Sipuncula |
| 1296 | Beihai picobirna-like virus 8 strain BHZC29883 RdRp gene complete cds | KX884076.1 | South China Sea | Mollusca | Bivalvia |
| 1297 | Hubei partiti-like virus 15 strain spider135364 RdRp gene partial cds | KX884167.1 | Terrestrial | Arthropoda | Chelicerata |
| 1298 | Beihai picorna-like virus 1 strain BHJJX25883 hypothetical protein 1 and hypothetical protein 2 genes complete cds | KX883371.1 | South China Sea | Arthropoda | Crustacea |
| 1299 | Beihai picorna-like virus 1 strain BHTSS18000 hypothetical protein 1 and hypothetical protein 2 genes complete cd | KX883421.1 | South China Sea | Arthropoda | Crustacea |
| 1300 | Beihai picorna-like virus 1 strain BHZC36988 hypothetical protein 1 and hypothetical protein 2 genes complete cds | KX884481.1 | South China Sea | Mollusca | Bivalvia |
| 1301 | Beihai picorna-like virus 1 strain SCJXSX39332 hypothetical protein 1 and hypothetical protein 2 genes complete c | KX883964.1 | South China Sea | Arthropoda | Crustacea |
| 1302 | Beihai picorna-like virus 1 strain BHBJDX18697 hypothetical protein 1 and hypothetical protein 2 genes complete | KX883332.1 | South China Sea | Arthropoda | Crustacea |
| 1303 | Beihai picorna-like virus 1 strain BHNXC41230 hypothetical protein 1 and hypothetical protein 2 genes complete c | KX883392.1 | South China Sea | Sipuncula | Sipuncula |
| 1304 | Beihai picorna-like virus 116 strain YYSZX16386 hypothetical protein 1 and hypothetical protein 2 genes complete | KX884385.1 | South China Sea | Arthropoda | Crustacea |
| 1305 | Wenling crustacean virus 7 strain WLJQ104070 putative nucleoprotein gene complete cds | KX884860.1 | East China Sea | Arthropoda | Crustacea |
| 1306 | Bivalvia Picorna-like virus D45 | GWHAOZG01000000 | East China Sea | Mollusca | Bivalvia |
| 1307 | Xinzhou nematode virus 2 strain XZSJSC33555 hypothetical protein 1 and hypothetical protein 2 genes complete c | KX884383.1 | Terrestrial | Nematoda | Mix |
| 1308 | Bivalvia Picorna-like virus D46 | GWHAOZJ01000000 | East China Sea | Mollusca | Bivalvia |
| 1309 | Hubei virga-like virus 16 strain arthropodmix13816 hypothetical protein RdRp and hypothetical protein genes comp | KX883739.1 | Terrestrial | Arthropoda | Mix |
| 1310 | Crustacea Picorna-like virus N16 | GWHAPDD01000000 | South China Sea | Arthropoda | Crustacea |
| 1311 | Gastropoda Unclassified virus D5 | GWHAPBL01000000 | East China Sea | Mollusca | Gastropoda |
| 1312 | Hubei virga-like virus 15 strain QCM619087 RdRp and hypothetical protein genes complete cds | KX883778.1 | Terrestrial | Arthropoda | Insecta |
| 1313 | Shayang virga-like virus 1 strain HC20676 hypothetical protein gene complete cds | KX883768.1 | Terrestrial | Nematoda | Rhabditida |
| 1314 | Shayang virga-like virus 1 strain SYJFC48301 RdRp and hypothetical protein genes complete cds | KX883805.1 | Terrestrial | Nematoda | Chromadorea |
| 1315 | Bivalvia Picorna-like virus N19 | GWHAPCJ01000000 | South China Sea | Mollusca | Bivalvia |
| 1316 | Beihai mantis shrimp virus 1 strain BHXG26050 hypothetical protein genes complete cds | KX883751.1 | South China Sea | Arthropoda | Crustacea |
| 1317 | Wenzhou picorna-like virus 6 strain WZSBei69574 hypothetical protein 1 and hypothetical protein 2 genes complet | KX884366.1 | East China Sea | Mollusca | Bivalvia |
| 1318 | Gastropoda Picorna-like virus D8 | GWHAPAQ01000000 | East China Sea | Mollusca | Gastropoda |
| 1319 | Wenzhou picorna-like virus 51 strain WZSBei69571 hypothetical protein 1 and hypothetical protein 2 genes comple | KX884364.1 | East China Sea | Mollusca | Bivalvia |
| 1320 | Hubei insect virus 1 strain arthropodmix14045 nucleoprotein gene complete cds | KX884736.1 | Terrestrial | Arthropoda | Mix |
| 1321 | Beihai shrimp virus 1 strain BHBJDX17672 hypothetical protein 1 and hypothetical protein 2 genes complete cds | KX883324.1 | South China Sea | Arthropoda | Crustacea |
| 1322 | Hubei bunya-like virus 3 strain QTM25853 putative nucleoprotein gene complete cds | KX884779.1 | Terrestrial | Arthropoda | Insecta |
| 1323 | Bivalvia Picorna-like virus N30 | GWHAPIW01000000 | South China Sea | Mollusca | Bivalvia |
| 1324 | Beihai picorna-like virus 14 strain BHHK130444 hypothetical protein 1 and hypothetical protein 2 genes complete | KX883348.1 | South China Sea | Cnidaria | Anthozoa |
| 1325 | Bivalvia Picorna-like virus D47 | GWHAOZX01000000 | East China Sea | Mollusca | Bivalvia |
| 1326 | Hubei myriapoda virus 4 strain WGML147816 hypothetical protein 1 and hypothetical protein 2 genes complete cds | KX884258.1 | Terrestrial | Arthropoda | Myriapoda |
| 1327 | Polychaeta Picorna-like virus H1 | GWHAPCY01000000 | Yellow China Sea | Annelida | Polychaeta |
| 1328 | Bivalvia Picorna-like virus D55 | GWHAOZH01000000 | East China Sea | Mollusca | Bivalvia |

| | | | | | |
|---|---|---|---|---|---|
| 1329 | Wenzhou bivalvia virus 1 strain beimix75829 hypothetical protein gene complete cds | KX883302.1 | East China Sea | Mollusca | Bivalvia |
| 1330 | Beihai picorna-like virus 11 strain BHZY60905 hypothetical protein 1 and hypothetical protein 2 genes complete cc | KX884519.1 | South China Sea | Mollusca | Cephalopoda |
| 1331 | Gastropoda Picorna-like virus H2 | GWHAPCL01000000 | Yellow China Sea | Mollusca | Gastropoda |
| 1332 | Shahe isopoda virus 1 strain SHWCII5332 hypothetical protein gene complete cds | KX883673.1 | Freshwater | Arthropoda | Mix |
| 1333 | Shahe isopoda virus 1 strain SHWC0209c12784 hypothetical protein gene complete cds | KX883643.1 | Freshwater | Arthropoda | Crustacea |
| 1334 | Beihai picorna-like virus 54 strain BHNXC41489 hypothetical protein 1 and hypothetical protein 2 genes complete | KX883407.1 | South China Sea | Sipuncula | Sipuncula |
| 1335 | Banna virus strain QTM104536 Banna VP7 gene complete cds | KX884644.1 | Terrestrial | Arthropoda | Insecta |
| 1336 | Bivalvia Picorna-like virus D76 | GWHAPKN01000000 | East China Sea | Mollusca | Bivalvia |
| 1337 | Beihai paphia shell virus 2 strain YYSZX17625 hypothetical protein 1 and hypothetical protein 2 genes complete cc | KX884389.1 | South China Sea | Arthropoda | Crustacea |
| 1338 | Beihai paphia shell virus 2 strain BWBFG38484 hypothetical protein 2 genes complete cds | KX884525.1 | South China Sea | Mollusca | Bivalvia |
| 1339 | Crustacea Picorna-like virus N19 | GWHAPDJ01000000 | South China Sea | Arthropoda | Crustacea |
| 1340 | Beihai picorna-like virus 93 strain BHBJDX18546 hypothetical protein 1 and hypothetical protein 2 genes complete | KX883328.1 | South China Sea | Arthropoda | Crustacea |
| 1341 | Bivalvia Picorna-like virus D38 | GWHAOYZ01000000 | East China Sea | Mollusca | Bivalvia |
| 1342 | Beihai picorna-like virus 82 strain YYSZX17728 hypothetical protein 1 and hypothetical protein 2 genes complete | KX884390.1 | South China Sea | Arthropoda | Crustacea |
| 1343 | Polychaeta Picorna-like virus H5 | GWHAPCU01000000 | Yellow China Sea | Annelida | Polychaeta |
| 1344 | Hubei picorna-like virus 61 strain mosHB235903 hypothetical protein gene complete cds | KX883915.1 | Terrestrial | Arthropoda | Insecta |
| 1345 | Polychaeta Picorna-like virus H6 | GWHAPCT01000000 | Yellow China Sea | Annelida | Polychaeta |
| 1346 | Bivalvia Picorna-like virus D40 | GWHAOYY0100000C | East China Sea | Mollusca | Bivalvia |
| 1347 | Wenling picorna-like virus 3 strain WLJQ100015 hypothetical protein 1 and hypothetical protein 2 genes complete | KX884309.1 | East China Sea | Arthropoda | Crustacea |
| 1348 | Wuhan cricket virus 2 strain WHXS3745 segment Seg 6 hypothetical protein gene complete cds | KX884213.1 | Terrestrial | Arthropoda | Insecta |
| 1349 | Wuhan cricket virus 2 strain ZCM19297 segment Seg 6 hypothetical protein gene complete cds | KX884248.1 | Terrestrial | Arthropoda | Insecta |
| 1350 | Changjiang hepe-like virus 1 strain CJLX30636 replicase and hypothetical protein genes complete cds | KX883762.1 | Freshwater | Arthropoda | Crustacea |
| 1351 | Wuhan cricket virus 2 strain ZCM19297 segment Seg 5 hypothetical protein gene complete cds | KX884247.1 | Terrestrial | Arthropoda | Insecta |
| 1352 | Wuhan cricket virus 2 strain WHXS3745 segment Seg 5 hypothetical protein gene complete cds | KX884212.1 | Terrestrial | Arthropoda | Insecta |
| 1353 | Hubei virga-like virus 1 strain mosHB236471 hypothetical protein genes complete cds | KX883773.1 | Terrestrial | Arthropoda | Insecta |
| 1354 | Shuangao partiti-like virus 1 strain mosZJ25114 RdRp gene complete cds | KX884102.1 | Terrestrial | Arthropoda | Insecta |
| 1355 | Shuangao partiti-like virus 1 strain insectZJ79891 RdRp gene complete cds | KX884100.1 | Terrestrial | Arthropoda | Insecta |
| 1356 | Beihai shrimp virus 6 strain BHBJDX18793 hypothetical protein 1 and hypothetical protein 2 genes complete cds | KX883023.1 | South China Sea | Arthropoda | Crustacea |
| 1357 | Beihai shrimp virus 6 strain BHWZXX15644 hypothetical protein 1 and hypothetical protein 2 genes complete cds | KX883051.1 | South China Sea | Arthropoda | Crustacea |
| 1358 | Bivalvia Picorna-like virus D48 | GWHAOZU01000000 | East China Sea | Mollusca | Bivalvia |
| 1359 | Bivalvia Picorna-like virus D59 | GWHAPAE01000000 | East China Sea | Mollusca | Bivalvia |
| 1360 | Ubei picorna-like virus 3 strain WHCC101453 hypothetical protein 1 and hypothetical protein 2 genes complete cds | KX884266.1 | Terrestrial | Arthropoda | Insecta |
| 1361 | Sanxia noda-like virus 1 strain SXXX12565 hypothetical protein gene complete cds | KX883188.1 | Freshwater | Arthropoda | Crustacea |
| 1362 | Hubei partiti-like virus 22 strain spider58365 RdRp gene partial cds | KX884168.1 | Terrestrial | Arthropoda | Chelicerata |
| 1363 | Hubei partiti-like virus 22 strain mosZJ34436 RdRp gene complete cds | KX884104.1 | Terrestrial | Arthropoda | Insecta |
| 1364 | Wenling partiti-like virus 4 strain WLJQ103127 RdRp gene complete cds | KX884228.1 | East China Sea | Arthropoda | Crustacea |
| 1365 | Hubei picorna-like virus 11 strain WHZHC53090 hypothetical protein gene complete cds | KX884306.1 | Terrestrial | Nematoda | Chromadorea |

| | | | | | |
|---|---|---|---|---|---|
| 1366 | Bivalvia Picorna-like virus D75 | GWHAPIA01000000 | East China Sea | Mollusca | Bivalvia |
| 1367 | Wuhan house centipede virus 4 strain arthropodmix8296 segment 2 hypothetical protein gene complete cds | KX882779.1 | Terrestrial | Arthropoda | Mix |
| 1368 | Wuhan house centipede virus 4 strain WHYY19909 segment 2 hypothetical protein gene complete cds | KX882924.1 | Terrestrial | Arthropoda | Myriapoda |
| 1369 | Wenzhou picorna-like virus 25 strain WZFSL75477 hypothetical protein gene complete cds | KX884338.1 | Freshwater | Mollusca | Gastropoda |
| 1370 | Beihai narna-like virus 7 strain BHZC36598 RNA-dependent RNA polymerase gene partial cds | KX883504.1 | South China Sea | Mollusca | Bivalvia |
| 1371 | Wuhan cricket virus 2 strain WHXS3745 segment Seg 4 hypothetical protein gene complete cds | KX884211.1 | Terrestrial | Arthropoda | Insecta |
| 1372 | Wuhan cricket virus 2 strain ZCM19297 segment Seg 4 hypothetical protein gene complete cds | KX884246.1 | Terrestrial | Arthropoda | Insecta |
| 1373 | Beihai picorna-like virus 119 strain BHJP60437 hypothetical protein 1 and hypothetical protein 2 genes complete c | KX883383.1 | South China Sea | Echinodermata | Echinozoa |
| 1374 | Beihai shrimp virus 4 strain BHWZXX19227 hypothetical protein 1 and hypothetical protein 2 genes complete cds | KX882795.1 | South China Sea | Arthropoda | Crustacea |
| 1375 | Hubei sobemo-like virus 24 strain WHLC5390 hypothetical protein 1 and hypothetical protein 2 genes complete cds | KX882912.1 | Terrestrial | Arthropoda | Insecta |
| 1376 | Wenling picorna-like virus 8 strain WLJQ103995 hypothetical protein gene complete cds | KX884318.1 | East China Sea | Arthropoda | Crustacea |
| 1377 | Sanxia narna-like virus 2 strain SXXX45823 RNA-dependent RNA polymerase gene partial cds | KX883567.1 | Freshwater | Arthropoda | Crustacea |
| 1378 | Sanxia water strider virus 20 strain SXSSP2584 hypothetical protein 1 and hypothetical protein 2 genes complete co | KX882976.1 | Freshwater | Arthropoda | Insecta |
| 1379 | Hubei partiti-like virus 6 strain spider131688 RdRp gene complete cds | KX884161.1 | Terrestrial | Arthropoda | Chelicerata |
| 1380 | Bivalvia Toli-like virus D1 | GWHAOZE01000000 | East China Sea | Mollusca | Bivalvia |
| 1381 | Wenzhou crab virus 5 strain WZRBX37749 hypothetical protein 1 hypothetical protein 2 hypothetical protein 3 hyp | KX882994.1 | East China Sea | Arthropoda | Crustacea |
| 1382 | Zhejiang mosquito virus 3 strain mosZJ35354 hypothetical protein and RNA-dependent RNA polymerase genes cor | KX883538.1 | Terrestrial | Arthropoda | Insecta |
| 1383 | Zhejiang mosquito virus 3 strain 3mos6196 hypothetical protein gene complete cds and RNA-dependent RNA poly | KX883461.1 | Terrestrial | Arthropoda | Insecta |
| 1384 | Zhejiang mosquito virus 3 strain mosHB236413 hypothetical protein gene complete cds and RNA-dependent RNA | KX883537.1 | Terrestrial | Arthropoda | Insecta |
| 1385 | Gastropoda Sobeli-like virus N1 | GWHAPFO01000000 | South China Sea | Mollusca | Gastropoda |
| 1386 | Crustacea Picorna-like virus N28 | GWHAPDE01000000 | South China Sea | Arthropoda | Crustacea |
| 1387 | Wenling crustacean virus 11 strain WLJQ201798 hypothetical protein 1 hypothetical protein 2 hypothetical protein | KX884456.1 | East China Sea | Arthropoda | Crustacea |
| 1388 | Beihai barnacle virus 9 strain BHTH10927 putative glycoprotein hypothetical protein and RNA-dependent RNA po | KX884409.1 | South China Sea | Arthropoda | Crustacea |
| 1389 | Wenling narna-like virus 1 strain WLJQ102793 RNA-dependent RNA polymerase gene complete cds | KX883607.1 | East China Sea | Arthropoda | Crustacea |
| 1390 | Hubei narna-like virus 16 strain fly55630 hypothetical protein gene partial cds | KX883526.1 | Terrestrial | Arthropoda | Insecta |
| 1391 | Beihai noda-like virus 23 strain BHHK113658 hypothetical protein gene complete cds | KX883025.1 | South China Sea | Cnidaria | Anthozoa |
| 1392 | Beihai picorna-like virus 108 strain BHTH16067 hypothetical protein gene complete cds | KX883414.1 | South China Sea | Arthropoda | Crustacea |
| 1393 | Beihai narna-like virus 24 strain BHXG26712 hypothetical protein and RNA-dependent RNA polymerase genes cor | KX883500.1 | South China Sea | Arthropoda | Crustacea |
| 1394 | Beihai weivirus-like virus 15 strain BWBFG27289 hypothetical protein 1 RNA-dependent RNA polymerase and hy | KX884021.1 | South China Sea | Mollusca | Bivalvia |
| 1395 | Wenling narna-like virus 8 strain WLJQ103992 RNA-dependent RNA polymerase gene partial cds and hypothetica | KX883605.1 | East China Sea | Arthropoda | Crustacea |
| 1396 | Wuhan insect virus 18 strain CC63583 RNA-dependent RNA polymerase gene complete cds | KX883516.1 | Terrestrial | Arthropoda | Insecta |
| 1397 | Wenling narna-like virus 7 strain WLJQ103612 RNA-dependent RNA polymerase and hypothetical protein genes c | KX883602.1 | East China Sea | Arthropoda | Crustacea |
| 1398 | Hubei qinvirus-like virus 1 strain SCM49583 hypothetical protein gene complete cds | KX883993.1 | Terrestrial | Arthropoda | Insecta |
| 1399 | Gastropoda Picorna-like virus N9 | GWHAPFP01000000 | South China Sea | Mollusca | Gastropoda |
| 1400 | Crustacea Picorna-like virus N32 | GWHAPDU01000000 | South China Sea | Arthropoda | Crustacea |
| 1401 | Gastropoda Picorna-like virus N10 | GWHAPFQ01000000 | South China Sea | Mollusca | Gastropoda |
| 1402 | Crustacea Ghabri-like virus N1 | GWHAPKW0100000( | South China Sea | Arthropoda | Crustacea |

| | | | | | |
|---|---|---|---|---|---|
| 1403 | Bivalvia Picorna-like virus H20 | GWHAPCS01000000 | Yellow China Sea | Mollusca | Bivalvia |
| 1404 | Wenzhou shrimp virus 4 strain WZRBX43306 hypothetical protein 1 and hypothetical protein 2 genes complete cds | KX884347.1 | East China Sea | Arthropoda | Crustacea |
| 1405 | Wenzhou shrimp virus 4 strain shrimp14546 hypothetical protein 1 and hypothetical protein 2 genes complete cds | KX883985.1 | East China Sea | Arthropoda | Crustacea |
| 1406 | Sanxia picorna-like virus 13 strain SXXX34014 hypothetical protein 1 and hypothetical protein 2 genes complete cc | KX883722.1 | Freshwater | Arthropoda | Crustacea |
| 1407 | Polychaeta Picorna-like virus H4 | GWHAPCW0100000( | Yellow China Sea | Annelida | Polychaeta |
| 1408 | Wenling partiti-like virus 10 strain WLJQ26001 RdRp gene complete cds | KX884224.1 | East China Sea | Arthropoda | Crustacea |
| 1409 | Crustacea Picorna-like virus N15 | GWHAPDB01000000 | South China Sea | Arthropoda | Crustacea |
| 1410 | Changjiang picorna-like virus 5 strain CJLX30750 hypothetical protein gene complete cds | KX884557.1 | Freshwater | Arthropoda | Crustacea |
| 1411 | Hubei coleoptera virus 6 strain QCM143720 RdRp gene complete cds | KX884108.1 | Terrestrial | Arthropoda | Insecta |
| 1412 | Tacheng Tick Virus 3 strain TC255 ORF1 (ORF1) and RNA-dependent RNA polymerase (L) genes complete cds | KM817640.1 | Terrestrial | Arthropoda | Chelicerata |
| 1413 | Beihai partiti-like virus 7 strain HOU343377 RdRp gene complete cds | KX884098.1 | South China Sea | Arthropoda | Chelicerata |
| 1414 | Tacheng Tick Virus 1 strain TC253 nucleopasid protein (N) gene complete cds | KM817743.1 | Terrestrial | Arthropoda | Chelicerata |
| 1415 | Beihai picorna-like virus 111 strain BHBei77071 hypothetical protein gene complete cds | KX883317.1 | South China Sea | Mollusca | Bivalvia |
| 1416 | Beihai picorna-like virus 111 strain BHZC37354 hypothetical protein gene complete cds | KX884490.1 | South China Sea | Mollusca | Bivalvia |
| 1417 | Beihai picorna-like virus 111 strain BHNXC41286 hypothetical protein gene complete cds | KX883394.1 | South China Sea | Sipuncula | Sipuncula |
| 1418 | Wuhan Mosquito Virus 4 strain XC3-4 PB1 (PB1) gene complete cds | KM817623.1 | Terrestrial | Arthropoda | Insecta |
| 1419 | Beihai picorna-like virus 73 strain BHHK131512 hypothetical protein gene complete cds | KX883352.1 | South China Sea | Cnidaria | Anthozoa |
| 1420 | Beihai picorna-like virus 73 strain BHJJX25960 hypothetical protein gene complete cds | KX883377.1 | South China Sea | Arthropoda | Crustacea |
| 1421 | Beihai picorna-like virus 73 strain BHTH16077 hypothetical protein gene complete cds | KX883415.1 | South China Sea | Arthropoda | Crustacea |
| 1422 | Wuhan Louse Fly Virus 2 strain BFJSC-11 nucleopasid protein (N) gene partial cds | KM817756.1 | Terrestrial | Arthropoda | Insecta |
| 1423 | Wuhan Mosquito Virus 4 strain 3mos5038 segment NP nucleocapsid protein gene complete cds | KX883842.1 | Terrestrial | Arthropoda | Insecta |
| 1424 | Wuhan Mosquito Virus 4 strain XC3-4 segment NP nucleocapsid protein gene complete cds | KX883867.1 | Terrestrial | Arthropoda | Insecta |
| 1425 | Wenzhou Crab Virus 2 strain ZCX13 segment 1 complete sequence | KM817601.1 | East China Sea | Arthropoda | Crustacea |
| 1426 | Wuhan spider virus 10 strain spider133290 RdRp gene complete cds | KX884164.1 | Terrestrial | Arthropoda | Chelicerata |
| 1427 | Wuhan spider virus 10 strain WHCCII13043 RdRp gene complete cds | KX884194.1 | Terrestrial | Arthropoda | Insecta |
| 1428 | Changjiang picorna-like virus 11 strain CJLX30672 hypothetical protein 1 and hypothetical protein 2 genes comple | KX884554.1 | Freshwater | Arthropoda | Crustacea |
| 1429 | Gastropoda Picorna-like virus N15 | GWHAPCC01000000 | South China Sea | Mollusca | Gastropoda |
| 1430 | Shahe arthropod virus 1 strain SHWC01c3692 hypothetical protein 1 and hypothetical protein 2 genes complete cds | KX883988.1 | Freshwater | Arthropoda | Insecta |
| 1431 | Shahe arthropod virus 1 strain SHWC13607 hypothetical protein 1 and hypothetical protein 2 genes complete cds | KX883662.1 | Freshwater | Arthropoda | Mix |
| 1432 | Shahe arthropod virus 1 strain SHWCII5326 hypothetical protein 1 and hypothetical protein 2 genes complete cds | KX883671.1 | Freshwater | Arthropoda | Mix |
| 1433 | Shahe arthropod virus 1 strain SHWC0209c12762 hypothetical protein 1 and hypothetical protein 2 genes complete | KX883641.1 | Freshwater | Arthropoda | Crustacea |
| 1434 | Sanxia Qinvirus-like virus 1 strain SXXX11646 hypothetical protein gene complete cds | KX883997.1 | Freshwater | Arthropoda | Crustacea |
| 1435 | Hubei partiti-like virus 2 strain arthropodmix12819 RdRp gene complete cds | KX884045.1 | Terrestrial | Arthropoda | Mix |
| 1436 | Hubei coleoptera virus 5 strain QCM105830 RdRp gene complete cds | KX884106.1 | Terrestrial | Arthropoda | Insecta |
| 1437 | Beihai razor shell virus 2 strain BHBei76898 hypothetical protein 1 and hypothetical protein 2 genes complete cds | KX883314.1 | South China Sea | Mollusca | Bivalvia |
| 1438 | Beihai razor shell virus 2 strain BHZC37307 hypothetical protein 1 and hypothetical protein 2 genes complete cds | KX884488.1 | South China Sea | Mollusca | Bivalvia |
| 1439 | Hubei partiti-like virus 42 strain spider134492 RdRp gene complete cds | KX884166.1 | Terrestrial | Arthropoda | Chelicerata |

| | | | | | |
|---|---|---|---|---|---|
| 1440 | Beihai picorna-like virus 107 strain BHZY60580 hypothetical protein gene complete cds | KX884508.1 | South China Sea | Mollusca | Cephalopoda |
| 1441 | Hubei coleoptera virus 4 strain QCM136227 RdRp gene complete cds | KX884107.1 | Terrestrial | Arthropoda | Insecta |
| 1442 | Beihai picorna-like virus 79 strain BHBJDX18844 hypothetical protein 1 and hypothetical protein 2 genes complete | KX883338.1 | South China Sea | Arthropoda | Crustacea |
| 1443 | Hubei partiti-like virus 37 strain QTM23479 RdRp gene complete cds | KX884118.1 | Terrestrial | Arthropoda | Insecta |
| 1444 | Hubei leech virus 2 strain SZmix21584 hypothetical protein 1 and hypothetical protein 2 genes complete cds | KX883726.1 | Freshwater | Annelida | Clitellata |
| 1445 | Hubei picorna-like virus 22 strain WHSF7472 hypothetical protein 1 and hypothetical protein 2 genes complete cds | KX884286.1 | Terrestrial | Arthropoda | Crustacea |
| 1446 | Bivalvia Amarillo-like virus N1 | GWHAPEV01000000 | South China Sea | Mollusca | Bivalvia |
| 1447 | Shuangao insect virus 7 strain SKC segment 4 complete sequence | KR902720.1 | Terrestrial | Arthropoda | Insecta |
| 1448 | Wuhan Mosquito Virus 7 strain XC1-5 PB1 (PB1) gene complete cds | KM817626.1 | Terrestrial | Arthropoda | Insecta |
| 1449 | Bivalvia Sobeli-like virus N2 | GWHAPHK01000000 | South China Sea | Mollusca | Bivalvia |
| 1450 | Beihai picorna-like virus 106 strain BHZY60875 hypothetical protein gene complete cds | KX884517.1 | South China Sea | Mollusca | Cephalopoda |
| 1451 | Shuangao insect virus 7 strain SKC segment 3 complete sequence | KR902719.1 | Terrestrial | Arthropoda | Insecta |
| 1452 | Bivalvia Picorna-like virus N12 | GWHAPLA01000000 | South China Sea | Mollusca | Bivalvia |
| 1453 | Wenzhou picorna-like virus 53 strain WZSLuoI86067 hypothetical protein 1 and hypothetical protein 2 genes comp | KX884373.1 | East China Sea | Mollusca | Gastropoda |
| 1454 | Beihai picorna-like virus 37 strain BHWZXX15436 hypothetical protein 1 gene complete cds and hypothetical prot | KX883426.1 | South China Sea | Arthropoda | Crustacea |
| 1455 | Changjiang crawfish virus 2 strain CJLX30764 hypothetical protein 1 and hypothetical protein 2 genes complete cd | KX884558.1 | Freshwater | Arthropoda | Crustacea |
| 1456 | Beihai partiti-like virus 6 strain BHZY110439 RdRp gene complete cds | KX884082.1 | South China Sea | Mollusca | Cephalopoda |
| 1457 | Beihai picorna-like virus 21 strain BHZY60822 hypothetical protein 1 and hypothetical protein 2 genes complete cc | KX884515.1 | South China Sea | Mollusca | Cephalopoda |
| 1458 | Banna virus strain QTM104536 Banna VP4 gene complete cds | KX884641.1 | Terrestrial | Arthropoda | Insecta |
| 1459 | Bivalvia Durna-like virus D3 | GWHAOYE01000000 | East China Sea | Mollusca | Bivalvia |
| 1460 | Beihai picobirna-like virus 13 strain BHZY58752 hypothetical protein gene complete cds and RdRp gene partial cd | KX884081.1 | South China Sea | Mollusca | Cephalopoda |
| 1461 | Changjiang tombus-like virus 18 strain CJLX21588 hypothetical protein 1 hypothetical protein 2 and hypothetical p | KX883083.1 | Freshwater | Arthropoda | Crustacea |
| 1462 | Changjiang narna-like virus 5 strain CJLX30442 RNA-dependent RNA polymerase gene partial cds | KX883519.1 | Freshwater | Arthropoda | Crustacea |
| 1463 | Wenzhou tombus-like virus 15 strain WZFSL70232 hypothetical protein and hypothetical protein 2 genes complete | KX883271.1 | Freshwater | Mollusca | Gastropoda |
| 1464 | Shahe levi-like virus 2 strain SHWC0209c10970 hypothetical protein genes complete cds | KX883561.1 | Freshwater | Arthropoda | Crustacea |
| 1465 | Wenzhou levi-like virus 3 strain WZSLuoI79939 hypothetical protein genes complete cds | KX883622.1 | East China Sea | Mollusca | Gastropoda |
| 1466 | Bivalvia Picorna-like virus D56 | GWHAPAB01000000 | East China Sea | Mollusca | Bivalvia |
| 1467 | Wuhan Louse Fly Virus 8 strain BFJSC-6 RNA-dependent RNA polymerase (L) gene complete cds | KM817655.1 | Terrestrial | Arthropoda | Insecta |
| 1468 | Jingshan Fly Virus 2 strain JSY-4 RNA-dependent RNA polymerase (L) gene partial cds | KM817631.1 | Terrestrial | Arthropoda | Insecta |
| 1469 | Hubei rhabdo-like virus 5 strain WHSFII19440 RNA-dependent RNA polymerase gene complete cds | KX884446.1 | Terrestrial | Arthropoda | Crustacea |
| 1470 | Hubei rhabdo-like virus 8 strain QTM19395 RNA-dependent RNA polymerase gene complete cds | KX884420.1 | Terrestrial | Arthropoda | Insecta |
| 1471 | Beihai picorna-like virus 97 strain BHZC37303 hypothetical protein gene partial cds | KX884487.1 | South China Sea | Mollusca | Bivalvia |
| 1472 | Hubei noda-like virus 8 strain WHLC5312 RNA-dependent RNA polymerase gene complete cds | KX883237.1 | Terrestrial | Arthropoda | Insecta |
| 1473 | Beihai picorna-like virus 101 strain BHBJDX18559 hypothetical protein 1 and hypothetical protein 2 genes comple | KX883329.1 | South China Sea | Arthropoda | Crustacea |
| 1474 | Hubei insect virus 2 strain arthropodmix13736 RNA-dependent RNA polymerase gene complete cds | KX884578.1 | Terrestrial | Arthropoda | Mix |
| 1475 | Sanxia picorna-like virus 6 strain SXXX37597 hypothetical protein gene partial cds | KX883713.1 | Freshwater | Arthropoda | Crustacea |
| 1476 | Hubei mosquito virus 5 strain mosHB233040 RdRp gene partial cds | KX884633.1 | Terrestrial | Arthropoda | Insecta |

| | | | | | |
|---|---|---|---|---|---|
| 1477 | Beihai picorna-like virus 103 strain BHBei77089 hypothetical protein gene complete cds | KX883318.1 | South China Sea | Mollusca | Bivalvia |
| 1478 | Hubei blood fluke virus 3 strain JSC69362 RNA-dependent RNA polymerase gene complete cds | KX884608.1 | Freshwater | Platyhelminthes | Trematoda |
| 1479 | Hubei blood fluke virus 3 strain RBXXC337523 RNA-dependent RNA polymerase gene complete cds | KX884687.1 | Freshwater | Platyhelminthes | Trematoda |
| 1480 | Beihai toti-like virus 4 strain HOU154008 hypothetical protein 1 and hypothetical protein 2 genes complete cds | KX882958.1 | South China Sea | Arthropoda | Chelicerata |
| 1481 | Hubei insect virus 2 strain arthropodmix13736 hypothetical protein 1 gene complete cds | KX884580.1 | Terrestrial | Arthropoda | Mix |
| 1482 | Hubei reo-like virus 8 strain mosHB236127 RdRp gene complete cds | KX884636.1 | Terrestrial | Arthropoda | Insecta |
| 1483 | Shahe levi-like virus 1 strain SHWC0209c12428 hypothetical protein genes complete cds | KX883557.1 | Freshwater | Arthropoda | Crustacea |
| 1484 | Bivalvia Unclassified virus D1 | GWHAOXZ01000000 | East China Sea | Mollusca | Bivalvia |
| 1485 | Gastropoda Picorna-like virus N14 | GWHAPCB01000000 | South China Sea | Mollusca | Gastropoda |
| 1486 | Gastropoda Sobeli-like virus D1 | GWHAPJW01000000 | East China Sea | Mollusca | Gastropoda |
| 1487 | Gastropoda Sobeli-like virus D2 | GWHAPAN01000000 | East China Sea | Mollusca | Gastropoda |
| 1488 | Bivalvia Sobeli-like virus N1 | GWHAPFC01000000 | South China Sea | Mollusca | Bivalvia |
| 1489 | Bivalvia Durna-like virus D4 | GWHAOYR01000000 | East China Sea | Mollusca | Bivalvia |
| 1490 | Beihai bunya-like virus 1 strain BHTSS17606 RNA-dependent RNA polymerase gene complete cds | KX884749.1 | South China Sea | Arthropoda | Crustacea |
| 1491 | Bivalvia Durna-like virus D6 | GWHAPES01000000 | East China Sea | Mollusca | Bivalvia |
| 1492 | Beihai hepe-like virus 4 strain BHZY60303 hypothetical protein genes complete cds | KX883756.1 | South China Sea | Mollusca | Cephalopoda |
| 1493 | Bivalvia Picorna-like virus N40 | GWHAPHP01000000 | South China Sea | Mollusca | Bivalvia |
| 1494 | Hubei diptera virus 7 strain SCM173098 RNA-dependent RNA polymerase gene complete cds | KX884807.1 | Terrestrial | Arthropoda | Insecta |
| 1495 | Wuhan Mosquito Virus 1 strain WT3-15 RNA-dependent RNA polymerase (L) gene complete cds | KM817697.1 | Terrestrial | Arthropoda | Insecta |
| 1496 | Hubei diptera virus 6 strain SCM95079 RNA-dependent RNA polymerase gene complete cds | KX884805.1 | Terrestrial | Arthropoda | Insecta |
| 1497 | Wuchang Cockraoch Virus 1 strain WCZL-5 RNA-dependent RNA polymerase (L) gene complete cds | KM817688.1 | Terrestrial | Arthropoda | Insecta |
| 1498 | Hubei odonate virus 9 strain QTM74767 RNA-dependent RNA polymerase gene complete cds | KX884786.1 | Terrestrial | Arthropoda | Insecta |
| 1499 | Hubei odonate virus 8 strain QTM19232 RNA-dependent RNA polymerase gene complete cds | KX884775.1 | Terrestrial | Arthropoda | Insecta |
| 1500 | Hubei bunya-like virus 1 strain SCM51446 RNA-dependent RNA polymerase gene complete cds | KX884804.1 | Terrestrial | Arthropoda | Insecta |
| 1501 | Wuhan ascaridia galli virus 1 strain WHJHC41208 RNA-dependent RNA polymerase gene complete cds | KX884847.1 | Terrestrial | Nematoda | Chromadorea |
| 1502 | Shayang ascaridia galli virus 1 strain HC21343 RNA-dependent RNA polymerase gene complete cds | KX884765.1 | Terrestrial | Nematoda | Rhabditida |
| 1503 | Shayang ascaridia galli virus 1 strain SYJFC11704 RNA-dependent RNA polymerase gene complete cds | KX884836.1 | Terrestrial | Nematoda | Chromadorea |
| 1504 | Jingmen bunya-like virus 1 strain JMYJCC72012 RNA-dependent RNA polymerase gene complete cds | KX884766.1 | Terrestrial | Nematoda | Chromadorea |
| 1505 | Xingshan nematode virus 3 strain XSNXC32933 RNA-dependent RNA polymerase gene complete cds | KX884867.1 | Terrestrial | Nematoda | Chromadorea |
| 1506 | Wuhan pillworm virus 1 strain WHSFII20185 hypothetical protein genes complete cds | KX884850.1 | Terrestrial | Arthropoda | Crustacea |
| 1507 | Shayang bunya-like virus 1 strain SYJFC48558 RNA-dependent RNA polymerase gene complete cds | KX884835.1 | Terrestrial | Nematoda | Chromadorea |
| 1508 | Wuhan Fly Virus 1 strain SYY1-9 glycoprotein precursor (G) gene complete cds | KM817722.1 | Terrestrial | Arthropoda | Insecta |
| 1509 | Hubei blood fluke virus 1 strain HBXXC56361 RNA-dependent RNA polymerase gene complete cds | KX884763.1 | Freshwater | Platyhelminthes | Trematoda |
| 1510 | Hubei blood fluke virus 1 strain JSC85519 RNA-dependent RNA polymerase gene complete cds | KX884770.1 | Freshwater | Platyhelminthes | Trematoda |
| 1511 | Hubei blood fluke virus 1 strain RBXXC303595 RNA-dependent RNA polymerase gene complete cds | KX884792.1 | Freshwater | Platyhelminthes | Trematoda |
| 1512 | Hubei toti-like virus 9 strain QTM27092 hypothetical protein gene complete cds | KX882967.1 | Terrestrial | Arthropoda | Insecta |
| 1513 | Lishi Spider Virus 1 strain LSZZ11 segment 2 complete sequence | KM817596.1 | Terrestrial | Arthropoda | Chelicerata |

| | | | | | |
|---|---|---|---|---|---|
| 1514 | Hubei hepe-like virus 3 strain WHYY20710 replicase putative capsid protein and hypothetical protein genes compl | KX883823.1 | Terrestrial | Arthropoda | Myriapoda |
| 1515 | Hubei lepidoptera virus 3 strain LCM141729 Cypovirus VP2 gene complete cds | KX884615.1 | Terrestrial | Arthropoda | Insecta |
| 1516 | Banna virus strain QTM104536 RNA-dependent RNA polymerase gene complete cds | KX884638.1 | Terrestrial | Arthropoda | Insecta |
| 1517 | Shuangao Insect Virus 6 strain QSA10 RNA-dependent RNA polymerase (L) gene partial cds | KM817639.1 | Terrestrial | Arthropoda | Insecta |
| 1518 | Shahe yuevirus-like virus 1 strain SHWC0209c11789 RNA-dependent RNA polymerase gene complete cds | KX884034.1 | Freshwater | Arthropoda | Crustacea |
| 1519 | Wuhan Fly Virus 1 strain SYY1-9 RNA-dependent RNA polymerase (L) gene complete cds | KM817689.1 | Terrestrial | Arthropoda | Insecta |
| 1520 | Hubei zhaovirus-like virus 1 strain WHBL52766 RNA-dependent RNA polymerase and putative capsid protein gen | KX884042.1 | Freshwater | Mollusca | Bivalvia |
| 1521 | Hubei insect virus 2 strain arthropodmix13736 Major core gene complete cds | KX884579.1 | Terrestrial | Arthropoda | Mix |
| 1522 | Beihai tombus-like virus 13 strain BHWZXX15674 hypothetical protein 1 hypothetical protein 2 and hypothetical p | KX883053.1 | South China Sea | Arthropoda | Crustacea |
| 1523 | Beihai tombus-like virus 13 strain BHZC37436 hypothetical protein 1 hypothetical protein 2 and hypothetical prote | KX883063.1 | South China Sea | Mollusca | Bivalvia |
| 1524 | Hubei reo-like virus 4 strain WGML144712 RNA-dependent RNA polymerase gene complete cds | KX884712.1 | Terrestrial | Arthropoda | Myriapoda |
| 1525 | Hubei picorna-like virus 10 strain QTM27287 hypothetical protein gene complete cds | KX883950.1 | Terrestrial | Arthropoda | Insecta |
| 1526 | Hubei reo-like virus 4 strain WGML144712 hypothetical protein 1 gene complete cds | KX884713.1 | Terrestrial | Arthropoda | Myriapoda |
| 1527 | Hubei picorna-like virus 82 strain spider133992 hypothetical protein 1 hypothetical protein 2 and hypothetical prote | KX883688.1 | Terrestrial | Arthropoda | Chelicerata |
| 1528 | Crustacea Bunya-like virus H1 | GWHAPGU01000000 | Yellow China Sea | Arthropoda | Crustacea |
| 1529 | Shuangao Insect Virus 3 strain QSA04 RNA-dependent RNA polymerase (L) gene complete cds | KM817681.1 | Terrestrial | Arthropoda | Insecta |
| 1530 | Wenling crustacean virus 7 strain WLJQ104070 RNA-dependent RNA polymerase gene complete cds | KX884859.1 | East China Sea | Arthropoda | Crustacea |
| 1531 | Wuhan horsefly Virus strain JJ2-1 RNA-dependent RNA polymerase (L) gene complete cds | KM817690.1 | Terrestrial | Arthropoda | Insecta |
| 1532 | Hubei diptera virus 5 strain SCM245062 RNA-dependent RNA polymerase gene complete cds | KX884799.1 | Terrestrial | Arthropoda | Insecta |
| 1533 | Wuhan Louse Fly Virus 1 strain BFJSC-10 RNA-dependent RNA polymerase (L) gene complete cds | KM817663.1 | Terrestrial | Arthropoda | Insecta |
| 1534 | Beihai sesarmid crab virus 5 strain SCJXSX36597 RNA-dependent RNA polymerase gene complete cds | KX884794.1 | South China Sea | Arthropoda | Crustacea |
| 1535 | Hubei lepidoptera virus 1 strain LCM141331 RNA-dependent RNA polymerase gene complete cds | KX884772.1 | Terrestrial | Arthropoda | Insecta |
| 1536 | Wuhan insect virus 16 strain arthropodmix13442 RNA-dependent RNA polymerase gene complete cds | KX884733.1 | Terrestrial | Arthropoda | Mix |
| 1537 | Wuhan insect virus 16 strain WHZM9931 RNA-dependent RNA polymerase gene complete cds | KX884854.1 | Terrestrial | Arthropoda | Insecta |
| 1538 | Beihai blue swimmer crab virus 2 strain YYSZX17850 RNA-dependent RNA polymerase gene complete cds | KX884870.1 | South China Sea | Arthropoda | Crustacea |
| 1539 | Sanxia Water Strider Virus 2 strain SXSSP02 RNA-dependent RNA polymerase (L) gene complete cds | KM817675.1 | Freshwater | Arthropoda | Insecta |
| 1540 | Shuangao Fly Virus 1 strain QSA06-1 polymerase gene complete cds and polymerase gene partial cds | KM817612.1 | Terrestrial | Arthropoda | Insecta |
| 1541 | Hubei bunya-like virus 3 strain QTM25853 RNA-dependent RNA polymerase gene complete cds | KX884778.1 | Terrestrial | Arthropoda | Insecta |
| 1542 | Wuhan Ant Virus strain WHMY02 RNA-dependent RNA polymerase (L) gene complete cds | KM817645.1 | Terrestrial | Arthropoda | Insecta |
| 1543 | Hubei chuvirus-like virus 4 strain SSZZ6973 RNA-dependent RNA polymerase gene partial cds | KX884438.1 | Freshwater | Arthropoda | Chelicerata |
| 1544 | Hubei lepidoptera virus 3 strain LCM141729 RNA-dependent RNA polymerase gene complete cds | KX884613.1 | Terrestrial | Arthropoda | Insecta |
| 1545 | Beihai picorna-like virus 65 strain BHZY60937 hypothetical protein gene complete cds | KX884523.1 | South China Sea | Mollusca | Cephalopoda |
| 1546 | Wuhan Insect virus 2 strain WHDL02 glycoprotein precursor (G) gene complete cds | KM817724.2 | Terrestrial | Arthropoda | Insecta |
| 1547 | Hubei bunya-like virus 10 strain QTM27070 putative glycoprotein gene complete cds | KX884783.1 | Terrestrial | Arthropoda | Insecta |
| 1548 | Beihai picorna-like virus 104 strain BHZC37407 hypothetical protein gene complete cds | KX884494.1 | South China Sea | Mollusca | Bivalvia |
| 1549 | Hubei lepidoptera virus 3 strain LCM141729 Cypovirus VP1 gene complete cds | KX884614.1 | Terrestrial | Arthropoda | Insecta |
| 1550 | Shayang Spider Virus 2 strain SYZZ-3 RNA-dependent RNA polymerase (L) gene complete cds | KM817677.1 | Terrestrial | Arthropoda | Chelicerata |

| | | | | | |
|---|---|---|---|---|---|
| 1551 | Hubei bunya-like virus 10 strain QTM27070 RNA-dependent RNA polymerase gene complete cds | KX884782.1 | Terrestrial | Arthropoda | Insecta |
| 1552 | Wuhan Insect virus 2 strain WHDL02 RNA-dependent RNA polymerase (L) gene complete cds | KM817692.1 | Terrestrial | Arthropoda | Insecta |
| 1553 | Gastropoda Picorna-like virus N12 | GWHAPCA01000000 | South China Sea | Mollusca | Gastropoda |
| 1554 | Beihai picorna-like virus 63 strain BHTSS17878 hypothetical protein 1 and hypothetical protein 2 genes complete c | KX883419.1 | South China Sea | Arthropoda | Crustacea |
| 1555 | Hubei virga-like virus 22 strain QCM140901 hypothetical protein gene complete cds | KX883777.1 | Terrestrial | Arthropoda | Insecta |
| 1556 | Beihai hepe-like virus 8 strain BHZY60925 replicase hypothetical protein and putative structural protein genes com | KX883759.1 | South China Sea | Mollusca | Cephalopoda |
| 1557 | Hubei picorna-like virus 35 strain QTM27260 hypothetical protein gene complete cds | KX883946.1 | Terrestrial | Arthropoda | Insecta |
| 1558 | Beihai tombus-like virus 9 strain BHZY58393 hypothetical protein 1 hypothetical protein 2 and hypothetical protein | KX883068.1 | South China Sea | Mollusca | Cephalopoda |
| 1559 | Hubei narna-like virus 9 strain WHBL72555 RNA-dependent RNA polymerase and hypothetical protein genes com | KX883579.1 | Freshwater | Mollusca | Bivalvia |
| 1560 | Beihai picorna-like virus 96 strain HOU158314 hypothetical protein gene complete cds | KX883901.1 | South China Sea | Arthropoda | Chelicerata |
| 1561 | Wuhan insect virus 13 strain CC64511 hypothetical protein gene complete cds | KX884537.1 | Terrestrial | Arthropoda | Insecta |
| 1562 | Hubei virga-like virus 11 strain fly114676 hypothetical protein genes complete cds | KX883766.1 | Terrestrial | Arthropoda | Insecta |
| 1563 | Hubei chuvirus-like virus 1 strain QTM26249 RNA-dependent RNA polymerase gene complete cds | KX884424.1 | Terrestrial | Arthropoda | Insecta |
| 1564 | Wenling zhaovirus-like virus 1 strain WLJQ103300 RNA-dependent RNA polymerase and hypothetical protein gen | KX884044.1 | East China Sea | Arthropoda | Crustacea |
| 1565 | Hubei coleoptera virus 1 strain QCM131202 hypothetical protein gene complete cds | KX883921.1 | Terrestrial | Arthropoda | Insecta |
| 1566 | Hubei picorna-like virus 16 strain SCM51353 hypothetical protein 1 and hypothetical protein 2 genes complete cds | KX883972.1 | Terrestrial | Arthropoda | Insecta |
| 1567 | Hubei hepe-like virus 2 strain WHCC101555 hypothetical protein genes complete cds | KX883808.1 | Terrestrial | Arthropoda | Insecta |
| 1568 | Hubei picorna-like virus 26 strain spider128568 hypothetical protein gene complete cds | KX883675.1 | Terrestrial | Arthropoda | Chelicerata |
| 1569 | Hubei diptera virus 1 strain SCM51289 hypothetical protein 1 and hypothetical protein 2 genes complete cds | KX883971.1 | Terrestrial | Arthropoda | Insecta |
| 1570 | Hubei odonate virus 14 strain QTM27275 RNA-dependent RNA polymerase gene complete cds | KX884683.1 | Terrestrial | Arthropoda | Insecta |
| 1571 | Changjiang crawfish virus 4 strain CLX8819 hypothetical protein gene complete cds | KX883889.1 | Freshwater | Arthropoda | Crustacea |
| 1572 | Hubei reo-like virus 6 strain WGML611819 hypothetical protein 1 gene complete cds | KX884717.1 | Terrestrial | Arthropoda | Myriapoda |
| 1573 | Hubei picorna-like virus 8 strain SCM51412 hypothetical protein gene complete cds | KX883973.1 | Terrestrial | Arthropoda | Insecta |
| 1574 | Hubei picorna-like virus 8 strain QTM27288 hypothetical protein 1 and hypothetical protein 2 genes complete cds | KX883951.1 | Terrestrial | Arthropoda | Insecta |
| 1575 | Changjiang tombus-like virus 21 strain CJLX8746 hypothetical protein 1 and hypothetical protein 2 genes complete | KX883081.1 | Freshwater | Arthropoda | Crustacea |
| 1576 | Beihai picorna-like virus 102 strain BHZC35144 hypothetical protein 1 gene complete cds | KX884475.1 | South China Sea | Mollusca | Bivalvia |
| 1577 | Hubei reo-like virus 6 strain WGML611819 RNA-dependent RNA polymerase gene complete cds | KX884716.1 | Terrestrial | Arthropoda | Myriapoda |
| 1578 | Hubei reo-like virus 11 strain arthropodmix13320 RNA-dependent RNA polymerase gene complete cds | KX884574.1 | Terrestrial | Arthropoda | Mix |
| 1579 | Hubei reo-like virus 11 strain QTM23921 RNA-dependent RNA polymerase gene complete cds | KX884671.1 | Terrestrial | Arthropoda | Insecta |
| 1580 | Hubei reo-like virus 1 strain QCM786512 RdRp gene complete cds | KX884637.1 | Terrestrial | Arthropoda | Insecta |
| 1581 | Wenzhou channeled applesnail virus 3 strain WZFSL85549 hypothetical protein gene complete cds | KX884327.1 | Freshwater | Mollusca | Gastropoda |
| 1582 | Wenzhou channeled applesnail virus 3 strain BHBei77067 hypothetical protein gene complete cds | KX883316.1 | South China Sea | Mollusca | Bivalvia |
| 1583 | Wenzhou channeled applesnail virus 3 strain BHZC37337 hypothetical protein gene complete cds | KX884489.1 | South China Sea | Mollusca | Bivalvia |
| 1584 | Wenzhou channeled applesnail virus 3 strain BHWZXX15670 hypothetical protein gene complete cds | KX883432.1 | South China Sea | Arthropoda | Crustacea |
| 1585 | Hubei insect virus 2 strain arthropodmix13736 Inner core protein gene complete cds | KX884582.1 | Terrestrial | Arthropoda | Mix |
| 1586 | Kadipiro virus strain QTM27331 Kadipiro VP2 gene complete cds | KX884651.1 | Terrestrial | Arthropoda | Insecta |
| 1587 | Wenzhou picorna-like virus 39 strain WZFSL83776 hypothetical protein gene complete cds | KX884332.1 | Freshwater | Mollusca | Gastropoda |

| | | | | | |
|---|---|---|---|---|---|
| 1588 | Beihai hepe-like virus 2 strain BHHZL10373 RdRp gene complete cds | KX883745.1 | South China Sea | Arthropoda | Crustacea |
| 1589 | Hubei tombus-like virus 28 strain SSZZ1762 hypothetical protein and RNA-dependent RNA polymerase genes com | KX883168.1 | Freshwater | Arthropoda | Chelicerata |
| 1590 | Sanxia tombus-like virus 8 strain SXSSP2544 hypothetical protein 1 and hypothetical protein 2 genes complete cds | KX883173.1 | Freshwater | Arthropoda | Insecta |
| 1591 | Shuangao alphatetra-like virus 1 strain insectZJ93971 replicase putative structural protein and hypothetical protein g | KX883771.1 | Terrestrial | Arthropoda | Insecta |
| 1592 | Wuchan romanomermis nematode virus 2 strain WCLSXC55347 hypothetical protein 1 hypothetical protein 2 hypo | KX884441.1 | Terrestrial | Nematoda | Enoplea |
| 1593 | Hubei insect virus 2 strain arthropodmix13736 hypothetical protein 7 gene complete cds | KX884586.1 | Terrestrial | Arthropoda | Mix |
| 1594 | Hubei reo-like virus 9 strain QTM27922 RdRp gene complete cds | KX884686.1 | Terrestrial | Arthropoda | Insecta |
| 1595 | Crustacea Picorna-like virus N20 | GWHAPDF01000000 | South China Sea | Arthropoda | Crustacea |
| 1596 | Xinzhou bunya-like virus 2 strain XZSJSC64965 RNA-dependent RNA polymerase gene complete cds | KX884869.1 | Terrestrial | Nematoda | Mix |
| 1597 | Xinzhou bunya-like virus 1 strain XZSJSC18544 RNA-dependent RNA polymerase gene complete cds | KX884868.1 | Terrestrial | Nematoda | Mix |
| 1598 | Beihai sobemo-like virus 2 strain BHZC35241 hypothetical protein 1 and hypothetical protein 2 genes complete cds | KX882796.1 | South China Sea | Mollusca | Bivalvia |
| 1599 | Hubei sobemo-like virus 31 strain WHCC117732 hypothetical protein 1 and hypothetical protein 2 genes complete | KX882908.1 | Terrestrial | Arthropoda | Insecta |
| 1600 | Hubei partiti-like virus 14 strain spider132884 RdRp gene complete cds | KX884163.1 | Terrestrial | Arthropoda | Chelicerata |
| 1601 | Wenzhou bivalvia virus 3 strain beimix38484 hypothetical protein 1 and hypothetical protein 2 genes complete cds | KX883014.1 | East China Sea | Mollusca | Bivalvia |
| 1602 | Changjiang sobemo-like virus 1 strain CJLX29826 hypothetical protein 1 hypothetical protein 2 and hypothetical pr | KX882816.1 | Freshwater | Arthropoda | Crustacea |
| 1603 | Sanxia sobemo-like virus 1 strain SXSSP20269 hypothetical protein 1 hypothetical protein 2 and hypothetical prote | KX882885.1 | Freshwater | Arthropoda | Insecta |
| 1604 | Beihai levi-like virus 28 strain BHXG26591 hypothetical protein genes complete cds | KX883499.1 | South China Sea | Arthropoda | Crustacea |
| 1605 | Beihai levi-like virus 29 strain BHZY57867 hypothetical protein genes complete cds | KX883511.1 | South China Sea | Mollusca | Cephalopoda |
| 1606 | Wuhan Millipede virus 4 strain GCM8225 segment Seg 1 RdRp gene complete cds | KX884093.1 | Terrestrial | Arthropoda | Insecta |
| 1607 | Wuhan Millipede virus 4 strain WHWG256022 segment Seg 1 RdRp gene complete cds | KX884202.1 | Terrestrial | Arthropoda | Myriapoda |
| 1608 | Shuangao chryso-like virus 1 strain insectZJ85542 hypothetical protein gene partial cds | KX882959.1 | Terrestrial | Arthropoda | Insecta |
| 1609 | Shuangao chryso-like virus 1 strain mosZJ34956 hypothetical protein gene partial cds | KX882964.1 | Terrestrial | Arthropoda | Insecta |
| 1610 | Hubei chryso-like virus 1 strain mosHB233224 hypothetical protein gene partial cds | KX882962.1 | Terrestrial | Arthropoda | Insecta |
| 1611 | Citrus tristeza virus strain WHWN54444 RNA-dependent RNA polymerase gene partial cds and p33 protein p65 pr | KX883820.1 | Freshwater | Mollusca | Gastropoda |
| 1612 | Hubei sobemo-like virus 44 strain WGML133391 hypothetical protein 1 and hypothetical protein 2 genes complete | KX882901.1 | Terrestrial | Arthropoda | Myriapoda |
| 1613 | Beihai tombus-like virus 3 strain BHZY54477 hypothetical protein 1 hypothetical protein 2 and hypothetical proteir | KX883067.1 | South China Sea | Mollusca | Cephalopoda |
| 1614 | Beihai tombus-like virus 2 strain BHNXC41455 hypothetical protein 1 and hypothetical protein 2 genes complete c | KX883035.1 | South China Sea | Sipuncula | Sipuncula |
| 1615 | Hubei chryso-like virus 2 strain SCM47694 hypothetical protein gene complete cds | KX882973.1 | Terrestrial | Arthropoda | Insecta |
| 1616 | Changjiang tombus-like virus 10 strain CJLX21891 hypothetical protein 1 hypothetical protein 2 and hypothetical p | KX883084.1 | Freshwater | Arthropoda | Crustacea |
| 1617 | Beihai tombus-like virus 18 strain BHZY60611 hypothetical protein 1 and hypothetical protein 3 genes complete cd | KX883074.1 | South China Sea | Mollusca | Cephalopoda |
| 1618 | Hubei diptera virus 16 strain SCM32007 hypothetical protein gene complete cds | KX883145.1 | Terrestrial | Arthropoda | Insecta |
| 1619 | Changjiang tombus-like virus 1 strain CJLX26263 hypothetical protein 1 and hypothetical protein 2 genes complete | KX883086.1 | Freshwater | Arthropoda | Crustacea |
| 1620 | Sanxia tombus-like virus 3 strain SXXX37381 hypothetical protein 1 and hypothetical protein 2 genes complete cds | KX883184.1 | Freshwater | Arthropoda | Crustacea |
| 1621 | Bivalvia Picorna-like virus D23 | GWHAOYU0100000C | East China Sea | Mollusca | Bivalvia |
| 1622 | Crustacea Picorna-like virus D3 | GWHAPBQ01000000 | East China Sea | Arthropoda | Crustacea |
| 1623 | Sanxia water strider virus 17 strain SXSSP1704 hypothetical protein 1 and hypothetical protein 2 genes complete cc | KX883174.1 | Freshwater | Arthropoda | Insecta |
| 1624 | Beihai weivirus-like virus 16 strain BHZY60618 RNA-dependent RNA polymerase and hypothetical protein genes | KX884011.1 | South China Sea | Mollusca | Cephalopoda |

| | | | | | |
|---|---|---|---|---|---|
| 1625 | Beihai weivirus-like virus 12 strain BWBFG40815 RNA-dependent RNA polymerase and hypothetical protein gene | KX884023.1 | South China Sea | Mollusca | Bivalvia |
| 1626 | Wenzhou sobemo-like virus 1 strain WZFSL61418 hypothetical protein gene complete cds | KX882931.1 | Freshwater | Mollusca | Gastropoda |
| 1627 | Wuhan insect virus 29 strain WHZM9836 hypothetical protein gene complete cds | KX882987.1 | Terrestrial | Arthropoda | Insecta |
| 1628 | Wuhan insect virus 26 strain WHZM10161 hypothetical protein 1 and hypothetical protein 2 genes complete cds | KX882983.1 | Terrestrial | Arthropoda | Insecta |
| 1629 | Beihai victori-like virus 1 strain BHZC37363 hypothetical protein 1 and hypothetical protein 2 genes complete cds | KX882948.1 | South China Sea | Mollusca | Bivalvia |
| 1630 | Hubei sobemo-like virus 2 strain WGML145690 hypothetical protein 1 hypothetical protein 2 and hypothetical prot | KX882902.1 | Terrestrial | Arthropoda | Myriapoda |
| 1631 | Beihai noda-like virus 24 strain BHBJDX12560 hypothetical protein gene complete cds | KX883016.1 | South China Sea | Arthropoda | Crustacea |
| 1632 | Wenzhou noda-like virus 4 strain WZFSL75166 hypothetical protein 1 and hypothetical protein 2 genes complete c | KX883269.1 | Freshwater | Mollusca | Gastropoda |
| 1633 | Beihai noda-like virus 3 strain BHZY60087 hypothetical protein 1 and hypothetical protein 2 genes complete cds | KX883065.1 | South China Sea | Mollusca | Cephalopoda |
| 1634 | Beihai noda-like virus 18 strain YYSZX13070 hypothetical protein gene complete cds | KX883282.1 | South China Sea | Arthropoda | Crustacea |
| 1635 | Beihai sobemo-like virus 3 strain BHZY60830 hypothetical protein 1 and hypothetical protein 2 genes complete cds | KX882807.1 | South China Sea | Mollusca | Cephalopoda |
| 1636 | Wenzhou levi-like virus 6 strain WZSLuoI133770 hypothetical protein genes complete cds | KX883620.1 | East China Sea | Mollusca | Gastropoda |
| 1637 | Wenling levi-like virus 2 strain WLJQ100415 hypothetical protein genes complete cds | KX883597.1 | East China Sea | Arthropoda | Crustacea |
| 1638 | Wenling levi-like virus 3 strain WLJQ101576 hypothetical protein genes complete cds | KX883598.1 | East China Sea | Arthropoda | Crustacea |
| 1639 | Changjiang levi-like virus 1 strain CJLX30166 hypothetical protein genes complete cds | KX883523.1 | Freshwater | Arthropoda | Crustacea |
| 1640 | Hubei levi-like virus 6 strain GCM8094 hypothetical protein genes complete cds | KX883527.1 | Terrestrial | Arthropoda | Insecta |
| 1641 | Hubei sobemo-like virus 22 strain ZCM21279 hypothetical protein 1 and hypothetical protein 2 genes complete cds | KX882937.1 | Terrestrial | Arthropoda | Insecta |
| 1642 | Wuhan house centipede virus 4 strain WHYY19909 segment 1 hypothetical protein 1 and hypothetical protein 2 ger | KX882923.1 | Terrestrial | Arthropoda | Myriapoda |
| 1643 | Wuhan house centipede virus 4 strain arthropodmix8296 segment 1 hypothetical protein genes complete cds | KX882778.1 | Terrestrial | Arthropoda | Mix |
| 1644 | Wenling tombus-like virus 3 strain WLJQ76752 hypothetical protein 1 hypothetical protein 2 and hypothetical prote | KX883255.1 | East China Sea | Arthropoda | Crustacea |
| 1645 | Sanxia water strider virus 10 strain SXSSP2571 segment 1 hypothetical protein 1 and hypothetical protein 2 genes c | KX882889.1 | Freshwater | Arthropoda | Insecta |
| 1646 | Sanxia water strider virus 10 strain SXXX37662 hypothetical protein 1 and hypothetical protein 2 genes complete c | KX882893.1 | Freshwater | Arthropoda | Crustacea |
| 1647 | Wenzhou tombus-like virus 7 strain WZFSL44625 hypothetical protein 1 and hypothetical protein 2 genes complete | KX883274.1 | Freshwater | Mollusca | Gastropoda |
| 1648 | Hubei diptera virus 12 strain SCM43654 segment 1 hypothetical protein 1 and hypothetical protein 2 genes complet | KX882853.1 | Terrestrial | Arthropoda | Insecta |
| 1649 | Shayang noda-like virus 1 strain SYJFC44697 hypothetical protein gene complete cds | KX883189.1 | Terrestrial | Nematoda | Chromadorea |
| 1650 | Hubei noda-like virus 18 strain HC16588 hypothetical protein gene complete cds | KX883110.1 | Terrestrial | Nematoda | Rhabditida |
| 1651 | Hubei noda-like virus 17 strain HC30049 hypothetical protein gene complete cds | KX883109.1 | Terrestrial | Nematoda | Rhabditida |
| 1652 | Hubei tombus-like virus 32 strain WHBL71139 hypothetical protein 1 and hypothetical protein 2 genes complete cc | KX883213.1 | Freshwater | Mollusca | Bivalvia |
| 1653 | Beihai sphaeromadae virus 2 strain BHTSS26432 hypothetical protein gene complete cds | KX883042.1 | South China Sea | Arthropoda | Crustacea |
| 1654 | Wuhan heteroptera virus 2 strain WHYCM467 segment 1 hypothetical protein 1 and hypothetical protein 2 genes cc | KX882917.1 | Terrestrial | Arthropoda | Insecta |
| 1655 | Wuhan heteroptera virus 2 strain arthropodmix10101 segment 1 hypothetical protein genes complete cds | KX882767.1 | Terrestrial | Arthropoda | Mix |
| 1656 | Wuhan heteroptera virus 2 strain SSZZ3534 segment 1 hypothetical protein 1 and hypothetical protein 2 genes com | KX882880.1 | Freshwater | Arthropoda | Chelicerata |
| 1657 | Sanxia water strider virus 14 strain SXSSP2565 hypothetical protein 1 hypothetical protein 2 hypothetical protein 3 | KX883176.1 | Freshwater | Arthropoda | Insecta |
| 1658 | Sanxia water strider virus 14 strain SXXX37453 hypothetical protein 1 hypothetical protein 2 hypothetical protein 3 | KX883183.1 | Freshwater | Arthropoda | Crustacea |
| 1659 | Crustacea Picorna-like virus N14 | GWHAPCZ01000000 | South China Sea | Arthropoda | Crustacea |
| 1660 | Wenzhou Nido-like virus 1 strain shrimp13441 RNA-dependent RNA polymerase gene partial cds | KX883631.1 | East China Sea | Arthropoda | Crustacea |
| 1661 | Hubei lepidoptera virus 5 strain LCM63289 hypothetical protein 3 gene complete cds | KX884626.1 | Terrestrial | Arthropoda | Insecta |

| | | | | | |
|---|---|---|---|---|---|
| **1662** | Hubei noda-like virus 6 strain SSZZ5803 hypothetical protein gene complete cds | KX883167.1 | Freshwater | Arthropoda | Chelicerata |
| **1663** | Beihai sesarmid crab virus 4 strain SCJXSX38901 hypothetical protein gene complete cds | KX883991.1 | South China Sea | Arthropoda | Crustacea |
| **1664** | Hubei zhaovirus-like virus 2 strain WHBL71389 RNA-dependent RNA polymerase and hypothetical protein genes | KX884043.1 | Freshwater | Mollusca | Bivalvia |
| **1665** | Beihai sesarmid crab virus 4 strain SCJXSX38901 RNA-dependent RNA polymerase gene complete cds | KX883990.1 | South China Sea | Arthropoda | Crustacea |
| **1666** | Wuhan insect virus 35 strain WHCCII10556 hypothetical protein 1 and hypothetical protein 2 genes complete cds | KX883233.1 | Terrestrial | Arthropoda | Insecta |
| **1667** | Changjiang tombus-like virus 20 strain CJLX30731 hypothetical protein 1 and hypothetical protein 2 genes complet | KX883100.1 | Freshwater | Arthropoda | Crustacea |
| **1668** | Hubei noda-like virus 9 strain WHLC5367 RNA-dependent RNA polymerase gene complete cds | KX883235.1 | Terrestrial | Arthropoda | Insecta |
| **1669** | Hubei noda-like virus 9 strain QTM26368 RNA-dependent RNA polymerase gene complete cds | KX883130.1 | Terrestrial | Arthropoda | Insecta |
| **1670** | Beihai narna-like virus 12 strain BHZC35702 RNA-dependent RNA polymerase gene complete cds | KX883505.1 | South China Sea | Mollusca | Bivalvia |
| **1671** | Hubei tombus-like virus 36 strain WGML125257 hypothetical protein 1 and hypothetical protein 2 genes complete | KX883191.1 | Terrestrial | Arthropoda | Myriapoda |
| **1672** | Beihai narna-like virus 5 strain BHNXC41323 RNA-dependent RNA polymerase gene partial cds and hypothetical | KX883478.1 | South China Sea | Sipuncula | Sipuncula |
| **1673** | Hubei tombus-like virus 37 strain WHBL72297 hypothetical protein gene complete cds | KX883222.1 | Freshwater | Mollusca | Bivalvia |
| **1674** | Wenzhou sobemo-like virus 3 strain mosZJ35256 hypothetical protein 1 hypothetical protein 2 and hypothetical pro | KX882830.1 | Terrestrial | Arthropoda | Insecta |
| **1675** | Shahe sobemo-like virus 1 strain SHWCII5101 hypothetical protein 1 and hypothetical protein 2 genes complete cd | KX882862.1 | Freshwater | Arthropoda | Mix |
| **1676** | Beihai barnacle virus 14 strain BHTH35712 hypothetical protein and RdRp genes complete cds | KX884071.1 | South China Sea | Arthropoda | Crustacea |
| **1677** | Hubei tombus-like virus 22 strain QTM26784 hypothetical protein 1 hypothetical protein 2 and hypothetical protein | KX883135.1 | Terrestrial | Arthropoda | Insecta |
| **1678** | Hubei tombus-like virus 40 strain QCM118690 hypothetical protein 1 hypothetical protein 2 and hypothetical protei | KX883127.1 | Terrestrial | Arthropoda | Insecta |
| **1679** | Hubei sobemo-like virus 45 strain QTM25680 hypothetical protein 1 and hypothetical protein 2 genes complete cds | KX882841.1 | Terrestrial | Arthropoda | Insecta |
| **1680** | Changjiang tombus-like virus 11 strain CJLX30677 hypothetical protein 1 hypothetical protein 2 and hypothetical p | KX883104.1 | Freshwater | Arthropoda | Crustacea |
| **1681** | Wenzhou tombus-like virus 5 strain WZFSL67420 hypothetical protein 1 hypothetical protein 2 and hypothetical pr | KX883272.1 | Freshwater | Mollusca | Gastropoda |
| **1682** | Changjiang tombus-like virus 8 strain CJLX29920 hypothetical protein 1 hypothetical protein 2 and hypothetical pr | KX883090.1 | Freshwater | Arthropoda | Crustacea |
| **1683** | Hubei noda-like virus 13 strain spider128934 hypothetical protein gene complete cds | KX883162.1 | Terrestrial | Arthropoda | Chelicerata |
| **1684** | Hubei noda-like virus 13 strain QTM161985 hypothetical protein gene complete cds | KX883139.1 | Terrestrial | Arthropoda | Insecta |
| **1685** | Beihai noda-like virus 5 strain BHZC36307 hypothetical protein 1 gene complete cds and hypothetical protein 2 ger | KX883059.1 | South China Sea | Mollusca | Bivalvia |
| **1686** | Beihai noda-like virus 5 strain HOU157766 hypothetical protein 1 and hypothetical protein 2 genes complete cds | KX883112.1 | South China Sea | Arthropoda | Chelicerata |
| **1687** | Beihai shrimp virus 6 strain BHWZXX15644 RNA-dependent RNA polymerase gene complete cds | KX883050.1 | South China Sea | Arthropoda | Crustacea |
| **1688** | Beihai shrimp virus 6 strain BHBJDX18793 RNA-dependent RNA polymerase gene complete cds | KX883022.1 | South China Sea | Arthropoda | Crustacea |
| **1689** | Hubei sobemo-like virus 8 strain QTM19779 hypothetical protein 1 and hypothetical protein 2 genes complete cds | KX882839.1 | Terrestrial | Arthropoda | Insecta |
| **1690** | Hubei mosquito virus 2 strain 3mos6212 segment 1 hypothetical protein 1 and RdRp genes complete cds | KX882764.1 | Terrestrial | Arthropoda | Insecta |
| **1691** | Hubei mosquito virus 2 strain spider133708 segment 1 hypothetical protein 1 and hypothetical protein 2 genes com | KX882873.1 | Terrestrial | Arthropoda | Chelicerata |
| **1692** | Hubei mosquito virus 2 strain mosZJ35453 segment 1 hypothetical protein 1 and hypothetical protein 2 genes comp | KX882832.1 | Terrestrial | Arthropoda | Insecta |
| **1693** | Wenzhou sobemo-like virus 4 strain mosZJ35391 hypothetical protein 1 and hypothetical protein 2 genes complete | KX882831.1 | Terrestrial | Arthropoda | Insecta |
| **1694** | Wuhan arthropod virus 4 strain WHCC117028 segment 1 hypothetical protein 1 and hypothetical protein 2 genes cc | KX882906.1 | Terrestrial | Arthropoda | Insecta |
| **1695** | Hubei tombus-like virus 25 strain WHBL69613 hypothetical protein 1 hypothetical protein 2 and hypothetical prote | KX883212.1 | Freshwater | Mollusca | Bivalvia |
| **1696** | Hubei narna-like virus 23 strain SZmix21602 RNA-dependent RNA polymerase gene complete cds | KX883574.1 | Freshwater | Annelida | Clitellata |
| **1697** | Beihai narna-like virus 18 strain HOU157597 RNA-dependent RNA polymerase gene complete cds | KX883533.1 | South China Sea | Arthropoda | Chelicerata |
| **1698** | Wenzhou tombus-like virus 12 strain WZFSL84432 hypothetical protein 1 hypothetical protein 2 hypothetical prote | KX883262.1 | Freshwater | Mollusca | Gastropoda |

| | | | | | |
|---|---|---|---|---|---|
| 1699 | Hubei sobemo-like virus 30 strain QTM19610 hypothetical protein 1 and hypothetical protein 2 genes complete cds | KX882838.1 | Terrestrial | Arthropoda | Insecta |
| 1700 | Hubei narna-like virus 25 strain SCM51430 RNA-dependent RNA polymerase gene complete cds | KX883546.1 | Terrestrial | Arthropoda | Insecta |
| 1701 | Beihai narna-like virus 17 strain SCJXSX39297 RNA-dependent RNA polymerase gene complete cds | KX883544.1 | South China Sea | Arthropoda | Crustacea |
| 1702 | Hubei sobemo-like virus 29 strain QTM12808 hypothetical protein 1 and hypothetical protein 2 genes complete cds | KX882837.1 | Terrestrial | Arthropoda | Insecta |
| 1703 | Sanxia water strider virus 15 strain SXSSP1894 hypothetical protein 1 and hypothetical protein 2 genes complete cc | KX883175.1 | Freshwater | Arthropoda | Insecta |
| 1704 | Sanxia water strider virus 15 strain SXXX18977 hypothetical protein 1 and hypothetical protein 2 genes complete c | KX883187.1 | Freshwater | Arthropoda | Crustacea |
| 1705 | Wuhan spider virus 9 strain spider112003 hypothetical protein 1 hypothetical protein 2 hypothetical protein 3 and h | KX883158.1 | Terrestrial | Arthropoda | Chelicerata |
| 1706 | Hubei tombus-like virus 18 strain QTM27278 hypothetical protein 1 hypothetical protein 2 hypothetical protein 3 ar | KX883138.1 | Terrestrial | Arthropoda | Insecta |
| 1707 | Wenling tombus-like virus 2 strain WLJQ104302 hypothetical protein 1 hypothetical protein 2 hypothetical protein | KX883254.1 | East China Sea | Arthropoda | Crustacea |
| 1708 | Hubei insect virus 2 strain arthropodmix13736 hypothetical protein 2 gene complete cds | KX884581.1 | Terrestrial | Arthropoda | Mix |
| 1709 | Beihai hermit crab virus 2 strain BHJJX25757 putative glycoprotein gene complete cds | KX884742.1 | South China Sea | Arthropoda | Crustacea |
| 1710 | Beihai sobemo-like virus 25 strain BHZC34084 hypothetical protein 1 hypothetical protein 2 and hypothetical prote | KX882797.1 | South China Sea | Mollusca | Bivalvia |
| 1711 | Hubei diptera virus 13 strain SCM94998 segment 1 hypothetical protein 1 and RdRp genes complete cds | KX882858.1 | Terrestrial | Arthropoda | Insecta |
| 1712 | Hubei diptera virus 13 strain arthropodmix8005 segment 1 hypothetical protein genes complete cds | KX882776.1 | Terrestrial | Arthropoda | Mix |
| 1713 | Hubei sobemo-like virus 20 strain CC64562 hypothetical protein 1 and hypothetical protein 2 genes complete cds | KX882814.1 | Terrestrial | Arthropoda | Insecta |
| 1714 | Hubei sobemo-like virus 28 strain QTM26976 hypothetical protein 1 and hypothetical protein 2 genes complete cds | KX882845.1 | Terrestrial | Arthropoda | Insecta |
| 1715 | Beihai levi-like virus 2 strain BHTSS13603 hypothetical protein genes complete cds | KX883486.1 | South China Sea | Arthropoda | Crustacea |
| 1716 | Beihai sobemo-like virus 16 strain BHCL80925 hypothetical protein 1 and hypothetical protein 2 genes complete cc | KX882786.1 | South China Sea | Mollusca | Gastropoda |
| 1717 | Hubei polero-like virus 1 strain WHCC118254 hypothetical protein 1 hypothetical protein 2 and hypothetical protei | KX882909.1 | Terrestrial | Arthropoda | Insecta |
| 1718 | Beihai noda-like virus 28 strain BHBJDX17825 hypothetical protein gene complete cds | KX883017.1 | South China Sea | Arthropoda | Crustacea |
| 1719 | Beihai noda-like virus 28 strain BHWZXX14359 hypothetical protein gene complete cds | KX883046.1 | South China Sea | Arthropoda | Crustacea |
| 1720 | Beihai tombus-like virus 15 strain BHBJDX18752 hypothetical protein 1 and hypothetical protein 2 genes complete | KX883020.1 | South China Sea | Arthropoda | Crustacea |
| 1721 | Hubei alphaflexi-like virus strain WHCC96629 hypothetical protein gene complete cds and hypothetical protein ger | KX883811.1 | Terrestrial | Arthropoda | Insecta |
| 1722 | Hubei tombus-like virus 16 strain spider58816 hypothetical protein 1 hypothetical protein 2 and hypothetical protei | KX883165.1 | Terrestrial | Arthropoda | Chelicerata |
| 1723 | Hubei tombus-like virus 4 strain WGML147612 hypothetical protein 1 hypothetical protein 2 and hypothetical prote | KX883201.1 | Terrestrial | Arthropoda | Myriapoda |
| 1724 | Wenzhou noda-like virus 5 strain WZFSL73727 hypothetical protein gene partial cds | KX883270.1 | Freshwater | Mollusca | Gastropoda |
| 1725 | Beihai narna-like virus 10 strain BHZY60512 RNA-dependent RNA polymerase and hypothetical protein genes cor | KX883510.1 | South China Sea | Mollusca | Cephalopoda |
| 1726 | Hubei sobemo-like virus 16 strain WGML146458 hypothetical protein 1 and hypothetical protein 2 genes complete | KX882903.1 | Terrestrial | Arthropoda | Myriapoda |
| 1727 | Hexanauplia Picorna-like virus D1 | GWHAPBX01000000 | East China Sea | Arthropoda | Hexanauplia |
| 1728 | Hubei toti-like virus 1 strain ZL17075 hypothetical protein gene partial cds | KX882998.1 | Terrestrial | Arthropoda | Insecta |
| 1729 | Bivalvia Picorna-like virus N9 | GWHAPGH01000000 | South China Sea | Mollusca | Bivalvia |
| 1730 | Hubei permutotetra-like virus 2 strain WHYY23891 RdRp and hypothetical protein genes complete cds | KX883455.1 | Terrestrial | Arthropoda | Myriapoda |
| 1731 | Hubei permutotetra-like virus 2 strain QTM26624 RdRp hypothetical protein 1 and hypothetical protein 2 genes cor | KX883443.1 | Terrestrial | Arthropoda | Insecta |
| 1732 | Sanxia permutotetra-like virus 1 strain SXSSP2461 RdRp and hypothetical protein genes complete cds | KX883450.1 | Freshwater | Arthropoda | Insecta |
| 1733 | Wuhan house centipede virus 6 strain QTM26514 RNA-dependent RNA polymerase gene complete cds | KX883132.1 | Terrestrial | Arthropoda | Insecta |
| 1734 | Wuhan house centipede virus 6 strain SCM50390 RNA-dependent RNA polymerase gene complete cds | KX883147.1 | Terrestrial | Arthropoda | Insecta |
| 1735 | Wuhan house centipede virus 6 strain WGML146150 RNA-dependent RNA polymerase gene complete cds | KX883198.1 | Terrestrial | Arthropoda | Myriapoda |

| | | | | | |
|---|---|---|---|---|---|
| 1736 | Wuhan house centipede virus 6 strain WHWG55397 RNA-dependent RNA polymerase gene complete cds | KX883241.1 | Terrestrial | Arthropoda | Myriapoda |
| 1737 | Wuhan house centipede virus 6 strain CJLX30267 RNA-dependent RNA polymerase gene complete cds | KX883092.1 | Freshwater | Arthropoda | Crustacea |
| 1738 | Wuhan house centipede virus 6 strain WHYY37237 RNA-dependent RNA polymerase gene complete cds | KX883245.1 | Terrestrial | Arthropoda | Myriapoda |
| 1739 | Wuhan house centipede virus 6 strain arthropodmix44324 RNA-dependent RNA polymerase gene complete cds | KX883011.1 | Terrestrial | Arthropoda | Mix |
| 1740 | Wuhan Mosquito Virus 2 strain QN2-7 RNA-dependent RNA polymerase (L) gene complete cds | KM817698.1 | Terrestrial | Arthropoda | Insecta |
| 1741 | Beihai picorna-like virus 109 strain YYSZX3390 hypothetical protein gene partial cds | KX884392.1 | South China Sea | Arthropoda | Crustacea |
| 1742 | Beihai tombus-like virus 10 strain BHZY59908 hypothetical protein 1 hypothetical protein 2 hypothetical protein 3 | KX883070.1 | South China Sea | Mollusca | Cephalopoda |
| 1743 | Beihai uca arcuata virus 1 strain BFZCX5633 hypothetical protein gene complete cds | KX882942.1 | South China Sea | Arthropoda | Crustacea |
| 1744 | Wuhan insect virus 30 strain WHZM9950 hypothetical protein gene partial cds | KX882986.1 | Terrestrial | Arthropoda | Insecta |
| 1745 | Wuhan insect virus 30 strain WHZM9934 hypothetical protein gene complete cds | KX882985.1 | Terrestrial | Arthropoda | Insecta |
| 1746 | Beihai sesarmid crab virus 3 strain SCJXSX39048 RNA-dependent RNA polymerase gene complete cds | KX884032.1 | South China Sea | Arthropoda | Crustacea |
| 1747 | Wenling crustacean virus 13 strain WLJQ104251 putative glycoprotein and hypothetical protein genes complete cds | KX884454.1 | East China Sea | Arthropoda | Crustacea |
| 1748 | Beihai sobemo-like virus 15 strain BHBJDX18383 hypothetical protein genes complete cds | KX882781.1 | South China Sea | Arthropoda | Crustacea |
| 1749 | Hubei permutotetra-like virus 5 strain WHYY24053 RdRp and hypothetical protein genes complete cds | KX883456.1 | Terrestrial | Arthropoda | Myriapoda |
| 1750 | Hubei permutotetra-like virus 3 strain spider127212 RdRp and hypothetical protein genes complete cds | KX883449.1 | Terrestrial | Arthropoda | Chelicerata |
| 1751 | Wuhan house centipede virus 9 strain WHYY23575 RdRp gene complete cds | KX883454.1 | Terrestrial | Arthropoda | Myriapoda |
| 1752 | Wuhan house centipede virus 9 strain arthropodmix13921 hypothetical protein gene complete cds | KX883436.1 | Terrestrial | Arthropoda | Mix |
| 1753 | Wuhan aphid virus 1 strain WHYC-1 segment 4 complete sequence | KR902724.1 | Terrestrial | Arthropoda | Insecta |
| 1754 | Wuhan aphid virus 1 strain WHYC-1 segment 1 complete sequence | KR902721.1 | Terrestrial | Arthropoda | Insecta |
| 1755 | Shahe bunya-like virus 1 strain SHWCII5304 RNA-dependent RNA polymerase gene complete cds | KX884828.1 | Freshwater | Arthropoda | Mix |
| 1756 | Shahe bunya-like virus 1 strain SHWC0209c12777 RNA-dependent RNA polymerase gene complete cds | KX884821.1 | Freshwater | Arthropoda | Crustacea |
| 1757 | Beihai paphia shell virus 4 strain BWBFG40859 hypothetical protein gene complete cds | KX884532.1 | South China Sea | Mollusca | Bivalvia |
| 1758 | Hubei bunya-like virus 4 strain WGML135976 RNA-dependent RNA polymerase gene complete cds | KX884846.1 | Terrestrial | Arthropoda | Myriapoda |
| 1759 | Whenzhou Shrimp Virus 2 strain BJDX-1 RNA-dependent RNA polymerase (L) gene complete cds | KM817687.1 | East China Sea | Arthropoda | Crustacea |
| 1760 | Shahe isopoda virus 4 strain SHWC0209c12776 putative glycoprotein gene complete cds | KX884817.1 | Freshwater | Arthropoda | Crustacea |
| 1761 | Shahe isopoda virus 4 strain SHWCII5333 putative glycoprotein gene complete cds | KX884827.1 | Freshwater | Arthropoda | Mix |
| 1762 | Qingnian Mosquito Virus strain QN3-7 RNA-dependent RNA polymerase (L) gene complete cds | KM817673.1 | Terrestrial | Arthropoda | Insecta |
| 1763 | Jingmen ascaridia virus 1 strain JMYJCC72107 RNA-dependent RNA polymerase gene complete cds | KX884767.1 | Terrestrial | Nematoda | Chromadorea |
| 1764 | Shahe bunya-like virus 4 strain SHWC0209c12661 RNA-dependent RNA polymerase gene partial cds | KX884825.1 | Freshwater | Arthropoda | Crustacea |
| 1765 | Xinzhou Mosquito Virus strain XC3-5 RNA-dependent RNA polymerase (L) gene complete cds | KM817701.1 | Terrestrial | Arthropoda | Insecta |
| 1766 | Hubei bunya-like virus 9 strain QTM75064 RNA-dependent RNA polymerase gene complete cds | KX884789.1 | Terrestrial | Arthropoda | Insecta |
| 1767 | Hubei bunya-like virus 8 strain SCM48138 RNA-dependent RNA polymerase gene complete cds | KX884802.1 | Terrestrial | Arthropoda | Insecta |
| 1768 | Beihai hermit crab virus 2 strain BHJJX25757 hypothetical protein genes complete cds | KX884743.1 | South China Sea | Arthropoda | Crustacea |
| 1769 | Beihai barnacle virus 6 strain BHTH41702 RNA-dependent RNA polymerase gene complete cds | KX884748.1 | South China Sea | Arthropoda | Crustacea |
| 1770 | Wuchang Cockraoch Virus 3 strain WCZL-1 segment 2 complete sequence | KM817605.1 | Terrestrial | Arthropoda | Insecta |
| 1771 | Wuhan pillworm virus 1 strain WHSFII20185 putative glycoprotein gene complete cds | KX884849.1 | Terrestrial | Arthropoda | Crustacea |
| 1772 | Hubei tombus-like virus 6 strain WGML12775 hypothetical protein 1 hypothetical protein 2 and hypothetical protei | KX883193.1 | Terrestrial | Arthropoda | Myriapoda |

| | | | | | |
|---|---|---|---|---|---|
| 1773 | Asian prunus virus 2 strain WHCCII13300 replicase gene partial cds and triple gene block protein 1 and triple gene | KX883815.1 | Terrestrial | Arthropoda | Insecta |
| 1774 | Changjiang tombus-like virus 17 strain CJLX30686 hypothetical protein 1 hypothetical protein 2 and hypothetical p | KX883097.1 | Freshwater | Arthropoda | Crustacea |
| 1775 | Changjiang tombus-like virus 19 strain CJLX57318 hypothetical protein 1 and hypothetical protein 2 genes complet | KX883082.1 | Freshwater | Arthropoda | Crustacea |
| 1776 | Beihai noda-like virus 1 strain BHZC37212 hypothetical protein 1 and hypothetical protein 2 genes complete cds | KX883062.1 | South China Sea | Mollusca | Bivalvia |
| 1777 | Beihai noda-like virus 21 strain BWBFG39999 hypothetical protein gene complete cds | KX883077.1 | South China Sea | Mollusca | Bivalvia |
| 1778 | Hubei tombus-like virus 38 strain WGML123693 hypothetical protein 1 hypothetical protein 2 and hypothetical pro | KX883192.1 | Terrestrial | Arthropoda | Myriapoda |
| 1779 | Hubei tombus-like virus 38 strain WHWG56034 hypothetical protein 1 hypothetical protein 2 and hypothetical prot | KX883243.1 | Terrestrial | Arthropoda | Myriapoda |
| 1780 | Beihai noda-like virus 7 strain BHWZXX2233 hypothetical protein gene partial cds | KX883043.1 | South China Sea | Arthropoda | Crustacea |
| 1781 | Wenzhou noda-like virus 1 strain WZFSL84567 hypothetical protein 1 and hypothetical protein 2 genes complete c | KX883258.1 | Freshwater | Mollusca | Gastropoda |
| 1782 | Hubei tombus-like virus 43 strain WGML146568 hypothetical protein 1 hypothetical protein 2 and hypothetical pro | KX883200.1 | Terrestrial | Arthropoda | Myriapoda |
| 1783 | Sanxia water strider virus 16 strain SXSSP3988 hypothetical protein gene complete cds | KX883179.1 | Freshwater | Arthropoda | Insecta |
| 1784 | Sanxia water strider virus 16 strain SXXX38069 hypothetical protein gene complete cds | KX883181.1 | Freshwater | Arthropoda | Crustacea |
| 1785 | Hubei partiti-like virus 12 strain QTM77172 RdRp gene complete cds | KX884144.1 | Terrestrial | Arthropoda | Insecta |
| 1786 | Hubei macula-like virus 2 strain QTM27299 hypothetical protein genes complete cds | KX883785.1 | Terrestrial | Arthropoda | Insecta |
| 1787 | Hubei tombus-like virus 26 strain QTM27120 hypothetical protein 1 and hypothetical protein 2 genes complete cds | KX883143.1 | Terrestrial | Arthropoda | Insecta |
| 1788 | Wenzhou tombus-like virus 14 strain WZFSL15005 hypothetical protein 1 hypothetical protein 2 and hypothetical p | KX883275.1 | Freshwater | Mollusca | Gastropoda |
| 1789 | Wenling reo-like virus 1 strain WLJQ76115 hypothetical protein 1 and RdRp genes complete cds | KX884731.1 | East China Sea | Arthropoda | Crustacea |
| 1790 | Hubei unio douglasiae virus 1 strain WHBL72336 RNA-dependent RNA polymerase and B2-like protein genes con | KX883207.1 | Freshwater | Mollusca | Bivalvia |
| 1791 | Muthill virus strain SCM51092 hypothetical protein gene complete cds and hypothetical protein gene partial cds | KX883790.1 | Terrestrial | Arthropoda | Insecta |
| 1792 | Wuhan insect virus 35 strain WHCCII10592 hypothetical protein gene complete cds | KX883234.1 | Terrestrial | Arthropoda | Insecta |
| 1793 | Hubei tombus-like virus 42 strain fly55787 hypothetical protein 1 and hypothetical protein 2 genes complete cds | KX883106.1 | Terrestrial | Arthropoda | Insecta |
| 1794 | Beihai noda-like virus 17 strain BHXG25071 hypothetical protein gene complete cds | KX883058.1 | South China Sea | Arthropoda | Crustacea |
| 1795 | Beihai noda-like virus 17 strain BHHK123677 hypothetical protein gene complete cds | KX883026.1 | South China Sea | Cnidaria | Anthozoa |
| 1796 | Hubei levi-like virus 2 strain WHWN39199 hypothetical protein genes complete cds | KX883588.1 | Freshwater | Mollusca | Gastropoda |
| 1797 | Hubei levi-like virus 8 strain WHWN54380 hypothetical protein genes complete cds | KX883590.1 | Freshwater | Mollusca | Gastropoda |
| 1798 | Hubei levi-like virus 7 strain WHWN54319 hypothetical protein genes complete cds | KX883589.1 | Freshwater | Mollusca | Gastropoda |
| 1799 | Hubei levi-like virus 10 strain WHYY22742 hypothetical protein genes complete cds | KX883593.1 | Terrestrial | Arthropoda | Myriapoda |
| 1800 | Changjiang levi-like virus 2 strain CJLX26414 hypothetical protein genes complete cds | KX883518.1 | Freshwater | Arthropoda | Crustacea |
| 1801 | Beihai levi-like virus 26 strain HOU143495 hypothetical protein genes complete cds | KX883532.1 | South China Sea | Arthropoda | Chelicerata |
| 1802 | Hubei levi-like virus 9 strain WGML141493 hypothetical protein genes complete cds | KX883575.1 | Terrestrial | Arthropoda | Myriapoda |
| 1803 | Beihai levi-like virus 33 strain HOU158604 hypothetical protein genes complete cds | KX883529.1 | South China Sea | Arthropoda | Chelicerata |
| 1804 | Hubei levi-like virus 4 strain WHYY19197 hypothetical protein genes complete cds | KX883595.1 | Terrestrial | Arthropoda | Myriapoda |
| 1805 | Beihai levi-like virus 24 strain BHTSS11091 hypothetical protein genes complete cds | KX883487.1 | South China Sea | Arthropoda | Crustacea |
| 1806 | Hubei levi-like virus 11 strain WHWN53995 hypothetical protein genes complete cds | KX883591.1 | Freshwater | Mollusca | Gastropoda |
| 1807 | Hubei levi-like virus 14 strain SZmix15676 hypothetical protein genes complete cds | KX883571.1 | Freshwater | Annelida | Clitellata |
| 1808 | Hubei levi-like virus 13 strain WHCC54021 hypothetical protein genes complete cds | KX883584.1 | Terrestrial | Arthropoda | Insecta |
| 1809 | Hubei noda-like virus 19 strain WHBL72681 hypothetical protein 1 and hypothetical protein 2 genes complete cds | KX883216.1 | Freshwater | Mollusca | Bivalvia |

| | | | | | |
|---|---|---|---|---|---|
| 1810 | Beihai narna-like virus 21 strain SCJXSX34167 RNA-dependent RNA polymerase gene complete cds | KX883545.1 | South China Sea | Arthropoda | Crustacea |
| 1811 | Bivalvia Picorna-like virus D50 | GWHAOZO01000000 | East China Sea | Mollusca | Bivalvia |
| 1812 | Wenzhou narna-like virus 5 strain WZFSL79329 RNA-dependent RNA polymerase and putative capsid protein gen | KX883610.1 | Freshwater | Mollusca | Gastropoda |
| 1813 | Hubei reo-like virus 14 strain fly110803 RdRp gene complete cds | KX884607.1 | Terrestrial | Arthropoda | Insecta |
| 1814 | Hubei reo-like virus 14 strain spider130579 RdRp gene partial cds | KX884702.1 | Terrestrial | Arthropoda | Chelicerata |
| 1815 | Bivalvia Amarillo-like virus N3 | GWHAPDV01000000 | South China Sea | Mollusca | Bivalvia |
| 1816 | Crustacea Picorna-like virus N30 | GWHAPDQ01000000 | South China Sea | Arthropoda | Crustacea |
| 1817 | Wenzhou levi-like virus 4 strain WZSLuoI85062 hypothetical protein genes complete cds and hypothetical protein g | KX883621.1 | East China Sea | Mollusca | Gastropoda |
| 1818 | Wenzhou levi-like virus 7 strain WZFSL85855 hypothetical protein genes complete cds | KX883616.1 | Freshwater | Mollusca | Gastropoda |
| 1819 | Wenling levi-like virus 4 strain WLJQ103632 hypothetical protein genes complete cds | KX883608.1 | East China Sea | Arthropoda | Crustacea |
| 1820 | Beihai levi-like virus 13 strain BHTSS2294 hypothetical protein genes complete cds | KX883494.1 | South China Sea | Arthropoda | Crustacea |
| 1821 | Bivalvia Bunya-like virus N1 | GWHAPHL01000000 | South China Sea | Mollusca | Bivalvia |
| 1822 | Bivalvia Bunya-like virus H1 | GWHAPGP01000000 | Yellow China Sea | Mollusca | Bivalvia |
| 1823 | Hubei hepe-like virus 1 strain WHCC116187 replicase and hypothetical protein genes complete cds | KX883809.1 | Terrestrial | Arthropoda | Insecta |
| 1824 | Hubei partiti-like virus 47 strain QTM46465 RdRp gene complete cds | KX884137.1 | Terrestrial | Arthropoda | Insecta |
| 1825 | Beihai tombus-like virus 11 strain BHHK128439 hypothetical protein 1 hypothetical protein 2 and hypothetical prot | KX883029.1 | South China Sea | Cnidaria | Anthozoa |
| 1826 | Hubei tombus-like virus 20 strain spider131362 hypothetical protein 1 hypothetical protein 2 hypothetical protein 3 | KX883163.1 | Terrestrial | Arthropoda | Chelicerata |
| 1827 | Beihai levi-like virus 5 strain BHTSS17971 hypothetical protein genes complete cds | KX883491.1 | South China Sea | Arthropoda | Crustacea |
| 1828 | Changjiang narna-like virus 4 strain CJLX30771 RNA-dependent RNA polymerase gene complete cds | KX883524.1 | Freshwater | Arthropoda | Crustacea |
| 1829 | Hubei narna-like virus 12 strain SZmix21586 RNA-dependent RNA polymerase gene complete cds | KX883573.1 | Freshwater | Annelida | Clitellata |
| 1830 | Wenling narna-like virus 6 strain WLJQ100912 RNA-dependent RNA polymerase gene partial cds | KX883609.1 | East China Sea | Arthropoda | Crustacea |
| 1831 | Beihai levi-like virus 30 strain HOU142311 hypothetical protein genes complete cds | KX883530.1 | South China Sea | Arthropoda | Chelicerata |
| 1832 | Wenzhou levi-like virus 5 strain WZSLuoI85245 hypothetical protein genes complete cds and hypothetical protein g | KX883623.1 | East China Sea | Mollusca | Gastropoda |
| 1833 | Beihai levi-like virus 27 strain BHTH15291 hypothetical protein genes complete cds | KX883483.1 | South China Sea | Arthropoda | Crustacea |
| 1834 | Beihai levi-like virus 32 strain BHSC167323 hypothetical protein genes complete cds | KX883479.1 | South China Sea | Annelida | Polychaeta |
| 1835 | Beihai levi-like virus 35 strain BHXun32039 hypothetical protein genes complete cds and hypothetical protein gene | KX883501.1 | South China Sea | Arthropoda | Crustacea |
| 1836 | Hubei virga-like virus 4 strain WHCCII12598 hypothetical protein gene complete cds and hypothetical protein gene | KX883814.1 | Terrestrial | Arthropoda | Insecta |
| 1837 | Hubei levi-like virus 12 strain WHCC110652 hypothetical protein genes complete cds | KX883585.1 | Terrestrial | Arthropoda | Insecta |
| 1838 | Hubei negev-like virus 2 strain CC64601 hypothetical protein gene partial cds | KX883760.1 | Terrestrial | Arthropoda | Insecta |
| 1839 | Beihai levi-like virus 34 strain BHBJDX37870 hypothetical protein genes complete cds | KX883466.1 | South China Sea | Arthropoda | Crustacea |
| 1840 | Beihai levi-like virus 14 strain BHXun31396 hypothetical protein genes complete cds | KX883502.1 | South China Sea | Arthropoda | Crustacea |
| 1841 | Beihai levi-like virus 25 strain BHSC163351 hypothetical protein genes complete cds | KX883480.1 | South China Sea | Annelida | Polychaeta |
| 1842 | Beihai levi-like virus 16 strain BHJP63773 hypothetical protein genes complete cds | KX883476.1 | South China Sea | Echinodermata | Echinozoa |
| 1843 | Beihai levi-like virus 15 strain HOU158632 hypothetical protein genes complete cds | KX883531.1 | South China Sea | Arthropoda | Chelicerata |
| 1844 | Beihai levi-like virus 19 strain BHBJDX18762 hypothetical protein genes complete cds | KX883464.1 | South China Sea | Arthropoda | Crustacea |
| 1845 | Sanxia levi-like virus 1 strain SXXX34761 hypothetical protein genes complete cds | KX883570.1 | Freshwater | Arthropoda | Crustacea |
| 1846 | Wenzhou levi-like virus 1 strain WZFSL78350 hypothetical protein genes complete cds | KX883612.1 | Freshwater | Mollusca | Gastropoda |

| | | | | | |
|---|---|---|---|---|---|
| ***1847*** | Beihai levi-like virus 21 strain BHZY58374 hypothetical protein genes complete cds | KX883506.1 | South China Sea | Mollusca | Cephalopoda |
| ***1848*** | Shahe levi-like virus 3 strain SHWC0209c12775 hypothetical protein genes complete cds | KX883558.1 | Freshwater | Arthropoda | Crustacea |
| ***1849*** | Hubei levi-like virus 3 strain WHYY22609 hypothetical protein genes complete cds and hypothetical protein gene p | KX883594.1 | Terrestrial | Arthropoda | Myriapoda |
| ***1850*** | Hubei narna-like virus 18 strain CC64130 hypothetical protein and RNA-dependent RNA polymerase genes comple | KX883517.1 | Terrestrial | Arthropoda | Insecta |
| ***1851*** | Sanxia narna-like virus 1 strain SXXX35820 RNA-dependent RNA polymerase and hypothetical protein genes com | KX883568.1 | Freshwater | Arthropoda | Crustacea |
| ***1852*** | Changjiang narna-like virus 3 strain SXXX34198 RNA-dependent RNA polymerase and hypothetical protein genes | KX883569.1 | Freshwater | Arthropoda | Crustacea |
| ***1853*** | Changjiang narna-like virus 3 strain CJLX30350 RNA-dependent RNA polymerase and hypothetical protein genes | KX883522.1 | Freshwater | Arthropoda | Crustacea |
| ***1854*** | Wenling levi-like virus 1 strain WLJQ101898 hypothetical protein genes complete cds | KX883606.1 | East China Sea | Arthropoda | Crustacea |
| ***1855*** | Beihai levi-like virus 7 strain BHJP61416 hypothetical protein gene partial cds hypothetical protein gene complete | KX883475.1 | South China Sea | Echinodermata | Echinozoa |
| ***1856*** | Beihai levi-like virus 8 strain BHTSS17667 hypothetical protein genes complete cds and hypothetical protein gene | KX883490.1 | South China Sea | Arthropoda | Crustacea |
| ***1857*** | Hubei levi-like virus 1 strain WHWN53257 hypothetical protein genes complete cds | KX883592.1 | Freshwater | Mollusca | Gastropoda |
| ***1858*** | Beihai levi-like virus 6 strain BHTSS17847 hypothetical protein genes complete cds | KX883496.1 | South China Sea | Arthropoda | Crustacea |
| ***1859*** | Changjiang levi-like virus 3 strain CJLX29558 hypothetical protein genes complete cds | KX883525.1 | Freshwater | Arthropoda | Crustacea |
| ***1860*** | Hubei narna-like virus 8 strain WGML132164 RNA-dependent RNA polymerase gene complete cds and hypothetic | KX883576.1 | Terrestrial | Arthropoda | Myriapoda |
| ***1861*** | Hubei narna-like virus 11 strain WHBL72829 RNA-dependent RNA polymerase and hypothetical protein genes cor | KX883582.1 | Freshwater | Mollusca | Bivalvia |
| ***1862*** | Beihai barnacle virus 10 strain BHTH18714 RNA-dependent RNA polymerase gene complete cds | KX883482.1 | South China Sea | Arthropoda | Crustacea |
| ***1863*** | Beihai levi-like virus 10 strain BHTSS17923 hypothetical protein genes complete cds | KX883492.1 | South China Sea | Arthropoda | Crustacea |
| ***1864*** | Beihai levi-like virus 22 strain BHTH16011 hypothetical protein genes complete cds | KX883481.1 | South China Sea | Arthropoda | Crustacea |
| ***1865*** | Beihai levi-like virus 22 strain BHJJX24892 hypothetical protein genes complete cds and hypothetical protein gene | KX883470.1 | South China Sea | Arthropoda | Crustacea |
| ***1866*** | Soybean yellow common mosaic virus strain spider132820 hypothetical protein 1 hypothetical protein 2 and hypoth | KX882870.1 | Terrestrial | Arthropoda | Chelicerata |
| ***1867*** | Hubei myriapoda virus 9 strain WGML401068 segment 1 hypothetical protein 1 hypothetical protein 2 and RdRp g | KX882898.1 | Terrestrial | Arthropoda | Myriapoda |
| ***1868*** | Hubei myriapoda virus 9 strain arthropodmix12082 segment 1 hypothetical protein genes complete cds | KX882770.1 | Terrestrial | Arthropoda | Mix |
| ***1869*** | Hubei myriapoda virus 9 strain WHYCM992 segment 1 hypothetical protein 1 hypothetical protein 2 and hypotheti | KX882919.1 | Terrestrial | Arthropoda | Insecta |
| ***1870*** | Hubei myriapoda virus 9 strain WHXS4007 segment 1 hypothetical protein 1 hypothetical protein 2 and hypothetica | KX882915.1 | Terrestrial | Arthropoda | Insecta |
| ***1871*** | Hubei myriapoda virus 9 strain ZCM18719 segment 1 hypothetical protein 1 hypothetical protein 2 and hypothetica | KX882935.1 | Terrestrial | Arthropoda | Insecta |
| ***1872*** | Beihai levi-like virus 1 strain BHTSS17154 hypothetical protein genes complete cds | KX883495.1 | South China Sea | Arthropoda | Crustacea |
| ***1873*** | Beihai narna-like virus 22 strain BHBJDX18651 RNA-dependent RNA polymerase gene complete cds | KX883463.1 | South China Sea | Arthropoda | Crustacea |
| ***1874*** | Shuangao sobemo-like virus 1 strain insectZJ66941 hypothetical protein 1 and hypothetical protein 2 genes complet | KX882824.1 | Terrestrial | Arthropoda | Insecta |
| ***1875*** | Hubei toti-like virus 2 strain arthropodmix13450 hypothetical protein 1 and hypothetical protein 2 genes complete c | KX882939.1 | Terrestrial | Arthropoda | Mix |
| ***1876*** | Wuhan insect virus 30 strain WHZM9752 hypothetical protein gene complete cds | KX882984.1 | Terrestrial | Arthropoda | Insecta |
| ***1877*** | Beihai blue swimmer crab virus 3 strain YYSZX17757 hypothetical protein 1 and hypothetical protein 2 genes com | KX882997.1 | South China Sea | Arthropoda | Crustacea |
| ***1878*** | Crustacea Picorna-like virus N29 | GWHAPDH01000000 | South China Sea | Arthropoda | Crustacea |
| ***1879*** | Huangshi Humpbacked Fly Virus strain HSCC01-1 RNA-dependent RNA polymerase (L) gene complete cds | KM817669.1 | Terrestrial | Arthropoda | Insecta |
| ***1880*** | Shahe qinvirus-like virus 1 strain SHWC0209c11359 RNA-dependent RNA polymerase gene complete cds | KX883994.1 | Freshwater | Arthropoda | Crustacea |
| ***1881*** | Beihai narna-like virus 1 strain BWBFG61141 RNA-dependent RNA polymerase gene complete cds | KX883515.1 | South China Sea | Mollusca | Bivalvia |
| ***1882*** | Wenzhou tapeworm virus 1 strain SGJSC14943 RNA-dependent RNA polymerase gene complete cds | KX884436.1 | Terrestrial | Platyhelminthes | Cestoda |
| ***1883*** | Hubei narna-like virus 7 strain WHBL72300 RNA-dependent RNA polymerase gene complete cds | KX883581.1 | Freshwater | Mollusca | Bivalvia |

| | | | | | |
|---|---|---|---|---|---|
| 1884 | Hubei permutotetra-like virus 4 strain QTM26287 RdRp putative capsid protein and hypothetical protein genes com | KX883445.1 | Terrestrial | Arthropoda | Insecta |
| 1885 | Hubei noda-like virus 5 strain spider124210 hypothetical protein gene complete cds | KX883161.1 | Terrestrial | Arthropoda | Chelicerata |
| 1886 | Hubei noda-like virus 5 strain CC74803 hypothetical protein gene complete cds | KX883080.1 | Terrestrial | Arthropoda | Insecta |
| 1887 | Hubei noda-like virus 11 strain SSZZ1919 hypothetical protein 1 and hypothetical protein 2 genes complete cds | KX883170.1 | Freshwater | Arthropoda | Chelicerata |
| 1888 | Hubei noda-like virus 11 strain arthropodmix22482 hypothetical protein 1 and hypothetical protein 2 genes complet | KX883010.1 | Terrestrial | Arthropoda | Mix |
| 1889 | Changjiang tombus-like virus 3 strain CJLX30495 hypothetical protein 1 hypothetical protein 2 hypothetical proteir | KX883095.1 | Freshwater | Arthropoda | Crustacea |
| 1890 | Wenling tombus-like virus 5 strain WLJQ104103 hypothetical protein 1 hypothetical protein 2 and hypothetical pro | KX883257.1 | East China Sea | Arthropoda | Crustacea |
| 1891 | Hubei sobemo-like virus 1 strain SZmix20791 hypothetical protein 1 hypothetical protein 2 and hypothetical proteir | KX882894.1 | Freshwater | Annelida | Clitellata |
| 1892 | Hubei sobemo-like virus 4 strain WHBL62463 hypothetical protein 1 gene partial cds and hypothetical protein 2 ge | KX882905.1 | Freshwater | Mollusca | Bivalvia |
| 1893 | Hubei mosquito virus 4 strain 3mos6213 hypothetical protein 1 hypothetical protein 2 and hypothetical protein 3 ge | KX883008.1 | Terrestrial | Arthropoda | Insecta |
| 1894 | Hubei mosquito virus 4 strain mosHB232766 hypothetical protein 1 hypothetical protein 2 hypothetical protein 3 an | KX883124.1 | Terrestrial | Arthropoda | Insecta |
| 1895 | Wenzhou tombus-like virus 11 strain mosZJ33874 hypothetical protein 1 hypothetical protein 2 and hypothetical pr | KX883126.1 | Terrestrial | Arthropoda | Insecta |
| 1896 | Sanxia tombus-like virus 9 strain SXXX37849 hypothetical protein 1 hypothetical protein 2 and hypothetical protei | KX883182.1 | Freshwater | Arthropoda | Crustacea |
| 1897 | Sanxia tombus-like virus 6 strain SXSSP2574 hypothetical protein 1 and hypothetical protein 2 genes complete cds | KX883178.1 | Freshwater | Arthropoda | Insecta |
| 1898 | Beihai shrimp virus 3 strain BHBJDX18821 putative glycoprotein gene complete cds | KX884738.1 | South China Sea | Arthropoda | Crustacea |
| 1899 | Beihai shrimp virus 3 strain BHWZXX15649 putative glycoprotein gene complete cds | KX884752.1 | South China Sea | Arthropoda | Crustacea |
| 1900 | Beihai bunya-like virus 4 strain BHCL81707 RNA-dependent RNA polymerase gene complete cds | KX884740.1 | South China Sea | Mollusca | Gastropoda |
| 1901 | Tacheng Tick Virus 2 strain TC252 nucleopasid protein (N) gene complete cds | KM817744.1 | Terrestrial | Arthropoda | Chelicerata |
| 1902 | Wenzhou channeled applesnail virus 4 strain WZFSL85898 putative glycoprotein gene complete cds | KX884864.1 | Freshwater | Mollusca | Gastropoda |
| 1903 | Dabieshan Tick Virus strain D3 RNA-dependent RNA polymerase (L) gene complete cds | KM817666.1 | Terrestrial | Arthropoda | Chelicerata |
| 1904 | Bole Tick Virus 1 strain BL075 RNA-dependent RNA polymerase (L) gene complete cds | KM817664.1 | Terrestrial | Arthropoda | Chelicerata |
| 1905 | Changping Tick Virus 1 strain CP1-2 RNA-dependent RNA polymerase (L) gene complete cds | KM817665.1 | Terrestrial | Arthropoda | Chelicerata |
| 1906 | Tacheng Tick Virus 2 strain TC252 RNA-dependent RNA polymerase (L) gene complete cds | KM817684.1 | Terrestrial | Arthropoda | Chelicerata |
| 1907 | Whenzhou Shrimp Virus 1 strain BJDX-5 glycoprotein precursor (G) gene complete cds | KM817719.1 | East China Sea | Arthropoda | Crustacea |
| 1908 | Zhee Mosquito virus strain XC1-8 RNA-dependent RNA polymerase (L) gene complete cds | KM817705.1 | Terrestrial | Arthropoda | Insecta |
| 1909 | Yichang Insect virus strain YCYC01 glycoprotein precursor (G) gene complete cds | KM817730.1 | Terrestrial | Arthropoda | Insecta |
| 1910 | Whenzhou Shrimp Virus 1 strain BJDX-5 RNA-dependent RNA polymerase (L) gene complete cds | KM817686.1 | East China Sea | Arthropoda | Crustacea |
| 1911 | Hubei insect virus 1 strain arthropodmix14045 RNA-dependent RNA polymerase gene complete cds | KX884735.1 | Terrestrial | Arthropoda | Mix |
| 1912 | Yichang Insect virus strain YCYC01 RNA-dependent RNA polymerase (L) gene complete cds | KM817703.1 | Terrestrial | Arthropoda | Insecta |
| 1913 | Wenling picorna-like virus 9 strain WLJQ104209 hypothetical protein gene complete cds | KX884320.1 | East China Sea | Arthropoda | Crustacea |
| 1914 | Beihai shrimp virus 3 strain BHBJDX18821 hypothetical protein genes complete cds | KX884739.1 | South China Sea | Arthropoda | Crustacea |
| 1915 | Beihai shrimp virus 3 strain BHWZXX15649 hypothetical protein genes complete cds | KX884753.1 | South China Sea | Arthropoda | Crustacea |
| 1916 | Beihai sipunculid worm virus 7 strain BHNXC41490 hypothetical protein and glycoprotein genes complete cds | KX884746.1 | South China Sea | Sipuncula | Sipuncula |
| 1917 | Huangpi Tick Virus 1 strain H124-1 glycoprotein precursor (G) gene complete cds | KM817706.1 | Terrestrial | Arthropoda | Chelicerata |
| 1918 | Hubei qinvirus-like virus 2 strain arthropodmix117161 RNA-dependent RNA polymerase gene partial cds | KX883989.1 | Terrestrial | Arthropoda | Mix |
| 1919 | Beihai sobemo-like virus 13 strain YYSZX17641 hypothetical protein 1 and hypothetical protein 2 genes complete | KX882934.1 | South China Sea | Arthropoda | Crustacea |
| 1920 | Beihai sobemo-like virus 17 strain HOU157151 hypothetical protein 1 and hypothetical protein 2 genes complete cc | KX882820.1 | South China Sea | Arthropoda | Chelicerata |

| | | | | | |
|---|---|---|---|---|---|
| 1921 | Beihai sobemo-like virus 24 strain BHNXC41208 hypothetical protein 1 and hypothetical protein 2 genes complete | KX882788.1 | South China Sea | Sipuncula | Sipuncula |
| 1922 | Beihai sobemo-like virus 24 strain BHBJDX18033 hypothetical protein genes complete cds | KX882780.1 | South China Sea | Arthropoda | Crustacea |
| 1923 | Tacheng Tick Virus 1 strain TC253 glycoprotein precursor (G) gene complete cds | KM817717.1 | Terrestrial | Arthropoda | Chelicerata |
| 1924 | Wenzhou Crab Virus 2 strain ZCX13 segment 2 complete sequence | KM817602.1 | East China Sea | Arthropoda | Crustacea |
| 1925 | Wenzhou shrimp virus 10 strain shrimp12509 RNA-dependent RNA polymerase gene complete cds | KX883551.1 | East China Sea | Arthropoda | Crustacea |
| 1926 | Wenzhou shrimp virus 10 strain WZRBX43419 RNA-dependent RNA polymerase gene complete cds | KX883617.1 | East China Sea | Arthropoda | Crustacea |
| 1927 | Hubei sobemo-like virus 25 strain QTM27214 hypothetical protein 1 and hypothetical protein 2 genes complete cds | KX882847.1 | Terrestrial | Arthropoda | Insecta |
| 1928 | Wuhan insect virus 15 strain WHCCII13252 RNA-dependent RNA polymerase gene complete cds | KX883998.1 | Terrestrial | Arthropoda | Insecta |
| 1929 | Jingmen Tick Virus isolate SY84 segment 3 complete sequence | KJ001581.1 | Terrestrial | Arthropoda | Chelicerata |
| 1930 | Hubei qinvirus-like virus 1 strain SCM49583 RNA-dependent RNA polymerase gene complete cds | KX883992.1 | Terrestrial | Arthropoda | Insecta |
| 1931 | Jingmen Tick Virus isolate SY84 segment 4 complete sequence | KJ001582.1 | Terrestrial | Arthropoda | Chelicerata |
| 1932 | Jingmen Tick Virus isolate SY84 segment 2 complete sequence | KJ001580.1 | Terrestrial | Arthropoda | Chelicerata |
| 1933 | Jingmen Tick Virus isolate SY84 segment 1 complete sequence | KJ001579.1 | Terrestrial | Arthropoda | Chelicerata |
| 1934 | Wenling noda-like virus 1 strain WLJQ102234 hypothetical protein gene partial cds | KX883250.1 | East China Sea | Arthropoda | Crustacea |
| 1935 | Beihai toti-like virus 5 strain BHBJDX8833 hypothetical protein 1 gene partial cds and hypothetical protein 2 gene | KX882943.1 | South China Sea | Arthropoda | Crustacea |
| 1936 | Beihai tombus-like virus 8 strain BHZY60516 hypothetical protein 1 and hypothetical protein 2 genes complete cds | KX883073.1 | South China Sea | Mollusca | Cephalopoda |
| 1937 | Wenling tombus-like virus 1 strain WLJQ104192 hypothetical protein 1 and hypothetical protein 2 genes complete | KX883252.1 | East China Sea | Arthropoda | Crustacea |
| 1938 | Changjiang tombus-like virus 9 strain CJLX30737 hypothetical protein 1 and hypothetical protein 2 genes complete | KX883101.1 | Freshwater | Arthropoda | Crustacea |
| 1939 | Hubei macula-like virus 3 strain spider128797 hypothetical protein gene partial cds and hypothetical protein gene c | KX883799.1 | Terrestrial | Arthropoda | Chelicerata |
| 1940 | Hubei macula-like virus 3 strain spider129741 hypothetical protein gene partial cds and hypothetical protein gene c | KX883800.1 | Terrestrial | Arthropoda | Chelicerata |
| 1941 | Beihai hepe-like virus 11 strain BHTH16134 hypothetical protein gene partial cds | KX883748.1 | South China Sea | Arthropoda | Crustacea |
| 1942 | Hubei permutotetra-like virus 7 strain QTM27196 RdRp and hypothetical protein genes complete cds | KX883444.1 | Terrestrial | Arthropoda | Insecta |
| 1943 | Beihai barnacle virus 5 strain BHTH16146 RNA-dependent RNA polymerase gene complete cds | KX884747.1 | South China Sea | Arthropoda | Crustacea |
| 1944 | Hubei birna-like virus strain SZmix18962 RNA-dependent RNA polymerase gene complete cds | KX883001.1 | Freshwater | Annelida | Clitellata |
| 1945 | Wenzhou qinvirus-like virus 1 strain WZSLuoI86141 RNA-dependent RNA polymerase gene complete cds | KX884002.1 | East China Sea | Mollusca | Gastropoda |
| 1946 | Wuhan insect virus 21 strain WHCCII13077 hypothetical protein 1 and hypothetical protein 2 genes complete cds | KX883227.1 | Terrestrial | Arthropoda | Insecta |
| 1947 | Beihai noda-like virus 13 strain HOU158094 hypothetical protein gene complete cds | KX883111.1 | South China Sea | Arthropoda | Chelicerata |
| 1948 | Beihai noda-like virus 22 strain BWBFG40757 hypothetical protein 1 and hypothetical protein 2 genes complete cd | KX883078.1 | South China Sea | Mollusca | Bivalvia |
| 1949 | Hubei tombus-like virus 7 strain WHBL69243 hypothetical protein 1 hypothetical protein 2 and hypothetical protei | KX883211.1 | Freshwater | Mollusca | Bivalvia |
| 1950 | Shuangao insect virus 11 strain insectZJ65889 hypothetical protein gene complete cds | KX883119.1 | Terrestrial | Arthropoda | Insecta |
| 1951 | Shuangao noda-like virus 1 strain insectZJ94204 hypothetical protein gene complete cds | KX883120.1 | Terrestrial | Arthropoda | Insecta |
| 1952 | Beihai weivirus-like virus 8 strain BHZC36478 RNA-dependent RNA polymerase and hypothetical protein genes c | KX884007.1 | South China Sea | Mollusca | Bivalvia |
| 1953 | Beihai weivirus-like virus 11 strain BWBFG40530 RNA-dependent RNA polymerase and hypothetical protein gene | KX884024.1 | South China Sea | Mollusca | Bivalvia |
| 1954 | Beihai weivirus-like virus 10 strain BHZY59101 RNA-dependent RNA polymerase gene complete cds and hypothe | KX884013.1 | South China Sea | Mollusca | Cephalopoda |
| 1955 | Beihai mantis shrimp virus 6 strain BHXG24422 RNA-dependent RNA polymerase hypothetical protein and B2-lik | KX883056.1 | South China Sea | Arthropoda | Crustacea |
| 1956 | Wenzhou noda-like virus 7 strain WZRBX38997 hypothetical protein 1 and hypothetical protein 2 genes complete | KX883277.1 | East China Sea | Arthropoda | Crustacea |
| 1957 | Hubei noda-like virus 10 strain arthropodmix12988 hypothetical protein gene complete cds | KX883009.1 | Terrestrial | Arthropoda | Mix |

| | | | | | |
|---|---|---|---|---|---|
| ***1958*** | Lake Sinai virus strain WHCC111282 hypothetical protein 1 hypothetical protein 2 hypothetical protein 3 and hypo | KX883223.1 | Terrestrial | Arthropoda | Insecta |
| ***1959*** | Sanxia water strider virus 13 strain SXSSP2578 RNA-dependent RNA polymerase gene complete cds | KX883566.1 | Freshwater | Arthropoda | Insecta |
| ***1960*** | Beihai noda-like virus 25 strain BHWZXX26411 hypothetical protein 1 and hypothetical protein 2 genes complete | KX883054.1 | South China Sea | Arthropoda | Crustacea |
| ***1961*** | Hubei noda-like virus 1 strain QTM26695 hypothetical protein gene complete cds | KX883134.1 | Terrestrial | Arthropoda | Insecta |
| ***1962*** | Hubei odonate virus 12 strain QTM3271 hypothetical protein 1 and hypothetical protein 2 genes complete cds | KX883142.1 | Terrestrial | Arthropoda | Insecta |
| ***1963*** | Hubei odonate virus 12 strain WGML403505 hypothetical protein gene partial cds | KX883203.1 | Terrestrial | Arthropoda | Myriapoda |
| ***1964*** | Hubei odonate virus 12 strain QTM25968 hypothetical protein 1 and hypothetical protein 2 genes complete cds | KX883141.1 | Terrestrial | Arthropoda | Insecta |
| ***1965*** | Hubei noda-like virus 14 strain QTM24683 hypothetical protein gene complete cds | KX883129.1 | Terrestrial | Arthropoda | Insecta |
| ***1966*** | Wenzhou tombus-like virus 8 strain beimix73523 hypothetical protein 1 and hypothetical protein 2 genes complete | KX883015.1 | East China Sea | Mollusca | Bivalvia |
| ***1967*** | Beihai narna-like virus 14 strain BHJP52694 RNA-dependent RNA polymerase gene complete cds | KX883473.1 | South China Sea | Echinodermata | Echinozoa |
| ***1968*** | Hubei noda-like virus 7 strain WHCC113193 hypothetical protein 1 gene complete cds | KX883225.1 | Terrestrial | Arthropoda | Insecta |
| ***1969*** | Wuhan insect virus 21 strain WHCCII13077 hypothetical protein and capsid protein genes complete cds | KX883228.1 | Terrestrial | Arthropoda | Insecta |
| ***1970*** | Hubei noda-like virus 12 strain mosZJ34483 hypothetical protein 1 and hypothetical protein 2 genes complete cds | KX883125.1 | Terrestrial | Arthropoda | Insecta |
| ***1971*** | Wenzhou tombus-like virus 18 strain beimix29339 hypothetical protein 1 hypothetical protein 2 and hypothetical pr | KX883013.1 | East China Sea | Mollusca | Bivalvia |
| ***1972*** | Wenzhou crab virus 4 strain WZRBX43276 hypothetical protein 1 hypothetical protein 2 and hypothetical protein 3 | KX883276.1 | East China Sea | Arthropoda | Crustacea |
| ***1973*** | Wenling toti-like virus 3 strain WLJQ62288 hypothetical protein 1 gene partial cds and hypothetical protein 2 gene | KX882993.1 | East China Sea | Arthropoda | Crustacea |
| ***1974*** | Beihai toti-like virus 1 strain BHXun20321 hypothetical protein gene partial cds and hypothetical protein 2 gene co | KX882947.1 | South China Sea | Arthropoda | Crustacea |
| ***1975*** | Beihai narna-like virus 25 strain BHJJX25414 RNA-dependent RNA polymerase gene partial cds | KX883471.1 | South China Sea | Arthropoda | Crustacea |
| ***1976*** | Beihai tombus-like virus 17 strain BHZC33014 hypothetical protein 1 and hypothetical protein 2 genes complete cd | KX883064.1 | South China Sea | Mollusca | Bivalvia |
| ***1977*** | Wenzhou narna-like virus 4 strain WZSLuoI82501 RNA-dependent RNA polymerase and hypothetical protein gene | KX883625.1 | East China Sea | Mollusca | Gastropoda |
| ***1978*** | Wenzhou levi-like virus 2 strain WZFSL82775 hypothetical protein genes complete cds and hypothetical protein ge | KX883611.1 | Freshwater | Mollusca | Gastropoda |
| ***1979*** | Hubei narna-like virus 17 strain mosHB204971 RNA-dependent RNA polymerase gene complete cds | KX883536.1 | Terrestrial | Arthropoda | Insecta |
| ***1980*** | Changjiang tombus-like virus 15 strain CJLX30583 hypothetical protein 1 and hypothetical protein 2 genes complet | KX883096.1 | Freshwater | Arthropoda | Crustacea |
| ***1981*** | Hubei tombus-like virus 3 strain WGML143241 hypothetical protein 1 hypothetical protein 2 and hypothetical prote | KX883196.1 | Terrestrial | Arthropoda | Myriapoda |
| ***1982*** | Xinzhou nematode virus 7 strain XZSJSC65582 hypothetical protein 1 and hypothetical protein 2 genes complete c | KX883280.1 | Terrestrial | Nematoda | Mix |
| ***1983*** | Changjiang tombus-like virus 16 strain CJLX25258 hypothetical protein 1 hypothetical protein 2 and hypothetical p | KX883085.1 | Freshwater | Arthropoda | Crustacea |
| ***1984*** | Beihai barnacle virus 15 strain BHTH16139 hypothetical protein 1 and hypothetical protein 2 genes complete cds | KX882946.1 | South China Sea | Arthropoda | Crustacea |
| ***1985*** | Beihai toti-like virus 2 strain BHCL81631 hypothetical protein 1 gene partial cds and hypothetical protein 2 gene c | KX882944.1 | South China Sea | Mollusca | Gastropoda |
| ***1986*** | Hubei noda-like virus 20 strain SZmix18793 hypothetical protein 1 and hypothetical protein 2 genes complete cds | KX883190.1 | Freshwater | Annelida | Clitellata |
| ***1987*** | Beihai sobemo-like virus 9 strain BHZY60769 hypothetical protein 1 and hypothetical protein 2 genes complete cd | KX882806.1 | South China Sea | Mollusca | Cephalopoda |
| ***1988*** | Beihai weivirus-like virus 17 strain BHZC37426 RNA-dependent RNA polymerase hypothetical protein and putativ | KX884008.1 | South China Sea | Mollusca | Bivalvia |
| ***1989*** | Beihai weivirus-like virus 2 strain BHZC36779 hypothetical protein RNA-dependent RNA polymerase and putative | KX884009.1 | South China Sea | Mollusca | Bivalvia |
| ***1990*** | Beihai weivirus-like virus 1 strain BWBFG40821 RNA-dependent RNA polymerase and hypothetical protein genes | KX884020.1 | South China Sea | Mollusca | Bivalvia |
| ***1991*** | Beihai weivirus-like virus 4 strain BWBFG40762 RNA-dependent RNA polymerase and hypothetical protein genes | KX884022.1 | South China Sea | Mollusca | Bivalvia |
| ***1992*** | Beihai sobemo-like virus 8 strain BWBFG36635 RdRp and putative capsid protein genes complete cds | KX882809.1 | South China Sea | Mollusca | Bivalvia |
| ***1993*** | Beihai weivirus-like virus 7 strain BHZY60887 RNA-dependent RNA polymerase and hypothetical protein genes c | KX884014.1 | South China Sea | Mollusca | Cephalopoda |
| ***1994*** | Beihai weivirus-like virus 3 strain BHZY60776 RNA-dependent RNA polymerase and hypothetical protein genes c | KX884010.1 | South China Sea | Mollusca | Cephalopoda |

| | | | | | |
|---|---|---|---|---|---|
| 1995 | Changjiang tombus-like virus 13 strain CJLX26767 hypothetical protein 1 and hypothetical protein 2 genes complet | KX883087.1 | Freshwater | Arthropoda | Crustacea |
| 1996 | Changjiang crawfish virus 7 strain CJLX29643 hypothetical protein 1 hypothetical protein 2 and hypothetical protei | KX883089.1 | Freshwater | Arthropoda | Crustacea |
| 1997 | Changjiang tombus-like virus 12 strain CJLX27860 hypothetical protein 1 and hypothetical protein 2 genes complet | KX883088.1 | Freshwater | Arthropoda | Crustacea |
| 1998 | Wenzhou tombus-like virus 4 strain shrimp14425 hypothetical protein 1 and hypothetical protein 2 genes complete | KX883152.1 | East China Sea | Arthropoda | Crustacea |
| 1999 | Changjiang tombus-like virus 4 strain CJLX30236 hypothetical protein 1 hypothetical protein 2 and hypothetical pr | KX883091.1 | Freshwater | Arthropoda | Crustacea |
| 2000 | Changjiang tombus-like virus 7 strain CJLX49320 hypothetical protein 2 and hypothetical protein 3 genes complete | KX883103.1 | Freshwater | Arthropoda | Crustacea |
| 2001 | Wuhan horsefly Virus 3 strain horsefly123863 RNA-dependent RNA polymerase gene complete cds | KX883528.1 | Terrestrial | Arthropoda | Insecta |
| 2002 | Beihai weivirus-like virus 13 strain BHZY60103 RNA-dependent RNA polymerase hypothetical protein 1 and hypo | KX884019.1 | South China Sea | Mollusca | Cephalopoda |
| 2003 | Hubei tombus-like virus 34 strain WHBL73047 hypothetical protein 1 hypothetical protein 2 and hypothetical prote | KX883218.1 | Freshwater | Mollusca | Bivalvia |
| 2004 | Beihai noda-like virus 10 strain HWRTX12795 hypothetical protein 1 and hypothetical protein 2 genes complete cd | KX883117.1 | South China Sea | Arthropoda | Crustacea |
| 2005 | Hubei unio douglasiae virus 3 strain WHBL73106 hypothetical protein 1 and hypothetical protein 2 genes complete | KX883220.1 | Freshwater | Mollusca | Bivalvia |
| 2006 | Beihai weivirus-like virus 21 strain BHZY60786 RNA-dependent RNA polymerase and hypothetical protein genes | KX884018.1 | South China Sea | Mollusca | Cephalopoda |
| 2007 | Beihai tombus-like virus 5 strain HOU131178 hypothetical protein 1 and hypothetical protein 2 genes complete cds | KX883116.1 | South China Sea | Arthropoda | Chelicerata |
| 2008 | Beihai weivirus-like virus 18 strain BHZY59250 RNA-dependent RNA polymerase gene complete cds | KX884016.1 | South China Sea | Mollusca | Cephalopoda |
| 2009 | Sanxia tombus-like virus 7 strain SXSSP2567 hypothetical protein 1 hypothetical protein 2 hypothetical protein 3 a | KX883177.1 | Freshwater | Arthropoda | Insecta |
| 2010 | Hubei permutotetra-like virus 10 strain ZCM16232 RdRp and putative capsid protein genes complete cds | KX883458.1 | Terrestrial | Arthropoda | Insecta |
| 2011 | Beihai permutotetra-like virus 2 strain HOU234589 RdRp and putative capsid protein genes complete cds | KX883438.1 | South China Sea | Arthropoda | Chelicerata |
| 2012 | Hubei tombus-like virus 13 strain WHYY18977 hypothetical protein 1 and hypothetical protein 2 genes complete c | KX883248.1 | Terrestrial | Arthropoda | Myriapoda |
| 2013 | Beihai sobemo-like virus 22 strain BHWZXX14614 hypothetical protein 1 and hypothetical protein 2 genes comple | KX882790.1 | South China Sea | Arthropoda | Crustacea |
| 2014 | Gastropoda Picorna-like virus D15 | GWHAPBD01000000 | East China Sea | Mollusca | Gastropoda |
| 2015 | Crustacea Picorna-like virus N22 | GWHAPDI01000000 | South China Sea | Arthropoda | Crustacea |
| 2016 | Wenling toti-like virus 1 strain WLJQ102546 hypothetical protein 1 and hypothetical protein 2 genes partial cds | KX882991.1 | East China Sea | Arthropoda | Crustacea |
| 2017 | Beihai sobemo-like virus 19 strain BHBJDX18617 hypothetical protein genes complete cds | KX882783.1 | South China Sea | Arthropoda | Crustacea |
| 2018 | Beihai zhaovirus-like virus 3 strain BHZY60564 polyprotein gene complete cds | KX884039.1 | South China Sea | Mollusca | Cephalopoda |
| 2019 | Hubei permutotetra-like virus 8 strain WHSWHC54546 RdRp putative capsid protein and hypothetical protein gene | KX883453.1 | Terrestrial | Nematoda | Chromadorea |
| 2020 | Hubei permutotetra-like virus 6 strain QCM123521 RdRp and putative capsid protein genes complete cds | KX883442.1 | Terrestrial | Arthropoda | Insecta |
| 2021 | Bivalvia Picorna-like virus D25 | GWHAOZC01000000 | East China Sea | Mollusca | Bivalvia |
| 2022 | Wenzhou tapeworm virus 1 strain SGJSC14943 hypothetical protein 1 hypothetical protein 2 hypothetical protein 3 | KX884437.1 | Terrestrial | Platyhelminthes | Cestoda |
| 2023 | Wenling crustacean virus 14 strain WLJQ104130 putative glycoprotein hypothetical protein 1 and hypothetical prot | KX884452.1 | East China Sea | Arthropoda | Crustacea |
| 2024 | Beihai levi-like virus 3 strain BHTSS17989 hypothetical protein genes complete cds | KX883488.1 | South China Sea | Arthropoda | Crustacea |
| 2025 | Beihai levi-like virus 4 strain BHTSS17991 hypothetical protein genes complete cds | KX883489.1 | South China Sea | Arthropoda | Crustacea |
| 2026 | Beihai levi-like virus 11 strain BHTH16058 hypothetical protein genes complete cds and hypothetical protein gene | KX883484.1 | South China Sea | Arthropoda | Crustacea |
| 2027 | Bivalvia Picorna-like virus D22 | GWHAOYN0100000C | East China Sea | Mollusca | Bivalvia |
| 2028 | Wenzhou narna-like virus 6 strain WZFSL27295 RNA-dependent RNA polymerase gene partial cds | KX883613.1 | Freshwater | Mollusca | Gastropoda |
| 2029 | Beihai noda-like virus 2 strain BHZY60456 hypothetical protein 1 gene complete cds and hypothetical protein 2 ge | KX883072.1 | South China Sea | Mollusca | Cephalopoda |
| 2030 | Wenzhou picorna-like virus 27 strain shrimp14502 hypothetical protein gene complete cds | KX883981.1 | East China Sea | Arthropoda | Crustacea |
| 2031 | Beihai noda-like virus 11 strain BHJJX49550 RNA-dependent RNA polymerase gene complete cds | KX883032.1 | South China Sea | Arthropoda | Crustacea |

| | | | | | |
|---|---|---|---|---|---|
| 2032 | Beihai noda-like virus 29 strain BHBJDX18138 hypothetical protein gene complete cds | KX883018.1 | South China Sea | Arthropoda | Crustacea |
| 2033 | Beihai noda-like virus 29 strain BHWZXX15622 hypothetical protein gene complete cds | KX883049.1 | South China Sea | Arthropoda | Crustacea |
| 2034 | Beihai sobemo-like virus 20 strain BHNXC40978 hypothetical protein 1 and hypothetical protein 2 genes complete | KX882787.1 | South China Sea | Sipuncula | Sipuncula |
| 2035 | Bivalvia Picorna-like virus D41 | GWHAOZK01000000 | East China Sea | Mollusca | Bivalvia |
| 2036 | Beihai noda-like virus 19 strain BHZY60825 hypothetical protein 1 and hypothetical protein 2 genes complete cds | KX883076.1 | South China Sea | Mollusca | Cephalopoda |
| 2037 | Changjiang narna-like virus 2 strain CJLX29055 RNA-dependent RNA polymerase and putative capsid protein gen | KX883520.1 | Freshwater | Arthropoda | Crustacea |
| 2038 | Hubei noda-like virus 26 strain GCM10516 hypothetical protein gene complete cds | KX883107.1 | Terrestrial | Arthropoda | Insecta |
| 2039 | Shahe tombus-like virus 1 strain SHWC0209c12685 hypothetical protein 1 and hypothetical protein 2 genes comple | KX883155.1 | Freshwater | Arthropoda | Crustacea |
| 2040 | Shahe picorna-like virus 4 strain SHWC13603 hypothetical protein gene complete cds | KX883659.1 | Freshwater | Arthropoda | Mix |
| 2041 | Shahe picorna-like virus 4 strain SHWC0209c12741 hypothetical protein gene complete cds | KX883645.1 | Freshwater | Arthropoda | Crustacea |
| 2042 | Wenzhou tombus-like virus 1 strain WZFSL78713 hypothetical protein 1 and hypothetical protein 2 genes complete | KX883268.1 | Freshwater | Mollusca | Gastropoda |
| 2043 | Wuhan Louse Fly Virus 7 strain BFJSC-3 segment 2 complete sequence | KM817608.1 | Terrestrial | Arthropoda | Insecta |
| 2044 | Hubei noda-like virus 2 strain WHBL70552 hypothetical protein gene complete cds | KX883205.1 | Freshwater | Mollusca | Bivalvia |
| 2045 | Wuhan Louse Fly Virus 6 strain BFJSC-2 segment 2 complete sequence | KM817607.1 | Terrestrial | Arthropoda | Insecta |
| 2046 | Gastropoda Picorna-like virus N5 | GWHAPFL01000000 | South China Sea | Mollusca | Gastropoda |
| 2047 | Hubei sobemo-like virus 3 strain WHBL72900 hypothetical protein 1 hypothetical protein 2 and hypothetical protei | KX882904.1 | Freshwater | Mollusca | Bivalvia |
| 2048 | Beihai sphaeromadae virus 3 strain BHHK126836 RNA-dependent RNA polymerase gene complete cds | KX883027.1 | South China Sea | Cnidaria | Anthozoa |
| 2049 | Beihai sphaeromadae virus 3 strain BHTSS17005 RNA-dependent RNA polymerase gene complete cds | KX883039.1 | South China Sea | Arthropoda | Crustacea |
| 2050 | Wenzhou noda-like virus 2 strain WZSLuoI54252 hypothetical protein gene complete cds | KX883278.1 | East China Sea | Mollusca | Gastropoda |
| 2051 | Beihai barnacle virus 11 strain BHTH10280 RNA-dependent RNA polymerase gene complete cds | KX883036.1 | South China Sea | Arthropoda | Crustacea |
| 2052 | Crustacea Toli-like virus N1 | GWHAPKY01000000 | South China Sea | Arthropoda | Crustacea |
| 2053 | Hubei tombus-like virus 12 strain WHBL71803 hypothetical protein 1 and hypothetical protein 2 genes complete cc | KX883221.1 | Freshwater | Mollusca | Bivalvia |
| 2054 | Hubei narna-like virus 3 strain spider124444 RNA-dependent RNA polymerase gene complete cds | KX883563.1 | Terrestrial | Arthropoda | Chelicerata |
| 2055 | Hubei narna-like virus 3 strain QTM25395 RNA-dependent RNA polymerase gene complete cds | KX883541.1 | Terrestrial | Arthropoda | Insecta |
| 2056 | Hubei tombus-like virus 5 strain WGML136364 hypothetical protein 1 and hypothetical protein 2 genes complete c | KX883194.1 | Terrestrial | Arthropoda | Myriapoda |
| 2057 | Wenling narna-like virus 2 strain WLJQ103952 RNA-dependent RNA polymerase gene complete cds | KX883603.1 | East China Sea | Arthropoda | Crustacea |
| 2058 | Beihai narna-like virus 20 strain BWBFG40664 RNA-dependent RNA polymerase gene complete cds | KX883514.1 | South China Sea | Mollusca | Bivalvia |
| 2059 | Hubei sobemo-like virus 35 strain QTM24286 hypothetical protein 1 and hypothetical protein 2 genes complete cds | KX882840.1 | Terrestrial | Arthropoda | Insecta |
| 2060 | Hubei sobemo-like virus 21 strain CC64469 segment 1 hypothetical protein 1 gene complete cds and hypothetical p | KX882812.1 | Terrestrial | Arthropoda | Insecta |
| 2061 | Hubei sobemo-like virus 40 strain QTM104806 hypothetical protein 1 and hypothetical protein 2 genes complete cc | KX882849.1 | Terrestrial | Arthropoda | Insecta |
| 2062 | Shuangao insect virus 9 strain insectZJ96499 hypothetical protein 1 and hypothetical protein 2 genes complete cds | KX882826.1 | Terrestrial | Arthropoda | Insecta |
| 2063 | Hubei sobemo-like virus 41 strain spider133517 hypothetical protein 1 and hypothetical protein 2 genes complete c | KX882872.1 | Terrestrial | Arthropoda | Chelicerata |
| 2064 | Hubei sobemo-like virus 41 strain 3mos6151 hypothetical protein genes complete cds | KX882766.1 | Terrestrial | Arthropoda | Insecta |
| 2065 | Hubei toti-like virus 22 strain CC64623 hypothetical protein gene partial cds | KX882954.1 | Terrestrial | Arthropoda | Insecta |
| 2066 | Xingshan nematode virus 5 strain XSNXC27943 segment 2 putative capsid protein gene complete cds | KX882933.1 | Terrestrial | Nematoda | Chromadorea |
| 2067 | Hubei toti-like virus 10 strain mosHB235560 hypothetical protein 1 and hypothetical protein 2 genes complete cds | KX882963.1 | Terrestrial | Arthropoda | Insecta |
| 2068 | Hubei toti-like virus 10 strain 3mos6210 hypothetical protein 1 and hypothetical protein 2 genes complete cds | KX882938.1 | Terrestrial | Arthropoda | Insecta |

| | | | | | |
|---|---|---|---|---|---|
| 2069 | Wuhan spider virus 8 strain spider134060 hypothetical protein 1 hypothetical protein 2 hypothetical protein 3 and h | KX883164.1 | Terrestrial | Arthropoda | Chelicerata |
| 2070 | Bole Tick Virus 1 strain BL075 nucleopasid protein (N) gene complete cds | KM817731.1 | Terrestrial | Arthropoda | Chelicerata |
| 2071 | Hubei toti-like virus 24 strain tick106628 hypothetical protein and hypothetical protein 2 genes complete cds | KX882977.1 | Terrestrial | Arthropoda | Chelicerata |
| 2072 | Yongjia Tick Virus 1 strain YJ1-1 nucleopasid protein (N) and nonstructural protein (NSs) genes complete cds | KM817764.1 | Terrestrial | Arthropoda | Chelicerata |
| 2073 | Wenzhou bivalvia virus 2 strain beimix75763 RNA-dependent RNA polymerase and hypothetical protein genes con | KX884028.1 | East China Sea | Mollusca | Bivalvia |
| 2074 | Hubei polero-like virus 2 strain QTM26674 hypothetical protein 1 hypothetical protein 2 hypothetical protein 3 hyp | KX882844.1 | Terrestrial | Arthropoda | Insecta |
| 2075 | Hubei partiti-like virus 54 strain SZmix20834 RdRp gene complete cds | KX884183.1 | Freshwater | Annelida | Clitellata |
| 2076 | Wuhan Millipede virus 4 strain WHWG256022 segment Seg 3 putative capsid protein gene complete cds | KX884204.1 | Terrestrial | Arthropoda | Myriapoda |
| 2077 | Wuhan Millipede virus 4 strain GCM8225 segment Seg 4 putative capsid protein gene complete cds | KX884095.1 | Terrestrial | Arthropoda | Insecta |
| 2078 | Wuhan cricket virus 2 strain WHXS3745 segment Seg 1 RdRp gene complete cds | KX884208.1 | Terrestrial | Arthropoda | Insecta |
| 2079 | Wuhan cricket virus 2 strain ZCM19297 segment Seg 1 RdRp gene complete cds | KX884243.1 | Terrestrial | Arthropoda | Insecta |
| 2080 | Hubei toti-like virus 20 strain CC62244 hypothetical protein 1 and hypothetical protein 2 genes complete cds | KX882952.1 | Terrestrial | Arthropoda | Insecta |
| 2081 | Hubei diptera virus 13 strain SCM94998 segment 2 putative capsid protein gene complete cds | KX882859.1 | Terrestrial | Arthropoda | Insecta |
| 2082 | Hubei diptera virus 13 strain arthropodmix8005 segment 2 hypothetical protein gene complete cds | KX882777.1 | Terrestrial | Arthropoda | Mix |
| 2083 | Hubei toti-like virus 19 strain horsefly123848 hypothetical protein 1 and hypothetical protein 2 genes complete cds | KX882957.1 | Terrestrial | Arthropoda | Insecta |
| 2084 | Hubei tetragnatha maxillosa virus 6 strain arthropodmix13964 segment 2 hypothetical protein gene complete cds | KX882774.1 | Terrestrial | Arthropoda | Mix |
| 2085 | Hubei tetragnatha maxillosa virus 6 strain SSZZ3520 segment 2 hypothetical protein 1 gene complete cds | KX882883.1 | Freshwater | Arthropoda | Chelicerata |
| 2086 | Hubei tetragnatha maxillosa virus 6 strain WHYY23923 segment 2 hypothetical protein 1 gene complete cds | KX882927.1 | Terrestrial | Arthropoda | Myriapoda |
| 2087 | Beihai hepe-like virus 1 strain BHZY58590 hypothetical protein gene partial cds | KX883755.1 | South China Sea | Mollusca | Cephalopoda |
| 2088 | Hubei toti-like virus 18 strain WHCC116098 hypothetical protein 1 and hypothetical protein 2 genes complete cds | KX882979.1 | Terrestrial | Arthropoda | Insecta |
| 2089 | Wenzhou qinvirus-like virus 2 strain WZRBX42684 hypothetical protein gene complete cds | KX884001.1 | East China Sea | Arthropoda | Crustacea |
| 2090 | Wenzhou toti-like virus 1 strain WZRBX43319 hypothetical protein 1 and hypothetical protein 2 genes complete cd | KX882995.1 | East China Sea | Arthropoda | Crustacea |
| 2091 | Dabieshan Tick Virus strain D3 nucleopasid protein (N) and nonstructural protein (NSs) genes complete cds | KM817733.1 | Terrestrial | Arthropoda | Chelicerata |
| 2092 | Beihai tombus-like virus 6 strain BHWZXX15351 hypothetical protein gene partial cds | KX883044.1 | South China Sea | Arthropoda | Crustacea |
| 2093 | Wenzhou qinvirus-like virus 1 strain WZSLuoI86141 hypothetical protein gene complete cds | KX884003.1 | East China Sea | Mollusca | Gastropoda |
| 2094 | Wuhan insect virus 15 strain WHCCII13252 hypothetical protein gene complete cds | KX883999.1 | Terrestrial | Arthropoda | Insecta |
| 2095 | Hubei partiti-like virus 5 strain QTM47485 RdRp gene complete cds | KX884139.1 | Terrestrial | Arthropoda | Insecta |
| 2096 | Beihai sobemo-like virus 26 strain BHTH8711 hypothetical protein 1 and hypothetical protein 2 genes complete cds | KX882789.1 | South China Sea | Arthropoda | Crustacea |
| 2097 | Hubei yanvirus-like virus 1 strain WHCC105213 hypothetical protein 1 RNA-dependent RNA polymerase and hypo | KX884030.1 | Terrestrial | Arthropoda | Insecta |
| 2098 | Xingshan nematode virus 6 strain XSNXC32793 hypothetical protein 1 hypothetical protein 2 and hypothetical prot | KX882996.1 | Terrestrial | Nematoda | Chromadorea |
| 2099 | Changjiang tombus-like virus 5 strain CJLX42586 hypothetical protein 1 hypothetical protein 2 and hypothetical pro | KX883102.1 | Freshwater | Arthropoda | Crustacea |
| 2100 | Hubei toti-like virus 5 strain WHSWHC37547 hypothetical protein gene complete cds | KX882980.1 | Terrestrial | Nematoda | Chromadorea |
| 2101 | Xingshan nematode virus 5 strain XSNXC27943 segment 1 hypothetical protein 1 and RdRp genes complete cds | KX882932.1 | Terrestrial | Nematoda | Chromadorea |
| 2102 | Beihai picorna-like virus 84 strain BHTH16053 hypothetical protein 1 and hypothetical protein 2 genes complete cc | KX883413.1 | South China Sea | Arthropoda | Crustacea |
| 2103 | Beihai narna-like virus 6 strain BHZC37402 RNA-dependent RNA polymerase gene complete cds | KX883503.1 | South China Sea | Mollusca | Bivalvia |
| 2104 | Beihai noda-like virus 20 strain BHZY60039 hypothetical protein 1 and hypothetical protein 2 genes complete cds | KX883071.1 | South China Sea | Mollusca | Cephalopoda |
| 2105 | Beihai narna-like virus 9 strain HOU146315 RNA-dependent RNA polymerase and hypothetical protein genes comp | KX883534.1 | South China Sea | Arthropoda | Chelicerata |

| | | | | | |
|---|---|---|---|---|---|
| 2106 | Hubei tombus-like virus 1 strain SCM51273 hypothetical protein 1 hypothetical protein 2 and hypothetical protein 3 | KX883149.1 | Terrestrial | Arthropoda | Insecta |
| 2107 | Hubei tombus-like virus 1 strain WHWN54407 hypothetical protein 1 hypothetical protein 2 hypothetical protein 3 | KX883244.1 | Freshwater | Mollusca | Gastropoda |
| 2108 | Hubei sobemo-like virus 5 strain spider59976 hypothetical protein 1 and hypothetical protein 2 genes complete cds | KX882877.1 | Terrestrial | Arthropoda | Chelicerata |
| 2109 | Wenling tombus-like virus 4 strain WLJQ100551 hypothetical protein 1 hypothetical protein 2 and hypothetical pro | KX883249.1 | East China Sea | Arthropoda | Crustacea |
| 2110 | Hubei narna-like virus 19 strain QTM26970 RNA-dependent RNA polymerase and hypothetical protein genes comp | KX883542.1 | Terrestrial | Arthropoda | Insecta |
| 2111 | Hubei mosquito virus 3 strain 3mos6141 RNA-dependent RNA polymerase and hypothetical protein genes complete | KX883460.1 | Terrestrial | Arthropoda | Insecta |
| 2112 | Wenzhou yanvirus-like virus 2 strain beimix74779 RNA-dependent RNA polymerase and hypothetical protein gene | KX884027.1 | East China Sea | Mollusca | Bivalvia |
| 2113 | Hubei narna-like virus 14 strain WHCCII12614 RNA-dependent RNA polymerase and hypothetical protein genes p | KX883586.1 | Terrestrial | Arthropoda | Insecta |
| 2114 | Beihai narna-like virus 19 strain BHNXC41285 RNA-dependent RNA polymerase gene complete cds | KX883477.1 | South China Sea | Sipuncula | Sipuncula |
| 2115 | Wenzhou narna-like virus 11 strain beimix75822 RNA-dependent RNA polymerase gene partial cds | KX883462.1 | East China Sea | Mollusca | Bivalvia |
| 2116 | Wenzhou narna-like virus 3 strain shrimp14503 RNA-dependent RNA polymerase gene complete cds | KX883550.1 | East China Sea | Arthropoda | Crustacea |
| 2117 | Wenzhou narna-like virus 2 strain shrimp13495 RNA-dependent RNA polymerase gene complete cds | KX883549.1 | East China Sea | Arthropoda | Crustacea |
| 2118 | Beihai narna-like virus 4 strain BHZY53599 RNA-dependent RNA polymerase gene complete cds | KX883508.1 | South China Sea | Mollusca | Cephalopoda |
| 2119 | Hubei toti-like virus 17 strain tick107859 hypothetical protein 1 hypothetical protein 2 and hypothetical protein 3 ge | KX882978.1 | Terrestrial | Arthropoda | Chelicerata |
| 2120 | Beihai tombus-like virus 19 strain BHTH16144 hypothetical protein 1 hypothetical protein 2 and hypothetical prote | KX883038.1 | South China Sea | Arthropoda | Crustacea |
| 2121 | Hubei narna-like virus 21 strain QCM13322 hypothetical protein and RNA-dependent RNA polymerase genes com | KX883539.1 | Terrestrial | Arthropoda | Insecta |
| 2122 | Hubei tombus-like virus 19 strain WHCCII13323 hypothetical protein 1 hypothetical protein 2 hypothetical protein | KX883232.1 | Terrestrial | Arthropoda | Insecta |
| 2123 | Wuhan Mosquito Virus 6 strain QN3-6 PB1 (PB1) gene complete cds | KM817625.1 | Terrestrial | Arthropoda | Insecta |
| 2124 | Hubei mosquito virus 2 strain mosZJ35453 segment 2 hypothetical protein 1 gene complete cds | KX882833.1 | Terrestrial | Arthropoda | Insecta |
| 2125 | Hubei mosquito virus 2 strain spider133708 segment 2 hypothetical protein 1 gene complete cds | KX882874.1 | Terrestrial | Arthropoda | Chelicerata |
| 2126 | Hubei mosquito virus 2 strain 3mos6212 segment 2 hypothetical protein gene complete cds | KX882765.1 | Terrestrial | Arthropoda | Insecta |
| 2127 | Beihai picorna-like virus 45 strain BHZC37433 hypothetical protein gene complete cds | KX884496.1 | South China Sea | Mollusca | Bivalvia |
| 2128 | Wuchan romanomermis nematode virus 3 strain WCLSXC83893 segment 1 hypothetical protein 1 and hypothetical | KX882896.1 | Terrestrial | Nematoda | Enoplea |
| 2129 | Sanxia picorna-like virus 12 strain SXXX24153 hypothetical protein 1 and hypothetical protein 2 genes complete c | KX883723.1 | Freshwater | Arthropoda | Crustacea |
| 2130 | Wuchan romanomermis nematode virus 3 strain WCLSXC83893 segment 2 hypothetical protein genes complete cd | KX882897.1 | Terrestrial | Nematoda | Enoplea |
| 2131 | Changjiang picorna-like virus 10 strain CJLX30646 hypothetical protein 1 and hypothetical protein 2 genes comple | KX884553.1 | Freshwater | Arthropoda | Crustacea |
| 2132 | Jingmen toti-like virus 1 strain JMYJCC57201 hypothetical protein gene partial cds | KX882961.1 | Terrestrial | Nematoda | Chromadorea |
| 2133 | Hubei diptera virus 12 strain SCM245722 segment 2 hypothetical protein gene complete cds | KX882854.1 | Terrestrial | Arthropoda | Insecta |
| 2134 | Beihai horseshoe crab virus 1 strain HOU157708 capsid protein gene complete cds | KX883115.1 | South China Sea | Arthropoda | Chelicerata |
| 2135 | Beihai picorna-like virus 66 strain BHHQ76843 hypothetical protein gene complete cds | KX883357.1 | South China Sea | Tunicata | Ascidiacea |
| 2136 | Wenzhou narna-like virus 8 strain WZRBX43201 RNA-dependent RNA polymerase gene complete cds | KX883618.1 | East China Sea | Arthropoda | Crustacea |
| 2137 | Beihai sobemo-like virus 14 strain HOU157842 hypothetical protein 1 and hypothetical protein 2 genes complete c | KX882821.1 | South China Sea | Arthropoda | Chelicerata |
| 2138 | Hubei sobemo-like virus 17 strain WGML125344 hypothetical protein 1 gene complete cds and hypothetical protei | KX882900.1 | Terrestrial | Arthropoda | Myriapoda |
| 2139 | Beihai razor shell virus 3 strain BHZC45609 hypothetical protein 1 and hypothetical protein 2 genes complete cds | KX882798.1 | South China Sea | Mollusca | Bivalvia |
| 2140 | Sanxia sobemo-like virus 4 strain SXSSP2021 hypothetical protein 1 and hypothetical protein 2 genes complete cds | KX882884.1 | Freshwater | Arthropoda | Insecta |
| 2141 | Beihai sobemo-like virus 6 strain BWBFG40829 hypothetical protein 1 and hypothetical protein 2 genes complete c | KX882811.1 | South China Sea | Mollusca | Bivalvia |
| 2142 | Hubei sobemo-like virus 7 strain spider134446 hypothetical protein 1 and hypothetical protein 2 genes complete cds | KX882876.1 | Terrestrial | Arthropoda | Chelicerata |

| | | | | | |
|---|---|---|---|---|---|
| 2143 | Hubei sobemo-like virus 6 strain spider128669 hypothetical protein 1 and hypothetical protein 2 genes complete cds | KX882869.1 | Terrestrial | Arthropoda | Chelicerata |
| 2144 | Beihai sobemo-like virus 10 strain BHZY60634 hypothetical protein 1 and hypothetical protein 2 genes complete co | KX882803.1 | South China Sea | Mollusca | Cephalopoda |
| 2145 | Beihai sobemo-like virus 11 strain BHZY59277 hypothetical protein 1 and hypothetical protein 2 genes complete co | KX882800.1 | South China Sea | Mollusca | Cephalopoda |
| 2146 | Shuangao sobemo-like virus 2 strain insectZJ89114 hypothetical protein 1 and hypothetical protein 2 genes complet | KX882825.1 | Terrestrial | Arthropoda | Insecta |
| 2147 | Hubei sobemo-like virus 15 strain tick96090 hypothetical protein 1 and hypothetical protein 2 genes complete cds | KX882895.1 | Terrestrial | Arthropoda | Chelicerata |
| 2148 | Shahe isopoda virus 5 strain SHWC0209c16708 hypothetical protein 1 hypothetical protein 2 hypothetical protein 3 | KX883154.1 | Freshwater | Arthropoda | Crustacea |
| 2149 | Shahe isopoda virus 5 strain SHWCII46393 hypothetical protein 1 hypothetical protein 2 hypothetical protein 3 and | KX883156.1 | Freshwater | Arthropoda | Mix |
| 2150 | Shahe isopoda virus 5 strain WZFSL84267 hypothetical protein 1 hypothetical protein 2 hypothetical protein 3 and | KX883263.1 | Freshwater | Mollusca | Gastropoda |
| 2151 | Beihai sobemo-like virus 1 strain HWRTX12503 hypothetical protein 1 and hypothetical protein 2 genes complete c | KX882822.1 | South China Sea | Arthropoda | Crustacea |
| 2152 | Beihai sobemo-like virus 1 strain BHZY60709 hypothetical protein 1 RdRp and putative capsid protein genes comp | KX882805.1 | South China Sea | Mollusca | Cephalopoda |
| 2153 | Wenling sobemo-like virus 2 strain WLJQ95081 hypothetical protein 1 and hypothetical protein 2 genes complete c | KX882929.1 | East China Sea | Arthropoda | Crustacea |
| 2154 | Narna-like virus 6 strain WGML136220 RNA-dependent RNA polymerase gene complete cds | KX883577.1 | Terrestrial | Arthropoda | Myriapoda |
| 2155 | Beihai permutotetra-like virus 1 strain SCJXSX31686 RdRp gene partial cds | KX883447.1 | South China Sea | Arthropoda | Crustacea |
| 2156 | Sanxia water strider virus 19 strain SXSSP2568 RdRp and hypothetical protein genes complete cds | KX883451.1 | Freshwater | Arthropoda | Insecta |
| 2157 | Sanxia water strider virus 19 strain SXXX37734 RdRp and hypothetical protein genes complete cds | KX883452.1 | Freshwater | Arthropoda | Crustacea |
| 2158 | Hubei permutotetra-like virus 9 strain ZCM21318 RdRp and hypothetical protein genes complete cds | KX883459.1 | Terrestrial | Arthropoda | Insecta |
| 2159 | Beihai narna-like virus 3 strain BHBJDX18174 RNA-dependent RNA polymerase gene complete cds | KX883465.1 | South China Sea | Arthropoda | Crustacea |
| 2160 | Changjiang polero-like virus 1 strain CJLX30310 hypothetical protein 1 hypothetical protein 2 and hypothetical pro | KX882817.1 | Freshwater | Arthropoda | Crustacea |
| 2161 | Hubei sobemo-like virus 40 strain spider66422 hypothetical protein 1 and hypothetical protein 2 genes complete cds | KX882864.1 | Terrestrial | Arthropoda | Chelicerata |
| 2162 | Wuchang cockroach Virus 4 strain ZL17511 RNA-dependent RNA polymerase gene complete cds | KX883626.1 | Terrestrial | Arthropoda | Insecta |
| 2163 | Hubei narna-like virus 15 strain QCM133922 RNA-dependent RNA polymerase gene complete cds | KX883540.1 | Terrestrial | Arthropoda | Insecta |
| 2164 | Beihai narna-like virus 8 strain BHZY59840 RNA-dependent RNA polymerase and hypothetical protein genes com | KX883509.1 | South China Sea | Mollusca | Cephalopoda |
| 2165 | Beihai sobemo-like virus 27 strain BHWZXX15541 hypothetical protein 1 and hypothetical protein 2 genes comple | KX882791.1 | South China Sea | Arthropoda | Crustacea |
| 2166 | Hubei levi-like virus 5 strain WHWN53876 hypothetical protein genes complete cds | KX883587.1 | Freshwater | Mollusca | Gastropoda |
| 2167 | Wenzhou narna-like virus 1 strain WZFSL81657 RNA-dependent RNA polymerase gene partial cds | KX883615.1 | Freshwater | Mollusca | Gastropoda |
| 2168 | Changjiang narna-like virus 1 strain CJLX30050 RNA-dependent RNA polymerase gene complete cds | KX883521.1 | Freshwater | Arthropoda | Crustacea |
| 2169 | Hubei narna-like virus 1 strain WHCC258047 RNA-dependent RNA polymerase gene partial cds | KX883583.1 | Terrestrial | Arthropoda | Insecta |
| 2170 | Shahe narna-like virus 2 strain SHWC0209c12594 RNA-dependent RNA polymerase gene complete cds | KX883559.1 | Freshwater | Arthropoda | Crustacea |
| 2171 | Shahe narna-like virus 1 strain SHWC0209c12582 RNA-dependent RNA polymerase gene complete cds | KX883553.1 | Freshwater | Arthropoda | Crustacea |
| 2172 | Beihai narna-like virus 11 strain BWBFG39775 RNA-dependent RNA polymerase and hypothetical protein genes c | KX883513.1 | South China Sea | Mollusca | Bivalvia |
| 2173 | Beihai narna-like virus 23 strain BHZY57139 RNA-dependent RNA polymerase and hypothetical protein genes cor | KX883507.1 | South China Sea | Mollusca | Cephalopoda |
| 2174 | Beihai levi-like virus 23 strain BHJP62133 hypothetical protein genes complete cds | KX883474.1 | South China Sea | Echinodermata | Echinozoa |
| 2175 | Changjiang tombus-like virus 22 strain CJLX30074 hypothetical protein 1 and hypothetical protein 2 genes complet | KX883105.1 | Freshwater | Arthropoda | Crustacea |
| 2176 | Bivalvia Durna-like virus D9 | GWHAOZF01000000 | East China Sea | Mollusca | Bivalvia |
| 2177 | Bivalvia Durna-like virus H4 | GWHAPGS01000000 | Yellow China Sea | Mollusca | Bivalvia |
| 2178 | Beihai picobirna-like virus 12 strain BHZC36678 hypothetical protein and RdRp genes complete cds | KX884078.1 | South China Sea | Mollusca | Bivalvia |
| 2179 | Beihai paphia shell virus 3 strain BWBFG40853 hypothetical protein gene complete cds | KX884531.1 | South China Sea | Mollusca | Bivalvia |

| | | | | | |
|---|---|---|---|---|---|
| 2180 | Bivalvia Picorna-like virus N3 | GWHAPIT01000000 | South China Sea | Mollusca | Bivalvia |
| 2181 | Wenling reo-like virus 2 strain WLJQ99210 RdRp gene complete cds | KX884732.1 | East China Sea | Arthropoda | Crustacea |
| 2182 | Hubei leech virus 4 strain CJLX30730 hypothetical protein 1 and hypothetical protein 2 genes complete cds | KX884556.1 | Freshwater | Arthropoda | Crustacea |
| 2183 | Hubei leech virus 4 strain SZmix20870 hypothetical protein 1 and hypothetical protein 2 genes complete cds | KX883725.1 | Freshwater | Annelida | Clitellata |
| 2184 | Wenzhou channeled applesnail virus 2 strain WZFSL85846 hypothetical protein 1 and hypothetical protein 2 genes | KX884325.1 | Freshwater | Mollusca | Gastropoda |
| 2185 | Changjiang picorna-like virus 2 strain CJLX30436 hypothetical protein gene complete cds | KX884547.1 | Freshwater | Arthropoda | Crustacea |
| 2186 | Hubei tetragnatha maxillosa virus 9 strain arthropodmix14049 hypothetical protein 1 gene partial cds | KX884599.1 | Terrestrial | Arthropoda | Mix |
| 2187 | Hubei tetragnatha maxillosa virus 9 strain SSZZ3444 hypothetical protein 2 gene partial cds | KX884705.1 | Freshwater | Arthropoda | Chelicerata |
| 2188 | Hubei tetragnatha maxillosa virus 9 strain QTM27093 hypothetical protein 1 gene complete cds | KX884676.1 | Terrestrial | Arthropoda | Insecta |
| 2189 | Hubei tetragnatha maxillosa virus 9 strain SSZZ3444 RNA-dependent RNA polymerase gene complete cds | KX884703.1 | Freshwater | Arthropoda | Chelicerata |
| 2190 | Hubei tetragnatha maxillosa virus 9 strain arthropodmix14049 RNA-dependent RNA polymerase gene complete cds | KX884598.1 | Terrestrial | Arthropoda | Mix |
| 2191 | Hubei tetragnatha maxillosa virus 9 strain QTM27093 RNA-dependent RNA polymerase gene complete cds | KX884675.1 | Terrestrial | Arthropoda | Insecta |
| 2192 | Bole hyalomma asiaticum virus 1 strain tick109109 hypothetical protein gene partial cds | KX883729.1 | Terrestrial | Arthropoda | Chelicerata |
| 2193 | Bole hyalomma asiaticum virus 1 strain liyanpi109109 hypothetical protein gene partial cds | KX883914.1 | Terrestrial | Arthropoda | Chelicerata |
| 2194 | Hubei tick virus 1 strain tick109202 hypothetical protein gene complete cds | KX883731.1 | Terrestrial | Arthropoda | Chelicerata |
| 2195 | Hubei odonate virus 15 strain QTM133059 RNA-dependent RNA polymerase gene complete cds | KX884664.1 | Terrestrial | Arthropoda | Insecta |
| 2196 | Hubei virga-like virus 14 strain spider133314 hypothetical protein gene partial cds | KX883802.1 | Terrestrial | Arthropoda | Chelicerata |
| 2197 | Xingshan nematode virus 1 strain XSNXC21749 hypothetical protein genes complete cds | KX883837.1 | Terrestrial | Nematoda | Chromadorea |
| 2198 | Hubei virga-like virus 13 strain spider57988 hypothetical protein gene complete cds and hypothetical protein gene p | KX883803.1 | Terrestrial | Arthropoda | Chelicerata |
| 2199 | Hubei reo-like virus 12 strain mosHB234257 RdRp gene complete cds | KX884634.1 | Terrestrial | Arthropoda | Insecta |
| 2200 | Wenzhou noda-like virus 6 strain WZFSL85309 hypothetical protein gene complete cds | KX883260.1 | Freshwater | Mollusca | Gastropoda |
| 2201 | Hubei noda-like virus 21 strain WHBL59860 hypothetical protein gene complete cds | KX883206.1 | Freshwater | Mollusca | Bivalvia |
| 2202 | Hubei noda-like virus 22 strain WGML142106 hypothetical protein gene complete cds | KX883195.1 | Terrestrial | Arthropoda | Myriapoda |
| 2203 | Hubei noda-like virus 23 strain QTM26991 hypothetical protein gene complete cds | KX883137.1 | Terrestrial | Arthropoda | Insecta |
| 2204 | Beihai noda-like virus 8 strain BHWZXX12837 hypothetical protein 1 gene complete cds and hypothetical protein 2 | KX883045.1 | South China Sea | Arthropoda | Crustacea |
| 2205 | Beihai noda-like virus 9 strain BHZC37179 hypothetical protein 1 and hypothetical protein 2 genes complete cds | KX883061.1 | South China Sea | Mollusca | Bivalvia |
| 2206 | Wuhan house centipede virus 6 strain SCM50390 capsid protein gene complete cds | KX883148.1 | Terrestrial | Arthropoda | Insecta |
| 2207 | Wuhan house centipede virus 6 strain QTM26514 capsid protein gene complete cds | KX883133.1 | Terrestrial | Arthropoda | Insecta |
| 2208 | Wuhan house centipede virus 6 strain WGML146150 capsid protein gene complete cds | KX883199.1 | Terrestrial | Arthropoda | Myriapoda |
| 2209 | Wuhan house centipede virus 6 strain arthropodmix44324 hypothetical protein 1 and hypothetical protein 2 genes co | KX883012.1 | Terrestrial | Arthropoda | Mix |
| 2210 | Wuhan house centipede virus 6 strain WHWG55397 capsid protein and hypothetical protein genes complete cds | KX883242.1 | Terrestrial | Arthropoda | Myriapoda |
| 2211 | Wuhan house centipede virus 6 strain WHYY37237 capsid protein gene complete cds | KX883246.1 | Terrestrial | Arthropoda | Myriapoda |
| 2212 | Wuhan house centipede virus 6 strain CJLX30267 capsid protein gene complete cds | KX883093.1 | Freshwater | Arthropoda | Crustacea |
| 2213 | Hubei tombus-like virus 2 strain WHSFII19265 hypothetical protein 1 hypothetical protein 2 hypothetical protein 3 | KX883240.1 | Terrestrial | Arthropoda | Crustacea |
| 2214 | Bivalvia Picorna-like virus D53 | GWHAOZP01000000 | East China Sea | Mollusca | Bivalvia |
| 2215 | Hubei tombus-like virus 10 strain WHCC112361 hypothetical protein 1 and hypothetical protein 2 genes complete c | KX883224.1 | Terrestrial | Arthropoda | Insecta |
| 2216 | Beihai noda-like virus 4 strain BHNXC40905 hypothetical protein gene complete cds | KX883034.1 | South China Sea | Sipuncula | Sipuncula |

| | | | | | |
|---|---|---|---|---|---|
| 2217 | Hubei tombus-like virus 15 strain WHYY21098 hypothetical protein 1 hypothetical protein 2 and hypothetical prote | KX883247.1 | Terrestrial | Arthropoda | Myriapoda |
| 2218 | Bivalvia Picorna-like virus D57 | GWHAPAF01000000 | East China Sea | Mollusca | Bivalvia |
| 2219 | Hubei tombus-like virus 21 strain spider121926 hypothetical protein 1 hypothetical protein 2 and hypothetical prote | KX883160.1 | Terrestrial | Arthropoda | Chelicerata |
| 2220 | Hubei toti-like virus 14 strain QTM6211 hypothetical protein 1 gene partial cds and hypothetical protein 2 gene con | KX882969.1 | Terrestrial | Arthropoda | Insecta |
| 2221 | Bivalvia Picorna-like virus D18 | GWHAPKB01000000 | East China Sea | Mollusca | Bivalvia |
| 2222 | Shahe isopoda virus 3 strain SHWC0209c10670 RNA-dependent RNA polymerase and hypothetical protein genes c | KX884029.1 | Freshwater | Arthropoda | Crustacea |
| 2223 | Hubei sobemo-like virus 43 strain arthropodmix11560 hypothetical protein genes complete cds | KX882769.1 | Terrestrial | Arthropoda | Mix |
| 2224 | Wenling picorna-like virus 7 strain WLJQ102051 hypothetical protein 1 and hypothetical protein 2 genes complete | KX884314.1 | East China Sea | Arthropoda | Crustacea |
| 2225 | Beihai sobemo-like virus 7 strain BHZY60654 hypothetical protein 1 and hypothetical protein 2 genes complete cds | KX882804.1 | South China Sea | Mollusca | Cephalopoda |
| 2226 | Hubei narna-like virus 20 strain SCM42484 RNA-dependent RNA polymerase gene partial cds and hypothetical pro | KX883548.1 | Terrestrial | Arthropoda | Insecta |
| 2227 | Wenzhou yanvirus-like virus 1 strain WZFSL78003 RNA-dependent RNA polymerase and hypothetical protein gen | KX884031.1 | Freshwater | Mollusca | Gastropoda |
| 2228 | Hubei narna-like virus 13 strain QTM27075 RNA-dependent RNA polymerase gene complete cds | KX883543.1 | Terrestrial | Arthropoda | Insecta |
| 2229 | Beihai noda-like virus 12 strain BHZY60795 hypothetical protein gene partial cds | KX883075.1 | South China Sea | Mollusca | Cephalopoda |
| 2230 | Beihai barnacle virus 2 strain BHTH16123 RdRp and hypothetical protein genes complete cds | KX883747.1 | South China Sea | Arthropoda | Crustacea |
| 2231 | Cucumber mosaic virus strain WHWN52449 RNA-dependent RNA polymerase and RNA silencing suppressor gene | KX883817.1 | Freshwater | Mollusca | Gastropoda |
| 2232 | Cucumber mosaic virus RNA2 strain fly60335 protein 2a and protein 2b genes complete cds | KX883764.1 | Terrestrial | Arthropoda | Insecta |
| 2233 | Cucumber mosaic virus RNA3 strain fly60335 protein 3 and coat protein genes complete cds | KX883765.1 | Terrestrial | Arthropoda | Insecta |
| 2234 | Cucumber mosaic virus strain WHWN52449 movement protein and coat protein genes complete cds | KX883819.1 | Freshwater | Mollusca | Gastropoda |
| 2235 | Cucumber mosaic virus RNA1 strain fly60335 protein 1a gene complete cds | KX883763.1 | Terrestrial | Arthropoda | Insecta |
| 2236 | Cucumber mosaic virus strain WHWN52449 helicase gene complete cds | KX883818.1 | Freshwater | Mollusca | Gastropoda |
| 2237 | Hubei tombus-like virus 24 strain GCM76561 hypothetical protein 1 and hypothetical protein 2 genes complete cds | KX883108.1 | Terrestrial | Arthropoda | Insecta |
| 2238 | Crustacea Sobeli-like virus N1 | GWHAPFY01000000 | South China Sea | Arthropoda | Crustacea |
| 2239 | Wenzhou shrimp virus 9 strain shrimp6189 segment 1 hypothetical protein 1 and RdRp genes complete cds | KX882860.1 | East China Sea | Arthropoda | Crustacea |
| 2240 | Wenzhou picorna-like virus 33 strain WZFSL76563 hypothetical protein 1 and hypothetical protein 2 genes comple | KX884337.1 | Freshwater | Mollusca | Gastropoda |
| 2241 | Beihai picorna-like virus 78 strain BHBei73249 hypothetical protein 1 gene complete cds and hypothetical protein 2 | KX883307.1 | South China Sea | Mollusca | Bivalvia |
| 2242 | Wuhan insect virus 17 strain WHCCII1277 hypothetical protein 1 and hypothetical protein 2 genes complete cds | KX882910.1 | Terrestrial | Arthropoda | Insecta |
| 2243 | Shahe tombus-like virus 2 strain SHWC01c3797 hypothetical protein 1 and hypothetical protein 2 genes complete c | KX883153.1 | Freshwater | Arthropoda | Insecta |
| 2244 | Changjiang tombus-like virus 6 strain CJLX30707 hypothetical protein 1 and hypothetical protein 2 genes complete | KX883099.1 | Freshwater | Arthropoda | Crustacea |
| 2245 | Changjiang tombus-like virus 14 strain CJLX30318 hypothetical protein 1 and hypothetical protein 2 genes complet | KX883094.1 | Freshwater | Arthropoda | Crustacea |
| 2246 | Hubei tombus-like virus 9 strain WHBL63242 hypothetical protein 1 and hypothetical protein 2 genes complete cds | KX883209.1 | Freshwater | Mollusca | Bivalvia |
| 2247 | Wenzhou noda-like virus 3 strain shrimp14297 hypothetical protein gene complete cds | KX883151.1 | East China Sea | Arthropoda | Crustacea |
| 2248 | Hubei sobemo-like virus 26 strain arthropodmix14077 hypothetical protein genes complete cds | KX882775.1 | Terrestrial | Arthropoda | Mix |
| 2249 | Wenzhou tombus-like virus 2 strain WZFSL84050 hypothetical protein 1 and hypothetical protein 2 genes complete | KX883264.1 | Freshwater | Mollusca | Gastropoda |
| 2250 | Wenzhou tombus-like virus 3 strain WZFSL80503 hypothetical protein 1 hypothetical protein 2 and hypothetical pr | KX883266.1 | Freshwater | Mollusca | Gastropoda |
| 2251 | Hubei leech virus 1 strain SZmix32467 putative protease RNA-dependent RNA polymerase and putative capsid pro | KX884571.1 | Freshwater | Annelida | Clitellata |
| 2252 | Shahe narna-like virus 4 strain SHWC0209c11478 RNA-dependent RNA polymerase gene partial cds | KX883556.1 | Freshwater | Arthropoda | Crustacea |
| 2253 | Shahe narna-like virus 4 strain SHWCII4709 RNA-dependent RNA polymerase gene partial cds | KX883562.1 | Freshwater | Arthropoda | Mix |

| | | | | | |
|---|---|---|---|---|---|
| 2254 | Beihai weivirus-like virus 5 strain BHZY60693 RNA-dependent RNA polymerase and hypothetical protein genes co | KX884012.1 | South China Sea | Mollusca | Cephalopoda |
| 2255 | Wenzhou weivirus-like virus 1 strain beimix75799 RNA-dependent RNA polymerase and hypothetical protein gene | KX884006.1 | East China Sea | Mollusca | Bivalvia |
| 2256 | Beihai tombus-like virus 12 strain BHTSS17936 hypothetical protein 1 hypothetical protein 2 hypothetical protein 3 | KX883041.1 | South China Sea | Arthropoda | Crustacea |
| 2257 | Beihai weivirus-like virus 19 strain BHZY59816 RNA-dependent RNA polymerase gene complete cds and hypothe | KX884015.1 | South China Sea | Mollusca | Cephalopoda |
| 2258 | Beihai weivirus-like virus 21 strain BHZY60867 RNA-dependent RNA polymerase and hypothetical protein genes | KX884017.1 | South China Sea | Mollusca | Cephalopoda |
| 2259 | Beihai weivirus-like virus 20 strain BWBFG40173 RNA-dependent RNA polymerase and hypothetical protein gene | KX884025.1 | South China Sea | Mollusca | Bivalvia |
| 2260 | Gastropoda Picorna-like virus D13 | GWHAPAV01000000 | East China Sea | Mollusca | Gastropoda |
| 2261 | Beihai noda-like virus 14 strain BHHK130223 hypothetical protein gene complete cds | KX883030.1 | South China Sea | Cnidaria | Anthozoa |
| 2262 | Wenling noda-like virus 3 strain WLJQ104201 hypothetical protein gene complete cds | KX883253.1 | East China Sea | Arthropoda | Crustacea |
| 2263 | Beihai noda-like virus 16 strain HOU155673 hypothetical protein gene complete cds | KX883113.1 | South China Sea | Arthropoda | Chelicerata |
| 2264 | Xinzhou nematode virus 3 strain XZSJSC65579 RNA-dependent RNA polymerase gene complete cds | KX884004.1 | Terrestrial | Nematoda | Mix |
| 2265 | Wenling noda-like virus 2 strain WLJQ103411 hypothetical protein gene complete cds | KX883251.1 | East China Sea | Arthropoda | Crustacea |
| 2266 | Hubei noda-like virus 15 strain QTM162292 hypothetical protein gene complete cds | KX883140.1 | Terrestrial | Arthropoda | Insecta |
| 2267 | Hubei toti-like virus 12 strain WHZHC75812 hypothetical protein 1 gene complete cds and hypothetical protein 2 g | KX882982.1 | Terrestrial | Nematoda | Chromadorea |
| 2268 | Hubei toti-like virus 12 strain HC21305 hypothetical protein 1 and hypothetical protein 2 genes complete cds | KX882956.1 | Terrestrial | Nematoda | Rhabditida |
| 2269 | Beihai noda-like virus 27 strain BHHK108774 hypothetical protein gene partial cds | KX883024.1 | South China Sea | Cnidaria | Anthozoa |
| 2270 | Beihai sobemo-like virus 21 strain BHZY58647 hypothetical protein 1 and hypothetical protein 2 genes complete co | KX882799.1 | South China Sea | Mollusca | Cephalopoda |
| 2271 | Bivalvia Durna-like virus D15 | GWHAPHZ01000000 | East China Sea | Mollusca | Bivalvia |
| 2272 | Crustacea Unclassified virus D1 | GWHAPBP01000000 | East China Sea | Arthropoda | Crustacea |
| 2273 | Lishi Spider Virus 1 strain LSZZ11 segment 1 complete sequence | KM817597.1 | Terrestrial | Arthropoda | Chelicerata |
| 2274 | Shahe picorna-like virus 8 strain SHWCII5209 hypothetical protein 1 and hypothetical protein 2 genes complete cds | KX883670.1 | Freshwater | Arthropoda | Mix |
| 2275 | Shahe picorna-like virus 8 strain SHWC12398 hypothetical protein 1 and hypothetical protein 2 genes complete cds | KX883654.1 | Freshwater | Arthropoda | Mix |
| 2276 | Shahe picorna-like virus 8 strain SHWC0209c12692 hypothetical protein 1 and hypothetical protein 2 genes comple | KX883648.1 | Freshwater | Arthropoda | Crustacea |
| 2277 | Beihai narna-like virus 13 strain BHWZXX14912 RNA-dependent RNA polymerase gene complete cds | KX883498.1 | South China Sea | Arthropoda | Crustacea |
| 2278 | Beihai picobirna-like virus 6 strain BHJJX16459 RdRp gene complete cds | KX884057.1 | South China Sea | Arthropoda | Crustacea |
| 2279 | Hubei narna-like virus 22 strain SZmix20730 RNA-dependent RNA polymerase gene complete cds | KX883572.1 | Freshwater | Annelida | Clitellata |
| 2280 | Wenzhou picorna-like virus 47 strain WHCCII11151 hypothetical protein gene complete cds | KX884273.1 | Terrestrial | Arthropoda | Insecta |
| 2281 | Wenzhou picorna-like virus 47 strain WZFSL80701 hypothetical protein gene complete cds | KX884323.1 | Freshwater | Mollusca | Gastropoda |
| 2282 | Wenzhou channeled applesnail virus 1 strain WZFSL85493 hypothetical protein 1 and hypothetical protein 2 genes | KX884329.1 | Freshwater | Mollusca | Gastropoda |
| 2283 | Hubei odonate virus 15 strain QTM133059 hypothetical protein 2 gene complete cds | KX884666.1 | Terrestrial | Arthropoda | Insecta |
| 2284 | Shahe picobirna-like virus 2 strain SHWC01c3537 hypothetical protein and RdRp genes complete cds | KX884154.1 | Freshwater | Arthropoda | Insecta |
| 2285 | Wenzhou picorna-like virus 17 strain WZSBei69283 hypothetical protein 1 and hypothetical protein 2 genes comple | KX884358.1 | East China Sea | Mollusca | Bivalvia |
| 2286 | Hubei blood fluke virus 3 strain JSC69362 hypothetical protein 1 gene complete cds | KX884609.1 | Freshwater | Platyhelminthes | Trematoda |
| 2287 | Hubei blood fluke virus 3 strain RBXXC337523 hypothetical protein 1 gene complete cds | KX884688.1 | Freshwater | Platyhelminthes | Trematoda |
| 2288 | Wenzhou picorna-like virus 41 strain WZFSL80064 hypothetical protein 1 and hypothetical protein 2 genes comple | KX884336.1 | Freshwater | Mollusca | Gastropoda |
| 2289 | Wuhan cricket virus strain WHXS-1 segment 3 complete sequence | KR902711.1 | Terrestrial | Arthropoda | Insecta |
| 2290 | Bivalvia Picorna-like virus N31 | GWHAPLL01000000 | South China Sea | Mollusca | Bivalvia |

| | | | | | |
|---|---|---|---|---|---|
| 2291 | Hubei myriapoda virus 3 strain WGML139652 hypothetical protein 1 hypothetical protein 2 hypothetical protein 3 a | KX883734.1 | Terrestrial | Arthropoda | Myriapoda |
| 2292 | Hubei reo-like virus 4 strain WGML144712 hypothetical protein 2 gene complete cds | KX884714.1 | Terrestrial | Arthropoda | Myriapoda |
| 2293 | Hubei picorna-like virus 54 strain WHSFII20272 hypothetical protein gene complete cds | KX884288.1 | Terrestrial | Arthropoda | Crustacea |
| 2294 | Hubei picorna-like virus 54 strain ZCM19837 hypothetical protein gene complete cds | KX884396.1 | Terrestrial | Arthropoda | Insecta |
| 2295 | Hubei picorna-like virus 54 strain WGML144949 hypothetical protein gene complete cds | KX884250.1 | Terrestrial | Arthropoda | Myriapoda |
| 2296 | Hubei diptera virus 20 strain SCM51404 hypothetical protein 2 gene complete cds | KX884695.1 | Terrestrial | Arthropoda | Insecta |
| 2297 | Hubei diptera virus 20 strain SCM51404 hypothetical protein 1 gene complete cds | KX884694.1 | Terrestrial | Arthropoda | Insecta |
| 2298 | Hubei picorna-like virus 51 strain QTM27291 hypothetical protein 1 and hypothetical protein 2 genes complete cds | KX883953.1 | Terrestrial | Arthropoda | Insecta |
| 2299 | Wuhan heteroptera virus 3 strain arthropodmix44000 RNA-dependent RNA polymerase gene complete cds | KX884588.1 | Terrestrial | Arthropoda | Mix |
| 2300 | Wuhan heteroptera virus 3 strain WHYCM468 RNA-dependent RNA polymerase gene complete cds | KX884721.1 | Terrestrial | Arthropoda | Insecta |
| 2301 | Wuhan spider virus 6 strain spider122561 hypothetical protein 1 hypothetical protein 2 hypothetical protein 3 and h | KX883680.1 | Terrestrial | Arthropoda | Chelicerata |
| 2302 | Wuhan Louse Fly Virus 5 strain BFJSC-5 nucleocapsid (N) phosphoprotein (P) matrix protein (M) glycoprotein (G) | KM817654.1 | Terrestrial | Arthropoda | Insecta |
| 2303 | Hubei reo-like virus 3 strain SCM43983 RdRp gene complete cds | KX884692.1 | Terrestrial | Arthropoda | Insecta |
| 2304 | Beihai picorna-like virus 43 strain BHBJDX18749 hypothetical protein 1 and hypothetical protein 2 genes complete | KX883335.1 | South China Sea | Arthropoda | Crustacea |
| 2305 | Beihai picorna-like virus 43 strain BHWZXX15634 hypothetical protein 1 and hypothetical protein 2 genes comple | KX883429.1 | South China Sea | Arthropoda | Crustacea |
| 2306 | Beihai picorna-like virus 43 strain BHNXC41212 hypothetical protein 1 and hypothetical protein 2 genes complete | KX883390.1 | South China Sea | Sipuncula | Sipuncula |
| 2307 | Beihai picorna-like virus 43 strain BHJJX25682 hypothetical protein 1 and hypothetical protein 2 genes complete c | KX883367.1 | South China Sea | Arthropoda | Crustacea |
| 2308 | Beihai picorna-like virus 43 strain SCJXSX39249 hypothetical protein 1 and hypothetical protein 2 genes complete | KX883962.1 | South China Sea | Arthropoda | Crustacea |
| 2309 | Hubei reo-like virus 6 strain WGML611819 hypothetical protein 3 gene complete cds | KX884719.1 | Terrestrial | Arthropoda | Myriapoda |
| 2310 | Beihai zhaovirus-like virus 2 strain BHTSS12281 polyprotein gene complete cds | KX884037.1 | South China Sea | Arthropoda | Crustacea |
| 2311 | Hubei tetragnatha maxillosa virus 5 strain SSZZ3471 hypothetical protein gene complete cds | KX883700.1 | Freshwater | Arthropoda | Chelicerata |
| 2312 | Hubei reo-like virus 11 strain arthropodmix13320 Major core capsid gene complete cds | KX884576.1 | Terrestrial | Arthropoda | Mix |
| 2313 | Hubei reo-like virus 11 strain QTM23921 major core capsid gene complete cds | KX884673.1 | Terrestrial | Arthropoda | Insecta |
| 2314 | Hubei picorna-like virus 33 strain QTM27281 hypothetical protein gene complete cds | KX883949.1 | Terrestrial | Arthropoda | Insecta |
| 2315 | Hubei odonate virus 14 strain QTM27275 Major core protein gene complete cds | KX884684.1 | Terrestrial | Arthropoda | Insecta |
| 2316 | Changjiang picorna-like virus 15 strain CJLX30633 hypothetical protein gene complete cds | KX884552.1 | Freshwater | Arthropoda | Crustacea |
| 2317 | Hubei reo-like virus 10 strain QTM136127 RNA-dependent RNA polymerase gene complete cds | KX884669.1 | Terrestrial | Arthropoda | Insecta |
| 2318 | Shuangao insect virus 12 strain insectZJ98124 hypothetical protein gene complete cds | KX883912.1 | Terrestrial | Arthropoda | Insecta |
| 2319 | Wuhan coneheads virus 1 strain ZCM4592 hypothetical protein gene complete cds | KX884402.1 | Terrestrial | Arthropoda | Insecta |
| 2320 | Wuhan coneheads virus 1 strain WHLC897 hypothetical protein gene complete cds | KX884279.1 | Terrestrial | Arthropoda | Insecta |
| 2321 | Sanxia reo-like virus 1 strain SXSSP2271 RdRp gene partial cds | KX884711.1 | Freshwater | Arthropoda | Insecta |
| 2322 | Shuangao sobemo-like virus 3 strain insectZJ66198 hypothetical protein 1 and hypothetical protein 2 genes complet | KX882823.1 | Terrestrial | Arthropoda | Insecta |
| 2323 | Wuhan arthropod virus 3 strain WHCC110739 hypothetical protein 1 hypothetical protein 2 and hypothetical protei | KX884267.1 | Terrestrial | Arthropoda | Insecta |
| 2324 | Hubei reo-like virus 11 strain QTM23921 minor core capsid gene complete cds | KX884674.1 | Terrestrial | Arthropoda | Insecta |
| 2325 | Hubei reo-like virus 11 strain arthropodmix13320 Minor core capsid 2 gene complete cds | KX884577.1 | Terrestrial | Arthropoda | Mix |
| 2326 | Wuhan fly virus 4 strain fly116586 hypothetical protein gene complete cds | KX883891.1 | Terrestrial | Arthropoda | Insecta |
| 2327 | Beihai picorna-like virus 44 strain BHXG26143 hypothetical protein 1 and hypothetical protein 2 genes complete cd | KX884466.1 | South China Sea | Arthropoda | Crustacea |

| | | | | | |
|---|---|---|---|---|---|
| 2328 | Changjiang Jingmen-like virus strain CJLX28200 NS5-like protein gene complete cds | KX883002.1 | Freshwater | Arthropoda | Crustacea |
| 2329 | Wenling picorna-like virus 5 strain WLJQ102796 hypothetical protein 1 and hypothetical protein 2 genes complete | KX884316.1 | East China Sea | Arthropoda | Crustacea |
| 2330 | Shuangao insect virus 8 strain insectZJ96360 hypothetical protein 1 hypothetical protein 2 hypothetical protein 3 an | KX883910.1 | Terrestrial | Arthropoda | Insecta |
| 2331 | Hubei reo-like virus 7 strain mosHB235771 RdRp gene complete cds | KX884635.1 | Terrestrial | Arthropoda | Insecta |
| 2332 | Hubei reo-like virus 6 strain WGML611819 hypothetical protein 2 gene complete cds | KX884718.1 | Terrestrial | Arthropoda | Myriapoda |
| 2333 | Beihai picorna-like virus 77 strain BHBei75652 hypothetical protein 1 and hypothetical protein 2 genes complete cc | KX883310.1 | South China Sea | Mollusca | Bivalvia |
| 2334 | Hubei chuvirus-like virus 1 strain QTM26249 putative glycoprotein and hypothetical protein genes complete cds | KX884425.1 | Terrestrial | Arthropoda | Insecta |
| 2335 | Changjiang picorna-like virus 17 strain CJLX30705 hypothetical protein gene partial cds | KX884555.1 | Freshwater | Arthropoda | Crustacea |
| 2336 | Wenzhou picorna-like virus 40 strain WZFSL83107 hypothetical protein 1 and hypothetical protein 2 genes comple | KX884334.1 | Freshwater | Mollusca | Gastropoda |
| 2337 | Hubei lepidoptera virus 3 strain LCM141729 Cypovirus VP4 gene complete cds | KX884616.1 | Terrestrial | Arthropoda | Insecta |
| 2338 | Hubei picorna-like virus 24 strain WGML142000 hypothetical protein 1 and hypothetical protein 2 genes complete | KX883736.1 | Terrestrial | Arthropoda | Myriapoda |
| 2339 | Wenzhou narna-like virus 7 strain WZFSL82100 RNA-dependent RNA polymerase gene complete cds | KX883614.1 | Freshwater | Mollusca | Gastropoda |
| 2340 | Wuhan cricket virus strain WHXS-1 segment 1 complete sequence | KR902709.1 | Terrestrial | Arthropoda | Insecta |
| 2341 | Wenling crustacean virus 8 strain WLJQ104142 RNA-dependent RNA polymerase gene complete cds | KX884861.1 | East China Sea | Arthropoda | Crustacea |
| 2342 | Wuhan Mosquito Virus 1 strain WT3-15 glycoprotein precursor (G) gene complete cds | KM817726.1 | Terrestrial | Arthropoda | Insecta |
| 2343 | Shuangao Insect Virus 5 strain QSA09 polymerase (L) gene complete cds | KM817614.1 | Terrestrial | Arthropoda | Insecta |
| 2344 | Hubei reo-like virus 11 strain QTM23921 RNA-binding protein gene complete cds | KX884672.1 | Terrestrial | Arthropoda | Insecta |
| 2345 | Hubei reo-like virus 11 strain arthropodmix13320 RNA-binding protein gene complete cds | KX884575.1 | Terrestrial | Arthropoda | Mix |
| 2346 | Hubei picorna-like virus 31 strain QTM26628 hypothetical protein gene complete cds | KX883927.1 | Terrestrial | Arthropoda | Insecta |
| 2347 | Changjiang crawfish virus 5 strain CJLX22437 hypothetical protein 1 and hypothetical protein 2 genes complete cd: | KX884539.1 | Freshwater | Arthropoda | Crustacea |
| 2348 | Wenzhou tombus-like virus 13 strain WZFSL80420 hypothetical protein 1 and hypothetical protein 2 genes comple | KX883267.1 | Freshwater | Mollusca | Gastropoda |
| 2349 | Hubei blood fluke virus 3 strain JSC69362 hypothetical protein 2 gene complete cds | KX884610.1 | Freshwater | Platyhelminthes | Trematoda |
| 2350 | Hubei blood fluke virus 3 strain RBXXC337523 hypothetical protein 2 gene complete cds | KX884689.1 | Freshwater | Platyhelminthes | Trematoda |
| 2351 | Hubei tombus-like virus 29 strain spider124630 hypothetical protein and hypothetical protein 2 genes complete cds | KX883157.1 | Terrestrial | Arthropoda | Chelicerata |
| 2352 | Hubei odonate virus 5 strain QTM27277 hypothetical protein gene complete cds | KX883948.1 | Terrestrial | Arthropoda | Insecta |
| 2353 | Shuangao Bedbug Virus 1 strain HKCC-1 glycoprotein precursor (G) gene complete cds | KM817713.1 | Terrestrial | Arthropoda | Insecta |
| 2354 | Wenling crustacean virus 2 strain WLJQ102135 hypothetical protein 1 and hypothetical protein 2 genes complete cc | KX884315.1 | East China Sea | Arthropoda | Crustacea |
| 2355 | Xinzhou Spider Virus strain XZZZ-2 RNA-dependent RNA polymerase (L) gene complete cds | KM817702.1 | Terrestrial | Arthropoda | Chelicerata |
| 2356 | Xinzhou Spider Virus strain XZZZ-2 glycoprotein precursor (G) gene partial cds | KM817729.1 | Terrestrial | Arthropoda | Chelicerata |
| 2357 | Shuangao Bedbug Virus 2 strain SACC-1 ORF1 (ORF1) glycoprotein (G) and RNA-dependent RNA polymerase (I | KM817637.1 | Terrestrial | Arthropoda | Insecta |
| 2358 | Hubei rhabdo-like virus 7 strain QTM26925 hypothetical protein putative glycoprotein and RNA-dependent RNA p | KX884428.1 | Terrestrial | Arthropoda | Insecta |
| 2359 | Jiangxia Mosquito Virus 2 strain JX1-14 RNA-dependent RNA polymerase (L) gene complete cds | KM817671.1 | Terrestrial | Arthropoda | Insecta |
| 2360 | Shahe bunya-like virus 2 strain SHWC0209c12742 RNA-dependent RNA polymerase gene complete cds | KX884822.1 | Freshwater | Arthropoda | Crustacea |
| 2361 | Shayang Spider Virus 1 strain SYZZ-4 glycoprotein precursor (G) gene complete cds | KM817712.1 | Terrestrial | Arthropoda | Chelicerata |
| 2362 | Shahe isopoda virus 4 strain SHWC0209c12776 RNA-dependent RNA polymerase gene complete cds | KX884816.1 | Freshwater | Arthropoda | Crustacea |
| 2363 | Shahe isopoda virus 4 strain SHWCII5333 RNA-dependent RNA polymerase gene complete cds | KX884826.1 | Freshwater | Arthropoda | Mix |
| 2364 | Shahe bunya-like virus 2 strain SHWC0209c12742 putative glycoprotein gene complete cds | KX884823.1 | Freshwater | Arthropoda | Crustacea |

| | | | | | |
|---|---|---|---|---|---|
| 2365 | Shuangao Bedbug Virus 1 strain HKCC-1 RNA-dependent RNA polymerase (L) gene complete cds | KM817678.1 | Terrestrial | Arthropoda | Insecta |
| 2366 | Shuangao Mosquito Virus strain WSA02 RNA-dependent RNA polymerase (L) gene partial cds | KM817682.1 | Terrestrial | Arthropoda | Insecta |
| 2367 | Hubei myriapoda virus 5 strain WGML140132 RNA-dependent RNA polymerase gene complete cds | KX884840.1 | Terrestrial | Arthropoda | Myriapoda |
| 2368 | Hubei myriapoda virus 5 strain GCM10499 RNA-dependent RNA polymerase gene complete cds | KX884758.1 | Terrestrial | Arthropoda | Insecta |
| 2369 | Lishi Spider Virus 2 strain LSZZ-4 ORF1 (ORF1) ORF2 (ORF2) and RNA-dependent RNA polymerase (L) genes c | KM817632.1 | Terrestrial | Arthropoda | Chelicerata |
| 2370 | Hubei lepidoptera virus 1 strain LCM141331 putative glycoprotein gene complete cds | KX884773.1 | Terrestrial | Arthropoda | Insecta |
| 2371 | Sanxia Water Strider Virus 2 strain SXSSP02 glycoprotein precursor (G) gene complete cds | KX650646.1 | Freshwater | Arthropoda | Insecta |
| 2372 | Hubei partiti-like virus 59 strain SCM33193 RdRp gene complete cds | KX884149.1 | Terrestrial | Arthropoda | Insecta |
| 2373 | Hubei picorna-like virus 42 strain arthropodmix13971 hypothetical protein 1 and hypothetical protein 2 genes comp | KX883291.1 | Terrestrial | Arthropoda | Mix |
| 2374 | Hubei picorna-like virus 42 strain WHYCM1191 hypothetical protein gene complete cds | KX884297.1 | Terrestrial | Arthropoda | Insecta |
| 2375 | Hubei sobemo-like virus 34 strain SSZZ41 hypothetical protein 1 and hypothetical protein 2 genes complete cds | KX882879.1 | Freshwater | Arthropoda | Chelicerata |
| 2376 | Wenzhou picorna-like virus 24 strain WZSBei69493 hypothetical protein gene complete cds | KX884362.1 | East China Sea | Mollusca | Bivalvia |
| 2377 | Wenling narna-like virus 9 strain WLJQ205243 RNA-dependent RNA polymerase gene complete cds | KX883604.1 | East China Sea | Arthropoda | Crustacea |
| 2378 | Hubei tombus-like virus 35 strain WGML3549 hypothetical protein 1 gene complete cds and hypothetical protein 2 | KX883202.1 | Terrestrial | Arthropoda | Myriapoda |
| 2379 | Changjiang picorna-like virus 13 strain CJLX40994 hypothetical protein 1 and hypothetical protein 2 genes comple | KX884563.1 | Freshwater | Arthropoda | Crustacea |
| 2380 | Sanxia water strider virus 9 strain SXSSP2367 hypothetical protein gene complete cds | KX883704.1 | Freshwater | Arthropoda | Insecta |
| 2381 | Beihai reo-like virus 1 strain BHXG2356 hypothetical protein 1 and RdRp genes complete cds | KX884606.1 | South China Sea | Arthropoda | Crustacea |
| 2382 | Shahe narna-like virus 3 strain SHWC01c3372 RNA-dependent RNA polymerase gene complete cds | KX883552.1 | Freshwater | Arthropoda | Insecta |
| 2383 | Shahe picorna-like virus 12 strain SHWC0209c11951 hypothetical protein 1 and hypothetical protein 2 genes comp | KX883650.1 | Freshwater | Arthropoda | Crustacea |
| 2384 | Hubei picorna-like virus 78 strain WHSF6879 hypothetical protein 1 and hypothetical protein 2 genes complete cds | KX884283.1 | Terrestrial | Arthropoda | Crustacea |
| 2385 | Shahe picobirna-like virus 1 strain SHWC0209c12590 hypothetical protein and RdRp genes complete cds | KX884156.1 | Freshwater | Arthropoda | Crustacea |
| 2386 | Sanxia Water Strider Virus 1 strain SXSSP08 glycoprotein precursor (G) gene complete cds | KM817711.1 | Freshwater | Arthropoda | Insecta |
| 2387 | Sanxia Water Strider Virus 1 strain SXXX37846 putative glycoprotein gene complete cds | KX884833.1 | Freshwater | Arthropoda | Crustacea |
| 2388 | Hubei rhabdo-like virus 6 strain QTM24798 hypothetical protein and RNA-dependent RNA polymerase genes com | KX884421.1 | Terrestrial | Arthropoda | Insecta |
| 2389 | Wenling crustacean virus 9 strain WLJQ100911 putative glycoprotein gene complete cds | KX884857.1 | East China Sea | Arthropoda | Crustacea |
| 2390 | Wuhan Millipede Virus 2 strain WHWG03 glycoprotein precursor (G) gene complete cds | KX650645.1 | Terrestrial | Arthropoda | Insecta |
| 2391 | Bivalvia Picorna-like virus D61 | GWHAOZZ01000000 | East China Sea | Mollusca | Bivalvia |
| 2392 | Wenzhou picorna-like virus 5 strain BHZC35527 hypothetical protein 1 gene complete cds and hypothetical protein | KX884476.1 | South China Sea | Mollusca | Bivalvia |
| 2393 | Changjiang picorna-like virus 12 strain CJLX29956 hypothetical protein 1 and hypothetical protein 2 genes comple | KX884545.1 | Freshwater | Arthropoda | Crustacea |
| 2394 | Wuhan insect virus 25 strain CC64706 RdRp gene complete cds | KX884087.1 | Terrestrial | Arthropoda | Insecta |
| 2395 | Beihai picorna-like virus 50 strain BHZC37234 hypothetical protein gene complete cds | KX884484.1 | South China Sea | Mollusca | Bivalvia |
| 2396 | Beihai picorna-like virus 50 strain BHBei76574 hypothetical protein gene complete cds | KX883311.1 | South China Sea | Mollusca | Bivalvia |
| 2397 | Beihai picobirna-like virus 7 strain BHNXC57916 segment Seg 2 hypothetical protein gene complete cds | KX884063.1 | South China Sea | Sipuncula | Sipuncula |
| 2398 | Kadipiro virus strain QTM27331 Kadipiro VP4 gene complete cds | KX884653.1 | Terrestrial | Arthropoda | Insecta |
| 2399 | Sanxia picorna-like virus 8 strain SXXX35540 hypothetical protein gene complete cds | KX883720.1 | Freshwater | Arthropoda | Crustacea |
| 2400 | Shuangao Insect Virus 1 strain QSA02 glycoprotein precursor (G) gene complete cds | KM817714.1 | Terrestrial | Arthropoda | Insecta |
| 2401 | Shuangao Insect Virus 1 strain QSA02 RNA-dependent RNA polymerase (L) gene complete cds | KM817679.1 | Terrestrial | Arthropoda | Insecta |

| | | | | | |
|---|---|---|---|---|---|
| ***2402*** | Wenling crustacean virus 8 strain WLJQ104142 putative glycoprotein gene complete cds | KX884862.1 | East China Sea | Arthropoda | Crustacea |
| ***2403*** | Hubei orthoptera virus 2 strain ZCM20815 RNA-dependent RNA polymerase gene complete cds | KX884871.1 | Terrestrial | Arthropoda | Insecta |
| ***2404*** | Hubei orthoptera virus 2 strain ZCM20815 putative glycoprotein gene complete cds | KX884872.1 | Terrestrial | Arthropoda | Insecta |
| ***2405*** | Bivalvia Picorna-like virus N46 | GWHAPDY01000000 | South China Sea | Mollusca | Bivalvia |
| ***2406*** | Beihai sphaeromadae virus 1 strain BHTSS18012 hypothetical protein gene complete cds | KX883422.1 | South China Sea | Arthropoda | Crustacea |
| ***2407*** | Hubei myriapoda virus 5 strain GCM10499 putative glycoprotein gene complete cds | KX884759.1 | Terrestrial | Arthropoda | Insecta |
| ***2408*** | Hubei myriapoda virus 5 strain WGML140132 putative glycoprotein gene complete cds | KX884841.1 | Terrestrial | Arthropoda | Myriapoda |
| ***2409*** | Wenzhou picorna-like virus 48 strain beimix75770 hypothetical protein gene complete cds | KX883301.1 | East China Sea | Mollusca | Bivalvia |
| ***2410*** | Wuhan cricket virus strain WHXS-1 segment 4 complete sequence | KR902712.1 | Terrestrial | Arthropoda | Insecta |
| ***2411*** | Kadipiro virus strain QTM27331 Kadipiro VP3 gene complete cds | KX884652.1 | Terrestrial | Arthropoda | Insecta |
| ***2412*** | Sanxia picorna-like virus 4 strain SXXX37816 hypothetical protein gene complete cds | KX883710.1 | Freshwater | Arthropoda | Crustacea |
| ***2413*** | Hubei myriapoda virus 1 strain WGML505798 hypothetical protein gene complete cds | KX884260.1 | Terrestrial | Arthropoda | Myriapoda |
| ***2414*** | Hubei reo-like virus 2 strain QTM25955 RNA-dependent RNA polymerase gene complete cds | KX884662.1 | Terrestrial | Arthropoda | Insecta |
| ***2415*** | Hubei picorna-like virus 40 strain WHYY23452 hypothetical protein gene complete cds | KX884303.1 | Terrestrial | Arthropoda | Myriapoda |
| ***2416*** | Hubei odonate virus 14 strain QTM27275 hypothetical protein gene complete cds | KX884685.1 | Terrestrial | Arthropoda | Insecta |
| ***2417*** | Shahe heteroptera virus 4 strain SHWC01c2975 hypothetical protein 1 and hypothetical protein 2 genes complete c | KX883640.1 | Freshwater | Arthropoda | Insecta |
| ***2418*** | Hubei orthoptera virus 1 strain ZCM21319 hypothetical protein 1 and hypothetical protein 2 genes complete cds | KX884400.1 | Terrestrial | Arthropoda | Insecta |
| ***2419*** | Hubei lepidoptera virus 3 strain LCM141729 Cypovirus VP5 gene complete cds | KX884617.1 | Terrestrial | Arthropoda | Insecta |
| ***2420*** | Wenzhou picorna-like virus 2 strain WZSBei58345 hypothetical protein 1 and hypothetical protein 2 genes complet | KX884353.1 | East China Sea | Mollusca | Bivalvia |
| ***2421*** | Wenzhou picorna-like virus 2 strain beimix73672 hypothetical protein 1 and hypothetical protein 2 genes complete | KX883299.1 | East China Sea | Mollusca | Bivalvia |
| ***2422*** | Hubei diptera virus 21 strain SCM95013 hypothetical protein 3 gene complete cds | KX884699.1 | Terrestrial | Arthropoda | Insecta |
| ***2423*** | Hubei odonate virus 4 strain QTM161803 hypothetical protein gene complete cds | KX883956.1 | Terrestrial | Arthropoda | Insecta |
| ***2424*** | Hubei diptera virus 21 strain SCM95013 RNA-dependent RNA polymerase gene complete cds | KX884696.1 | Terrestrial | Arthropoda | Insecta |
| ***2425*** | Wuhan arthropod virus 2 strain WHSFII47608 hypothetical protein 1 and hypothetical protein 2 genes complete cds | KX884287.1 | Terrestrial | Arthropoda | Crustacea |
| ***2426*** | Wuhan arthropod virus 2 strain WHSF0 hypothetical protein 1 and hypothetical protein 2 genes complete cds | KX884282.1 | Terrestrial | Arthropoda | Crustacea |
| ***2427*** | Wuhan arthropod virus 2 strain WHCC257917 hypothetical protein 1 and hypothetical protein 2 genes complete cds | KX884271.1 | Terrestrial | Arthropoda | Insecta |
| ***2428*** | Hubei diptera virus 21 strain SCM95013 hypothetical protein 1 gene complete cds | KX884697.1 | Terrestrial | Arthropoda | Insecta |
| ***2429*** | Wenzhou picorna-like virus 36 strain WZRBX14225 hypothetical protein gene complete cds | KX884343.1 | East China Sea | Arthropoda | Crustacea |
| ***2430*** | Beihai sobemo-like virus 23 strain BHZY181484 hypothetical protein 1 and hypothetical protein 2 genes complete | KX882808.1 | South China Sea | Mollusca | Cephalopoda |
| ***2431*** | Beihai picorna-like virus 122 strain BHZY60922 hypothetical protein gene complete cds | KX884521.1 | South China Sea | Mollusca | Cephalopoda |
| ***2432*** | Wenling picorna-like virus 6 strain WLJQ101512 hypothetical protein gene complete cds | KX884313.1 | East China Sea | Arthropoda | Crustacea |
| ***2433*** | Hubei sobemo-like virus 19 strain SCM51527 hypothetical protein 1 and hypothetical protein 2 genes complete cds | KX882855.1 | Terrestrial | Arthropoda | Insecta |
| ***2434*** | Wuhan aphid virus 2 strain WHYC-2 segment 4 complete sequence | KR902728.1 | Terrestrial | Arthropoda | Insecta |
| ***2435*** | Hubei picorna-like virus 50 strain spider113255 hypothetical protein gene complete cds | KX883679.1 | Terrestrial | Arthropoda | Chelicerata |
| ***2436*** | Hubei noda-like virus 24 strain WHBL73074 hypothetical protein 1 and hypothetical protein 2 genes complete cds | KX883219.1 | Freshwater | Mollusca | Bivalvia |
| ***2437*** | Shuangao Insect Virus 2 strain QSA03 RNA-dependent RNA polymerase (L) gene partial cds | KM817680.1 | Terrestrial | Arthropoda | Insecta |
| ***2438*** | Changping bunya-like virus 1 strain CPQY18609 RNA-dependent RNA polymerase gene partial cds | KX884757.1 | Terrestrial | Annelida | Clitellata |

| | | | | | |
|---|---|---|---|---|---|
| 2439 | Bivalvia Picorna-like virus D78 | GWHAPID01000000 | East China Sea | Mollusca | Bivalvia |
| 2440 | Changjiang crawfish virus 1 strain CJLX30787 hypothetical protein 1 and hypothetical protein 2 genes complete cds | KX884562.1 | Freshwater | Arthropoda | Crustacea |
| 2441 | Hubei insect virus 2 strain arthropodmix13736 hypothetical protein 3 gene complete cds | KX884583.1 | Terrestrial | Arthropoda | Mix |
| 2442 | Shahe heteroptera virus 3 strain SHWC01c3680 putative glycoprotein gene complete cds | KX884814.1 | Freshwater | Arthropoda | Insecta |
| 2443 | Shahe heteroptera virus 3 strain SHWCII5319 putative glycoprotein gene complete cds | KX884830.1 | Freshwater | Arthropoda | Mix |
| 2444 | Shahe heteroptera virus 3 strain SHWC0209c8767 putative glycoprotein gene complete cds | KX884819.1 | Freshwater | Arthropoda | Crustacea |
| 2445 | Wuhan snail virus 2 strain WHWN46660 RNA-dependent RNA polymerase gene complete cds | KX884852.1 | Freshwater | Mollusca | Gastropoda |
| 2446 | Hubei myriapoda virus 6 strain WGML140341 putative glycoprotein gene complete cds | KX884844.1 | Terrestrial | Arthropoda | Myriapoda |
| 2447 | Hubei bunya-like virus 7 strain WGML112345 putative glycoprotein gene complete cds | KX884839.1 | Terrestrial | Arthropoda | Myriapoda |
| 2448 | Wenzhou channeled applesnail virus 4 strain WZFSL85898 RNA-dependent RNA polymerase gene complete cds | KX884863.1 | Freshwater | Mollusca | Gastropoda |
| 2449 | Beihai sipunculid worm virus 7 strain BHNXC41490 RNA-dependent RNA polymerase gene complete cds | KX884745.1 | South China Sea | Sipuncula | Sipuncula |
| 2450 | Wuhan snail virus 2 strain WHWN46660 putative glycoprotein gene complete cds | KX884853.1 | Freshwater | Mollusca | Gastropoda |
| 2451 | Whenzhou Shrimp Virus 2 strain BJDX-1 glycoprotein precursor (G) gene complete cds | KM817720.2 | East China Sea | Arthropoda | Crustacea |
| 2452 | Wenzhou bunya-like virus 1 strain WZSLuoI85881 RNA-dependent RNA polymerase gene complete cds | KX884865.1 | East China Sea | Mollusca | Gastropoda |
| 2453 | Wenling chuvirus-like virus 1 strain WLJQ101487 RNA-dependent RNA polymerase gene complete cds | KX884449.1 | East China Sea | Arthropoda | Crustacea |
| 2454 | Wuhan Mosquito Virus 2 strain QN2-7 glycoprotein precursor (G) gene complete cds | KM817727.1 | Terrestrial | Arthropoda | Insecta |
| 2455 | Beihai picorna-like virus 120 strain BHSC167783 hypothetical protein 1 hypothetical protein 2 hypothetical protein | KX883411.1 | South China Sea | Annelida | Polychaeta |
| 2456 | Wuhan heteroptera virus 2 strain SSZZ3534 segment 2 hypothetical protein gene complete cds | KX882881.1 | Freshwater | Arthropoda | Chelicerata |
| 2457 | Wuhan heteroptera virus 2 strain WHYCM467 segment 2 hypothetical protein gene complete cds | KX882918.1 | Terrestrial | Arthropoda | Insecta |
| 2458 | Wuhan heteroptera virus 2 strain arthropodmix10101 segment 2 hypothetical protein gene complete cds | KX882768.1 | Terrestrial | Arthropoda | Mix |
| 2459 | Hubei tombus-like virus 39 strain WGML9144 hypothetical protein 1 hypothetical protein 2 and hypothetical protei | KX883204.1 | Terrestrial | Arthropoda | Myriapoda |
| 2460 | Wenling crustacean virus 12 strain WLJQ47777 putative nucleoprotein hypothetical protein and RNA-dependent R | KX884457.1 | East China Sea | Arthropoda | Crustacea |
| 2461 | Wuhan flea virus strain WHZM segment 4 complete sequence | KR902716.1 | Terrestrial | Arthropoda | Insecta |
| 2462 | Hubei tombus-like virus 11 strain WHSFII11317 hypothetical protein 1 and hypothetical protein 2 genes complete c | KX883239.1 | Terrestrial | Arthropoda | Crustacea |
| 2463 | Beihai astro-like virus strain BHWZXX13371 putative protease RNA-dependent RNA polymerase and putative cap | KX884564.1 | South China Sea | Arthropoda | Crustacea |
| 2464 | Sanxia tombus-like virus 5 strain SXSSP12445 hypothetical protein 1 and hypothetical protein 2 genes complete cds | KX883172.1 | Freshwater | Arthropoda | Insecta |
| 2465 | Shuangao insect virus 10 strain insectZJ94627 hypothetical protein gene complete cds | KX883121.1 | Terrestrial | Arthropoda | Insecta |
| 2466 | Wuhan flea virus strain WHZM segment 3 complete sequence | KR902715.1 | Terrestrial | Arthropoda | Insecta |
| 2467 | Bivalvia Picorna-like virus H1 | GWHAPGO01000000 | Yellow China Sea | Mollusca | Bivalvia |
| 2468 | Beihai sobemo-like virus 12 strain BHWZXX15654 hypothetical protein 1 and hypothetical protein 2 genes comple | KX882794.1 | South China Sea | Arthropoda | Crustacea |
| 2469 | Beihai sobemo-like virus 12 strain BHBJDX18408 hypothetical protein genes complete cds | KX882782.1 | South China Sea | Arthropoda | Crustacea |
| 2470 | Beihai sobemo-like virus 12 strain HOU156693 hypothetical protein 1 and hypothetical protein 2 genes complete cc | KX882819.1 | South China Sea | Arthropoda | Chelicerata |
| 2471 | Hubei bunya-like virus 7 strain WGML112345 RNA-dependent RNA polymerase gene complete cds | KX884838.1 | Terrestrial | Arthropoda | Myriapoda |
| 2472 | Hubei myriapoda virus 6 strain WGML140341 RNA-dependent RNA polymerase gene complete cds | KX884843.1 | Terrestrial | Arthropoda | Myriapoda |
| 2473 | Hubei diptera virus 4 strain SCM94992 RNA-dependent RNA polymerase gene complete cds | KX884809.1 | Terrestrial | Arthropoda | Insecta |
| 2474 | Wutai Mosquito Virus strain QN3-5 RNA-dependent RNA polymerase (L) gene complete cds | KM817700.1 | Terrestrial | Arthropoda | Insecta |
| 2475 | Wutai Mosquito Virus strain QN3-5 glycoprotein precursor (G) gene complete cds | KM817728.1 | Terrestrial | Arthropoda | Insecta |

| | | | | | |
|---|---|---|---|---|---|
| 2476 | Hubei blood fluke virus 2 strain HBXXC60124 RNA-dependent RNA polymerase gene complete cds | KX884761.1 | Freshwater | Platyhelminthes | Trematoda |
| 2477 | Hubei blood fluke virus 2 strain RBXXC137009 RNA-dependent RNA polymerase gene complete cds | KX884790.1 | Freshwater | Platyhelminthes | Trematoda |
| 2478 | Hubei blood fluke virus 2 strain JSC69896 RNA-dependent RNA polymerase gene complete cds | KX884768.1 | Freshwater | Platyhelminthes | Trematoda |
| 2479 | Hubei diptera virus 4 strain SCM94992 putative glycoprotein gene complete cds | KX884810.1 | Terrestrial | Arthropoda | Insecta |
| 2480 | Wuhan Spider Virus strain SYZZ-2 RNA-dependent RNA polymerase (L) gene complete cds | KM817699.1 | Terrestrial | Arthropoda | Chelicerata |
| 2481 | Hubei diptera virus 3 strain SCM17647 RNA-dependent RNA polymerase gene complete cds | KX884796.1 | Terrestrial | Arthropoda | Insecta |
| 2482 | Hubei bunya-like virus 6 strain SZmix21338 RNA-dependent RNA polymerase gene partial cds | KX884837.1 | Freshwater | Annelida | Clitellata |
| 2483 | Beihai bunya-like virus 2 strain BHZC35426 RNA-dependent RNA polymerase gene complete cds | KX884754.1 | South China Sea | Mollusca | Bivalvia |
| 2484 | Changping earthworm virus 2 strain CPQY18140 segment 1 RNA-dependent RNA polymerase gene complete cds | KX883849.1 | Terrestrial | Annelida | Clitellata |
| 2485 | Changping earthworm virus 2 strain CPQY18140 segment 2 hypothetical protein gene complete cds | KX883850.1 | Terrestrial | Annelida | Clitellata |
| 2486 | Hubei tetragnatha maxillosa virus 6 strain arthropodmix13964 segment 1 hypothetical protein genes complete cds | KX882773.1 | Terrestrial | Arthropoda | Mix |
| 2487 | Hubei tetragnatha maxillosa virus 6 strain SSZZ3520 segment 1 hypothetical protein 1 and hypothetical protein 2 ge | KX882882.1 | Freshwater | Arthropoda | Chelicerata |
| 2488 | Hubei tetragnatha maxillosa virus 6 strain WHYY23923 segment 1 hypothetical protein 1 and hypothetical protein 2 | KX882926.1 | Terrestrial | Arthropoda | Myriapoda |
| 2489 | Shahe heteroptera virus 3 strain SHWCII5319 RNA-dependent RNA polymerase gene complete cds | KX884829.1 | Freshwater | Arthropoda | Mix |
| 2490 | Shahe heteroptera virus 3 strain SHWC01c3680 RNA-dependent RNA polymerase gene complete cds | KX884813.1 | Freshwater | Arthropoda | Insecta |
| 2491 | Shahe heteroptera virus 3 strain SHWC0209c8767 RNA-dependent RNA polymerase gene complete cds | KX884818.1 | Freshwater | Arthropoda | Crustacea |
| 2492 | Wuhan aphid virus 1 strain WHYC-1 segment 3 complete sequence | KR902723.1 | Terrestrial | Arthropoda | Insecta |
| 2493 | Wuhan Insect virus 1 strain WHYC03 RNA-dependent RNA polymerase (L) gene complete cds | KM817691.1 | Terrestrial | Arthropoda | Insecta |
| 2494 | Wuhan Millipede Virus 1 strain WHWG04 RNA-dependent RNA polymerase (L) gene complete cds | KM817695.1 | Terrestrial | Arthropoda | Myriapoda |
| 2495 | Shuangao insect virus 7 strain SKC segment 1 complete sequence | KR902717.1 | Terrestrial | Arthropoda | Insecta |
| 2496 | Cephalopoda Picorna-like virus H1 | GWHAPHA01000000 | Yellow China Sea | Mollusca | Cephalopoda |
| 2497 | Beihai picorna-like virus 76 strain HOU156700 hypothetical protein gene partial cds | KX883903.1 | South China Sea | Arthropoda | Chelicerata |
| 2498 | Hubei lepidoptera virus 4 strain LCM63323 hypothetical protein 2 gene complete cds | KX884630.1 | Terrestrial | Arthropoda | Insecta |
| 2499 | Hubei picorna-like virus 80 strain WHCCII10252 hypothetical protein gene complete cds | KX884272.1 | Terrestrial | Arthropoda | Insecta |
| 2500 | Hubei sobemo-like virus 36 strain spider124913 hypothetical protein 1 and hypothetical protein 2 genes complete c | KX882867.1 | Terrestrial | Arthropoda | Chelicerata |
| 2501 | Hubei sobemo-like virus 27 strain WHCCII13324 hypothetical protein 1 and hypothetical protein 2 genes complete | KX882911.1 | Terrestrial | Arthropoda | Insecta |
| 2502 | Wuhan aphid virus 2 strain WHYC-2 segment 1 complete sequence | KR902725.1 | Terrestrial | Arthropoda | Insecta |
| 2503 | Shuangao toti-like virus strain insectZJ97483 hypothetical protein 1 and hypothetical protein 2 genes complete cds | KX882960.1 | Terrestrial | Arthropoda | Insecta |
| 2504 | Beihai picorna-like virus 68 strain HOU158652 hypothetical protein gene complete cds | KX883899.1 | South China Sea | Arthropoda | Chelicerata |
| 2505 | Beihai sea slater virus 3 strain BHHZL10233 hypothetical protein 1 hypothetical protein 2 and hypothetical protein | KX882945.1 | South China Sea | Arthropoda | Crustacea |
| 2506 | Kadipiro virus strain QTM27331 RNA-dependent RNA polymerase gene complete cds | KX884650.1 | Terrestrial | Arthropoda | Insecta |
| 2507 | Hubei tombus-like virus 33 strain WHBL67076 hypothetical protein 1 and hypothetical protein 2 genes complete c | KX883210.1 | Freshwater | Mollusca | Bivalvia |
| 2508 | Bivalvia Picorna-like virus D5 | GWHAPEO01000000 | East China Sea | Mollusca | Bivalvia |
| 2509 | Wenling crustacean virus 14 strain WLJQ104130 RNA-dependent RNA polymerase gene complete cds | KX884451.1 | East China Sea | Arthropoda | Crustacea |
| 2510 | Wuhan Millipede virus 4 strain WHWG256022 segment Seg 2 hypothetical protein gene complete cds | KX884203.1 | Terrestrial | Arthropoda | Myriapoda |
| 2511 | Wuhan Millipede virus 4 strain GCM8225 segment Seg 2 hypothetical protein gene complete cds | KX884094.1 | Terrestrial | Arthropoda | Insecta |
| 2512 | Cactus virus X strain SCM51431 ORF1 ORF2 ORF3 and ORF4 genes complete cds | KX883791.1 | Terrestrial | Arthropoda | Insecta |

| | | | | | |
|---|---|---|---|---|---|
| 2513 | Hubei reo-like virus 5 strain WHCCII11806 RdRp and hypothetical protein genes complete cds | KX884720.1 | Terrestrial | Arthropoda | Insecta |
| 2514 | Hubei permutotetra-like virus 1 strain QTM46259 RdRp and hypothetical protein genes complete cds | KX883446.1 | Terrestrial | Arthropoda | Insecta |
| 2515 | Hubei permutotetra-like virus 1 strain WZFSL84790 RdRp and hypothetical protein genes complete cds | KX883457.1 | Freshwater | Mollusca | Gastropoda |
| 2516 | Hubei permutotetra-like virus 1 strain spider133365 RdRp and hypothetical protein genes complete cds | KX883448.1 | Terrestrial | Arthropoda | Chelicerata |
| 2517 | Hubei permutotetra-like virus 1 strain mosZJ32939 RdRp hypothetical protein 1 and hypothetical protein 2 genes cc | KX883441.1 | Terrestrial | Arthropoda | Insecta |
| 2518 | Wuhan insect virus 27 strain WHZM10130 hypothetical protein 1 and hypothetical protein 2 genes complete cds | KX882990.1 | Terrestrial | Arthropoda | Insecta |
| 2519 | Wuhan Millipede virus 4 strain GCM8225 segment Seg 3 hypothetical protein gene complete cds | KX884096.1 | Terrestrial | Arthropoda | Insecta |
| 2520 | Wuhan Millipede virus 4 strain WHWG256022 segment Seg 4 hypothetical protein gene complete cds | KX884205.1 | Terrestrial | Arthropoda | Myriapoda |
| 2521 | Beihai tombus-like virus 16 strain BHWZXX15284 hypothetical protein 1 hypothetical protein 2 and hypothetical p | KX883047.1 | South China Sea | Arthropoda | Crustacea |
| 2522 | Bivalvia Picorna-like virus N45 | GWHAPHU01000000 | South China Sea | Mollusca | Bivalvia |
| 2523 | Beihai sphaeromadae virus 4 strain BHTSS17969 RdRp and putative capsid protein genes complete cds | KX883437.1 | South China Sea | Arthropoda | Crustacea |
| 2524 | Wuhan aphid virus 1 strain WHYC-1 segment 2 complete sequence | KR902722.1 | Terrestrial | Arthropoda | Insecta |
| 2525 | Bivalvia Sobeli-like virus H3 | GWHAPHI01000000 | Yellow China Sea | Mollusca | Bivalvia |
| 2526 | Bivalvia Durna-like virus D12 | GWHAPAL01000000 | East China Sea | Mollusca | Bivalvia |
| 2527 | Wuhan Fly Virus 2 strain SYY1-3 nucleocapsid (N) ORF2 (ORF2) X protein (X) matrix protein (M) glycoprotein (( | KM817646.1 | Terrestrial | Arthropoda | Insecta |
| 2528 | Hubei rhabdo-like virus 3 strain QCM135517 hypothetical protein 1 hypothetical protein 2 hypothetical protein 3 pu | KX884417.1 | Terrestrial | Arthropoda | Insecta |
| 2529 | Drosophila immigrans Nora virus strain SCM51502 hypothetical protein 1 hypothetical protein 2 hypothetical prote | KX883975.1 | Terrestrial | Arthropoda | Insecta |
| 2530 | Beihai paphia shell virus 5 strain BWBFG36411 hypothetical protein gene partial cds | KX882951.1 | South China Sea | Mollusca | Bivalvia |
| 2531 | Wuhan pillworm virus 4 strain WHSFII2200 RdRp gene complete cds | KX884199.1 | Terrestrial | Arthropoda | Crustacea |
| 2532 | Hubei odonate virus 6 strain QTM11719 hypothetical protein 1 hypothetical protein 2 hypothetical protein 3 and hy | KX883928.1 | Terrestrial | Arthropoda | Insecta |
| 2533 | Hubei lepidoptera virus 3 strain LCM141729 Cypovirus VP8 gene complete cds | KX884620.1 | Terrestrial | Arthropoda | Insecta |
| 2534 | Wuhan Mosquito Virus 8 strain XC2-7 complete genome | KM817610.1 | Terrestrial | Arthropoda | Insecta |
| 2535 | Sanxia Water Strider Virus 5 strain SXSSP11 nucleocapsid (N) ORF2 (ORF2) ORF3 (ORF3) glycoprotein (G) and | KM817634.1 | Freshwater | Arthropoda | Insecta |
| 2536 | Beihai sesarmid crab virus 3 strain SCJXSX39048 hypothetical protein gene complete cds | KX884033.1 | South China Sea | Arthropoda | Crustacea |
| 2537 | Bivalvia Picorna-like virus D15 | GWHAOYH0100000C | East China Sea | Mollusca | Bivalvia |
| 2538 | Bivalvia Picorna-like virus H7 | GWHAPJA01000000 | Yellow China Sea | Mollusca | Bivalvia |
| 2539 | Shuangao lacewing virus 2 strain SCL polyprotein gene complete cds | KR902734.1 | Terrestrial | Arthropoda | Insecta |
| 2540 | Hubei diptera virus 4 strain SCM94992 putative nucleoprotein gene complete cds | KX884811.1 | Terrestrial | Arthropoda | Insecta |
| 2541 | Porcine epidemic diarrhea virus strain WHZHC72223 ORF1ab S protein ORF3 protein envelope protein M protein | KX883635.1 | Terrestrial | Nematoda | Chromadorea |
| 2542 | Cephalopoda Durna-like virus H1 | GWHAPGX01000000 | Yellow China Sea | Mollusca | Cephalopoda |
| 2543 | Xinzhou toro-like virus strain XZSJSC65757 1ab putative glycoprotein and hypothetical protein genes complete cds | KX883638.1 | Terrestrial | Nematoda | Mix |
| 2544 | Wuhan Insect virus 1 strain WHYC03 nucleopasid protein (N) gene partial cds | KM817752.1 | Terrestrial | Arthropoda | Insecta |
| 2545 | Bivalvia Picorna-like virus D88 | GWHAPIK01000000 | East China Sea | Mollusca | Bivalvia |
| 2546 | Xinzhou spider virus 2 strain XZZZ-7 polyprotein gene complete cds | KR902735.1 | Terrestrial | Arthropoda | Chelicerata |
| 2547 | Bivalvia Picorna-like virus D89 | GWHAPIN01000000 | East China Sea | Mollusca | Bivalvia |
| 2548 | Beihai Nido-like virus 2 strain BHXun32263 1ab putative glycoprotein 1 putative glycoprotein 2 and hypothetical p | KX883628.1 | South China Sea | Arthropoda | Crustacea |
| 2549 | Shahe yuevirus-like virus 1 strain SHWC0209c11789 hypothetical protein gene complete cds | KX884035.1 | Freshwater | Arthropoda | Crustacea |

| | | | | | |
|---|---|---|---|---|---|
| 2550 | Bivalvia Picorna-like virus N39 | GWHAPHR01000000 | South China Sea | Mollusca | Bivalvia |
| 2551 | Tacheng tick virus 8 strain TCP-2 polyprotein gene complete cds | KR902741.1 | Terrestrial | Arthropoda | Chelicerata |
| 2552 | Bivalvia Picorna-like virus N57 | GWHAPJR01000000 | South China Sea | Mollusca | Bivalvia |
| 2553 | Beihai barnacle viurs 1 strain BHGZ polyprotein gene complete cds | KR902731.1 | South China Sea | Arthropoda | Crustacea |
| 2554 | Shayang fly virus 4 strain SYCY polyprotein gene complete cds | KR902732.1 | Terrestrial | Arthropoda | Insecta |
| 2555 | Bivalvia Picorna-like virus N8 | GWHAPEY01000000 | South China Sea | Mollusca | Bivalvia |
| 2556 | Kadipiro virus strain QTM27331 Kadipiro VP12 gene complete cds | KX884661.1 | Terrestrial | Arthropoda | Insecta |
| 2557 | Bivalvia Picorna-like virus N49 | GWHAPLN01000000 | South China Sea | Mollusca | Bivalvia |
| 2558 | Hubei tetragnatha maxillosa virus 7 strain SSZZ3468 1ab putative structural polyprotein and hypothetical protein ge | KX883633.1 | Freshwater | Arthropoda | Chelicerata |
| 2559 | Jingshan Fly Virus 1 strain SCM40889 segment NP nucleocapsid protein (NP) gene complete cds | KX883878.1 | Terrestrial | Arthropoda | Insecta |
| 2560 | Jingshan Fly Virus 1 strain SYY1-7 segment NP nucleocapsid protein gene complete cds | KX883857.1 | Terrestrial | Arthropoda | Insecta |
| 2561 | Hubei lepidoptera virus 3 strain LCM141729 Cypovirus VP9 gene complete cds | KX884621.1 | Terrestrial | Arthropoda | Insecta |
| 2562 | Jiangxia Mosquito Virus 2 strain JX1-14 glycoprotein precursor (G) gene partial cds | KM817709.1 | Terrestrial | Arthropoda | Insecta |
| 2563 | Shayang spider virus 4 strain SYZZ-1 polyprotein gene complete cds | KR902739.1 | Terrestrial | Arthropoda | Chelicerata |
| 2564 | Beihai orthomyxo-like virus 1 strain BHTSS17872 segment 4 hemagglutinin gene complete cds | KX883847.1 | South China Sea | Arthropoda | Crustacea |
| 2565 | Hubei toti-like virus 4 strain WHWG8762 hypothetical protein gene partial cds | KX882981.1 | Terrestrial | Arthropoda | Myriapoda |
| 2566 | Beihai hermit crab virus 4 strain BHJJX25971 1ab putative glycoprotein 1 hypothetical protein putative glycoprotei | KX883627.1 | South China Sea | Arthropoda | Crustacea |
| 2567 | Hubei diptera virus 6 strain SCM95079 putative nucleoprotein gene complete cds | KX884806.1 | Terrestrial | Arthropoda | Insecta |
| 2568 | Wuhan Louse Fly Virus 1 strain BFJSC-10 glycoprotein precursor (G) gene partial cds | KM817725.1 | Terrestrial | Arthropoda | Insecta |
| 2569 | Xinzhou spider virus 3 strain XZZZ-3 polyprotein gene complete cds | KR902730.1 | Terrestrial | Arthropoda | Chelicerata |
| 2570 | Hubei toti-like virus 7 strain SCM411298 hypothetical protein gene partial cds | KX882971.1 | Terrestrial | Arthropoda | Insecta |
| 2571 | Bivalvia Picorna-like virus N25 | GWHAPLF01000000 | South China Sea | Mollusca | Bivalvia |
| 2572 | Beihai toti-like virus 3 strain BWBFG40422 hypothetical protein gene partial cds | KX882950.1 | South China Sea | Mollusca | Bivalvia |
| 2573 | Wuhan centipede virus strain WHWG-11 polyprotein gene complete cds | KR902737.1 | Terrestrial | Arthropoda | Myriapoda |
| 2574 | Hubei bunya-like virus 12 strain SCM26604 RNA-dependent RNA polymerase gene partial cds | KX884795.1 | Terrestrial | Arthropoda | Insecta |
| 2575 | Sanxia water strider virus 6 strain SYSZZ-2 polyprotein gene complete cds | KR902740.1 | Freshwater | Arthropoda | Insecta |
| 2576 | Wuhan heteroptera virus 3 strain WHYCM468 hypothetical protein 7 gene complete cds | KX884729.1 | Terrestrial | Arthropoda | Insecta |
| 2577 | Wuhan heteroptera virus 3 strain arthropodmix44000 hypothetical protein 8 gene complete cds | KX884596.1 | Terrestrial | Arthropoda | Mix |
| 2578 | Bivalvia Picorna-like virus D21 | GWHAOYM0100000( | East China Sea | Mollusca | Bivalvia |
| 2579 | Sanxia Water Strider Virus 1 strain SXXX37846 RNA-dependent RNA polymerase gene complete cds | KX884832.1 | Freshwater | Arthropoda | Crustacea |
| 2580 | Sanxia Water Strider Virus 1 strain SXSSP08 RNA-dependent RNA polymerase (L) gene complete cds | KM817674.1 | Freshwater | Arthropoda | Insecta |
| 2581 | Wuhan Insect virus 4 strain YCYC03 nucleocapsid (N) phosphoprotein (P) 4b protein (4b) putative matrix protein ( | KM817650.1 | Terrestrial | Arthropoda | Insecta |
| 2582 | Hubei lepidoptera virus 1 strain LCM141331 putative nucleoprotein gene complete cds | KX884774.1 | Terrestrial | Arthropoda | Insecta |
| 2583 | Gastropoda Picorna-like virus N13 | GWHAPCD01000000 | South China Sea | Mollusca | Gastropoda |
| 2584 | Bivalvia Picorna-like virus H6 | GWHAPJB01000000 | Yellow China Sea | Mollusca | Bivalvia |
| 2585 | Shayang Fly Virus 1 strain SYY3-1 glycoprotein (G) putative nucleoprotein (N) and polymerase (L) genes complete | KM817598.1 | Terrestrial | Arthropoda | Insecta |
| 2586 | Wuchang Cockraoch Virus 1 strain WCZL-5 nucleopasid protein (N) gene complete cds | KM817748.1 | Terrestrial | Arthropoda | Insecta |

| | | | | | |
|---|---|---|---|---|---|
| 2587 | Shahe endorna-like virus 1 strain SHWC0209c12790 hypothetical protein gene complete cds | KX883795.1 | Freshwater | Arthropoda | Crustacea |
| 2588 | Drosophila immigrans sigmavirus strain SCM45623 nucleocapsid protein polymerase-associated protein PP3 matrix | KX884434.1 | Terrestrial | Arthropoda | Insecta |
| 2589 | Hubei earwig virus 1 strain GCM19162 segment 5 hemagglutinin (HA) gene complete cds | KX883862.1 | Terrestrial | Arthropoda | Insecta |
| 2590 | Bivalvia Picorna-like virus N5 | GWHAPEZ01000000 | South China Sea | Mollusca | Bivalvia |
| 2591 | Beihai rhabdo-like virus 2 strain BHTSS7258 hypothetical protein 1 hypothetical protein 2 putative glycoprotein an | KX884413.1 | South China Sea | Arthropoda | Crustacea |
| 2592 | Bivalvia Picorna-like virus H9 | GWHAPJF01000000 | Yellow China Sea | Mollusca | Bivalvia |
| 2593 | Beihai hepe-like virus 4 strain YYSZX17852 hypothetical protein genes complete cds | KX883839.1 | South China Sea | Arthropoda | Crustacea |
| 2594 | Sanxia Water Strider Virus 2 strain SXSSP02 nucleopasid protein (N) gene complete cds | KX650647.1 | Freshwater | Arthropoda | Insecta |
| 2595 | Sanxia atyid shrimp virus 4 strain SXXX37205 RNA-dependent RNA polymerase putative glycoprotein and hypoth | KX884439.1 | Freshwater | Arthropoda | Crustacea |
| 2596 | Hubei diptera virus 3 strain SCM17647 putative nucleoprotein gene complete cds | KX884798.1 | Terrestrial | Arthropoda | Insecta |
| 2597 | Bivalvia Picorna-like virus D68 | GWHAPKQ01000000 | East China Sea | Mollusca | Bivalvia |
| 2598 | Crustacea Picorna-like virus D4 | GWHAPBT01000000 | East China Sea | Arthropoda | Crustacea |
| 2599 | Beihai sipunculid worm virus 5 strain BHNXC41492 hypothetical protein 1 hypothetical protein 2 hypothetical prot | KX883408.1 | South China Sea | Sipuncula | Sipuncula |
| 2600 | Bivalvia Picorna-like virus D29 | GWHAOYT01000000 | East China Sea | Mollusca | Bivalvia |
| 2601 | Hubei orthoptera virus 6 strain ZCM21154 segment 5 nucleocapsid protein gene complete cds | KX883888.1 | Terrestrial | Arthropoda | Insecta |
| 2602 | Bivalvia Picorna-like virus D63 | GWHAPAM0100000 | East China Sea | Mollusca | Bivalvia |
| 2603 | Gastropoda Picorna-like virus N3 | GWHAPFE01000000 | South China Sea | Mollusca | Gastropoda |
| 2604 | Sanxia atyid shrimp virus 1 strain SXXX35475 RdRp and hypothetical protein genes complete cds | KX883804.1 | Freshwater | Arthropoda | Crustacea |
| 2605 | Hubei partiti-like virus 39 strain QTM24152 RdRp gene complete cds | KX884120.1 | Terrestrial | Arthropoda | Insecta |
| 2606 | Hubei virga-like viurs 8 strain WGML111093 RdRp and hypothetical protein genes complete cds and hypothetical | KX883807.1 | Terrestrial | Arthropoda | Myriapoda |
| 2607 | Gastropoda Unclassified virus D6 | GWHAPBM0100000 | East China Sea | Mollusca | Gastropoda |
| 2608 | Xinzhou dimarhabdovirus virus 1 strain XZSJSC65538 hypothetical protein putative glycoprotein and RNA-depend | KX884460.1 | Terrestrial | Nematoda | Mix |
| 2609 | Beihai narna-like virus 16 strain BHJP26864 RNA-dependent RNA polymerase gene complete cds | KX883472.1 | South China Sea | Echinodermata | Echinozoa |
| 2610 | Hubei lepidoptera virus 4 strain LCM63323 hypothetical protein 4 gene complete cds | KX884632.1 | Terrestrial | Arthropoda | Insecta |
| 2611 | Beihai picorna-like virus 74 strain SCJXSX39031 hypothetical protein 1 and hypothetical protein 2 genes complete | KX883961.1 | South China Sea | Arthropoda | Crustacea |
| 2612 | Beihai picorna-like virus 74 strain BHHK130969 hypothetical protein 1 and hypothetical protein 2 genes complete | KX883349.1 | South China Sea | Cnidaria | Anthozoa |
| 2613 | Hubei picorna-like virus 58 strain SSZZ391 hypothetical protein gene complete cds | KX883701.1 | Freshwater | Arthropoda | Chelicerata |
| 2614 | Hubei picobirna-like virus 2 strain WHBL68697 RdRp gene complete cds | KX884190.1 | Freshwater | Mollusca | Bivalvia |
| 2615 | Hubei partiti-like virus 28 strain spider113121 RdRp gene complete cds | KX884158.1 | Terrestrial | Arthropoda | Chelicerata |
| 2616 | Bivalvia Durna-like virus N3 | GWHAPLE01000000 | South China Sea | Mollusca | Bivalvia |
| 2617 | Crustacea Picorna-like virus N3 | GWHAPFX01000000 | South China Sea | Arthropoda | Crustacea |
| 2618 | Sanxia picorna-like virus 5 strain SXXX34146 hypothetical protein and hypothetical protein 2 genes complete cds | KX883721.1 | Freshwater | Arthropoda | Crustacea |
| 2619 | Wenling picorna-like virus 1 strain WLJQ101464 hypothetical protein 1 and hypothetical protein 2 genes complete | KX884311.1 | East China Sea | Arthropoda | Crustacea |
| 2620 | Wenling partiti-like virus 2 strain WLJQ102434 RdRp gene complete cds | KX884227.1 | East China Sea | Arthropoda | Crustacea |
| 2621 | Hubei noda-like virus 9 strain WHLC5367 capsid protein gene complete cds | KX883236.1 | Terrestrial | Arthropoda | Insecta |
| 2622 | Hubei noda-like virus 9 strain QTM26368 capsid protein gene complete cds | KX883131.1 | Terrestrial | Arthropoda | Insecta |
| 2623 | Wenzhou hepe-like virus 2 strain WZFSL85851 replicase and putative structural protein genes complete cds | KX883832.1 | Freshwater | Mollusca | Gastropoda |

| | | | | | |
|---|---|---|---|---|---|
| ***2624*** | Beihai picobirna-like virus 7 strain BHBJDX17263 RdRp gene complete cds | KX884053.1 | South China Sea | Arthropoda | Crustacea |
| ***2625*** | Beihai picobirna-like virus 7 strain BHJJX15399 RdRp gene complete cds | KX884056.1 | South China Sea | Arthropoda | Crustacea |
| ***2626*** | Beihai picobirna-like virus 7 strain BHNXC57916 segment Seg 1 RdRp gene complete cds | KX884062.1 | South China Sea | Sipuncula | Sipuncula |
| ***2627*** | Beihai picobirna-like virus 7 strain BHZC50960 RdRp gene complete cds | KX884080.1 | South China Sea | Mollusca | Bivalvia |
| ***2628*** | Beihai picobirna-like virus 7 strain BHWZXX14123 RdRp gene complete cds | KX884073.1 | South China Sea | Arthropoda | Crustacea |
| ***2629*** | Wenzhou picorna-like virus 42 strain WZFSL74289 hypothetical protein 1 and hypothetical protein 2 genes comple | KX884339.1 | Freshwater | Mollusca | Gastropoda |
| ***2630*** | Hubei partiti-like virus 18 strain SCM50556 RdRp gene complete cds | KX884152.1 | Terrestrial | Arthropoda | Insecta |
| ***2631*** | Beihai picorna-like virus 25 strain BHJJX20433 hypothetical protein 1 gene complete cds and hypothetical protein [illegible] | KX883361.1 | South China Sea | Arthropoda | Crustacea |
| ***2632*** | Wenling partiti-like virus 12 strain WLJQ98567 RdRp gene complete cds | KX884226.1 | East China Sea | Arthropoda | Crustacea |
| ***2633*** | Bivalvia Picorna-like virus D79 | GWHAPET01000000 | East China Sea | Mollusca | Bivalvia |
| ***2634*** | Hubei arthropod virus 3 strain WGML144461 hypothetical protein gene complete cds | KX883738.1 | Terrestrial | Arthropoda | Myriapoda |
| ***2635*** | Hubei arthropod virus 3 strain GCM7473 hypothetical protein gene complete cds | KX883895.1 | Terrestrial | Arthropoda | Insecta |
| ***2636*** | Hubei arthropod virus 3 strain ZCM127409 hypothetical protein gene complete cds | KX884393.1 | Terrestrial | Arthropoda | Insecta |
| ***2637*** | Bivalvia Picorna-like virus D90 | GWHAPIO01000000 | East China Sea | Mollusca | Bivalvia |
| ***2638*** | Hubei picorna-like virus 5 strain WHBL70549 hypothetical protein 1 and hypothetical protein 2 genes complete cds | KX884261.1 | Freshwater | Mollusca | Bivalvia |
| ***2639*** | Crustacea Durna-like virus N1 | GWHAPFI01000000 | South China Sea | Arthropoda | Crustacea |
| ***2640*** | Hubei picorna-like virus 59 strain spider133744 hypothetical protein gene complete cds | KX883682.1 | Terrestrial | Arthropoda | Chelicerata |
| ***2641*** | Bivalvia Picorna-like virus N41 | GWHAPHT01000000 | South China Sea | Mollusca | Bivalvia |
| ***2642*** | Bivalvia Picorna-like virus N44 | GWHAPHQ01000000 | South China Sea | Mollusca | Bivalvia |
| ***2643*** | Xinzhou nematode virus 6 strain XZSJSC61041 1ab gene partial cds and S protein hypothetical protein M protein h | KX883637.1 | Terrestrial | Nematoda | Mix |
| ***2644*** | Hubei partiti-like virus 27 strain QTM26091 RdRp gene partial cds | KX884128.1 | Terrestrial | Arthropoda | Insecta |
| ***2645*** | Gastropoda Durna-like virus D1 | GWHAPKJ01000000 | East China Sea | Mollusca | Gastropoda |
| ***2646*** | Wuhan insect virus 11 strain CJLX30623 hypothetical protein 1 and hypothetical protein 2 genes complete cds | KX884551.1 | Freshwater | Arthropoda | Crustacea |
| ***2647*** | Wuhan insect virus 11 strain ZCM21223 hypothetical protein 1 and hypothetical protein 2 genes complete cds | KX884398.1 | Terrestrial | Arthropoda | Insecta |
| ***2648*** | Wuhan insect virus 11 strain arthropodmix14080 hypothetical protein 1 and hypothetical protein 2 genes complete c | KX883293.1 | Terrestrial | Arthropoda | Mix |
| ***2649*** | Wuhan insect virus 11 strain WHZM10185 hypothetical protein 1 and hypothetical protein 2 genes complete cds | KX884307.1 | Terrestrial | Arthropoda | Insecta |
| ***2650*** | Bivalvia Durna-like virus H8 | GWHAPCQ01000000 | Yellow China Sea | Mollusca | Bivalvia |
| ***2651*** | Hubei lepidoptera virus 3 strain LCM141729 Cypovirus VP7 gene complete cds | KX884619.1 | Terrestrial | Arthropoda | Insecta |
| ***2652*** | Behai endorna-like virus 1 strain BHTSS17827 hypothetical protein gene complete cds | KX883750.1 | South China Sea | Arthropoda | Crustacea |
| ***2653*** | Hubei endorna-like virus 1 strain QCM148882 hypothetical protein gene complete cds | KX883776.1 | Terrestrial | Arthropoda | Insecta |
| ***2654*** | Wuhan millipede virus 3 strain WHWG56524 hypothetical protein 1 and hypothetical protein 2 genes complete cds | KX884293.1 | Terrestrial | Arthropoda | Myriapoda |
| ***2655*** | Hubei lepidoptera virus 3 strain LCM141729 Cypovirus VP6 gene complete cds | KX884618.1 | Terrestrial | Arthropoda | Insecta |
| ***2656*** | Hubei dimarhabdovirus virus 2 strain QTM26629 putative nucleoprotein hypothetical protein 2 hypothetical protein | KX884426.1 | Terrestrial | Arthropoda | Insecta |
| ***2657*** | Hubei toti-like virus 11 strain QTM2810 hypothetical protein gene partial cds | KX882968.1 | Terrestrial | Arthropoda | Insecta |
| ***2658*** | Hubei odonate virus 8 strain QTM19232 putative glycoprotein gene complete cds | KX884776.1 | Terrestrial | Arthropoda | Insecta |
| ***2659*** | Beihai rhabdo-like virus 1 strain BHTSS15727 hypothetical protein 1 hypothetical protein 2 and RNA-dependent R | KX884412.1 | South China Sea | Arthropoda | Crustacea |
| ***2660*** | Hubei diptera virus 5 strain SCM245062 putative glycoprotein gene complete cds | KX884800.1 | Terrestrial | Arthropoda | Insecta |

| | | | | | |
|---|---|---|---|---|---|
| 2661 | Bivalvia Picorna-like virus H17 | GWHAPJI01000000 | Yellow China Sea | Mollusca | Bivalvia |
| 2662 | Wuhan house centipede virus 3 strain WHYY18014 hypothetical protein gene complete cds | KX884300.1 | Terrestrial | Arthropoda | Myriapoda |
| 2663 | Wuhan house centipede virus 3 strain arthropodmix13505 hypothetical protein gene complete cds | KX883287.1 | Terrestrial | Arthropoda | Mix |
| 2664 | Beihai picobirna-like virus 3 strain BHTH13093 RdRp gene complete cds | KX884066.1 | South China Sea | Arthropoda | Crustacea |
| 2665 | Hubei picorna-like virus 55 strain spider134013 hypothetical protein 1 and hypothetical protein 2 genes complete cc | KX883691.1 | Terrestrial | Arthropoda | Chelicerata |
| 2666 | Hubei picorna-like virus 39 strain SCM51450 hypothetical protein gene complete cds | KX883974.1 | Terrestrial | Arthropoda | Insecta |
| 2667 | Hubei insect virus 2 strain arthropodmix13736 hypothetical protein 5 and hypothetical protein 6 genes complete cds | KX884585.1 | Terrestrial | Arthropoda | Mix |
| 2668 | Hubei blood fluke virus 3 strain RBXXC337523 hypothetical protein 4 gene complete cds | KX884691.1 | Freshwater | Platyhelminthes | Trematoda |
| 2669 | Hubei blood fluke virus 3 strain JSC69362 hypothetical protein 4 gene complete cds | KX884612.1 | Freshwater | Platyhelminthes | Trematoda |
| 2670 | Hexanauplia Picorna-like virus D3 | GWHAPKD01000000 | East China Sea | Arthropoda | Hexanauplia |
| 2671 | Bivalvia Picorna-like virus H4 | GWHAPCO01000000 | Yellow China Sea | Mollusca | Bivalvia |
