## Supplementary material for "Viromes in Marine Ecosystems Reveal Remarkable Invertebrate RNA Virus Diversity": The information of primers used for PCR confirmation

**Table S4 The information of primers used for PCR confirmation**

| *Virus Name* | *Primers Name* | *Primers Sequence* |
|---|---|---|
| Crustacea Hepeli-like virus D1 | 1_TRINITY_DN13810_c0_g1_i1_725F | CGTGGTTGACCATGAAAACTGACG |
| Crustacea Hepeli-like virus D1 | 1_TRINITY_DN13810_c0_g1_i1_1166R | AAAAGGGTATCCGCTCTACCAGAC |
| Crustacea Hepeli-like virus D2 | 1_TRINITY_DN8232_c0_g1_i1_2004F | AGGTTCTGAGCCTTGAAGAACCAC |
| Crustacea Hepeli-like virus D2 | 1_TRINITY_DN8232_c0_g1_i1_2427R | ACCTGTTTCTCCGCTTCTATGGTG |
| Crustacea Hepeli-like virus D3 | 1_TRINITY_DN83828_c0_g1_i1_376F | GTTTCGGGCGACTTCTTCAACCTT |
| Crustacea Hepeli-like virus D3 | 1_TRINITY_DN83828_c0_g1_i1_994R | GGTAACGCCCATTGGATGCAAACA |
| Bivalvia Picorna-like virus D2 | 11_TRINITY_DN101233_c0_g1_i3_298F | GCGAGATCCGGAAAGAATCCGAAA |
| Bivalvia Picorna-like virus D2 | 11_TRINITY_DN101233_c0_g1_i3_745R | CCCTGCATGTAGCACGACAGATAA |
| Bivalvia Picorna-like virus D1 | 11_TRINITY_DN2073_c0_g1_i2_651F | GAGAGCTATGTTGACATCCACAGC |
| Bivalvia Picorna-like virus D1 | 11_TRINITY_DN2073_c0_g1_i2_1194R | CCTAAGCTCTTTTGTGTGCACACG |
| Bivalvia Picorna-like virus D4 | 12_TRINITY_DN47756_c0_g1_i2_353F | GTCACTAGAATGGCAACAATGGCG |
| Bivalvia Picorna-like virus D4 | 12_TRINITY_DN47756_c0_g1_i2_792R | CGCACAAATGGATCTCCAGTAGTC |
| Bivalvia Picorna-like virus D5 | 12_TRINITY_DN80727_c0_g1_i4_432F | ATTGAACACTCTACGAGCAGGTGG |
| Bivalvia Picorna-like virus D5 | 12_TRINITY_DN80727_c0_g1_i4_862R | TTCAGCAAGGAATGTTGTGCGTGG |
| Bivalvia Durna-like virus D1 | 12_TRINITY_DN962_c0_g1_i2_360F | AGCTCTGTACGTAGGATAGATGCC |
| Bivalvia Durna-like virus D1 | 12_TRINITY_DN962_c0_g1_i2_866R | CTCATCTACCATGCTCGTAGTCAG |
| Bivalvia Picorna-like virus D13 | 13_TRINITY_DN106217_c0_g2_i1_94F | CAAGGTCGTTTGATGGTGACTTGG |
| Bivalvia Picorna-like virus D13 | 13_TRINITY_DN106217_c0_g2_i1_783R | CTCGATCCCAGCATCAGTTCTACG |
| Bivalvia Picorna-like virus D7 | 13_TRINITY_DN10961_c0_g1_i4_1359F | GGGTGAGTGAAAACCAAATCTGGC |
| Bivalvia Picorna-like virus D7 | 13_TRINITY_DN10961_c0_g1_i4_1829R | GTTCCAAAGGTTCGAGGAGCTTTC |
| Bivalvia Picorna-like virus D9 | 13_TRINITY_DN1723_c0_g1_i6_455F | CCTCGCATCGAATATCGATACGTC |
| Bivalvia Picorna-like virus D9 | 13_TRINITY_DN1723_c0_g1_i6_912R | ATTAGTTGGAGGACAGGGAACTGG |
| Bivalvia Unclassified virus D1 | 13_TRINITY_DN19948_c0_g1_i16_2103F | CTTGTTCAGTAGCAGGAGCAACAG |
| Bivalvia Unclassified virus D1 | 13_TRINITY_DN19948_c0_g1_i16_2613R | ACTGCCCAGTTTCCGCTGAAAATG |
| Bivalvia Picorna-like virus D10 | 13_TRINITY_DN34518_c0_g1_i2_423F | GATCCCTAGCGGGTTAAAGAGATG |
| Bivalvia Picorna-like virus D10 | 13_TRINITY_DN34518_c0_g1_i2_1098R | CCTTTTGGCACGAACAGAGATTCC |
| Bivalvia Picorna-like virus D11 | 13_TRINITY_DN43256_c0_g2_i1_280F | GGTGATAACACTTTCAAGGCGTGG |
| Bivalvia Picorna-like virus D11 | 13_TRINITY_DN43256_c0_g2_i1_834R | CTCAAGCAAAGAGTCGTACCATGC |
| Bivalvia Picorna-like virus D6 | 13_TRINITY_DN7096_c0_g1_i1_367F | GTCTCAGGAGTAGTTGCCGAGTTT |
| Bivalvia Picorna-like virus D6 | 13_TRINITY_DN7096_c0_g1_i1_1049R | GATAGGGTCCATTCAGCCTAGAGT |
| Bivalvia Picorna-like virus D12 | 13_TRINITY_DN7205_c0_g1_i6_250F | CTAAATCTCCTACACCTCGAGCCA |
| Bivalvia Picorna-like virus D12 | 13_TRINITY_DN7205_c0_g1_i6_767R | GGTGACGCAGTTTGTTGATGATGC |
| Bivalvia Durna-like virus D3 | 14_TRINITY_DN1066_c0_g1_i4_1221F | CAAAGTAACCCCGATCCCATTTGC |
| Bivalvia Durna-like virus D3 | 14_TRINITY_DN1066_c0_g1_i4_1672R | CGGATGTCTTAAGCGAAGTAGTCG |
| Bivalvia Picorna-like virus D14 | 14_TRINITY_DN127071_c0_g1_i1_258F | CCGTGCTTGTAGAGGTTCAGTTAG |
| Bivalvia Picorna-like virus D14 | 14_TRINITY_DN127071_c0_g1_i1_711R | CAGCGTGGGGACATTGACGAAAAA |
| Bivalvia Durna-like virus D2 | 14_TRINITY_DN2705_c0_g1_i3_269F | TAGTCCTGAAGAGATACTCCGTGG |
| Bivalvia Durna-like virus D2 | 14_TRINITY_DN2705_c0_g1_i3_708R | TTTCATCGAGGACAGTTGCTAGCG |
| Bivalvia Picorna-like virus D15 | 14_TRINITY_DN51343_c0_g2_i1_370F | CTGATTCGGATGCTATGCAGTTCG |
| Bivalvia Picorna-like virus D15 | 14_TRINITY_DN51343_c0_g2_i1_758R | GAGACGCTCATCACTGCAATCGAA |
| Bivalvia Picorna-like virus D16 | 16_TRINITY_DN68194_c0_g1_i1_254F | CTTTGCCCAATGTGACGCACTCAA |

| | | |
|---|---|---|
| Bivalvia Picorna-like virus D16 | 16_TRINITY_DN68194_c0_g1_i1_731R | GCGGACACGTTGATCGTACTTCTT |
| Gastropoda Picorna-like virus D4 | 17_TRINITY_DN105295_c0_g1_i1_204F | GGGGCTTTTGCAGCTTACTATCGT |
| Gastropoda Picorna-like virus D4 | 17_TRINITY_DN105295_c0_g1_i1_630R | CTATACTGGGGGCATGTCCAAAAG |
| Gastropoda Unclassified virus D1 | 17_TRINITY_DN106166_c0_g1_i2_258F | CTTTGCGCTGACACCCTGGATATA |
| Gastropoda Unclassified virus D1 | 17_TRINITY_DN106166_c0_g1_i2_653R | GAATCCACTGGCGAAATCTCTTCC |
| Gastropoda Picorna-like virus D3 | 17_TRINITY_DN1818_c0_g1_i6_604F | TGACCTTTTTCTTCACAGGCAGCC |
| Gastropoda Picorna-like virus D3 | 17_TRINITY_DN1818_c0_g1_i6_892R | GTCCTGCTAGAATAGGTAAGTCCG |
| Gastropoda Sobeli-like virus D1 | 17_TRINITY_DN24209_c0_g1_i2_1174F | GGAGGATATCGCATAACATCAGCG |
| Gastropoda Sobeli-like virus D1 | 17_TRINITY_DN24209_c0_g1_i2_1626R | TCGAGACACTGTTGAAGGAGTCTC |
| Gastropoda Picorna-like virus D2 | 17_TRINITY_DN24637_c0_g1_i1_49F | GCCTACGATCAGGTAAACAATCGC |
| Gastropoda Picorna-like virus D2 | 17_TRINITY_DN24637_c0_g1_i1_473R | CCGTTGGTTACTGTGTAGATAGCC |
| Gastropoda Unclassified virus D2 | 17_TRINITY_DN87839_c0_g1_i2_278F | GGATGGATTGCTAGCGCAGAAAAG |
| Gastropoda Unclassified virus D2 | 17_TRINITY_DN87839_c0_g1_i2_734R | GCAGTACTGGAGAGTCTGGAATTC |
| Bivalvia Picorna-like virus D17 | 18_TRINITY_DN46869_c0_g1_i1_335F | CCAACGTGGGCAAGTTCAGTACTT |
| Bivalvia Picorna-like virus D17 | 18_TRINITY_DN46869_c0_g1_i1_906R | CATTTTCTGGTTGGCGCGGATCAA |
| Bivalvia Picorna-like virus D18 | 18_TRINITY_DN95612_c0_g1_i4_171F | GATAATAACCGTTTAGCTCCGCCG |
| Bivalvia Picorna-like virus D18 | 18_TRINITY_DN95612_c0_g1_i4_604R | AGTAGAAGCTCCATCGGTATGACC |
| Bivalvia Picorna-like virus D19 | 19_TRINITY_DN10_c0_g1_i3_511F | GACGAGAGTGATGAGCAAATTCCC |
| Bivalvia Picorna-like virus D19 | 19_TRINITY_DN10_c0_g1_i3_1070R | GCTCGTGGTTTCAATCCACTAGAC |
| Bivalvia Picorna-like virus D35 | 19_TRINITY_DN125270_c0_g1_i2_278F | CTCCCATGAACAAGGACCGAGAAA |
| Bivalvia Picorna-like virus D35 | 19_TRINITY_DN125270_c0_g1_i2_717R | CCCTTCACCTTTGAGGGTATCGAT |
| Bivalvia Picorna-like virus D33 | 19_TRINITY_DN138578_c0_g1_i2_145F | TGGCAGAACAACTGTGTATCGGAG |
| Bivalvia Picorna-like virus D33 | 19_TRINITY_DN138578_c0_g1_i2_716R | AAGAACAGGTTTCTCAAGAGCTCC |
| Bivalvia Durna-like virus D6 | 19_TRINITY_DN1456_c0_g1_i5_727F | GTGATGTTCCCCAAACATCCCTAG |
| Bivalvia Durna-like virus D6 | 19_TRINITY_DN1456_c0_g1_i5_1179R | CACAGGCATCTAGGATTGGTGATG |
| Bivalvia Picorna-like virus D28 | 19_TRINITY_DN1578_c0_g1_i14_2317F | CCGATGCAAGACAGATGGATACTC |
| Bivalvia Picorna-like virus D28 | 19_TRINITY_DN1578_c0_g1_i14_2740R | CTGTTTCCGCGTGAGAAGAAGAAG |
| Bivalvia Durna-like virus D5 | 19_TRINITY_DN182296_c0_g1_i1_248F | TTCGAACATTTTGATCATACGCGG |
| Bivalvia Durna-like virus D5 | 19_TRINITY_DN182296_c0_g1_i1_722R | AATGCCGCAAACCGAGTTTTAAAC |
| Bivalvia Picorna-like virus D21 | 19_TRINITY_DN197365_c0_g1_i1_186F | TGTACGATAGGTTGGTTTAATCGC |
| Bivalvia Picorna-like virus D21 | 19_TRINITY_DN197365_c0_g1_i1_541R | AAGCTGTTATGGAAACATTGGGAC |
| Bivalvia Picorna-like virus D22 | 19_TRINITY_DN24867_c0_g1_i2_2487F | AAACGCACCTTGCTTGACGACTTC |
| Bivalvia Picorna-like virus D22 | 19_TRINITY_DN24867_c0_g1_i2_2929R | ACAATCTCAAGGCGTTCCGCTATC |
| Bivalvia Picorna-like virus D20 | 19_TRINITY_DN3173_c0_g1_i4_2689F | TCAATCAACTCCTCACATGCTGCG |
| Bivalvia Picorna-like virus D20 | 19_TRINITY_DN3173_c0_g1_i4_3120R | GTCGGTTTTGTCCATTATCGCTGG |
| Bivalvia Picorna-like virus D30 | 19_TRINITY_DN39273_c0_g1_i6_384F | GAGCTTCGACATATGATGTGCCAG |
| Bivalvia Picorna-like virus D30 | 19_TRINITY_DN39273_c0_g1_i6_902R | CCGATCTATTGAACTCAGCGATGG |
| Bivalvia Durna-like virus D4 | 19_TRINITY_DN5_c0_g1_i9_1009F | TTGACTCCTCCATTCTGTCCAAGG |
| Bivalvia Durna-like virus D4 | 19_TRINITY_DN5_c0_g1_i9_1465R | CTGCTGCACAAGTTTTAGAGTGGC |
| Bivalvia Picorna-like virus D29 | 19_TRINITY_DN58808_c0_g1_i1_225F | TCAATATCGCAATGGAAGCACCGG |
| Bivalvia Picorna-like virus D29 | 19_TRINITY_DN58808_c0_g1_i1_649R | GTCCAAGGTTCTTGCTTTTGGAGC |
| Bivalvia Picorna-like virus D23 | 19_TRINITY_DN64758_c0_g1_i1_1263F | GCTCTCCACCTTGATCCACTTTAG |
| Bivalvia Picorna-like virus D23 | 19_TRINITY_DN64758_c0_g1_i1_1696R | TGAGGACCGAACTTTCTGGTTCTG |

| | | |
|---|---|---|
| Bivalvia Picorna-like virus D27 | 19_TRINITY_DN67445_c0_g1_i2_335F | CCTGTCTATCCAAGATGGGAATCC |
| Bivalvia Picorna-like virus D27 | 19_TRINITY_DN67445_c0_g1_i2_967R | GCCAGAATATTCGCTGCATCGAAC |
| Bivalvia Picorna-like virus D24 | 19_TRINITY_DN67864_c0_g1_i1_298F | CGACAACTCCATCCCTTTGGAGTT |
| Bivalvia Picorna-like virus D24 | 19_TRINITY_DN67864_c0_g1_i1_896R | CATTTCGCTGGAACGCCAAACTCT |
| Bivalvia Picorna-like virus D34 | 19_TRINITY_DN68127_c0_g1_i2_268F | CAAATAGTGGAACCAGTGCAGCCA |
| Bivalvia Picorna-like virus D34 | 19_TRINITY_DN68127_c0_g1_i2_673R | CCGCTGCAAGCAAGTTATAGATCC |
| Bivalvia Picorna-like virus D40 | 19_TRINITY_DN71737_c0_g1_i2_294F | GGAGGTTTTCCCTTCTTGGATTCG |
| Bivalvia Picorna-like virus D40 | 19_TRINITY_DN71737_c0_g1_i2_717R | TACTCCTTGTGAGTGGTGAAGAGG |
| Bivalvia Picorna-like virus D38 | 19_TRINITY_DN72857_c0_g1_i1_243F | TGGATTTGGCTATGAGTACGAGCG |
| Bivalvia Picorna-like virus D38 | 19_TRINITY_DN72857_c0_g1_i1_670R | CGAATGAGACTGTACCCTCTCAAC |
| Bivalvia Picorna-like virus D26 | 19_TRINITY_DN76323_c0_g1_i1_204F | TCTTCTTTACAAGGTGATGGCCCG |
| Bivalvia Picorna-like virus D26 | 19_TRINITY_DN76323_c0_g1_i1_628R | GAACTTCCATGCTTTGCGGAAGAG |
| Bivalvia Picorna-like virus D25 | 19_TRINITY_DN8971_c0_g1_i2_794F | ACCGGATGGATGAGTTTAAGGTCC |
| Bivalvia Picorna-like virus D25 | 19_TRINITY_DN8971_c0_g1_i2_1217R | TATTTCCCAGCTACGAGCAGGTTG |
| Bivalvia Toli-like virus D1 | 20_TRINITY_DN10818_c0_g1_i3_403F | ACCGGTGCATTTGGATACATGAGG |
| Bivalvia Toli-like virus D1 | 20_TRINITY_DN10818_c0_g1_i3_843R | TTGGAAGTACACCGCTGTTGGTAC |
| Bivalvia Durna-like virus D9 | 20_TRINITY_DN119_c0_g1_i1_1180F | GGTTTCGCATCAACCTCTAATCCG |
| Bivalvia Durna-like virus D9 | 20_TRINITY_DN119_c0_g1_i1_1685R | TCAATAGGGTAGAAGCCAGGTTGC |
| Bivalvia Picorna-like virus D45 | 20_TRINITY_DN147687_c0_g1_i3_290F | CCTGCTAAAGTCAAAGATGAGGGC |
| Bivalvia Picorna-like virus D45 | 20_TRINITY_DN147687_c0_g1_i3_846R | GAATGAGGATTCATTCCCACTGCG |
| Bivalvia Picorna-like virus D55 | 20_TRINITY_DN162235_c0_g1_i1_90F | TAATTGCACCCAAACCGATTGTGG |
| Bivalvia Picorna-like virus D55 | 20_TRINITY_DN162235_c0_g1_i1_589R | TTTGAGGGCCGTTATTCAACTCTC |
| Bivalvia Picorna-like virus D42 | 20_TRINITY_DN1696_c0_g1_i7_2834F | GTTGCCTTTGTCTCCCACAAATCG |
| Bivalvia Picorna-like virus D42 | 20_TRINITY_DN1696_c0_g1_i7_3301R | TGGGGCGTTGATTTGACAGTGATG |
| Bivalvia Picorna-like virus D41 | 20_TRINITY_DN28415_c0_g1_i1_163F | TCTGGAAATGCAAAGCGAGCAGTG |
| Bivalvia Picorna-like virus D41 | 20_TRINITY_DN28415_c0_g1_i1_760R | TTCAGATGTGAATGCAGCGGAAGG |
| Bivalvia Durna-like virus D10 | 20_TRINITY_DN3714_c0_g1_i2_130F | CGACGCGATTTGTTTCGAATGTCG |
| Bivalvia Durna-like virus D10 | 20_TRINITY_DN3714_c0_g1_i2_696R | ATCGCTTAAGGGCACAGAATGCAC |
| Bivalvia Picorna-like virus D44 | 20_TRINITY_DN4026_c0_g1_i1_743F | GTTCTTCATAGGCGTTCAACCCAG |
| Bivalvia Picorna-like virus D44 | 20_TRINITY_DN4026_c0_g1_i1_1195R | CGATGATGCGTTTGTCCAATGGAG |
| Bivalvia Picorna-like virus D50 | 20_TRINITY_DN49_c0_g2_i1_728F | TCCACAAACTGTACTGGCCATAGC |
| Bivalvia Picorna-like virus D50 | 20_TRINITY_DN49_c0_g2_i1_1158R | TTGATCCACATTCTGATGGTGGCG |
| Bivalvia Picorna-like virus D53 | 20_TRINITY_DN493_c0_g1_i5_662F | TTGGCTTGGGTACGCAGAATAGTG |
| Bivalvia Picorna-like virus D53 | 20_TRINITY_DN493_c0_g1_i5_1127R | CGCGCAATTATCAGCAGTACAAGG |
| Bivalvia Durna-like virus D7 | 20_TRINITY_DN67_c0_g1_i4_414F | AGCTCTGTACGTAGGATAGATGCC |
| Bivalvia Durna-like virus D7 | 20_TRINITY_DN67_c0_g1_i4_920R | CTCATCTACCATGCTCGTAGTCAG |
| Bivalvia Picorna-like virus D52 | 20_TRINITY_DN67819_c0_g1_i1_217F | CAACTCTCGGAGATCAATTGCTGG |
| Bivalvia Picorna-like virus D52 | 20_TRINITY_DN67819_c0_g1_i1_644R | CGAATCAATACCGCAGTATACCCC |
| Bivalvia Picorna-like virus D49 | 20_TRINITY_DN75302_c0_g2_i2_615F | GACGACTTGATCATGCGTCATGGT |
| Bivalvia Picorna-like virus D49 | 20_TRINITY_DN75302_c0_g2_i2_1071R | GGGTTTTCTGGCATGTGAAAGTGC |
| Bivalvia Picorna-like virus D48 | 20_TRINITY_DN75695_c0_g1_i1_150F | TCGACTCCGCGACACAAATGTATG |
| Bivalvia Picorna-like virus D48 | 20_TRINITY_DN75695_c0_g1_i1_641R | TGTACAGAGGAAAATGAGCGCGTC |
| Bivalvia Picorna-like virus D51 | 20_TRINITY_DN76783_c0_g1_i3_193F | AGTCTCTCTCGAATTGCTGGATCG |

| | | |
|---|---|---|
| Bivalvia Picorna-like virus D51 | 20_TRINITY_DN76783_c0_g1_i3_827R | AATGCGAACTCCTAACGCGTTCAC |
| Bivalvia Hepeli-like virus D1 | 21_TRINITY_DN1218_c0_g1_i8_1740F | AATGCATCCGTCATCACCAGTGTC |
| Bivalvia Hepeli-like virus D1 | 21_TRINITY_DN1218_c0_g1_i8_2214R | TGGTCAAGGAATTAGTGCTAGGCC |
| Bivalvia Durna-like virus D11 | 21_TRINITY_DN23877_c0_g2_i2_174F | GGAAGACTACGACCTGTTAGTCTC |
| Bivalvia Durna-like virus D11 | 21_TRINITY_DN23877_c0_g2_i2_601R | CCTAATGATGCATCGTAGCCACTG |
| Bivalvia Picorna-like virus D58 | 21_TRINITY_DN45675_c0_g2_i1_230F | GCATCACATTCAAAGTGGCGCTCT |
| Bivalvia Picorna-like virus D58 | 21_TRINITY_DN45675_c0_g2_i1_673R | CCGGTGCGTTGTACTTCTTGCTTT |
| Bivalvia Picorna-like virus D60 | 21_TRINITY_DN66916_c0_g1_i3_373F | GCATGGCATCTTATCGCTTGGTGA |
| Bivalvia Picorna-like virus D60 | 21_TRINITY_DN66916_c0_g1_i3_1062R | GTTGTACTCGGCGACACTGTTTGT |
| Bivalvia Picorna-like virus D59 | 21_TRINITY_DN66916_c0_g2_i1_207F | GAATAACCGCCAAGCAAGTTCACC |
| Bivalvia Picorna-like virus D59 | 21_TRINITY_DN66916_c0_g2_i1_634R | GCAGCTGCCGACGATAATCAAATC |
| Bivalvia Picorna-like virus D62 | 22_TRINITY_DN17786_c0_g1_i8_1371F | CATTCACACACCTTTCATCCGTCC |
| Bivalvia Picorna-like virus D62 | 22_TRINITY_DN17786_c0_g1_i8_1872R | GCCCCGGGTAAAATTGCAAATAGG |
| Bivalvia Durna-like virus D13 | 22_TRINITY_DN28221_c0_g1_i1_322F | CACTACTGTGTTTGCCCAAAGGAG |
| Bivalvia Durna-like virus D13 | 22_TRINITY_DN28221_c0_g1_i1_811R | GTTAGCGGAACGGTTGAACCAATC |
| Bivalvia Hepeli-like virus D2 | 22_TRINITY_DN33039_c0_g1_i1_72F | ACTGAAAGTGTATTTCAATGCACG |
| Bivalvia Hepeli-like virus D2 | 22_TRINITY_DN33039_c0_g1_i1_706R | GATTCTTCCCAAGACTCTTCGTCC |
| Bivalvia Picorna-like virus D63 | 22_TRINITY_DN5657_c0_g1_i7_320F | CCTTTTCCTTTGAAACCCCAACCG |
| Bivalvia Picorna-like virus D63 | 22_TRINITY_DN5657_c0_g1_i7_743R | ATATGAGGGGTTTACCAGAGCACG |
| Gastropoda Sobeli-like virus D2 | 25_TRINITY_DN1292_c0_g1_i7_1081F | CAGATTCATTGGGTGGGATACACG |
| Gastropoda Sobeli-like virus D2 | 25_TRINITY_DN1292_c0_g1_i7_1508R | CCATGATGCAAATTCCAGAGCGAG |
| Gastropoda Picorna-like virus D11 | 25_TRINITY_DN14243_c0_g2_i2_86F | GGCATTGAGACCACAAAGTTTGCC |
| Gastropoda Picorna-like virus D11 | 25_TRINITY_DN14243_c0_g2_i2_535R | AGTCAGGGCAAGAGTACTGTGATG |
| Gastropoda Picorna-like virus D9 | 25_TRINITY_DN14243_c0_g3_i1_260F | ACTTTGATGAAACCTTGCGGCACC |
| Gastropoda Picorna-like virus D9 | 25_TRINITY_DN14243_c0_g3_i1_686R | TCGAAGAACGTGATAAGCAGCCAG |
| Gastropoda Picorna-like virus D8 | 25_TRINITY_DN14399_c0_g1_i1_233F | CGATTTCTGAACGTCCTGTGCCTT |
| Gastropoda Picorna-like virus D8 | 25_TRINITY_DN14399_c0_g1_i1_693R | CGGCACCATGTCAATCCAAACATC |
| Gastropoda Picorna-like virus D5 | 25_TRINITY_DN15438_c0_g3_i2_258F | GTTTGGTCTAATTCCTCGGCACCA |
| Gastropoda Picorna-like virus D5 | 25_TRINITY_DN15438_c0_g3_i2_776R | CACAAGTGCTCTGTCTACTGTTCC |
| Gastropoda Picorna-like virus D6 | 25_TRINITY_DN1564_c0_g1_i5_3601F | CATTGTTGCACACCTCAAAGAGGC |
| Gastropoda Picorna-like virus D6 | 25_TRINITY_DN1564_c0_g1_i5_4037R | GTAGGAAGAGGATGGACAGAGTTG |
| Gastropoda Picorna-like virus D10 | 25_TRINITY_DN4029_c0_g1_i1_112F | ACCGGCAATAACAGGCTGTTTACC |
| Gastropoda Picorna-like virus D10 | 25_TRINITY_DN4029_c0_g1_i1_552R | GGCATTGTGTGGTTTCCAAGACAC |
| Gastropoda Picorna-like virus D7 | 25_TRINITY_DN5970_c0_g1_i8_410F | TACGTTGAACCTCGGTAGCTTACC |
| Gastropoda Picorna-like virus D7 | 25_TRINITY_DN5970_c0_g1_i8_847R | GCATCTGCTGATAGCGCGTTTATC |
| Gastropoda Picorna-like virus D13 | 26_TRINITY_DN16980_c0_g1_i8_1013F | GAGTTCTGACGAATGCAACCACAG |
| Gastropoda Picorna-like virus D13 | 26_TRINITY_DN16980_c0_g1_i8_1439R | AGGTCCACGTCTTCATAGCTCATG |
| Gastropoda Unclassified virus D3 | 26_TRINITY_DN32994_c0_g1_i1_259F | GACCAAATCATATTGACGCGGACG |
| Gastropoda Unclassified virus D3 | 26_TRINITY_DN32994_c0_g1_i1_692R | ATAAACAAGGTCAGGTGGGCCTTC |
| Gastropoda Picorna-like virus D14 | 26_TRINITY_DN38383_c0_g1_i3_134F | GTACGAAATCACGAAGCCAAGTGC |
| Gastropoda Picorna-like virus D14 | 26_TRINITY_DN38383_c0_g1_i3_597R | AAACCCACTGTGACCAAAGCACAC |
| Gastropoda Picorna-like virus D16 | 26_TRINITY_DN38671_c0_g2_i1_122F | TCGAAGCCTACATCAGTTGCTGTC |
| Gastropoda Picorna-like virus D16 | 26_TRINITY_DN38671_c0_g2_i1_605R | TGACCGCTTCTGACCTAACAGATG |

| | | |
|---|---|---|
| Gastropoda Picorna-like virus D17 | 26_TRINITY_DN45248_c0_g2_i2_232F | CCGGTAAGCTTCAAACATCTCACG |
| Gastropoda Picorna-like virus D17 | 26_TRINITY_DN45248_c0_g2_i2_683R | GATATCAGATCCAGAGTGGTGCGA |
| Gastropoda Picorna-like virus D12 | 26_TRINITY_DN5270_c0_g1_i5_579F | ATGTGGGAACTTGGAGGTTCATGG |
| Gastropoda Picorna-like virus D12 | 26_TRINITY_DN5270_c0_g1_i5_1017R | AATGAGCGTTTCAGCTGGCGATTC |
| Gastropoda Picorna-like virus D18 | 26_TRINITY_DN63407_c0_g1_i1_258F | TCAGTCACACTTATGGTTGGCACC |
| Gastropoda Picorna-like virus D18 | 26_TRINITY_DN63407_c0_g1_i1_685R | TCAAGATGAAAGGGGGTGAGTCAC |
| Gastropoda Unclassified virus D4 | 26_TRINITY_DN67898_c0_g1_i1_723F | GGAACGCACATCGGTGTAGTAATC |
| Gastropoda Unclassified virus D4 | 26_TRINITY_DN67898_c0_g1_i1_1194R | TTTTACTCCGGTAGCCCTTGTGTG |
| Crustacea Picorna-like virus D1 | 3_TRINITY_DN13688_c0_g1_i1_906F | AAATCTGCTACTGCTGCATGTCCG |
| Crustacea Picorna-like virus D1 | 3_TRINITY_DN13688_c0_g1_i1_1394R | TCGCTTATGGCCGTTAGAACTACG |
| Crustacea Picorna-like virus D5 | 3_TRINITY_DN141_c0_g1_i11_1670F | AGTGTGACAGCTCTTATTCCGGAG |
| Crustacea Picorna-like virus D5 | 3_TRINITY_DN141_c0_g1_i11_2165R | TGAAGTACGTGGATGGAGAGATGC |
| Crustacea Unclassified virus D1 | 3_TRINITY_DN19160_c0_g1_i1_856F | GCCCTCCCATACCTACAAGATATC |
| Crustacea Unclassified virus D1 | 3_TRINITY_DN19160_c0_g1_i1_1296R | TCGTACCAAGGTGCTTGTTCACAG |
| Crustacea Picorna-like virus D3 | 3_TRINITY_DN35153_c0_g1_i1_908F | AACAACCTTGTGAATCGTGTCGCC |
| Crustacea Picorna-like virus D3 | 3_TRINITY_DN35153_c0_g1_i1_1359R | AATACGATGGTAACATGGCGCGAG |
| Crustacea Picorna-like virus D6 | 3_TRINITY_DN4609_c0_g1_i3_314F | CTTTCTCCAGTCCAGATCTTCACC |
| Crustacea Picorna-like virus D6 | 3_TRINITY_DN4609_c0_g1_i3_874R | CTGTTTCTGCTTACGGAGGACCTT |
| Crustacea Picorna-like virus D2 | 3_TRINITY_DN61560_c0_g1_i1_54F | AGAACCATGCTTATACCGTCCTCC |
| Crustacea Picorna-like virus D2 | 3_TRINITY_DN61560_c0_g1_i1_530R | TCTCTAACCACTCTTTGGGAGAGG |
| Crustacea Picorna-like virus D4 | 3_TRINITY_DN84776_c0_g1_i1_363F | GATCAGTTGATGCGTGAGCTACAC |
| Crustacea Picorna-like virus D4 | 3_TRINITY_DN84776_c0_g1_i1_912R | CGTGAGGTGTCTCATGCTATGTCT |
| Crustacea Picorna-like virus D7 | 4_TRINITY_DN10052_c0_g1_i1_353F | GGAAAAGTGACTTGTAGTCACCCC |
| Crustacea Picorna-like virus D7 | 4_TRINITY_DN10052_c0_g1_i1_778R | TGGAAGTTCTCAATGGAGACCACG |
| Crustacea Picorna-like virus D8 | 4_TRINITY_DN612_c0_g1_i2_3587F | GGCTCAAAGACTTATCGTCTGTCC |
| Crustacea Picorna-like virus D8 | 4_TRINITY_DN612_c0_g1_i2_4010R | TTTGCGTGTAGCAAAGTCTGCTGG |
| Bivalvia Durna-like virus D14 | 5_TRINITY_DN64381_c0_g1_i1_376F | GACAAATGCGGGACTACCTTCGAT |
| Bivalvia Durna-like virus D14 | 5_TRINITY_DN64381_c0_g1_i1_970R | GTCACCCTGGACTTGAGAGTACTT |
| Hexanauplia Picorna-like virus D1 | 6_TRINITY_DN104099_c0_g1_i1_326F | GCCTATGCTGTCATAAGTGTGAGC |
| Hexanauplia Picorna-like virus D1 | 6_TRINITY_DN104099_c0_g1_i1_802R | ATCCCTTACATGCAGAATACCCCC |
| Hexanauplia Bunya-like virus D1 | 6_TRINITY_DN13006_c0_g1_i1_5797F | GATAACAGGGTCAGAAAGGAGGTC |
| Hexanauplia Bunya-like virus D1 | 6_TRINITY_DN13006_c0_g1_i1_6241R | AACAGACTCTCTCAAGAGCACAGC |
| Hexanauplia Picorna-like virus D2 | 6_TRINITY_DN2661_c0_g3_i1_378F | CTGGGCTGCAATTTTGTGTGGGTT |
| Hexanauplia Picorna-like virus D2 | 6_TRINITY_DN2661_c0_g3_i1_1064R | GCAACACCCGTATCAAGACACAGT |
| Hexanauplia Picorna-like virus D3 | 6_TRINITY_DN51494_c0_g1_i1_128F | GCTGACCAGAGAGTTAAACTTCGG |
| Hexanauplia Picorna-like virus D3 | 6_TRINITY_DN51494_c0_g1_i1_759R | CGAAGTACTGCAAGATCGCTAGTC |
| Hexanauplia Picorna-like virus D4 | 6_TRINITY_DN80338_c0_g3_i1_226F | GAAACCGTGGTTGTGGTGAAGAAC |
| Hexanauplia Picorna-like virus D4 | 6_TRINITY_DN80338_c0_g3_i1_662R | TTCTTTATCTGGTGGCGGTTGGTC |
| Gastropoda Picorna-like virus D29 | 7_TRINITY_DN105202_c0_g1_i1_229F | CATTACGGTTCCTCGGAGGAAAGT |
| Gastropoda Picorna-like virus D29 | 7_TRINITY_DN105202_c0_g1_i1_695R | GAACTCTGTGGGTCAACATCTCTC |
| Gastropoda Picorna-like virus D27 | 7_TRINITY_DN111908_c0_g2_i1_279F | GACAACTTGCTCTGACCCATCTCT |
| Gastropoda Picorna-like virus D27 | 7_TRINITY_DN111908_c0_g2_i1_780R | CCGGTAGCCTTTTTCCTTTCTTCG |
| Gastropoda Picorna-like virus D28 | 7_TRINITY_DN217501_c0_g1_i1_410F | CAAAGGACCCATACAAGCGCCATA |

| | | |
|---|---|---|
| Gastropoda Picorna-like virus D28 | 7_TRINITY_DN217501_c0_g1_i1_1041R | GTGGTGTAAGCCCACAAAGTGTTG |
| Gastropoda Durna-like virus D2 | 7_TRINITY_DN22197_c0_g1_i1_291F | ATGAGGATGCTCACATGCGATCAC |
| Gastropoda Durna-like virus D2 | 7_TRINITY_DN22197_c0_g1_i1_721R | ATTCCGGTCCAATAACGTTGGTCG |
| Gastropoda Durna-like virus D1 | 7_TRINITY_DN65869_c0_g1_i2_252F | ATAGTCGGCACCTATTATCCCCTC |
| Gastropoda Durna-like virus D1 | 7_TRINITY_DN65869_c0_g1_i2_858R | TGGCCCGGAAGAAGTTGACAAAAG |
| Gastropoda Picorna-like virus D26 | 7_TRINITY_DN91604_c0_g1_i1_1883F | AGGGCATTACGGTTATGGAAACCG |
| Gastropoda Picorna-like virus D26 | 7_TRINITY_DN91604_c0_g1_i1_2319R | TCCCGGAAAACCATTAGTCTGTGC |
| Bivalvia Picorna-like virus D73 | 8-1_TRINITY_DN118249_c0_g4_i1_86F | AGCCTGATGCATCTAATGGAACTG |
| Bivalvia Picorna-like virus D73 | 8-1_TRINITY_DN118249_c0_g4_i1_548R | TCAACAACACCAACTTCAAAGTCC |
| Bivalvia Unclassified virus D3 | 8-1_TRINITY_DN124617_c0_g2_i1_239F | ATGCAAGAGGATTTTTGGCATGTG |
| Bivalvia Unclassified virus D3 | 8-1_TRINITY_DN124617_c0_g2_i1_610R | GAGCGGCAATTTCTTCCACATTC |
| Bivalvia Picorna-like virus D76 | 8-1_TRINITY_DN15500_c0_g3_i1_391F | ACGCAAGTAGCTGCTCAGC |
| Bivalvia Picorna-like virus D76 | 8-1_TRINITY_DN15500_c0_g3_i1_938R | CTAATCTCAATGGGCTTTTCGTGC |
| Bivalvia Picorna-like virus D69 | 8-1_TRINITY_DN160154_c0_g2_i1_291F | CAAAGTTCAGGGTTAGCAGCGTGA |
| Bivalvia Picorna-like virus D69 | 8-1_TRINITY_DN160154_c0_g2_i1_816R | GTAAACACGGTCACACAGCAATGC |
| Bivalvia Picorna-like virus D80 | 8-1_TRINITY_DN163267_c0_g1_i1_337F | CGAAAAGAAAGGAGCAGGGGAACA |
| Bivalvia Picorna-like virus D80 | 8-1_TRINITY_DN163267_c0_g1_i1_691R | GACCTATGTGGAAGAAGCCATTGG |
| Bivalvia Picorna-like virus D68 | 8-1_TRINITY_DN166016_c0_g1_i1_266F | GGAGGTTCAAGCTAATTGGGCACT |
| Bivalvia Picorna-like virus D68 | 8-1_TRINITY_DN166016_c0_g1_i1_674R | CTCGATACACTGTGCTAGCAAAGC |
| Bivalvia Picorna-like virus D67 | 8-1_TRINITY_DN182373_c0_g1_i1_392F | GTAGATTTTCCAAGACCGGCTCCT |
| Bivalvia Picorna-like virus D67 | 8-1_TRINITY_DN182373_c0_g1_i1_950R | CCTTCACGAAACCAGAAAGTGGGA |
| Bivalvia Toli-like virus D2 | 8-1_TRINITY_DN18524_c0_g1_i1_180F | CGGGGCCATTACCTCCATTTATTC |
| Bivalvia Toli-like virus D2 | 8-1_TRINITY_DN18524_c0_g1_i1_614R | ACGTAGGGTTAGAAGATACCCGAG |
| Bivalvia Picorna-like virus D70 | 8-1_TRINITY_DN190690_c0_g1_i1_95F | TGATGTGCCTAATCCGTGGAAAGC |
| Bivalvia Picorna-like virus D70 | 8-1_TRINITY_DN190690_c0_g1_i1_769R | TAAGCGGTCTCAAACTGCTTCTGC |
| Bivalvia Picorna-like virus D77 | 8-1_TRINITY_DN24911_c0_g2_i1_250F | ACATTTTGATTCCGGTGAACCCCG |
| Bivalvia Picorna-like virus D77 | 8-1_TRINITY_DN24911_c0_g2_i1_678R | AAGGGGCGCTACACTAAGTACATC |
| Bivalvia Picorna-like virus D79 | 8-1_TRINITY_DN38244_c0_g1_i1_51F | AGCAATAACCTCAGCAACTGAGGC |
| Bivalvia Picorna-like virus D79 | 8-1_TRINITY_DN38244_c0_g1_i1_484R | TCTCACGTACTTTGGACTCAGACG |
| Bivalvia Durna-like virus D15 | 8-1_TRINITY_DN4556_c0_g1_i1_333F | CCTCAGTAAAGCTCGGTACAAAGG |
| Bivalvia Durna-like virus D15 | 8-1_TRINITY_DN4556_c0_g1_i1_831R | TAGTCGCGGAAAGGTCCTTCTAAG |
| Bivalvia Picorna-like virus D75 | 8-1_TRINITY_DN46313_c0_g1_i2_3512F | GCATGGTTAGTTCCCTTGAAACGC |
| Bivalvia Picorna-like virus D75 | 8-1_TRINITY_DN46313_c0_g1_i2_3941R | TGGGTTACCATCAAGGCATGCATC |
| Bivalvia Picorna-like virus D74 | 8-1_TRINITY_DN62_c0_g1_i1_3730F | AAGGGGCTACTCGAAAAGATCGAG |
| Bivalvia Picorna-like virus D74 | 8-1_TRINITY_DN62_c0_g1_i1_4400R | CGGAAATGCTCAGAATGGATTCGC |
| Bivalvia Picorna-like virus D78 | 8-1_TRINITY_DN66185_c0_g3_i1_759F | TGAATAGAGTCCAGCGTCCAAGTC |
| Bivalvia Picorna-like virus D78 | 8-1_TRINITY_DN66185_c0_g3_i1_1238R | CAATCCCTTCGGGGTTTGGATAAG |
| Bivalvia Picorna-like virus D71 | 8-1_TRINITY_DN7678_c0_g2_i1_341F | CCGCAAAGAATTTCGGAGGCTTCT |
| Bivalvia Picorna-like virus D71 | 8-1_TRINITY_DN7678_c0_g2_i1_981R | GAATGTCGCTGATATGAGCGCTTC |
| Bivalvia Unclassified virus D4 | 8-1_TRINITY_DN86975_c0_g1_i2_119F | TTGTCGATAGTCACTTGTCAAGCG |
| Bivalvia Unclassified virus D4 | 8-1_TRINITY_DN86975_c0_g1_i2_576R | CCCCTGTTATATGGGACGGAAGAG |
| Bivalvia Durna-like virus D16 | 8-1_TRINITY_DN96678_c0_g1_i1_383F | CGGTAGGTTACGACCTTTACTCCT |
| Bivalvia Durna-like virus D16 | 8-1_TRINITY_DN96678_c0_g1_i1_1027R | CCCTGTACTTGAGAGAACTCGAGA |

| | | |
|---|---|---|
| Bivalvia Hepeli-like virus D3 | 8-2_TRINITY_DN128170_c0_g1_i2_141F | AAGCGGTCGAAAGACAAAGGAGAG |
| Bivalvia Hepeli-like virus D3 | 8-2_TRINITY_DN128170_c0_g1_i2_614R | CGGCTCCATACCTTTCGGAAATTG |
| Bivalvia Picorna-like virus D81 | 8-2_TRINITY_DN148753_c0_g3_i1_338F | GATGGTTCACAGCCAACTGGACAA |
| Bivalvia Picorna-like virus D81 | 8-2_TRINITY_DN148753_c0_g3_i1_1007R | GTGCAAGTGTACTCTGGACCTTCT |
| Bivalvia Picorna-like virus D86 | 8-2_TRINITY_DN320196_c0_g1_i1_131F | TTGTCCAATCCATTGGAGCACCTG |
| Bivalvia Picorna-like virus D86 | 8-2_TRINITY_DN320196_c0_g1_i1_616R | AATGTGGTCCTCCTGTGATGAAGG |
| Bivalvia Picorna-like virus D88 | 8-2_TRINITY_DN358270_c0_g1_i1_336F | GCGGATGCACTATGTATTGACGGT |
| Bivalvia Picorna-like virus D88 | 8-2_TRINITY_DN358270_c0_g1_i1_773R | CCGCCTTGCTTACTGAATTTAGGC |
| Bivalvia Picorna-like virus D83 | 8-2_TRINITY_DN369983_c0_g1_i1_234F | CAGCTCCCTATCTGTAGGAGTTGA |
| Bivalvia Picorna-like virus D83 | 8-2_TRINITY_DN369983_c0_g1_i1_644R | CCACCTAATCCACGGATATCCTCA |
| Bivalvia Picorna-like virus D84 | 8-2_TRINITY_DN41441_c0_g1_i2_237F | GGACATTTTGCACACGCCTATAGG |
| Bivalvia Picorna-like virus D84 | 8-2_TRINITY_DN41441_c0_g1_i2_676R | CCCAGATTGCCGTAATATTCCAGG |
| Bivalvia Picorna-like virus D89 | 8-2_TRINITY_DN44242_c0_g3_i2_134F | AAGTAGTGGTGTTCGTAAAGCTCC |
| Bivalvia Picorna-like virus D89 | 8-2_TRINITY_DN44242_c0_g3_i2_568R | GTTATCACTGCATGCATACGCC |
| Bivalvia Picorna-like virus D90 | 8-2_TRINITY_DN50636_c0_g1_i2_262F | CTGTGAAGGTAGACGCTGACGATA |
| Bivalvia Picorna-like virus D90 | 8-2_TRINITY_DN50636_c0_g1_i2_736R | CCTCAACCAGTTCGCATATGGCAA |
| Bivalvia Picorna-like virus D85 | 8-2_TRINITY_DN60228_c0_g1_i1_329F | CGTCGCGGCAACACCTTTAGATAT |
| Bivalvia Picorna-like virus D85 | 8-2_TRINITY_DN60228_c0_g1_i1_866R | GTATATCTGGTGTGCGGATCCTTC |
| Bivalvia Picorna-like virus D82 | 8-2_TRINITY_DN88231_c0_g2_i1_156F | GTAGGGTCCAAAATTGACTGCGTG |
| Bivalvia Picorna-like virus D82 | 8-2_TRINITY_DN88231_c0_g2_i1_593R | ACGGAAGACTTGTAGCAATAGGCC |
| Bivalvia Picorna-like virus N1 | BH05_TRINITY_DN217842_c0_g1_i1_336F | GGTCGGCATGTGAAAATAGCGCTT |
| Bivalvia Picorna-like virus N1 | BH05_TRINITY_DN217842_c0_g1_i1_855R | GATTGTGTAGACAGCGATGTCCGA |
| Bivalvia Durna-like virus N1 | BH05_TRINITY_DN238840_c0_g1_i1_109F | CTTTCCGAGGTTGCGTTGC |
| Bivalvia Durna-like virus N1 | BH05_TRINITY_DN238840_c0_g1_i1_639R | GGTAACCATGGTGTACCATCTGG |
| Bivalvia Amarillo-like virus N1 | BH05_TRINITY_DN2720_c0_g1_i2_5069F | TGGAAACCAGTGGGAGTATATGGC |
| Bivalvia Amarillo-like virus N1 | BH05_TRINITY_DN2720_c0_g1_i2_5514R | GCCATTTCCATTCCACTCTCATCC |
| Bivalvia Picorna-like virus N3 | BH05_TRINITY_DN3549_c0_g1_i8_1177F | GATGGAGATGGTACTGATCCAGTG |
| Bivalvia Picorna-like virus N3 | BH05_TRINITY_DN3549_c0_g1_i8_1623R | TGGAGTGGTCGCTGAAGATTTGTC |
| Bivalvia Picorna-like virus N2 | BH05_TRINITY_DN83599_c0_g1_i1_359F | GTCTTTGCAACCAGCACGCACAAT |
| Bivalvia Picorna-like virus N2 | BH05_TRINITY_DN83599_c0_g1_i1_916R | GACCATTGCAAGTGGTGTCATGTG |
| Bivalvia Picorna-like virus N4 | BH05_TRINITY_DN9263_c0_g2_i1_275F | GAACCAATTCGCAGAAGCCTGCAA |
| Bivalvia Picorna-like virus N4 | BH05_TRINITY_DN9263_c0_g2_i1_780R | GTGAACCCACACTTTGTTCAGGCT |
| Bivalvia Picorna-like virus N5 | BH06_TRINITY_DN17841_c0_g1_i7_284F | CACATCCCCAACTCATTGGAAGAG |
| Bivalvia Picorna-like virus N5 | BH06_TRINITY_DN17841_c0_g1_i7_863R | GGGGGGAATACGTATCTCTATCTC |
| Bivalvia Sobeli-like virus N1 | BH06_TRINITY_DN89577_c0_g1_i2_1062F | AAAGGGAATGCCAGGTGTGGAATC |
| Bivalvia Sobeli-like virus N1 | BH06_TRINITY_DN89577_c0_g1_i2_1486R | GCAAATACACCTCCTACACCTACG |
| Gastropoda Picorna-like virus N3 | BH09_TRINITY_DN25791_c0_g1_i4_343F | GTCTCCAGCTGTACCTCTCCTAAT |
| Gastropoda Picorna-like virus N3 | BH09_TRINITY_DN25791_c0_g1_i4_798R | GAGGACTCGAGTGATTTTCTGTGC |
| Gastropoda Toli-like virus N1 | BH10_TRINITY_DN92075_c0_g1_i2_370F | GCAGCGGAATGGTGGAATCTCATT |
| Gastropoda Toli-like virus N1 | BH10_TRINITY_DN92075_c0_g1_i2_1049R | CGAGATTTGACTCTGTGACTGGAG |
| Crustacea Jingchu-like virus N1 | BH11_TRINITY_DN113_c0_g1_i4_5401F | CTAGTATCATTGTTCTGGTGGGGG |
| Crustacea Jingchu-like virus N1 | BH11_TRINITY_DN113_c0_g1_i4_5856R | AAAGACAAGACTACCAGGACACCC |
| Crustacea Durna-like virus N1 | BH11_TRINITY_DN16288_c0_g1_i5_199F | GAACTACCACTAGCTAACGCTTCC |

| | | |
|---|---|---|
| Crustacea Durna-like virus N1 | BH11_TRINITY_DN16288_c0_g1_i5_631R | TCCACCCACAAAAGGGAGAAGATG |
| Crustacea Durna-like virus N2 | BH11_TRINITY_DN75_c0_g1_i12_589F | GTAGGTACTACAAAGACAGCGTCG |
| Crustacea Durna-like virus N2 | BH11_TRINITY_DN75_c0_g1_i12_1095R | CGGTAGCACTAAGCAAAGGAGTAC |
| Gastropoda Stella-like virus N1 | BH12_TRINITY_DN116276_c0_g1_i1_165F | GCGTCGTTGAAAGAATTTAGCCTG |
| Gastropoda Stella-like virus N1 | BH12_TRINITY_DN116276_c0_g1_i1_583R | ACTGTTGTGGATAACACTTTGGTG |
| Gastropoda Picorna-like virus N6 | BH12_TRINITY_DN38608_c0_g2_i1_310F | CCTCAACTTCAGTGTGAGCGTTGA |
| Gastropoda Picorna-like virus N6 | BH12_TRINITY_DN38608_c0_g2_i1_731R | CGCGGAAATTTTGCAGGTGTGCAA |
| Gastropoda Picorna-like virus N4 | BH12_TRINITY_DN89012_c0_g1_i2_324F | GCAAGGTCCCTCTGTAGATGAGTT |
| Gastropoda Picorna-like virus N4 | BH12_TRINITY_DN89012_c0_g1_i2_950R | GACTTGCCAATACCTGTGGTACCA |
| Gastropoda Sobeli-like virus N1 | BH12_TRINITY_DN91413_c0_g1_i1_98F | TGACCTTCCTGCATTCCATGATGG |
| Gastropoda Sobeli-like virus N1 | BH12_TRINITY_DN91413_c0_g1_i1_528R | TTTGGCGGAATTGGTGGAGAGAAG |
| Gastropoda Picorna-like virus N9 | BH13_TRINITY_DN15351_c0_g1_i7_512F | GGCGATTTTTGACGTGACCAAAGG |
| Gastropoda Picorna-like virus N9 | BH13_TRINITY_DN15351_c0_g1_i7_938R | ATTGAAGCGGTCTACAGCAAGCTG |
| Gastropoda Picorna-like virus N10 | BH13_TRINITY_DN16706_c0_g1_i1_256F | TCAGCCTTCAGTGACTGGTGTTTG |
| Gastropoda Picorna-like virus N10 | BH13_TRINITY_DN16706_c0_g1_i1_691R | AGCTAGAGTCAGACTACTGGACAG |
| Gastropoda Picorna-like virus N8 | BH13_TRINITY_DN262263_c0_g1_i1_214F | CTTCCGGTGGTATTTTGCCAGCTT |
| Gastropoda Picorna-like virus N8 | BH13_TRINITY_DN262263_c0_g1_i1_659R | GTACACTGGTCTTGAAACCACCAG |
| Gastropoda Durna-like virus N1 | BH13_TRINITY_DN48022_c0_g1_i3_384F | GCAACGGTCTTCGCTAAGCAAATC |
| Gastropoda Durna-like virus N1 | BH13_TRINITY_DN48022_c0_g1_i3_976R | CGCTTCGCACTAAAGTGAAGCTTC |
| Crustacea Picorna-like virus N1 | G06_TRINITY_DN10475_c0_g1_i3_202F | ACAGCTGGGATTGCGAAATTCCAG |
| Crustacea Picorna-like virus N1 | G06_TRINITY_DN10475_c0_g1_i3_706R | AGTAAAGGCACCCTTGTTGGGTTG |
| Crustacea Picorna-like virus N2 | G06_TRINITY_DN4195_c0_g1_i3_17F | TTCGATTCAGACCACCTTGTAGCC |
| Crustacea Picorna-like virus N2 | G06_TRINITY_DN4195_c0_g1_i3_448R | GTGAGTGTCTGCGTAATGTCCTAC |
| Crustacea Picorna-like virus N5 | G06_TRINITY_DN42055_c0_g1_i1_304F | CGTATTGCACAAGCAGACGTCCTT |
| Crustacea Picorna-like virus N5 | G06_TRINITY_DN42055_c0_g1_i1_665R | CACCCACACTGAAGGTATTGACCA |
| Crustacea Picorna-like virus N3 | G06_TRINITY_DN5219_c0_g1_i2_409F | GTGCAGATCTTCCACCTGCTTGTA |
| Crustacea Picorna-like virus N3 | G06_TRINITY_DN5219_c0_g1_i2_1122R | CATGCGAGAGAAAACCCTCTCGTT |
| Crustacea Sobeli-like virus N1 | G06_TRINITY_DN705_c0_g1_i3_784F | GTGTATGGAGTCCACACAAATGGC |
| Crustacea Sobeli-like virus N1 | G06_TRINITY_DN705_c0_g1_i3_1232R | AATTCCTCTATGTACGGGGACTCC |
| Crustacea Picorna-like virus N4 | G06_TRINITY_DN8627_c0_g1_i1_251F | CAGAGAGGAGAGGACAATTCCAAC |
| Crustacea Picorna-like virus N4 | G06_TRINITY_DN8627_c0_g1_i1_772R | CTTGCAGTTTGCTCATCGTAGCCT |
| Crustacea Picorna-like virus N6 | G07_TRINITY_DN120_c0_g1_i4_4612F | GGGAGAAGCTCAGATTTGAGCTTG |
| Crustacea Picorna-like virus N6 | G07_TRINITY_DN120_c0_g1_i4_5079R | AGTTCTCGAATTCGACCAGATGGC |
| Crustacea Picorna-like virus N7 | G07_TRINITY_DN3356_c0_g1_i2_304F | CAAGAGTACCAGCGAAAGGAAGAG |
| Crustacea Picorna-like virus N7 | G07_TRINITY_DN3356_c0_g1_i2_837R | GCCCTGACATTAACGGATATCTCG |
| Crustacea Picorna-like virus N8 | G07_TRINITY_DN3356_c0_g2_i2_377F | GAAGAGGCACCTAGACTCTCCAAT |
| Crustacea Picorna-like virus N8 | G07_TRINITY_DN3356_c0_g2_i2_1059R | CGATTTACAACACATCGGGAGCAG |
| Crustacea Picorna-like virus N9 | G07_TRINITY_DN4331_c0_g1_i3_282F | CATCAAGCGAGATATTGAGGCCGT |
| Crustacea Picorna-like virus N9 | G07_TRINITY_DN4331_c0_g1_i3_860R | CTGACGAGTTGGATGACAACTGTG |
| Crustacea Ghabri-like virus N1 | G08_TRINITY_DN464_c0_g1_i1_3494F | ATAGGGATGGGGCGTTGCTTTAAG |
| Crustacea Ghabri-like virus N1 | G08_TRINITY_DN464_c0_g1_i1_3920R | ACTCCAGCGGTTATTCAGATGCTG |
| Crustacea Picorna-like virus N10 | G10_TRINITY_DN11011_c0_g1_i1_248F | TCACCGATTGTCTTGACCTTACGC |
| Crustacea Picorna-like virus N10 | G10_TRINITY_DN11011_c0_g1_i1_751R | CTGCGATTCTGTTCGGACAATGAG |

| | | |
|---|---|---|
| Crustacea Toli-like virus N1 | G10_TRINITY_DN2373_c0_g1_i8_1992F | AGGTCACAGAATTCTCGGTGACAC |
| Crustacea Toli-like virus N1 | G10_TRINITY_DN2373_c0_g1_i8_2432R | AGTCTCCAAGGACTTTGTCGAGAC |
| Crustacea Ghabri-like virus N2 | G10_TRINITY_DN3803_c0_g1_i5_268F | CACCGCTGTTGGTAGGTCAAAAAC |
| Crustacea Ghabri-like virus N2 | G10_TRINITY_DN3803_c0_g1_i5_691R | GATTTCTTTGGCGACATCAGCAGG |
| Crustacea Picorna-like virus N11 | G10_TRINITY_DN55839_c0_g1_i1_325F | CCAAAATATGGGACGTAGGTGTGG |
| Crustacea Picorna-like virus N11 | G10_TRINITY_DN55839_c0_g1_i1_926R | GTTAAACTGTCCATCGAAGGCGTG |
| Crustacea Picorna-like virus N12 | G10_TRINITY_DN88610_c0_g1_i1_152F | CTTGAAACATGCCGCTCCTTCATG |
| Crustacea Picorna-like virus N12 | G10_TRINITY_DN88610_c0_g1_i1_568R | ATGTGTAGAGCTAGGCAGGATAGG |
| Gastropoda Picorna-like virus N11 | G11_TRINITY_DN11242_c0_g1_i1_344F | GGTAGCTCTCTTGAGCGCTTTGTT |
| Gastropoda Picorna-like virus N11 | G11_TRINITY_DN11242_c0_g1_i1_1010R | CTTCACGGCATTGGTTCTGACGTT |
| Gastropoda Unclassified virus N1 | G11_TRINITY_DN1339_c0_g1_i3_123F | ACATTGACGGAAATGAGGGGGTTC |
| Gastropoda Unclassified virus N1 | G11_TRINITY_DN1339_c0_g1_i3_549R | TCAGAGTTATCCCCATCATGCCAG |
| Gastropoda Picorna-like virus N14 | G11_TRINITY_DN1500_c0_g1_i3_1050F | AGGTGAGCCAACGATTTGAGTAGC |
| Gastropoda Picorna-like virus N14 | G11_TRINITY_DN1500_c0_g1_i3_1559R | TGTGAATTCACGGACGTTGCACAC |
| Gastropoda Picorna-like virus N15 | G11_TRINITY_DN1861_c0_g1_i1_218F | TTTATGTACACTGCGTGCAGCTGG |
| Gastropoda Picorna-like virus N15 | G11_TRINITY_DN1861_c0_g1_i1_717R | CCACAAGCCAGCTCAAGAATACAG |
| Gastropoda Picorna-like virus N13 | G11_TRINITY_DN5796_c0_g1_i2_469F | GGATTGTGTTATGATGGCCATGCC |
| Gastropoda Picorna-like virus N13 | G11_TRINITY_DN5796_c0_g1_i2_841R | CATACCCTCACGATTGTTACCTGC |
| Bivalvia Picorna-like virus N14 | G12_TRINITY_DN134571_c0_g2_i1_545F | CGTCAACTTCCTACCCTCTATTGG |
| Bivalvia Picorna-like virus N14 | G12_TRINITY_DN134571_c0_g2_i1_1004R | GGCTGGTAGTGAATCTAGGGCTTT |
| Bivalvia Picorna-like virus N9 | G12_TRINITY_DN13497_c0_g1_i2_405F | ATCGATTCCCGAAGAACCTGTGAG |
| Bivalvia Picorna-like virus N9 | G12_TRINITY_DN13497_c0_g1_i2_887R | CCTCTTACGCTAGCACGTCTTATC |
| Bivalvia Picorna-like virus N13 | G12_TRINITY_DN1475_c0_g1_i1_2383F | GGTGGCTGTGAAAGTGACATTTGG |
| Bivalvia Picorna-like virus N13 | G12_TRINITY_DN1475_c0_g1_i1_2808R | ATGTCGCCCACAGACCAAAGAAAC |
| Bivalvia Picorna-like virus N15 | G12_TRINITY_DN36494_c0_g1_i1_138F | TCCACCAGTGATCAAGGGTCG |
| Bivalvia Picorna-like virus N15 | G12_TRINITY_DN36494_c0_g1_i1_585R | AGCTTATGTGCCTCAAACCCATG |
| Bivalvia Picorna-like virus N10 | G12_TRINITY_DN45079_c0_g1_i3_298F | CATTGCGCAGATTTCCCTACGTTG |
| Bivalvia Picorna-like virus N10 | G12_TRINITY_DN45079_c0_g1_i3_750R | TAGCCAGTTCTCCTGTCGTAAGTG |
| Bivalvia Picorna-like virus N11 | G12_TRINITY_DN73867_c0_g1_i1_213F | CCTCCATCAGCTTAACGTGTTCTG |
| Bivalvia Picorna-like virus N11 | G12_TRINITY_DN73867_c0_g1_i1_628R | CATGTTCGACACCATAGCGGTTTC |
| Bivalvia Picorna-like virus N16 | G13_TRINITY_DN20817_c0_g1_i1_226F | GAAGAAGAAGAGTAAGGGCCGTAC |
| Bivalvia Picorna-like virus N16 | G13_TRINITY_DN20817_c0_g1_i1_650R | CTGAACTCACACGGTCGAAATGAG |
| Bivalvia Wolfram-like virus N1 | G13_TRINITY_DN7700_c0_g1_i6_17F | CTGTGCTTCGACCGAAGTTATCTG |
| Bivalvia Wolfram-like virus N1 | G13_TRINITY_DN7700_c0_g1_i6_443R | TGATATAGGATCCACCGACTCCAG |
| Bivalvia Durna-like virus N2 | G13_TRINITY_DN81622_c0_g1_i1_526F | CGCCCTGGACTTGAGAGTACATAA |
| Bivalvia Durna-like virus N2 | G13_TRINITY_DN81622_c0_g1_i1_753R | CCTTCGAAGTGTATTTCCCAAGCC |
| Bivalvia Durna-like virus N3 | G14_TRINITY_DN11170_c0_g1_i1_494F | CTGGATCAAAGGTTGCATACGCAC |
| Bivalvia Durna-like virus N3 | G14_TRINITY_DN11170_c0_g1_i1_930R | CAGACCAATCCTTGAGTATCAGCG |
| Bivalvia Picorna-like virus N25 | G14_TRINITY_DN125042_c0_g1_i1_100F | CCTGAATTTGAGGCTCTTGTAGCC |
| Bivalvia Picorna-like virus N25 | G14_TRINITY_DN125042_c0_g1_i1_749R | CGACGATGCAAGGCATCAATACAC |
| Bivalvia Picorna-like virus N22 | G14_TRINITY_DN128452_c0_g1_i1_266F | TTATGGTTGTCAGAGGATTGCCGG |
| Bivalvia Picorna-like virus N22 | G14_TRINITY_DN128452_c0_g1_i1_704R | CCTCGTCACATCAACAAGACTGAC |
| Bivalvia Picorna-like virus N17 | G14_TRINITY_DN134_c0_g1_i12_4417F | CCTACGTTTCATTGCACGGTTCTG |

| | | |
|---|---|---|
| Bivalvia Picorna-like virus N17 | G14_TRINITY_DN134_c0_g1_i12_4862R | GTCCGGTTCATCGACCTAGATTTC |
| Bivalvia Picorna-like virus N18 | G14_TRINITY_DN134_c1_g1_i4_415F | CTTTGATTGGCAGGAAGCTATGGC |
| Bivalvia Picorna-like virus N18 | G14_TRINITY_DN134_c1_g1_i4_979R | CGCTCTTGCCGAAACATTTTGGAC |
| Bivalvia Picorna-like virus N24 | G14_TRINITY_DN135456_c0_g1_i1_287F | GCGCGGCTCCACAACAAATTTGTT |
| Bivalvia Picorna-like virus N24 | G14_TRINITY_DN135456_c0_g1_i1_775R | GTCGACAATGTCGCTTCATGGTTC |
| Bivalvia Picorna-like virus N30 | G14_TRINITY_DN143872_c0_g3_i1_434F | CTAGGGGTTGTGCTGGTGTATTGT |
| Bivalvia Picorna-like virus N30 | G14_TRINITY_DN143872_c0_g3_i1_1082R | GCTTCACAACGGTAAACTTCCTCC |
| Bivalvia Picorna-like virus N20 | G14_TRINITY_DN1548_c0_g1_i3_592F | AACCAATGCGGGTAAACGCAAAGG |
| Bivalvia Picorna-like virus N20 | G14_TRINITY_DN1548_c0_g1_i3_1226R | CTCACGCTAGTCATCATTCTGAGG |
| Bivalvia Picorna-like virus N21 | G14_TRINITY_DN1548_c1_g1_i3_250F | AGGAGACCAAAATGCAAGAGCAGG |
| Bivalvia Picorna-like virus N21 | G14_TRINITY_DN1548_c1_g1_i3_677R | CATCACCATCGGCAACAAAGTAGG |
| Bivalvia Picorna-like virus N27 | G14_TRINITY_DN7_c0_g1_i3_153F | TCATCTAGCACAAACCGCATCGAC |
| Bivalvia Picorna-like virus N27 | G14_TRINITY_DN7_c0_g1_i3_816R | AGGTAGAAGAGGAAGTGCAGCTAC |
| Bivalvia Picorna-like virus N26 | G14_TRINITY_DN89352_c0_g2_i2_283F | GATCCATAGTCATCGTCTACTGGG |
| Bivalvia Picorna-like virus N26 | G14_TRINITY_DN89352_c0_g2_i2_757R | CTCCAGGTGCTATTAATGCACGAC |
| Bivalvia Picorna-like virus N28 | G14_TRINITY_DN9013_c0_g1_i2_440F | GGAGATGTTGAGCCTGAACAAGCA |
| Bivalvia Picorna-like virus N28 | G14_TRINITY_DN9013_c0_g1_i2_861R | GCTATCTCCGGTGATGCCAAAGTT |
| Bivalvia Picorna-like virus N19 | G14_TRINITY_DN92252_c0_g2_i2_392F | CACTGGACTGTTTCAGCGCTCAAT |
| Bivalvia Picorna-like virus N19 | G14_TRINITY_DN92252_c0_g2_i2_1030R | CCCGGAAAGTAAACGTGGTATGGT |
| Gastropoda Picorna-like virus H1 | H01_TRINITY_DN119159_c0_g2_i1_79F | CACTAAACAGGAATCTCCTCCAGC |
| Gastropoda Picorna-like virus H1 | H01_TRINITY_DN119159_c0_g2_i1_515R | AATCCGAAAGAGGGTGAATAGCAG |
| Gastropoda Picorna-like virus H2 | H01_TRINITY_DN60146_c0_g5_i1_50F | GCTCGTAACAAACAACAGCGAGTG |
| Gastropoda Picorna-like virus H2 | H01_TRINITY_DN60146_c0_g5_i1_781R | AGGAGGTCTCCGATAGCTAGAAAC |
| Bivalvia Picorna-like virus H5 | H02_TRINITY_DN1250_c0_g1_i3_1978F | AAGGAATTTCCACGGCATCTGACG |
| Bivalvia Picorna-like virus H5 | H02_TRINITY_DN1250_c0_g1_i3_2405R | GAGGGTTGATTTCCCAATACCAGG |
| Bivalvia Durna-like virus H2 | H02_TRINITY_DN12567_c0_g1_i4_161F | TACTTTTCCGTCGACTCTGGGAAG |
| Bivalvia Durna-like virus H2 | H02_TRINITY_DN12567_c0_g1_i4_714R | GCGCTTGGGAAGGTTTTAGTCAAG |
| Bivalvia Picorna-like virus H4 | H02_TRINITY_DN34742_c0_g1_i2_286F | GGAAAGAACGGCTGGTTCATCAGT |
| Bivalvia Picorna-like virus H4 | H02_TRINITY_DN34742_c0_g1_i2_683R | GGTACTGGCTTTATGATCCCAGGA |
| Bivalvia Durna-like virus H1 | H02_TRINITY_DN4_c0_g1_i2_260F | CCCGAATCTATTCTATCAGGGTGG |
| Bivalvia Durna-like virus H1 | H02_TRINITY_DN4_c0_g1_i2_688R | TAAGATGGCAGCAAGAGGGTAACC |
| Bivalvia Picorna-like virus H2 | H02_TRINITY_DN43443_c0_g1_i1_43F | TCAGCAGCTTTGGAAGCAAAGTCC |
| Bivalvia Picorna-like virus H2 | H02_TRINITY_DN43443_c0_g1_i1_469R | ATTTCGTGCCTGCGACCTCTTTAG |
| Bivalvia Picorna-like virus H3 | H02_TRINITY_DN5051_c0_g1_i3_114F | GTCTTGCCGTCAATAACTTCCTCC |
| Bivalvia Picorna-like virus H3 | H02_TRINITY_DN5051_c0_g1_i3_561R | CGCGTTCGAAATTCACAACTGCTG |
| Bivalvia Picorna-like virus H7 | H02_TRINITY_DN5152_c0_g1_i1_408F | GATGATGGTAGAACACGCACGATG |
| Bivalvia Picorna-like virus H7 | H02_TRINITY_DN5152_c0_g1_i1_826R | GGCACTTCTGTAACTCCCATCCAA |
| Bivalvia Picorna-like virus H6 | H02_TRINITY_DN60471_c0_g1_i1_549F | CGGAAAAGGAGATCCAAGAGGAAC |
| Bivalvia Picorna-like virus H6 | H02_TRINITY_DN60471_c0_g1_i1_921R | CGCTTCCCGAGTAAGAAGCAAATG |
| Bivalvia Amarillo-like virus H1 | H02_TRINITY_DN67462_c0_g1_i21_277F | CATTTGGCTGCTTGATCGAGAGGT |
| Bivalvia Amarillo-like virus H1 | H02_TRINITY_DN67462_c0_g1_i21_793R | GGTCCCTCTAGAGCATTGTTTCTC |
| Bivalvia Picorna-like virus H1 | H02_TRINITY_DN7318_c0_g1_i15_681F | GCACCATTTGTCTTTCGTTGAGGC |
| Bivalvia Picorna-like virus H1 | H02_TRINITY_DN7318_c0_g1_i15_1108R | GAATGGAACCTTAACGAGGTCTCC |

| | | |
|---|---|---|
| Bivalvia Bunya-like virus H1 | H02_TRINITY_DN84_c0_g1_i3_1621F | GTCTCCGACATATATGCCTCTGTC |
| Bivalvia Bunya-like virus H1 | H02_TRINITY_DN84_c0_g1_i3_2055R | ACAACACTCAAAGAGTACGGCGAG |
| Bivalvia Durna-like virus H5 | H03_TRINITY_DN4623_c0_g1_i1_334F | AGCTCTGTACGTAGGATAGATGCC |
| Bivalvia Durna-like virus H5 | H03_TRINITY_DN4623_c0_g1_i1_840R | CTCATCTACCATGCTCGTAGTCAG |
| Bivalvia Durna-like virus H3 | H03_TRINITY_DN72722_c0_g1_i1_256F | GTGAGGTACTCGAAGGTTGGGATA |
| Bivalvia Durna-like virus H3 | H03_TRINITY_DN72722_c0_g1_i1_764R | GTCTCTTAATGCTGCTCTCCAAGG |
| Bivalvia Durna-like virus H4 | H03_TRINITY_DN78573_c0_g1_i1_1135F | TACGGGTGAAGCTGCAACGTATTC |
| Bivalvia Durna-like virus H4 | H03_TRINITY_DN78573_c0_g1_i1_1611R | TCAATAGGGTAGAAGCCAGGTTGC |
| Crustacea Ghabri-like virus H3 | H04_TRINITY_DN100462_c0_g1_i1_365F | GCAGTGGTAGGCTGATGGTCAAAT |
| Crustacea Ghabri-like virus H3 | H04_TRINITY_DN100462_c0_g1_i1_1065R | GAGCTTTTCCTCACTGCAGCTAAG |
| Crustacea Unclassified virus H1 | H04_TRINITY_DN134457_c0_g1_i1_383F | CGGATTCGGCAAAACACAAACTGG |
| Crustacea Unclassified virus H1 | H04_TRINITY_DN134457_c0_g1_i1_863R | GCGTCCGATTCGAATGTTTGTGGA |
| Crustacea Bunya-like virus H1 | H04_TRINITY_DN202600_c0_g1_i1_4050F | GATCTGTGGACTGTTGTCAATGCC |
| Crustacea Bunya-like virus H1 | H04_TRINITY_DN202600_c0_g1_i1_4868R | TAGATTAATACTTACTGATAAAAC |
| Crustacea Ghabri-like virus H1 | H04_TRINITY_DN38222_c0_g1_i6_449F | TAAGAAGTTGAGGCCGTGGACATC |
| Crustacea Ghabri-like virus H1 | H04_TRINITY_DN38222_c0_g1_i6_905R | TATCCGCTGAGCTTTTAGGCTGTG |
| Crustacea Ghabri-like virus H2 | H04_TRINITY_DN993_c0_g1_i1_2651F | GCAGAACACGTTGCTAACTTACGC |
| Crustacea Ghabri-like virus H2 | H04_TRINITY_DN993_c0_g1_i1_3120R | TGTGAGCATGCCAATGTATCTGGG |
| Crustacea Picorna-like virus N14 | ZH03_TRINITY_DN1649_c0_g1_i3_2169F | TATGTCTGGTACTACAGCGGCATC |
| Crustacea Picorna-like virus N14 | ZH03_TRINITY_DN1649_c0_g1_i3_2598R | CCTCAGCATATTGACAAGGCCATC |
| Crustacea Picorna-like virus N13 | ZH03_TRINITY_DN1695_c0_g1_i10_912F | TCAGTGGCCAACGTTCTGTCAAAC |
| Crustacea Picorna-like virus N13 | ZH03_TRINITY_DN1695_c0_g1_i10_1337R | ACCAGAAGGTGGTAGGAAATAGGG |
| Crustacea Picorna-like virus N15 | ZH03_TRINITY_DN1759_c0_g1_i6_219F | CTTGCTATGACTTTACAACCGCCG |
| Crustacea Picorna-like virus N15 | ZH03_TRINITY_DN1759_c0_g1_i6_651R | CGAGAACTTGTGAGCCAAAAGACC |
| Crustacea Picorna-like virus N27 | ZH04_TRINITY_DN12125_c0_g1_i2_61F | CTACTCTTTAAGACGGCAGTCTCG |
| Crustacea Picorna-like virus N27 | ZH04_TRINITY_DN12125_c0_g1_i2_499R | ATGTAGATTCGCTCGTTTCCGAGC |
| Crustacea Picorna-like virus N16 | ZH04_TRINITY_DN14124_c0_g1_i1_33F | CCTTAGCATGATCTTGAAGGATGT |
| Crustacea Picorna-like virus N16 | ZH04_TRINITY_DN14124_c0_g1_i1_501R | ATCACTGAACAGATCAAGGAGTCG |
| Crustacea Picorna-like virus N28 | ZH04_TRINITY_DN17951_c0_g1_i2_104F | CGAGACTGTGTATCCTCCAGTATC |
| Crustacea Picorna-like virus N28 | ZH04_TRINITY_DN17951_c0_g1_i2_559R | ATTGTCTACGGACCTCCTGCTAAC |
| Crustacea Picorna-like virus N20 | ZH04_TRINITY_DN1912_c0_g1_i11_1813F | GCTACCGACAACTCTGTGTCTATG |
| Crustacea Picorna-like virus N20 | ZH04_TRINITY_DN1912_c0_g1_i11_2250R | AACGAATTCATCGTCACTCTCCGC |
| Crustacea Picorna-like virus N17 | ZH04_TRINITY_DN25765_c0_g1_i2_284F | GTTTGTCAATGTTGGCAAACGGGG |
| Crustacea Picorna-like virus N17 | ZH04_TRINITY_DN25765_c0_g1_i2_786R | CGGAGAGCTCTATGACAACGATGA |
| Crustacea Picorna-like virus N29 | ZH04_TRINITY_DN4100_c0_g1_i20_794F | CTGGTTGCGTCACAACCATAAAGG |
| Crustacea Picorna-like virus N29 | ZH04_TRINITY_DN4100_c0_g1_i20_1223R | TTGAGGATGCGTCTAGTCATGAGG |
| Crustacea Picorna-like virus N22 | ZH04_TRINITY_DN450_c0_g1_i9_1315F | CCCGAGCAGAGTATCAATTAACGG |
| Crustacea Picorna-like virus N22 | ZH04_TRINITY_DN450_c0_g1_i9_1739R | ACTCCACATCACTGAGTTCTCACC |
| Crustacea Picorna-like virus N19 | ZH04_TRINITY_DN45473_c0_g1_i1_323F | GAAGTATTCCACGAATCGCCCTAG |
| Crustacea Picorna-like virus N19 | ZH04_TRINITY_DN45473_c0_g1_i1_906R | GAAAACCAGCTGCGCTAGTATCAG |
| Crustacea Picorna-like virus N21 | ZH04_TRINITY_DN494_c0_g1_i5_110F | CTCCACCATTGAAACTGTGAAGCC |
| Crustacea Picorna-like virus N21 | ZH04_TRINITY_DN494_c0_g1_i5_563R | TGGCAGATACTGCTTCTGGAATCC |
| Crustacea Picorna-like virus N24 | ZH04_TRINITY_DN641_c0_g1_i16_703F | GACATGATAGCACCTTCTGTGGTG |

| | | |
|---|---|---|
| Crustacea Picorna-like virus N24 | ZH04_TRINITY_DN641_c0_g1_i16_1134R | AAATCCCAAGCTTGTTTGCGACGC |
| Crustacea Picorna-like virus N18 | ZH04_TRINITY_DN7212_c0_g1_i4_457F | AGGTTTGTTACATGGCATCCCTCC |
| Crustacea Picorna-like virus N18 | ZH04_TRINITY_DN7212_c0_g1_i4_905R | TCTTCGTCACTCAGTCTGTTGTCG |
| Crustacea Picorna-like virus N25 | ZH04_TRINITY_DN782_c0_g1_i5_1024F | TACTCGGTGGAACCGAGTTCAATC |
| Crustacea Picorna-like virus N25 | ZH04_TRINITY_DN782_c0_g1_i5_1453R | CACACATCATCCAGTTAGCCAAGG |
| Crustacea Picorna-like virus N26 | ZH04_TRINITY_DN903_c0_g1_i5_365F | CAAAGTTCGTCCACAGTTACGCGT |
| Crustacea Picorna-like virus N26 | ZH04_TRINITY_DN903_c0_g1_i5_1015R | CATCGCTACCACACAAGGAAGTGT |
| Crustacea Picorna-like virus N23 | ZH04_TRINITY_DN960_c0_g1_i9_75F | CGATCTCTTCACTTCGAACTCCAC |
| Crustacea Picorna-like virus N23 | ZH04_TRINITY_DN960_c0_g1_i9_548R | GCTGGGATTGCAGAATTTCAGTGT |
| Crustacea Picorna-like virus N30 | ZH05_TRINITY_DN22607_c0_g1_i2_607F | GCAACCATGGAACTAACTTCACGG |
| Crustacea Picorna-like virus N30 | ZH05_TRINITY_DN22607_c0_g1_i2_1031R | GTTGAGCGCTTTGCTAGCATCTTC |
| Crustacea Picorna-like virus N31 | ZH05_TRINITY_DN3251_c0_g1_i5_538F | CTTATGCCTGAACGTGTCAAAGCC |
| Crustacea Picorna-like virus N31 | ZH05_TRINITY_DN3251_c0_g1_i5_962R | TGCTTAGGCCCACTTCCTCTATTC |
| Crustacea Picorna-like virus N33 | ZH06_TRINITY_DN18407_c0_g1_i3_390F | GTTTTGGGCTTGCTCAGAGTAGAG |
| Crustacea Picorna-like virus N33 | ZH06_TRINITY_DN18407_c0_g1_i3_1091R | GAAGACGGCATACGAGATGCCAAT |
| Crustacea Unclassified virus N1 | ZH06_TRINITY_DN2721_c0_g1_i9_160F | GCGAAAGTACTCGTATACACGGAC |
| Crustacea Unclassified virus N1 | ZH06_TRINITY_DN2721_c0_g1_i9_586R | AAAAGAGGTGGTCTCGCAGAATGC |
| Crustacea Picorna-like virus N32 | ZH06_TRINITY_DN546_c0_g1_i1_3795F | AAACTCCACCTGTGCATCTGTTCC |
| Crustacea Picorna-like virus N32 | ZH06_TRINITY_DN546_c0_g1_i1_4223R | TGGGTTACTAACGACGACATTCCC |
| Bivalvia Amarillo-like virus N3 | ZH07_TRINITY_DN11327_c0_g1_i3_1077F | TGCCATAAGTTGCGTGCATGAACC |
| Bivalvia Amarillo-like virus N3 | ZH07_TRINITY_DN11327_c0_g1_i3_1502R | AAGCCCCACCTTTTGAAGTTCGTC |
| Bivalvia Bunya-like virus N2 | ZH07_TRINITY_DN19819_c0_g1_i5_362F | GTGGCATCTTCACCTCCGTCTATT |
| Bivalvia Bunya-like virus N2 | ZH07_TRINITY_DN19819_c0_g1_i5_1103R | CAAAAGGGATGCAAGGACCTGAAC |
| Bivalvia Picorna-like virus N31 | ZH07_TRINITY_DN22405_c0_g1_i5_695F | AACACCAGCTGTTACTGTACTGGC |
| Bivalvia Picorna-like virus N31 | ZH07_TRINITY_DN22405_c0_g1_i5_1247R | AGGTGCTAGCCCAGTCATTAGAAG |
| Bivalvia Picorna-like virus N32 | ZH07_TRINITY_DN40582_c0_g1_i10_201F | AATCTGTACTGCTGGTTGTGACGG |
| Bivalvia Picorna-like virus N32 | ZH07_TRINITY_DN40582_c0_g1_i10_697R | TATCCTGTCCTATGCAGAGTGCAC |
| Bivalvia Sobeli-like virus N2 | ZH07_TRINITY_DN473_c0_g1_i9_452F | TATCAATTCCGACGCAGACCGAAC |
| Bivalvia Sobeli-like virus N2 | ZH07_TRINITY_DN473_c0_g1_i9_876R | TCACTACTCCTCTTTGGTAGGCAC |
| Bivalvia Bunya-like virus N1 | ZH07_TRINITY_DN8577_c0_g1_i4_1286F | GGCTTTCAAATCGCCAGGTTTCAC |
| Bivalvia Bunya-like virus N1 | ZH07_TRINITY_DN8577_c0_g1_i4_1710R | ATCCAGAGTACGTGAGAATCGCAG |
| Bivalvia Amarillo-like virus N2 | ZH07_TRINITY_DN8886_c0_g1_i5_3549F | GGCGCGACTAGATAGGAATCATTG |
| Bivalvia Amarillo-like virus N2 | ZH07_TRINITY_DN8886_c0_g1_i5_3978R | ATTGGAGACAAGCCTGTCATTGCG |
| Bivalvia Picorna-like virus N36 | ZH08_TRINITY_DN168754_c0_g1_i1_36F | ACATGTCAATTCCAAAGTGTCACC |
| Bivalvia Picorna-like virus N36 | ZH08_TRINITY_DN168754_c0_g1_i1_536R | TCTGCATGTTTTGCATGAGTGCAG |
| Bivalvia Picorna-like virus N34 | ZH08_TRINITY_DN176904_c0_g1_i1_279F | CAAAACCCAACGGCGGAACTAGTA |
| Bivalvia Picorna-like virus N34 | ZH08_TRINITY_DN176904_c0_g1_i1_774R | GACGCTGCCGCAAAATCTGTGAAT |
| Bivalvia Picorna-like virus N33 | ZH08_TRINITY_DN61069_c0_g1_i1_53F | ATGTTAGACGCTTCTAGCTGCTCG |
| Bivalvia Picorna-like virus N33 | ZH08_TRINITY_DN61069_c0_g1_i1_490R | GTACACTGGCAGTGTCTTTAAGGC |
| Bivalvia Picorna-like virus N37 | ZH08_TRINITY_DN79368_c0_g1_i1_320F | GTGGGCAAGAAAACGACTGACAAG |
| Bivalvia Picorna-like virus N37 | ZH08_TRINITY_DN79368_c0_g1_i1_832R | GCTGGGTTCCACAGAAAGGTATCA |
| Bivalvia Picorna-like virus N38 | ZH08_TRINITY_DN81541_c0_g1_i1_197F | CCTCCAATTCAGCACGGATATAGG |
| Bivalvia Picorna-like virus N38 | ZH08_TRINITY_DN81541_c0_g1_i1_649R | TTAGTTTGGTTGATGCTGCGGTGC |

| | | |
|---|---|---|
| Bivalvia Bunya-like virus N3 | ZH09_TRINITY_DN73110_c0_g1_i1_218F | TCTCGGCTAGGGTTGTCGTATATG |
| Bivalvia Bunya-like virus N3 | ZH09_TRINITY_DN73110_c0_g1_i1_687R | TGCCGAAAGGTGATCACATCCTAC |
| Bivalvia Picorna-like virus N40 | ZH10_TRINITY_DN12413_c0_g1_i6_1337F | CCTACTGCTGCAAGCTTATCTTGC |
| Bivalvia Picorna-like virus N40 | ZH10_TRINITY_DN12413_c0_g1_i6_1872R | GCTGATGGTTCTGATGCTGTTGAC |
| Bivalvia Picorna-like virus N44 | ZH10_TRINITY_DN22369_c0_g1_i4_353F | TGGAGCAAAACAGACCTTACCCTC |
| Bivalvia Picorna-like virus N44 | ZH10_TRINITY_DN22369_c0_g1_i4_965R | CGGTGTAGGCAATGATAAAGAGGC |
| Bivalvia Picorna-like virus N42 | ZH10_TRINITY_DN31476_c0_g1_i1_18F | GACGGCCATAGTTGTGTTAGGATC |
| Bivalvia Picorna-like virus N42 | ZH10_TRINITY_DN31476_c0_g1_i1_501R | AGTTCCAGCGTCTGTTTTGACACC |
| Bivalvia Picorna-like virus N41 | ZH10_TRINITY_DN4534_c0_g1_i1_278F | AGATACGAAGACTACCGCTATGCC |
| Bivalvia Picorna-like virus N41 | ZH10_TRINITY_DN4534_c0_g1_i1_828R | GGATGATGGAACCGAATTTCGTGG |
| Bivalvia Picorna-like virus N43 | ZH10_TRINITY_DN79622_c0_g1_i1_461F | GCGTAAGGATTATGAGCAGGAACC |
| Bivalvia Picorna-like virus N43 | ZH10_TRINITY_DN79622_c0_g1_i1_826R | GGTGCCACATTAAACACGCGTGTT |
| Bivalvia Picorna-like virus N45 | ZH11_TRINITY_DN12131_c0_g3_i4_420F | CCCAACTGGCAAAAACCAAAGCAG |
| Bivalvia Picorna-like virus N45 | ZH11_TRINITY_DN12131_c0_g3_i4_843R | GGACGCTGGGAATTGAGAGTTTTG |
| Bivalvia Durna-like virus N4 | ZH11_TRINITY_DN15550_c0_g1_i1_55F | GATTCCATACTGCACAACTGAACC |
| Bivalvia Durna-like virus N4 | ZH11_TRINITY_DN15550_c0_g1_i1_581R | CGATGCAATCGAAAGGTTTGTCAT |
| Bivalvia Ghabri-like virus N1 | ZH11_TRINITY_DN212836_c0_g1_i1_145F | ACTGGTTGATCTTACGCCAGATGG |
| Bivalvia Ghabri-like virus N1 | ZH11_TRINITY_DN212836_c0_g1_i1_573R | ATGAGCAAGCTTAGGTTGGCTGTC |
| Bivalvia Picorna-like virus N46 | ZH13_TRINITY_DN28503_c0_g1_i11_718F | TTTGCGATGCTACGGGTTCAGATC |
| Bivalvia Picorna-like virus N46 | ZH13_TRINITY_DN28503_c0_g1_i11_1178R | CCAACCCAATTCAACTCCTACACC |
| Bivalvia Durna-like virus N5 | ZH13_TRINITY_DN92793_c0_g1_i1_243F | CCAGTATAGATAGCGTGCGAATCG |
| Bivalvia Durna-like virus N5 | ZH13_TRINITY_DN92793_c0_g1_i1_669R | TTAGTGACACCCGATGGGGTATTG |
| Bivalvia Picorna-like virus N52 | ZH14_TRINITY_DN10501_c0_g1_i20_1854F | ACTAAATTCTTCCCTACGCCAGGG |
| Bivalvia Picorna-like virus N52 | ZH14_TRINITY_DN10501_c0_g1_i20_2285R | GAAGACAGTTTTAGTAGCCTGGGG |
| Bivalvia Picorna-like virus N54 | ZH14_TRINITY_DN10919_c0_g1_i2_536F | ATGCGACGGTAAAAAGCTTCTGGG |
| Bivalvia Picorna-like virus N54 | ZH14_TRINITY_DN10919_c0_g1_i2_1052R | ATTCCCGTGCACCAGAACTTAGTC |
| Bivalvia Picorna-like virus N58 | ZH14_TRINITY_DN20009_c0_g1_i3_698F | AGATGGTTATTATGGGCCTGAGGG |
| Bivalvia Picorna-like virus N58 | ZH14_TRINITY_DN20009_c0_g1_i3_1223R | TTCCGTCGGATTTACGGAAGATGG |
| Bivalvia Unclassified virus N2 | ZH14_TRINITY_DN2139_c0_g1_i7_250F | CATACCCTCCCCATTCTTGTTGTC |
| Bivalvia Unclassified virus N2 | ZH14_TRINITY_DN2139_c0_g1_i7_734R | GGTATGCCAAATGGACAGTGGAGA |
| Bivalvia Picorna-like virus N57 | ZH14_TRINITY_DN22339_c0_g1_i2_488F | AACGGACCAAAGAAACGAGAACAT |
| Bivalvia Picorna-like virus N57 | ZH14_TRINITY_DN22339_c0_g1_i2_1010R | GACTTCTTCGGTGTAACTGGTGGA |
| Bivalvia Picorna-like virus N47 | ZH14_TRINITY_DN2243_c0_g1_i6_825F | CTTCTAGACTAGCCATGGGTAACG |
| Bivalvia Picorna-like virus N47 | ZH14_TRINITY_DN2243_c0_g1_i6_1295R | GATTCACCCCAAGGGAATGTATCG |
| Bivalvia Picorna-like virus N55 | ZH14_TRINITY_DN23605_c0_g2_i1_367F | CTTCATCTGCTTTGGCAACCGGTA |
| Bivalvia Picorna-like virus N55 | ZH14_TRINITY_DN23605_c0_g2_i1_850R | CGCACTTTCTTGCCTAGAGTGTCA |
| Bivalvia Unclassified virus N1 | ZH14_TRINITY_DN248443_c0_g1_i1_461F | GCCCATTGAATACACTCAGCCATG |
| Bivalvia Unclassified virus N1 | ZH14_TRINITY_DN248443_c0_g1_i1_663R | GGTGTCGATTCTACGTTAGTTCGG |
| Bivalvia Picorna-like virus N48 | ZH14_TRINITY_DN32447_c0_g1_i3_272F | CACAGTACCTCTCGTAAAACGACG |
| Bivalvia Picorna-like virus N48 | ZH14_TRINITY_DN32447_c0_g1_i3_986R | TGTGAGTGAGCAGCCATGTACAAG |
| Bivalvia Toli-like virus N2 | ZH14_TRINITY_DN34716_c0_g1_i6_250F | TGGCGTCAGGTAACCGAATCAAAG |
| Bivalvia Toli-like virus N2 | ZH14_TRINITY_DN34716_c0_g1_i6_691R | AATGCCACATCTACACCTTCTGGC |
| Bivalvia Picorna-like virus N49 | ZH14_TRINITY_DN4424_c0_g1_i17_426F | GAAGGCCACTTACCACTGATGAAG |

| | | |
|---|---|---|
| Bivalvia Picorna-like virus N49 | ZH14_TRINITY_DN4424_c0_g1_i17_760R | CCTACCTGCAGCTACTTTCTCATG |
| Bivalvia Picorna-like virus N50 | ZH14_TRINITY_DN520_c0_g1_i6_3785F | TCGAAGTCAACCTCACAAAACGCC |
| Bivalvia Picorna-like virus N50 | ZH14_TRINITY_DN520_c0_g1_i6_4235R | GCGCTGCTACAATCTTCTTTGAGG |
| Bivalvia Picorna-like virus N51 | ZH14_TRINITY_DN60154_c0_g2_i4_458F | ATGGTCAATACCCGTACGATCGTG |
| Bivalvia Picorna-like virus N51 | ZH14_TRINITY_DN60154_c0_g2_i4_937R | CCGAACGTTTGGGCTGCAAATTAC |
| Bivalvia Picorna-like virus N56 | ZH14_TRINITY_DN7333_c0_g1_i6_1235F | AGAATGCCTAACTGCGTCCATCAC |
| Bivalvia Picorna-like virus N56 | ZH14_TRINITY_DN7333_c0_g1_i6_1704R | GGATAGGTAAGACGTTGAAGGAGC |
| Bivalvia Unclassified virus N3 | ZH14_TRINITY_DN7640_c0_g1_i2_481F | TCATTAGGGTATCCCTCAGGTCTC |
| Bivalvia Unclassified virus N3 | ZH14_TRINITY_DN7640_c0_g1_i2_956R | TATGTGTACAGGTGTCCTACTCGC |
| Bivalvia Picorna-like virus N53 | ZH14_TRINITY_DN7791_c0_g1_i7_1446F | CAACCTTCTCCATTTGGGTCTCTC |
| Bivalvia Picorna-like virus N53 | ZH14_TRINITY_DN7791_c0_g1_i7_1920R | AAAAGGCCTCCCAGAGTTATGGAG |
| Cephalopoda Durna-like virus H1 | H05_TRINITY_DN1216_c0_g1_i1_117F | CCATCTTGCTTCTCACGCATCTTA |
| Cephalopoda Durna-like virus H1 | H05_TRINITY_DN1216_c0_g1_i1_549R | TAATGTATTCTCAGGTCCAGGGCG |
| Cephalopoda Durna-like virus H2 | H05_TRINITY_DN3806_c0_g1_i3_334F | CTTCTGCGCTCATTGACATCATGC |
| Cephalopoda Durna-like virus H2 | H05_TRINITY_DN3806_c0_g1_i3_975R | CTTCCGATCTCCCTAAGAAGGATG |
| Cephalopoda Mononega-like virus H1 | H06_TRINITY_DN19025_c0_g2_i1_252F | CAGGAATGATTCTTGTCAACGAGG |
| Cephalopoda Mononega-like virus H1 | H06_TRINITY_DN19025_c0_g2_i1_783R | GCCTTCATTGGAGAATCTGCTCTC |
| Cephalopoda Bunya-like virus H1 | H06_TRINITY_DN2012_c0_g1_i1_7F | TTTAATGATACGGCGACCACCGAG |
| Cephalopoda Bunya-like virus H1 | H06_TRINITY_DN2012_c0_g1_i1_452R | AGGAGTAGGCATCCACAGGTTTAG |
| Bivalvia Picorna-like virus H12 | H07_TRINITY_DN208108_c0_g2_i1_44F | GGTTTGCCATATTGCTCTTCTGGC |
| Bivalvia Picorna-like virus H12 | H07_TRINITY_DN208108_c0_g2_i1_650R | TTTTCTTCCCTCGGAAAGGTGTGG |
| Bivalvia Picorna-like virus H18 | H07_TRINITY_DN338148_c0_g1_i1_48F | AAACTGGCATTGGAATATTGCCAG |
| Bivalvia Picorna-like virus H18 | H07_TRINITY_DN338148_c0_g1_i1_544R | AGGGTGTGTCTTCATCAACTTCAA |
| Bivalvia Durna-like virus H6 | H07_TRINITY_DN339057_c0_g1_i1_153F | TCAACCTATTGAGTGCTCTCCAGG |
| Bivalvia Durna-like virus H6 | H07_TRINITY_DN339057_c0_g1_i1_677R | ACGTGAATACCGCTATGACTCGTC |
| Bivalvia Picorna-like virus H8 | H07_TRINITY_DN3435_c0_g4_i4_112F | ACTGTGATATCTGGCCTATCGTCC |
| Bivalvia Picorna-like virus H8 | H07_TRINITY_DN3435_c0_g4_i4_541R | ATGTTGATACGCAAGATTTGGTCG |
| Bivalvia Picorna-like virus H16 | H07_TRINITY_DN352121_c0_g1_i1_392F | CAACGGGAACCGGATGTATATTGG |
| Bivalvia Picorna-like virus H16 | H07_TRINITY_DN352121_c0_g1_i1_1114R | CATATTGGCCAGTCCCATTGGGAA |
| Bivalvia Picorna-like virus H9 | H07_TRINITY_DN38393_c0_g1_i1_47F | AATGATGCTTTATCGCATCCTGGT |
| Bivalvia Picorna-like virus H9 | H07_TRINITY_DN38393_c0_g1_i1_641R | CCATATCTGGGCAGACCAACCTAT |
| Bivalvia Sobeli-like virus H1 | H07_TRINITY_DN38542_c0_g1_i3_1102F | GTGATCTCGCAAGTTTCAGCCTTG |
| Bivalvia Sobeli-like virus H1 | H07_TRINITY_DN38542_c0_g1_i3_1530R | AATACACTTGGGGGAAATGGGAGC |
| Bivalvia Picorna-like virus H10 | H07_TRINITY_DN5264_c0_g1_i7_62F | TCAGAAGAGTGAGAGTGATGTGGC |
| Bivalvia Picorna-like virus H10 | H07_TRINITY_DN5264_c0_g1_i7_505R | TTATCTTGGGGCTTACGCCTCAAG |
| Bivalvia Picorna-like virus H15 | H07_TRINITY_DN579_c0_g1_i13_670F | AAACACACGAATGGCCAAGTGACC |
| Bivalvia Picorna-like virus H15 | H07_TRINITY_DN579_c0_g1_i13_1364R | CTCACGCTAGTCATCATTCTGAGG |
| Bivalvia Picorna-like virus H11 | H07_TRINITY_DN8311_c0_g1_i5_427F | CTTTATCGTCCAATGCCGCCTGAT |
| Bivalvia Picorna-like virus H11 | H07_TRINITY_DN8311_c0_g1_i5_912R | GATCCAAGAACACTACATCAGCCG |
| Bivalvia Durna-like virus H7 | H09_TRINITY_DN214833_c0_g1_i1_256F | TAGTCCTGAAGAGATACTCCGTGG |
| Bivalvia Durna-like virus H7 | H09_TRINITY_DN214833_c0_g1_i1_695R | TTTCATCGAGGACAGTTGCTAGCG |
| Bivalvia Durna-like virus H8 | H09_TRINITY_DN29861_c0_g1_i3_242F | CCGGAAGAAATCCTTTCAGGTTGG |
| Bivalvia Durna-like virus H8 | H09_TRINITY_DN29861_c0_g1_i3_683R | ATCTCATCTCATCGACGACAGCTG |

| | | |
|---|---|---|
| Bivalvia Picorna-like virus H19 | H09_TRINITY_DN38653_c0_g1_i1_122F | AGAACTGATAAACAGGGGCAACCC |
| Bivalvia Picorna-like virus H19 | H09_TRINITY_DN38653_c0_g1_i1_564R | ATCTAGATCTGTGCGAGTCCTCTG |
| Bivalvia Sobeli-like virus H3 | H09_TRINITY_DN40946_c0_g1_i1_2672F | GTCCTATGCTGGATGCTCTTGAAG |
| Bivalvia Sobeli-like virus H3 | H09_TRINITY_DN40946_c0_g1_i1_3101R | CGCACCTGTGCATAATAAGCTTCC |
| Bivalvia Picorna-like virus H20 | H09_TRINITY_DN59781_c0_g1_i2_423F | CACCAGGACAGTACCAAACAGCAA |
| Bivalvia Picorna-like virus H20 | H09_TRINITY_DN59781_c0_g1_i2_931R | GGTCTACCGTGACCTTTCTGATGA |
| Gastropoda Picorna-like virus H3 | H13_TRINITY_DN7_c0_g1_i3_470F | CTTGATTTACCTATCTCCACCCGC |
| Gastropoda Picorna-like virus H3 | H13_TRINITY_DN7_c0_g1_i3_937R | TGGGACCTAAGCACACAACTAACG |
| Polychaeta Picorna-like virus H6 | H14_TRINITY_DN15661_c0_g1_i2_244F | TTCTGAAGTCAGCGTCTGGGAAAC |
| Polychaeta Picorna-like virus H6 | H14_TRINITY_DN15661_c0_g1_i2_690R | ACATCAATCCGGCCATCATTGTCC |
| Polychaeta Picorna-like virus H5 | H14_TRINITY_DN20355_c0_g1_i3_319F | TGTAGCAATGCGAGCATCCAAACG |
| Polychaeta Picorna-like virus H5 | H14_TRINITY_DN20355_c0_g1_i3_756R | ATTGCTTGGGAATACACGCGGTAC |
| Polychaeta Picorna-like virus H2 | H14_TRINITY_DN20824_c0_g1_i1_333F | GGCATAAAGACCAGGATGGTACGT |
| Polychaeta Picorna-like virus H2 | H14_TRINITY_DN20824_c0_g1_i1_882R | GAATGGTGTTGCTTCGGTTGCTAG |
| Polychaeta Picorna-like virus H4 | H14_TRINITY_DN2661_c0_g1_i1_337F | ACCTGAAGGAGAAAGTCTAGCCAC |
| Polychaeta Picorna-like virus H4 | H14_TRINITY_DN2661_c0_g1_i1_771R | TTTGTTGCTAGTGGTCCTAGCACG |
| Polychaeta Picorna-like virus H3 | H14_TRINITY_DN32077_c0_g1_i1_322F | GTTTCCTGCTCTTATGGTTCCACC |
| Polychaeta Picorna-like virus H3 | H14_TRINITY_DN32077_c0_g1_i1_628R | GTGAGACTTGCAATCCGTTGGGAT |
| Polychaeta Picorna-like virus H1 | H14_TRINITY_DN32628_c0_g1_i2_353F | GAGTTTTCACCACAGACAGGGGAA |
| Polychaeta Picorna-like virus H1 | H14_TRINITY_DN32628_c0_g1_i2_1010R | CTTGCACACGAGTCTGTCGAAAAC |
| Bivalvia Picorna-like virus D3 | 12_2747F_[Beihai_sipunculid_worm_vir...]_1 | TGGAGAACATAGAGTCGTAGCAGG |
| Bivalvia Picorna-like virus D3 | 12_3188R_[Beihai_sipunculid_worm_vir...]_1 | ATCACACTCGGCTAATACACGTGC |
| Bivalvia Picorna-like virus D8 | 13_1886F_[Marine_RNA_virus_BC-1] | GCAGGAACAAGTTGTTGACAGAGC |
| Bivalvia Picorna-like virus D8 | 13_2338R_[Marine_RNA_virus_BC-1] | CGCCTGAAATATGCAGAACCCAAC |
| Gastropoda Picorna-like virus D1 | 17_2163F_[Beihai_picorna-like_virus_33]_1 | ACTCAAAAGTCAAGGCCTGAGCTC |
| Gastropoda Picorna-like virus D1 | 17_2587R_[Beihai_picorna-like_virus_33]_1 | TTAGCAAGGTATTCTGCGACCTCG |
| Gastropoda Picorna-like virus D1 | 17_599F_[Beihai_picorna-like_virus_33]_2 | GACAGTCGTAAGCCTCTTATGTGG |
| Gastropoda Picorna-like virus D1 | 17_1043R_[Beihai_picorna-like_virus_33]_2 | ATTACGAGCTTCCACTGCTCGTTG |
| Bivalvia Picorna-like virus D31 | 19_843F_[Beihai_picorna-like_virus_16] | CACAGAGACCTCATGTGTACTTGG |
| Bivalvia Picorna-like virus D31 | 19_1311R_[Beihai_picorna-like_virus_16] | GGCAATGGAGAATTTACGCAAGGC |
| Bivalvia Picorna-like virus D36 | 19_489F_[Hubei_picorna-like_virus_45] | CTTGGAATCGTGTAATCGAGACCG |
| Bivalvia Picorna-like virus D36 | 19_939R_[Hubei_picorna-like_virus_45] | GCAGTGCGGTAATTTTCCTCAGTC |
| Bivalvia Unclassified virus D2 | 19_511F_[Beihai_octopus_virus_1] | CTTAAGACGGTTGGAGTCTATGGG |
| Bivalvia Unclassified virus D2 | 19_936R_[Beihai_octopus_virus_1] | GGGCTATAAGGGACGTTGTTACAC |
| Bivalvia Picorna-like virus D31 | 19_3409F_[Beihai_paphia_shell_virus_2] | GCCCTCACCTTTGCTAAATACGAC |
| Bivalvia Picorna-like virus D31 | 19_3851R_[Beihai_paphia_shell_virus_2] | TTGTTGACTTCTACAGTGAGGGGC |
| Bivalvia Picorna-like virus D32 | 19_696F_[Beihai_picorna-like_virus_41] | AAACAGGAAGAGCGAGTGTGAGAC |
| Bivalvia Picorna-like virus D32 | 19_1242R_[Beihai_picorna-like_virus_41] | CCCGTGGGTTTTTATTCCCACAAG |
| Bivalvia Picorna-like virus D37 | 19_774F_[Beihai_sipunculid_worm_vir...] | GGATCAAGAAACATCAGGTGTGGG |
| Bivalvia Picorna-like virus D37 | 19_1370R_[Beihai_sipunculid_worm_vir...] | TCATCATCAGAGTGAGCTACCAGC |
| Bivalvia Picorna-like virus D39 | 19_697F_[European_brown_hare_sy...] | CAGTTTTGGCTTATGGAGCACCAC |
| Bivalvia Picorna-like virus D39 | 19_1208R_[European_brown_hare_sy...] | GCCTGTTCCTTTTGGACCTCTAAG |
| Bivalvia Picorna-like virus D37 | 19_2514F_[Marine_RNA_virus_PAL438] | TTGTTGCTTCGCGTAGTGCTTTGG |

| | | |
|---|---|---|
| Bivalvia Picorna-like virus D37 | 19_2980R_[Marine_RNA_virus_PAL438] | GAGAGCAACATTCCGTGCAAAAGG |
| Bivalvia Picorna-like virus D46 | 20_762F_[Bat_dicibavirus] | CGCATGTATGCGAATATGGCTGAG |
| Bivalvia Picorna-like virus D46 | 20_1231R_[Bat_dicibavirus] | AGCACAATCTTGCCGGCTAGATTC |
| Bivalvia Picorna-like virus D57 | 21_1088F_[Hubei_picorna-like_virus_11] | GACATGCATGCTGAAGGTGGAAAC |
| Bivalvia Picorna-like virus D57 | 21_1514R_[Hubei_picorna-like_virus_11] | AAGGCAACTCGTATTTCCGCTCAC |
| Bivalvia Picorna-like virus D56 | 21_577F_[Shuangao_insect_virus_12] | GCCGGAGCTTCTACTGAATTGAAG |
| Bivalvia Picorna-like virus D56 | 21_1055R_[Shuangao_insect_virus_12] | CGTCGTCCTACTTGATCTGTGTTC |
| Bivalvia Picorna-like virus D56 | 21_577F_[Shuangao_insect_virus_12] | GCCGGAGCTTCTACTGAATTGAAG |
| Bivalvia Picorna-like virus D56 | 21_1055R_[Shuangao_insect_virus_12] | CGTCGTCCTACTTGATCTGTGTTC |
| Bivalvia Picorna-like virus D57 | 21_1193F_[Wenzhou_gastropodes_virus_2] | GGATCTCCAGGCAAGCAGAAAAAC |
| Bivalvia Picorna-like virus D57 | 21_1631R_[Wenzhou_gastropodes_virus_2] | AACTGCCGGCATTCGTATTACCAG |
| Bivalvia Picorna-like virus D61 | 21_2404F_[Beihai_picorna-like_virus_4] | TCTAGCGATGATACCGTACCACTG |
| Bivalvia Picorna-like virus D61 | 21_2827R_[Beihai_picorna-like_virus_4] | AAACGCTATCCTACGCACGTGAAG |
| Bivalvia Picorna-like virus D61 | 21_2404F_[Beihai_picorna-like_virus_4] | TCTAGCGATGATACCGTACCACTG |
| Bivalvia Picorna-like virus D61 | 21_2827R_[Beihai_picorna-like_virus_4] | AAACGCTATCCTACGCACGTGAAG |
| Bivalvia Picorna-like virus D56 | 21_2577F_[Wenzhou_picorna-like_virus...] | TTCATGCTTTGACAGCACTAGCCG |
| Bivalvia Picorna-like virus D56 | 21_3096R_[Wenzhou_picorna-like_virus...] | GTGAAAAGACTATGCGTGGCATCC |
| Gastropoda Picorna-like virus D15 | 26_1896F_[Beihai_picorna-like_virus_75] | ATGGATTCTGCGTTCAGGAAGTGC |
| Gastropoda Picorna-like virus D15 | 26_2319R_[Beihai_picorna-like_virus_75] | AAGCTCCCTCATAGCTGACCTATG |
| Gastropoda Picorna-like virus D21 | 27_452F_[Wenzhou_picorna-like_virus...] | AGTTCCGGTGTTGGAAAATCAGGC |
| Gastropoda Picorna-like virus D21 | 27_876R_[Wenzhou_picorna-like_virus...] | TTCACTTGAACCTCTCCAACGAGG |
| Bivalvia Picorna-like virus D72 | 8-1_3933F_[Hubei_odonate_virus_5] | TGTTACCAGATGGAGAGAAGGTCG |
| Bivalvia Picorna-like virus D72 | 8-1_4378R_[Hubei_odonate_virus_5] | AACCAGACTTCCACATGATGCGTC |
| Bivalvia Picorna-like virus D66 | 8-1_466F_[Sanxia_picorna-like_virus_13] | ATCCAGGAACCATCAGGATAGTGC |
| Bivalvia Picorna-like virus D66 | 8-1_905R_[Sanxia_picorna-like_virus_13] | ATACAGAGGATACAGCCGATGCTG |
| Bivalvia Picorna-like virus D87 | 8-2_924F_[Wenzhou_picorna-like_virus_24] | AAGTCGTATTGGAGGTGGAGTCTG |
| Bivalvia Picorna-like virus D87 | 8-2_1348R_[Wenzhou_picorna-like_virus_24] | CAACGACACAAGACCTAACACCAG |
| Bivalvia Picorna-like virus N6 | BH06_472F_[Beihai_picorna-like_virus_122]_1 | CACTTAGCTGAGAAATCAGCTGCG |
| Bivalvia Picorna-like virus N6 | BH06_914R_[Beihai_picorna-like_virus_122]_1 | TGCTGCACTGCCAAGTTGTTCTTG |
| Bivalvia Picorna-like virus N7 | BH06_617F_[Beihai_picorna-like_virus_122]_2 | AGGAGATCAAAAAGCTCGTGGTGG |
| Bivalvia Picorna-like virus N7 | BH06_1058R_[Beihai_picorna-like_virus_122]_2 | GGGTGCTCTCACTACTGATTTCTC |
| Bivalvia Picorna-like virus N8 | BH06_2419F_[Wenzhou_gastropodes_virus_2] | TGATGGGCAATTATTGCAGGCAGC |
| Bivalvia Picorna-like virus N8 | BH06_2979R_[Wenzhou_gastropodes_virus_2] | CCATAACTCTGGGAGGCCTTTTTC |
| Bivalvia Picorna-like virus N8 | BH06_2419F_[Wenzhou_gastropodes_virus_2] | TGATGGGCAATTATTGCAGGCAGC |
| Bivalvia Picorna-like virus N8 | BH06_2979R_[Wenzhou_gastropodes_virus_2] | CCATAACTCTGGGAGGCCTTTTTC |
| Bivalvia Picorna-like virus N8 | BH06_2419F_[Wenzhou_gastropodes_virus_2] | TGATGGGCAATTATTGCAGGCAGC |
| Bivalvia Picorna-like virus N8 | BH06_2979R_[Wenzhou_gastropodes_virus_2] | CCATAACTCTGGGAGGCCTTTTTC |
| Gastropoda Picorna-like virus N1 | BH09_449F_[Beihai_mollusks_virus_2] | CTCCATCCCCCTTGAAGAAATTGG |
| Gastropoda Picorna-like virus N1 | BH09_875R_[Beihai_mollusks_virus_2] | GGTCGTTCCTCAAGCTTTTGGAAC |
| Gastropoda Picorna-like virus N1 | BH09_449F_[Beihai_mollusks_virus_2] | CTCCATCCCCCTTGAAGAAATTGG |
| Gastropoda Picorna-like virus N1 | BH09_875R_[Beihai_mollusks_virus_2] | GGTCGTTCCTCAAGCTTTTGGAAC |
| Gastropoda Picorna-like virus N1 | BH09_449F_[Beihai_mollusks_virus_2] | CTCCATCCCCCTTGAAGAAATTGG |
| Gastropoda Picorna-like virus N1 | BH09_875R_[Beihai_mollusks_virus_2] | GGTCGTTCCTCAAGCTTTTGGAAC |

| | | |
|---|---|---|
| Gastropoda Picorna-like virus N1 | BH09_449F_[Beihai_mollusks_virus_2] | CTCCATCCCCCTTGAAGAAATTGG |
| Gastropoda Picorna-like virus N1 | BH09_875R_[Beihai_mollusks_virus_2] | GGTCGTTCCTCAAGCTTTTGGAAC |
| Gastropoda Picorna-like virus N1 | BH09_449F_[Beihai_mollusks_virus_2] | CTCCATCCCCCTTGAAGAAATTGG |
| Gastropoda Picorna-like virus N1 | BH09_875R_[Beihai_mollusks_virus_2] | GGTCGTTCCTCAAGCTTTTGGAAC |
| Gastropoda Picorna-like virus N2 | BH09_2920F_[Marine_RNA_virus_JP-A] | CAGATCTATCTGTGGTGTGGGTTC |
| Gastropoda Picorna-like virus N2 | BH09_3392R_[Marine_RNA_virus_JP-A] | ATTGATCGGGCTCTTAGGATGCAC |
| Gastropoda Picorna-like virus N2 | BH09_2920F_[Marine_RNA_virus_JP-A] | CAGATCTATCTGTGGTGTGGGTTC |
| Gastropoda Picorna-like virus N2 | BH09_3392R_[Marine_RNA_virus_JP-A] | ATTGATCGGGCTCTTAGGATGCAC |
| Gastropoda Picorna-like virus N2 | BH09_2920F_[Marine_RNA_virus_JP-A] | CAGATCTATCTGTGGTGTGGGTTC |
| Gastropoda Picorna-like virus N2 | BH09_3392R_[Marine_RNA_virus_JP-A] | ATTGATCGGGCTCTTAGGATGCAC |
| Gastropoda Picorna-like virus N2 | BH09_2920F_[Marine_RNA_virus_JP-A] | CAGATCTATCTGTGGTGTGGGTTC |
| Gastropoda Picorna-like virus N2 | BH09_3392R_[Marine_RNA_virus_JP-A] | ATTGATCGGGCTCTTAGGATGCAC |
| Gastropoda Picorna-like virus N2 | BH09_653F_[Bat_dicibavirus] | TTCCCATGATGGGTATTTCGGGAG |
| Gastropoda Picorna-like virus N2 | BH09_1132R_[Bat_dicibavirus] | TATCGTCAGCTTTCCACATGTGGC |
| Gastropoda Picorna-like virus N2 | BH09_653F_[Bat_dicibavirus] | TTCCCATGATGGGTATTTCGGGAG |
| Gastropoda Picorna-like virus N2 | BH09_1132R_[Bat_dicibavirus] | TATCGTCAGCTTTCCACATGTGGC |
| Gastropoda Picorna-like virus N2 | BH09_653F_[Bat_dicibavirus] | TTCCCATGATGGGTATTTCGGGAG |
| Gastropoda Picorna-like virus N2 | BH09_1132R_[Bat_dicibavirus] | TATCGTCAGCTTTCCACATGTGGC |
| Gastropoda Picorna-like virus N5 | BH12_641F_[Cricket_paralysis_virus] | CATTGTTGAGTCGTCTCTGCATGG |
| Gastropoda Picorna-like virus N5 | BH12_1074R_[Cricket_paralysis_virus] | AAACGACATCAGGCACATCACCTC |
| Gastropoda Picorna-like virus N5 | BH12_921F_[Himetobi_P_virus] | ATCGGATATGACTATGACGCCCTC |
| Gastropoda Picorna-like virus N5 | BH12_1351R_[Himetobi_P_virus] | TCTCCAAGGAAGCGGCATTCAATC |
| Gastropoda Picorna-like virus N7 | BH13_435F_[Beihai_octopus_virus_1] | GCGTTTTGAGACATGCCCTACTAG |
| Gastropoda Picorna-like virus N7 | BH13_870R_[Beihai_octopus_virus_1] | ATCACCAACCATACAGGTGGTGAC |
| Gastropoda Picorna-like virus N7 | BH13_3492F_[Marine_RNA_virus_JP-A] | AGGCTTTTAGGGTTACTCCGGTAC |
| Gastropoda Picorna-like virus N7 | BH13_3927R_[Marine_RNA_virus_JP-A] | TTCCAATCACCACCATTGGGAACC |
| Gastropoda Picorna-like virus N7 | BH13_3492F_[Marine_RNA_virus_JP-A] | AGGCTTTTAGGGTTACTCCGGTAC |
| Gastropoda Picorna-like virus N7 | BH13_3927R_[Marine_RNA_virus_JP-A] | TTCCAATCACCACCATTGGGAACC |
| Gastropoda Picorna-like virus N7 | BH13_3492F_[Marine_RNA_virus_JP-A] | AGGCTTTTAGGGTTACTCCGGTAC |
| Gastropoda Picorna-like virus N7 | BH13_3927R_[Marine_RNA_virus_JP-A] | TTCCAATCACCACCATTGGGAACC |
| Gastropoda Picorna-like virus N12 | G11_2142F_[Wenling_crustacean_virus_2] | CTTAAAGAGAGTGATGCTTCCCGC |
| Gastropoda Picorna-like virus N12 | G11_2622R_[Wenling_crustacean_virus_2] | GACAAGCTGGAATGCGTACATTCG |
| Gastropoda Picorna-like virus N12 | G11_2142F_[Wenling_crustacean_virus_2] | CTTAAAGAGAGTGATGCTTCCCGC |
| Gastropoda Picorna-like virus N12 | G11_2622R_[Wenling_crustacean_virus_2] | GACAAGCTGGAATGCGTACATTCG |
| Gastropoda Picorna-like virus N12 | G11_2142F_[Wenling_crustacean_virus_2] | CTTAAAGAGAGTGATGCTTCCCGC |
| Gastropoda Picorna-like virus N12 | G11_2622R_[Wenling_crustacean_virus_2] | GACAAGCTGGAATGCGTACATTCG |
| Gastropoda Picorna-like virus N12 | G11_2142F_[Wenling_crustacean_virus_2] | CTTAAAGAGAGTGATGCTTCCCGC |
| Gastropoda Picorna-like virus N12 | G11_2622R_[Wenling_crustacean_virus_2] | GACAAGCTGGAATGCGTACATTCG |
| Gastropoda Picorna-like virus N12 | G11_2142F_[Wenling_crustacean_virus_2] | CTTAAAGAGAGTGATGCTTCCCGC |
| Gastropoda Picorna-like virus N12 | G11_2622R_[Wenling_crustacean_virus_2] | GACAAGCTGGAATGCGTACATTCG |
| Bivalvia Picorna-like virus N12 | G12_543F_[Beihai_sipunculid_worm_vir...] | AACCGATCACCTTCATCAACCTCC |
| Bivalvia Picorna-like virus N12 | G12_1033R_[Beihai_sipunculid_worm_vir...] | AAGCGCTTCTTCCGATAGTGCTAC |
| Bivalvia Picorna-like virus N29 | G14_4350F_[Beihai_sesarmid_crab_virus_2]_2 | ATACCTGACTCCTGATACAGCAGC |

| | | |
|---|---|---|
| Bivalvia Picorna-like virus N29 | G14_4836R_[Beihai_sesarmid_crab_virus_2]_2 | AGGAGAATACCACATTCCTCCTCG |
| Bivalvia Picorna-like virus N23 | G14_767F_[Biomphalaria_virus_1] | ACAACAAGCTTGTGATCGCTCAGC |
| Bivalvia Picorna-like virus N23 | G14_1545R_[Biomphalaria_virus_1] | AATCTCCAGGAGGGAGATTTGTCG |
| Cephalopoda Unclassified virus H1 | H05_846F_[Wenling_crustacean_virus_3] | CGCCGAAGCAAGAATTTACGACAC |
| Cephalopoda Unclassified virus H1 | H05_1293R_[Wenling_crustacean_virus_3] | GATTCGAACACGATCACGTTGAGC |
| Cephalopoda Unclassified virus H1 | H05_846F_[Wenling_crustacean_virus_3] | CGCCGAAGCAAGAATTTACGACAC |
| Cephalopoda Unclassified virus H1 | H05_1293R_[Wenling_crustacean_virus_3] | GATTCGAACACGATCACGTTGAGC |
| Cephalopoda Unclassified virus H1 | H05_846F_[Wenling_crustacean_virus_3] | CGCCGAAGCAAGAATTTACGACAC |
| Cephalopoda Unclassified virus H1 | H05_1293R_[Wenling_crustacean_virus_3] | GATTCGAACACGATCACGTTGAGC |
| Cephalopoda Unclassified virus H1 | H05_846F_[Wenling_crustacean_virus_3] | CGCCGAAGCAAGAATTTACGACAC |
| Cephalopoda Unclassified virus H1 | H05_1293R_[Wenling_crustacean_virus_3] | GATTCGAACACGATCACGTTGAGC |
| Cephalopoda Unclassified virus H1 | H05_846F_[Wenling_crustacean_virus_3] | CGCCGAAGCAAGAATTTACGACAC |
| Cephalopoda Unclassified virus H1 | H05_1293R_[Wenling_crustacean_virus_3] | GATTCGAACACGATCACGTTGAGC |
| Cephalopoda Picorna-like virus H1 | H05_699F_[Wenling_picorna-like_virus_2] | TAAGTTCGGATGTGGTGCGTAAGC |
| Cephalopoda Picorna-like virus H1 | H05_1213R_[Wenling_picorna-like_virus_2] | ATGTGTTGGCATGTTGACGAGAGG |
| Cephalopoda Picorna-like virus H1 | H05_699F_[Wenling_picorna-like_virus_2] | TAAGTTCGGATGTGGTGCGTAAGC |
| Cephalopoda Picorna-like virus H1 | H05_1213R_[Wenling_picorna-like_virus_2] | ATGTGTTGGCATGTTGACGAGAGG |
| Cephalopoda Picorna-like virus H1 | H05_699F_[Wenling_picorna-like_virus_2] | TAAGTTCGGATGTGGTGCGTAAGC |
| Cephalopoda Picorna-like virus H1 | H05_1213R_[Wenling_picorna-like_virus_2] | ATGTGTTGGCATGTTGACGAGAGG |
| Bivalvia Picorna-like virus H13 | H07_1295F_[Beihai_picorna-like_virus_122] | AGGCGATTCCCGATTTCCTACAAG |
| Bivalvia Picorna-like virus H13 | H07_1734R_[Beihai_picorna-like_virus_122] | TTTGCAACACCTAGCTCTTCAGCC |
| Bivalvia Picorna-like virus H13 | H07_1295F_[Beihai_picorna-like_virus_122] | AGGCGATTCCCGATTTCCTACAAG |
| Bivalvia Picorna-like virus H13 | H07_1734R_[Beihai_picorna-like_virus_122] | TTTGCAACACCTAGCTCTTCAGCC |
| Bivalvia Picorna-like virus H14 | H07_1295F_[Beihai_picorna-like_virus_122] | AGGCGATTCCCGATTTCCTACAAG |
| Bivalvia Picorna-like virus H14 | H07_1734R_[Beihai_picorna-like_virus_122] | TTTGCAACACCTAGCTCTTCAGCC |
| Bivalvia Picorna-like virus H17 | H07_481F_[Wenzhou_gastropodes_virus_2] | AGTCCATGGTGATTGGCATGAGAG |
| Bivalvia Picorna-like virus H17 | H07_1238R_[Wenzhou_gastropodes_virus_2] | ACACTGGGTCAAATCCTTGTGCTG |
| Bivalvia Picorna-like virus H17 | H07_481F_[Wenzhou_gastropodes_virus_2] | AGTCCATGGTGATTGGCATGAGAG |
| Bivalvia Picorna-like virus H17 | H07_1238R_[Wenzhou_gastropodes_virus_2] | ACACTGGGTCAAATCCTTGTGCTG |
| Bivalvia Picorna-like virus H17 | H07_481F_[Wenzhou_gastropodes_virus_2] | AGTCCATGGTGATTGGCATGAGAG |
| Bivalvia Picorna-like virus H17 | H07_1238R_[Wenzhou_gastropodes_virus_2] | ACACTGGGTCAAATCCTTGTGCTG |
| Bivalvia Picorna-like virus H17 | H07_481F_[Wenzhou_gastropodes_virus_2] | AGTCCATGGTGATTGGCATGAGAG |
| Bivalvia Picorna-like virus H17 | H07_1238R_[Wenzhou_gastropodes_virus_2] | ACACTGGGTCAAATCCTTGTGCTG |
| Bivalvia Sobeli-like virus H2 | H09_938F_[Norovirus_GI] | TAAAACAAGGAGCCTCTTGGGCTG |
| Bivalvia Sobeli-like virus H2 | H09_1362R_[Norovirus_GI] | TTCCCGCACATCATCAGAGACTTG |
| Bivalvia Sobeli-like virus H2 | H09_938F_[Norovirus_GI] | TAAAACAAGGAGCCTCTTGGGCTG |
| Bivalvia Sobeli-like virus H2 | H09_1362R_[Norovirus_GI] | TTCCCGCACATCATCAGAGACTTG |
| Bivalvia Picorna-like virus N35 | ZH08_662F_[Beihai_picorna-like_virus_41] | TGACTCTTTATGCTGCTGGCAAGG |
| Bivalvia Picorna-like virus N35 | ZH08_1115R_[Beihai_picorna-like_virus_41] | TTTCGAGCCCATAAATTGGCCTGC |
| Bivalvia Picorna-like virus N35 | ZH08_662F_[Beihai_picorna-like_virus_41] | TGACTCTTTATGCTGCTGGCAAGG |
| Bivalvia Picorna-like virus N35 | ZH08_1115R_[Beihai_picorna-like_virus_41] | TTTCGAGCCCATAAATTGGCCTGC |
| Bivalvia Picorna-like virus N35 | ZH08_662F_[Beihai_picorna-like_virus_41] | TGACTCTTTATGCTGCTGGCAAGG |
| Bivalvia Picorna-like virus N35 | ZH08_1115R_[Beihai_picorna-like_virus_41] | TTTCGAGCCCATAAATTGGCCTGC |

| | | |
|---|---|---|
| Bivalvia Picorna-like virus N35 | ZH08_1579F_[Marine_RNA_virus_PAL438] | GGTTTTGATACTCGAATGGCAGCC |
| Bivalvia Picorna-like virus N35 | ZH08_2005R_[Marine_RNA_virus_PAL438] | ATACGGCTCCTTTGTCAGCCATTG |
| Bivalvia Toli-like virus N1 | ZH10_1502F_[Jingmen_tombus-like_virus_1]_1 | GGAGCCTATTTGGTCGGAAAACAC |
| Bivalvia Toli-like virus N1 | ZH10_1930R_[Jingmen_tombus-like_virus_1]_1 | CATACCTTCGCCTTTTTGCTCCAC |
| Bivalvia Picorna-like virus N39 | ZH10_659F_[Wenzhou_picorna-like_virus_5] | ATACGTGCCAAGCAAGGTTATGCC |
| Bivalvia Picorna-like virus N39 | ZH10_1186R_[Wenzhou_picorna-like_virus_5] | AAGAAAAGCATCAGCCATTGGGGG |
| Bivalvia Picorna-like virus N39 | ZH10_659F_[Wenzhou_picorna-like_virus_5] | ATACGTGCCAAGCAAGGTTATGCC |
| Bivalvia Picorna-like virus N39 | ZH10_1186R_[Wenzhou_picorna-like_virus_5] | AAGAAAAGCATCAGCCATTGGGGG |
